# Multi-Platform Assessment of DNA Sequencing Performance using Human and Bacterial Reference Genomes in the ABRF Next-Generation Sequencing Study

**DOI:** 10.1101/2020.07.23.218602

**Authors:** Jonathan Foox, Scott W. Tighe, Charles M. Nicolet, Justin M. Zook, Marta Byrska-Bishop, Wayne E. Clarke, Michael M. Khayat, Medhat Mahmoud, Phoebe K. Laaguiby, Zachary T. Herbert, Derek Warner, George S. Grills, Jin Jen, Shawn Levy, Jenny Xiang, Alicia Alonso, Gary P. Schroth, Fritz J. Sedlazeck, Giuseppe Narzisi, William Farmerie, Don A. Baldwin, Christopher E. Mason

**Affiliations:** Department of Physiology and Biophysics, Weill Cornell Medicine, New York, New York, USA; The HRH Prince Alwaleed Bin Talal Bin Abdulaziz Alsaud Institute for Computational Biomedicine, Weill Cornell Medicine, New York, New York, USA; University of Vermont Cancer Center, Vermont Integrative Genomics Resource, University of Vermont, Burlington, Vermont, USA; Keck School of Medicine, University of Southern California, Los Angeles, California, USA; Biosystems and Biomaterials Division, National Institute of Standards and Technology, Gaithersburg, Maryland, USA; New York Genome Center, New York, NY, 10013, USA; Human Genome Sequencing Center, Baylor College of Medicine, Houston, TX, USA; Department of Molecular and Human Genetics, Baylor College of Medicine, Houston, Texas; Molecular Biology Core Facilities, Dana-Farber Cancer Institute, Boston, Massachusetts, USA; DNA Sequencing Core, University of Utah, Salt Lake City, Utah, USA; Sylvester Comprehensive Cancer Center, University of Miami, Miami, FL, USA; Department of Laboratory Medicine and Pathology, Mayo Clinic, Rochester, MN; HudsonAlpha Institute for Biotechnology, Huntsville, Alabama, USA; Illumina, Inc., San Diego, CA, USA; Interdisciplinary Center for Biotechnology Research, University of Florida, Gainesville, Florida, USA; Department of Pathology, Fox Chase Cancer Center, Philadelphia, Pennsylvania, USA; The Feil Family Brain and Mind Research Institute, New York, New York, USA; The WorldQuant Initiative for Quantitative Prediction, Weill Cornell Medicine, New York, NY, USA

## Abstract

Massively parallel DNA sequencing is a critical tool for genomics research and clinical diagnostics. Here, we describe the Association of Biomolecular Resource Facilities (ABRF) Next-Generation Sequencing Phase II Study to measure quality and reproducibility of DNA sequencing. Replicates of human and bacterial reference DNA samples were generated across multiple sequencing platforms, including well-established technologies such as Illumina, ThermoFisher Ion Torrent, and Pacific Biosciences, as well as emerging technologies such as BGI, Genapsys, and Oxford Nanopore. A total of 202 datasets were generated to investigate the performance of a total of 16 sequencing platforms, including mappability of reads, coverage and error rates in difficult genomic regions, and detection of small-scale polymorphisms and large-scale structural variants. This study provides a comprehensive baseline resource for continual benchmarking as chemistries, methods, and platforms evolve for DNA sequencing.

## Introduction

High-throughput next-generation DNA sequencing (DNA-seq) is an essential analytical method for clinical and basic biomedical research [1, 2]. DNA-seq has numerous experimental applications, including (but not limited to) genotyping and variant discovery within individuals [3], population- and species-level characterization of genomes [4], and resolving the taxonomic diversity within a metagenomic mixture [5]. Genome sequencing has become ubiquitous, owing to the significant decrease in cost [6], which has led to rapid diversification of sample collection, library preparation, sequencing chemistry, and downstream bioinformatic pipelines. As sequencing technologies continually evolve, a broad, multi-dimensional collection of DNA-seq data can serve as a robust benchmarking tool set to assess our ability to capture genomes accurately, to investigate performance in the most and least reproducible regions of the genome, and to provide a valuable resource for laboratories and sequencing centers to evaluate new methods, chemistries, and protocols.

Prior examinations of DNA-seq have provided such valuable baselines. These studies have focused on single gene amplicon sequencing [7], multilocus/core genome bacterial typing [8], and on a select number of then-emerging platforms [9]. Previous large-scale studies of RNA-seq, such as the Microarray Quality Control (MAQC) Consortium, have established reference sets by characterizing variability within and between microarray platforms for reproducible detection of gene expression [10, 11]. This work was followed by an Association of Biomolecular Resource Facilities (ABRF) study, which conducted large-scale NGS to examine reproducibility in RNA-seq data [12]. This “Phase I” study characterized a range of RNA-seq libraries from reference RNA prepared with four preparation protocols and sequenced across five platforms and multiple labs. Concurrent reports described RNA-seq quality control [13], concordance with microarrays [14], and best practices for data processing [15] and normalization [16]. Together, these studies profiled intra- and inter-lab reproducibility, as well as baseline performance for RNA-seq, but there is not yet an analogous study for DNA-seq.

The National Institute of Standards and Technology (NIST), in partnership with the Genome In A Bottle (GIAB) Consortium, has enabled DNA-seq benchmarking by developing a series of reference materials (RM) [17], benchmarking tools [18], and associated ultra-deep sequence data [19]. A pilot RM, genome NA12878, is widely used as a technical standard, but is a single human genome collected with more limited donor consent. The recently released RM 8392 provides a family trio of genomes to support benchmarking of haplotypes and structural variants [20] and is consented through the Personal Genome Project [21] for broad use, publication, and inclusion in commercial products. However, it does not provide insights into the general variability between technologies and approaches that need to be considered for experimental designs.

Here, the ABRF NGS Phase II DNA-seq Study gives detailed insights for each currently commonly used sequencing technology by studying inter- and intra-lab replicates of a family trio of human genomes (NIST RM 8392, known as the Ashkenazi Trio;Mother (HG004), Father (HG003), and Son (HG002), three individual bacterial strains, and a metagenomic mixture often bacterial species, across well-established technologies including six Illumina and three ThermoFisher Ion Torrent platforms, as well as emerging sequencing technologies including BGI, Oxford Nanopore, and Genapsys platforms. The data generated by this consortium were examined for performance and reproducibility over a range of base compositions and GC-content profiles. Human libraries were synthesized using PCR-free whole genome and targeted exome capture methods. Microbial libraries were synthesized using specified techniques by each platform, including Nextera (Illumina), NEBNextUltra II (Genapsys), and LSK109 with and without native barcoding (Oxford Nanopore). Instrument and laboratory replicates were processed and compared through platform-dependent bioinformatic pipelines. Collectively, these data provide one of the largest inter-laboratory benchmarking resources for human and bacterial DNA-seq NGS across a wealth of platforms generated to date and a comprehensive resource for improving alignment, variant calling, and sequencing methods.

## Results

### Study Design and Data Quality

Individual human and bacterial genomes, as well as an equimolar mixture of ten bacterial species, were sequenced across an array of platforms, including Illumina, Ion Torrent, Oxford Nanopore, PacBio, MGI, and Genapsys (Figure 1; Supplementary Table 1). The overall quality of the sequencing data was consistently high across all replicates, seen in the insert size distributions for paired end reactions (Figure S1) and in the base quality distribution for all samples (Figure S2). To ensure comparability between data, all replicates were processed through the same bioinformatic pipeline, as appropriate to the data type, from raw read alignment through variant calling (see methods). For human reads, alignment was done against the GRCh37 reference that includes the hs37d5 decoy sequence in order to filter out spurious, easily misaligned reads, which successfully captured 5% of reads across samples (Figure S3).

**Figure 1:**
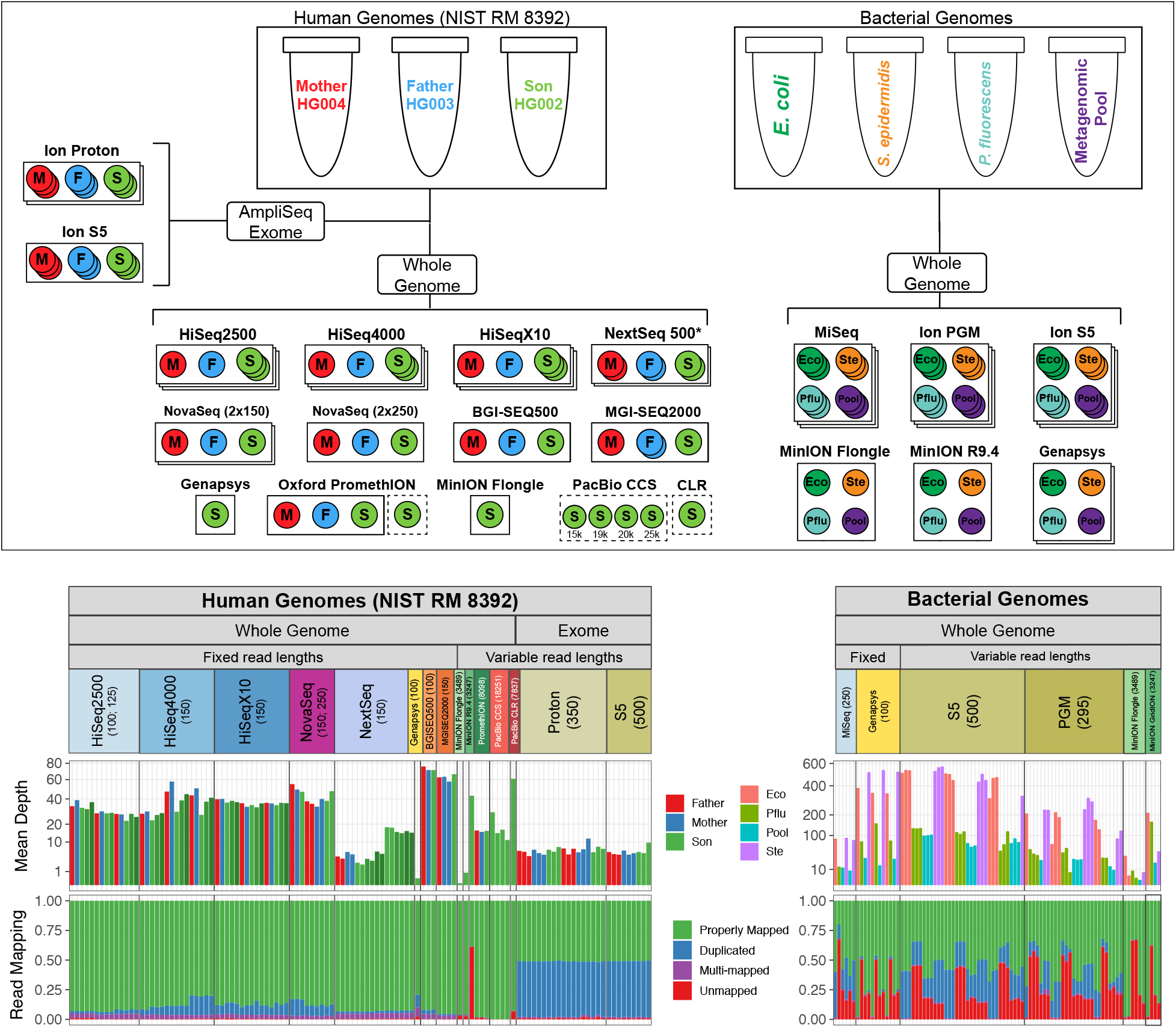
Experimental design and mapping results. Three standard human genomic DNA samples from the NIST Reference Material 8392 were used to prepare libraries, including TruSeq PCR-Free whole genome libraries and AmpliSeq exome libraries, for sequencing on multiple Illumina (HiSeq 2500, HiSeq 4000, HiSeq X10, NextSeq, NovaSeq) ThermoFisher (Ion Proton, Ion S5), MGI (BGISEQ500, MGISEQ2000), Oxford (MinION with R9.4 and Flongle flow cells;PromethION), and PacBio (CCS, CLR) platforms, Three bacterial species (*E. coli, S. epidermidis, P. fluorescens*) and one metagenomic mixture of ten bacterial species (Metagenomic Pool) were used to prepare libraries, including TruSeq PCR-Free whole genome libraries for Illumina MiSeq and Ion Xpress Plus libraries for Personal Genome Machine (PGM) and Ion S5. The number of intralab replicates is indicated by each circle, and the number of interlab replicates by the stacked rectangles. Mean genomic depth capture is plotted below, followed by a proportional bar plot representing the distribution of mapping outcomes for all reads per replicate.

Depth of sequencing and proportional distribution of read mapping outcomes was calculated for each replicate (Figure 1). The variability of sequencing depth was high between platforms, with data sets ranging from as many as 252 billion bases sequenced for one BGISEQ500 replicate to as few as 51 million bases sequenced for one MinION replicate using the Flongle flow cell. Variable depth of sequencing was also seen from different laboratories using the same platform;for example, with one HiSeq 4000 replicate yielding twice as much coverage as another (Figure 1; Supplementary Table 1). Despite differences in throughput, the rate at which reads mapped was very consistent between platforms across all human replicates. An average of 96.1% (93.0–97.7%) of reads mapped to the reference and were properly paired in paired-end reactions. An average of 3.72% (3.12–4.35%) of reads mapped to multiple sites in the reference;and an average of 0.79% (0.37–1.84%) of reads went unmapped at all (Supplementary Table 2). For AmpliSeq Exome panels, the rate of on-target mapping was high, ranging from 84.6–96.6%, with little variation between replicates (Supplementary Table 3).

Three individual bacterial species and one metagenomic mixture comprising ten bacterial species were sequenced on Illumina, Ion Torrent, Oxford Nanopore, and Genapsys platforms. One laboratory sequenced triplicates of each on an Illumina MiSeq, while three laboratories sequenced triplicates of each for both the Personal Genome Machine (PGM) and Ion S5. Each bacterial sample was only once with the Genapsys and Oxford nanopore platforms, however two two types of Oxford Nanopore flow cells and library methods were used in this study. For the Oxford MinION MK1b sequencer, the Flongle flow cell was paired with a single bacterial species using the LSK109 library method and run individually, while the larger format 9.4 flow cell allowed combining all samples on one flow cell using the LSK109 with native barcoding. This strategy was selected because Flongles are designed for low output (0.5-1 Gb) whereas the 9.4 flow cell has much higher read depth (15 Gb). The ten-species metagenomic mixture was included in the sequencing design in order to understand the influence that a mixed sample has on the capture of individual genomes. These species, which include the three species that were sequenced individually, plus seven other species comprising a wide variety of genome sizes, GC content, Gram staining responses, ecological niches, and posing physiological challenges for capture, such as high saline affinity (Supplementary Table 4). Overall, mappability was found to be directly related to the reference genome to which reads were mapped, across all platforms examined. For *E. coli* replicates, an average of 0.64% (0.14–2.08%) of reads went unmapped; for *S. epidermidis*, 5.11% (3.86-7.33%);for *P. fluorescens*, 57.19% (51.84-65.41%), and for the metagenomic mixed pool, 18.99% (17.46-24.30%). Notably, the finished genome was still in progress for the ATCC strain of *P. fluorescens*, so lower mapping rates were not unexpected. Both inter- and intralab replicates indicated consistent mapping distributions across species, library preparation, and sequencing platform, suggesting that reference genome is the limiting factor with respect to rate of mapping.

### Normalized Coverage Analysis

Depth of coverage was calculated within the repeat classes using the normalized 25x coverage alignments across all replicates with adequate depth of sequencing, using a mapping quality cutoff of MQ20. The distributions were very consistent among technologies, including short and long reads (Figure 2A). Bimodal distributions were seen in satellite regions, including a mode of increased coverage in the long read technologies and decreased coverage in the short reads, with a notable proportion of zero coverage regions as well. Different platforms exhibited different strengths and weaknesses of coverage as compared to the average of all other platforms (Figure 2B). BGISEQ500, HiSeq4000, and NovaSeq 2×150bp captured Alus most reliably;HiSeq 25000 replicates performed best in L1, L2, and Low Complexity regions, along with HiSeq X10 and NovaSeq 2×150bp;PacBio CCS and both NovaSeq chemistries performed best in satellite and Simple Repeat regions;and PromethION, alongside other long read platforms, performed best in telomeric regions. All-vs-all comparisons provide a more detailed profile of any one platform’s coverage capture versus any other (Figure S10), and every possible pairwise comparison is shown in Supplementary Data 1, exemplified by Figure S11.

**Figure 2:**
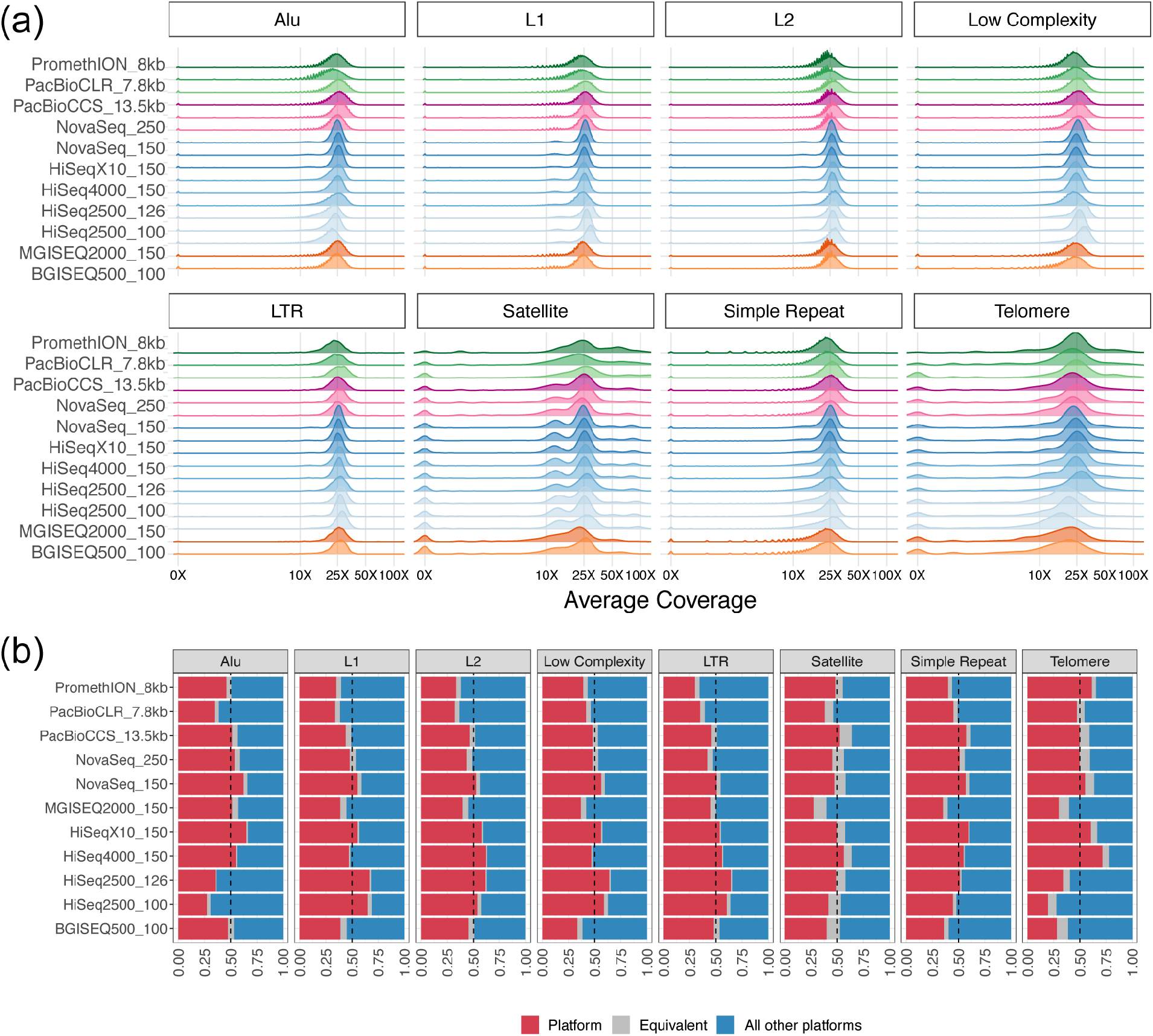
Distribution of genomic coverage across sequencing technologies for all replicates. Platforms are labeled with their mean read length. (a) Aligned BAMs were downsampled to 25x mean read depth, and the distribution of coverage of each locus in the UCSC RepeatMask regions was plotted. Platforms are listed alongside their mean read lengths. (b) Average genomic coverage of each platform against all other platforms, stratified by UCSC RepeatMask region, using the normalized 25x coverage alignment files.

### Rates of Sequencing Error

Rates of sequencing error were calculated for every human replicate, both genome-wide and in localized, difficult-to-capture regions of the genome, based on alignment of reads to the hs37d5 reference genome. Globally, reads were binned into windows corresponding to GC content from 0-100%. AT- (0-25%) and GC- (75%-100%) rich regions showed elevated rates of error, including nucleotide substitutions, insertions, and deletions, for all platforms analyzed (Figure 3A). In particular, Illumina and MGI platforms showed a propensity for substitution errors, while Genapsys (v1 chemistry) and long read platforms showed higher rates of insertion/deletions. Long read platforms showed elevated error rates at GC extremes in a similar manner to short read platforms, though it should be noted that most replicates have consistent GC composition (except for low coverage samples such as NextSeq replicates) and relatively few reads contain extreme GC bias (Figure S4). In the PacBio CCS replicate, error rates in middling GC windows are in line with the rates found in short read platforms. The low depth long read sequencing replicates from Flongle and R9.4 flow cells on the MinION MK1b showed very similar error profiles to more deeply sequenced replicates, with the exception of loss of reads at high/low GC content windows (0-25% and 75-100%), likely indicating that a certain threshold of sequencing is required before these extreme read compositions can be achieved.

**Figure 3:**
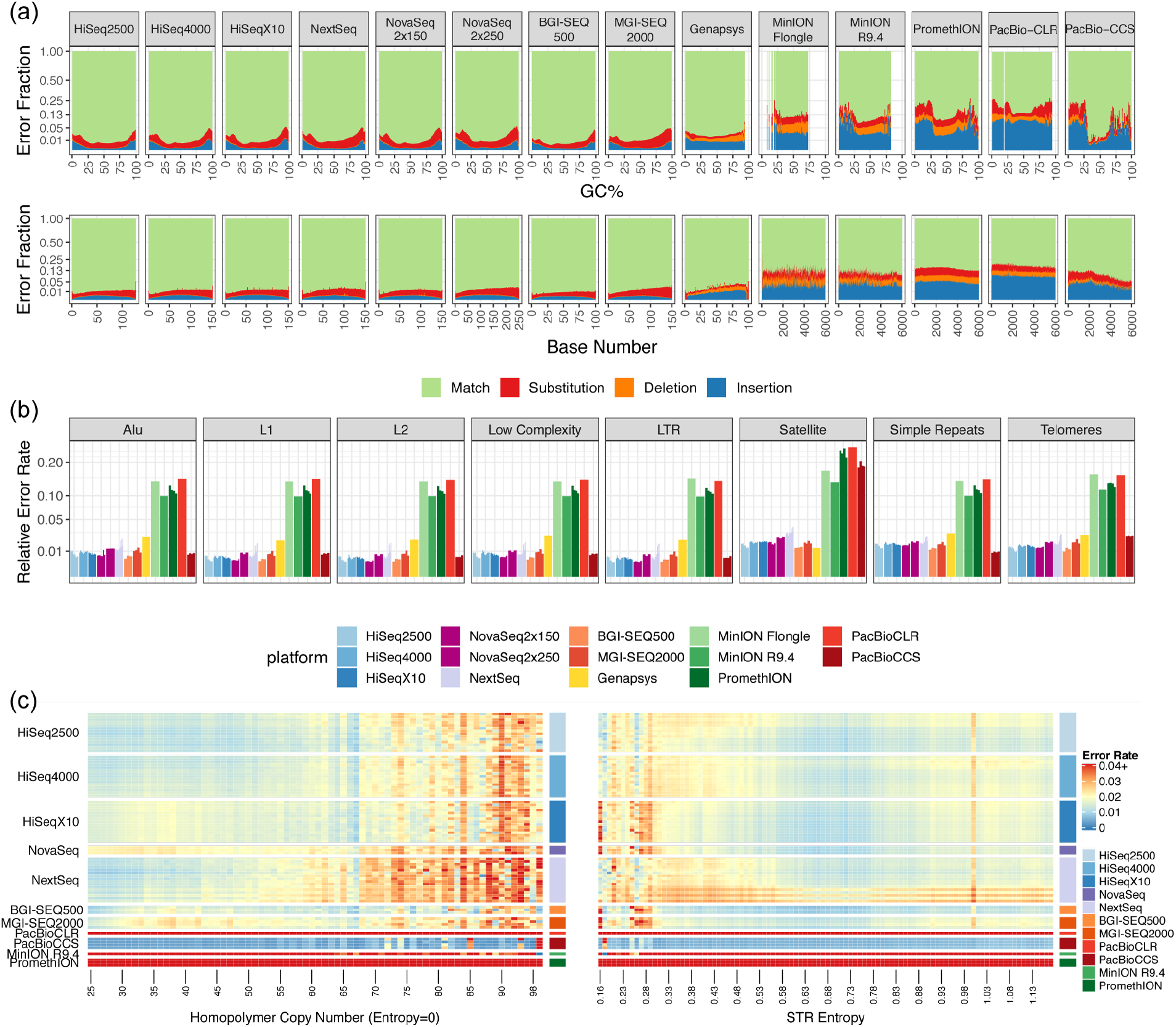
Rates of sequencing error per platform stratified by UCSC RepeatMask regions. (a) Proprortional error rates across GC windows and base number. Values at each window are averaged across all reads from all replicates, Missing columns are indicative of zero reads containing that GC composition. For long read platforms, read length is capped at 6kbp. Y-axis is plotted as square root. (b) Bar plot showing total average error rate within each region type. Individual replicates per platform are shown as separate bars, Y-axis is plotted as square root. (c) Error rate in homopolymer (n=72.687) and short tandem repeat (n=928.143) (STR) regions. On the left, true homopolymers are shown at increasing copy number. On the right, STRs are plotted by entropy, a measure of complexity of the motif. For PacBioCLR, MinION R9.4, and PromethION replicates, error rates consistently ranged between 0.10 and 0.15 per window, but the scale for the error rate is capped at 0.04 in order to show the differences within short read platforms.

Across the length of a given read, all platforms showed very consistent error profiles (Figure 3B), including long reads and short reads, as well as the novel NovaSeq 2×250bp chemistry. The MGI2000 platform showed some elevation of substitution errors towards the final cycles of each read. Although showing some fluctuation, owing to the smaller pool of reads and thus lower power of sensitivity, the Flongle and 9.4 flow cells on the MinION performed comparably to the 9.4 flow cell on the higher throughput PromethION platform. The PacBio CCS showed decreased error rate further into each read, which may reflect fidelity of accuracy at the ends of the subset of reads that extend that far, which may not have benefitted from the multiple rounds of coverage per read afforded in circular consensus sequencing within shorter reads.

Reads were stratified by the UCSC RepeatMask reference regions(Haeussler et al. 2019), split into different repeat classes (Alu, L1, L2, LTR, Satellite, simple repeats, and telomere regions). Error rates were very consistent within each platform across each repeat context (Figure 3C). Short read technologies, including Illumina platforms, MGI platforms, and the Genapsys, all ranged between 1-3% error rate across all repeat types, As expected, errors were more elevated in Oxford Nanopore platforms, although the Flongle flow cell performed comparably to the MinION and PromethION standard-use R9.4 flow cells. The PacBio CCS matched the performance of, and in some cases outperformed, the short read platforms, One exception was within the satellite regions, where PaBio CCS returned error rates more in line with other long read platforms in the 15-20% range, though it should be noted that satellites are highly variable between individuals and are not considered to be well represented in reference genomes.

Reads were then stratified against the UCSC Table Browser Simple Repeat Schema as defined by Tandem Repeat Finder [22]. Repeats were split into true homopolymers (stretches of poly-N in the reference genome) and other short tandem repeats (STRs), ordered by their entropy, a measurement of complexity of the STR motif (Figure 3C). The NovaSeq and MGISEQ2000 showed elevated error rates (2-4% per window) in the short homopolymer repeat stretches compared to other short read platforms (1% per window), while the HiSeq2500, HiSeq4000, HiSeqX10, and NextSeq showed elevated error rates in longer homopolymer repeat regions. The PacBio (CLR), MinION, and PromethION platforms showed uniform elevated rates (range 0.10-0.15) across all STR types and windows, while the PacBio CCS was robust against errors in nearly all windows. Although not included due to shallow coverage, the Flongle flow cell and Genapsys data showed comparable performance in the few STRs that each platform captured with confidence.

### SNV and INDEL Detection

In the previous section we detailed the error characteristics of each sequencing platform. It is also relevant to investigate consensus errors in the context of nucleotide variant detection compared to the human reference genome. Variant call sets, including short nucleotide polymorphism (SNPs) and insertion/deletion (INDEL) events, were characterized against the Genome in a Bottle (GIAB) high confidence truth set (v3.3.2 [20] for every replicate of the Ashkenazi Son (HG002) genome with adequate depth of coverage, and each alignment normalized to mean 25x coverage. By comparing these downsampled call sets against the GIAB high confidence set, we could calculate the sensitivity and specificity of SNP/INDEL detection for a given HG002 replicate for each platform (Figure 4A). All short read platforms exhibited a trade-off between precision and recall along the gradient of genotype quality cutoffs supplied by RTG vcfeval analysis, with the highest balance point occurring at around GQ=40, and falling off with greater thresholds (Figure S5). Within SNPs, BGI/MGISEQ replicates achieved the highest scores, followed by NovaSeq 2×250bp. NovaSeq 2×150bp, HiSeq 2500, HiSeq X10, and HiSeq4000. Among INDELs, BGI/MGISEQ replicates were roughly equal to NovaSeq replicates, outperforming the other Illumina platforms, though all platforms achieved >99.5% precision and reached close to 100% recall with at least one GQ cutoff. The same could not be said for long read PacBio/Nanopore platforms, which are not yet precise enough to capture these variants with the same level of fidelity (Figure S6). All platforms exhibited low power of detection of variants within repeat regions filtered for GIAB high confidence calls, though it should be noted that each context had relatively few high confidence calls, as compared to the global distribution (Figure S7). Within the exome, performance was more variable, with all Illumina and replicates achieving relatively high sensitivity and specificity scores, BGI/MGI slightly lower for INDEL detection, and long reads and Ion Torrent platforms with less power to detect GIAB high confidence exomic variants (Figure S9). Mendelian violations were captured via Mendelian Violation Detection as described in a previous publication ([23]) as a further measure of concordance among the family trio, as well as to characterize potential de novo germline mutations in the Ashkehanzi son cell line. The overall rate of Mendelian impossibility (i.e. both parents were homozygous reference while the son showed a homo- or heterozygous variant) ranged among platforms, from as low as 0.16% of variants in GIAB defined high confidence regions in the MGISEQ2000 trio and 0.24% in the BGISEQ500 trio, to an average of 1.2% among HiSeq instruments, to 17% in the one Nanopore trio, reflecting current limitations of the platform (Supplementary Table 5). Intersection of Mendelian violation regions shows most are platform-specific, rather than shared globally, even when stratified by different SNP/INDEL classes (Figure S8).

**Figure 4:**
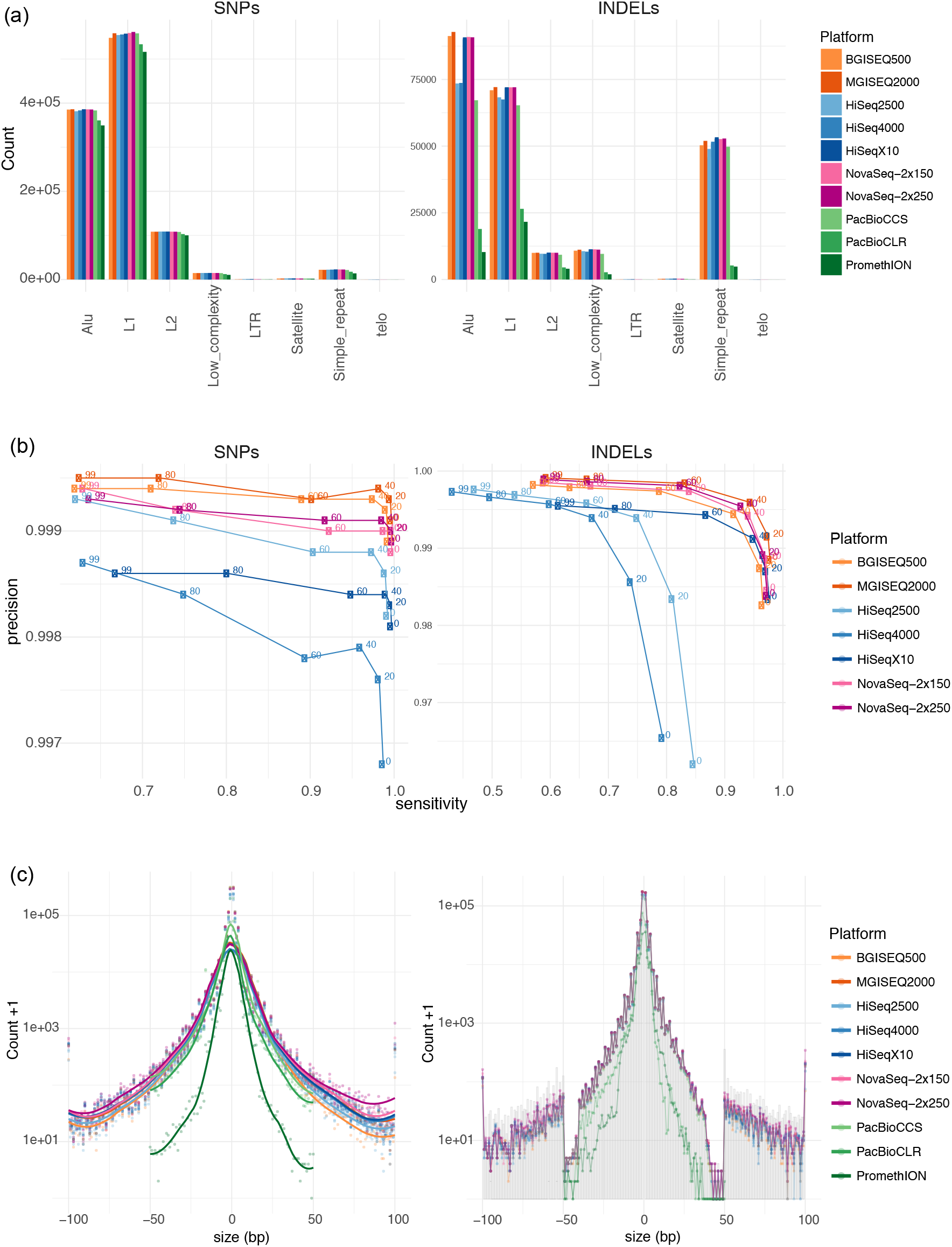
Validating short nucleotide polymorphisms (SNPs) and insertion/deletion (INDEL) events from short read datasets against the Genome in a Bottle (GIAB) high confidence truth set as determined by RTG vcfeval. One replicate of the HG002 genome was chosen per platform, and the call sets were generated from normalized 25x coverage alignments. (a) Counts of variant calls per UCSC RepeatMask context, colored by sequencing platform. (b) Precision and sensitivity of calls per platform against the Genome in a Bottle high confidence truth set. Genotype Quality was used as a Receiver-Operator Curve score, and values at different cutoffs are plotted. (c) Distribution of sizes of INDELs capture per sequencing platform. On the left, genome-wide capture. On the right, filtered by true positives in the GIAB truth set.

The variants captured were further stratified by the UCSC RepeatMasker classes, and some differences could be detected between sequencing platforms (Figure 4B). Within SNP calls, all short read technologies and PacBio CCS performed comparably, capturing slightly more variants than PacBio CLR and PromethION in Alu, L1, L2, and Simple Repeat regions. More variability was seen in INDEL regions, where BGI/MGI had the advantage in Alu regions, as well as L1s alongside HiSeqX10 and NovaSeq platforms, outperforming the HiSeq 2500/4000 replicates. In Alus, L1s, L2s, and Simple Repeat regions, the long read platforms, including PacBio CCS, had difficulty capturing INDEL events. No platform successfully captured telomeric variants, likely due to the very few telomeric high confidence regions available for benchmarking (n=405).

INDEL calling was also analyzed as a function of the size of the polymorphism event (Figure 4C). Very few INDELs could accurately be called beyond a size of 100bp, so INDEL counts at sizes >= 100bp were aggregated as one value. The NovaSeq 2×250bp chemistry outperformed other platforms across all INDEL sizes, reflecting the power of increased read length with per-base accuracy. Other Illumina and BGI platforms performed comparably, followed by PacBio CCS, and then CLR and PromethION, both in terms of global INDEL capture and with respect to the GIAB high confidence true positive set for HG002. We also compared indel size distribution among high confidence true positive INDEL calls only, as defined by the GIAB (merged SNV/INDEL v3.3.2 and SV Tier 1 truth sets) and noticed similar trend across platforms as seen for the genome-wide distribution with NovaSeq 2×250bp chemistry outperforming the rest. A large drop-off in capture with INDEL sizes approaching 50 bp reflects the strict filtering strategy applied by the GIAB to create the SNV/INDEL truth set, which includes events up to 50bp only. Due to limitations of the SNV caller, no long read platform could accurately detect INDELs beyond 50bp, requiring different analysis methods for larger, structural events. A full compilation of pairwise INDEL capture comparisons between each downsampled library is available as Supplementary Data 2.

### Structural Variant Detection

We further investigated the ability to reproducibly detect and characterize structural variations (SVs). This was achieved using a multi-caller consensus approach, combining the outputs of individual SV callers and filtering for consensus using SURVIVOR [24] (see Methods), and applied only to HG002 samples, as this was the cell line with greatest representation across platforms. An average of 12,602 SVs were identified across 27 samples from HG002. The majority (80.95%) of these SVs overlapped with the GIAB high confidence region set ([20]). The majority of events called were deletions (8169 SVs), followed by translocations (2828 SVs), duplications (889 SVs), inversions (705 SVs) and insertions (10 SVs), which matched the expected distribution both in terms of size and event type [25]. The high number of translocation calls were ignored, as they often represent false positives [26]. The filtered dataset contained an average of 10,201 SVs per replicate. Of these, 27.58% SVs (2814) were deletions and insertions that overlapped with the GIAB Consortium HG002 SV call set (v0.6). The overall performance metrics among all datasets, as well as the distribution of SV calls per sample, showed high concordance between samples (Figure 5A). No significant correlation was observed between the total number of SVs and an increase in the average coverage or insert size across the datasets. However, for true positive SV calls, positive correlations were observed with respect to coverage (mean: 39.10, cor: 0.76, p-value: 3.26e-07, standard deviation: 67.78, cor: 0.72, p-value: 2.41e-06), insert size (mean: 348.58, cor: 0.60, p-value: 0.00023, standard deviation: 123.74 cor: 0.55, p-value: 0.00085) and read length (mean: 141.67, cor: 0.63, p-value: 8.23e-05, standard deviation: 0.00012, cor: −0.26, p-value: 0.14).

**Figure 5:**
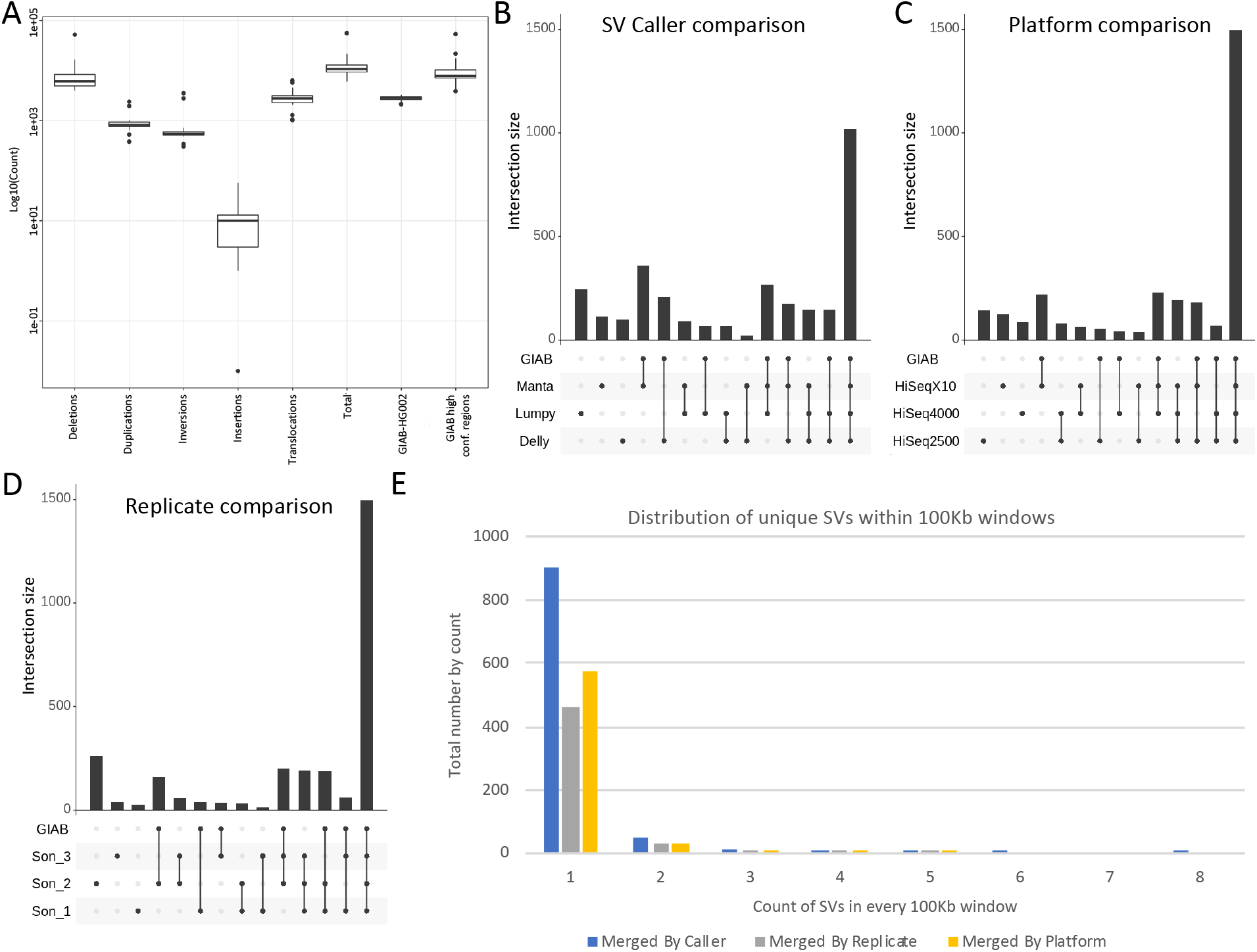
Variability of HG002 structural variant (SV) capture across platforms that had more than one replicate with adequate coverage (HiSeq2500, HiSeq4000, HiSeqX10). (a) Number of SVs across sequencing reactions, including deletions, duplications, inversions, insertions, translocations, total number of events called, SVs that overlap with the Genome in a Bottle (GIAB) HG002 SV set, and SVs that overlap with the GIAB high confidence regions. (b-d) Variability attributed to caller algorithms (b), sequencing platforms (c), and laboratories (d), visualized with UpSet overlaps. (e) Distribution of single support (unique) SVs in 100KB windows across different stratification strategies.

To identify the cause of the discrepancy between the number of SVs called among the samples, we investigated reproducibility among the different sequencing and analysis steps: SV callers (Figure 5B), platforms (Figure 5C), and replicates (Figure 5D). Overall, the SV callers themselves contributed the most to individual variability (697 SVs, 43.86%). The second most variability came from the sequencing platforms (374 SVs, 23.54%), followed by intra-lab replicates (268 SVs, 16.87%). The majority of consensus SV calls overlapped the GIAB HG002 truth set (89.96% for SV callers 83.16% for platforms, and 84.33% for replicates). Thus, interestingly, false negatives were predominantly observed (i.e. captured in one or more but missed by others), instead of the expected false positive SV calls. To identify potential variability biases in the characterization of SVs, we further investigated the SVs that could be assigned solely to a certain analysis step. SV call sets did not show any clustering in a particular region of the genome and seemed to be distributed throughout (Figure 5E).

Additionally, for SV callers, it is interesting to note that the majority of SV calls that were specific to Delly or Manta are in fact true positives. In parallel to this, it is evident that most false positives from SV caller variability are attributed to SV calls from Lumpy, followed by Delly, and then Manta (Figure S14). The summarized results for all strategies in terms of false positive, negative and true positive were aggregated in Supplementary Table 5. Among platforms, the HiSeqX10 returned the largest number of SVs (2552 SVs), followed by HiSeq4000 (2260 SVs) and HiSeq2500 (2256 SVs). The HiSeq2500 was observed to produce the largest number of unique false positive SVs (143 SVs), followed by HiSeqX10 (127 SVs), and HiSeq4000 (84 SVs). Interestingly, 63.29% (219 SVs) of unique HiSeqX10 SVs are false negatives compared to HiSeq4000 33.33% (42 SVs) and HiSeq2500 27.41% (54 SVs) (Figure S15). For replicates, 15.67% of unique replicate SVs were false positives that are not concordant with the GIAB HG002 truth set. Overall, 91.78% of non-unique SVs overlapped with GIAB-HG002 SV call set, indicating a smaller number of false positives and high concordance between the replicates (Figure S16).

### Bacterial Genome Capture

In addition to the relatively GC-balanced human genome, analysis of sequencer performance at high and low GC content genomes was evaluated. A variety of bacterial isolates, as well as a combination metagenomic mixture of ten bacterial species, were sequenced across an array of platforms in order to assess reproducibility of genomic capture with variable GC content, Gram stain, ecology, and physiology. In particular for the metagenomic pool (ATCC MSA-3001 mix), taxonomic composition was found to be quite variable both within and between platforms, with MiSeq and Genapsys showing the most reproducible and balanced compositions. Flongle and R9.4 flow cell data on the MinION indicated similar but imbalanced compositions, and the Ion S5 and Personal Genome Machine (PGM) showing more variability and taxonomic imbalance (Figure 6A). Although the ATCC MSA-3001 ten strain mix is designed to contain 10% representation of each bacterium, the observed composition of each taxon was highly variable. Generally, AT- and GC-rich genomes were less represented, with balanced GC content genomes overrepresented, and generally Gram positive genomes were better captured than Gram negative (Figure 6B). Error rates were also calculated per genome per platform, and showed some taxon-specific bias, with P. fluorescens having the highest substitution rate, followed by the ten strain Pool, and then *S. aureus* and *E. coli* (Figure 6C). Platform-specific bias was also detected, with individual Flongle data and native barcoded 9.4 data generated on the MinION MK1b exhibiting higher insertion/deletion rates, MiSeq and Genapsys showing almost exclusively substitution errors, and the S5 and PGM showing low overall error but biased towards insertion/deletions. Unlike the human trio, there was not a clear spike of sequencing error at extreme GC windows. Both Flongle and R9.4 flow cells showed narrower windows of GC% capture, owing again to the shallower depth of sequencing with longer reads that tend to average out in middle GC ranges.

**Figure 6:**
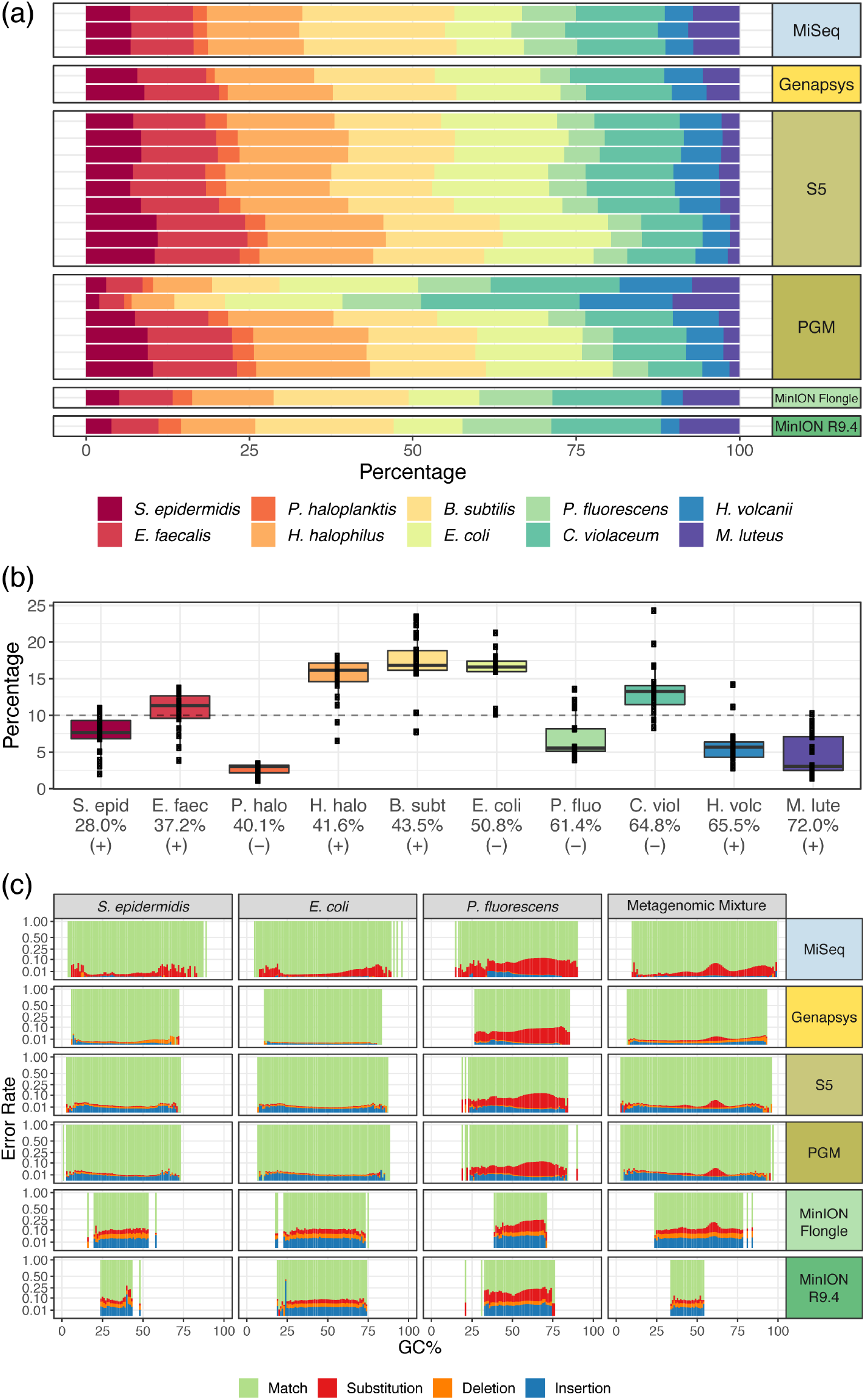
Reproducibility of sequencing of bacterial genomes. (a) Distribution of taxonomic assignment of strains present in the metagenomic mixture, per replicate per sequencing platform. (b) Distribution of presence of each taxon across replicates, ordered by GC content and with Gram stain indicated. (c) Error rates across each individual strain and the metagenomic pool, per sequencing platform.

## Discussion

The ABRF-NGS Phase II study is a comprehensive DNA-seq resource, providing multiple inter- and intra-lab replicates of whole-genome and exome sequencing across multiple established and emerging platforms. The data comprise well-characterized and publicly accessible human genomes that have become cornerstones of genomics research, as well as bacterial genomes that span a diversity of genome sizes and nucleotide compositions. Analyses of this multi-dimensional dataset reveal characteristic differences between sequencing platforms, as well as degrees of variability and reproducibility within replicates from the same platform. Together, these provide a valuable resource for the genomics research community.

Reproducibility analyses within this study revealed the degree to which performance can vary within and across sequencing sites and platforms, controlled for library preparation and bioinformatic pipelines. Several factors contributed to variability within these data. GC content was a major factor, as elevated error rates were correlated with extreme compositions, both in human and bacterial genomes. Genomic context played an even larger role, as all analyses stratified by UCSC RepeatMask repetitive regions shed light on differences between sequencing technologies. These difficult regions of the genome (referred to as the “Classes of Evil”) revealed the relative performance strengths of each platform, such as outsized capture of Alu regions by BGI, highest fidelity of variant capture by the NovaSeq 2×250bp chemistry, and Nanopore detection of telomeric regions. Together, these results provide a reference for which platform to choose given a particular region of interest for a genomics study.

Depth of coverage is a major contributor to the accuracy of any sequencing product, and although deeper sequencing provided more robust variant detection and reduced sequencing error, particularly in short read datasets forthe human genomes, deeper sequencing did not ameliorate mappability in a proportional sense for certain bacterial species. In particular, for *Pseudomonas fìuorescens*, some replicates with triple the coverage of others showed roughly 55% of reads not mapped to the reference *P. fìuorescens* genome, indicating the possibility that the strain was not *Pseudomonas fìourescens* as originally purchased or a culture collection error. The reference genome and ATCC designation for this strain of *P. fìuorescens* is under review and needs further investigation. Ultimately, the ability to map a pool of reads is limited by the quality of the reference genome, and these data show a clear need for improved assemblies for commonly studied bacterial references.

The distribution of reads in a DNA-seq reaction was seen to be highly reproducible when sequencing an individual genome, across all three members of the Ashkenazi Trio. Across laboratories and sequencing platforms, error rates were consistent per platform, including in repetitive and low complexity regions. In particular, emerging platforms from BGI, Genapsys, and Oxford Nanopore performed comparably to well-established platforms, providing promising results as the genomics landscape continues to grow and diversify. More complex metagenomic samples were less consistent, showing compositional bias and elevated variance of normalized coverage, indicating a challenge for future metagenomic studies. Notably, all platforms were able to identify all strains in each mix, and showed robustness in identifying the presence of taxa in metagenomic samples. At the same time, the degree of variability within metagenomic capture remains a clear confounding variable that should be tracked and examined in future work, along with the other components of metagenomics analysis [27].

The Genome in a Bottle consortium has provided benchmark variant call sets for the RM 8392 references examined within this study, as well as other standard, well-characterized genomes [19]. Our results complement that work with a more in-depth intra-platform comparison leveraging multiple replicates per platform, providing an unbiased evaluation of current and emerging sequencing technologies. Building on the resources provided by GIAB, the Global Alliance for Genomic Health (GA4GH), and UCSC, this study provides a community resource with many intra-platform replicates of DNA-seq data to help improve future genomics research efforts. Specifically, these data can assist sequencing facilities in self-assessment of the performance of their local platform. The results provide independently generated benchmarks against which new protocols, chemistries, instruments, and bioinformatics can be judged during development. The data generated by this study are freely available on NCBI, and can act as a baseline as sequencing technologies continue to evolve rapidly. Finally, these findings can inform the evolution of new best practices in sequencing and analysis, serving as highly characterized reference material data designed to support a variety of genomic analyses and methods, which will be essential as new methods emerge.

## Methods

The complete set of code used to analyze and visualize all data is available as Supplementary Data 3.

### Human DNA

DNA from cell lines derived from a family trio in the Personal Genome Project (PGP) are distributed as National Institutes for Standards and Technology (NIST) reference material RM 8392, which serves as source material for genomic DNA sequencing. These DNA samples were developed for the Genome in a Bottle (GIAB) consortium to provide a set of highly characterized standards for genetic analyses, and are approved for all research uses under the terms of the PGP. Standardized human genomic DNA samples were obtained by a single laboratory from NIST, and whole genome sequencing (WGS) libraries were prepared at a single laboratory site (HudsonAlpha Institute for Biotechnology, Huntsville, AL), then distributed to individual laboratories for sequencing.

In a few cases, libraries were not prepared at HudsonAlpha but rather at the sequencing core facility, although libraries were prepared using the same NIST stock and library synthesis kits as at HudsonAlpha. This was the case for the NovaSeq platform replicates, where one set of replicates was prepared and sequenced by Illumina on a NovaSeq 6000, while another set of replicates was sequenced also on a NovaSeq 6000 using paired-end 150 bp and a novel 250 bp reaction at a second lab. Single sets of replicates from Genapsys, BGISEQ500, and MGISEQ2000 were generated by one respective laboratory for each case. Oxford Nanopore PromethION R9.4 replicates were prepared using the PCR-free Ligation Sequencing Kit (LSK109) at one site, while the MinION libraries were prepared for the Flongle using the PCR-free Ligation Sequencing Kit (LSK109) and combined the with native barcoding kit for the R9.4 flow cells at a different core lab, Finally, all replicates of PacBio Circular Consensus Sequencing (CCS), Continuous Long Reads (CLR), and one Oxford Nanopore PromethION replicate were downloaded from the public repository generated by the Genome in a Bottle (GIAB) Consortium and hosted by the National Center for Biotechnology Information (NCBI).

### Microbial DNA

Microbial reference gDNA was prepared from bacteria obtained from the American Type Culture Collection (ATCC-Manassas, VA). Pure agar cultures were grown to early log phase and harvested prior to gDNA extraction using the Omega Metagenomics DNA kit (Omega BioTek Norcross GA, M5633-00). Briefly, cell mass was resuspended in dPBS pH 7.5 and digested with Metapolyzyme (MAC4L Millipore Sigma, St. Louis, MO) for 8 hours before dual detergent lysis with CTAB and SDS and farther lysis and clean up using phenol chloroform + isoamyl alcohol and RQ magnetic beads. DNA was evaluated using Qubit spectrofluorometry (Thermo Fisher, Waltham, MA), Agilent Bioanalyzer 2100 (Santa Clara, CA), RTqPCR (Applied Biosystems, Foster City, CA), and Nanodrop spectrophotometry (Thermo Fisher). Sequencing QC was performed using both Sanger sequencing of the entire 16s rDNA (Primer 27f and 1492r) as previously described (Innis et al, 2012), and whole genome sequencing using Oxford Nanopore and Illumina sequencing. DNA for the 10 species combined mixtures was combined as an equimolar pool at approximately 10% each. This gDNA material is deposited at ATCC as product MSA-3001 and is publicly available.

### Library Preparation

For non-exome libraries, each laboratory used the Ion Xpress Fragment Library kit (part 4471269) per the manufacturer’s protocol, using 100ng of input DNA. For Ion Ampliseq exome sequencing, DNA was amplified through a massively multiplexed PCR reaction to create the library following the Ion Ampliseq Exome protocol (kit 4489061).

All libraries were templated onto beads (Ion PI Hi-Q Template OT2 kit A26434 for Proton, Ion PGM Hi-Q OT2 200 Kit A27739 and S5 (part A27751 for bacterial libraries and A27753 for Exome libraries). The Exome libraries were sequenced on either the Ion S5 or Ion Proton instruments used standard 200bp chemistries and protocols (Proton kit A26771, S5 kit A27753). The bacterial libraries were sequenced on the Ion PGM or Ion S5 using 400bp chemistries (Ion PGM Hi-Q Seq Kit A25592 and Ion S5 kit A27751).

For the Genapsys GS110 sequencer, library synthesis was performed using a two step approach by first synthesizing a standard NGS library followed by a Genapsys clonal amplified library. 100 ng of microbial gDNA was fragmented using Covaris S2 instrument to a mean size of 250 bp and used as input to the NEBNext Ultra II kit (E7645 New England Biolabs Ipswich, MA) and checked for quality using the Agilent Bioanalyzer 2100 and Qubit spectrofluotometer. This NEBNext library was used as input to the version 1 chemistry of the fully manual Genapsys clonal amplification kit (1002000) which required 1.0 x 108 molecules (33 pol) before hybridizing to the G3 electronic sequencing chip (1000737 Genapsys Redwood City, CA) and sequencing on the GS111 Genius Sequencing Platform.

Microbial gDNA was prepared for Nanopore sequencing using two library methods. For the Flongle flow cell runs, the direct ligation sequencing library kit (LSK-109 Oxford Nanopore UK) was used on individual bacteria and sequenced on dedicated flow cells. For the R9.4 flow cells runs, the individual bacteria strains as well as the 10 species mix was prepared using the LSK109 method with the native barcoding expansion kit (EXP-NBD104) and combined into one final library pool and sequenced together on a single flow cell, This ligation sequencing method is a non-PCR based library method that allows for direct sequencing of native DNA. Briefly, gDNA is “repaired” using the NEBNext FFPE DNA Repair reagents (M6630, New England Biolabs Ipswich Ma) followed by dA-tailed using the NEBNext End Repair/dA-tailing module, and ligated to nanopore specific sequencing adapters, Sequencing was performed immediately after library synthesis.

TruSeq PCR-Free libraries were prepared according to manufacturer’s protocols for Illumina libraries. The high MW genomic DNA from NIST was fragmented using an LE Series Covaris sonicator (Woburn MA) with a targeted average size of 350 bp. Libraries were then synthesized at HudsonAlpha Biotechnology Institute robotically using 1 ug of DNA. Library quality was evaluated by Qubit quantification and Agilent Bioanalyzer 2100. After passing QC, libraries were shipped to different sites (core facilities) for sequencing. Twelve sites participated in sequencing, and twelve independent libraries from the mother and father were generated and distributed. For the child, thirty six independent libraries were synthesized and three of each were distributed to the twelve sites. This latter triplicate helps serve as a control for technical variability in library generation.

### DNA Sequencing

For each of the Illumina HiSeq 2500, 4000, X10, and NextSeq platforms, WGS was performed on each of the human genomes across multiple laboratories (not necessarily the same labs in each case). AmpliSeq exome panels (Thermo Fisher) of RM 8392 genomes were prepared and distributed in the same manner as WGS libraries.

TruSeq PCR-Free libraries were sequenced on the Illumina HiSeq 2500, HiSeq 4000, HiSeq X10, MiSeq, and NovaSeq 6000 with Xp loading. For the bacterial samples, libraries were created at each facility from the samples in the table above. Libraries were constructed using the Ion Torrent Ion Xpress Library Kit per standard kit protocols which uses enzymatic fragmentation for the library build. Libraries were run separately on the PGM (other than the pool) and were multiplexed on the Proton PI and S5 540 chips. Standard protocols were used for 400bp read lengths on the PGM and S5 520 and 530 chips. The bacterial libraries were run using 200bp reads on the Proton and S5 540 chip using standard protocols. The different read lengths were due to the availability of 400bp chemistry on the smaller chips for both PGM and S5 whereas the larger PI and 540 chips run 200bp chemistry. All libraries were run in triplicate (on different runs). All libraries were synthesized using 1ug of DNA.

The exomes were run only on the Proton PI and S5 540 chips because of the read numbers requirements. Libraries were created using the Ion Torrent Ampliseq Exome Ready kit. Briefly, the exomes are amplified in a massively mulitplexed PCR reaction and the resulting libraries are sequenced per standard sequencing protocols. Samples were run in triplicate with two samples per chip to accommodate read numbers needed for analysis.

For Genapsys sequencing, successful clonal libraries were loaded onto the G3 electronic sequencing chip according to manufacture protocol (GS111 User Guide 1000698 Rev C Oct 2019) following an initial priming step including buffer washes. The electronic flow cell was injected with 35ul of the sequencing bead library followed by 40ul of a DNA polymerase solution. Sequencing was initiated on the GS111 Genius sequencer and run for 48 hours to achieve 10-15 million reads of single end 150 bp data.

Oxford nanopore sequencing was performed using the Promethion sequencer for the human samples with the standard use R9.4flow cell. For bacterial genomes, the MinION MK1B sequencer was used with both Flongles and R9.4 flow cells. Flongles were injected with 20 fmol of each library on the with the slight modification of a 20% reduction of loading beads to increase Q-score performance. Sequencing was perfomed up to 48hrs. 9.4 flow cells were injected with 50 fmol of the pooled native barcoded library according to the manufactures example protocol (NBE_9065_v109_revJ_23May2018) and allowed to sequence for 72hrs.

### Alignment and Variant Calling

All short read datasets, including human and bacterial samples, were aligned and had variants called using Sentieon ([28]), except for exomic data which was aligned/variant called with Torrent Suite (see below). Long read datasets were aligned using minimap2 (v2.17) ([29]). Short nucleotide variants were called with Clair (v2) ([30]) while structural variants were called using a multi-algorithmic approach (Delly ([31]), Lumpy ([32]) and Manta ([33]), and validated with SURVIVOR [24]). Whole genome human samples were aligned against hs37d5, an integrated reference genome combining the GRCh37 primary assembly (including canonical chromosomes plus unlocalized and unplaced contigs), the rCRS mitochondrial sequence (AC:NC_012920), Human herpesvirus 4 type 1 (AC:NC_007605), and concatenated decoy sequences to improve variant calling, Indeed, approximately 5% of reads across sequencing platforms were captured by the decoy hs37d5 sequence, helping to filter out spurious, low complexity reads (Figure S3).

More specifically: Illumina-derived reads were aligned using bwa mem (bwa mem -M -R $readgroup -K 10000000 -t 4 $reference $fastqR1 $fastqR2), Sorting, deduplication, indel realignment, base quality score recalibration, and variant calling were performed using the DNASeq workflow within Sentieon build 201808.0329 with default parameters. LifeTech-derived reads were processed through the Torrent Suite version 5.10, including tmap mapall (tmap mapall -f $reference -r $input -n 20 -v -u -o 1 stage1 map4) for alignment and variant_caller_pipeline.py within tvc for variant calling, using default parameters. For human data, all reads for all platforms were aligned to the 1000 Genomes build of hs37d5 (GRCh37 plus decoy sequences). For bacterial data, all reads were aligned to genome builds of respective species derived from the NCBI Genome portal (Supplementary Table 3). Depth of coverage was calculated using mosdepth ([34]) with the -n flag.

Base quality distributions, insert size distributions, and GC bias metrics were calculated using default values within Picard v2.10.10-SNAPSHOT, Read mapping metrics, on-target mapping rates, species distributions in metagenomic mixtures, conversion of BAMs to FASTQs, BAM indexing, and BAM header alterations were performed using samtools v1.930. BAMs were downsampled to a normalized 25x coverage using the -s flag within samtools, with 0 as the leading seed, and the fraction calculated based on mosdepth-inferred depth divided to achieve 25x 3.1Gbp coverage.

Genomic intersection and low complexity region masking was done with bedtools v2.27.131. VCF statistics were summarized using vcftools v0.1.1532, and merging was done with bcftools v1.633. Variant allele frequency marix generation was done with bcftools using the −012 flag. UpSet plots were generated with the UpSetR package [35]. t-SNE analysis was done with the Rtsne library with the starting seed 42, perplexity of 20, theta of 0.5, and 2 dimensions. Heatmaps with colored annotation tracks were created using the ComplexHeatmap library. Error rates were calculated using BBtools (https://sourceforge.net/projects/bbmap/). The BAM from each sample was run through bbmap to filter into 100 sub-BAMs (one per GC window), with mappings to known variants from the Genome in a Bottle (GIAB) benchmark set filtered using bedtools intersect. Error rates were then inferred from the mhist table derived for each sub-BAM.

High confidence variants were analyzed using hap.py against the GIAB truth variant sets for each of the RM 8392 genomes (see below for RTG vcfeval analysis of SNPs and INDELs). Conversion of VCF data to allele frequency matrices, extraction of mapping and mismatch statistics across GC contents and normalized base position from BAM files, conversion of hap.py outputs into matrices, generating UpSet matrices, and homopolymer detection and SNP/indel assignment were all performed using Python 3.7.0 scripts, and all visualizations were performed in R 3.6.0.

All custom scripts and R markdown notebooks are available as Supplementary Material X.

### SNP and INDEL Benchmarking

Sensitivity vs. precision analyses were performed using the short nucleotide variant (SNV) and insertion/deletion (INDEL) call sets generated by each of the sequencing platforms (based on downsampled BAMs;see above) for the HG002 sample. Each replicate was compared against the GIAB SNV/INDEL HG002 truth set (v3.3.2) [17] using the Real Time Genomics (RTG) vcfeval tool [36]. True positive, false positive, and false negative calls were identified within the high confidence regions of the genome (as defined by GIAB) using Genotype Quality (GQ) as a Receiver-Operator Curve (ROC) score, stratified by variant type. For the three long read datasets (PacBioCCS, PacBioCLR, and PromethION) processed through the Clair pipeline, we only report sensitivity and precision value based on all SNVs/INDELs (equivalent of GQ >=0) because Clair does not output normalized Phred-scaled GQ scores.

To compare genome-wide INDEL size distribution across sequencing platforms and replicates of sample HG002, we used the RTG vcfstats tool with the option -allele-length, which outputs a histogram of variant length for a given VCF file. Vcfstats increments counts for each called allele, therefore a heterozygous call increases a count of an appropriate size bin by 1, while a homozygous alternate call increases a count by 2. To ensure that differences in distribution of INDEL sizes across platforms are not driven by the differences in mean coverage, we used INDEL call sets generated using alignment files downsampled to mean coverage of 25X.

In addition to comparing the distribution of INDEL sizes genome-wide (i.e. including all INDELs called by the SNV/INDEL Sentieon or Clair pipeline), we also restricted the analysis to high confidence genomic regions, as well as high confidence true positive INDEL calls, as defined by the GIAB for sample HG002 [17, 20]. Due to a filtering strategy that GIAB applied when generating their benchmarking truth sets, the SNV/INDEL truth set (v3.3.2) includes INDELs only up to 50bp in length, whereas the SV truth set (Tier 1) includes SVs starting at 50bp. Because the Sentieon SNV Haplotyper Caller generates INDEL calls larger than 50bp, we merged the GIAB SNV/INDEL and the SV truth sets (as well as high confidence bed files) and identified true positive calls across the entire size range. True positive calls made by each platform were again identified using the RTG vcfeval tool.

To facilitate a more detailed comparison, we also generated genome-wide, as well as high confidence true positive pairwise comparisons of INDEL size distributions for all possible pairs of sequencing platforms and replicates of the HG002 sample, stratified by shared and unique INDEL calls. Shared and unique INDEL calls for each pair of datasets were identified using the RTG vcfeval tool by treating one of the datasets as a truth set and the other as an evaluation set. High confidence true positive subsets were identified using the merged GIAB SNV/INDEL and SV truth set, as described above.

To compare numbers of SNV and INDEL calls that fall within different classes of repetitive and low complexity regions of the genome across all sequencing platforms we first restricted the analysis to the HG002 true positives that match the GIAB high confidence calls. We then annotated the true positive SNV and INDEL calls from each platform using the UCSC RepeatMask BED files.

### Structural Variant Detection

The aligned short reads were analyzed using Delly [31] (v0.8.2), Lumpy[32] (v0.2.13) and Manta[33] (v1.4.0), each with default parameters. The SV call sets generated per sample were merged using SURVIVOR[24] (v1.0.7) using the following parameters: “1000 2 1 0 0 0”, which requires a maximum of 1kbp allowance on the start or stop breakpoint;that the SVs to be merged are of the same type;and that at least two out of the three callers need to agree on a SV to keep it. Overlaps with GIAB high confidence regions (v.0.6) were established using bedtools (v2.29.2).

The long-read datasets were aligned using minimap2 (v2.17) [29] using the following parameters: -MD, -a, and -x map-ont for Nanopore replicates. For PacBio replicates, -x map-pb was used instead, and flag -H for homopolymer-compressed k-mer representation. The human genome assembly (hg37) downloaded from Ensembl was used as a reference, with removal of all decoy and alt contigs, This reference matches the non-alt or decoy contigs within hs37d5. The aligned BAM files were sorted and indexed using samtools (v1.9) [37]. Subsequently, Structural Variants were identified using Sniffles (v1.0.11) [26] based on the BAM files from the previous mapping steps. Sniffles was run with a minimum requirement of two reads (-s) to identify SVs, and –num_reads_report −1 –genotype –report_seq parameters were used.

Clair [30] (v2) was used with default parameters and a two read minimum to identify short nucleotide variants (SNVs). The appropriate model was chosen depending on the sequencing technology. Subsequently, the SNV call sets were filtered using RTG [36] based on the hg37 truth SNV set from GIAB.

## Acknowledgments

We thank Illumina and ThermoFisher for providing reagents allowing the study to take place. We also thank NIST for providing the Genome in a Bottle DNA samples necessary to carry out the study. We acknowledge the HudsonAlpha Institute of Biotechnology for expert assistance in Illumina DNA library preparation. The Association of Biomolecular Resource Facilities also provided funding, logistical support and project oversight. We are particularly grateful for the assistance provided by multiple core facilities spending their own time and resources in order to participate in this research. We would like to thank the Epigenomics Core Facility and Scientific Computing Unit at Weill Cornell Medicine, as well as the Starr Cancer Consortium (I9-A9-071) and funding from the Irma T. Hirschl and Monique Weill-Caulier Charitable Trusts, Bert L and N Kuggie Vallee Foundation, the WorldQuant Foundation, The Pershing Square Sohn Cancer Research Alliance, NASA (NNX14AH50G, NNX17AB26G), the National Institutes of Health (R25EB020393, R01NS076465, R01A1125416, R01ES021 006, 1R21A1129851, 1R01MH117406),the Bill and Melinda Gates Foundation (OPP1151054), TRISH (NNX16A069A:0107, NNX16A069A:0061), the Leukemia and Lymphoma Society (LLS) grants (LLS 9238-16, Mak, LLS-MCL-982, Chen-Kiang) the Alfred P. Sloan Foundation (G-2015-13964). Certain commercial equipment, instruments, or materials are identified to adequately specify experimental conditions or reported results. Such identification does not imply recommendation or endorsement by the National Institute of Standards, nor does it imply that the equipment, instruments, or materials identified are necessarily the best available for the purpose. FJS and MM are supported by NIH (UM1 HG008898).

## Author Contributions

C.E.M., S.W.T., C.M.N, D.A.B. conceived and designed the study. C.E.M., S.W.T., Z.H., W.F., G.S.G., S.L.,P.L., D.W. implemented the protocols. J.M.Z., W.E.C, M.B.B, and G.N assisted with analysis design. J.F. aggregated and processed data, led data analysis and figure generation, and wrote the manuscript. W.E.C, M.B.B., G.N., M.M.K, M.M., S.T performed data analysis, figure generation, and manuscript editing. The ABRF-NGS Study members contributed to the design and execution of the project.

## Data Availability

The genome sequences in this study are available as EBV-immortalized B-lymphocyte cell lines (from Coriell) as well as from DNA (from Coriell and NIST). All data generated within this study from these genomes are publicly available on NCBI Sequence Read Archive (SRA) under the BioProject PRJNA646948.

## Supplementary materials

### Variability of Structural Variants

#### Variability among Callers

The variability of structural variant (SV) calling is impacted by multiple factors, including caller algorithm, sequencing platform, biological replicates (see Supplementary Table 6). We analyzed the variability observed among the data sets in greater detail highlighting the false positive (i.e. artificial SV calls) vs. false negative (i.e. often missed SV calls) rates per variability. For this we used the GIAB high confidence regions and SV call sets 6.

For HG002, we investigated the variability attributed to SV callers while stratifying for variability from platforms (HiSeqX10, HiSeq2000 and HiSeq4000), replicates, and centers. To stratify for other factors, we merged across platforms, replicates, and then centers, requiring that SVs be concordant at each step between the respective SV call sets (Figure S14A). SURVIVOR was used for concatenation at each step, requiring SVs to be of the same type, have a pairwise overlap smaller than 1000bp and the length of the SV > 30bp. In the last step, we performed a union merge using SURVIVOR across the different SV callers (Manta, Lumpy, Delly) requiring SVs to be larger than 50bp. The SV call set was then filtered using the GIAB high confidence regions and compared to the SV calls from GIAB (Figure S14B). After stratification for other sources of variability and filtering, the resulting set had a total of 3.017 SVs that showed variability due to the SV callers. The majority of these SV calls are deletions (70.73%), most of which are between 100-1000bp in size (Figure S14C). Other SVs are spuriously called including translocations (12.46%), insertions (7.36%), duplications (5.17%,) and inversions (4.28%). We further filtered the SV call set to only include deletions and insertions to a total of 2.356 SVs. This made the SV call set comparable to the GIAB HG002 Truth Set and eased the interpretation of the majority of SVs observed to be variable due to the SV caller.

Next, we investigated the overlap with GIAB SV calls to identify potential false positives (i.e. Identified by a SV caller but absent in GIAB HG002 Truth Set) vs. false negatives (i.e. missed by one or more SV callers but present in GIAB HG002 Truth Set). We observed that 697 (29.58%) insertions and deletions were unique to a single SV caller, Interestingly, we observe that the majority of unique SVs in the short read SV call set (89.96%) overlap with GIAB-HG002 data set. This indicates that the majority of unique SVs are false negatives missed in the SV call sets of different SV callers. Additionally, 10.04% of the unique SV call set do not overlap with GIAB HG002 and are likely sequencing artifacts (false positives). Specifically, we observed that Manta (472) had the most unique SV calls, followed by Lumpy (312) and Delly (308), We observed that 67.86% (unique by Delly), 21.47% (unique by Lumpy), and 75.85% (unique by Manta) of these SV call sets respectively overlap with GIAB HG002 Truth Set. Thus, it is interesting to note that the majority of SV calls that are specific to Delly or Manta are in fact true positives. In parallel to this, it is evident that most false positives from SV caller variability are attributed to SV calls from Lumpy, followed by Delly and Manta. Further, we find that most non-unique SVs called are concordant with the GIAB SVs call set, On average, these are 84.31% concordant with GIAB-HG002 compared to unique caller SVs (55.06%).

#### Variability among Platforms

We investigated the variability attributed to different short read sequencing platforms (HiSeqX10, HiSeq2000 and HiSeq4000) while stratifying for variability from SV callers and replicates, SV call sets were merged across SV callers per sequencing run, requiring agreement of two or more SV callers for an SV to pass (Figure S15A). We then stratified for center and replicate variability by merging, sequentially requiring the SVs to be concordant in each merge respectively. SURVIVOR was used for each step, requiring SVs to be of the same type, have pairwise overlap smaller than 1000bp and be larger than 30 bp in size. A final SV call set was then created using a union merge across platforms followed by filtration using the GIAB high confidence regions.

We observe a total of 3023 SVs in the GIAB filtered SV call set that included 2411 deletions, 152 duplications, 117 inversions, 1 insertion and 342 translocations. For this SV call set, the HiSeqX10 has the largest number of SVs (2552), followed by HiSeq4000 (2260) and HiSeq2500 (2256) (Figure S15B). Across platforms, 81.49% on average of HiSeqX10 (83.39%), HiSeq4000 (81.33%), and HiSeq2500 (79.74%) SV call sets overlap with GIAB HG002 Truth Set (Figure S15C). Most non-unique SV calls (85.15%) overlap with the GIAB HG002 Truth Set. Thus, 14.85% of non-unique SVs were considered false positives in the SV call set where the remaining majority are true positives or false negatives missed by a platform’s SV call set. We observe that the HiSeq2500 produced the largest number of unique false positive SVs (143), followed by HiSeqX10 (127) and HiSeq4000 (84), possibly owing to the depth of sequencing achieved within each platform, Interestingly, 63.29% (219) of unique HiSeqX10 SVs are false negatives compared to HiSeq4000 33.33% (42) and HiSeq2500 27.41% (54).

We further filtered our SV call set to include only insertions and deletions in a similar manner to the GIAB SV call set (Figure S15D). The majority of SVs in the filtered SV set are deletions sized between 100-1000bp, which is comparable to other studies, including those from GIAB. We observe 374 insertions and deletions that are supported by a single platform, 311 (83.16%) of which overlap the GIAB HG002 call set, This is consistent with the SVs being false negatives missed by the two other platforms respectively, A total of 16.84% of SV calls do not overlap GIAB HG002 and are false positives due to individual platforms. We observed an increase in translocations and overall in variability likely due to the relaxed filtering strategy that had to be implemented for this comparison (Figure S15E).

#### Variability among Replicates

We investigated the variability among replicates for HG002 (HiSeqX10, HiSeq2000 and HiSeq4000 SV call sets) after stratifying for variability from SV callers, platforms and centers (Figure S16A). SV call sets were filtered across SV callers per sequencing run, requiring agreement of greater than two for an SV to pass. Subsequently, we merged these SVs across the three platforms requiring the SVs to be concordant, in order to stratify for platform variability. The SV call sets were then merged across centers requiring an overlap between all three, in order to stratify for center variability. A union merge using SURVIVOR was used across resulting replicates SV call sets with 50bp as the cutoff for SV size while maintaining previously described parameters, The SV call set was then filtered using the GIAB high confidence regions and overlapped with GIAB SV call set (Figure S16B). We identified 2797 SVs including 2267 deletions, 130 duplications, 103 inversions, 0 insertions, and 297 translocations (Figure S16C). We filtered the SV call set to only include insertions and deletions (2267 SVs) for comparisons with GIAB-HG002 truth set, A total of 268 SVs were supported by a single replicate of which 226 SVs (84.33%) overlap with the GIAB-HG002. The majority of the variability observed is thus due to false negatives in other replicate SV call sets compared to the individual one that has it. A total of 15.67% of unique replicate SVs are false positives that were not concordant with the GIAB HG002 Truth Set. Overall, 91.78% of non-unique SVs overlapped with GIAB-HG002 SV call set indicating a smaller number of false positives and high concordance between the replicates.

#### Clustering analysis of unique SVs

We investigated the possibility of an enrichment of these outlier SVs in certain genomic regions. Thus, for each strategy we took only the unique SVs that were supported only by one SV caller (blue), one replicate (gray), or one sequencing platform (yellow) (Figure S17). Here we counted the number of SVs starting in 100kbp windows across the genome in the GIAB high confidence regions. Overall we did not see significant clustering also supporting the observations that the majority of the unique calls are often true positives rather than false positives. Only very few SVs seem to cluster together, potentially explained as random noise, Only in one window did we observe eight SVs that are unique to a certain SV caller to cluster within 100kbp.

#### Variability due to Sequencing Factors

In the previous sections we investigated variability with respect to SV caller, platform, replicates and sequencing center, However, variant SV calling is often thought to be impacted by coverage, read length (e.g. impacting the mappability) and insert size of the paired end reads. Thus, we quantified for the first time the contribution of these three factors for SVs calling specifically across multiple replicates and with respect to true positive calls based on GIAB.

First, we investigated the impact of varying coverage on the ability to detect SVs (Figure S18). For this we computed the mean coverage along the genome as a representation and the standard deviation and compared both to the number of SVs and false positives based on GIAB SV calls (Figure S17 top row). It is clear to see that the increase in coverage has a positive effect with the number of SVs detected. This is also reflected in the number of true positives based on the GIAB SV calls, but not as clearly as the overall calls, While it is obvious why the true positive increases with the average coverage, given that more evidence can be found to support each SV, it is not clear how and if the standard deviation for coverage has a direct impact on the SV calling. This can be explained by the improvement in calling of SVs from paired-end reads with higher coverage.

Next, we investigated the possible impact of insertion size and variability of the insertion size for SV detection. We hypothesized that the variability of insert sizes plays an important role for the detection of SVs, since SV callers leverage the abnormal spacing of paired end reads to detect deletions[20, 25]. However, we could not identify a trend to support this, neither when investigating the average insert size vs, total number of SVs (**??** middle row), nor for the standard deviation of insert sizes. No trend was observed when focusing only on the true positive SVs based on the overlap with the GIAB SV call set). Thus, we conclude that the insert size variability seems not to impact directly the ability to identify SVs, at least not within the GIAB high confidence regions. This is possibly a result of the ability of the SV callers to leverage split read information.

Finally, we investigated the impact of read length on SV calling. We hypothesized that with an increase in read length, the mapping is more robust, and the ability of the mapper to characterize a split in the alignment should be improved. However, the trend that we observed with respect to the average read length and the standard deviation is minimal (Figure S17 bottom row). Since these are all Illumina data sets, we ignore for now the standard deviation of the read length. For the average read length we have 3 categories (100bp, 150bp and 250bp) that we can investigate. Among these bins, we don’t observe a clear pattern for the total number of SVs identified based on the average read length. Nevertheless, when filtered for the GIAB SV calls we do indeed see an improvement for true positive SV calls compared to the increase in read length.

## Supplementary figures

**Figure S1:**
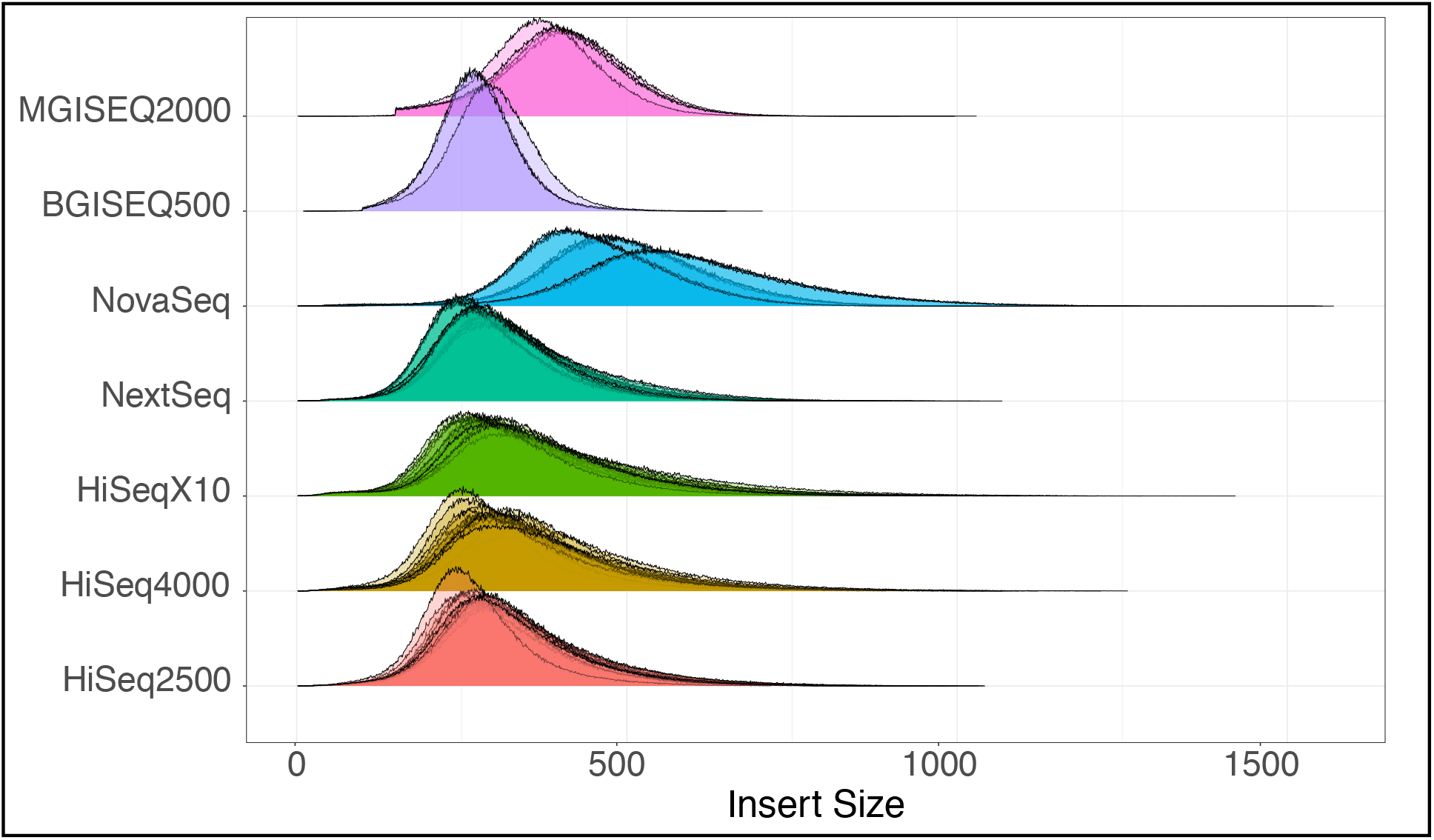
Insert size distribution per platform.

**Figure S2:**
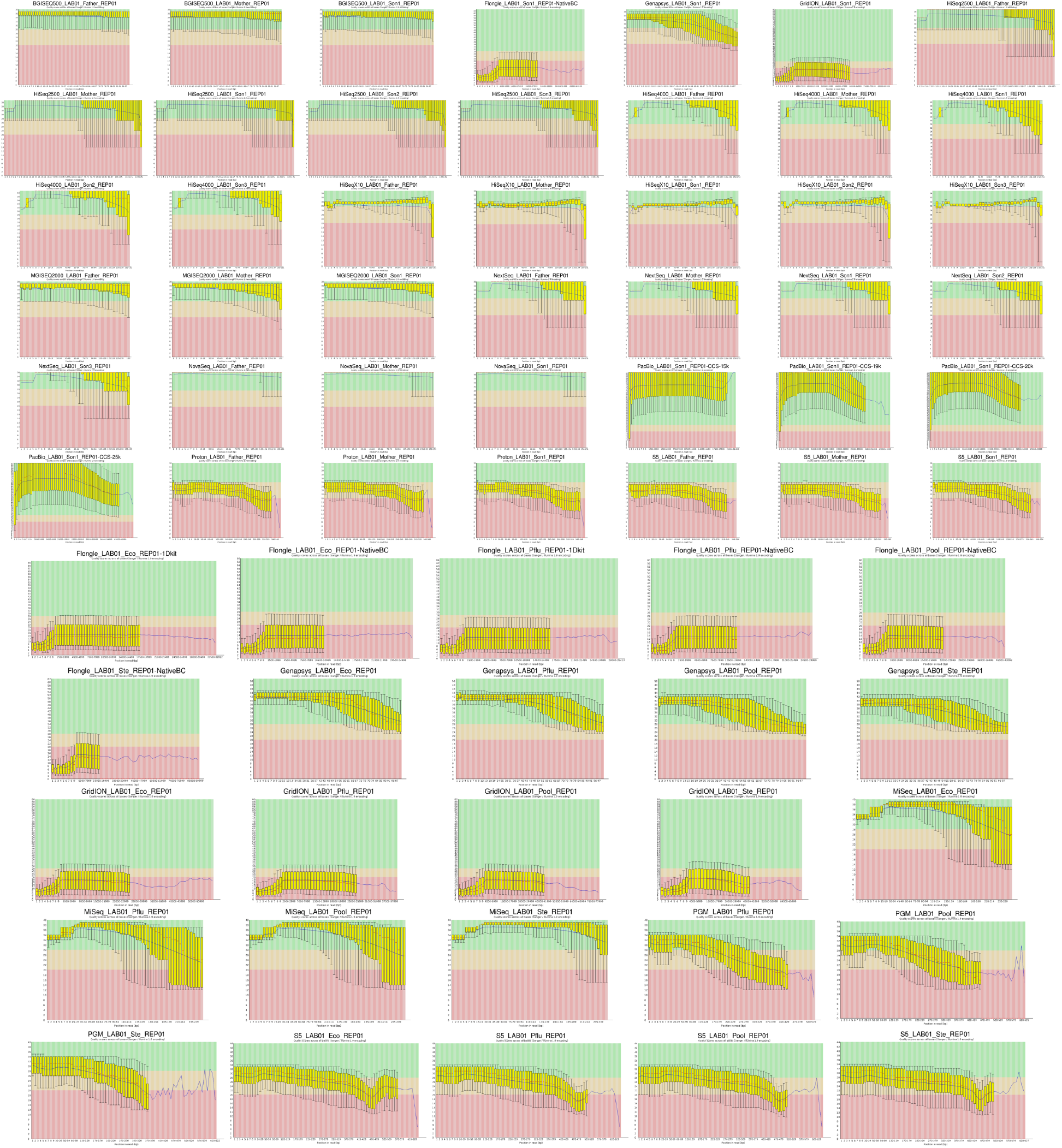
Base quality plots per replicate.

**Figure S3:**
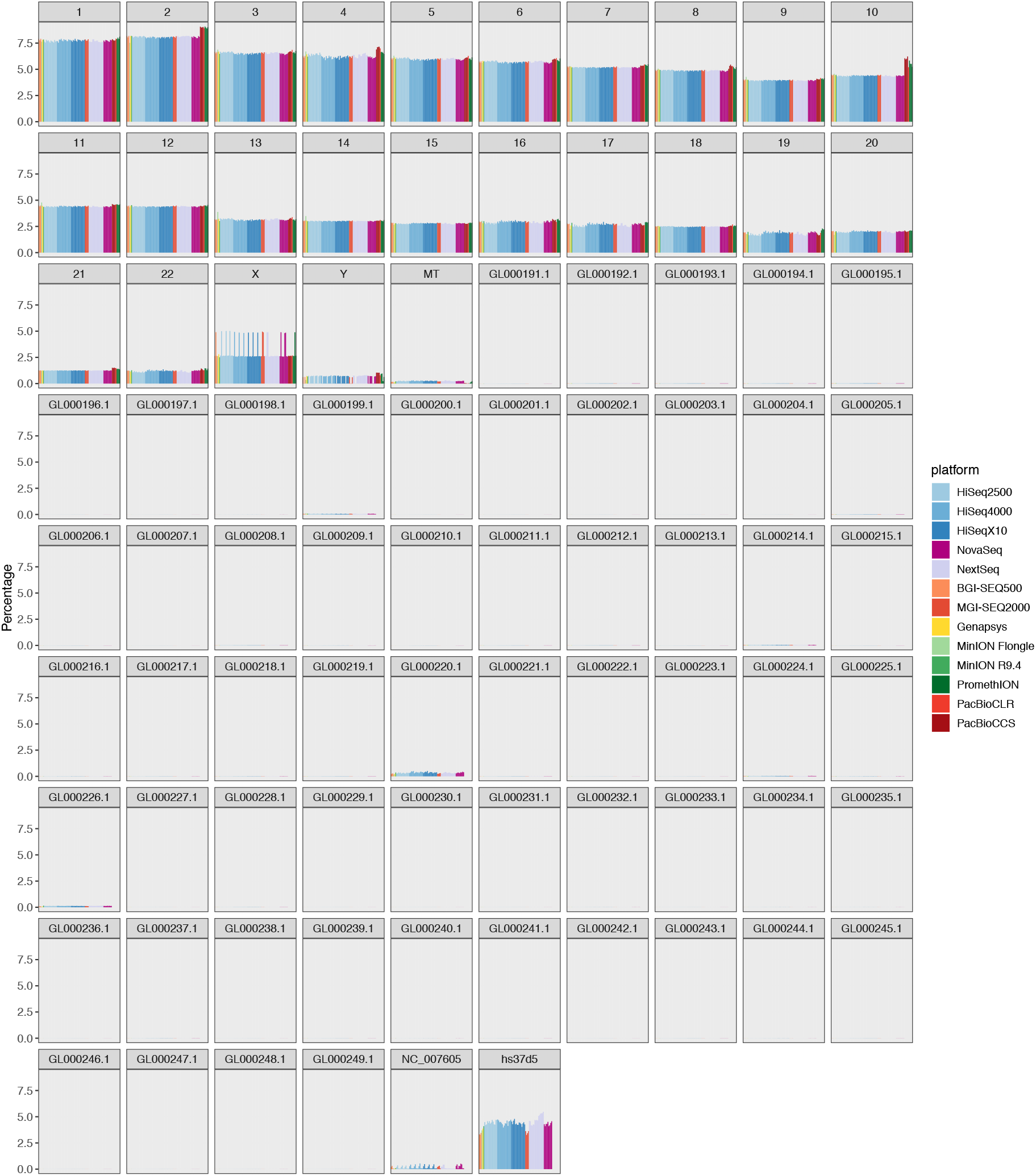
Percentage of coverage by chromosome for all human replicates against the hs37d5 reference genome, including canonical chromosomes and decoy sequences.

**Figure S4:**
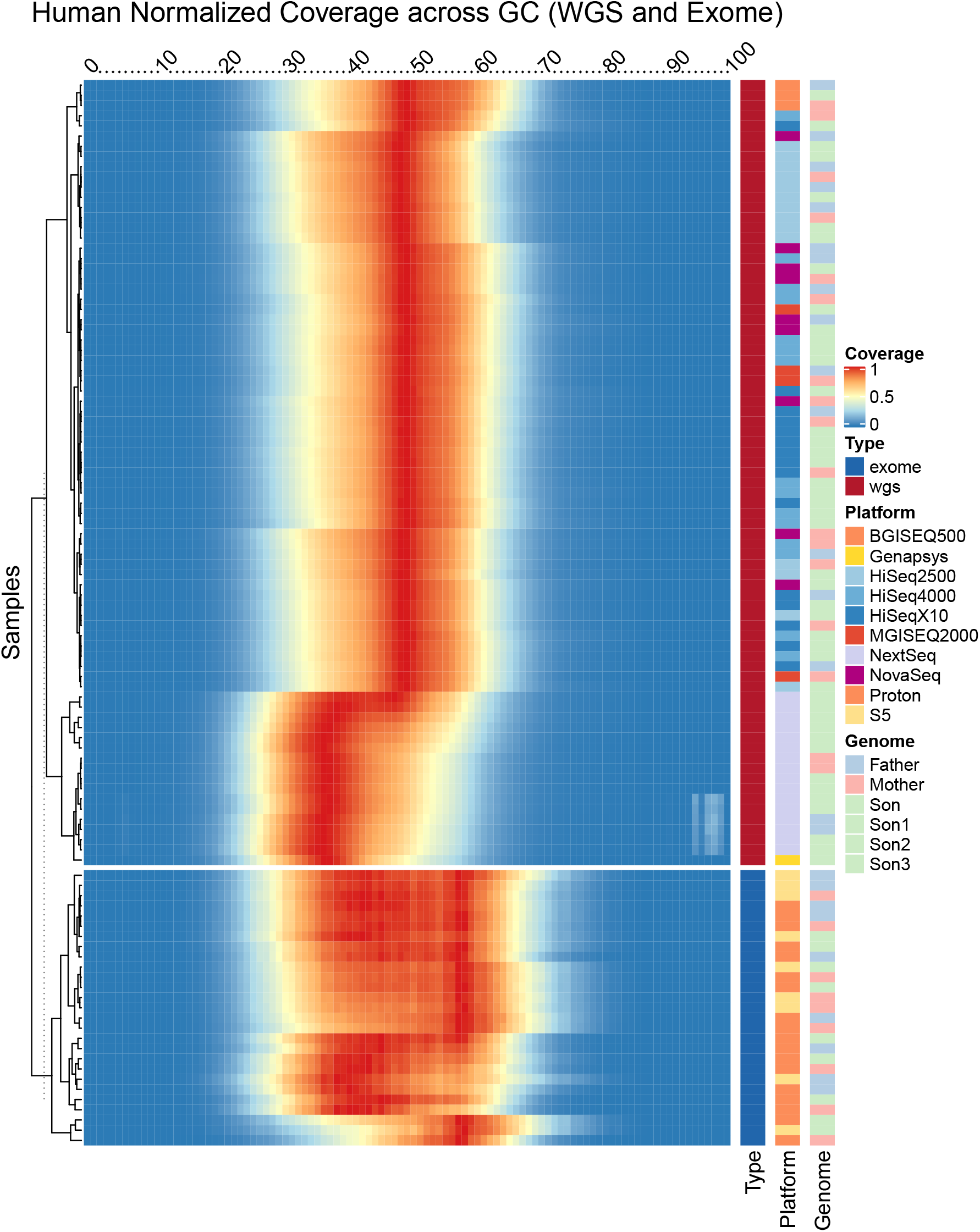
Distribution of reads by GC window across human whole genome and exome samples.

**Figure S5:**
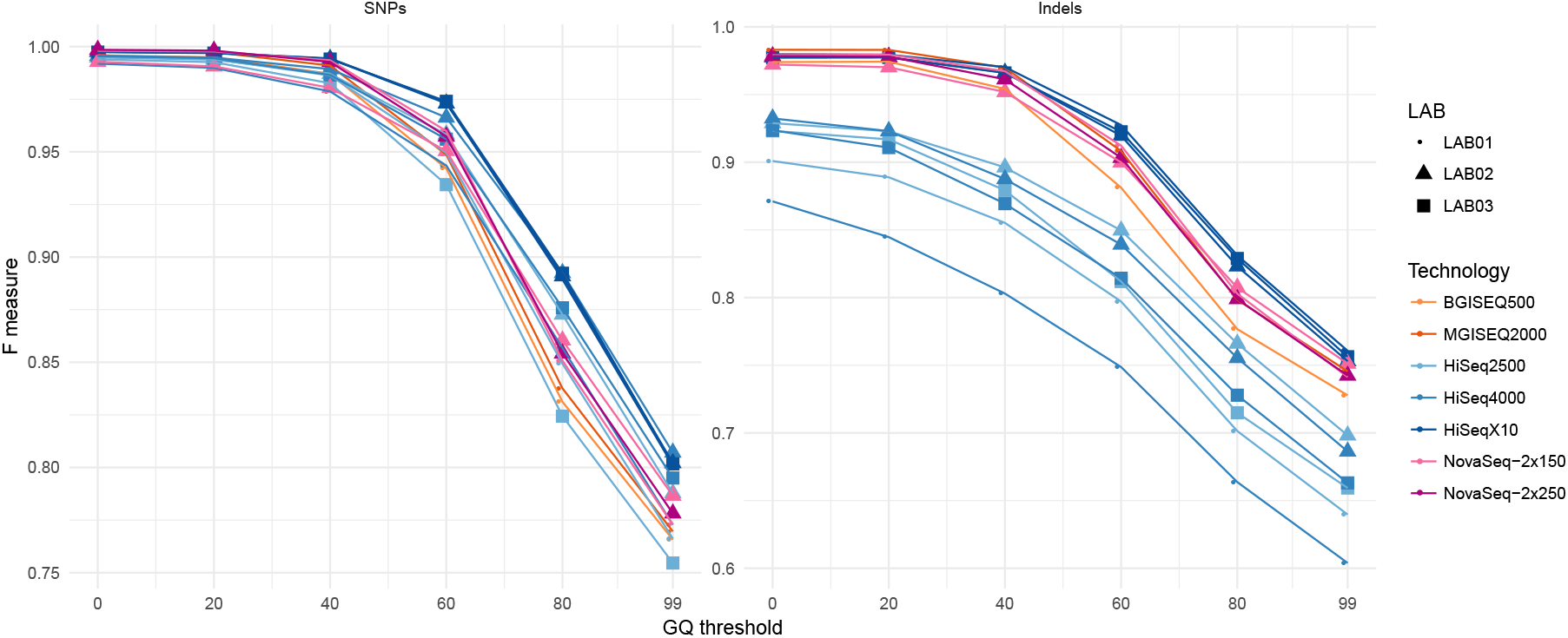
Distribution of F1 scores (harmonized metric combining sensitivity and specificity of concordance) at each Genotype Quality (GQ) cutoff per sequencing platform, separated by event type (SNPs and INDELs), only including replicates of the HG002 genome.

**Figure S6:**
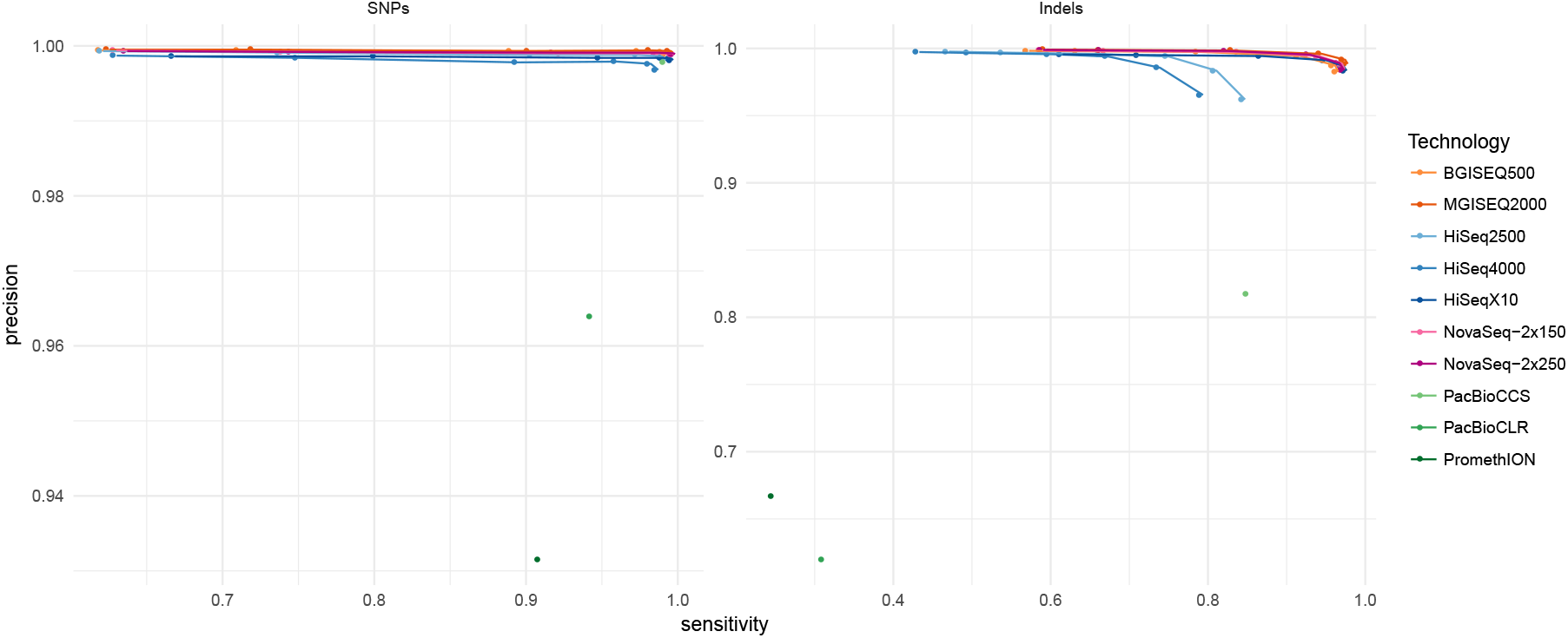
Sensitivity and precision metrics as derived from RTG’s vcfeval for all sequencing platforms, including long reads. Values at different Genotype Quality cutoff are plotted and connected, but not labeled. One replicate of the HG002 genome was chosen to represent each sequencing technology.

**Figure S7:**
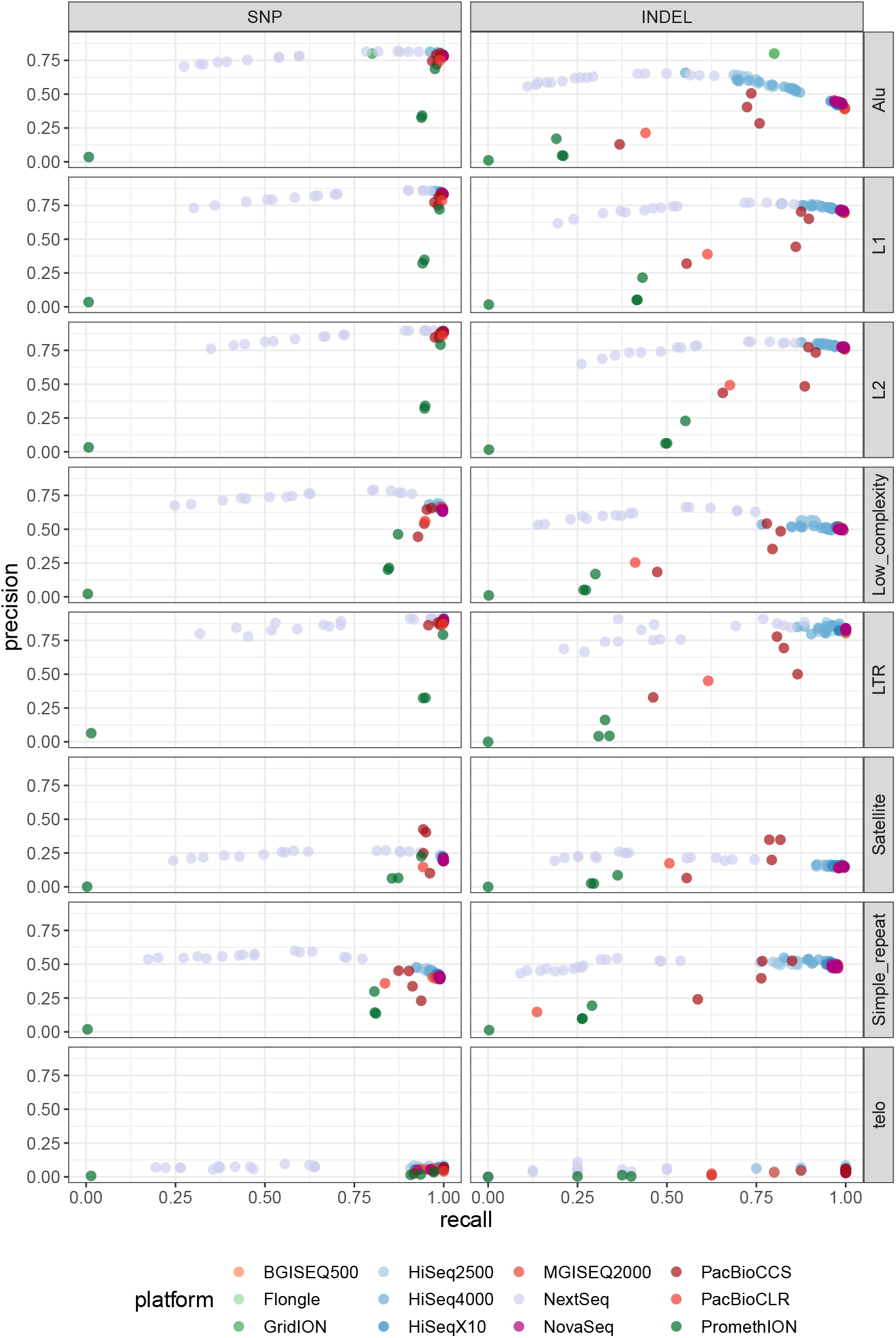
Precision and sensitivity scores as derived from RTG’s vcfeval, stratified by the UCSC RepeatMask contexts, and filtered by regions that cover the Genome in a Bottle (GIAB) high confidence regions for each respective genome within the Ashkenazi Trio.

**Figure S8:**
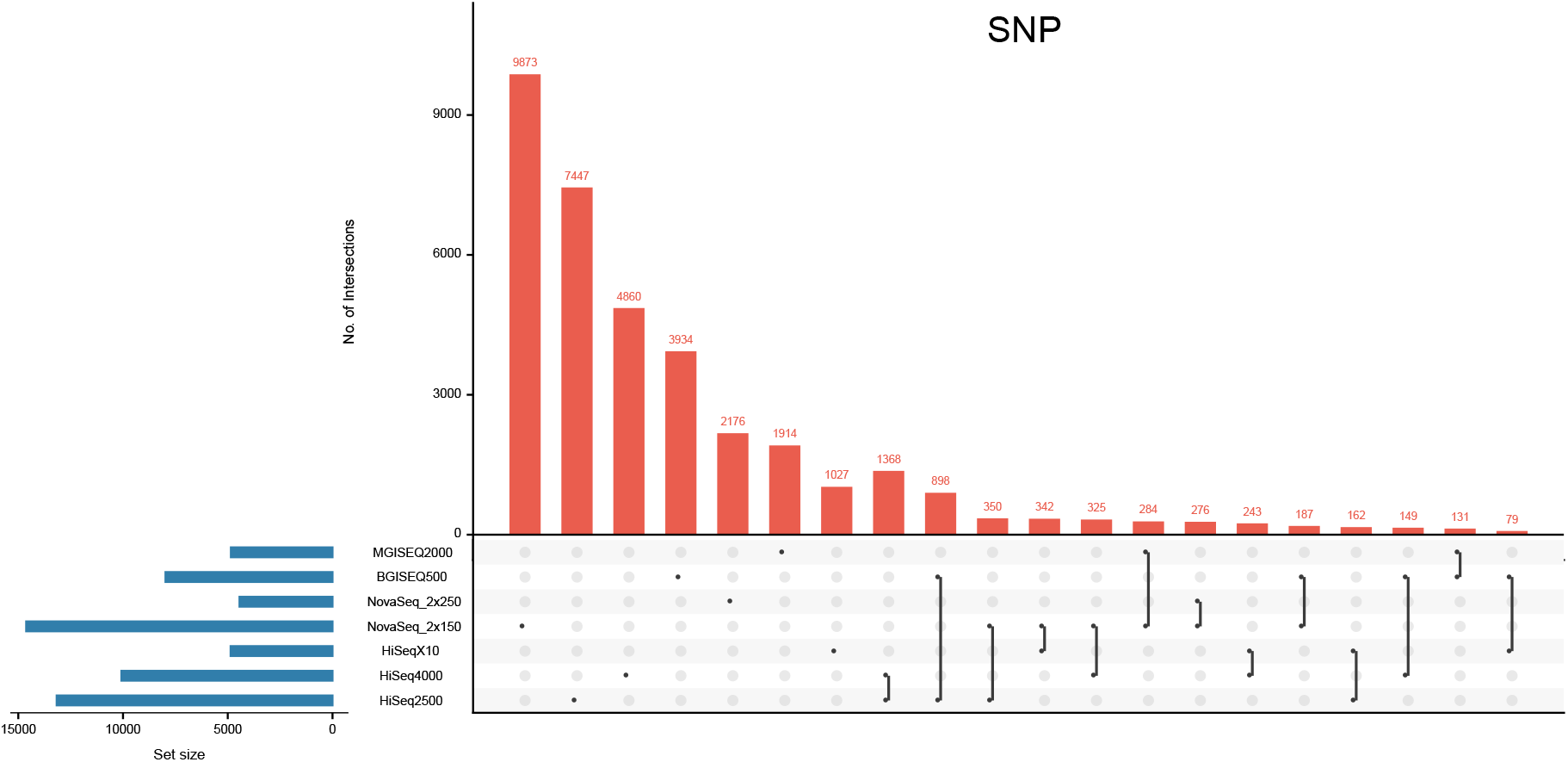
Intersection of SNP Mendelian violations summarized by platform.

**Figure S9:**
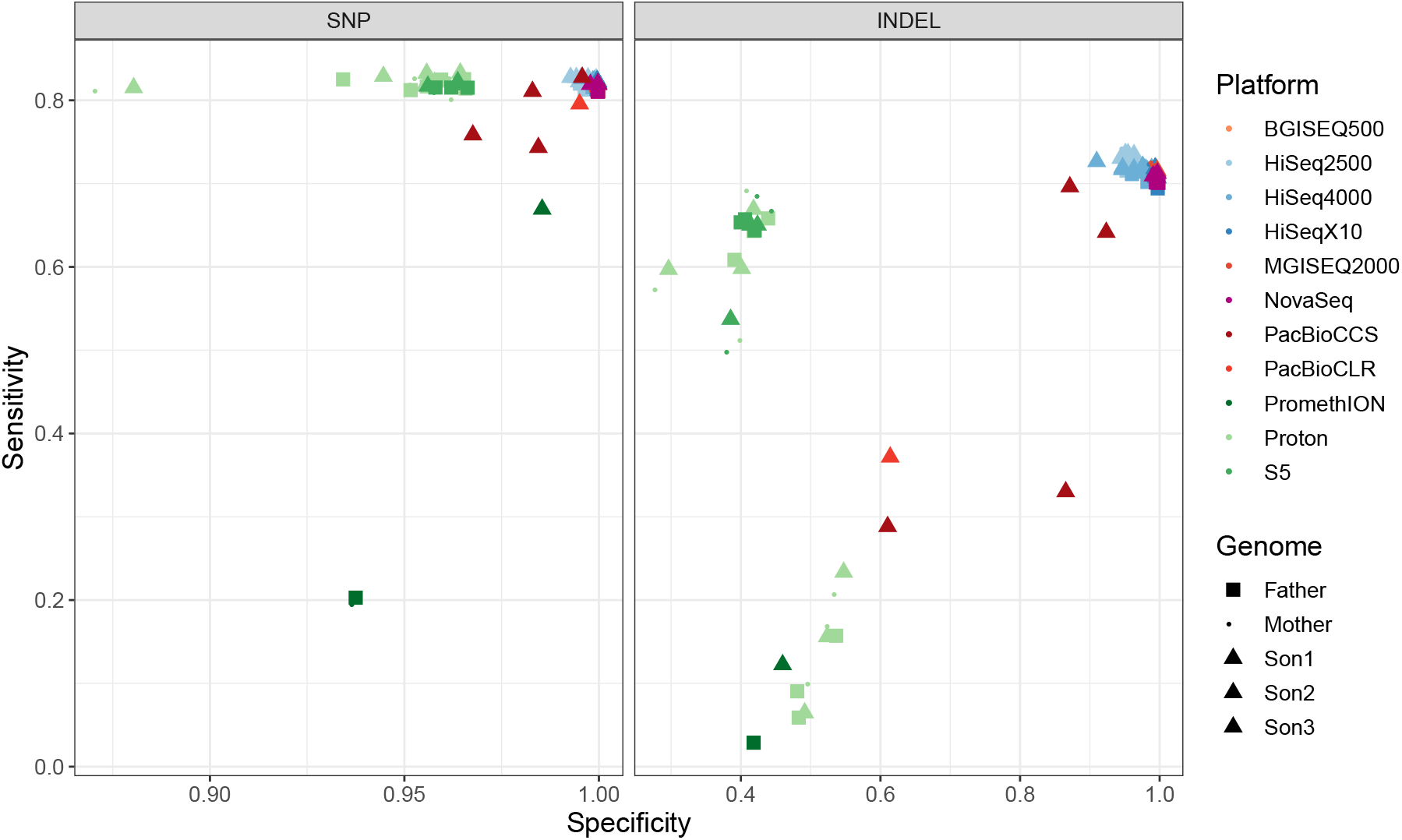
Sensitivity and precision metrics as derived from RTG’s vcfeval within the exome. Replicates included all Ion Torrent S5 and Proton data generated with the Ion Ampliseq protocol, and genomic samples filtered to exomic regions.

**Figure S10:**
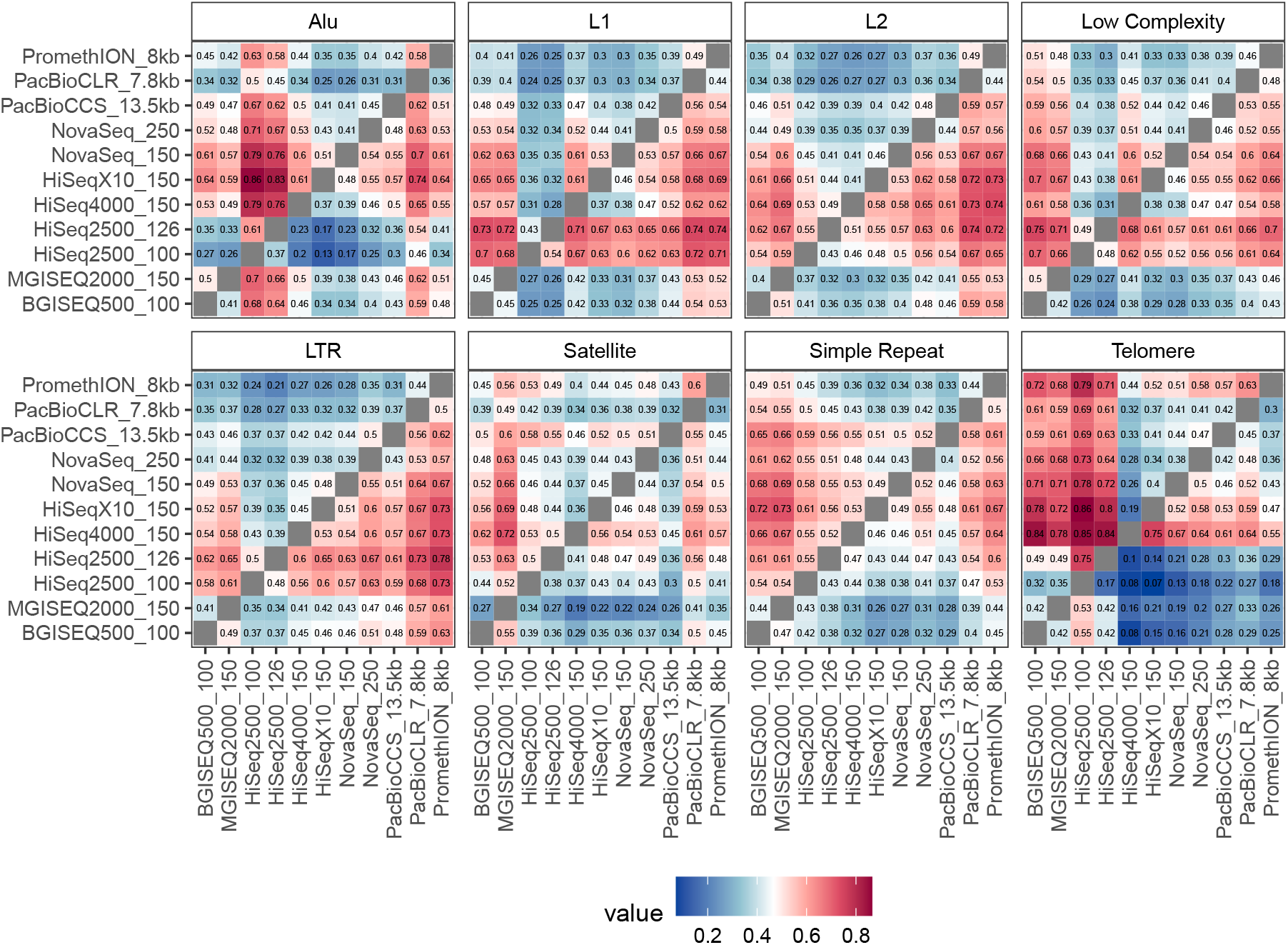
Pairwise comparison of coverage per genomic coordinate in each UCSC RepeatMask context, as captured by each sequencing technology. Comparisons of coverage were made for every pairwise combination in every context (Supplementary File X). The values in the heatmap represent the fraction of coordinates that are more highly covered in the Y-axis platform as compared to the X-axis platform, for any coordinate that is covered at 30x or below in both platforms (in order to avoid the very small percentage of points that have unexpected, very high coverage). Cells are non-reciprocal due to points which are equally covered by both platforms.

**Figure S11:**
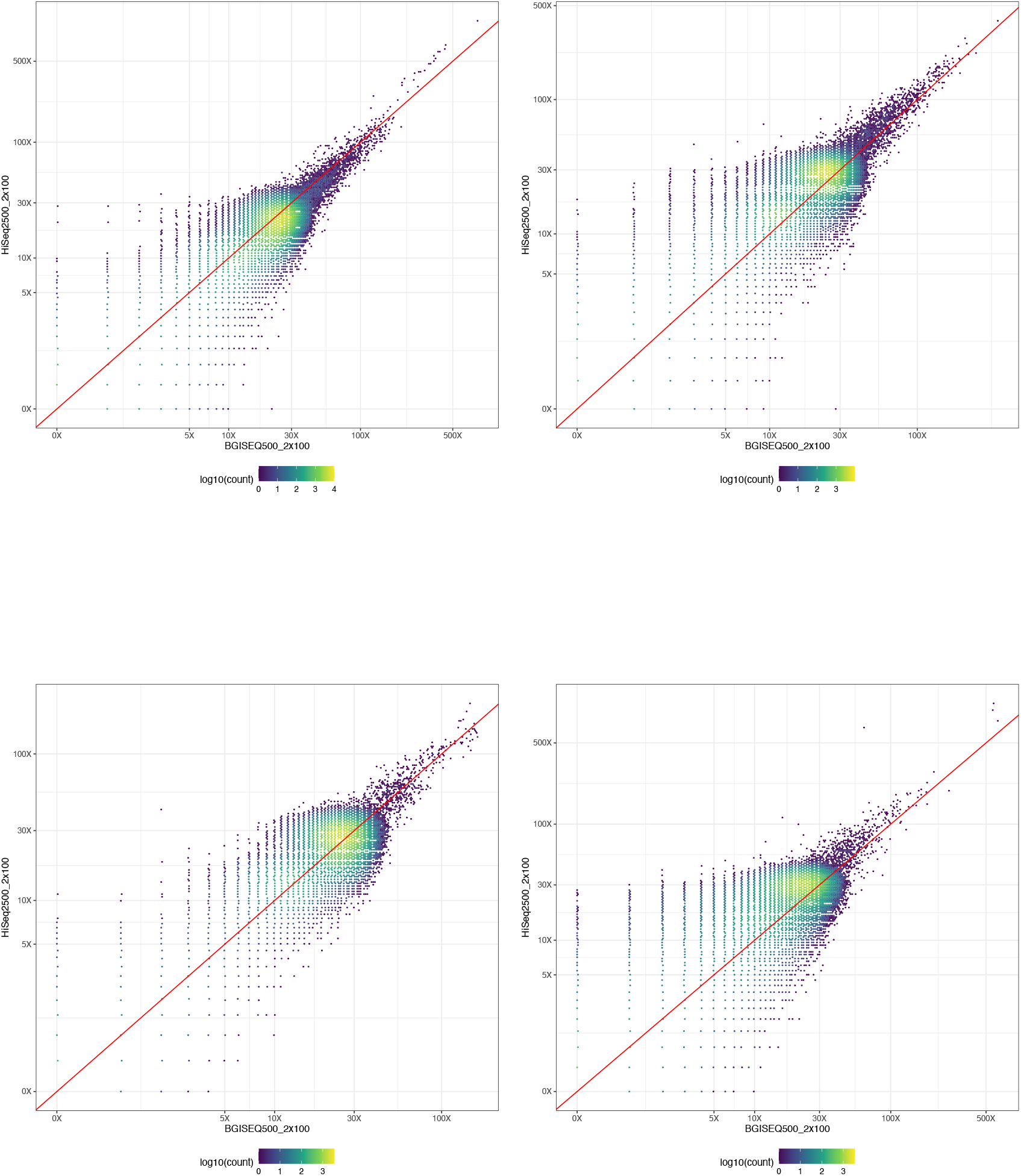
Examples of pairwise comparisons made between sequencing technologies of coverage in UCSC RepeatMask regions, using normalized 25x coverage alignments. Coverage of one platform versus another is shown, colored by frequency, and with the red a=b line indicating regions where coverage is equal. Repeat types clockwise from the upper left are Alu, L1, L2, and LTR.

**Figure S12:**
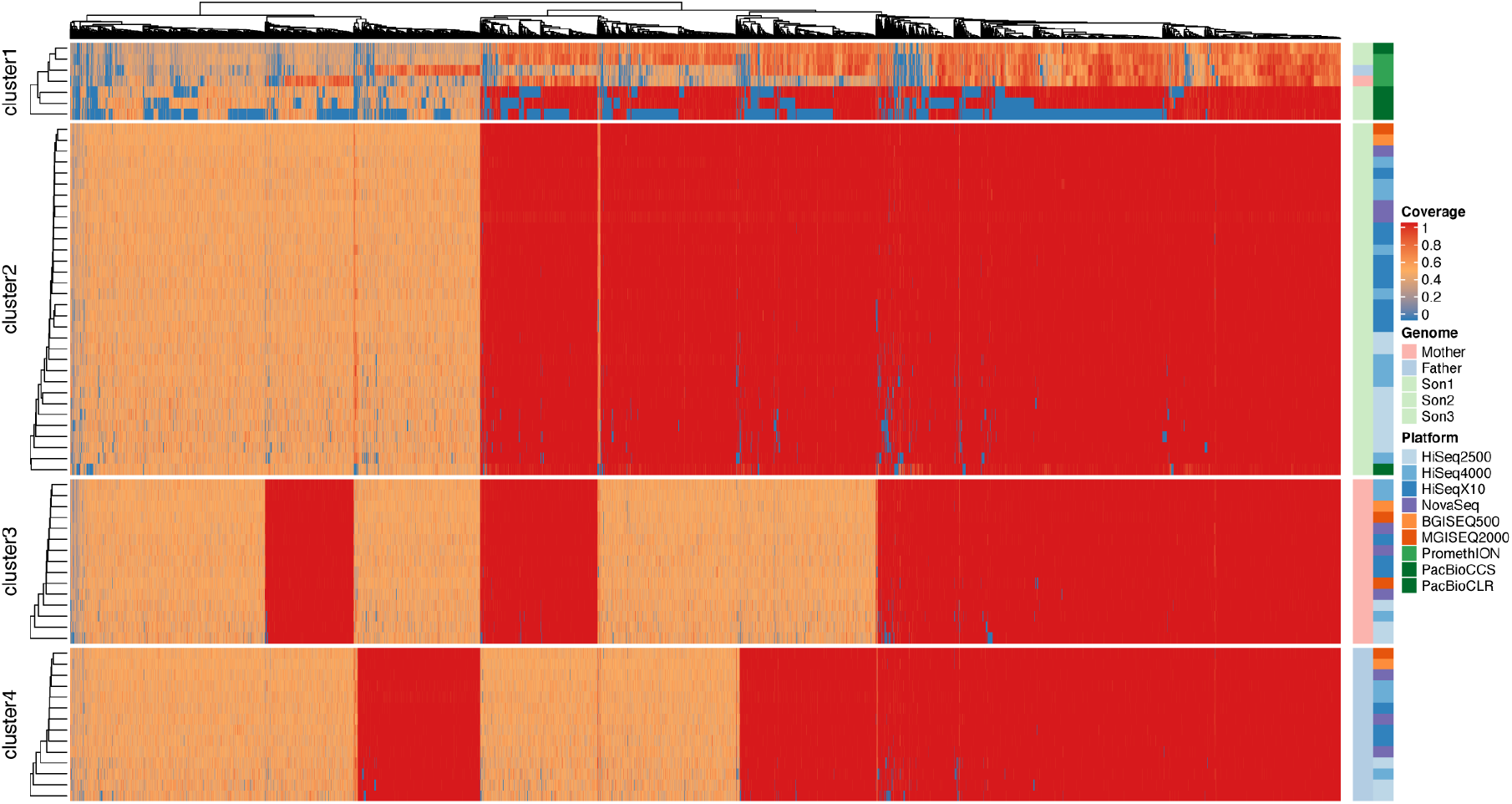
Heatmap of genotype of variant alleles across all human replicates across all sequencing platforms, as measured against the Genome in a Bottle high confidence variant call sets for each genome. Heterozygous variant alleles are shaded in orange, homozygous variants in red, and missing data in blue. Clusters 2, 3, and 4 represent the Ashkenazi Son (HG002), Mother (HG004), and Father (HG003) respectively, while cluster 1 is a combination of all three from long read platforms, which show much more variability in call type and a greater proportion of missing data.

**Figure S13:**
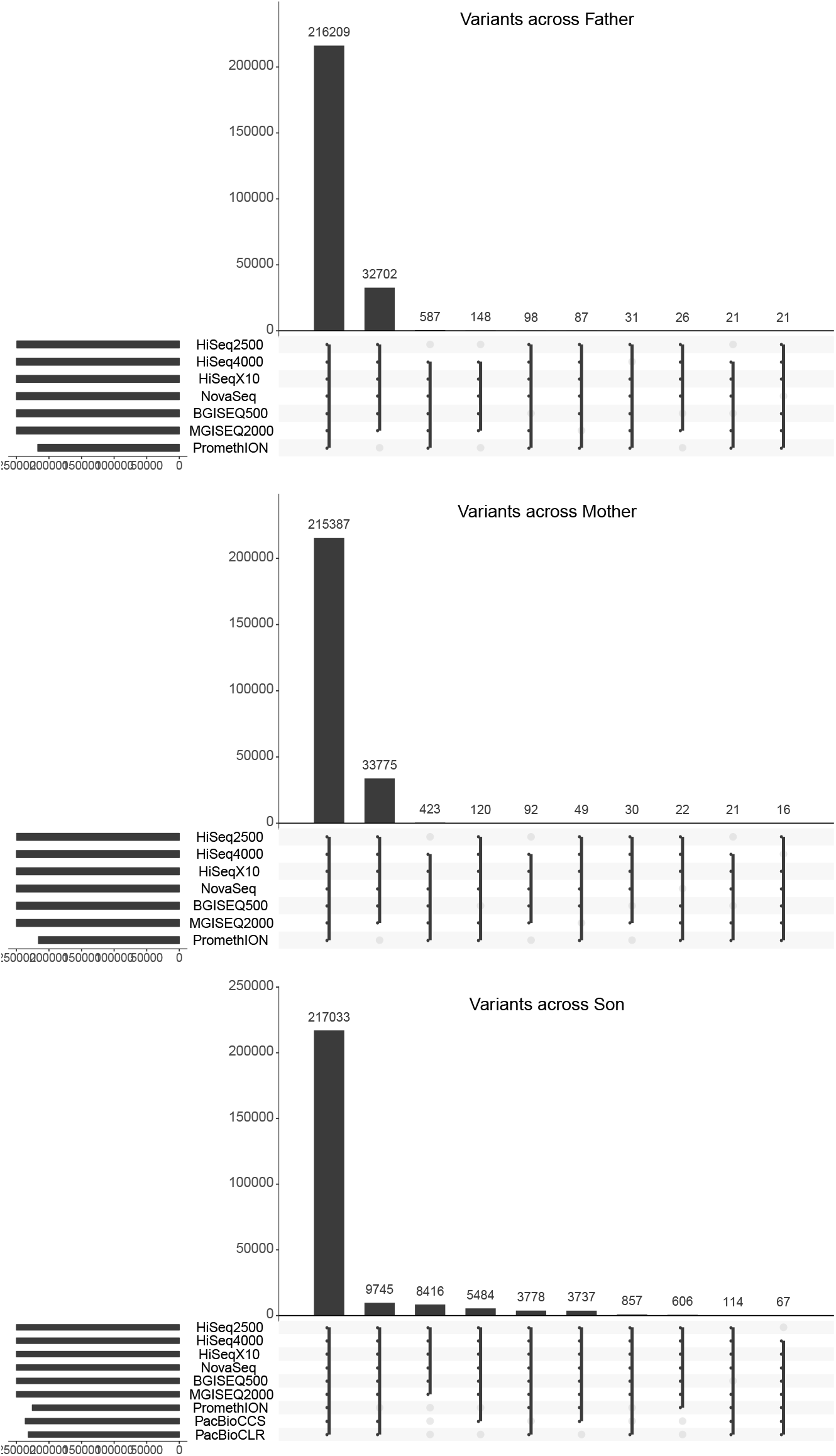
Distribution of overlap of variant allele calls per platform for each genome (taken from a random 10% subsample of the global distribution of calls). Overlaps were counted irrespective of variant type (homozygous or heterozygous). A variant allele was counted for a particular sequencing platform if the majority of replicates captured that variant, Only the top ten overlap sets are shown for each comparison.

**Figure S14:**
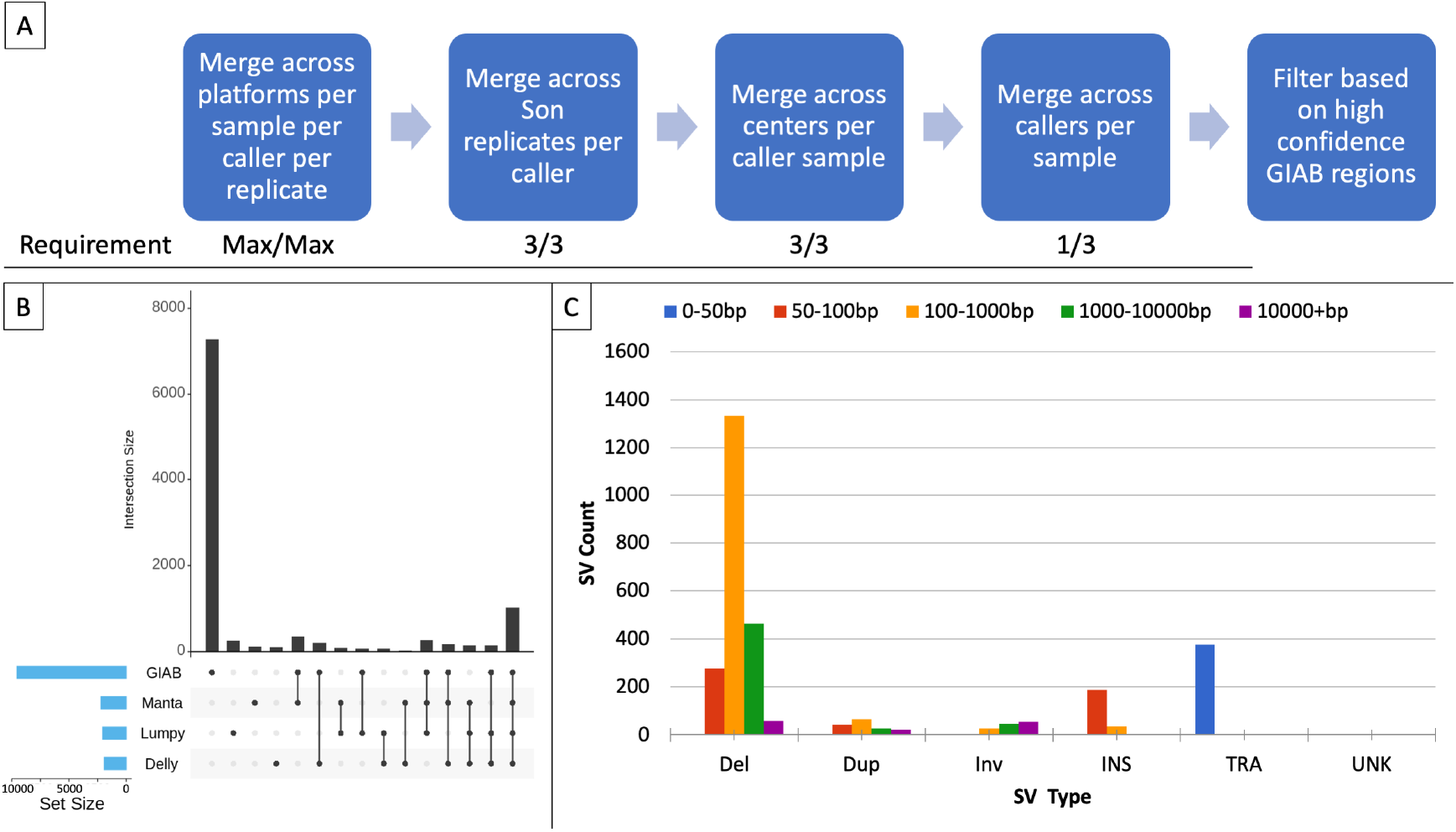
Structural Variant (SV) variability by caller using data from short read platforms with more than one HG002 replicate (HiSeq2500, HiSeq4000, HiSeqX10). (a) The strategy employed to examine SV caller variability, after stratifying for platforms, replicates, and centers. (b) the SV call set sizes and overlap with the Genome in a Bottle (GIAB) SV call set for HG002. (c) types and sizes of SVs across callers, with translocations set to 50bp by default by SURVIVOR for the purposes of visualization.

**Figure S15:**
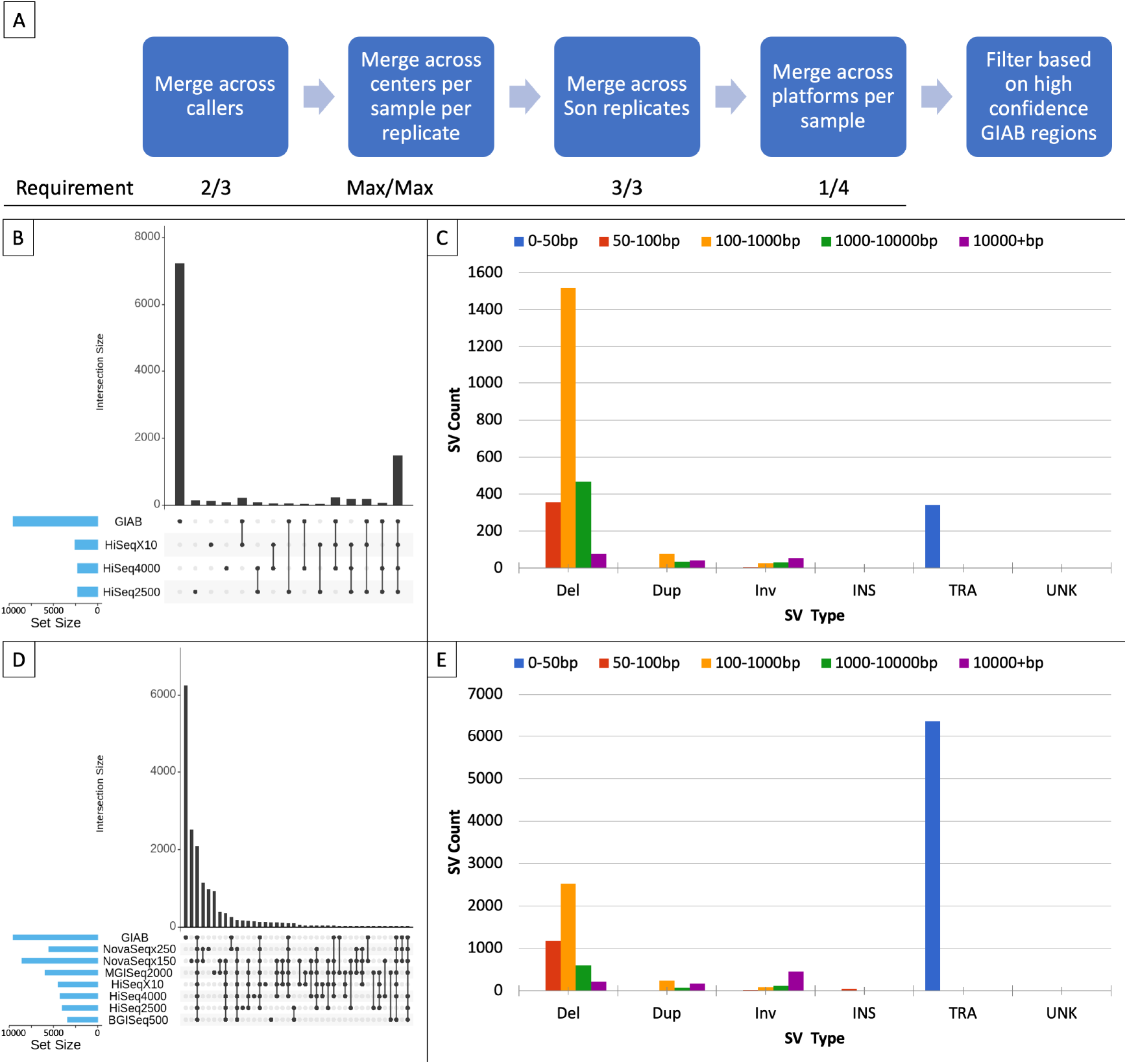
Structural Variant (SV) variability by platform. (a) The strategy employed to examine SV caller variability, after stratifying for callers, replicates, and centers. (b) the SV call set sizes and overlap with the Genome in a Bottle (GIAB) SV call set for HG002. (c) types and sizes of SVs across callers, with translocations set to 50bp by default by SURVIVOR for the purposes of visualization. (d) same as (b). but including more platforms (BGISEQ500, MGISEQ2000, and both NovaSeq types, 2×150bp and 2×250bp). (e) same as (c). but including the platforms represented in (d).

**Figure S16:**
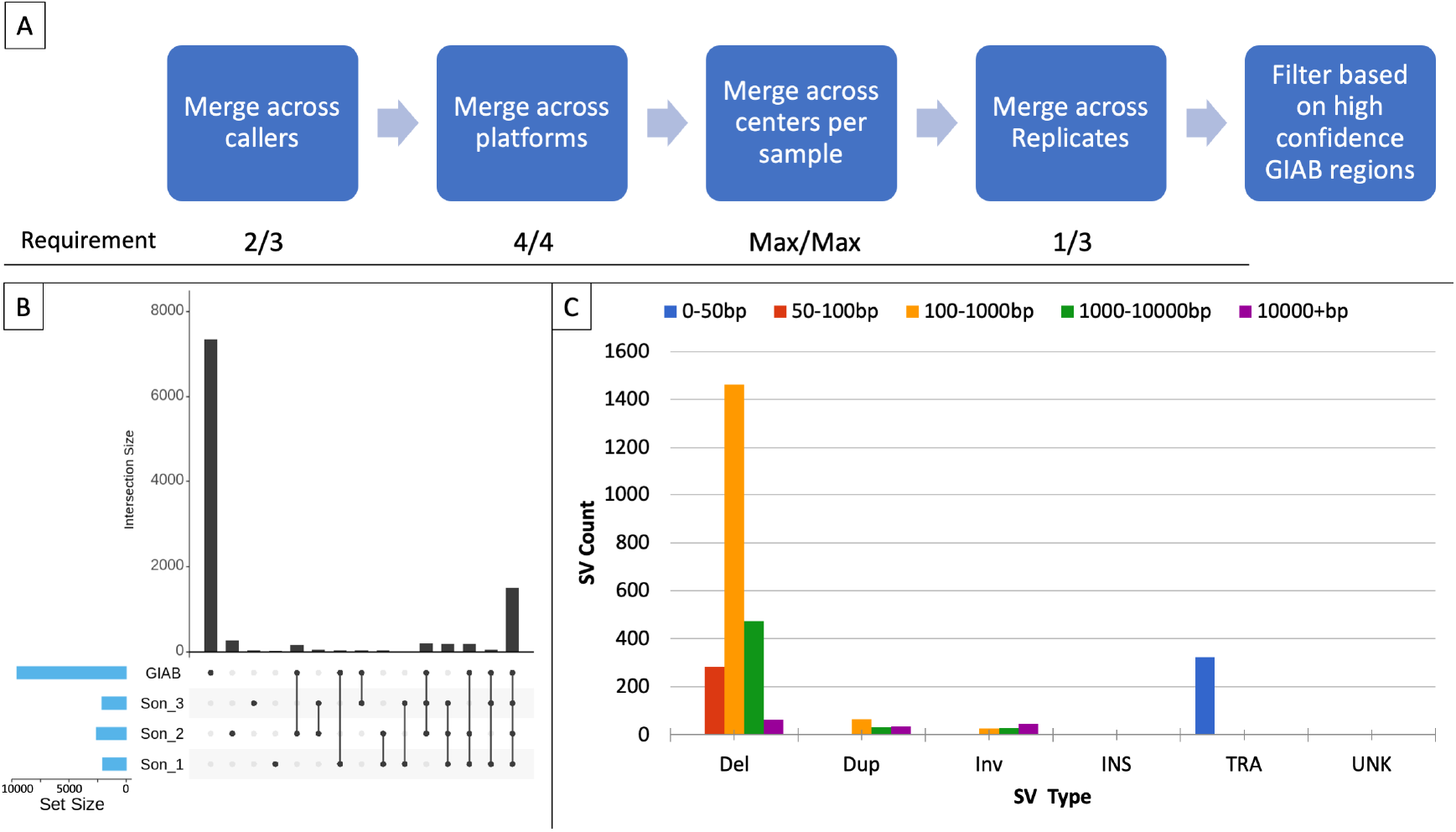
Structural Variant (SV) variability by replicate of the HG002 genome using data from short read platforms with more than one HG002 replicate (HiSeq2500, HiSeq4000, HiSeqX10). (a) The strategy employed to examine SV caller variability, after stratifying for platforms, replicates, and centers. (b) the SV call set sizes and overlap with the Genome in a Bottle (GIAB) SV call set for HG002. (c) types and sizes of SVs across callers, with translocations set to 50bp by default by SURVIVOR for the purposes of visualization.

**Figure S17:**
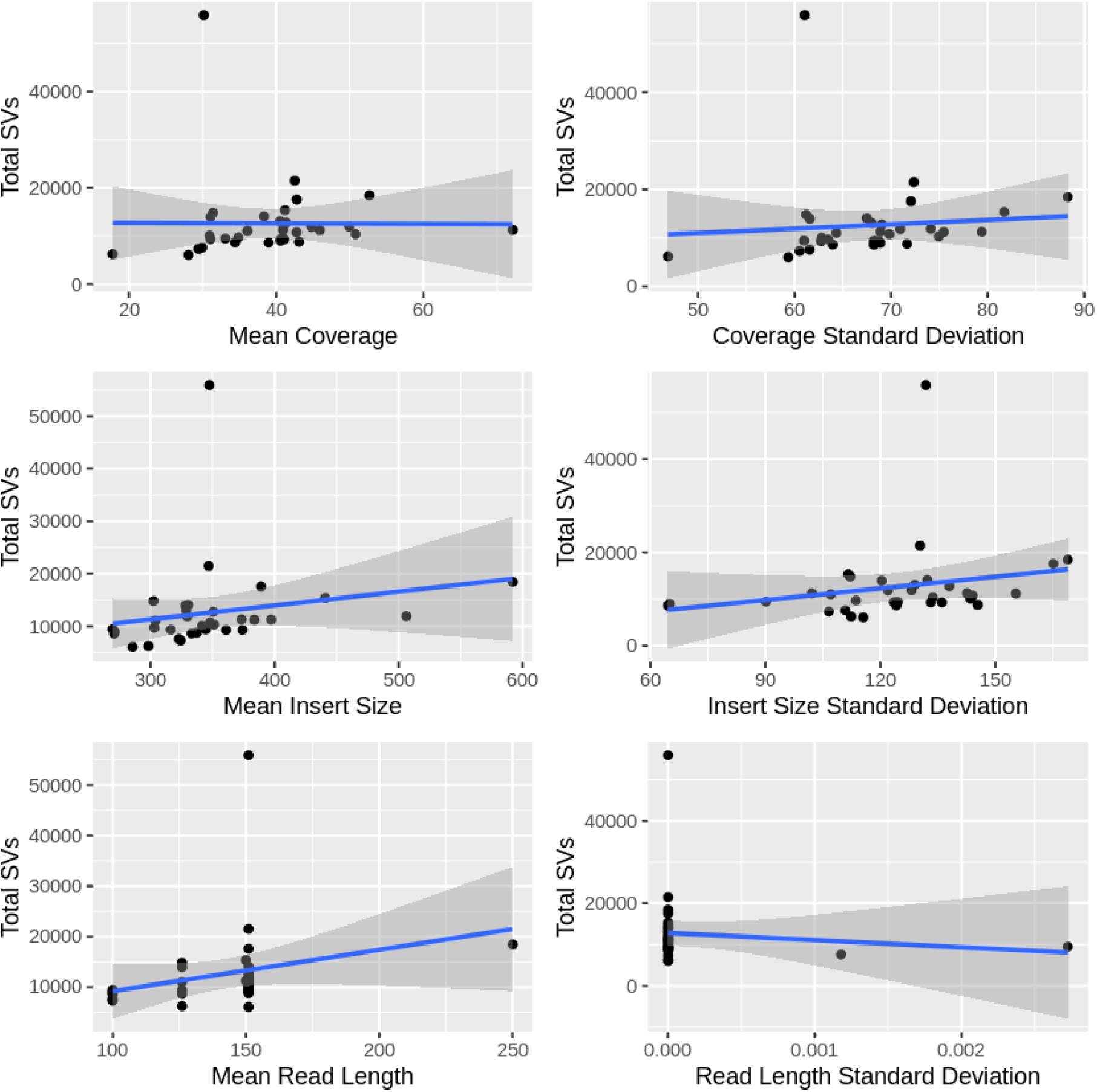
Coverage, insert size, and read length mean and standard deviation across total structural variants (SVs) overlapping with the Genome in a Bottle HG002 truth set, with no filtration.

**Figure S18:**
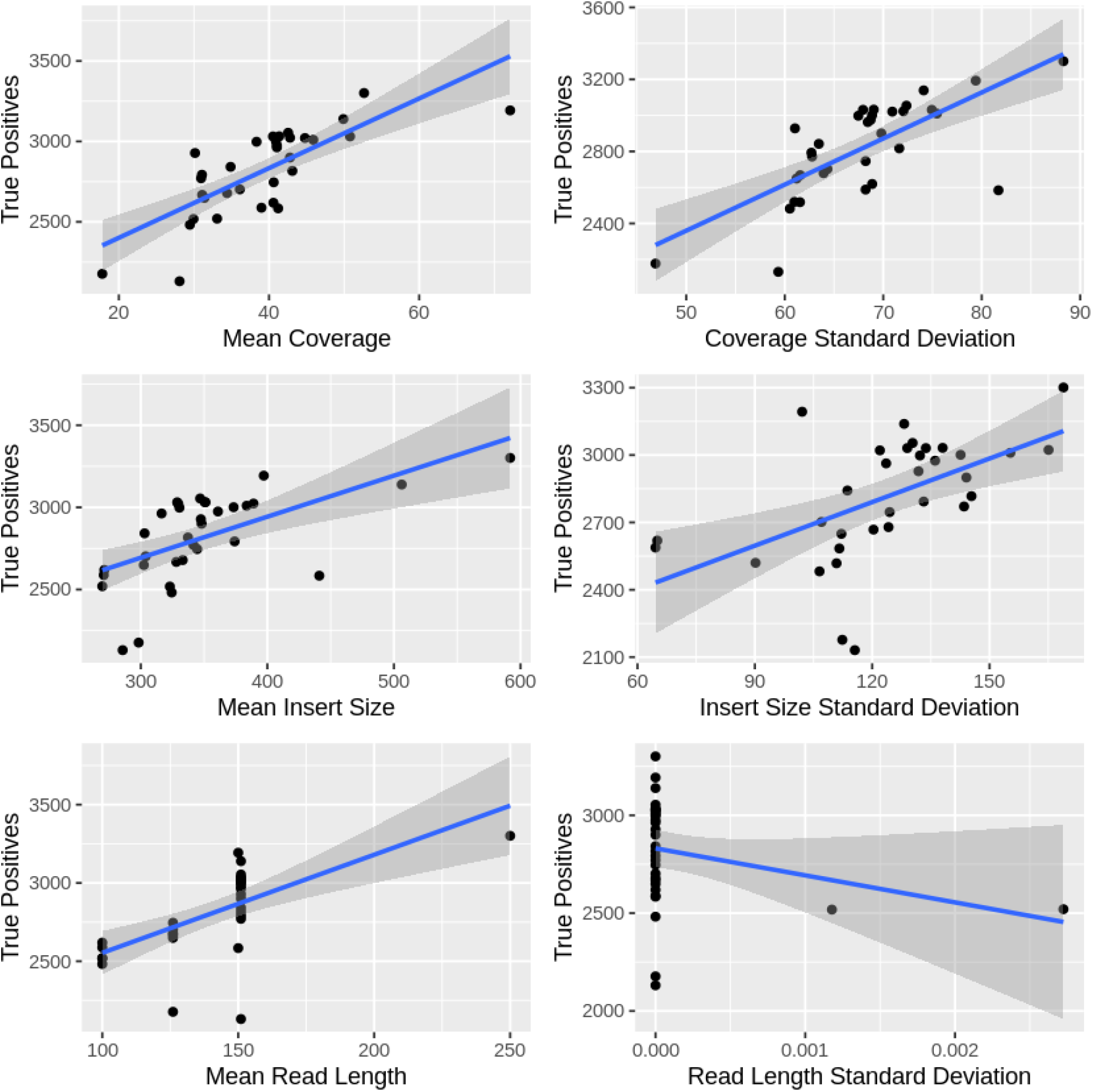
Coverage, insert size, and read length mean and standard deviation across total structural variants (SVs) overlapping with the Genome in a Bottle HG002 truth set, filtered to only true positives.

**Figure S19:**
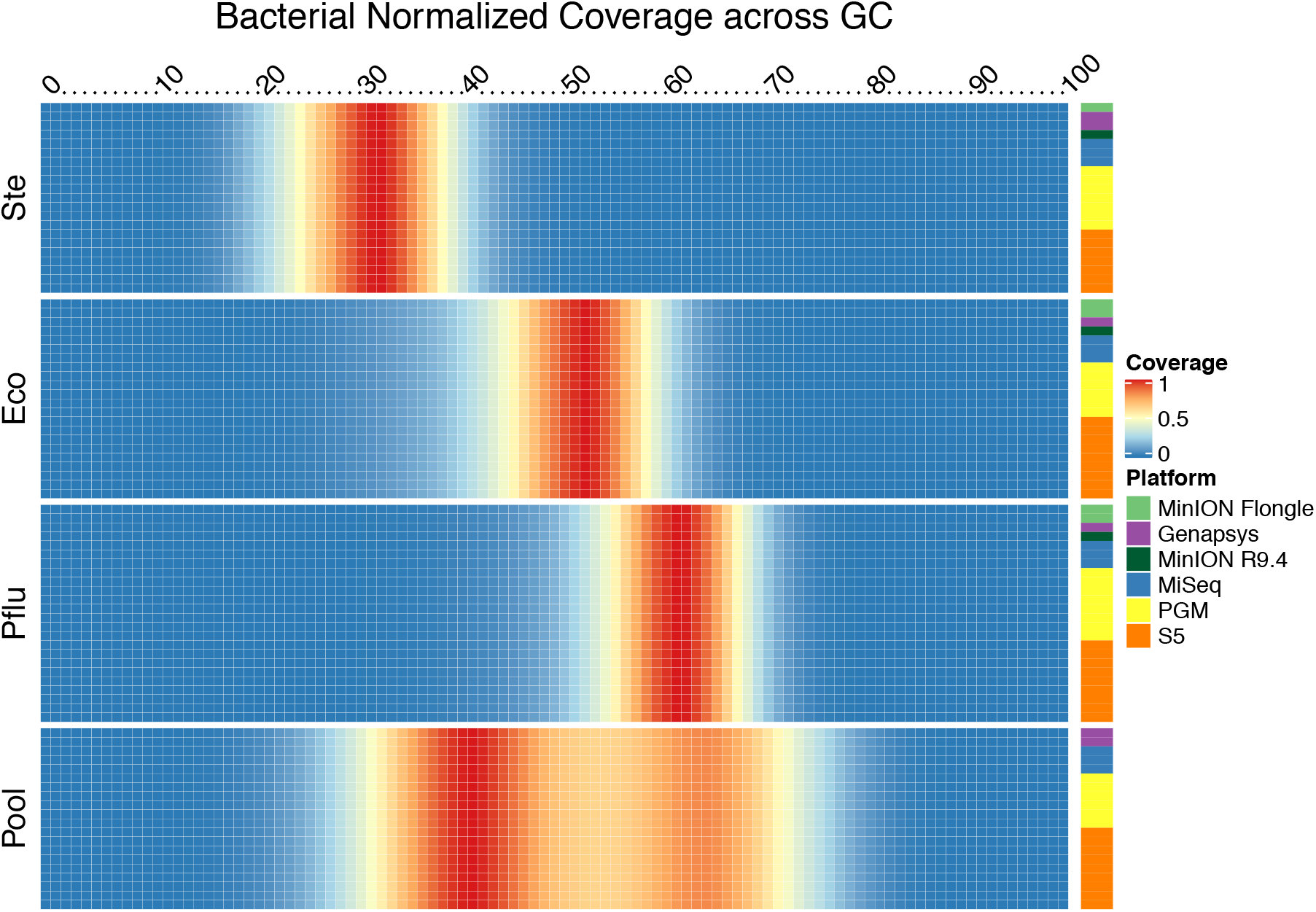
Distribution of coverage across GC content windows per bacterial genome using the number of reads that match a particular GC and normalized by total reads per replicate such that a value of 1 matches the bin with the greatest numer of reads. All replicates are shown, and are colored by sequencing platform. Ste=*S. aureus*; Eco=*E. coli*; Pflu=*P fìuorescens*; Pool=Metagenomic mixture often bacterial species.

## References

1. Schuster, S. C. Next-generation sequencing transforms today’s biology. Nature methods 5, 16–18 (2008).

2. Shendure, J. & Ji, H. Next-generation DNA sequencing. Nature biotechnology 26, 1135 (2008).

3. DePristo, M. A. et al. A framework for variation discovery and genotyping using next-generation DNA sequencing data. Nature genetics 43, 491 (2011).

4. Mardis, E. R. The impact of next-generation sequencing technology on genetics. Trends in genetics 24, 133–141 (2008).

5. MacLean, D., Jones, J. D. & Studholme, D. J. Application of’next-generation’sequencing technologies to microbial genetics. Nature Reviews Microbiology 7, 96–97 (2009).

6. Glenn, T. C. Field guide to next-generation DNA sequencers. Molecular ecology resources 11, 759–769 (2011).

7. Zhou, J. et al. Reproducibility and quantitation of amplicon sequencing-based detection. The ISME journal 5, 1303–1313 (2011).

8. Mellmann, A. et al. High interlaboratory reproducibility and accuracy of next-generation-sequencing-based bacterial genotyping in a ring trial. Journal of clinical microbiology 55, 908–913 (2017).

9. Quail, M. A. et al. A tale of three next generation sequencing platforms: comparison of Ion Torrent, Pacific Biosciences and Illumina MiSeq sequencers. BMC genomics 13, 341 (2012).

10. Shi, L. et al. The MicroArray Quality Control (MAQC) project shows inter-and intraplatform reproducibility of gene expression measurements. Nature biotechnology 24, 1151 (2006).

11. Shi, L. et al. The MicroArray Quality Control (MAQC)-II study of common practices for the development and validation of microarray-based predictive models. Nature biotechnology 28, 827 (2010).

12. Li, S. et al. Multi-platform assessment of transcriptome profiling using RNA-seq in the ABRF next-generation sequencing study. Nature Biotechnology 32, 915 (2014).

13. Su, Z. et al. A comprehensive assessment of RNA-seq accuracy, reproducibility and information content by the Sequencing Quality Control Consortium. Nature biotechnology 32, 903 (2014).

14. Wang, C. et al. The concordance between RNA-seq and microarray data depends on chemical treatment and transcript abundance. Nature biotechnology 32, 926 (2014).

15. Li, S. et al. Detecting and correcting systematic variation in large-scale RNA sequencing data. Nature biotechnology 32, 888 (2014).

16. Risso, D., Ngai, J., Speed, T. P. & Dudoit, S. Normalization of RNA-seq data using factor analysis of control genes or samples. Nature biotechnology 32, 896 (2014).

17. Zook, J. M. et al. An open resource for accurately benchmarking small variant and reference calls. Nature biotechnology 37, 561–566 (2019).

18. Krusche, P. et al. Best practices for benchmarking germline small-variant calls in human genomes. Nature biotechnology 37, 555–560 (2019).

19. Zook, J. M. et al. Extensive sequencing of seven human genomes to characterize benchmark reference materials. Scientific data 3, 1–26 (2016).

20. Zook, J. M. et al. A robust benchmark for detection of germline large deletions and insertions. Nature biotechnology (2020).

21. Ball, M. P. et al. A public resource facilitating clinical use of genomes. Proceedings of the National Academy of Sciences 109, 11920–11927 (2012).

22. Benson, G. Tandem repeats finder: a program to analyze DNA sequences. Nucleic acids research 27, 573–580 (1999).

23. Toptaş, B. Ç., Rakocevic, G., Kómár, P. & Kural, D. Comparing complex variants in family trios. Bioinformatics 34, 4241–4247 (2018).

24. Jeffares, D. C. et al. Transient structural variations have strong effects on quantitative traits and reproductive isolation in fission yeast. Nature communications 8, 1–11 (2017).

25. Mahmoud, M. et al. Structural variant calling: the long and the short of it. Genome biology 20, 246 (2019).

26. Sedlazeck, F. J. et al. Accurate detection of complex structural variations using single-molecule sequencing. Nature methods 15, 461–468 (2018).

27. McIntyre, A. B. et al. Comprehensive benchmarking and ensemble approaches for metagenomic classifiers. Genome biology 18, 182 (2017).

28. Freed, D. N., Aldana, R., Weber, J. A. & Edwards, J. S. The Sentieon Genomics Tools-A fast and accurate solution to variant calling from next-generation sequence data. BioRxiv, 115717 (2017).

29. Li, H. Minimap2: pairwise alignment for nucleotide sequences. Bioinformatics 34, 3094–3100 (2018).

30. Luo, R. et al. Exploring the limit of using a deep neural network on pileup data for germline variant calling. Nature Machine Intelligence 2, 220–227 (2020).

31. Rausch, T. et al. DELLY: structural variant discovery by integrated paired-end and split-read analysis. Bioinformatics 28, i333–i339 (2012).

32. Layer, R. M., Chiang, C., Quinlan, A. R. & Hall, I. M. LUMPY: a probabilistic framework for structural variant discovery. Genome biology 15, R84 (2014).

33. Chen, X. et al. Manta: rapid detection of structural variants and indels for germline and cancer sequencing applications. Bioinformatics 32, 1220–1222 (2016).

34. Pedersen, B. S. & Quinlan, A. R. Mosdepth: quick coverage calculation for genomes and exomes. Bioinformatics 34, 867–868 (2018).

35. Conway, J. R., Lex, A. & Gehlenborg, N. UpSetR: an R package for the visualization of intersecting sets and their properties. Bioinformatics 33, 2938–2940 (2017).

36. Cleary, J. G. et al. Comparing variant call files for performance benchmarking of next-generation sequencing variant calling pipelines. BioRxiv, 023754 (2015).

37. Li, H. et al. The sequence alignment/map format and SAMtools. Bioinformatics 25, 2078–2079 (2009).

