## Supplementary figures and images for "Multi-Platform Assessment of DNA Sequencing Performance using Human and Bacterial Reference Genomes in the ABRF Next-Generation Sequencing Study"

### GIAB_L1_NovaSeq_2x150-v-NovaSeq_2x250_hex.pdf

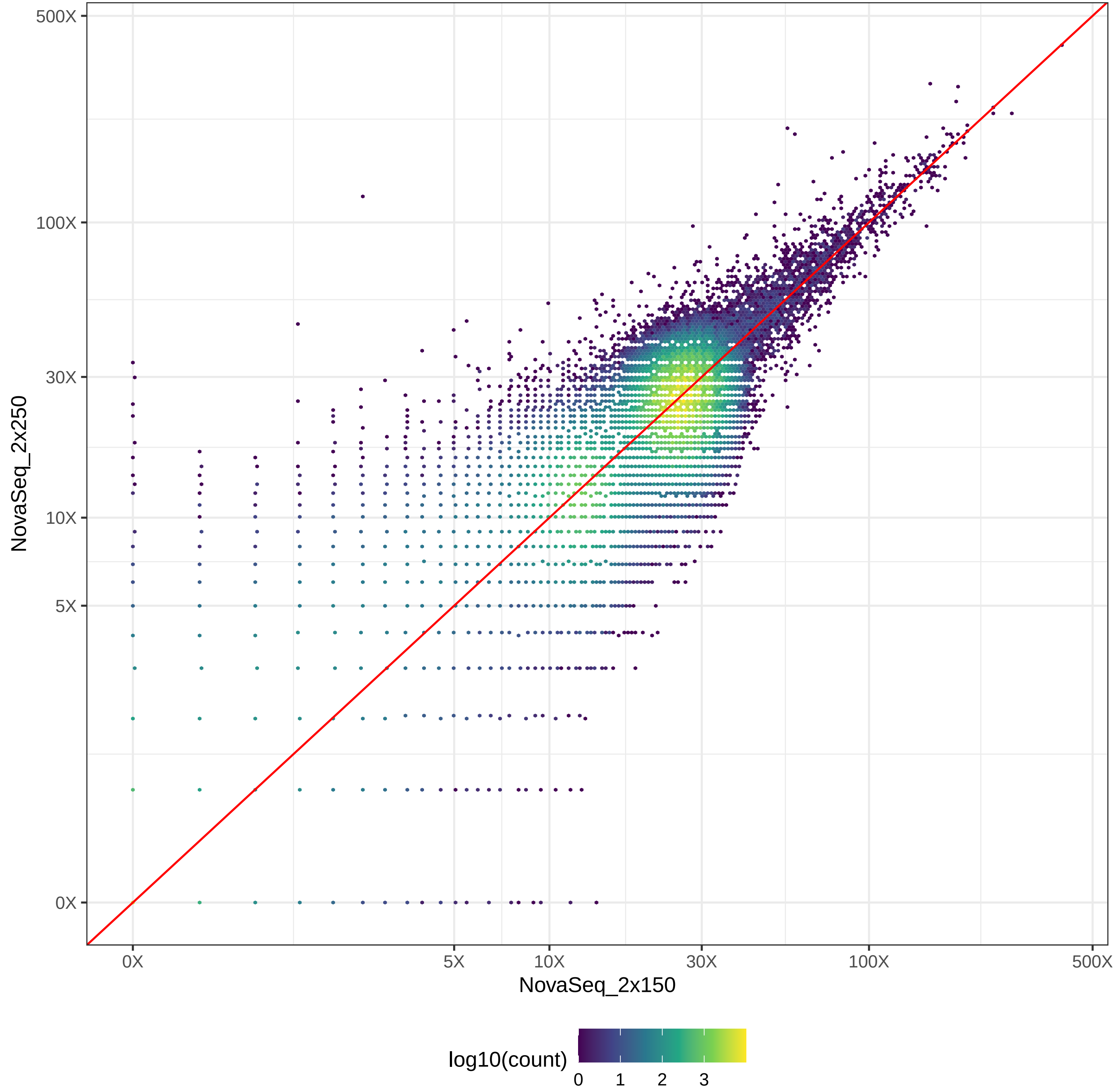

### GIAB_L1_NovaSeq_2x150-v-PacBio_CCS-15k_hex.pdf

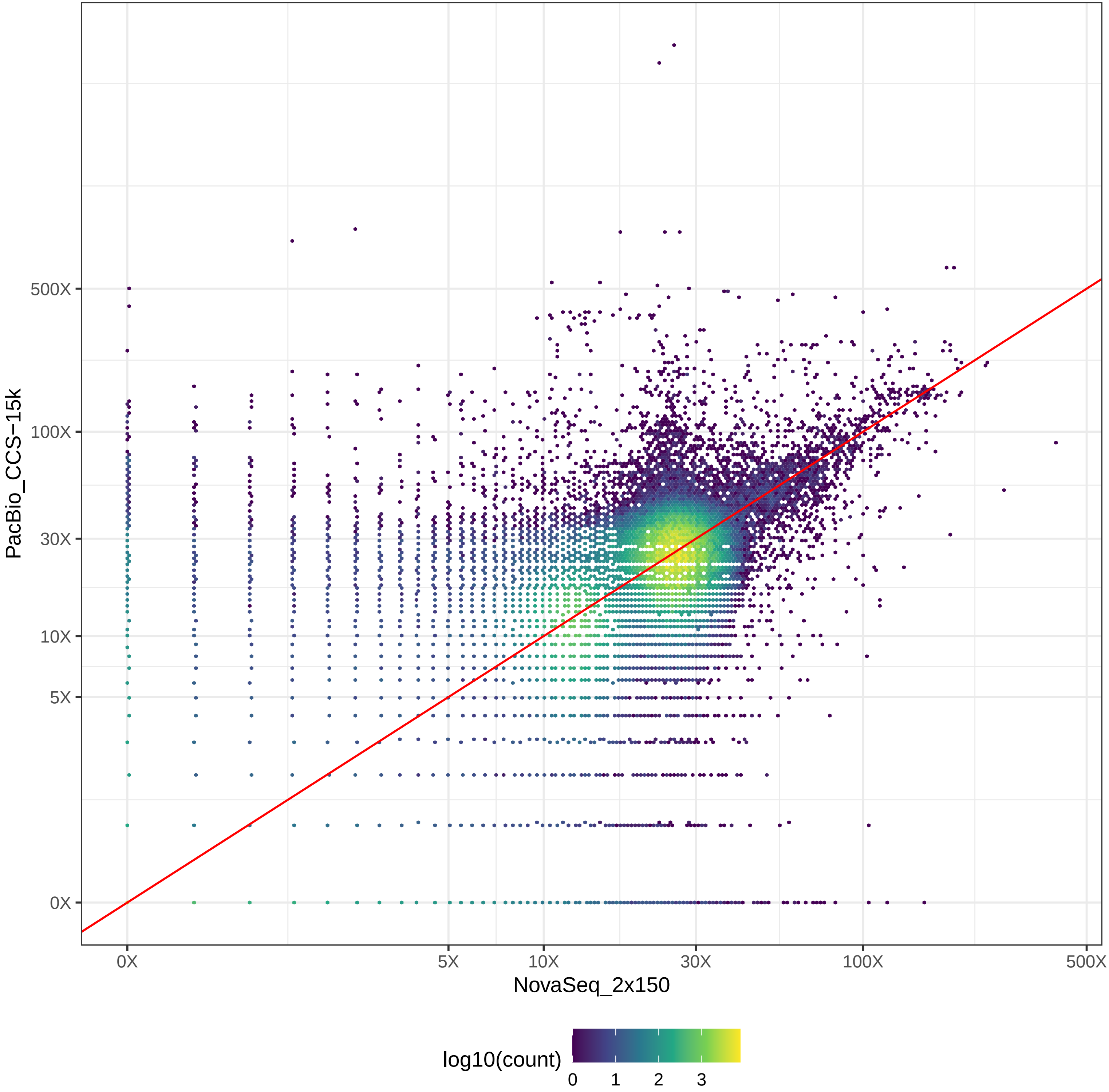

### GIAB_L1_NovaSeq_2x150-v-PacBio_CLR_hex.pdf

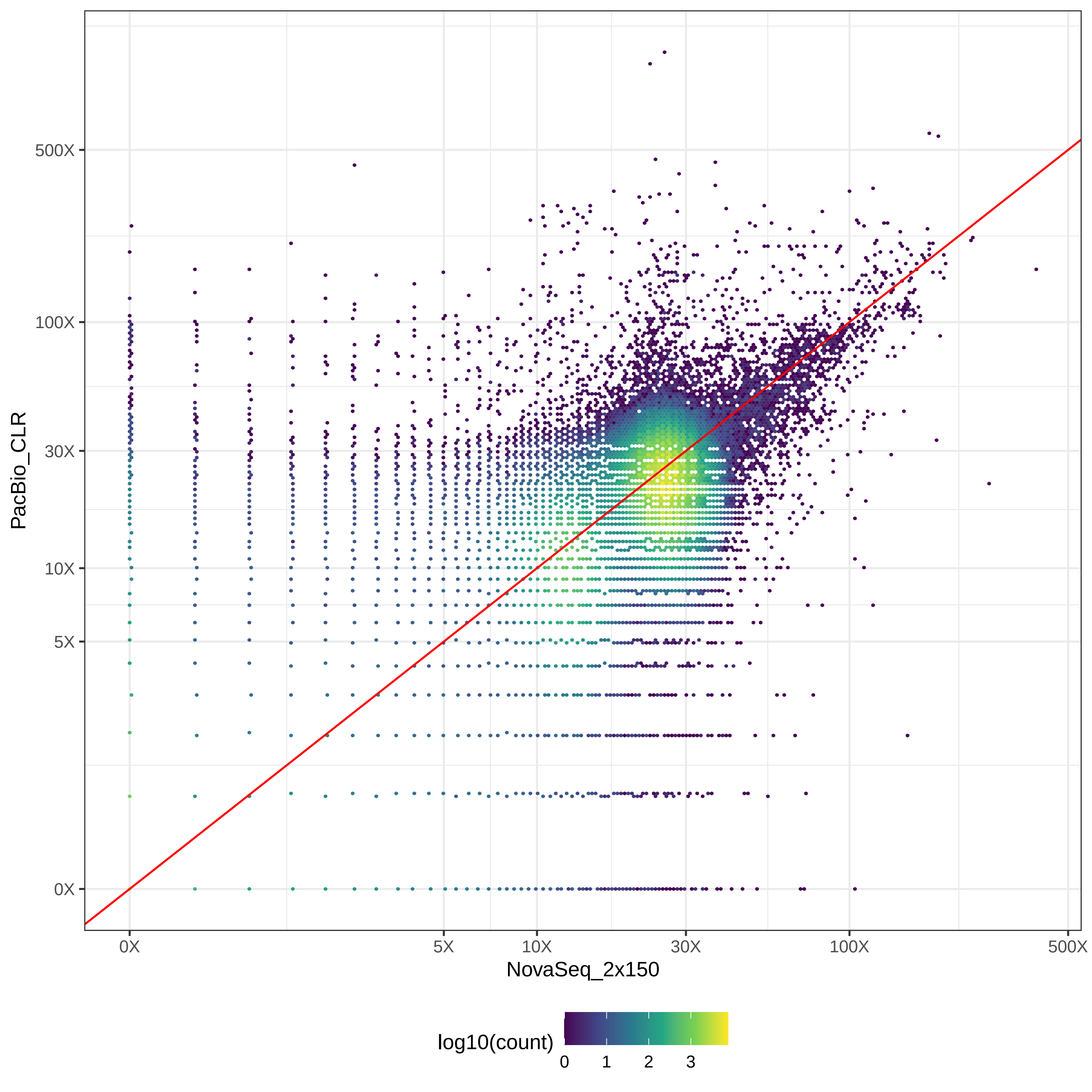

### GIAB_L1_NovaSeq_2x150-v-PromethION_μ=8092_hex.pdf

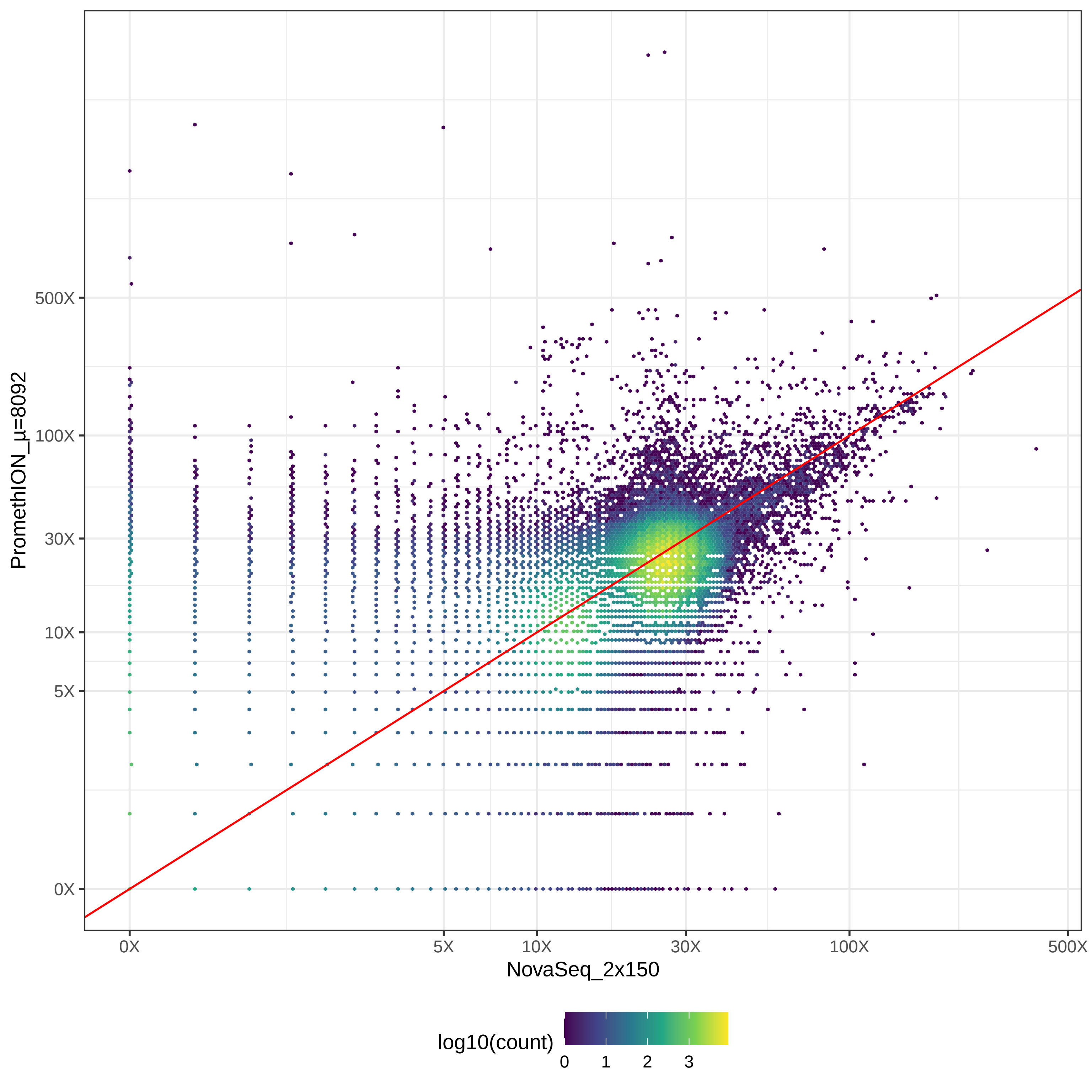

### GIAB_L1_NovaSeq_2x250-v-PacBio_CCS-15k_hex.pdf

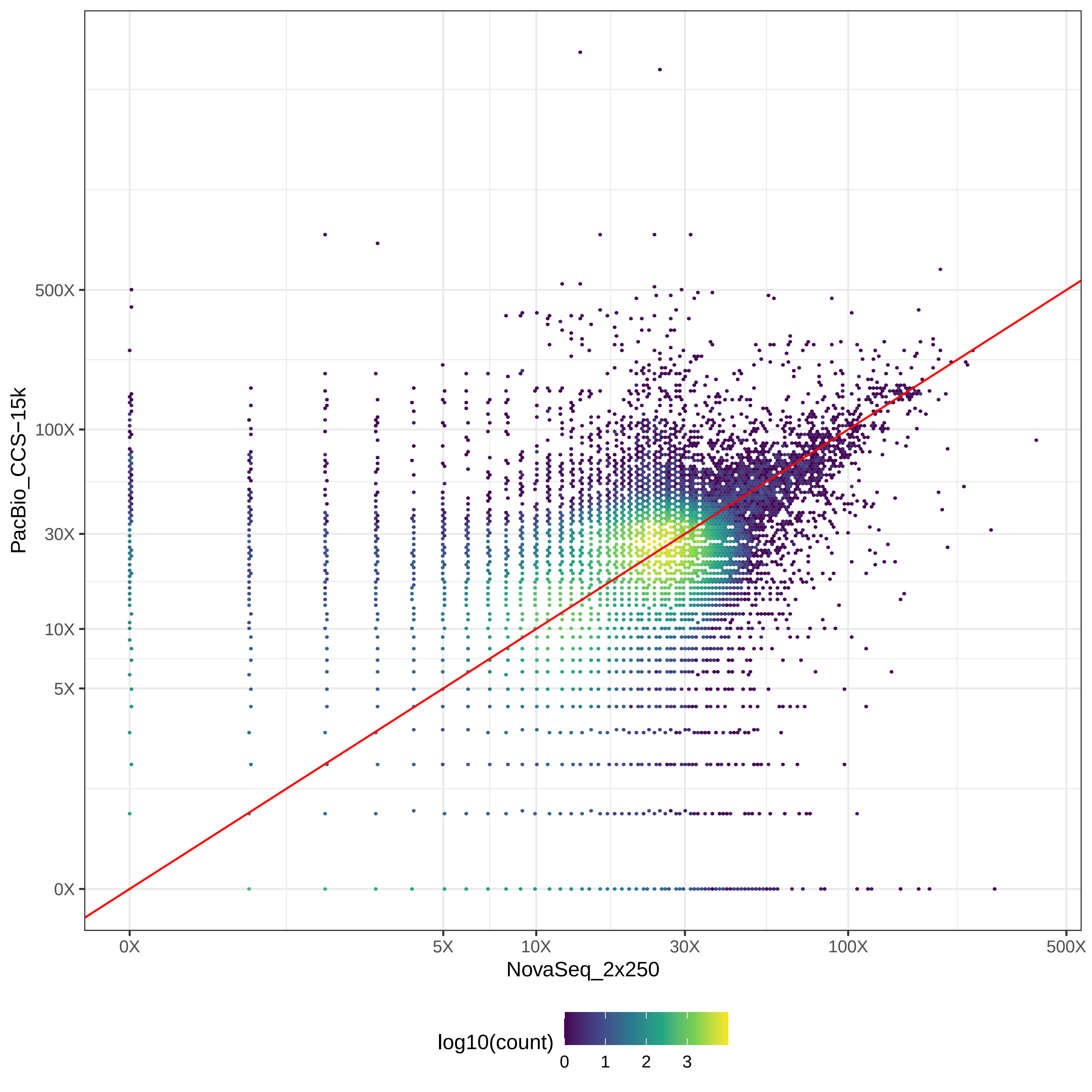

### GIAB_L1_NovaSeq_2x250-v-PacBio_CLR_hex.pdf

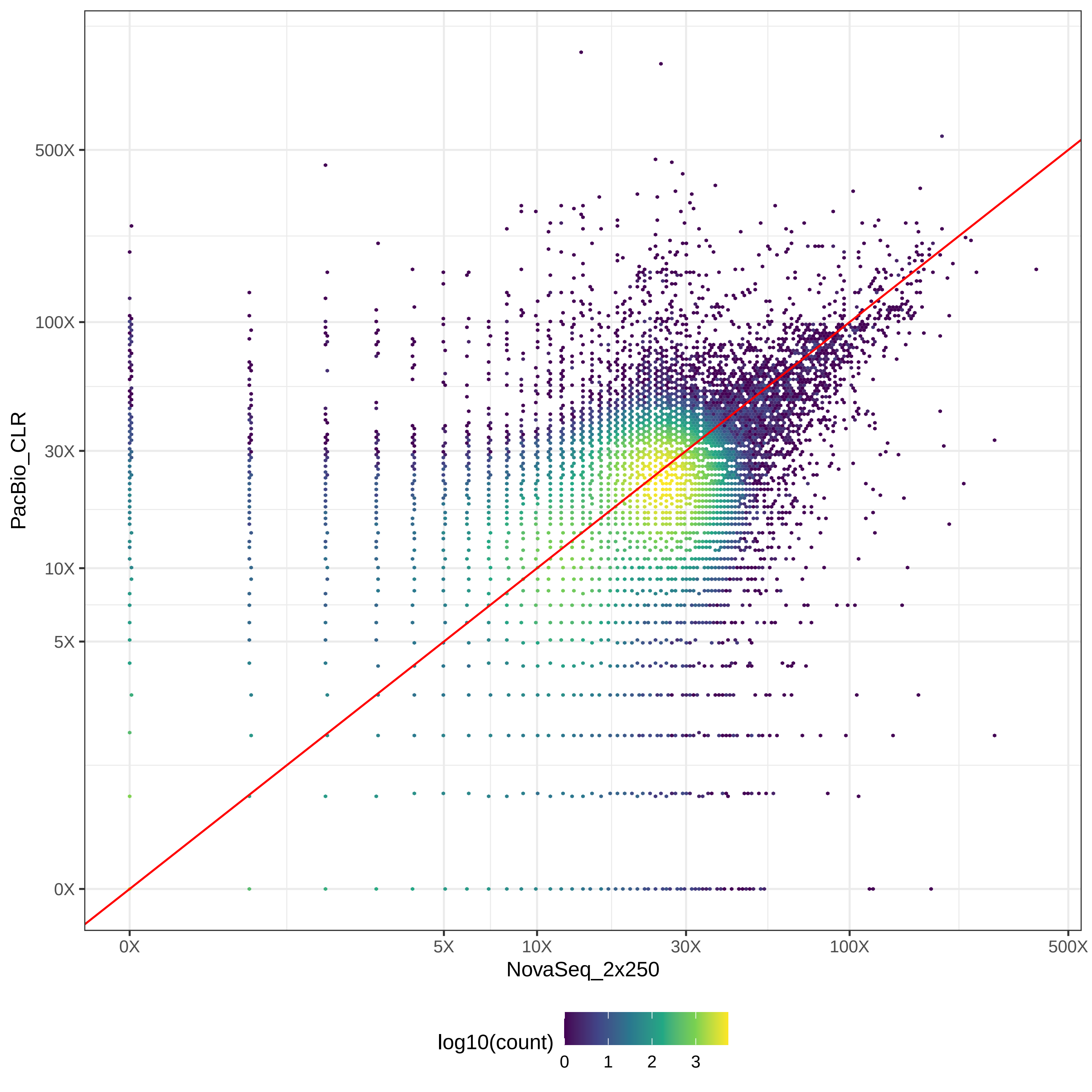

### GIAB_L1_NovaSeq_2x250-v-PromethION_μ=8092_hex.pdf

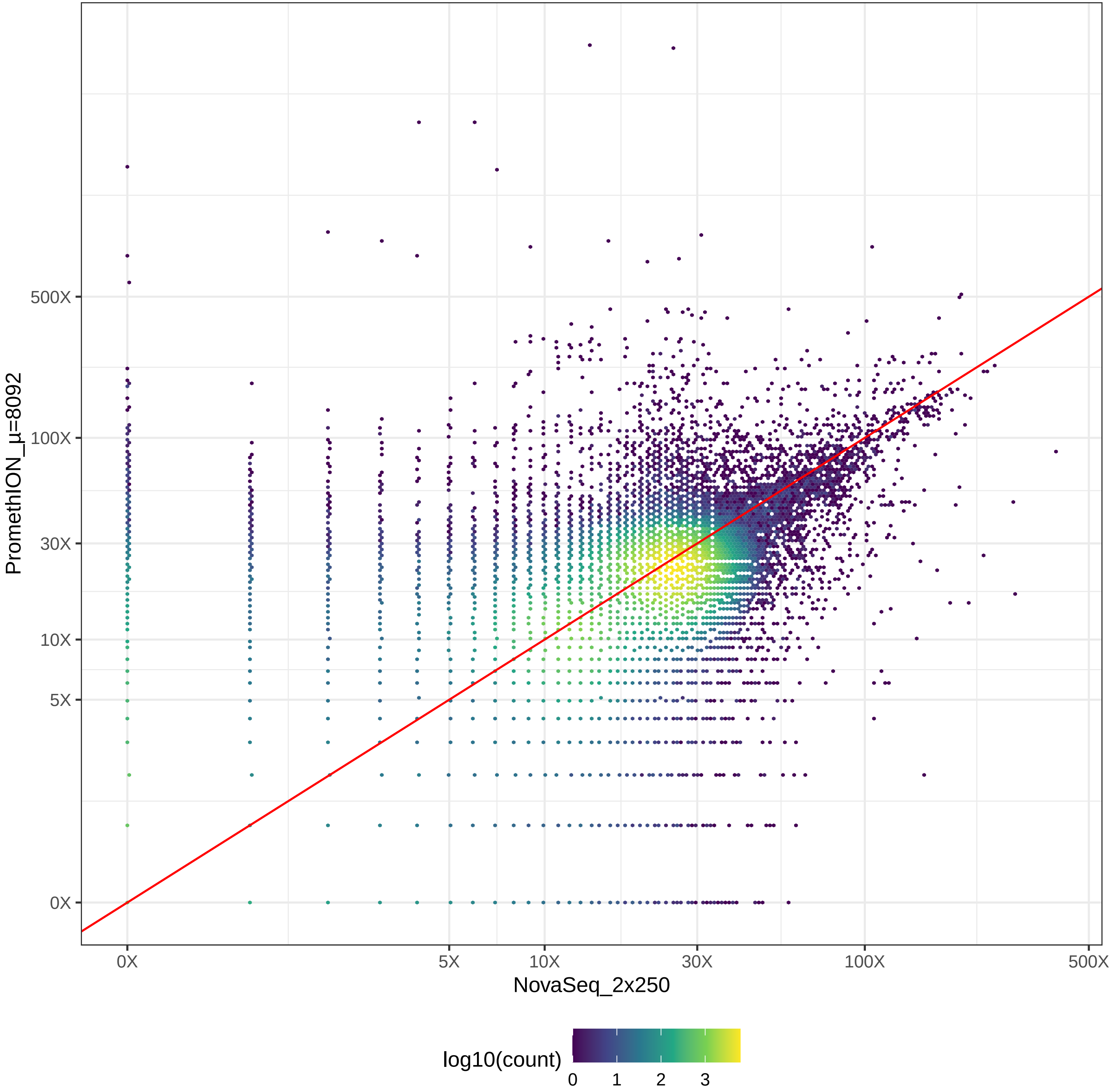

### GIAB_L1_PacBio_CCS-15k-v-PacBio_CLR_hex.pdf

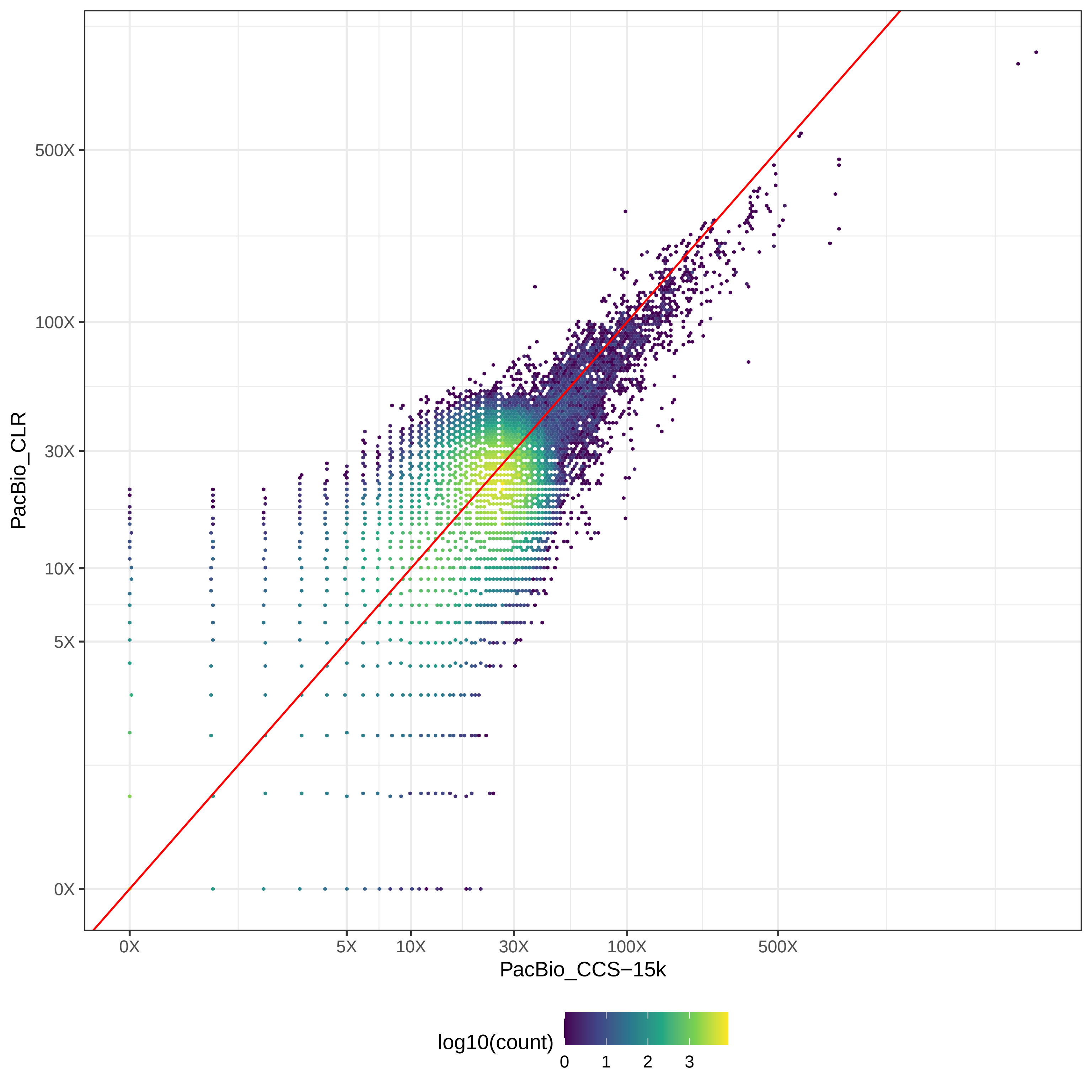

### GIAB_L1_PacBio_CCS-15k-v-PromethION_μ=8092_hex.pdf

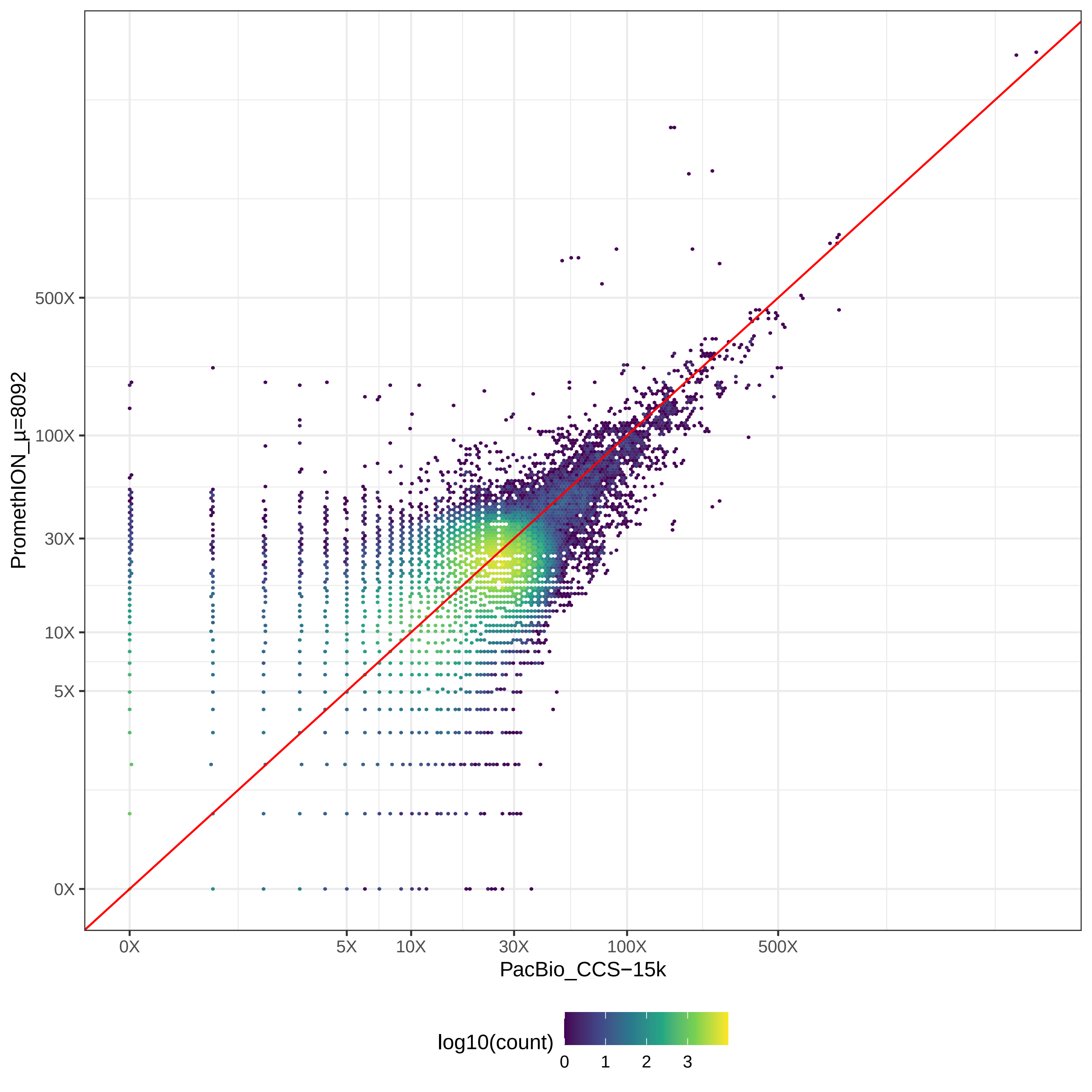

### GIAB_L1_PacBio_CLR-v-PromethION_μ=8092_hex.pdf

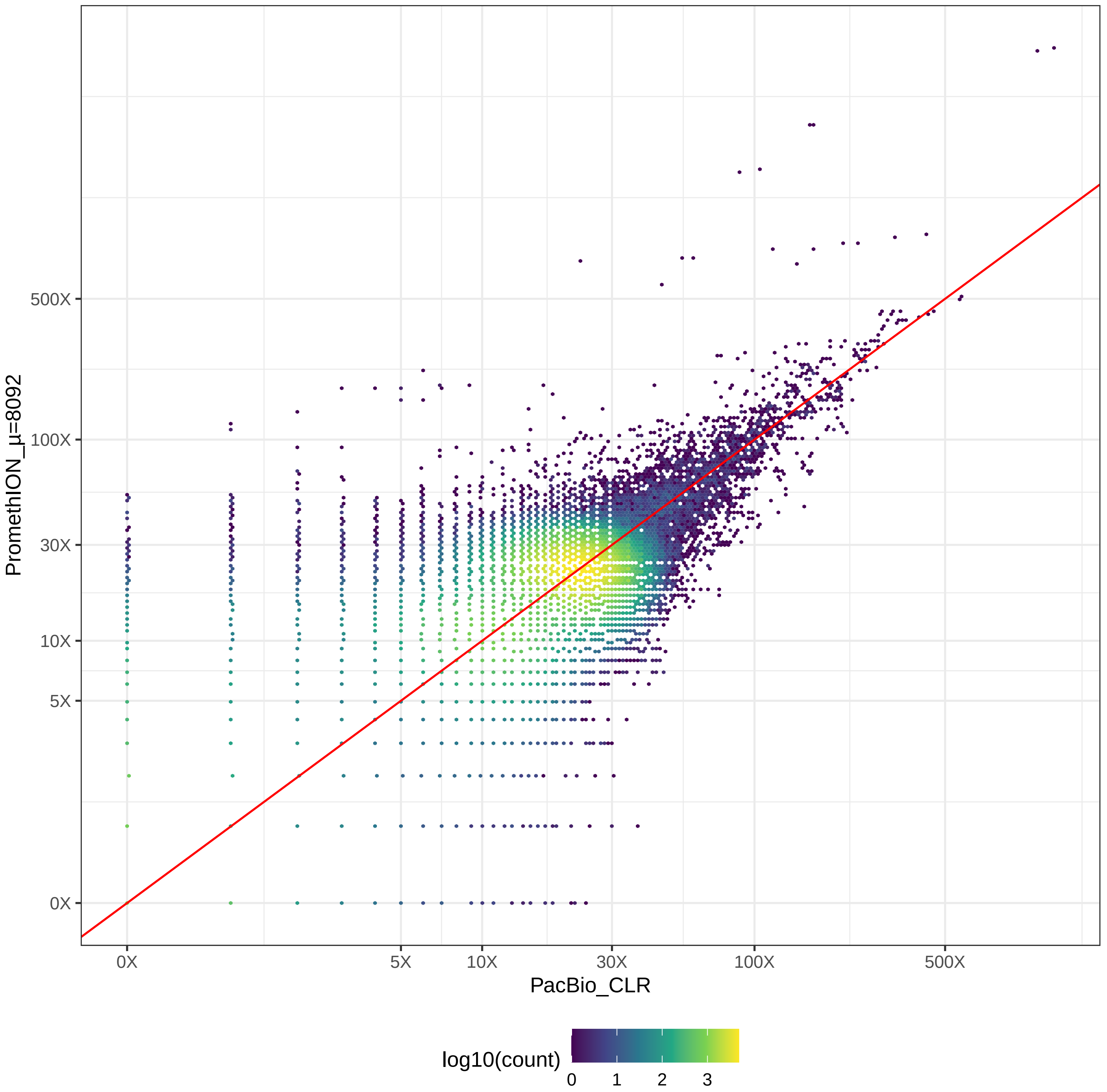

### GIAB_L2_BGISEQ500_2x100-v-HiSeq2500_2x100_hex.pdf

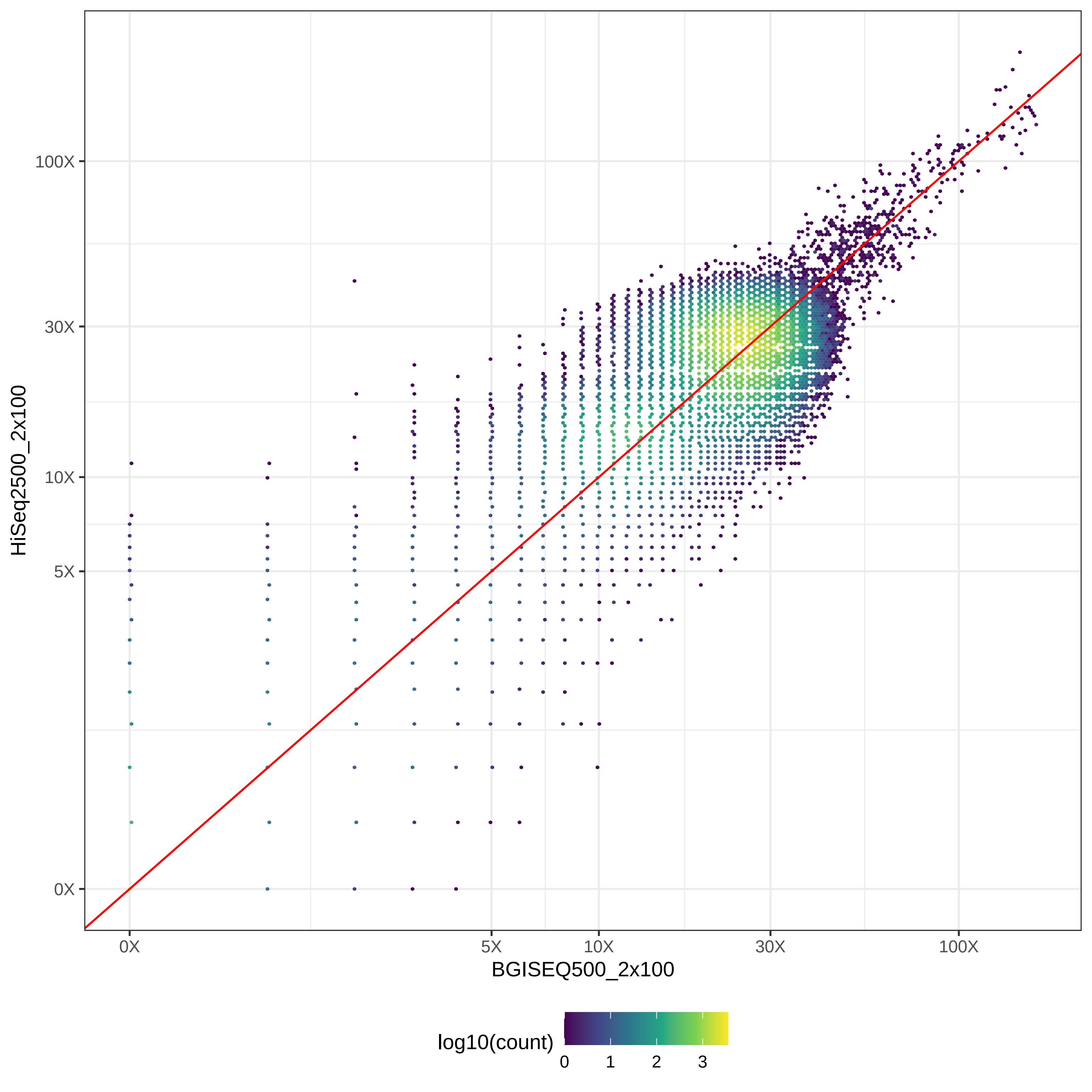

### GIAB_L2_BGISEQ500_2x100-v-HiSeq2500_2x126_hex.pdf

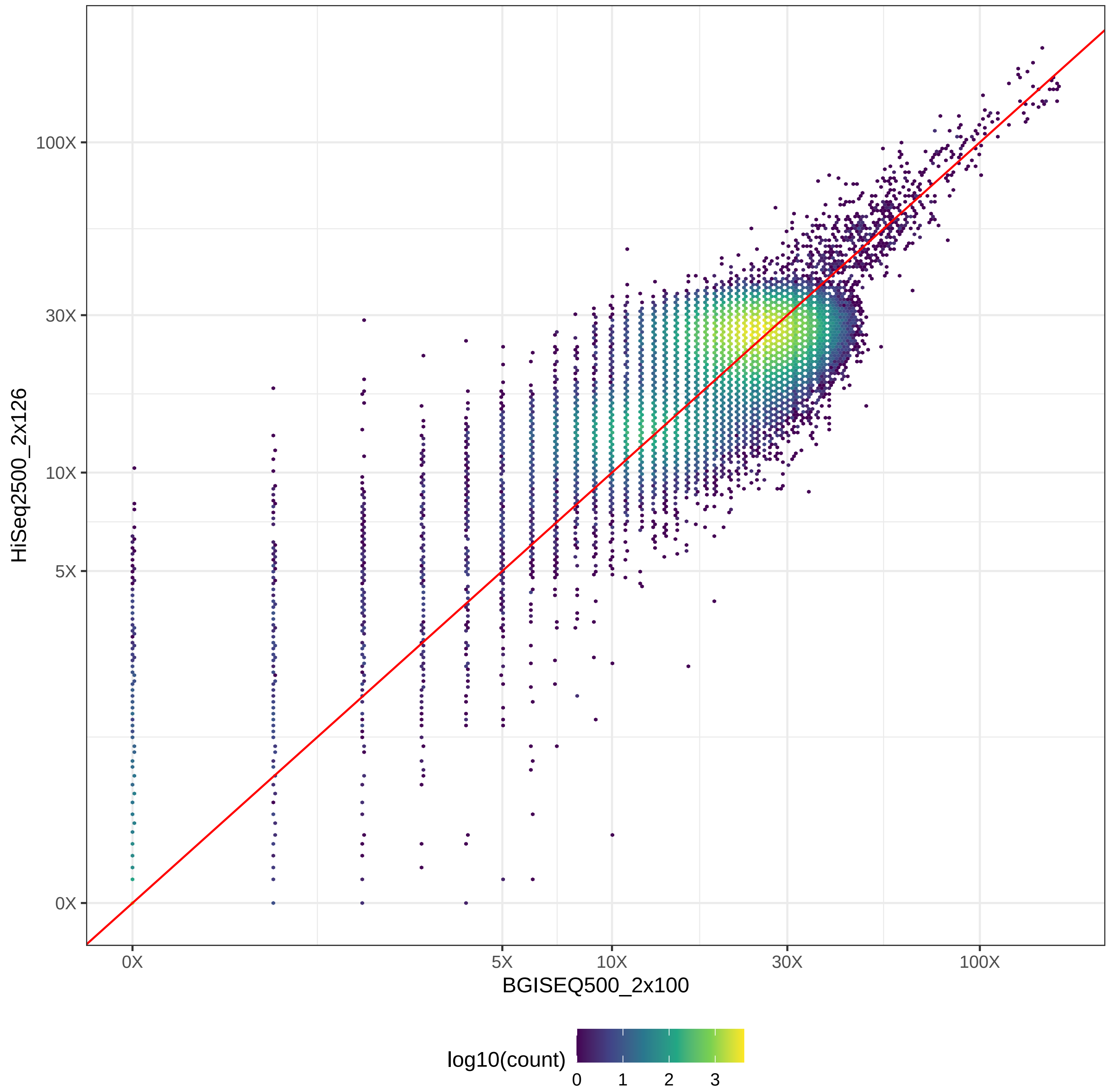

### GIAB_L2_BGISEQ500_2x100-v-HiSeq4000_2x150_hex.pdf

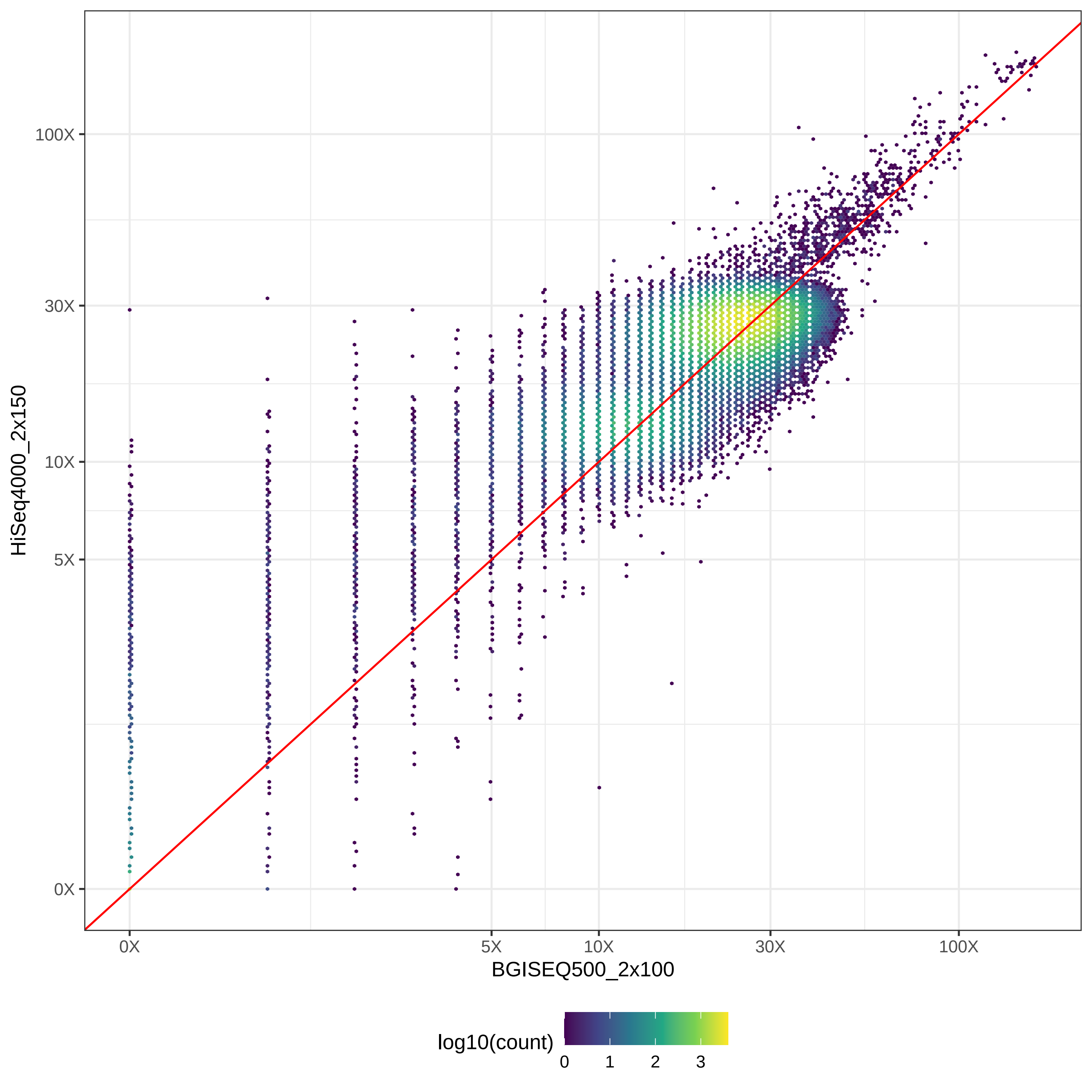

### GIAB_L2_BGISEQ500_2x100-v-HiSeqX10_2x150_hex.pdf

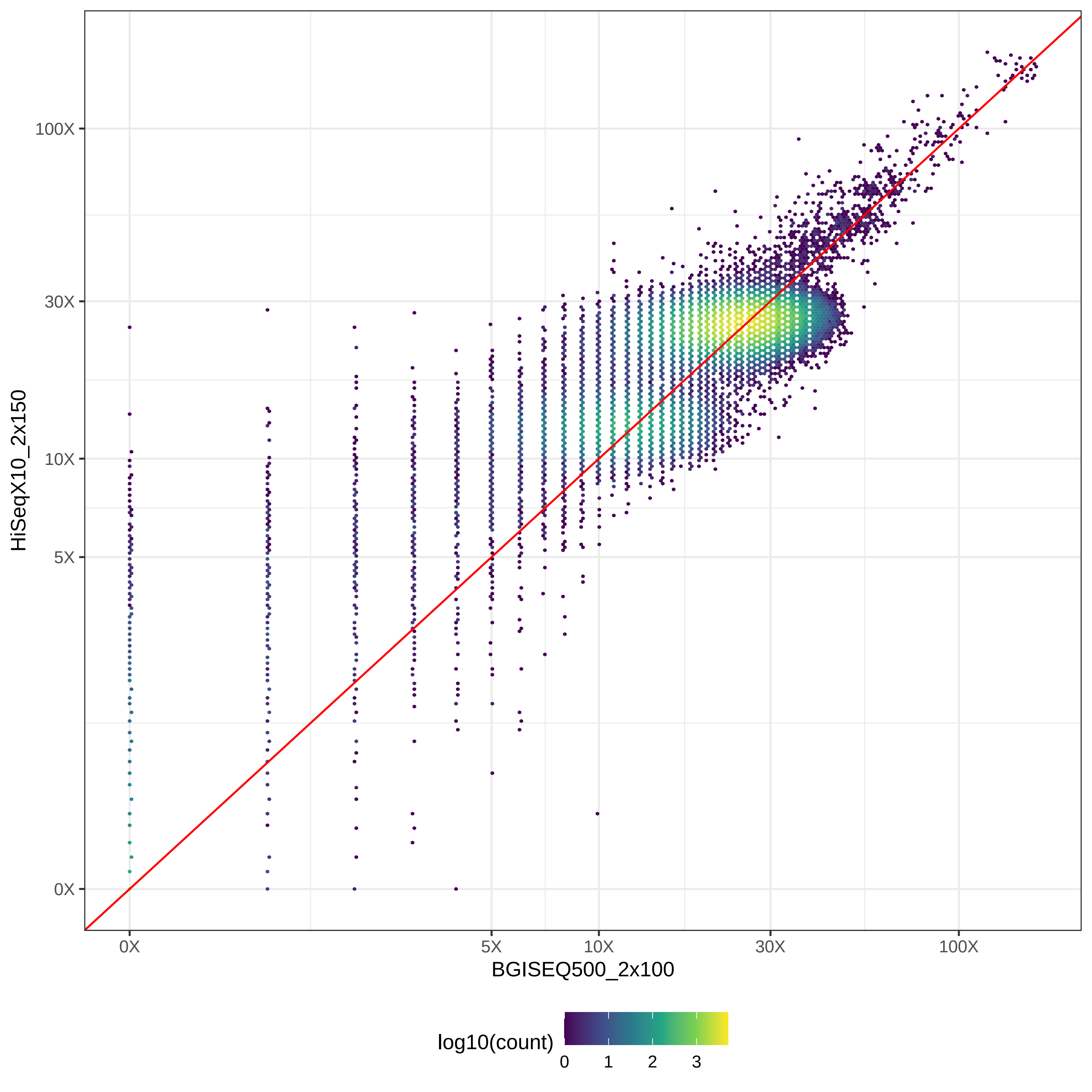

### GIAB_L2_BGISEQ500_2x100-v-MGISEQ2000_2x150_hex.pdf

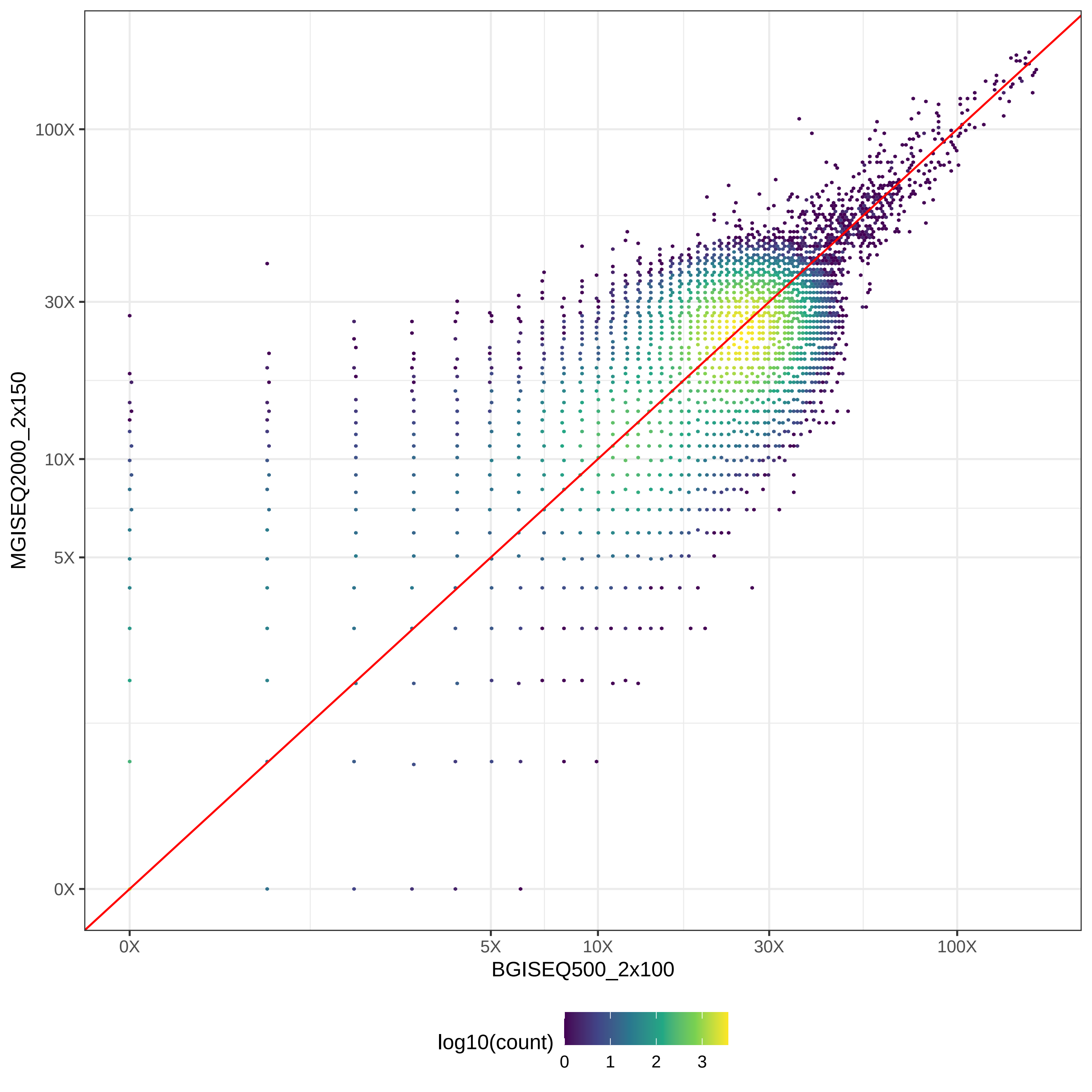

### GIAB_L2_BGISEQ500_2x100-v-NovaSeq_2x150_hex.pdf

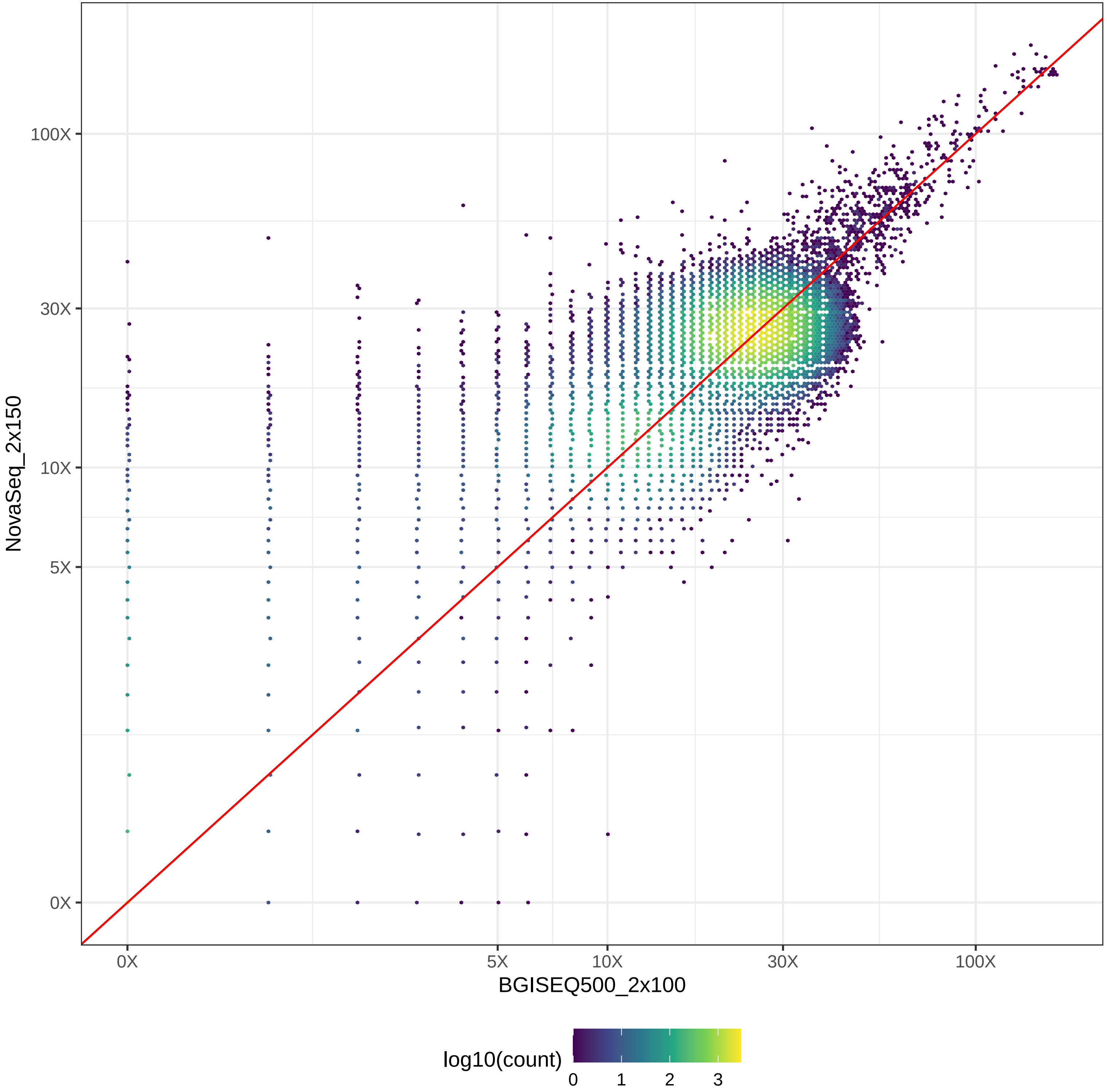

### GIAB_L2_BGISEQ500_2x100-v-NovaSeq_2x250_hex.pdf

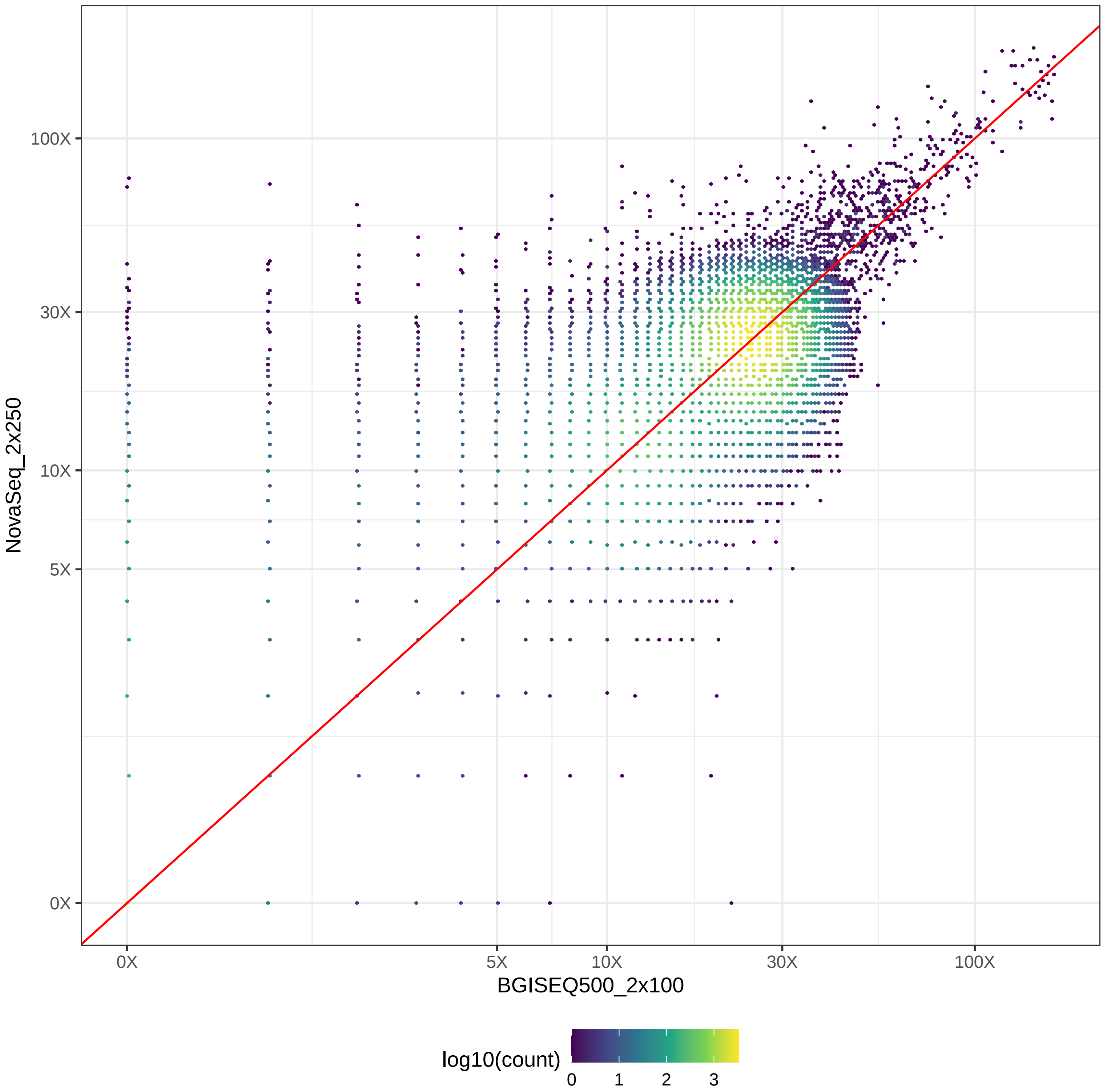

### GIAB_L2_BGISEQ500_2x100-v-PacBio_CCS-15k_hex.pdf

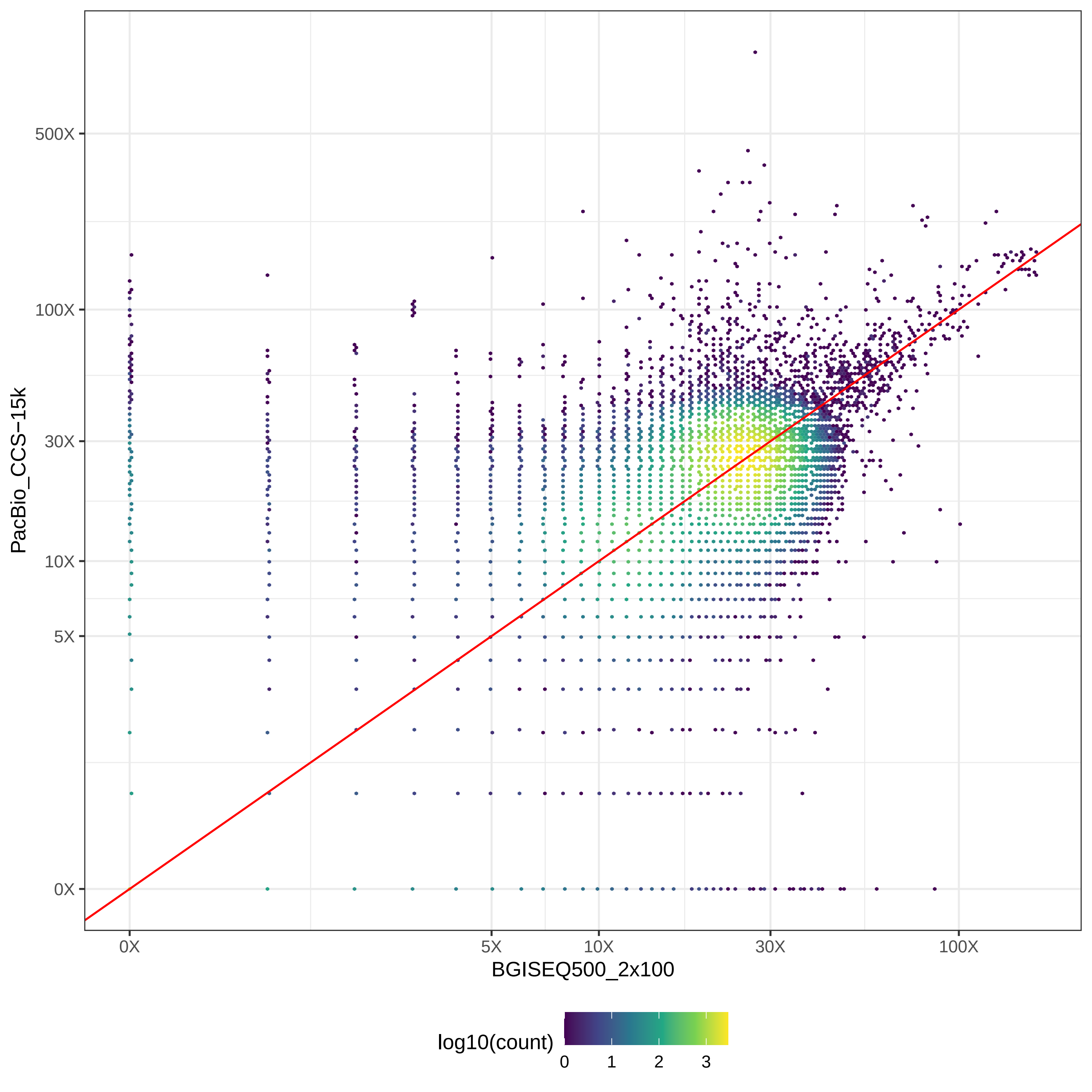

### GIAB_L2_BGISEQ500_2x100-v-PacBio_CLR_hex.pdf

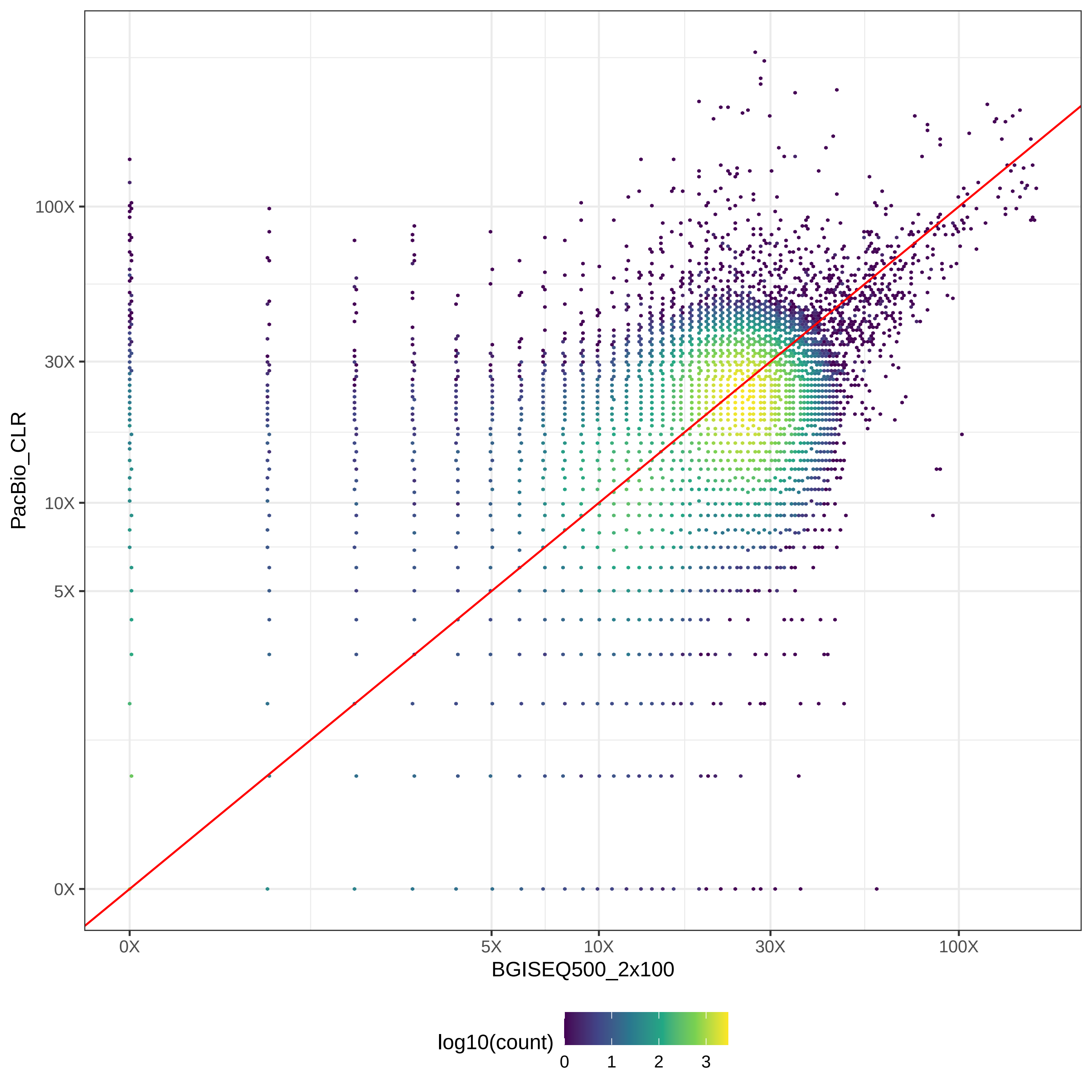

### GIAB_L2_BGISEQ500_2x100-v-PromethION_μ=8092_hex.pdf

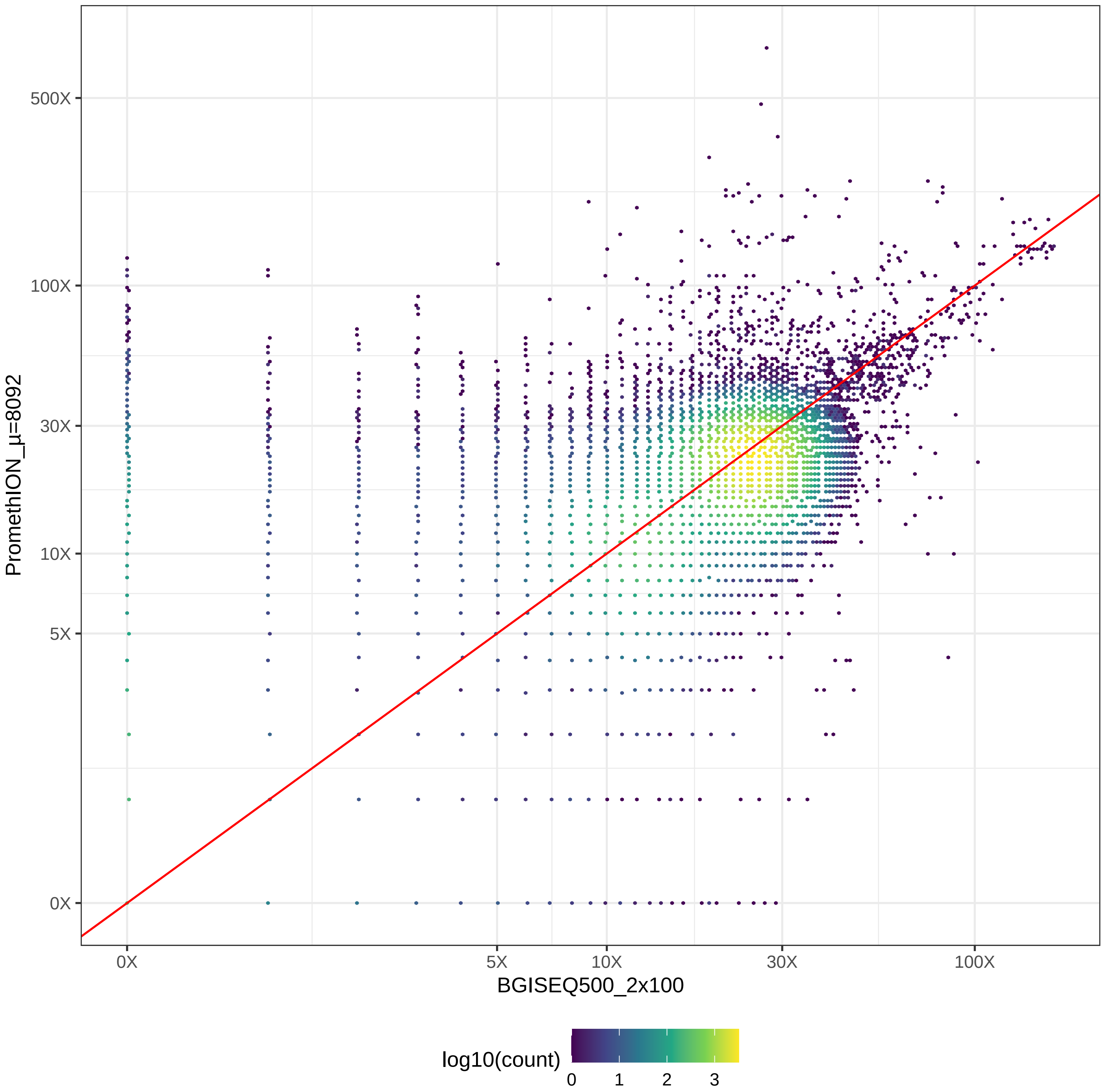

### GIAB_L2_HiSeq2500_2x100-v-HiSeq2500_2x126_hex.pdf

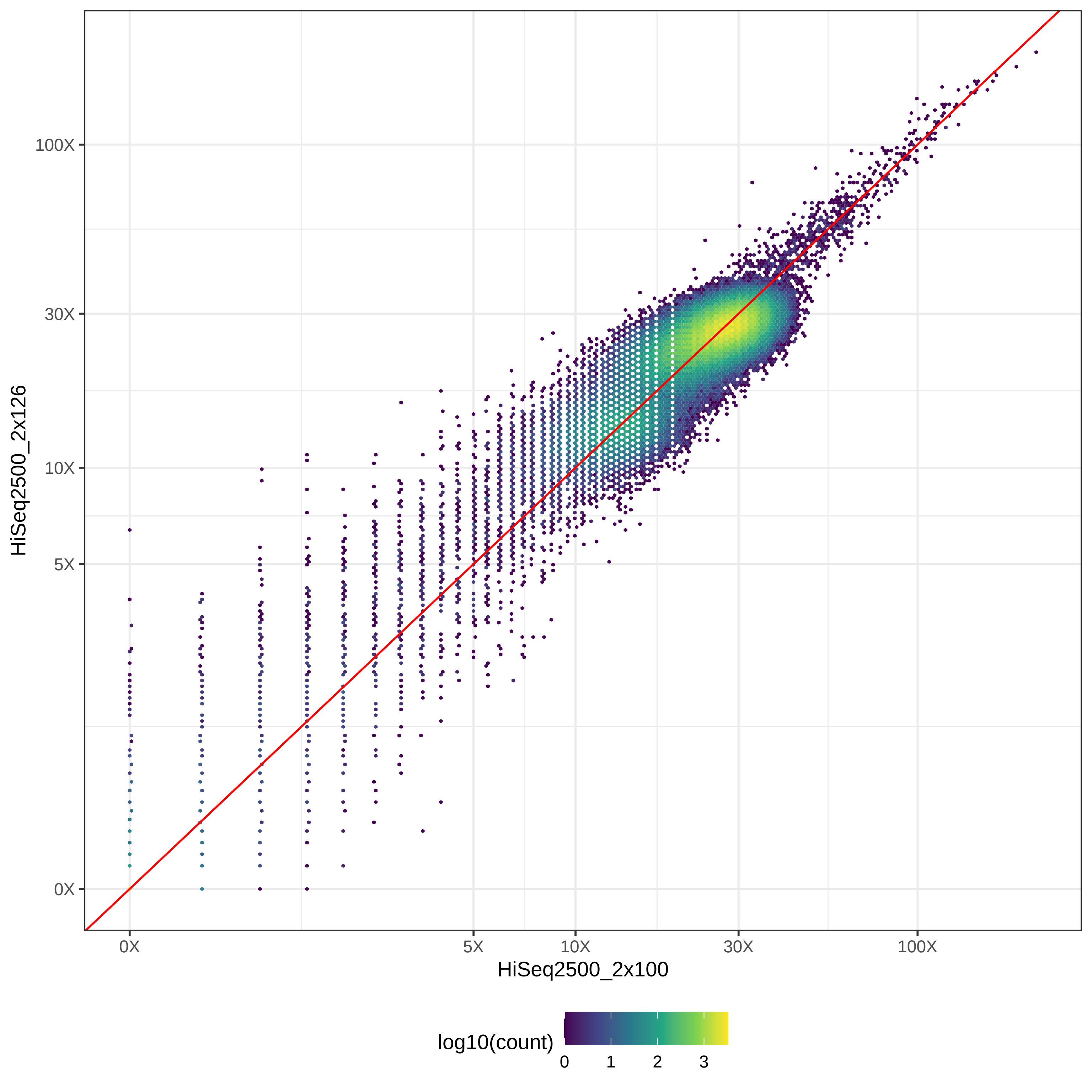

### GIAB_L2_HiSeq2500_2x100-v-HiSeq4000_2x150_hex.pdf

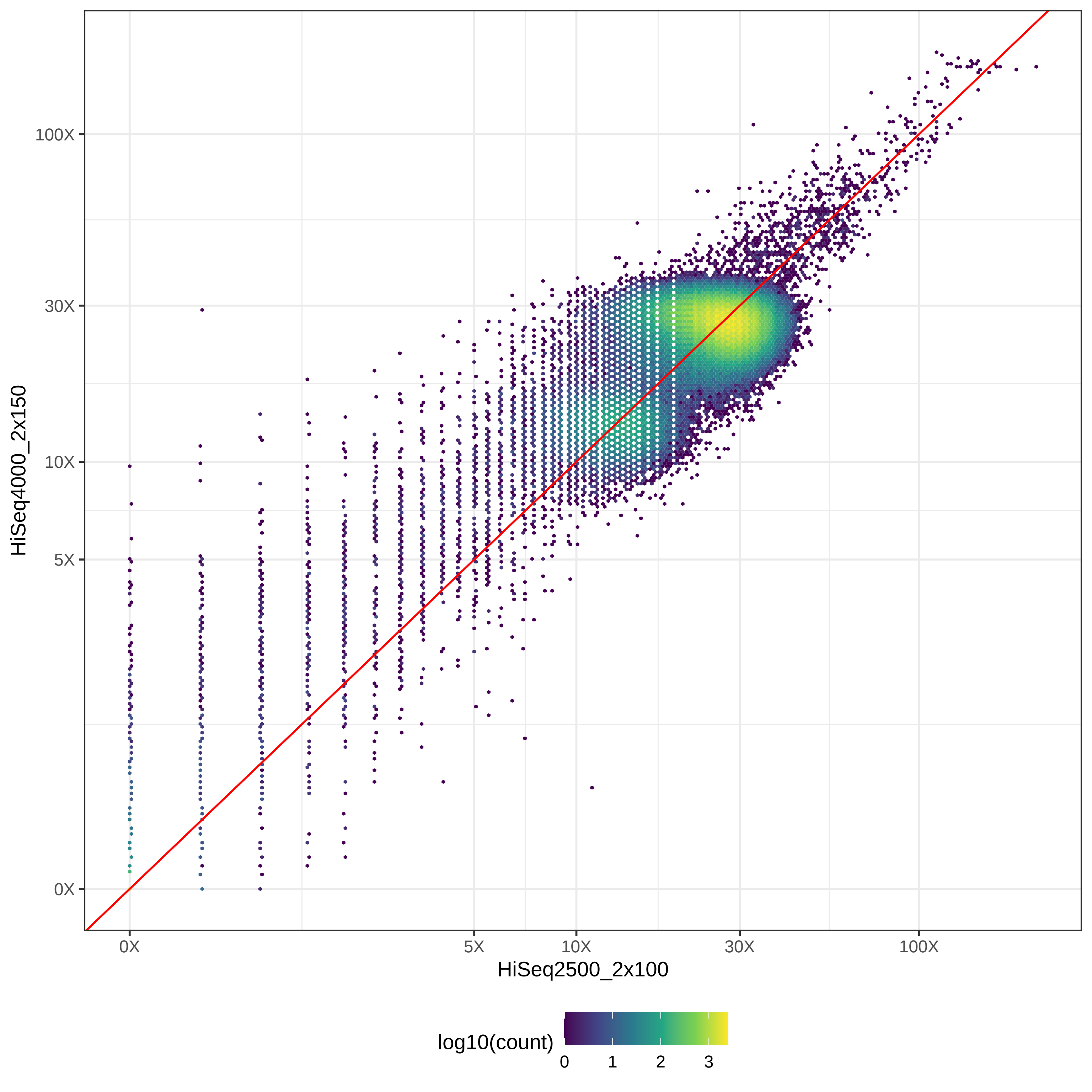

### GIAB_L2_HiSeq2500_2x100-v-HiSeqX10_2x150_hex.pdf

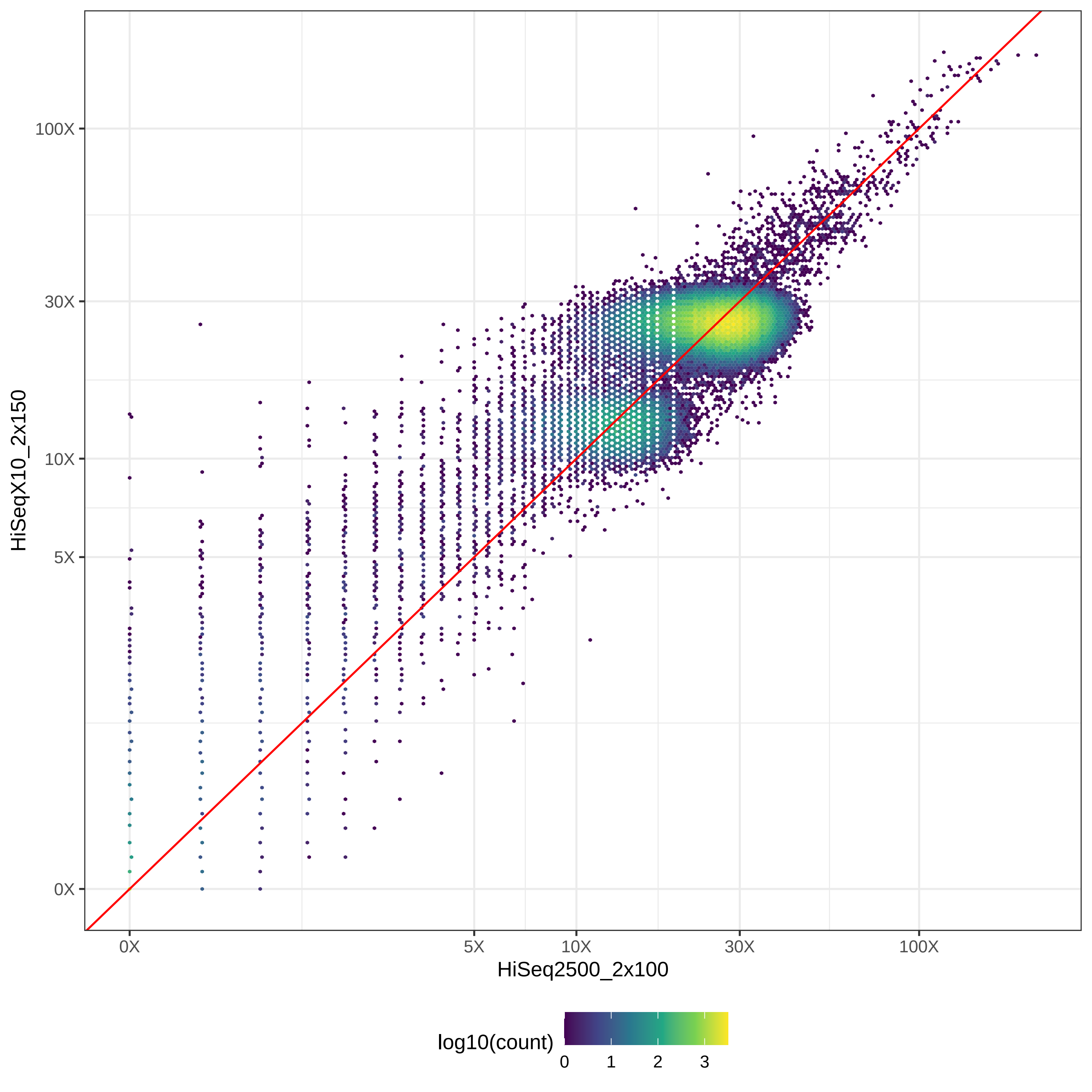

### GIAB_L2_HiSeq2500_2x100-v-MGISEQ2000_2x150_hex.pdf

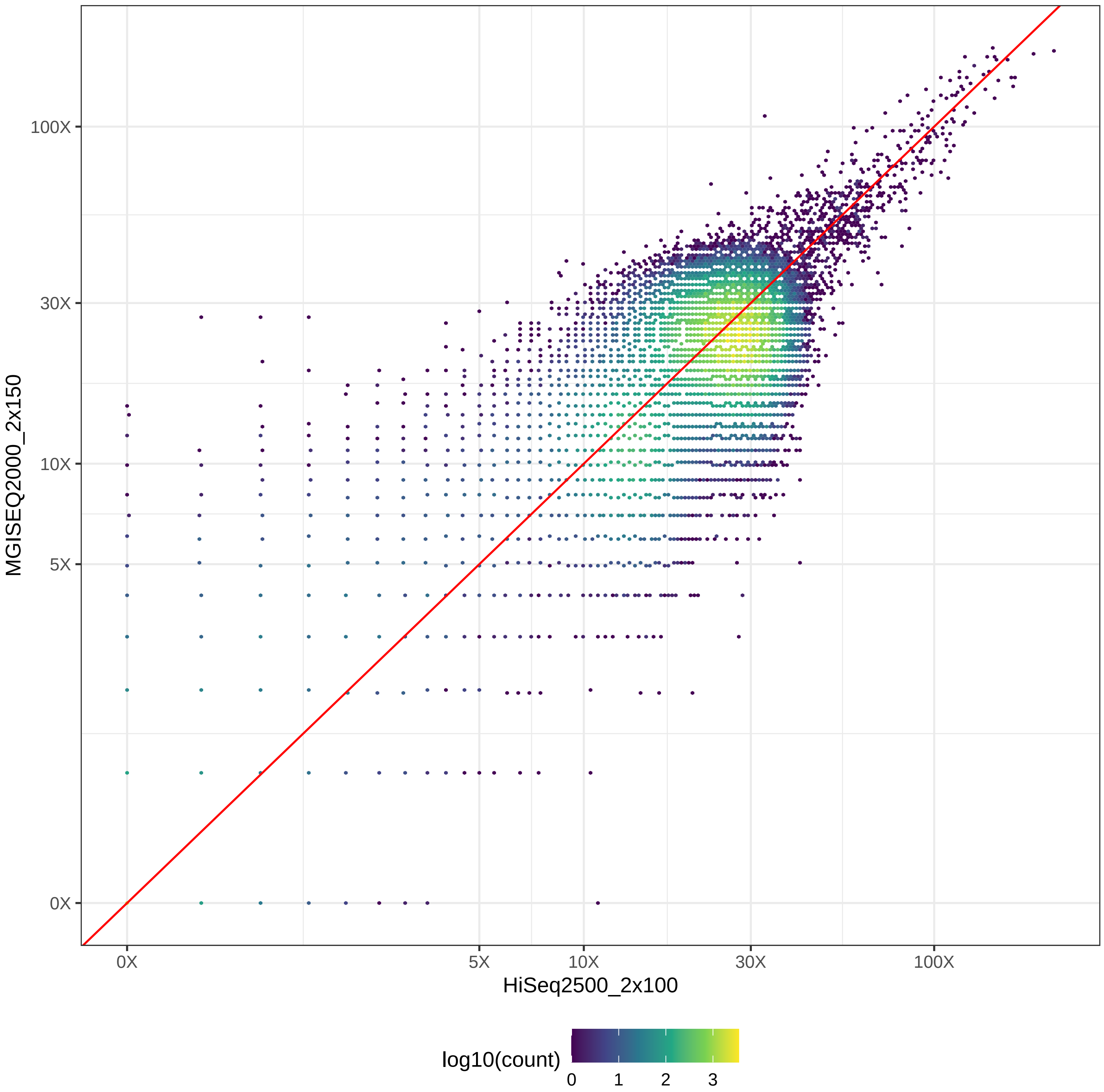

### GIAB_L2_HiSeq2500_2x100-v-NovaSeq_2x150_hex.pdf

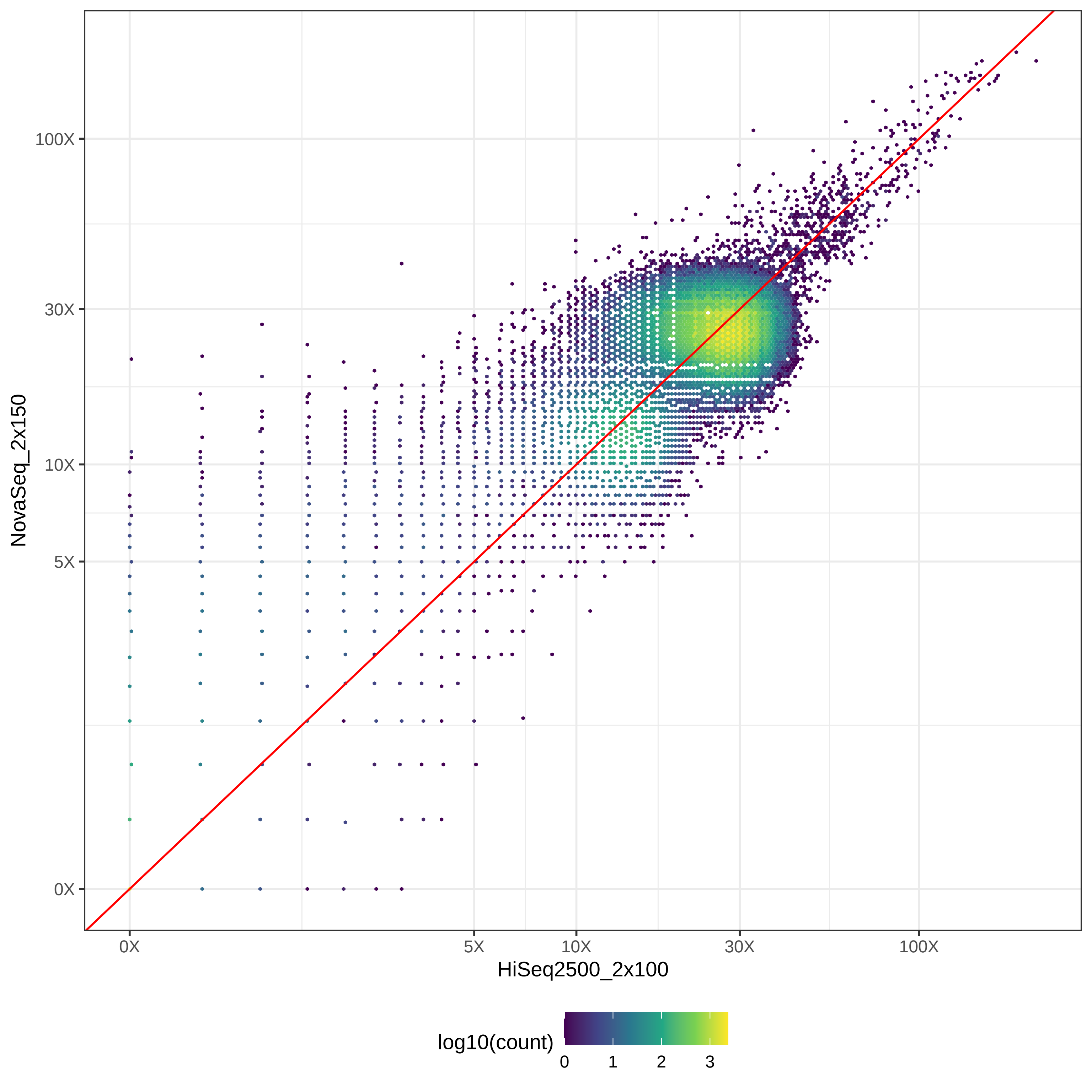

### GIAB_L2_HiSeq2500_2x100-v-NovaSeq_2x250_hex.pdf

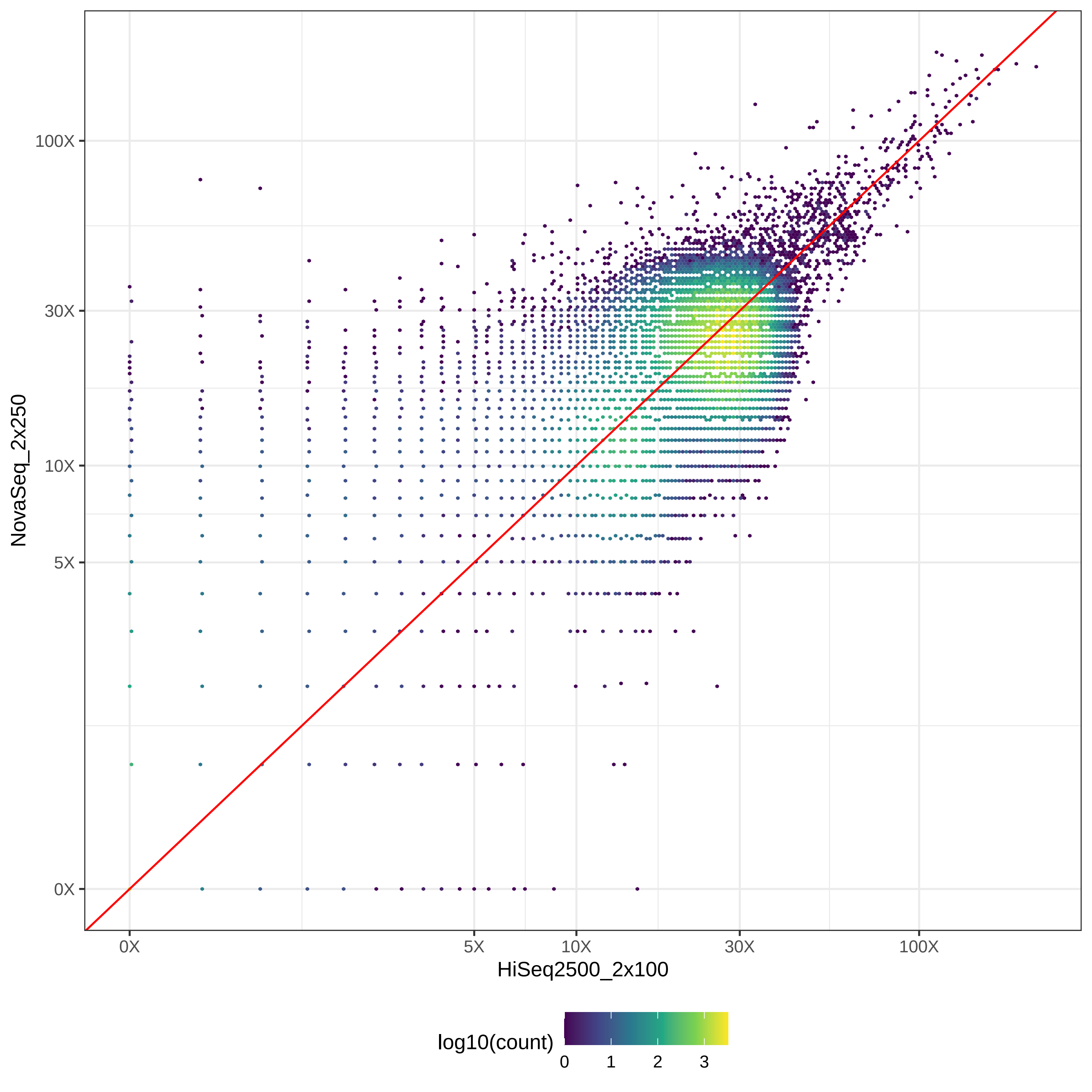

### GIAB_L2_HiSeq2500_2x100-v-PacBio_CCS-15k_hex.pdf

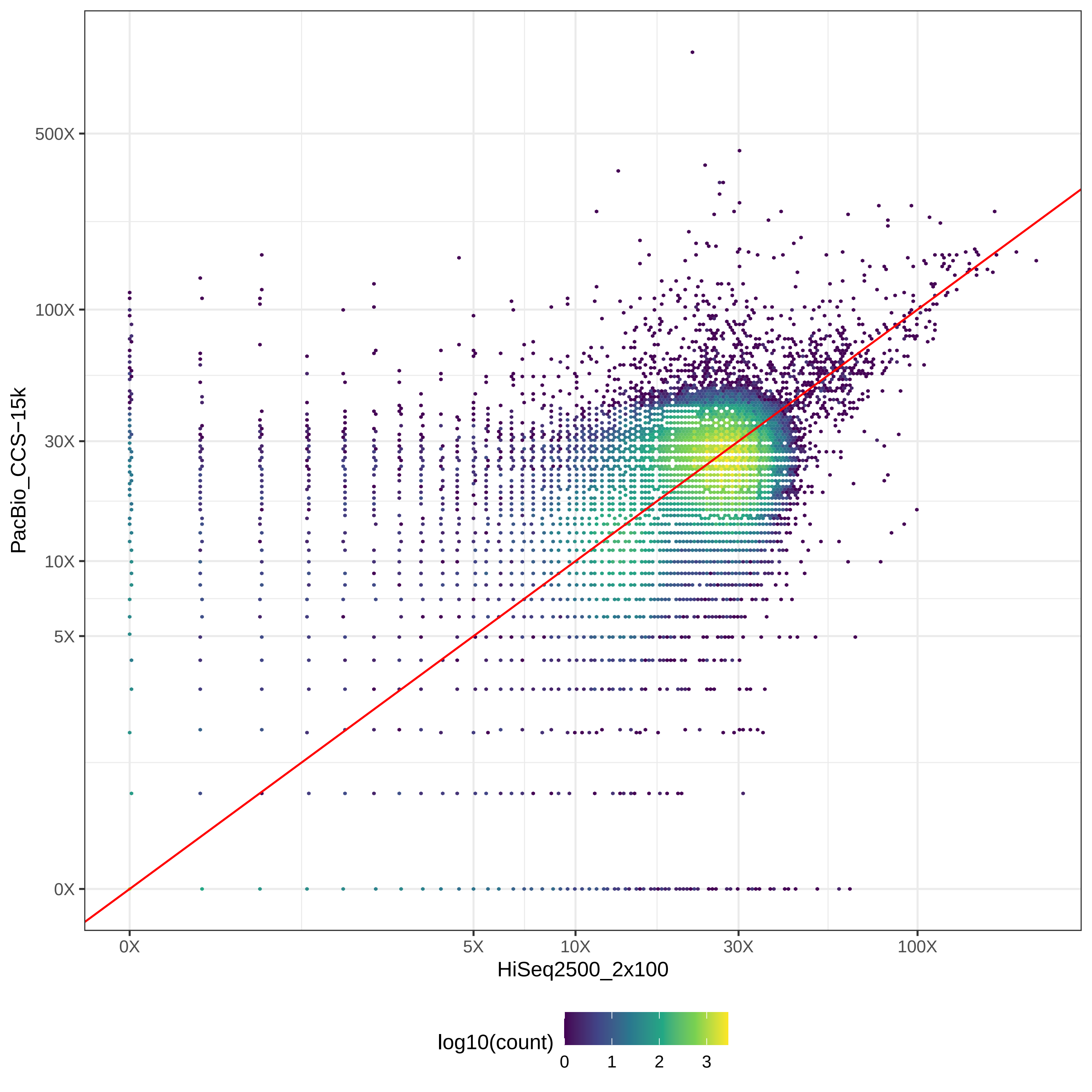

### GIAB_L2_HiSeq2500_2x100-v-PacBio_CLR_hex.pdf

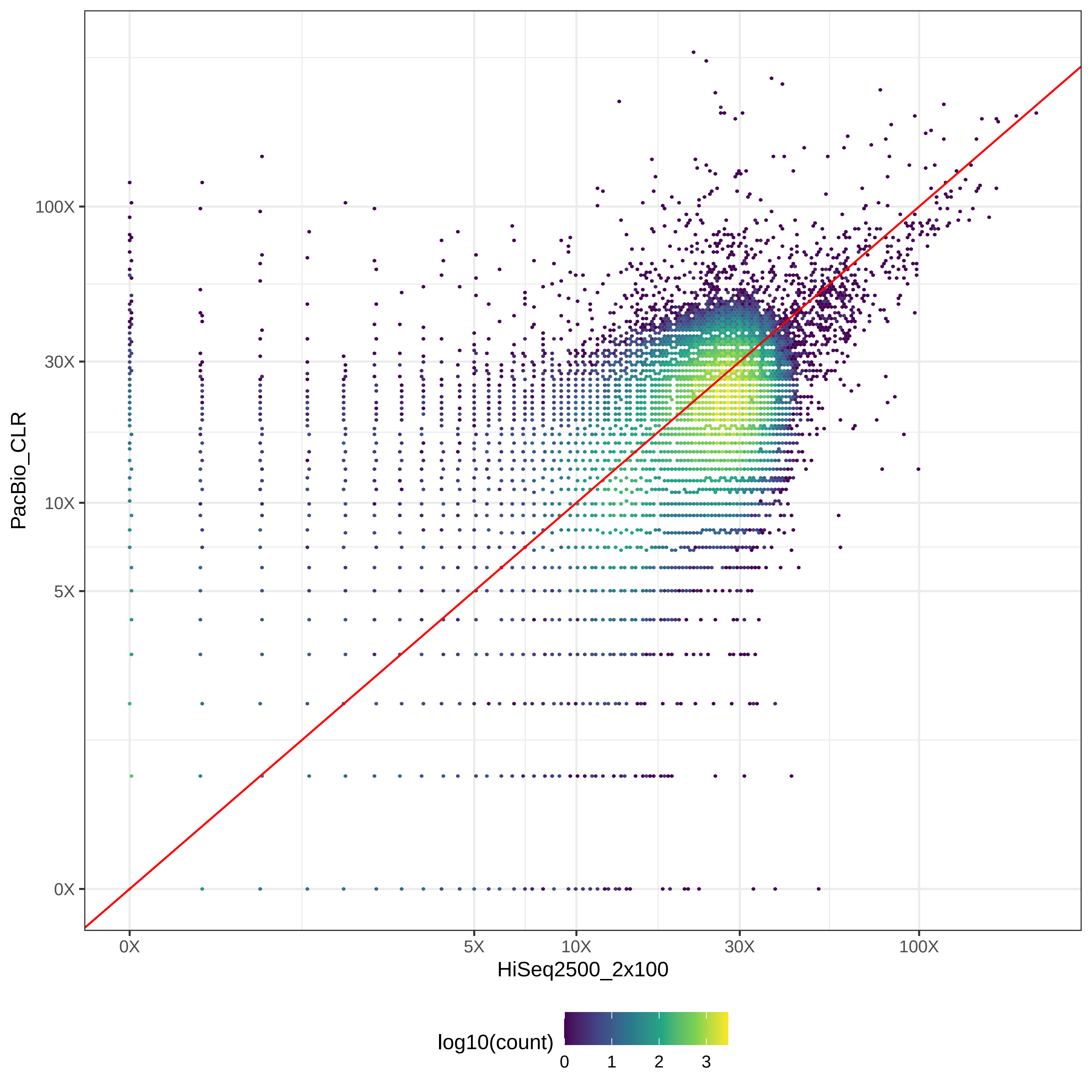

### GIAB_L2_HiSeq2500_2x100-v-PromethION_μ=8092_hex.pdf

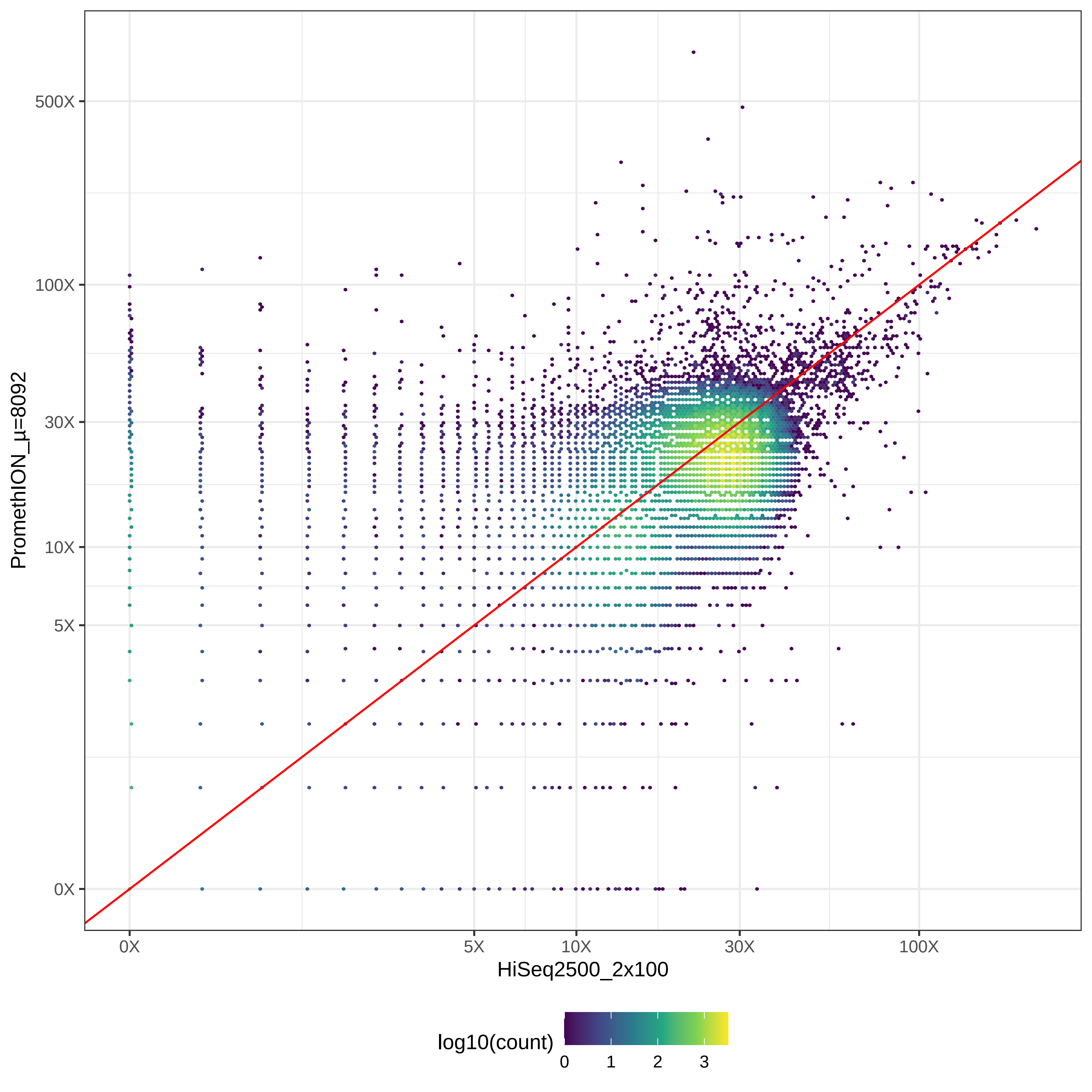

### GIAB_L2_HiSeq2500_2x126-v-HiSeq4000_2x150_hex.pdf

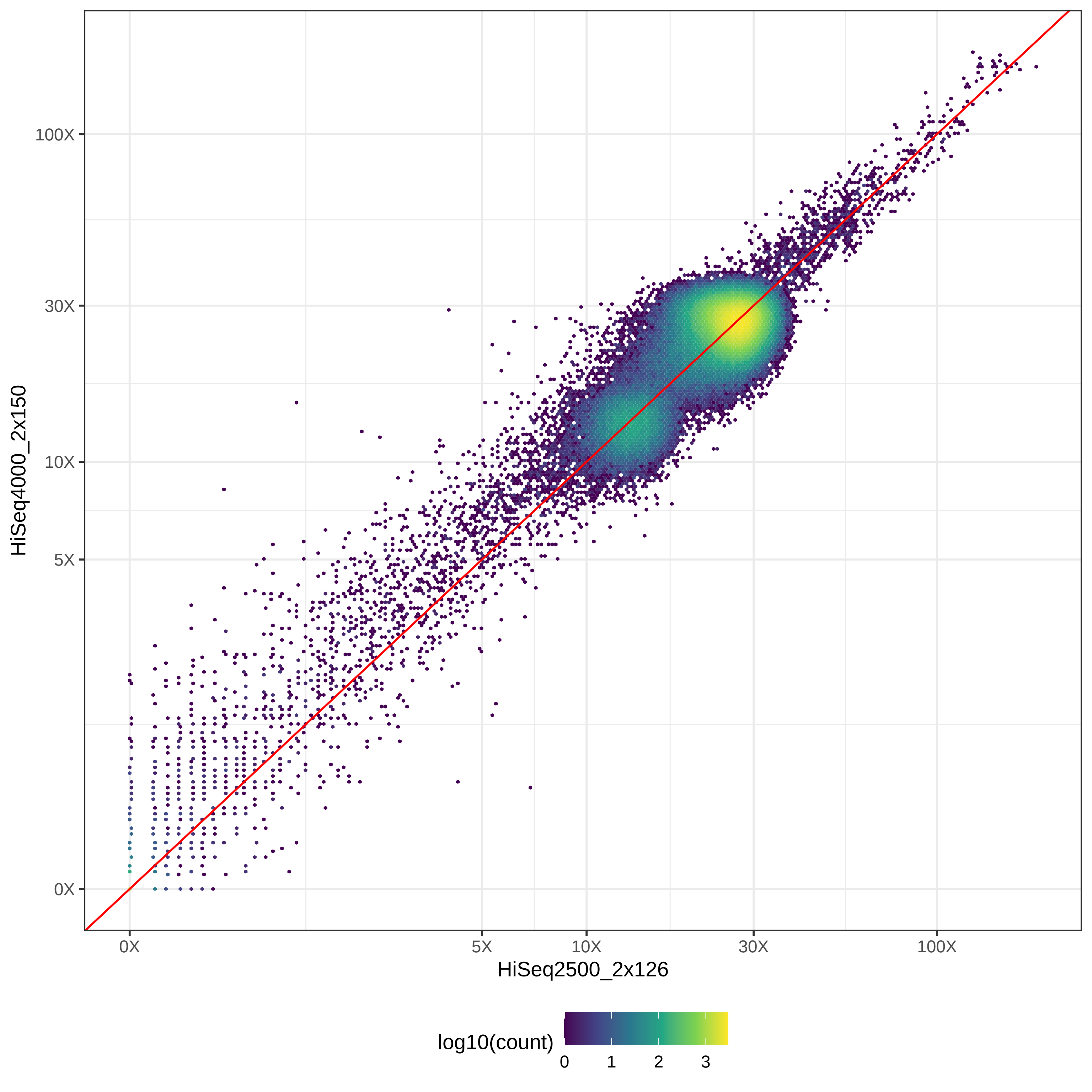

### GIAB_L2_HiSeq2500_2x126-v-HiSeqX10_2x150_hex.pdf

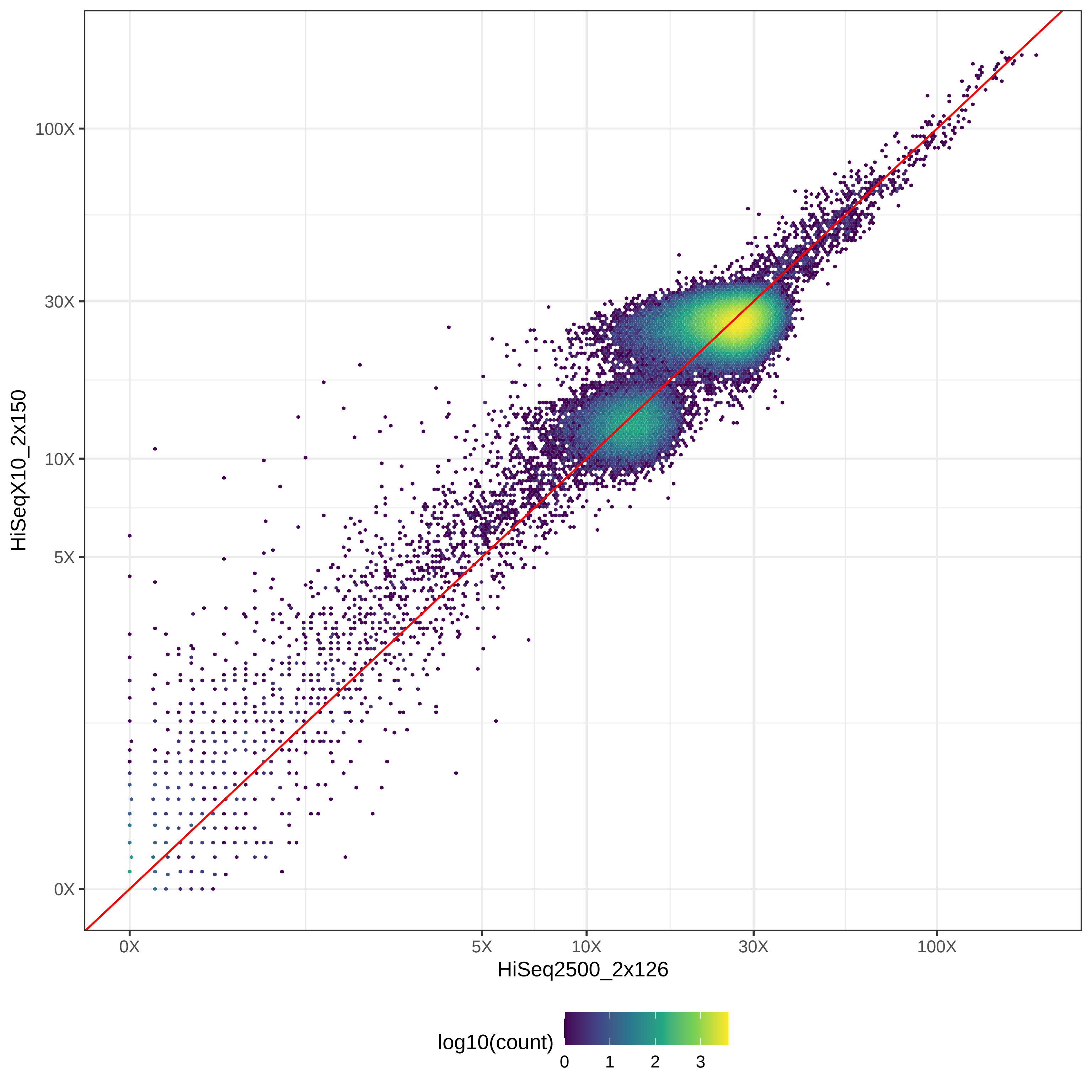

### GIAB_L2_HiSeq2500_2x126-v-MGISEQ2000_2x150_hex.pdf

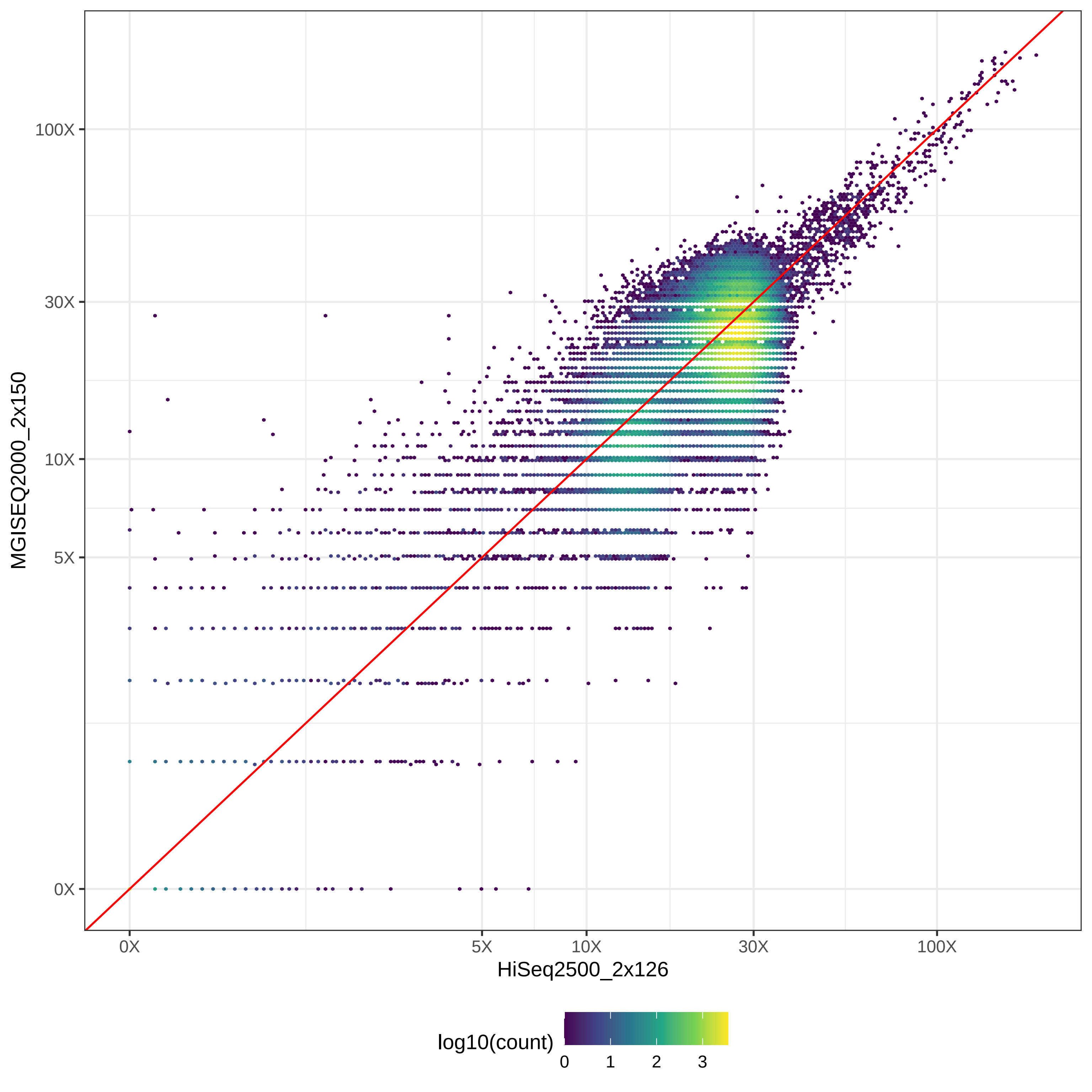

### GIAB_L2_HiSeq2500_2x126-v-NovaSeq_2x150_hex.pdf

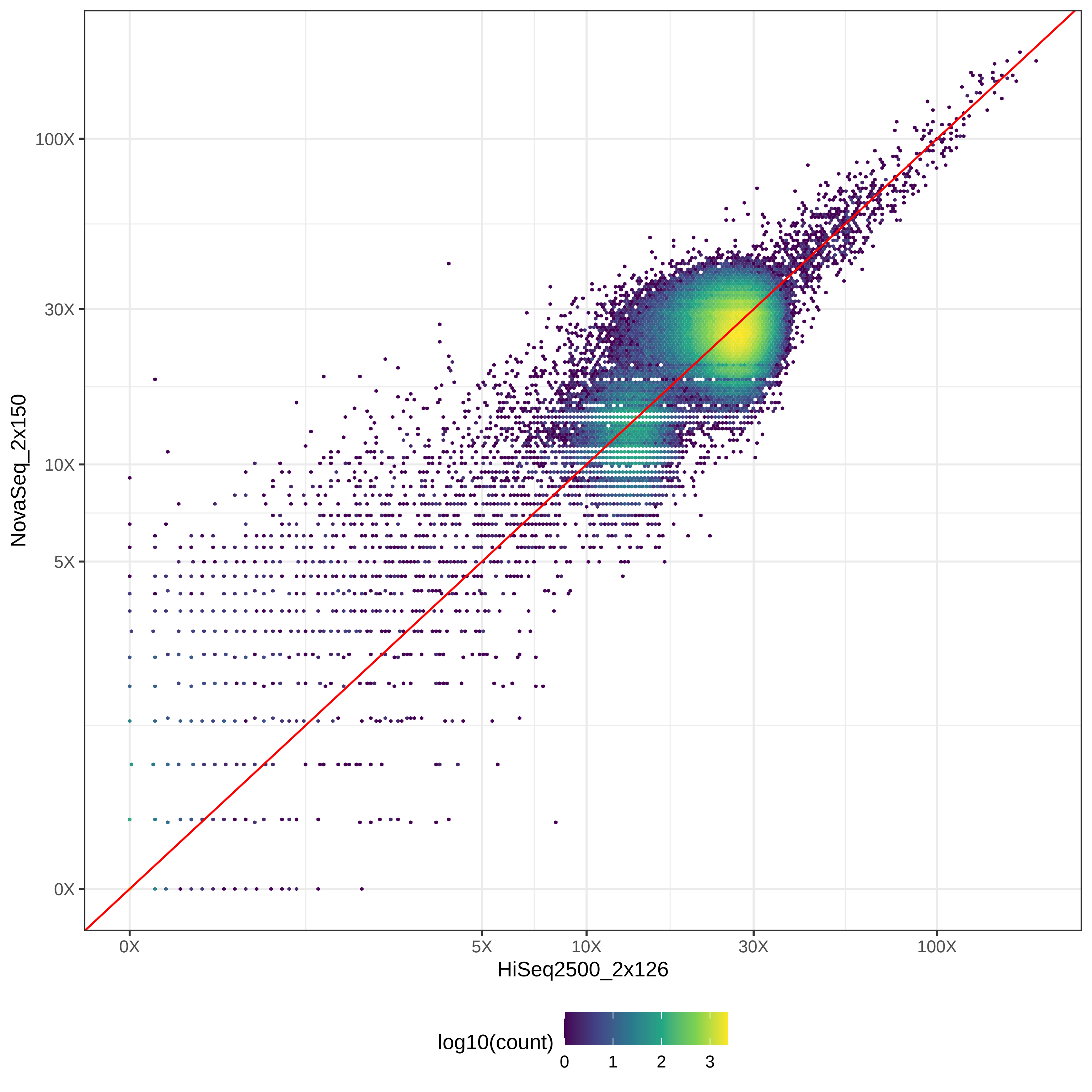

### GIAB_L2_HiSeq2500_2x126-v-NovaSeq_2x250_hex.pdf

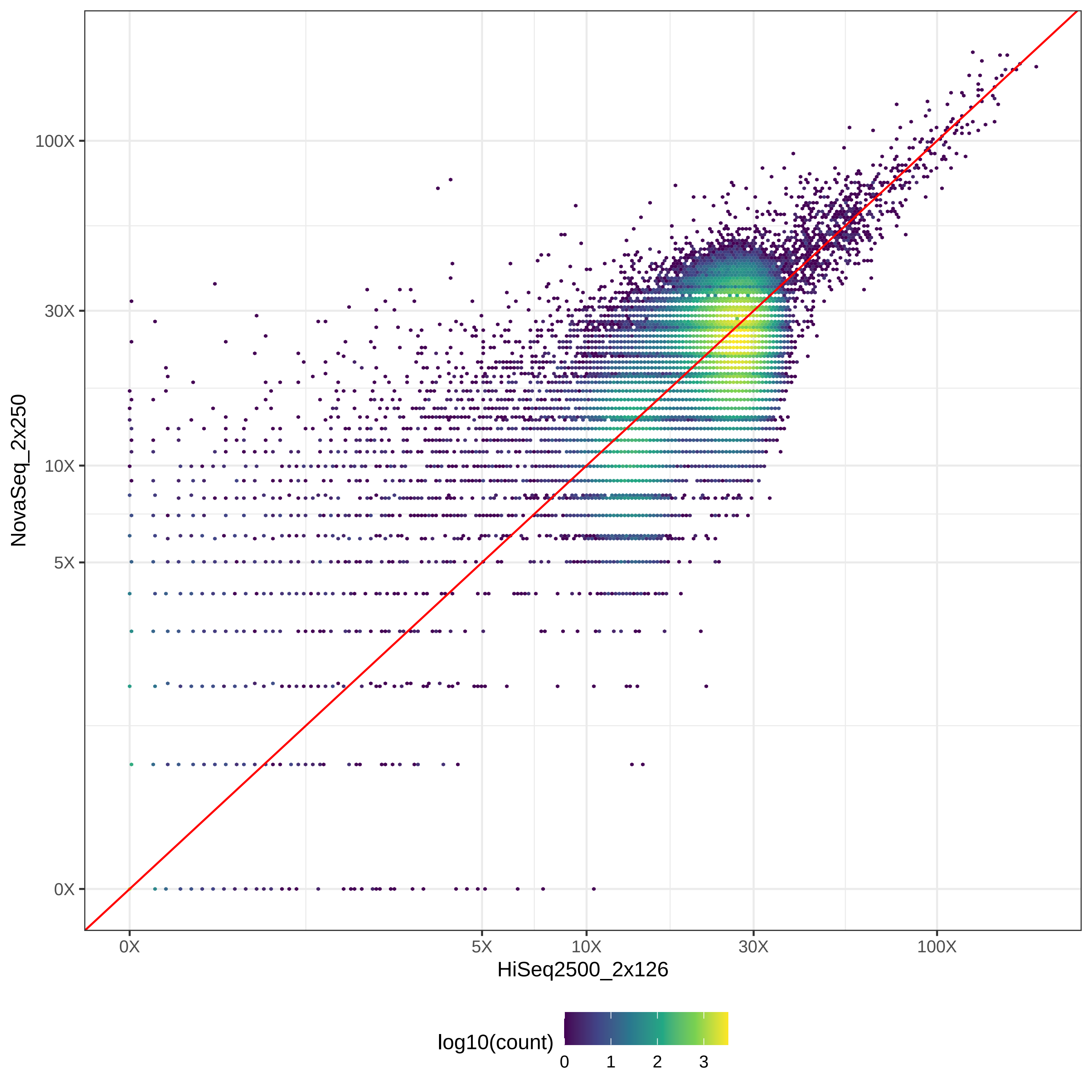

### GIAB_L2_HiSeq2500_2x126-v-PacBio_CCS-15k_hex.pdf

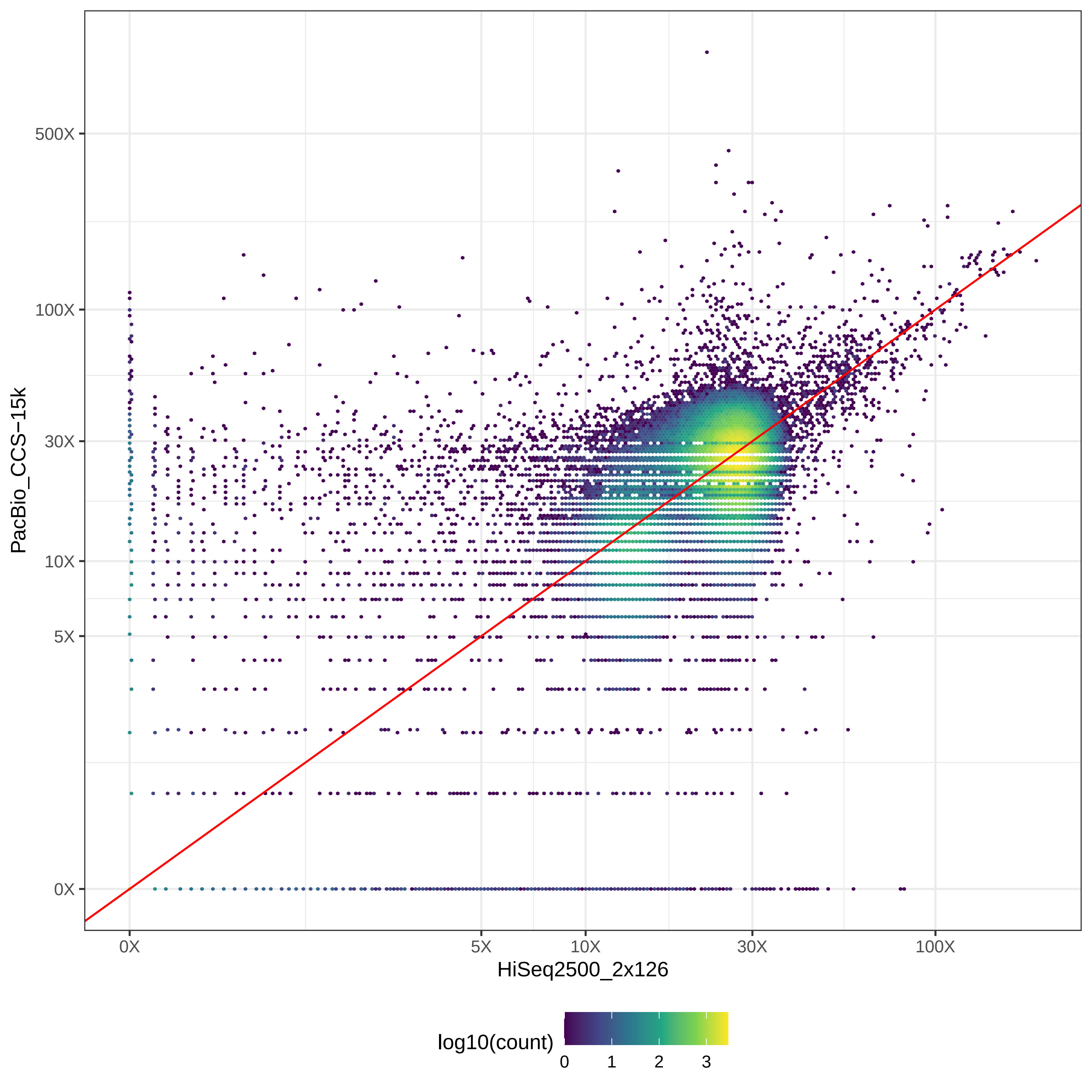

### GIAB_L2_HiSeq2500_2x126-v-PacBio_CLR_hex.pdf

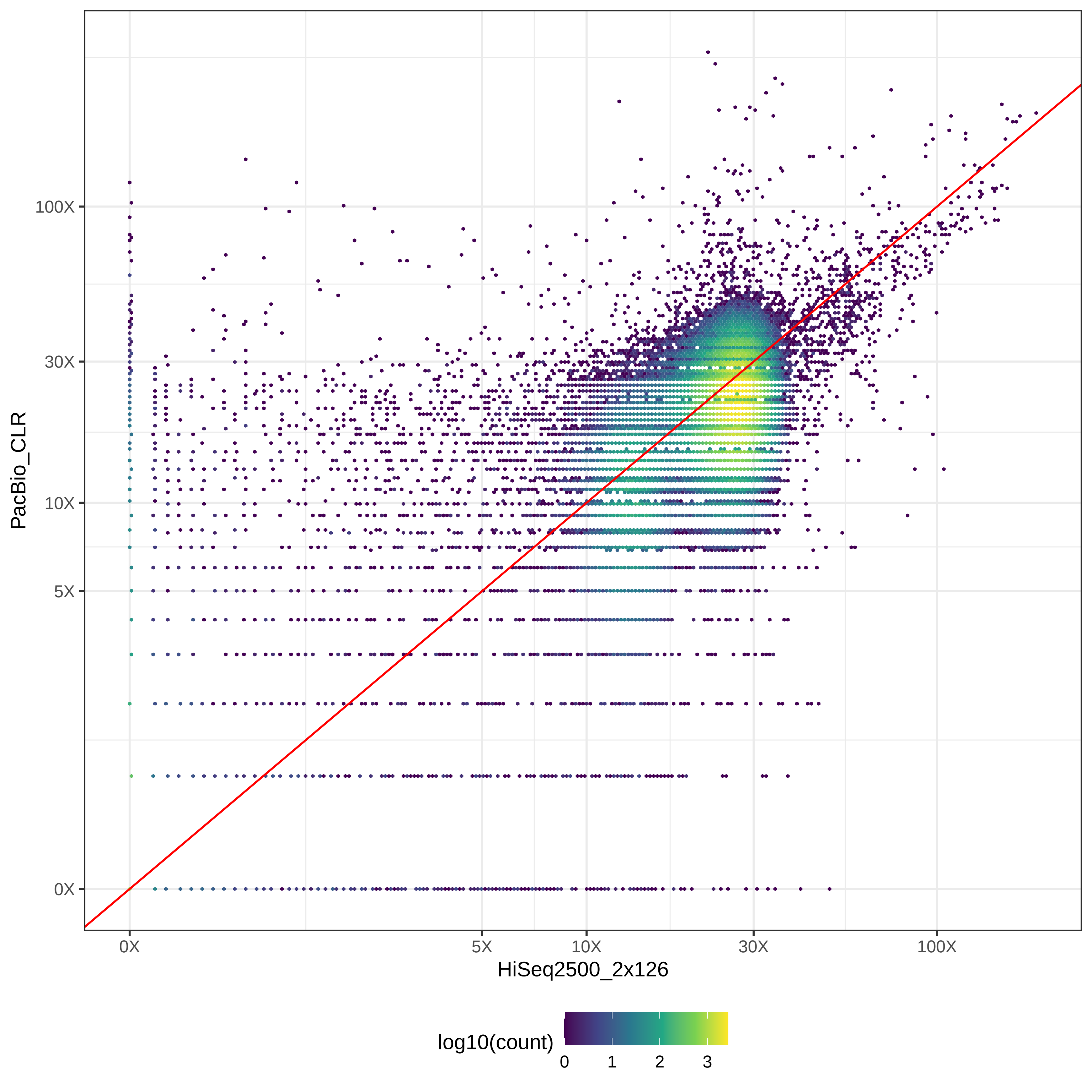

### GIAB_L2_HiSeq2500_2x126-v-PromethION_μ=8092_hex.pdf

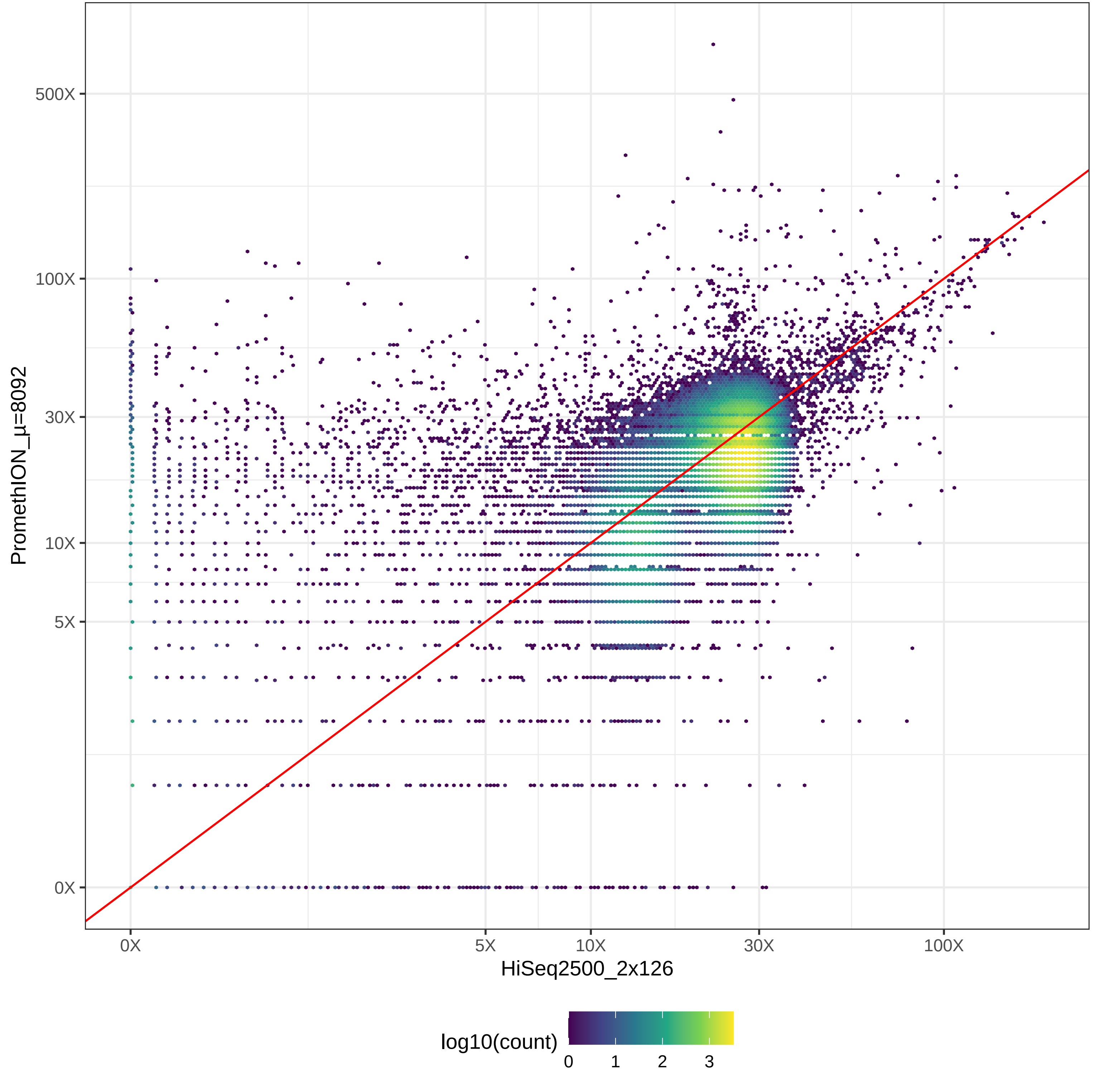

### GIAB_L2_HiSeq4000_2x150-v-HiSeqX10_2x150_hex.pdf

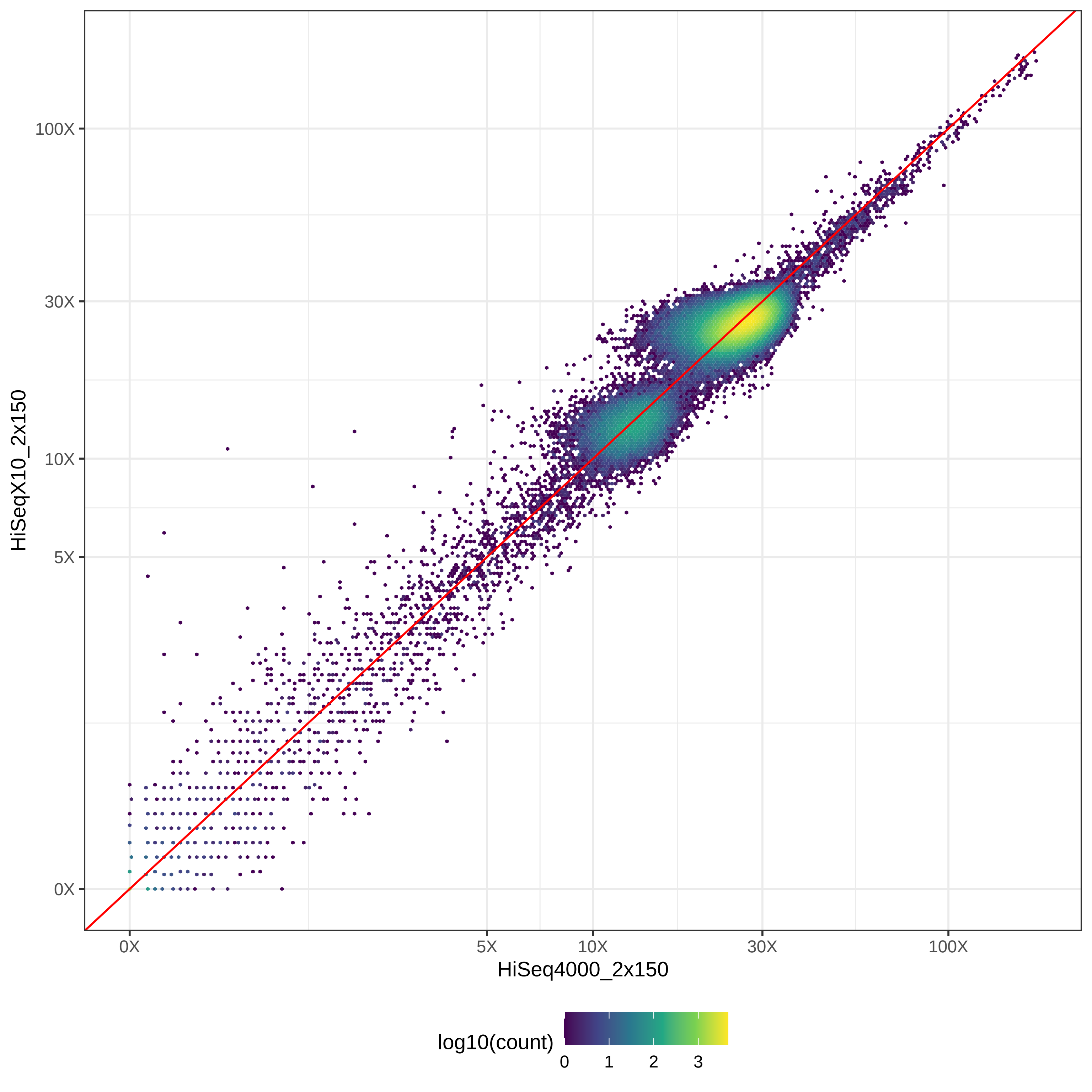

### GIAB_L2_HiSeq4000_2x150-v-MGISEQ2000_2x150_hex.pdf

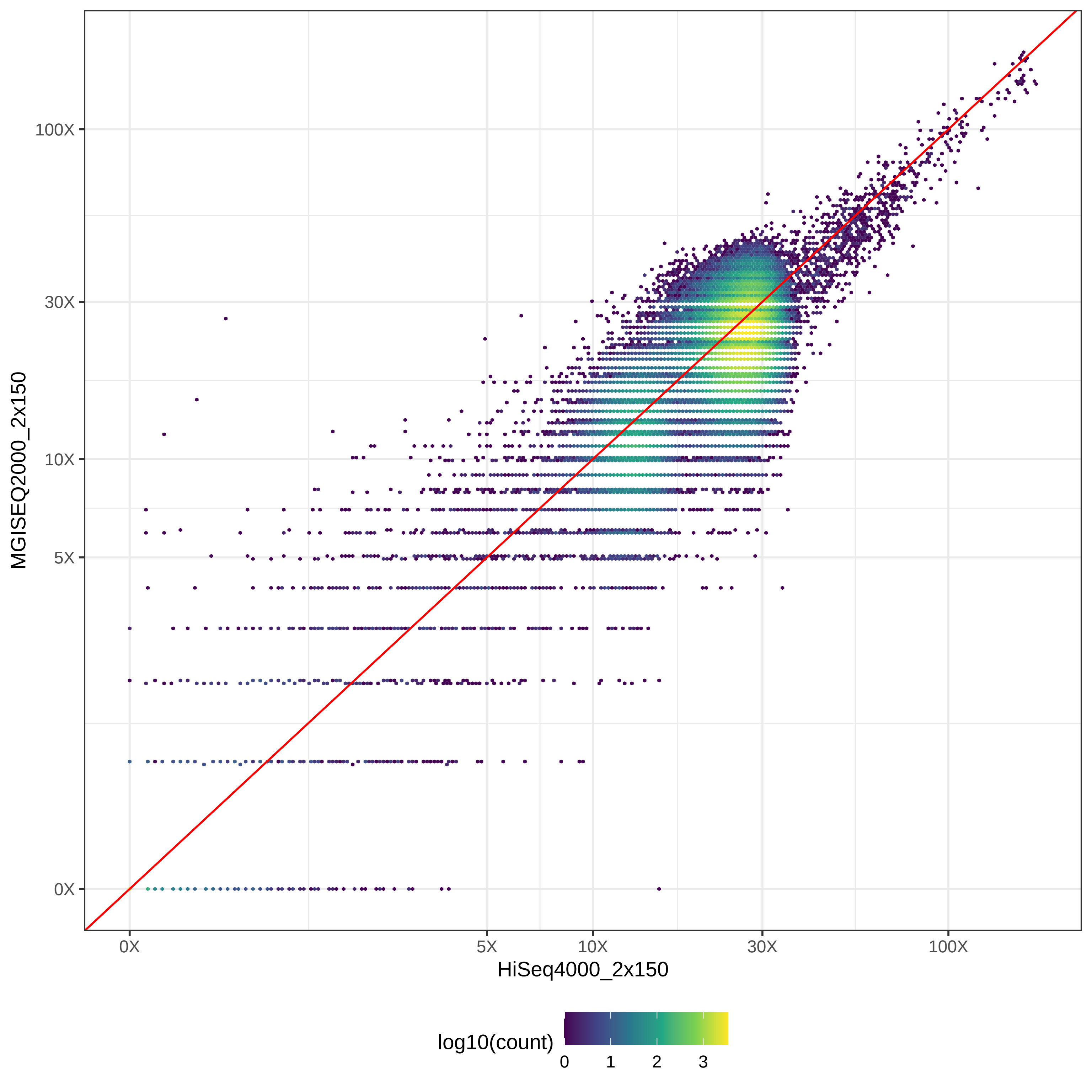

### GIAB_L2_HiSeq4000_2x150-v-NovaSeq_2x150_hex.pdf

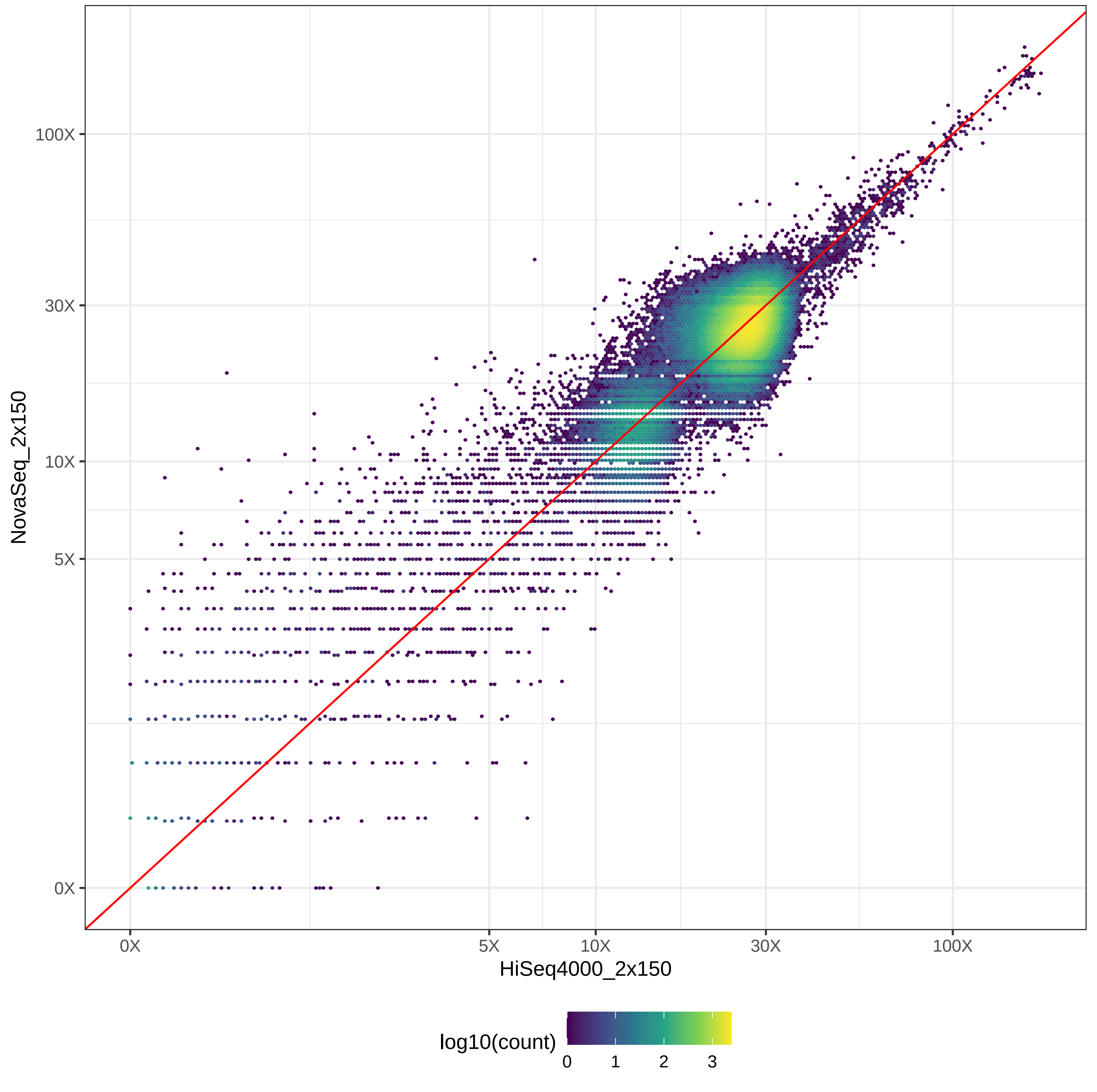

### GIAB_L2_HiSeq4000_2x150-v-NovaSeq_2x250_hex.pdf

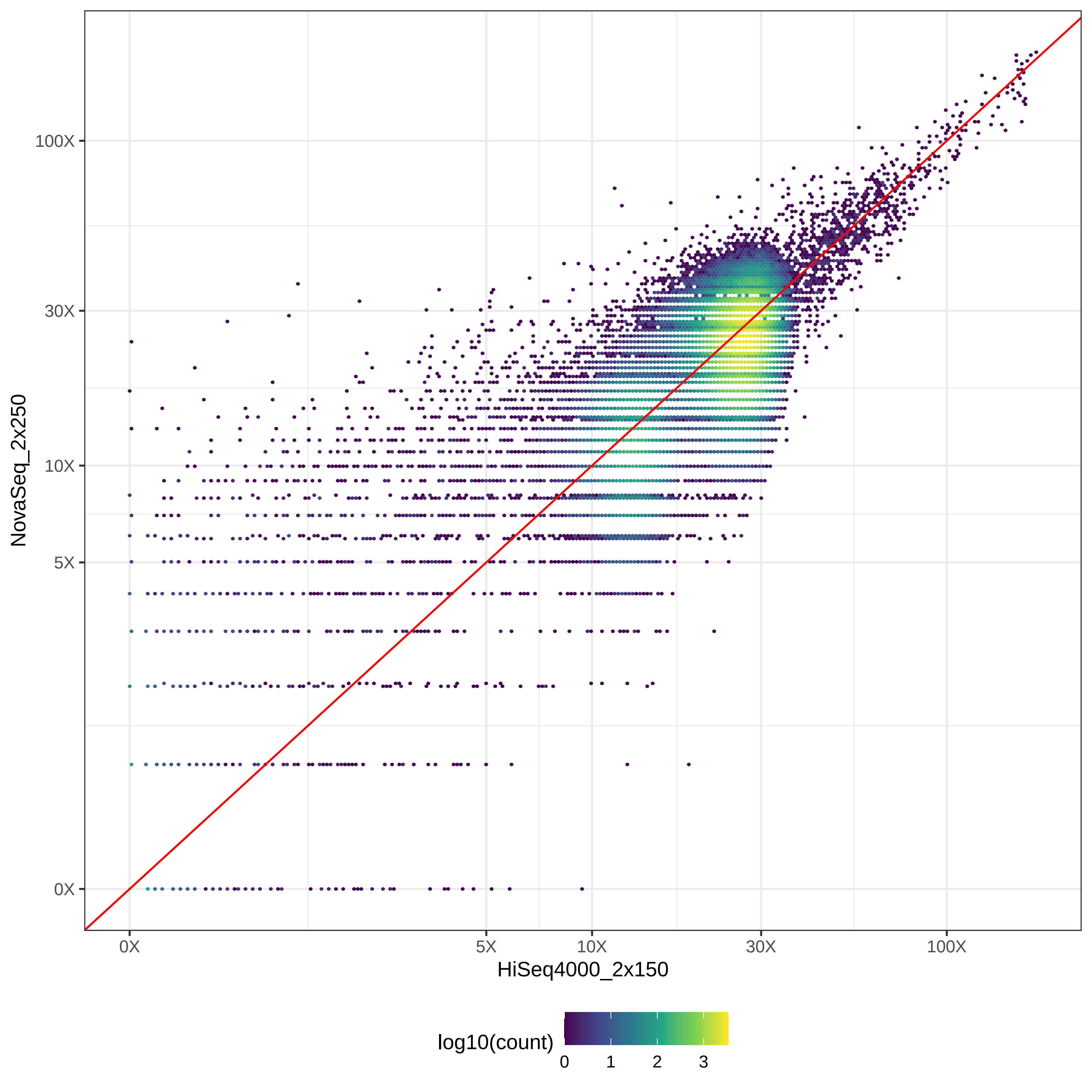

### GIAB_L2_HiSeq4000_2x150-v-PacBio_CCS-15k_hex.pdf

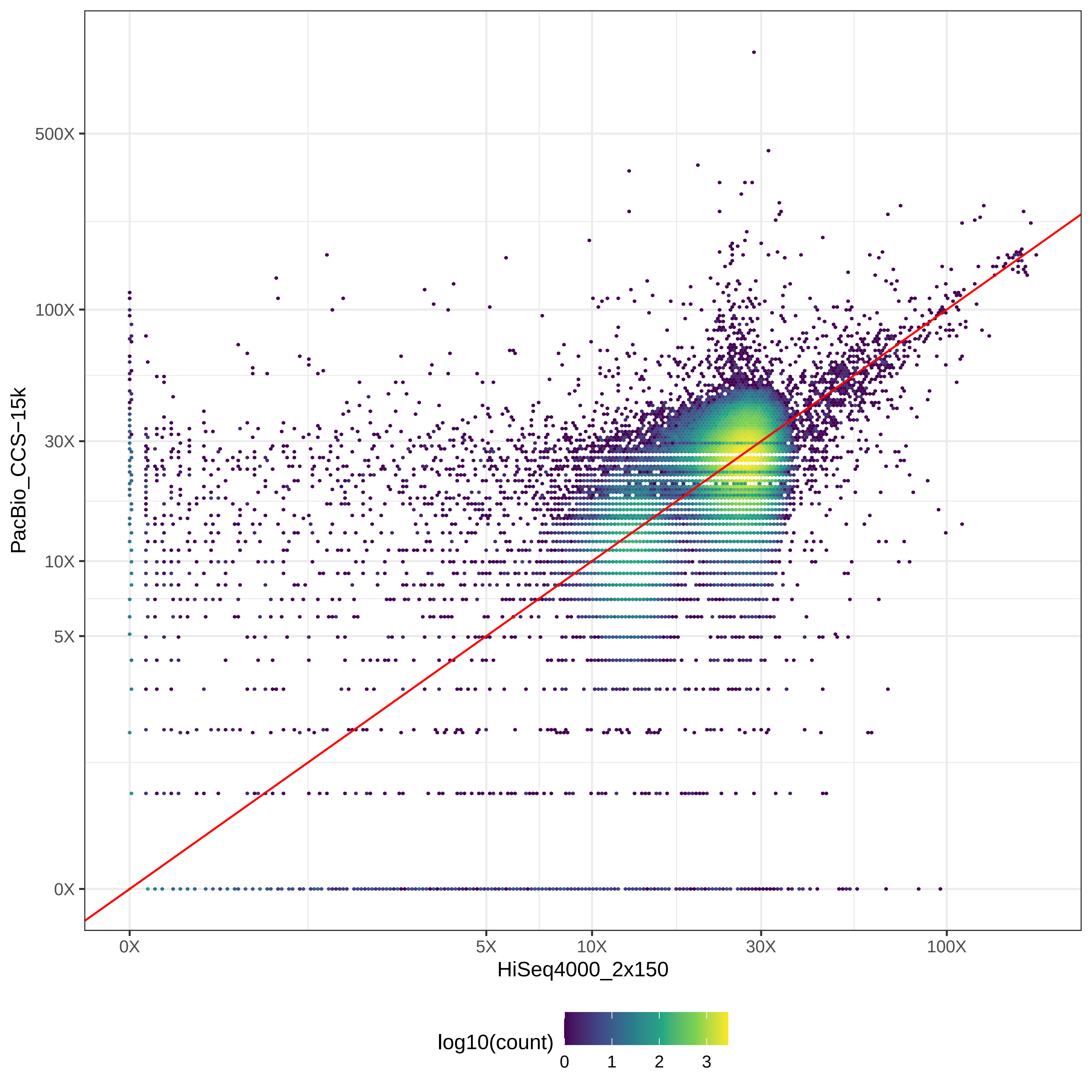

### GIAB_L2_HiSeq4000_2x150-v-PacBio_CLR_hex.pdf

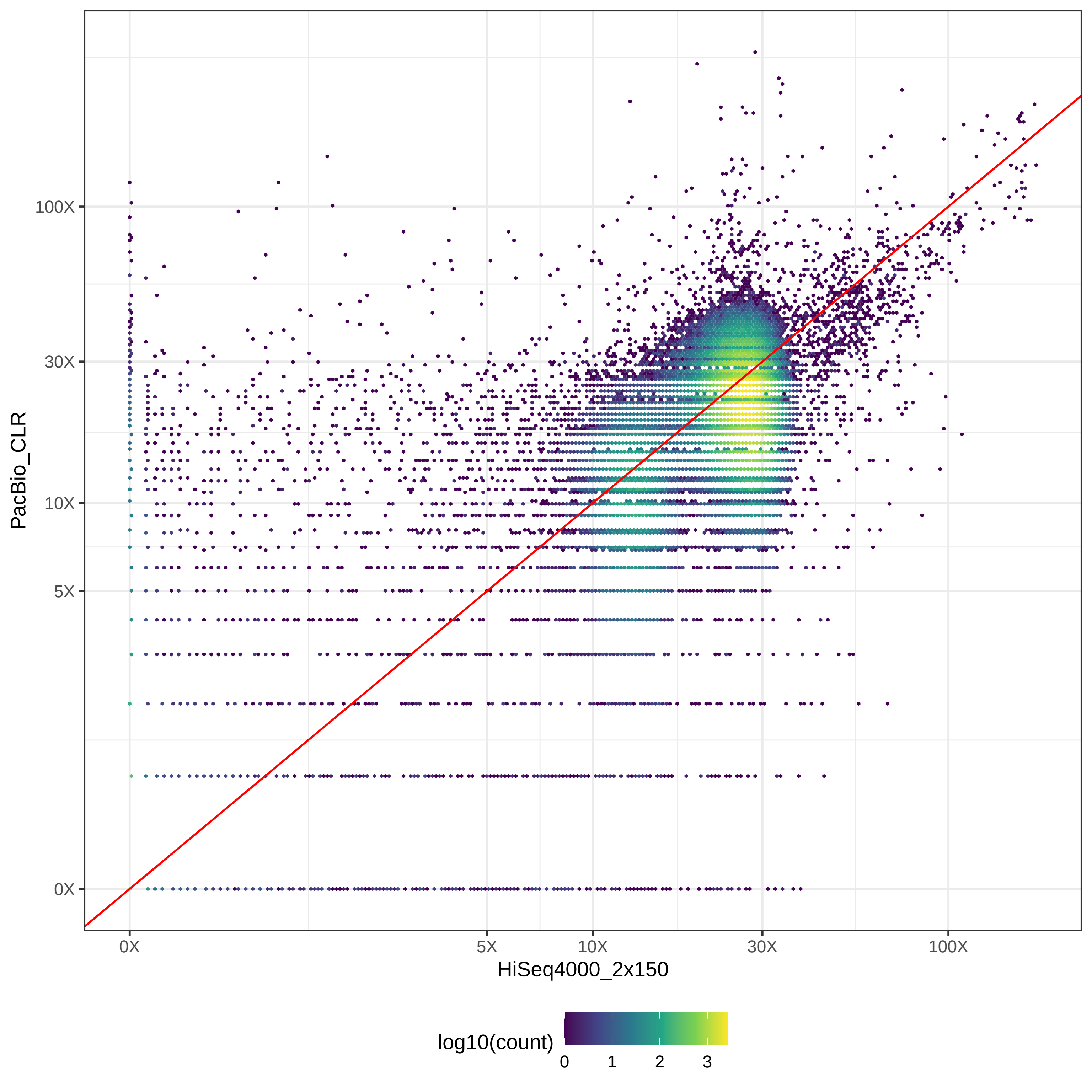

### GIAB_L2_HiSeq4000_2x150-v-PromethION_μ=8092_hex.pdf

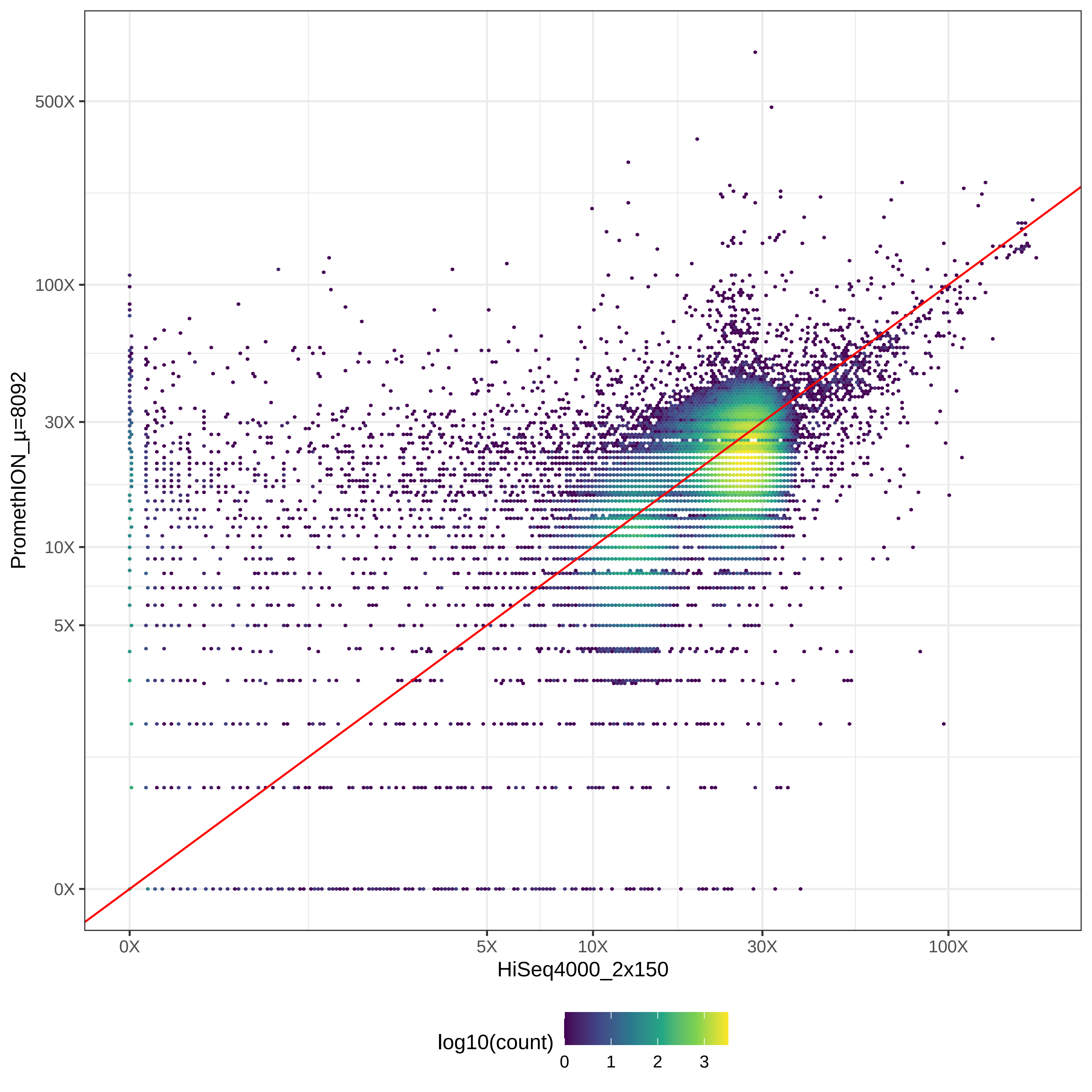

### GIAB_L2_HiSeqX10_2x150-v-MGISEQ2000_2x150_hex.pdf

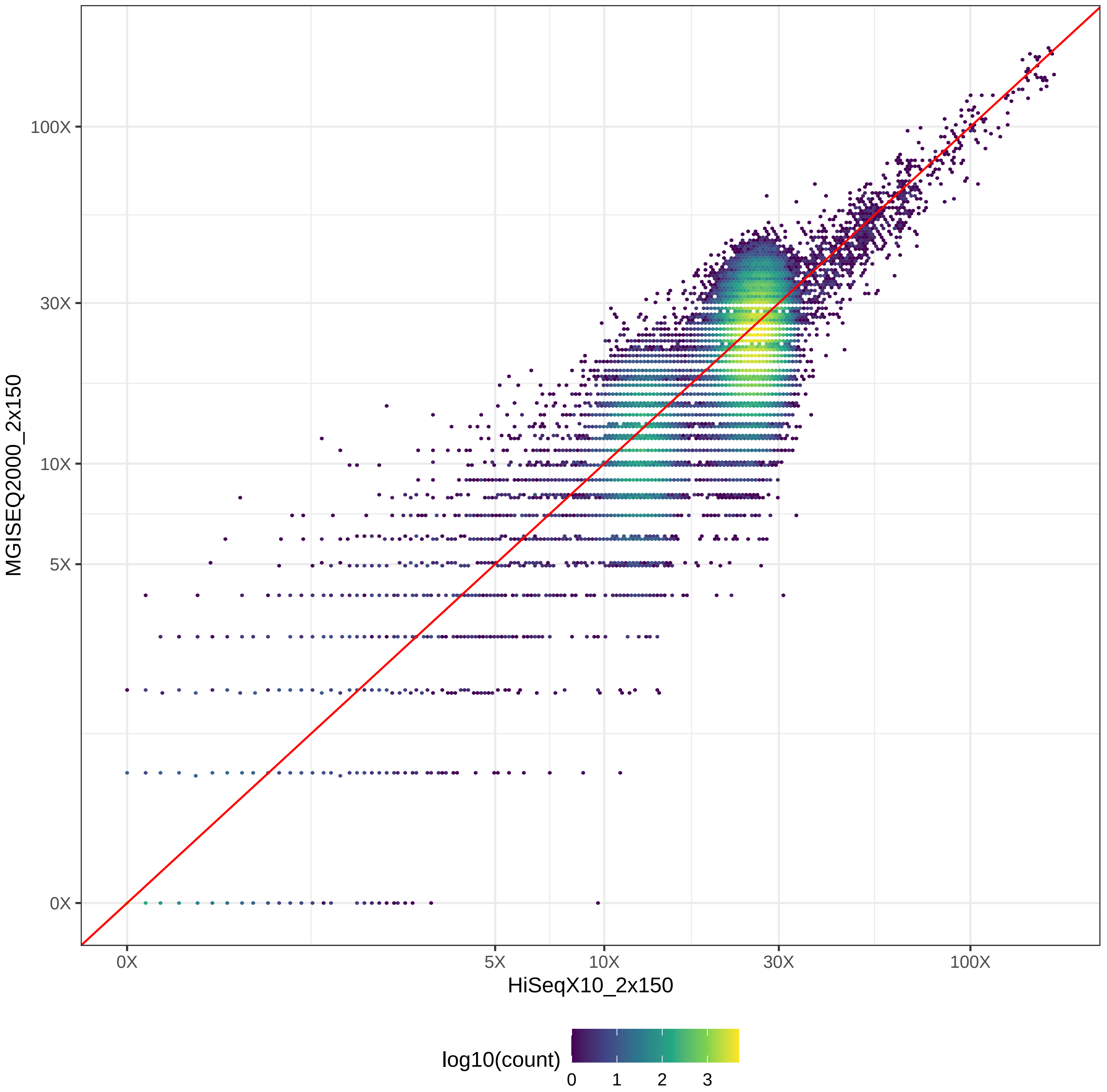

### GIAB_L2_HiSeqX10_2x150-v-NovaSeq_2x150_hex.pdf

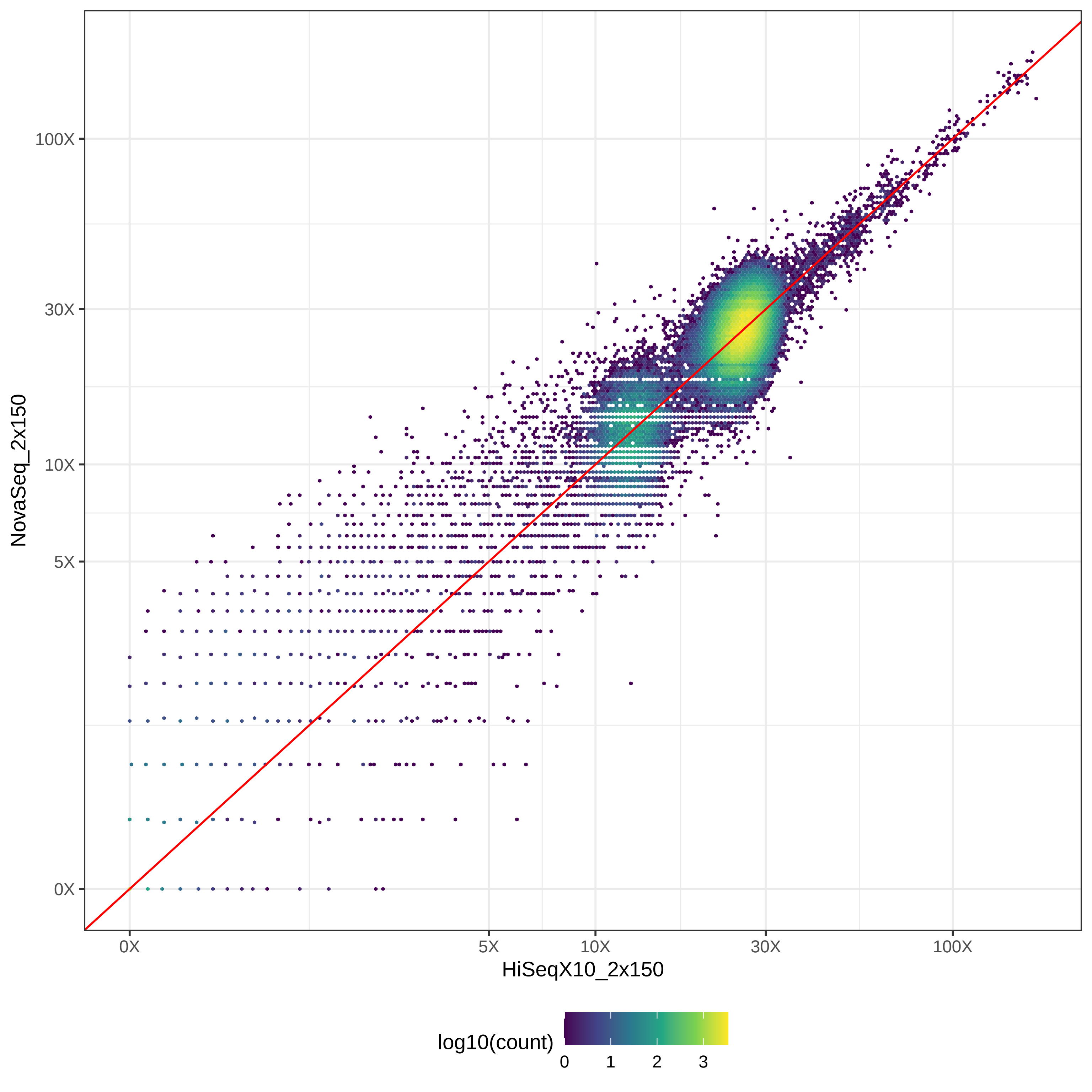

### GIAB_L2_HiSeqX10_2x150-v-NovaSeq_2x250_hex.pdf

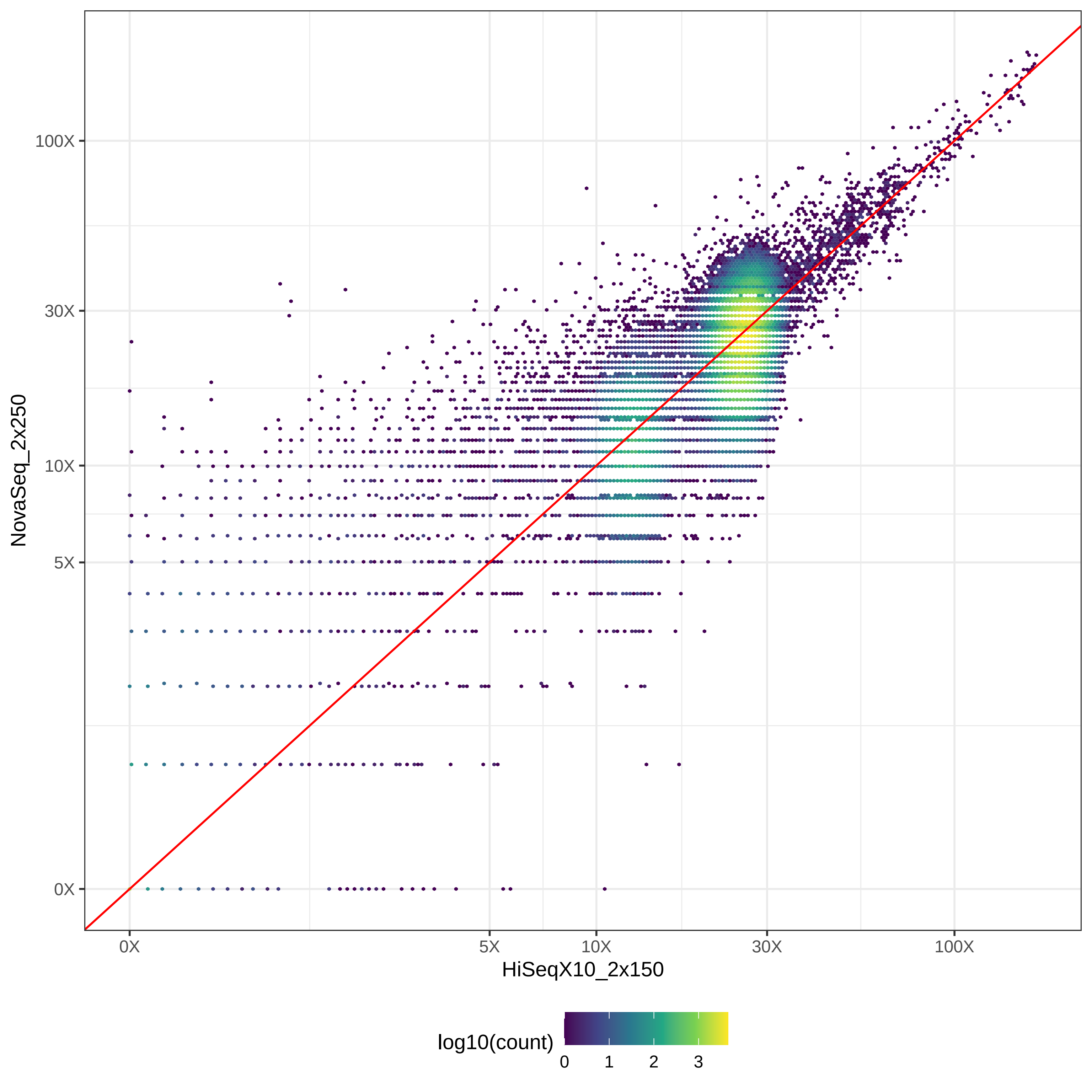

### GIAB_L2_HiSeqX10_2x150-v-PacBio_CCS-15k_hex.pdf

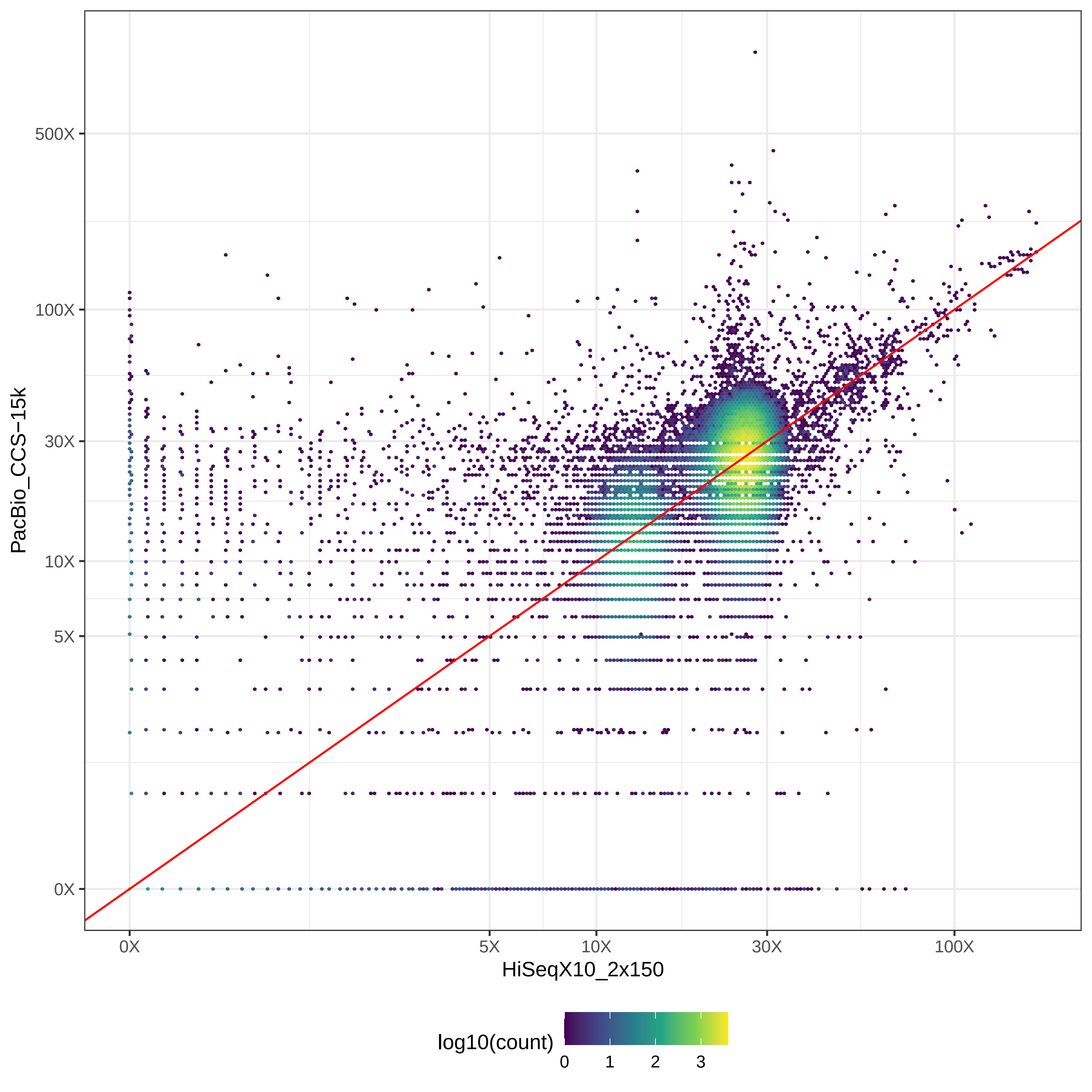

### GIAB_L2_HiSeqX10_2x150-v-PacBio_CLR_hex.pdf

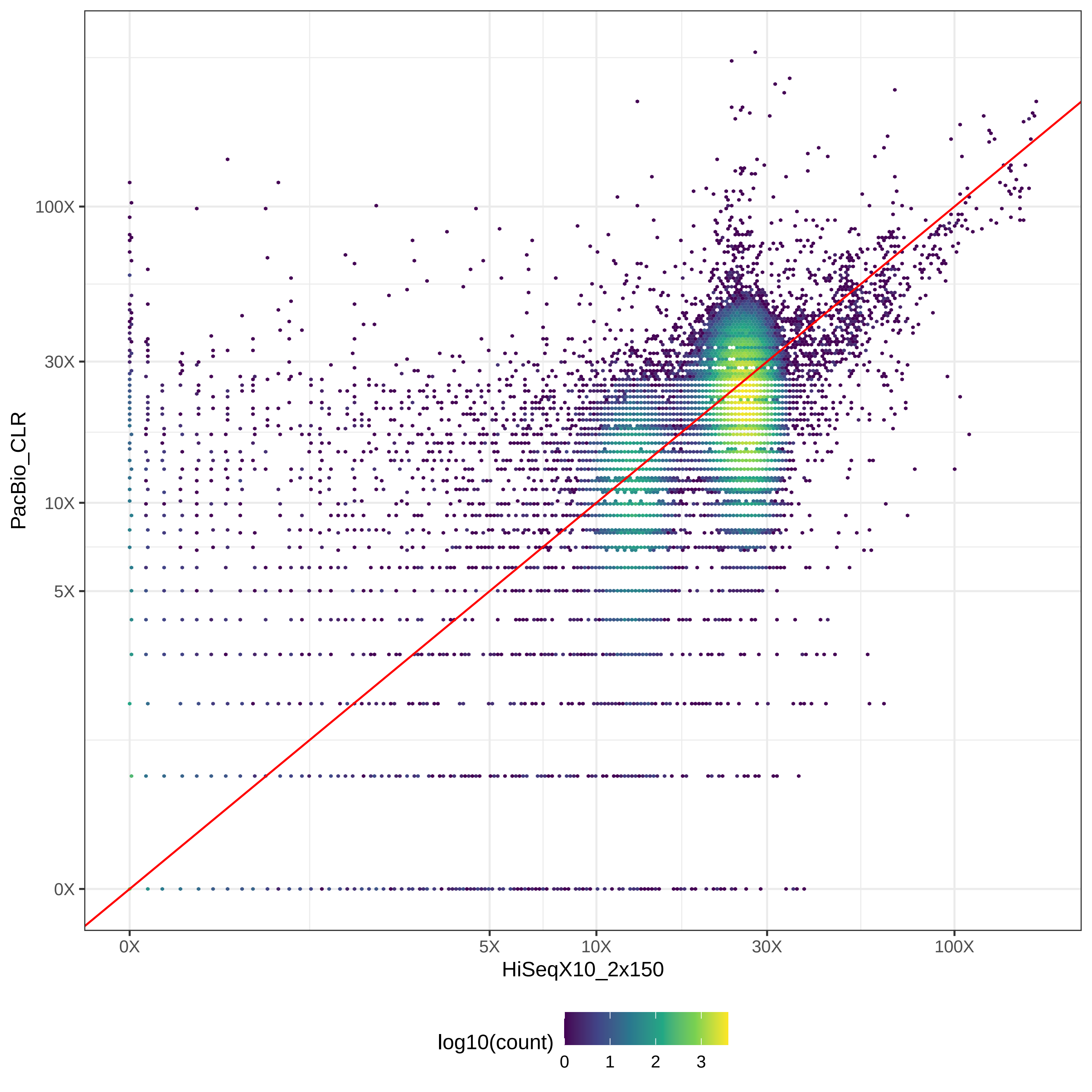

### GIAB_L2_HiSeqX10_2x150-v-PromethION_μ=8092_hex.pdf

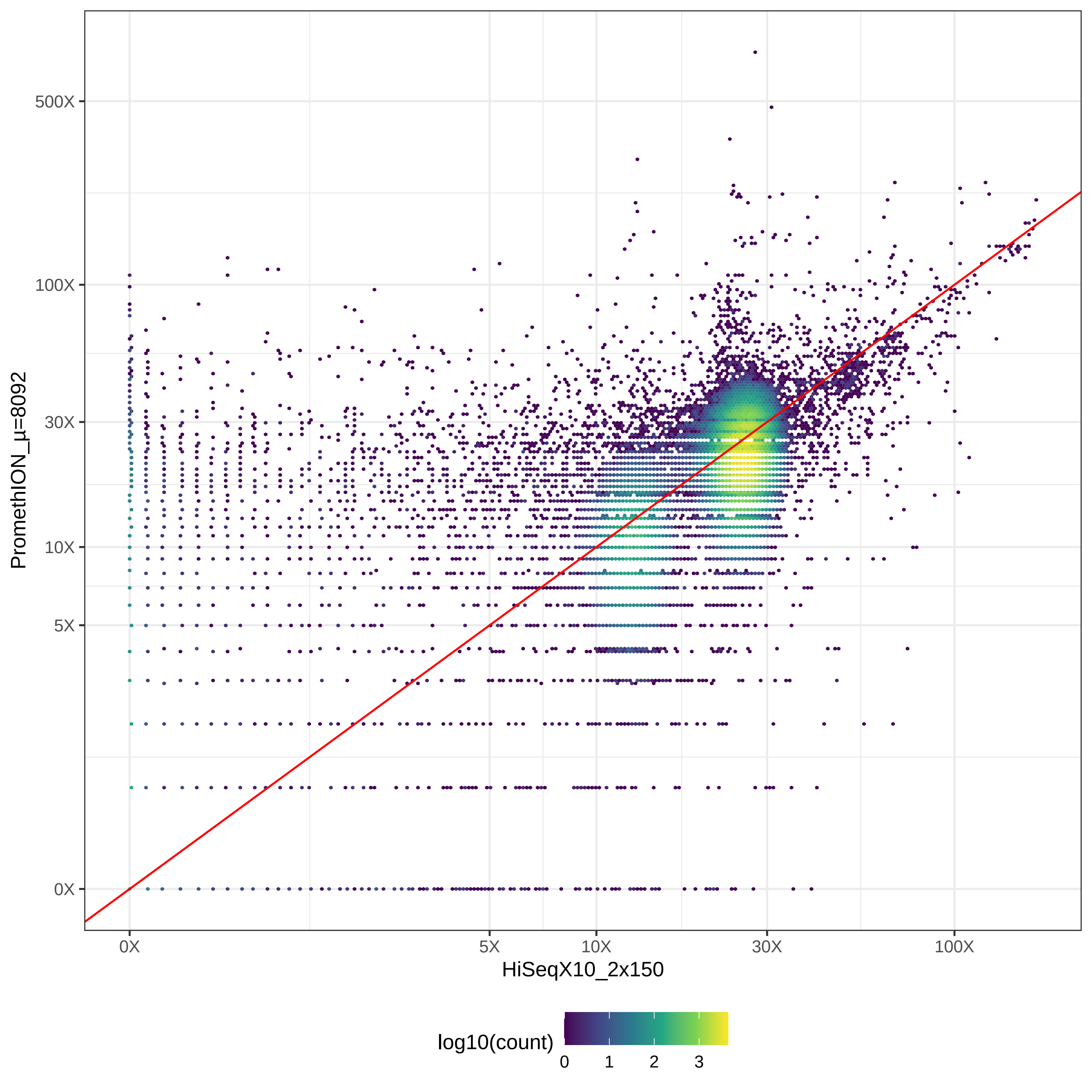

### GIAB_L2_MGISEQ2000_2x150-v-NovaSeq_2x150_hex.pdf

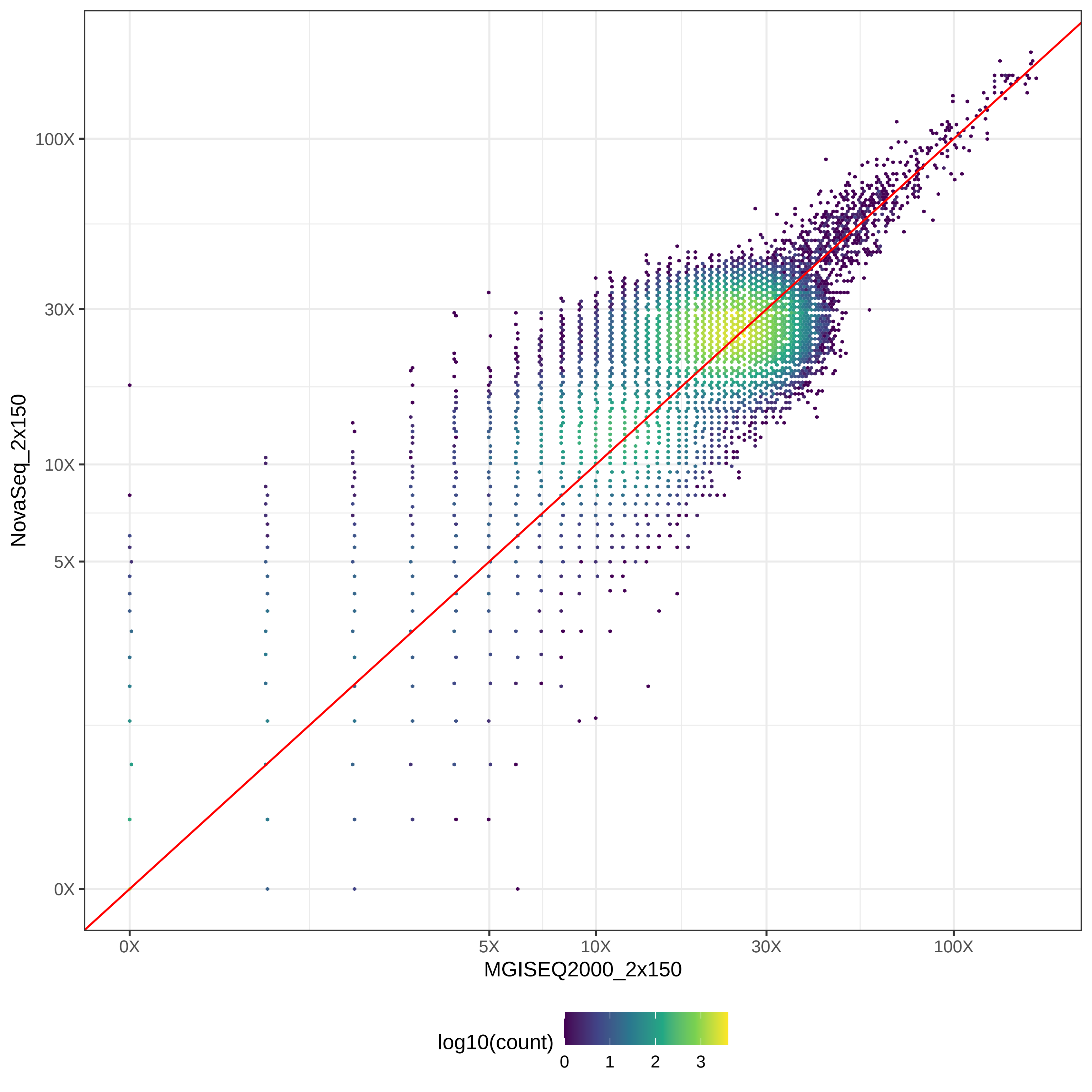

### GIAB_L2_MGISEQ2000_2x150-v-NovaSeq_2x250_hex.pdf

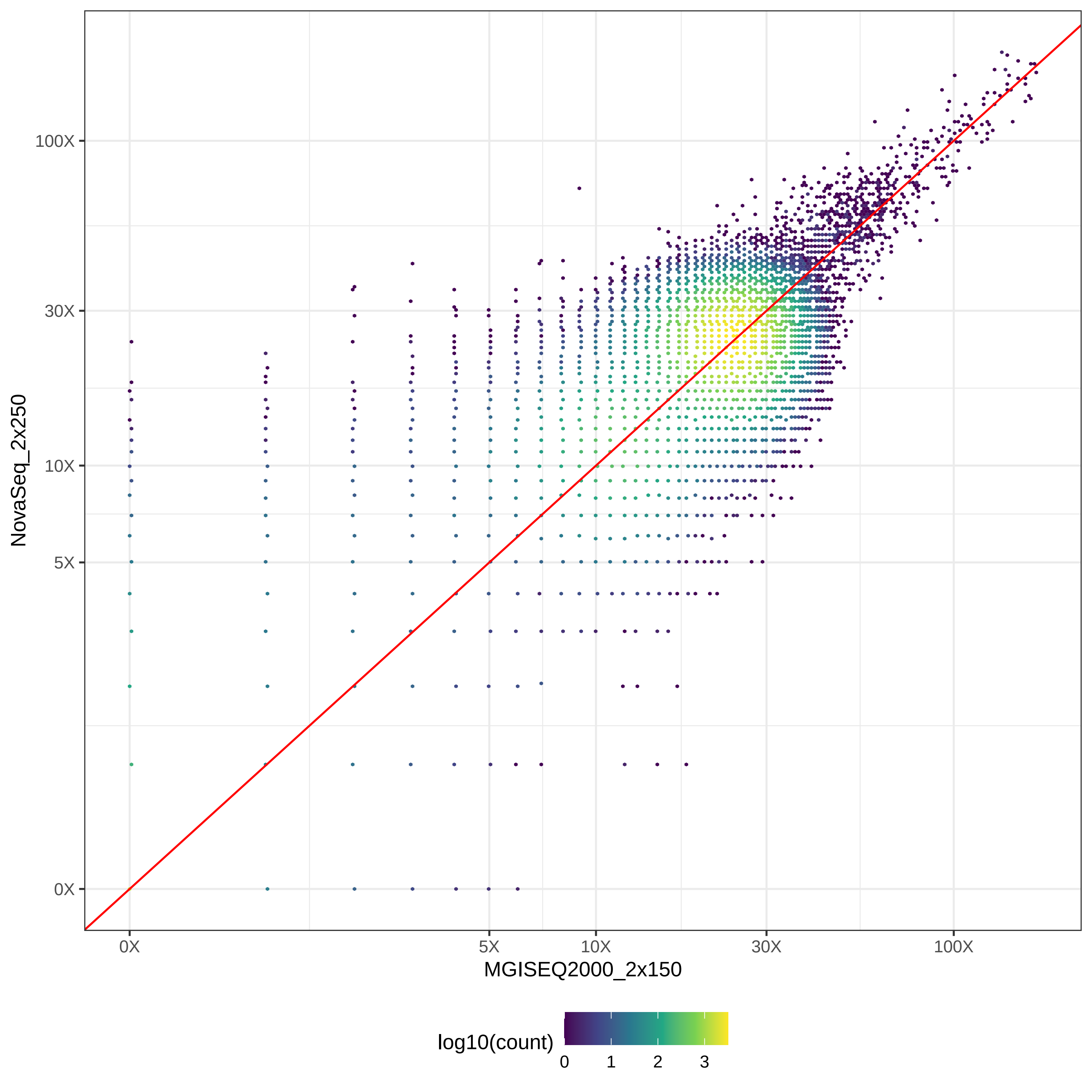

### GIAB_L2_MGISEQ2000_2x150-v-PacBio_CCS-15k_hex.pdf

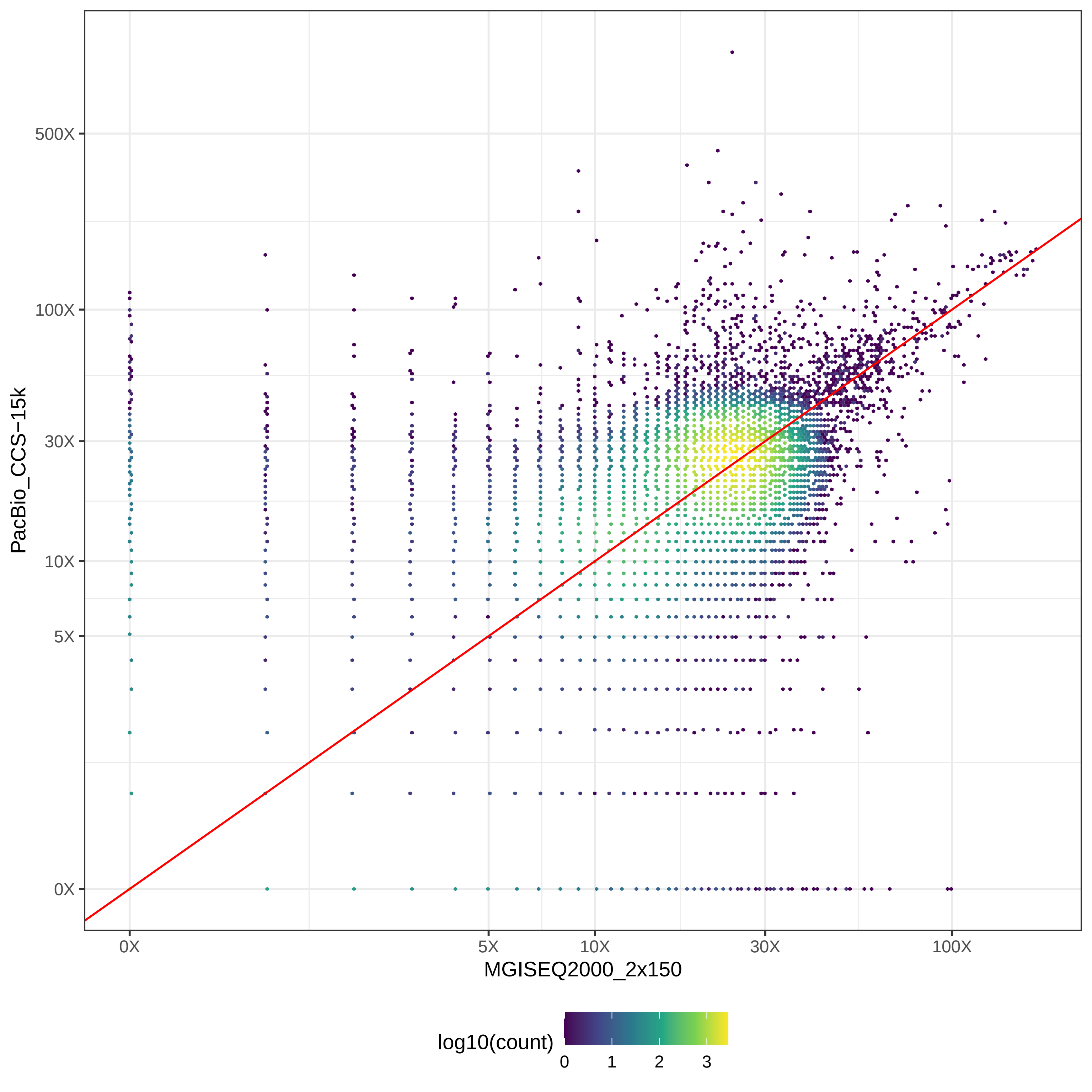

### GIAB_L2_MGISEQ2000_2x150-v-PacBio_CLR_hex.pdf

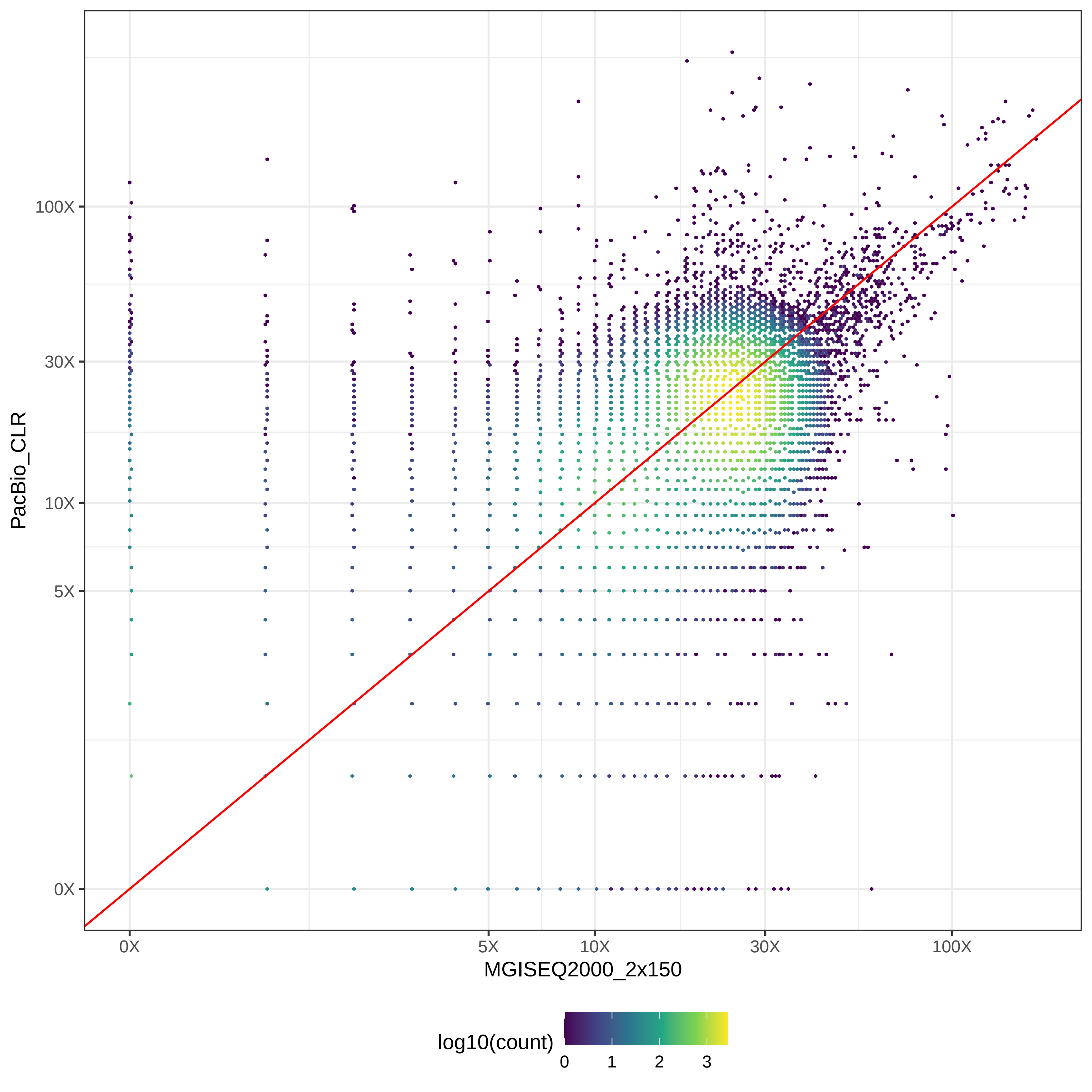

### GIAB_L2_MGISEQ2000_2x150-v-PromethION_μ=8092_hex.pdf

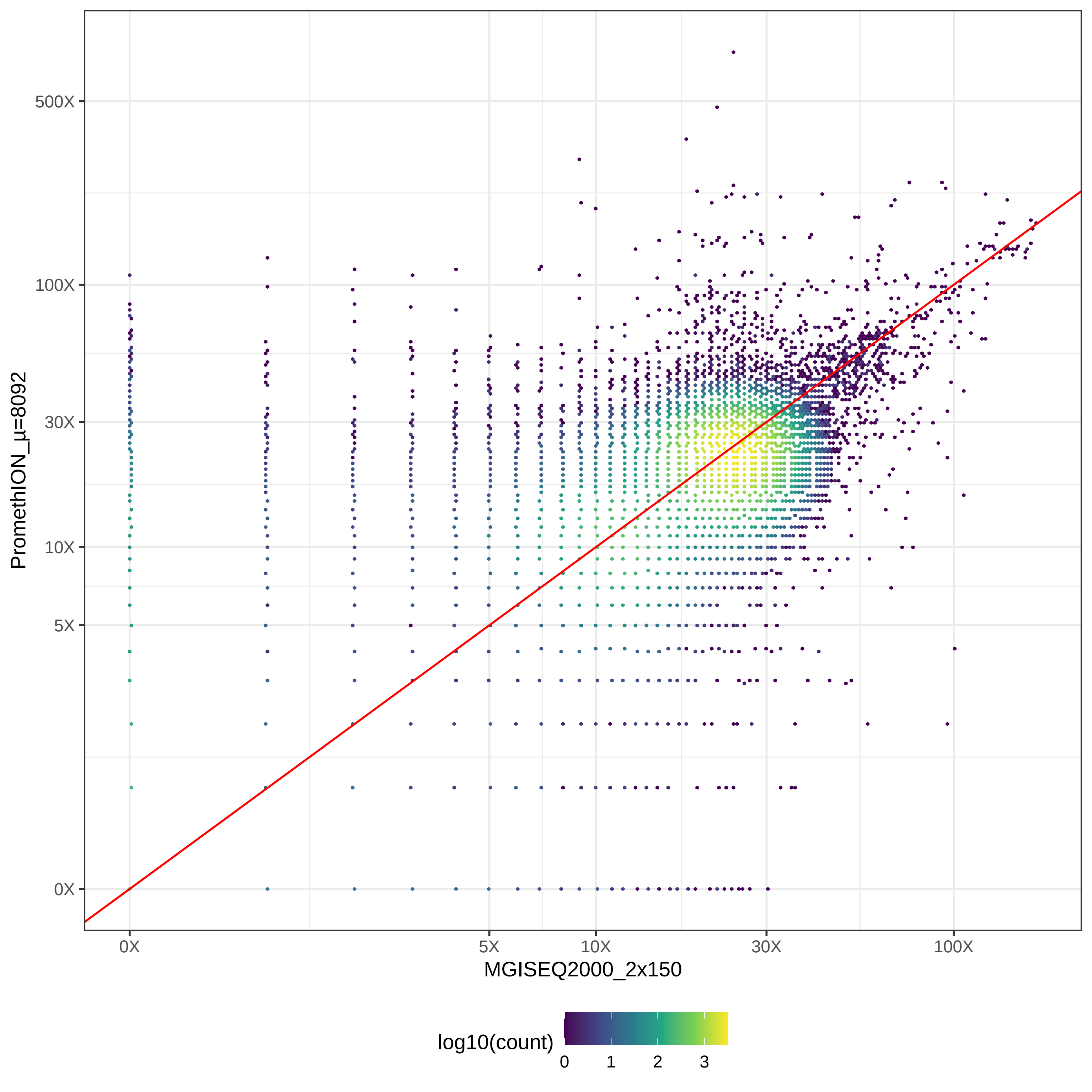

### GIAB_L2_NovaSeq_2x150-v-NovaSeq_2x250_hex.pdf

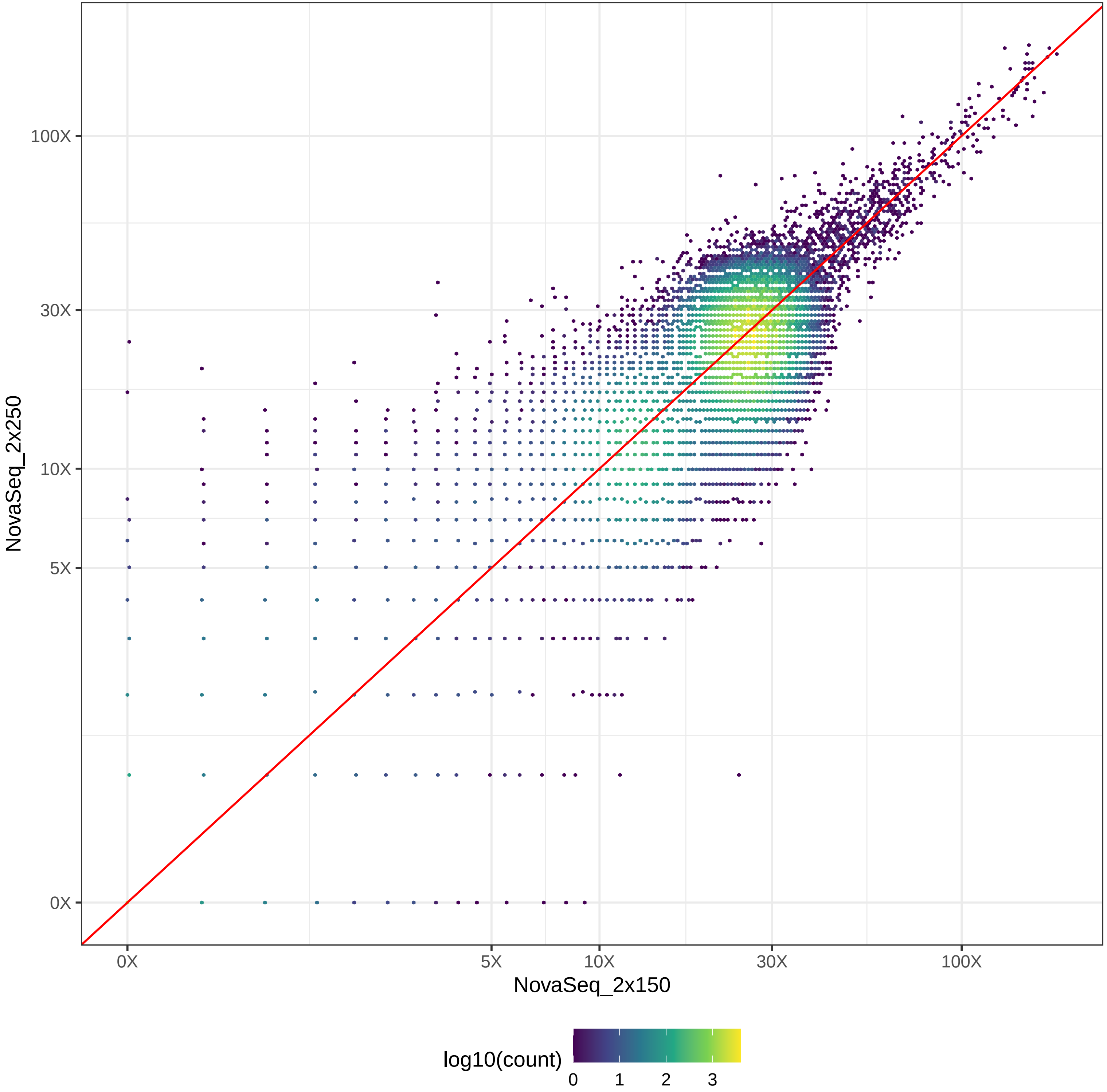

### GIAB_L2_NovaSeq_2x150-v-PacBio_CCS-15k_hex.pdf

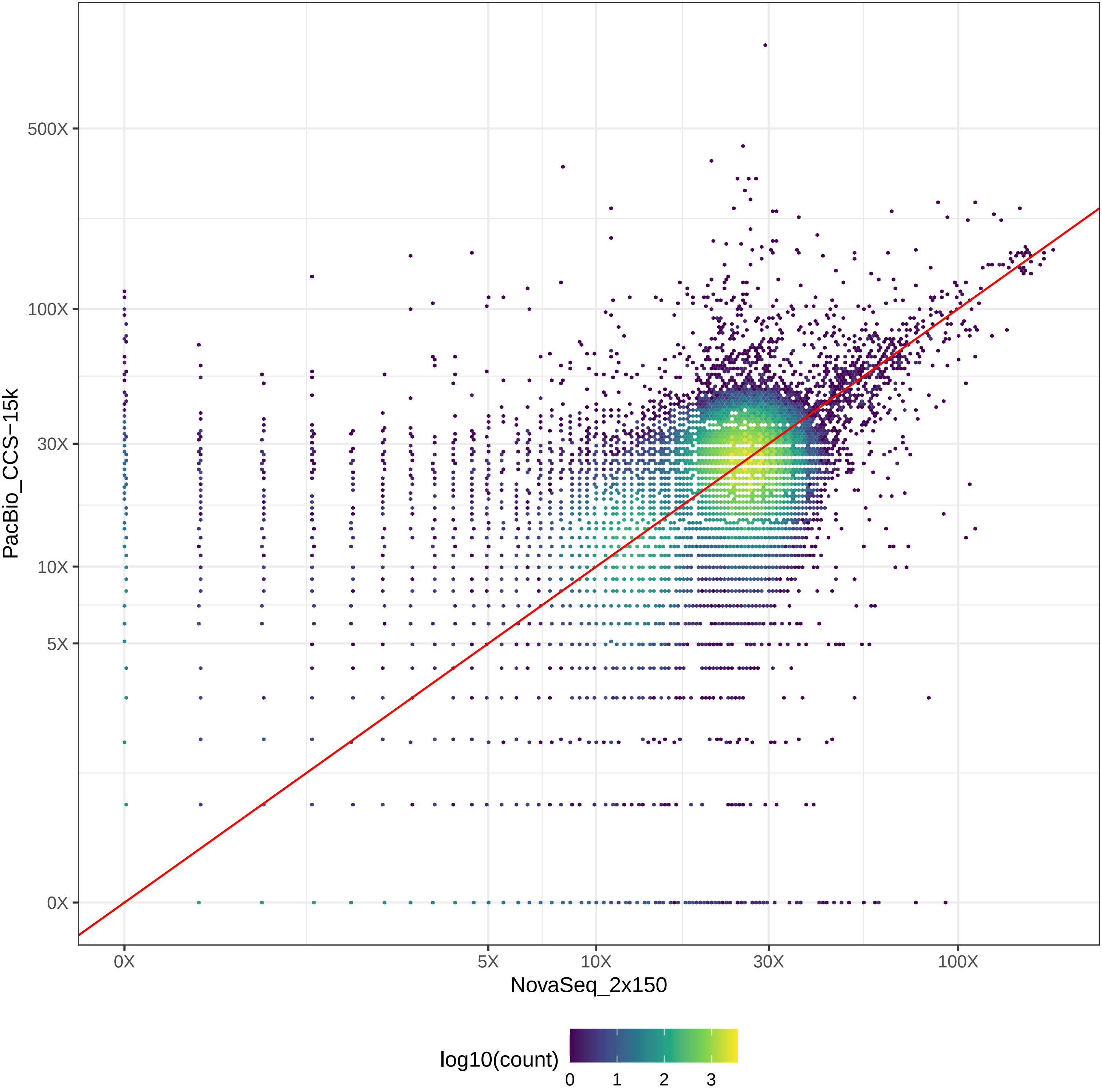

### GIAB_L2_NovaSeq_2x150-v-PacBio_CLR_hex.pdf

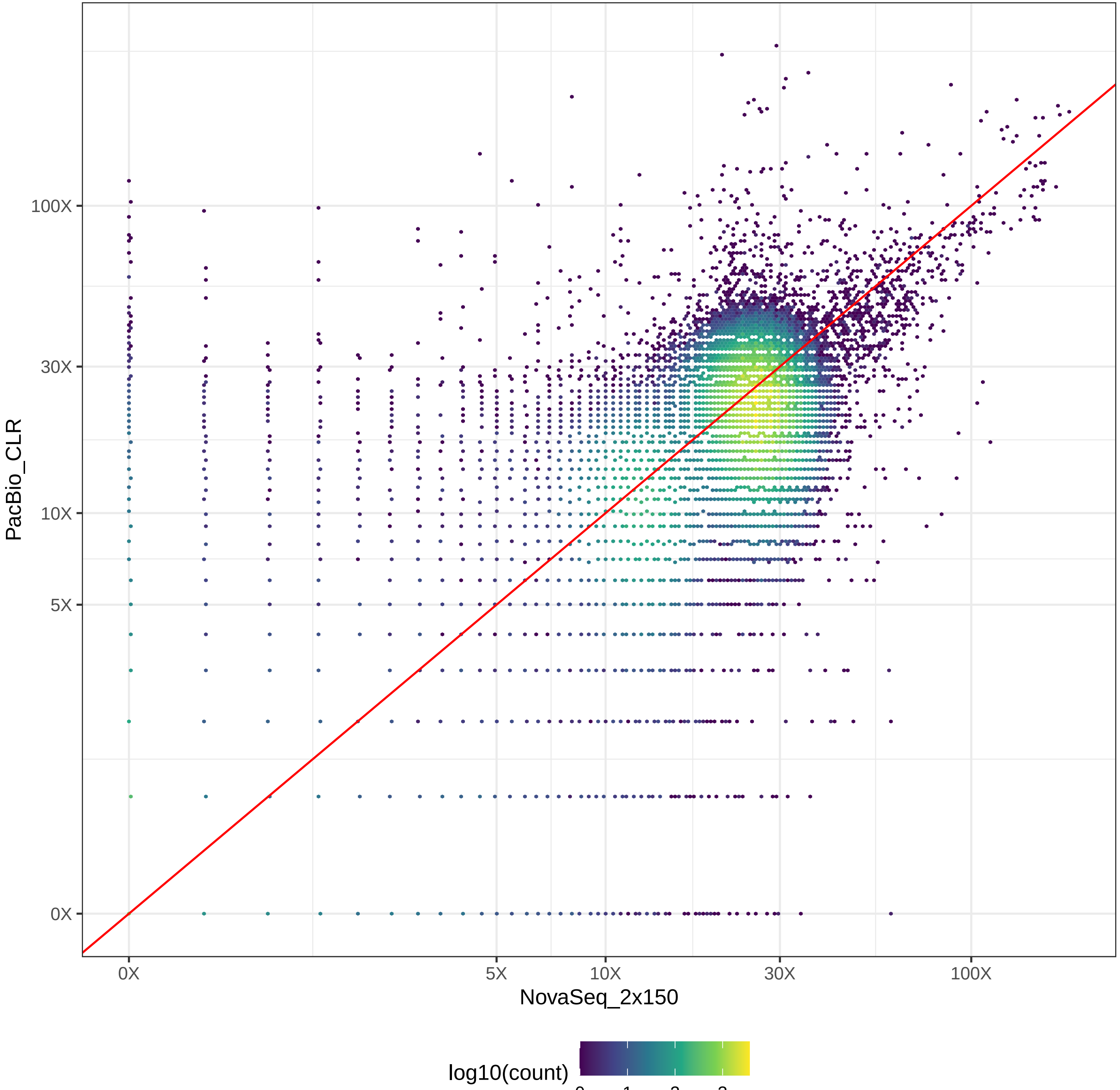

### GIAB_L2_NovaSeq_2x150-v-PromethION_μ=8092_hex.pdf

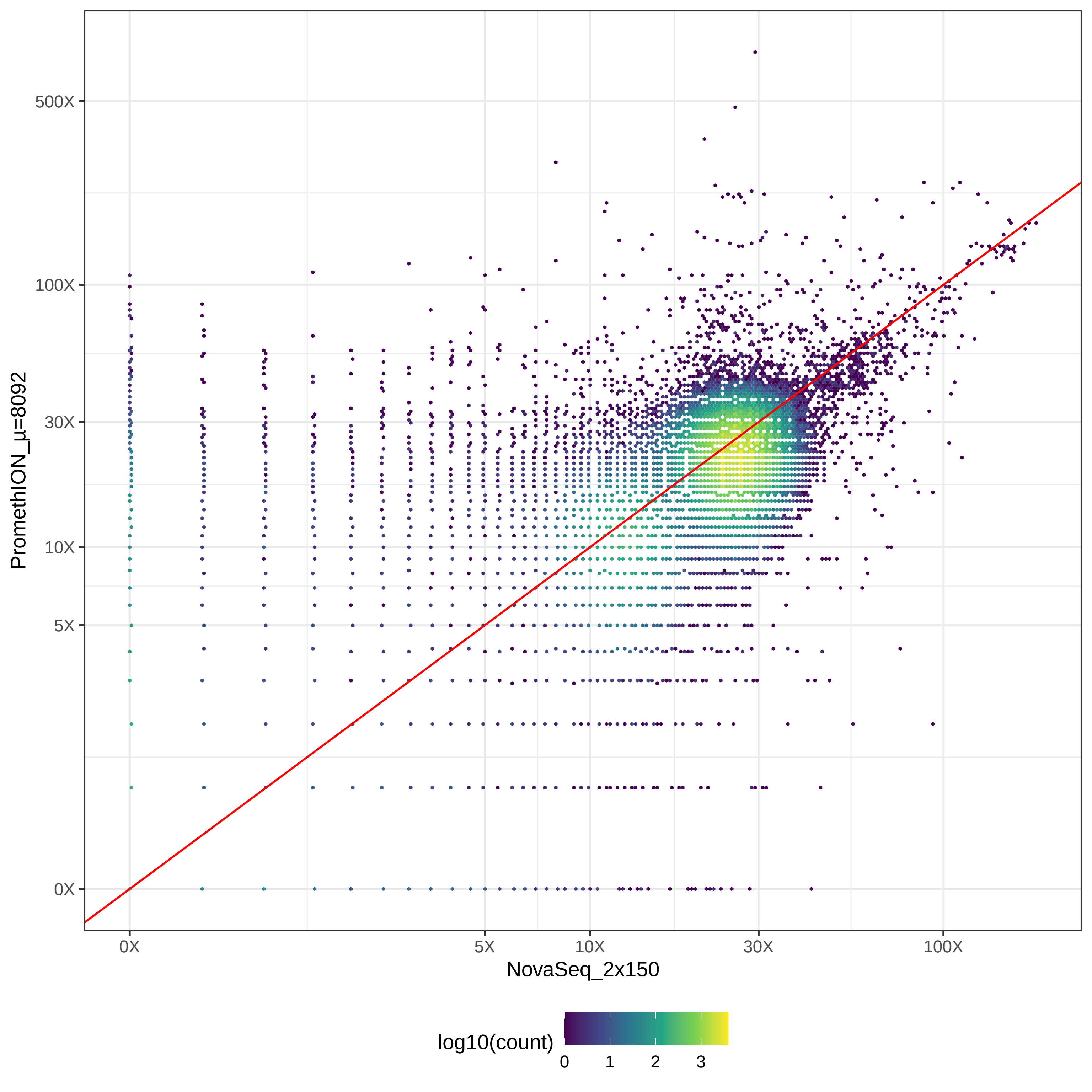

### GIAB_L2_NovaSeq_2x250-v-PacBio_CCS-15k_hex.pdf

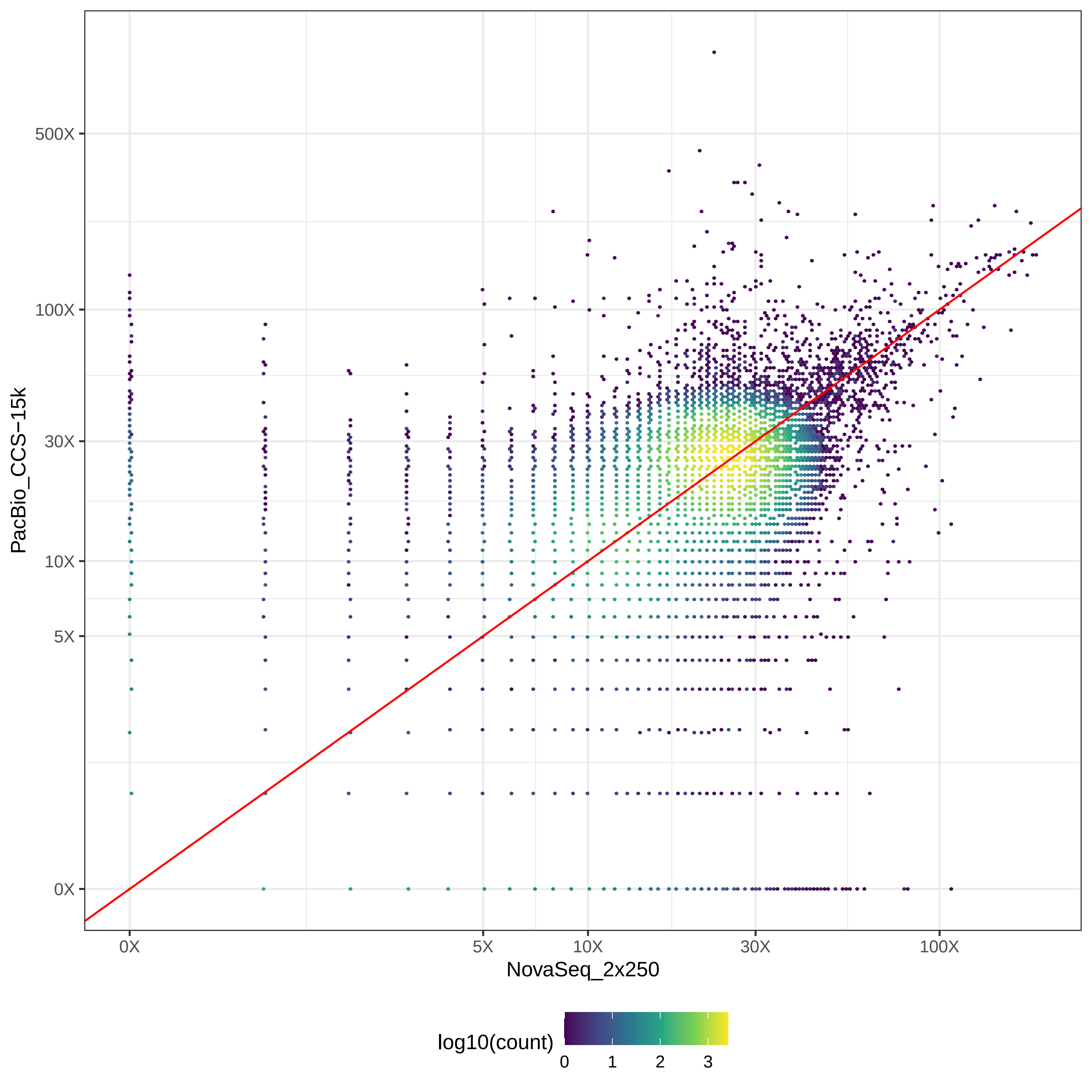

### GIAB_L2_NovaSeq_2x250-v-PacBio_CLR_hex.pdf

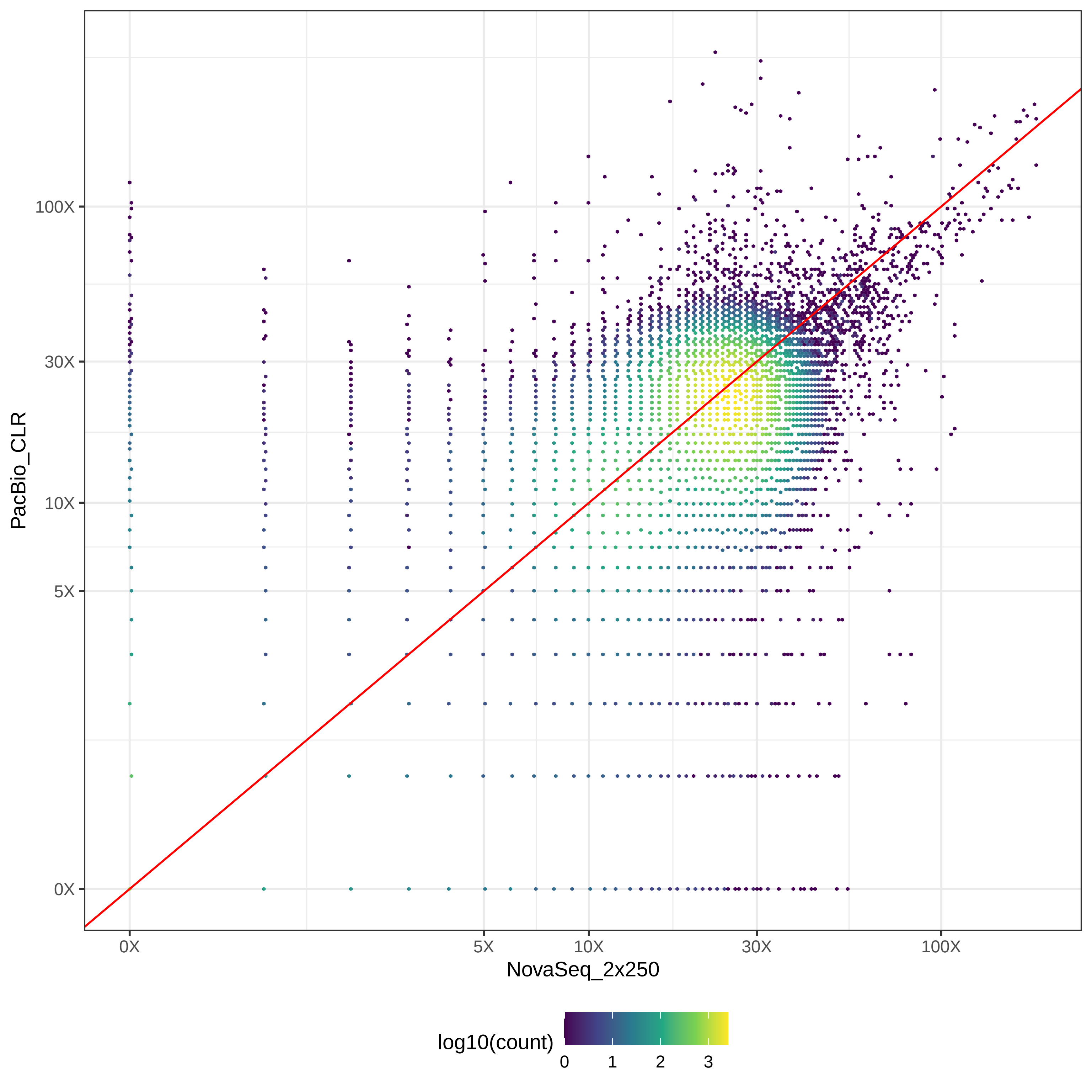

### GIAB_L2_NovaSeq_2x250-v-PromethION_μ=8092_hex.pdf

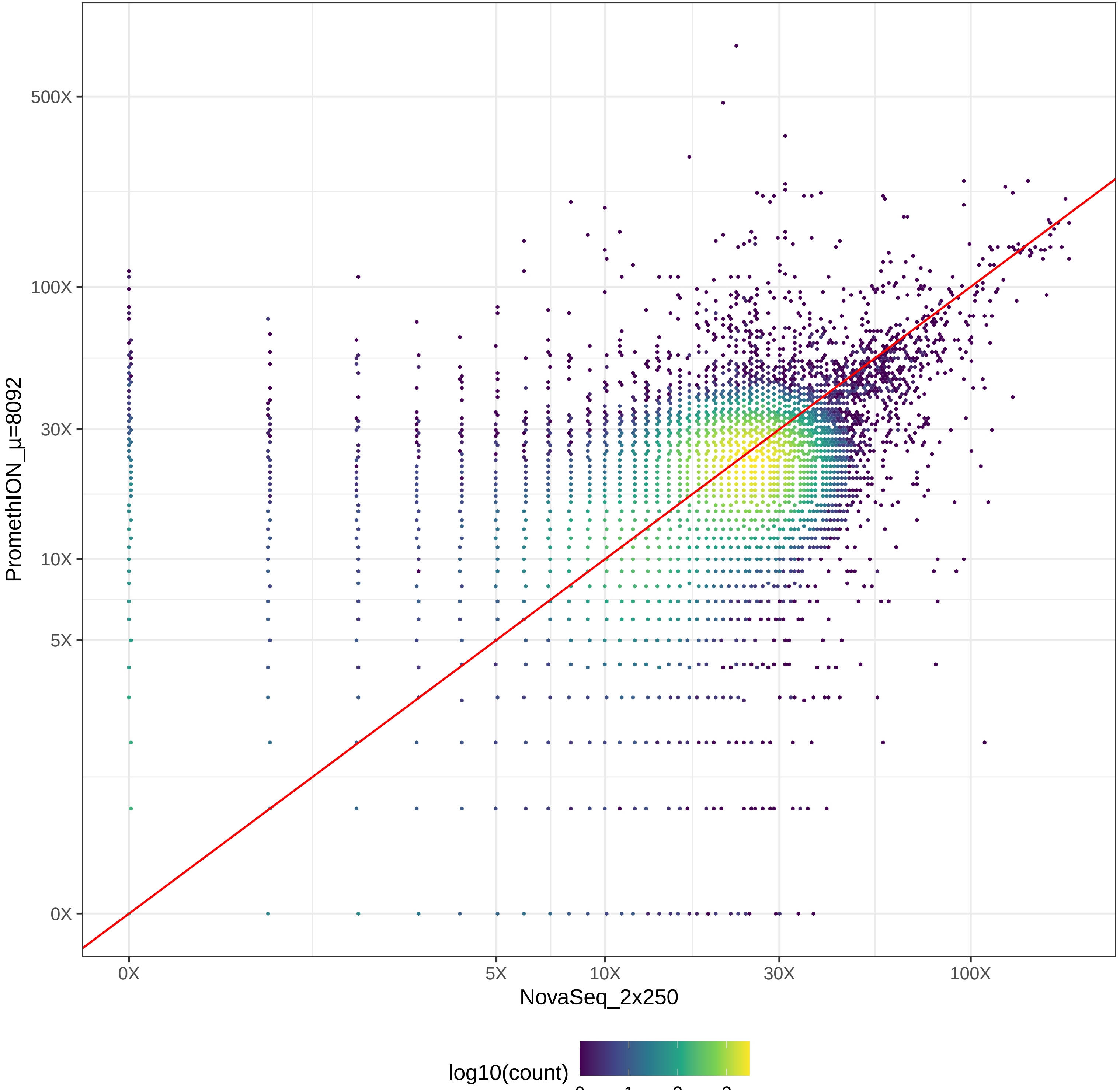

### GIAB_L2_PacBio_CCS-15k-v-PacBio_CLR_hex.pdf

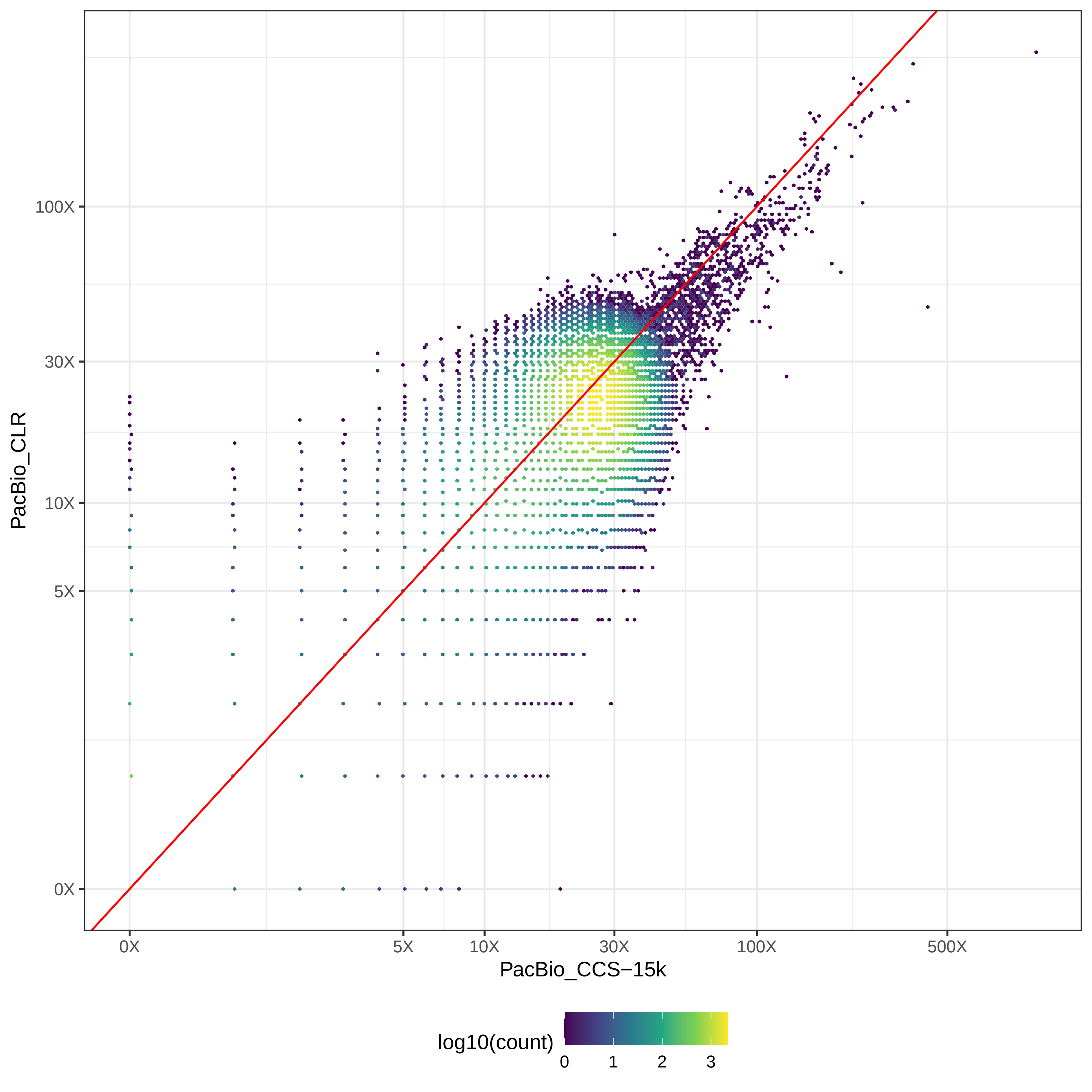

### GIAB_L2_PacBio_CCS-15k-v-PromethION_μ=8092_hex.pdf

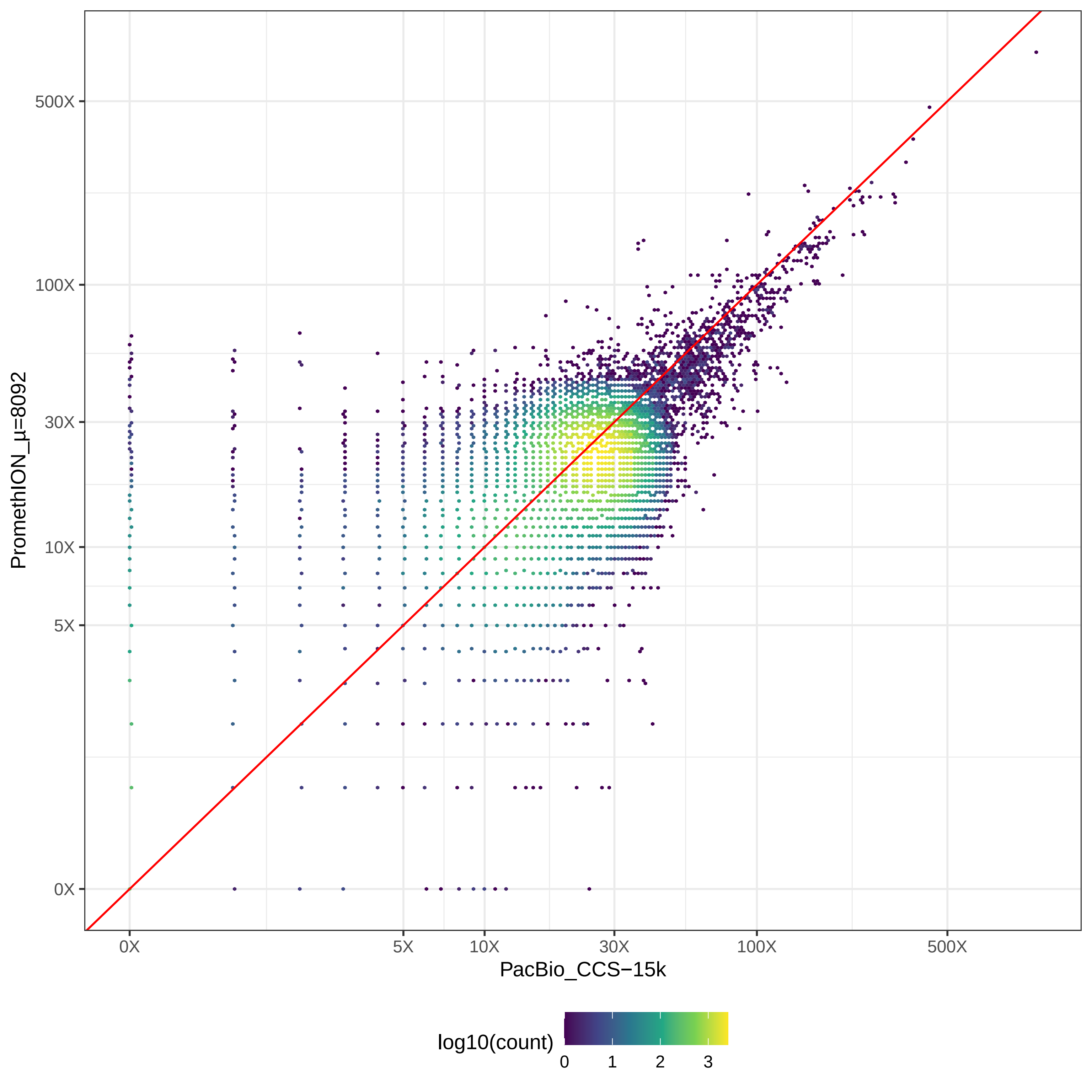

### GIAB_L2_PacBio_CLR-v-PromethION_μ=8092_hex.pdf

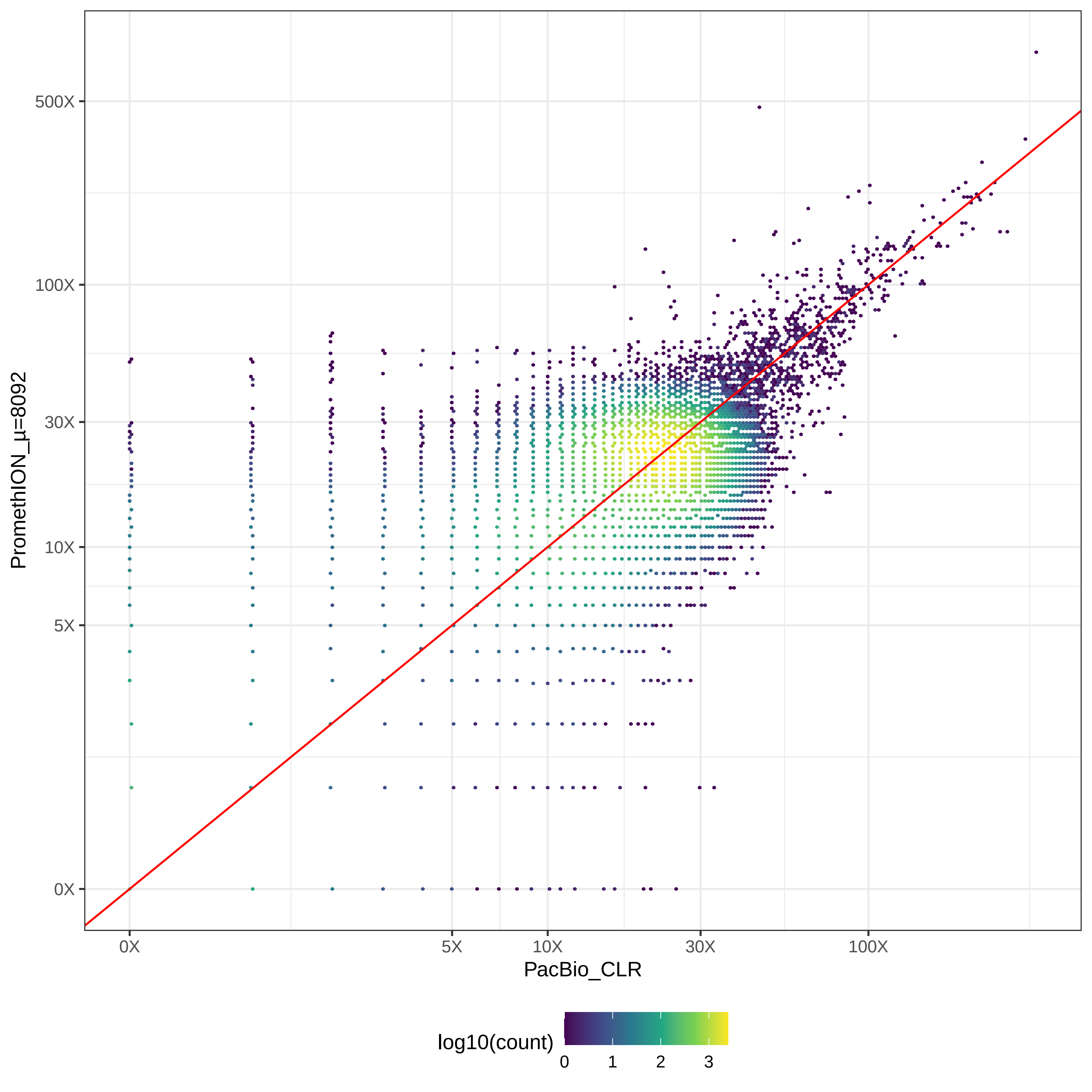

### GIAB_Low_Complexity_BGISEQ500_2x100-v-HiSeq2500_2x100_hex.pdf

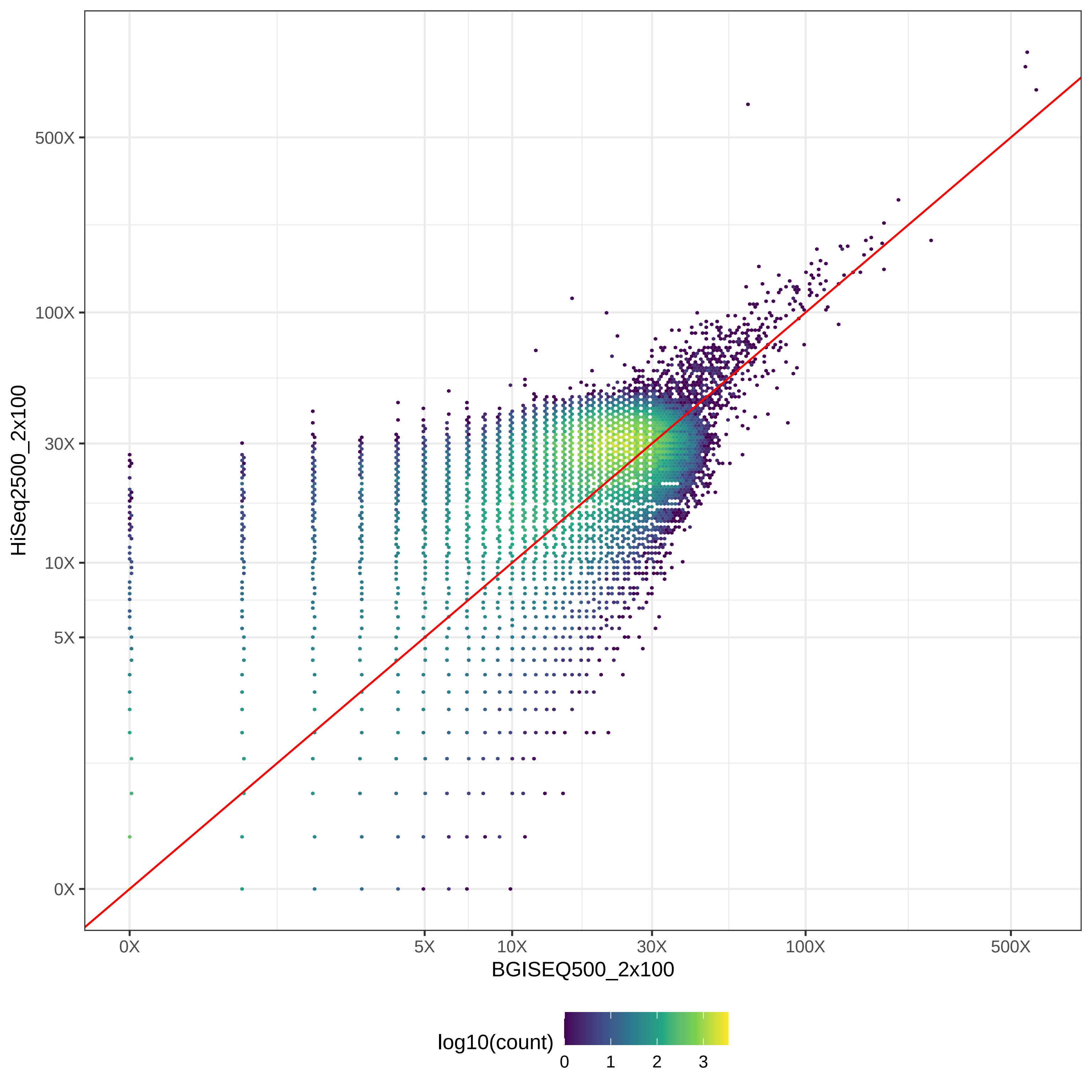

### GIAB_Low_Complexity_BGISEQ500_2x100-v-HiSeq2500_2x126_hex.pdf

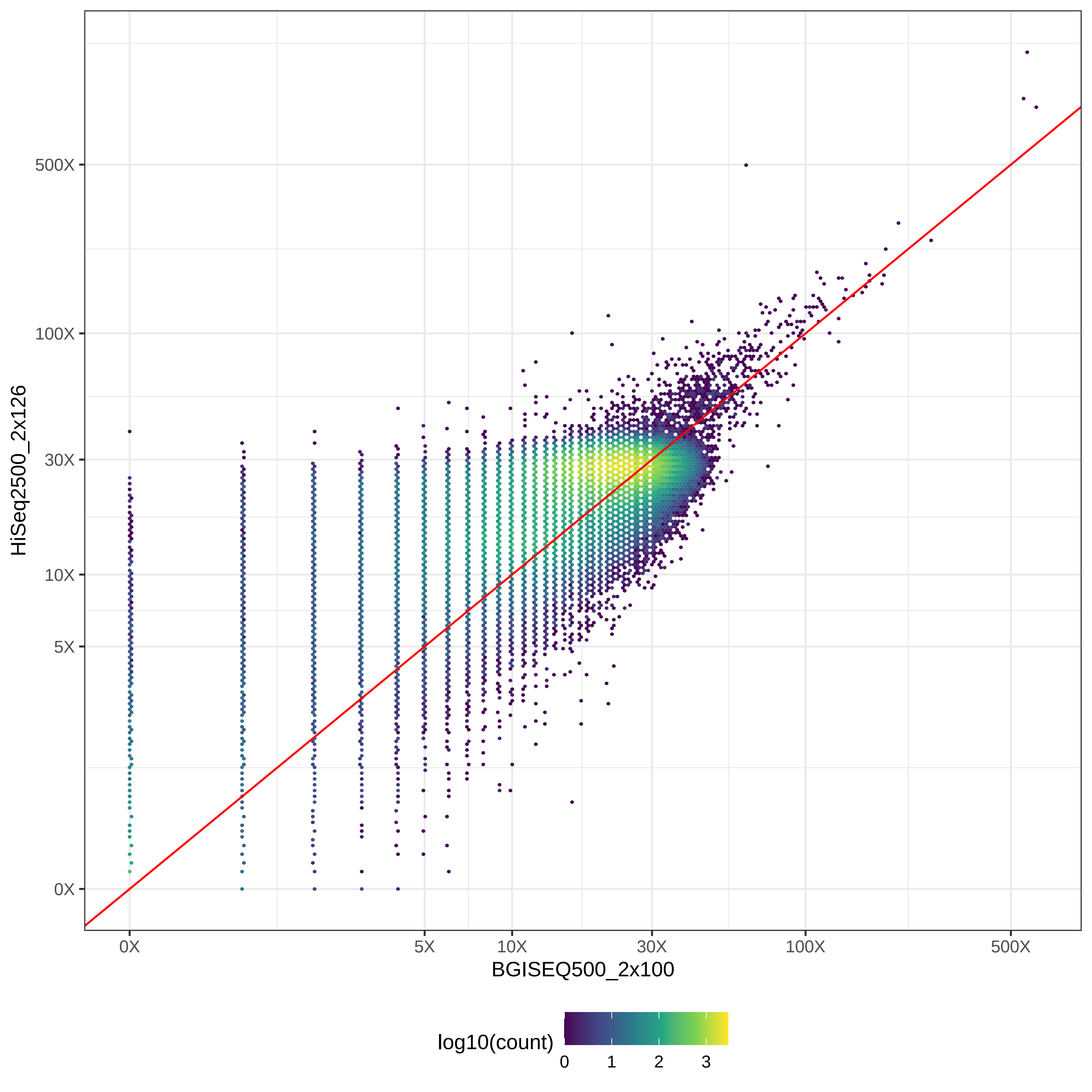

### GIAB_Low_Complexity_BGISEQ500_2x100-v-HiSeq4000_2x150_hex.pdf

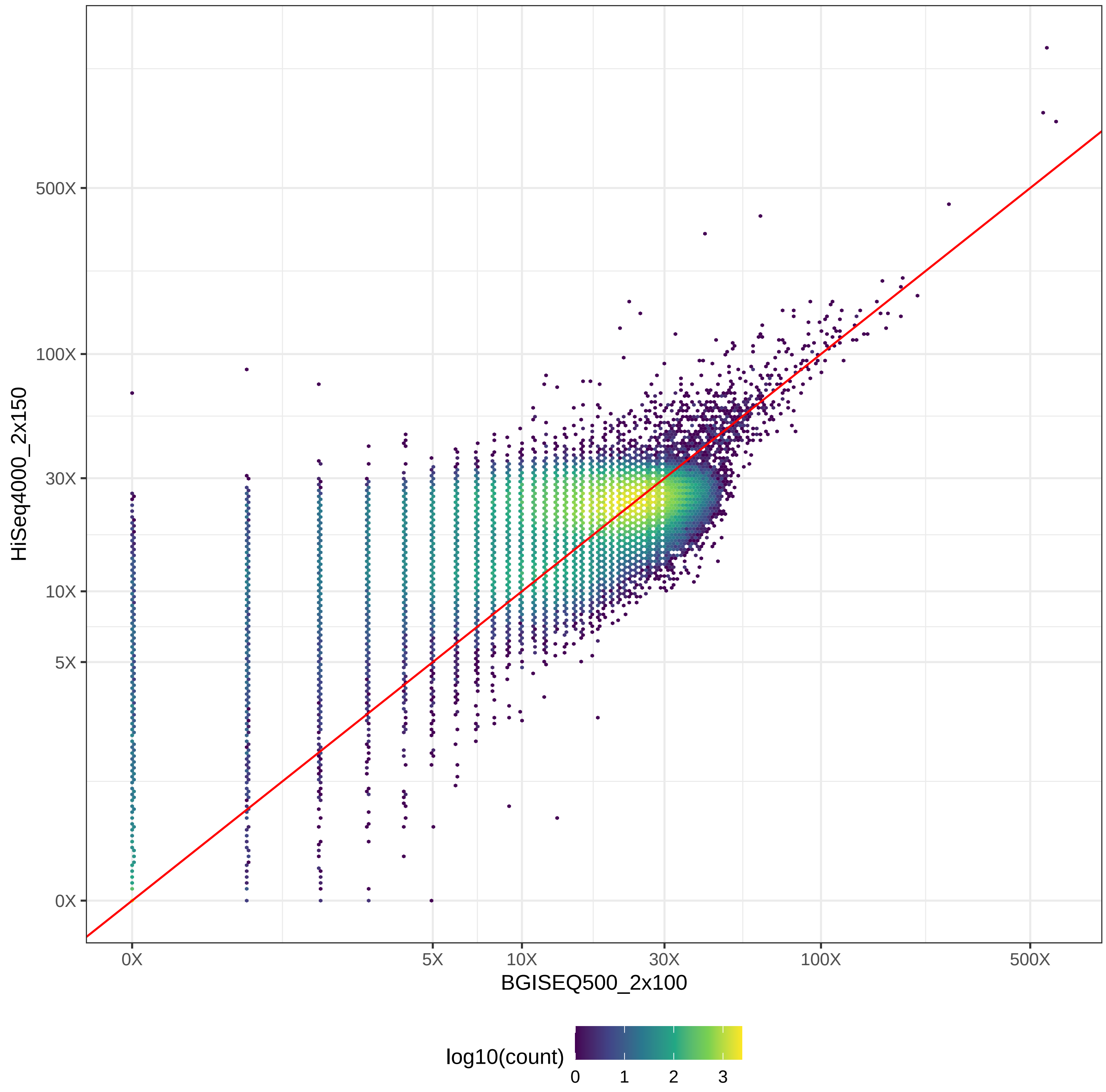

### GIAB_Low_Complexity_BGISEQ500_2x100-v-HiSeqX10_2x150_hex.pdf

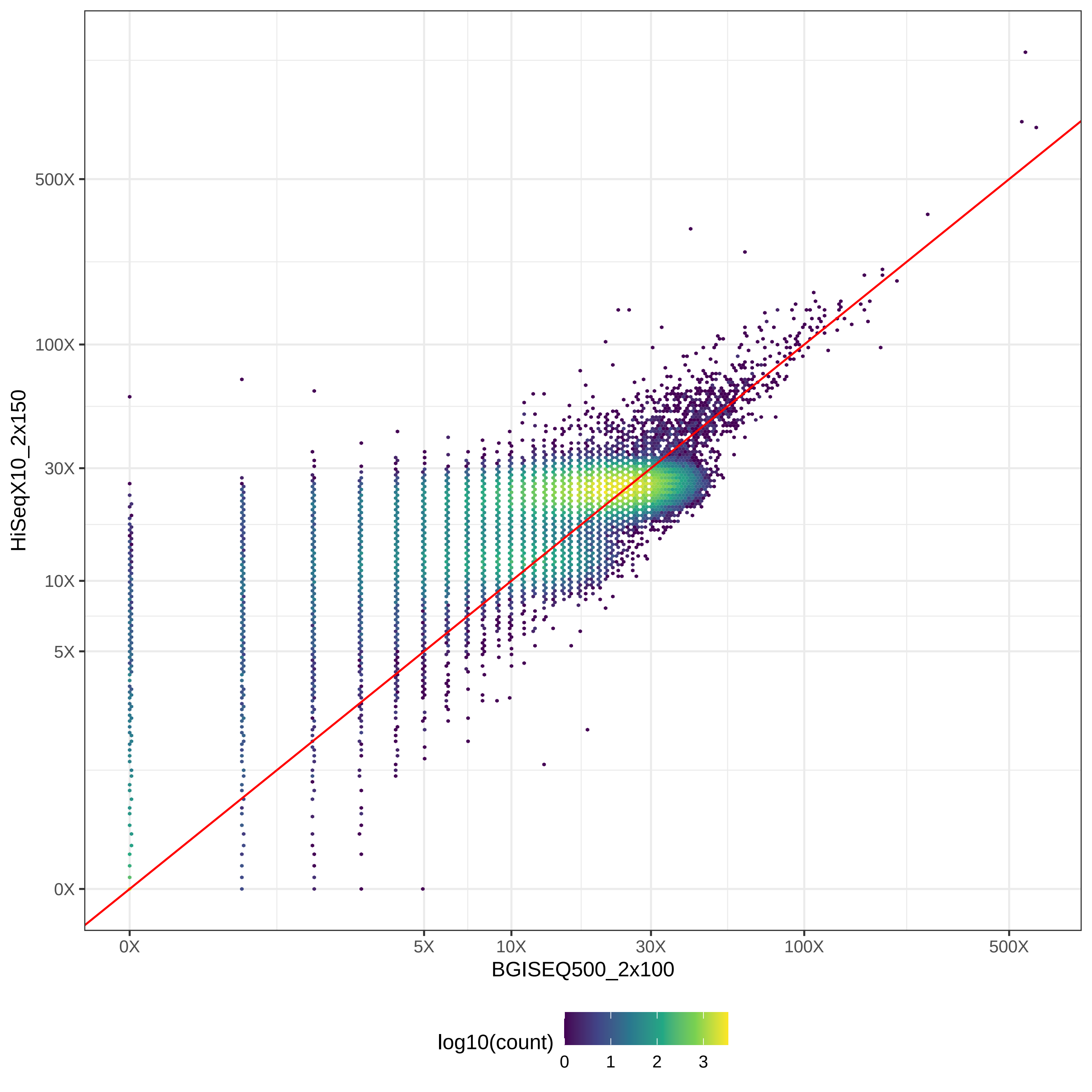

### GIAB_Low_Complexity_BGISEQ500_2x100-v-MGISEQ2000_2x150_hex.pdf

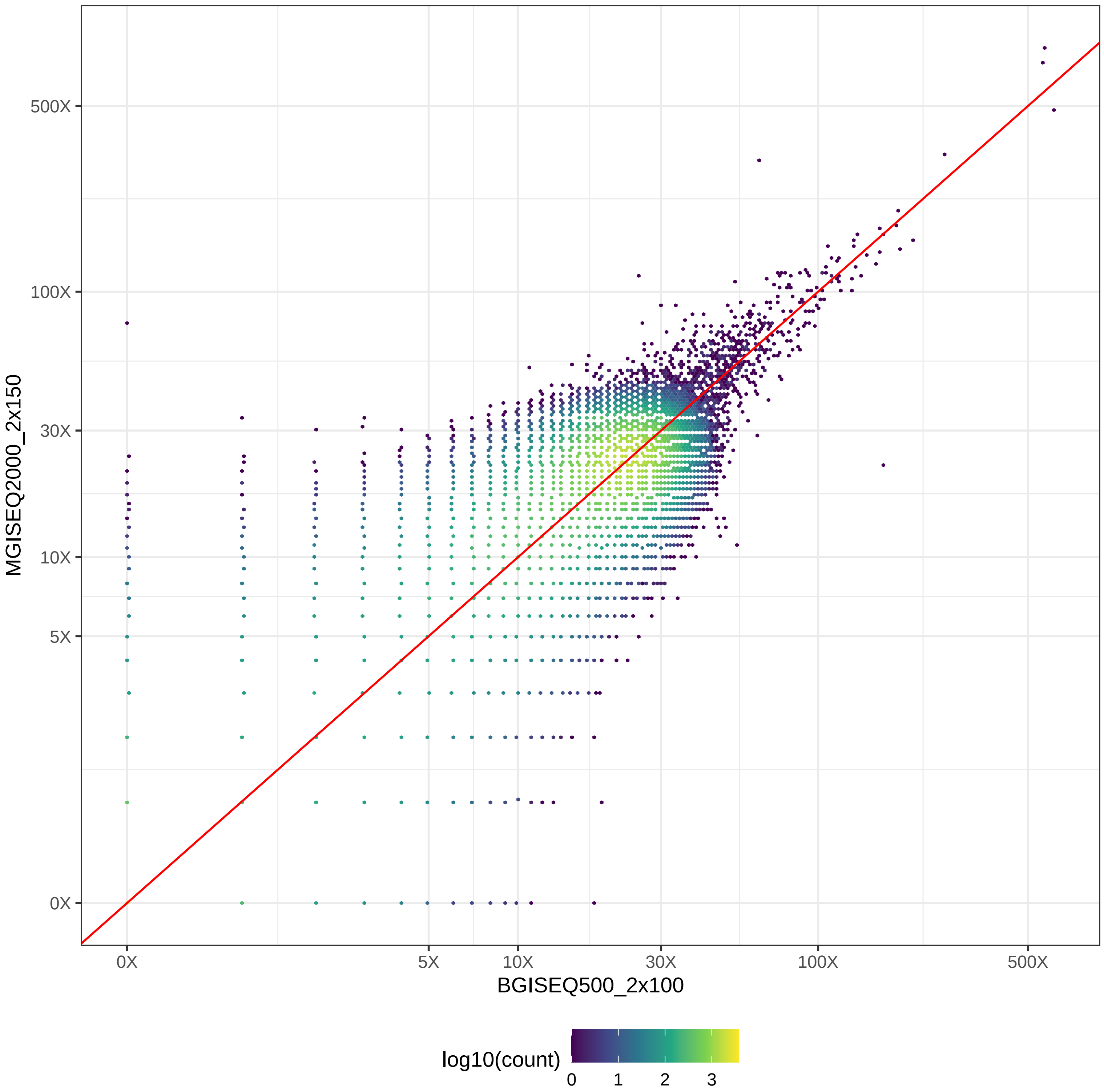

### GIAB_Low_Complexity_BGISEQ500_2x100-v-NovaSeq_2x150_hex.pdf

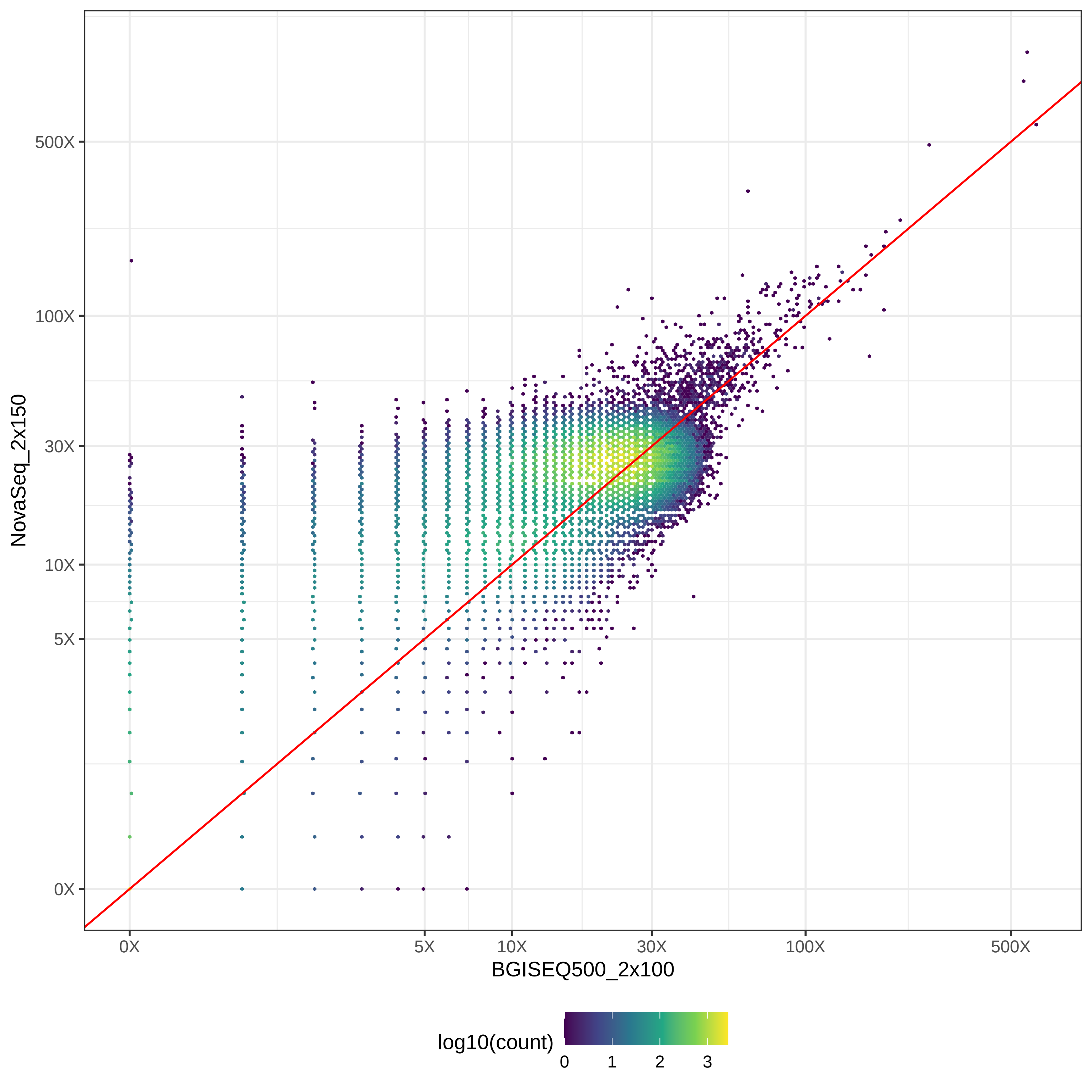

### GIAB_Low_Complexity_BGISEQ500_2x100-v-NovaSeq_2x250_hex.pdf

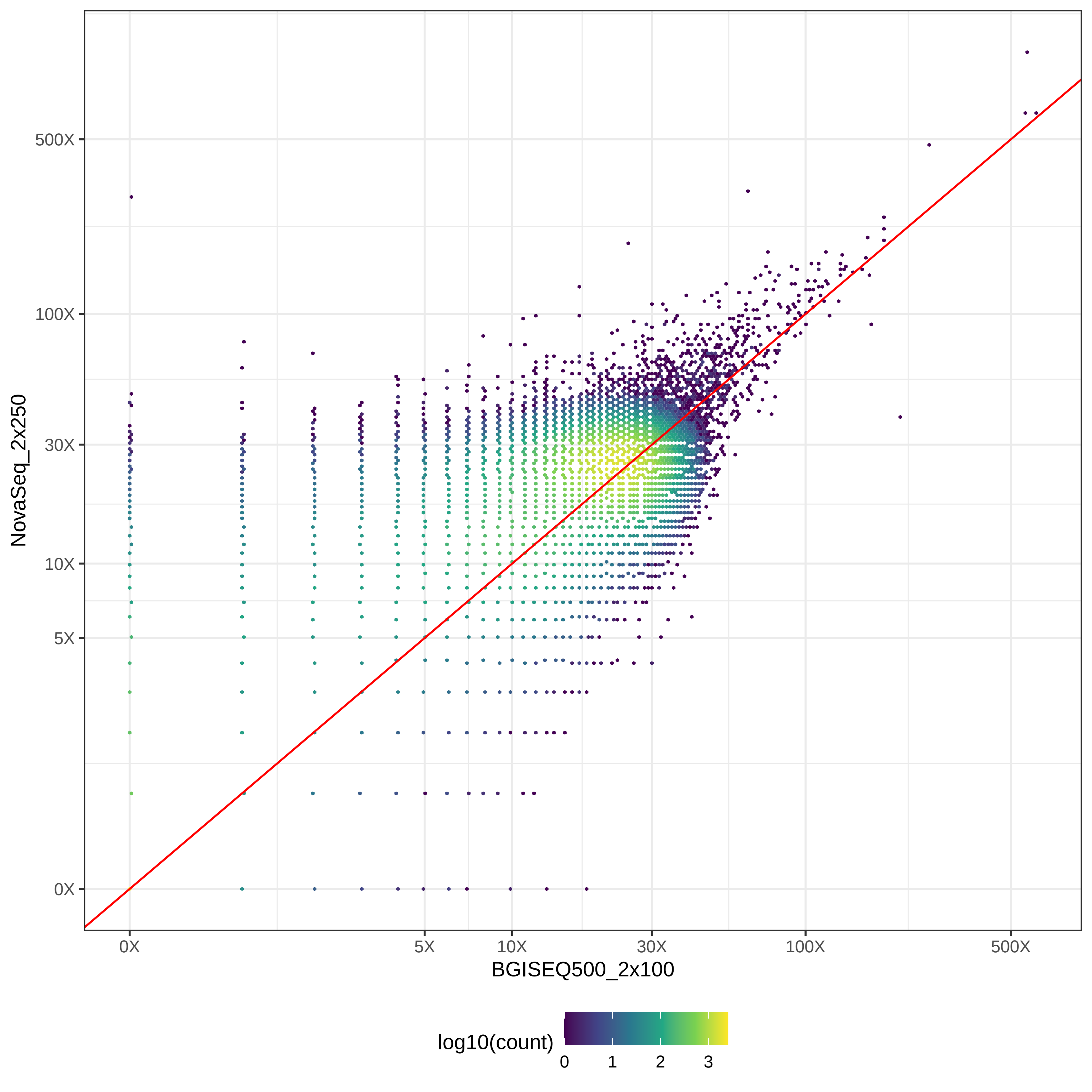

### GIAB_Low_Complexity_BGISEQ500_2x100-v-PacBio_CCS-15k_hex.pdf

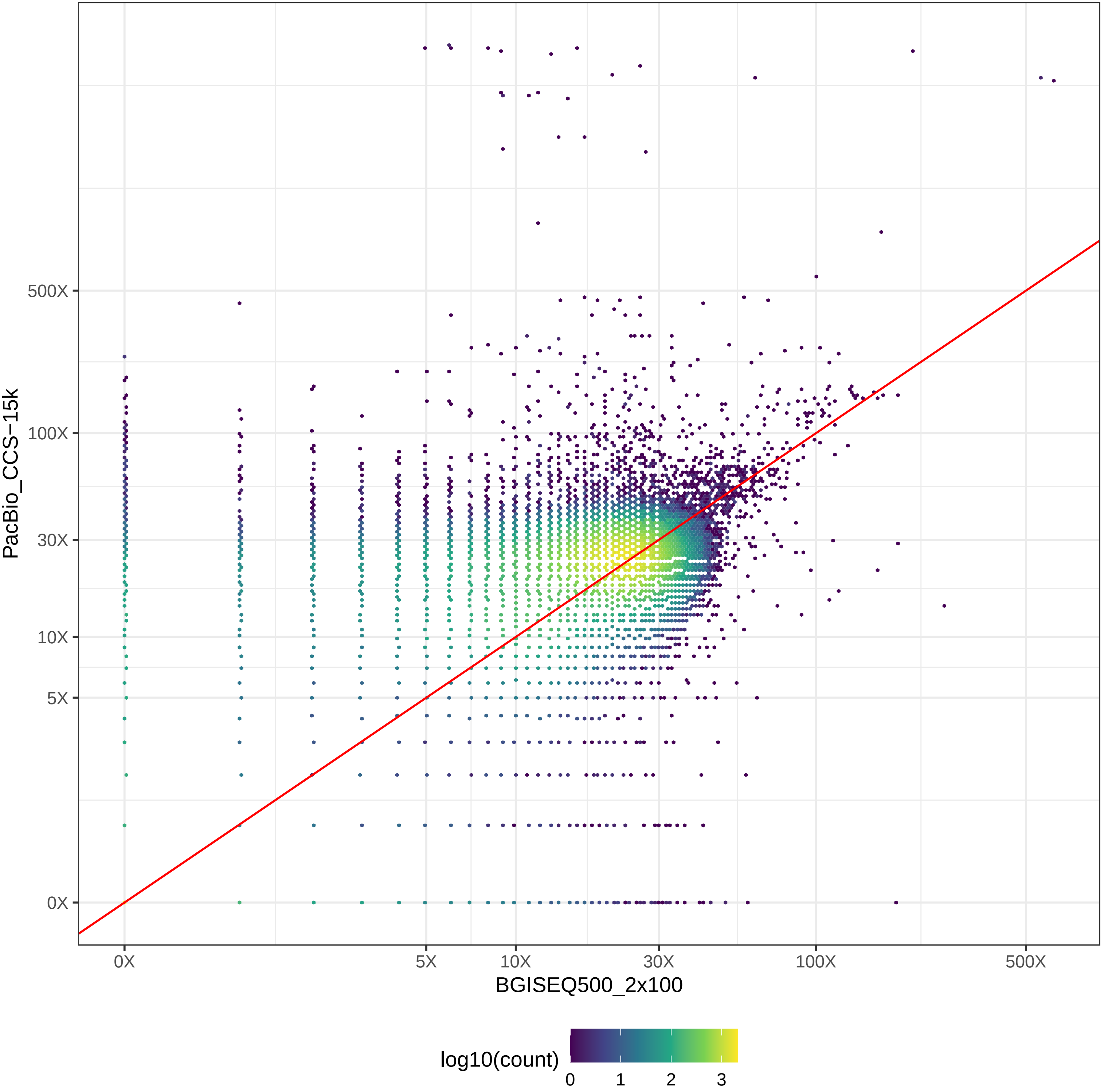

### GIAB_Low_Complexity_BGISEQ500_2x100-v-PacBio_CLR_hex.pdf

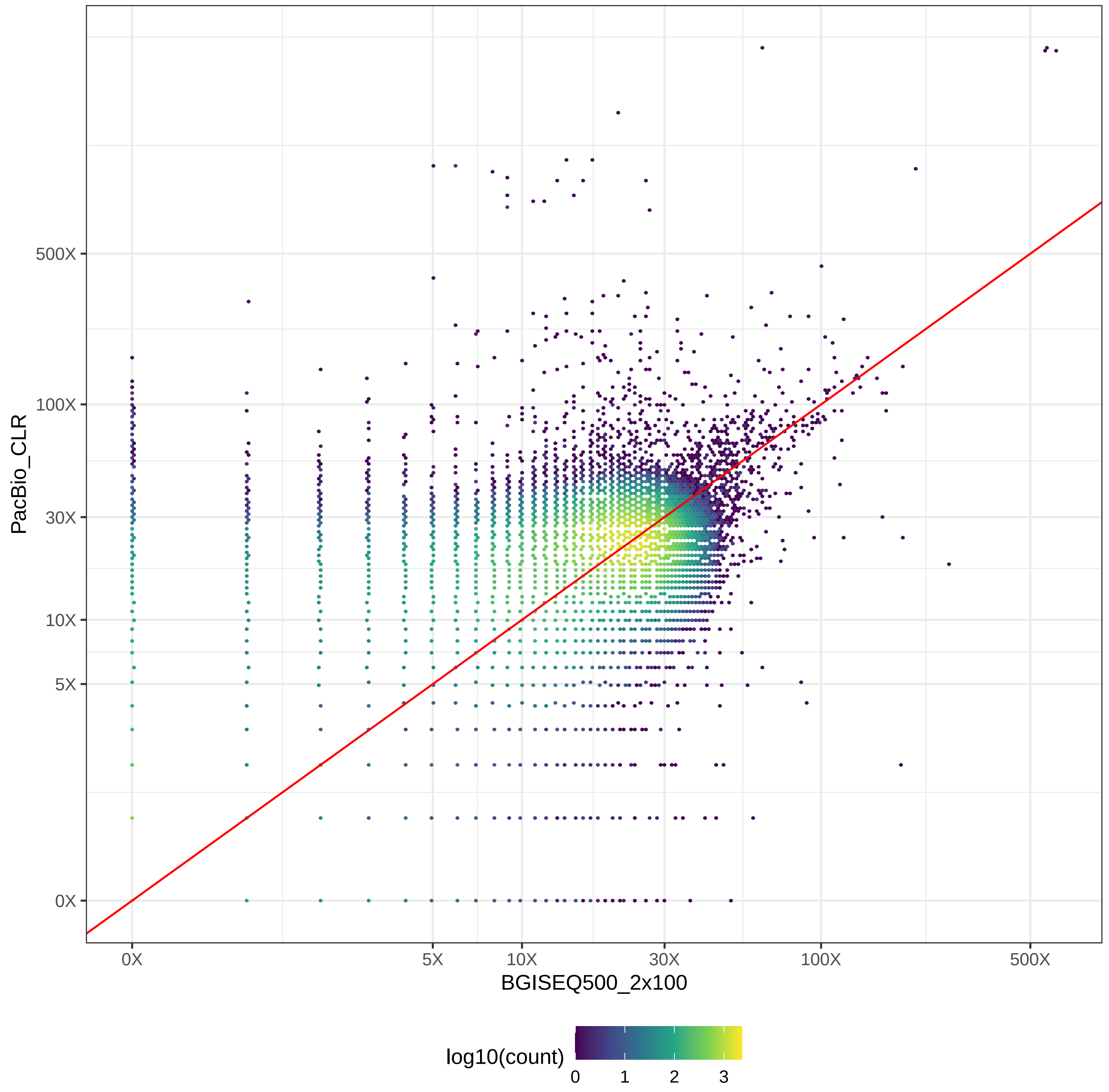

### GIAB_Low_Complexity_BGISEQ500_2x100-v-PromethION_μ=8092_hex.pdf

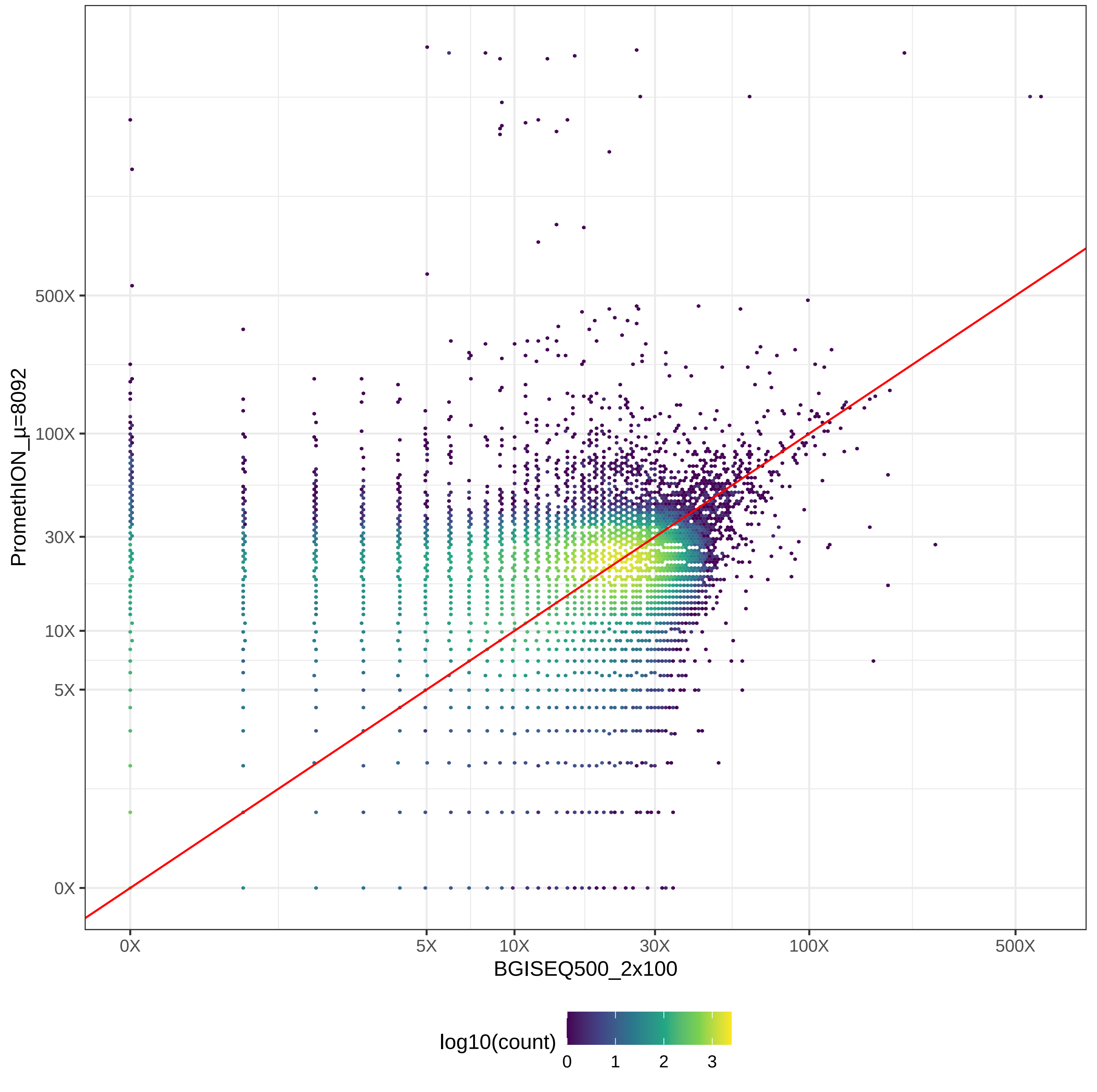

### GIAB_Low_Complexity_HiSeq2500_2x100-v-HiSeq2500_2x126_hex.pdf

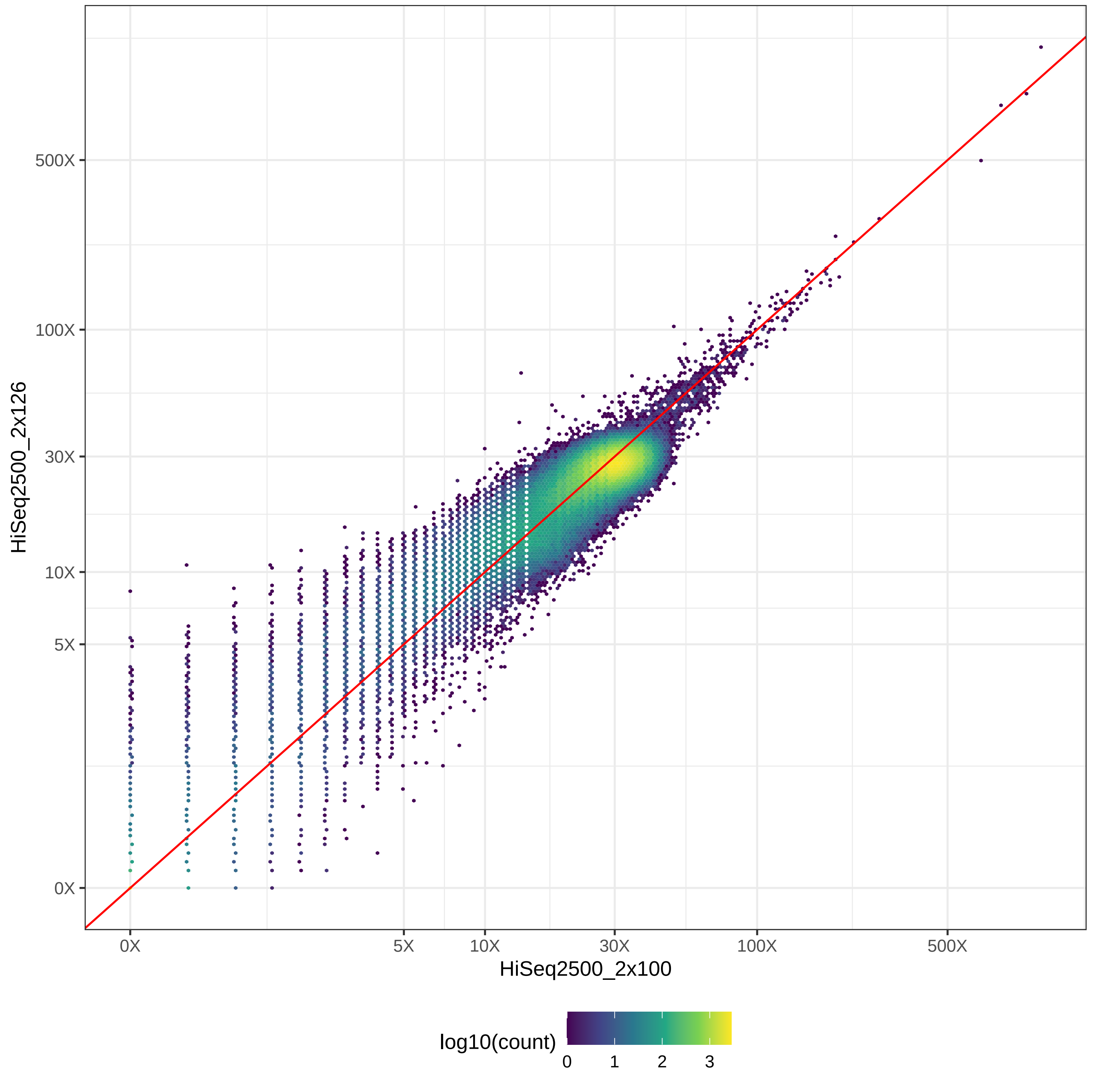

### GIAB_Low_Complexity_HiSeq2500_2x100-v-HiSeq4000_2x150_hex.pdf

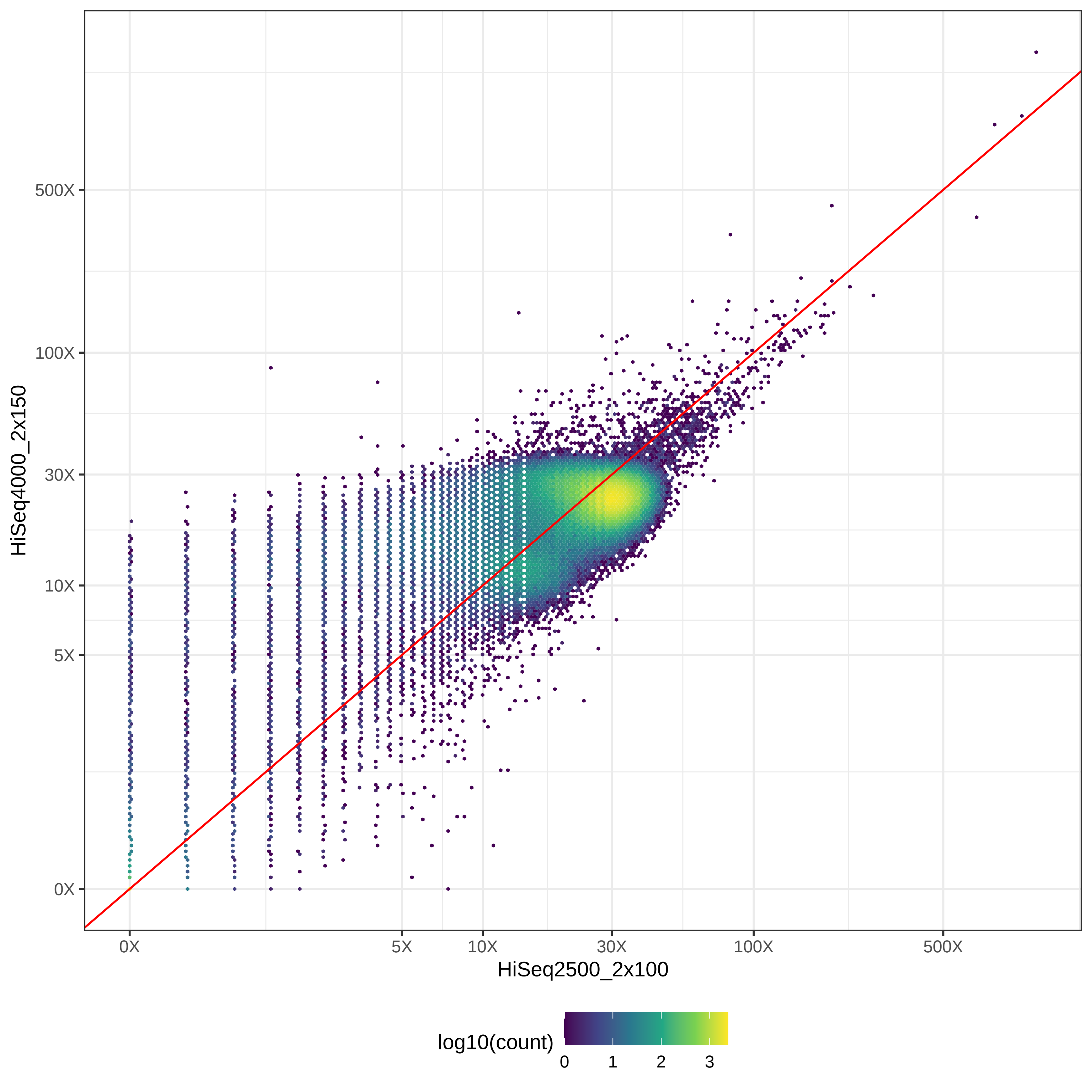

### GIAB_Low_Complexity_HiSeq2500_2x100-v-HiSeqX10_2x150_hex.pdf

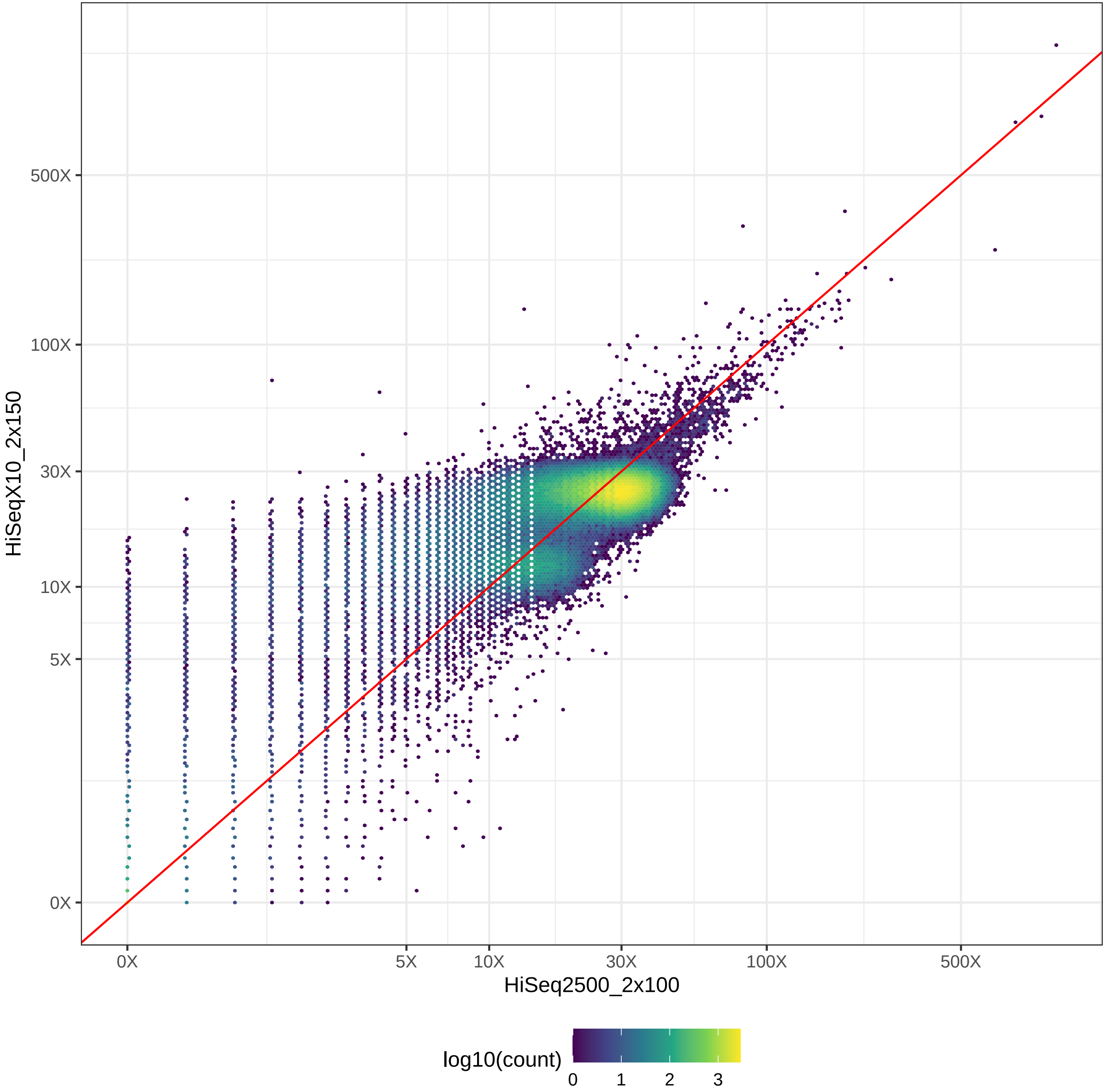

### GIAB_Low_Complexity_HiSeq2500_2x100-v-MGISEQ2000_2x150_hex.pdf

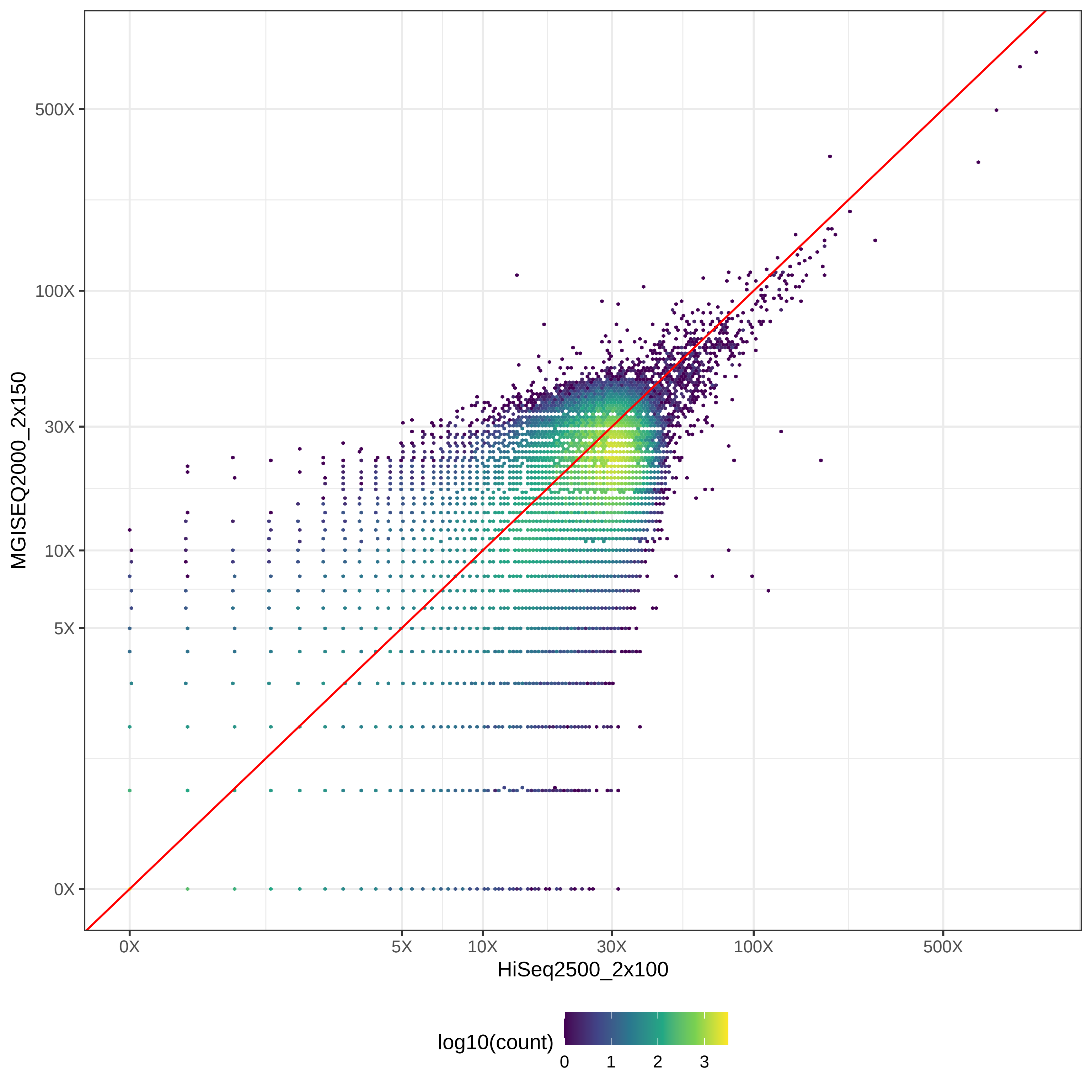

### GIAB_Low_Complexity_HiSeq2500_2x100-v-NovaSeq_2x150_hex.pdf

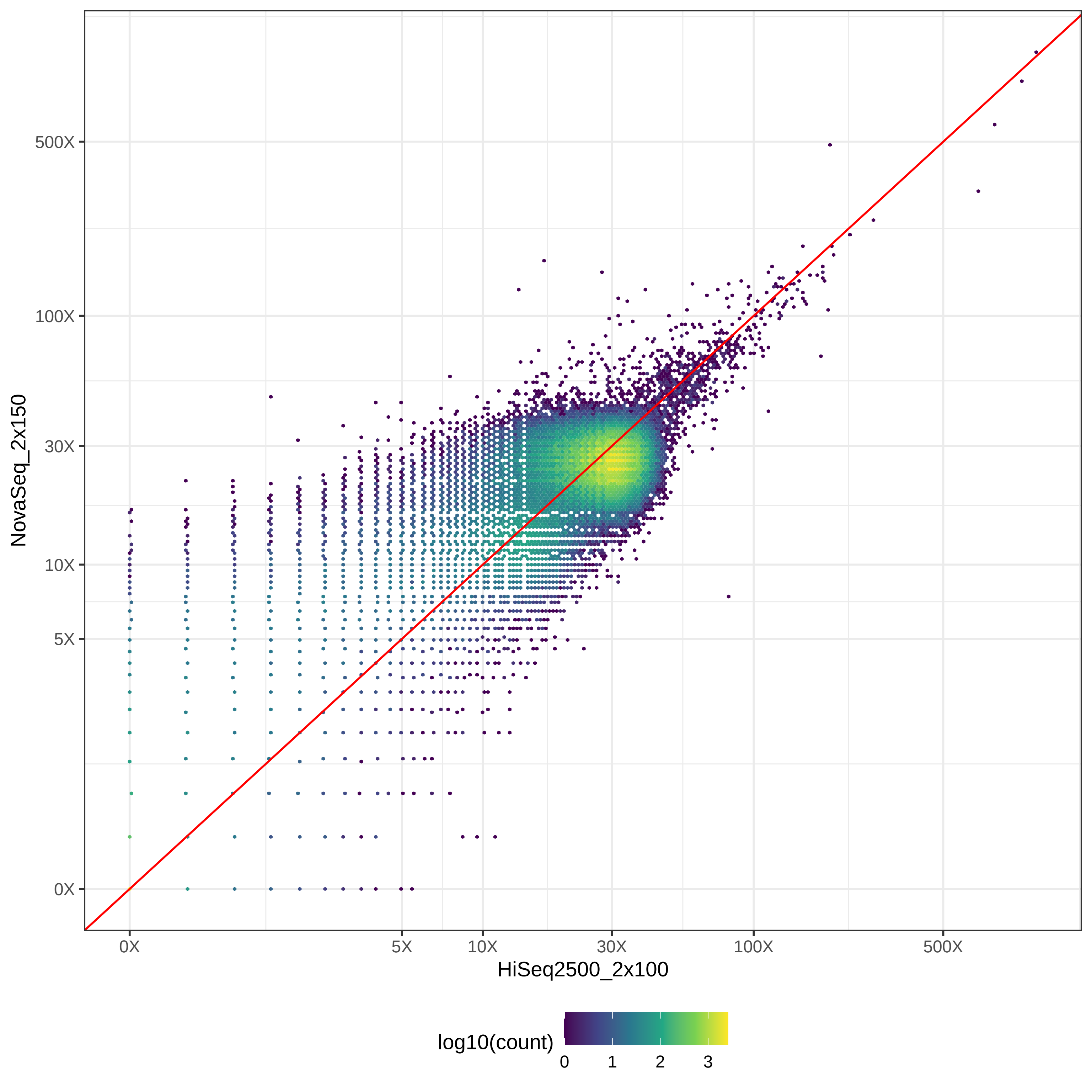

### GIAB_Low_Complexity_HiSeq2500_2x100-v-NovaSeq_2x250_hex.pdf

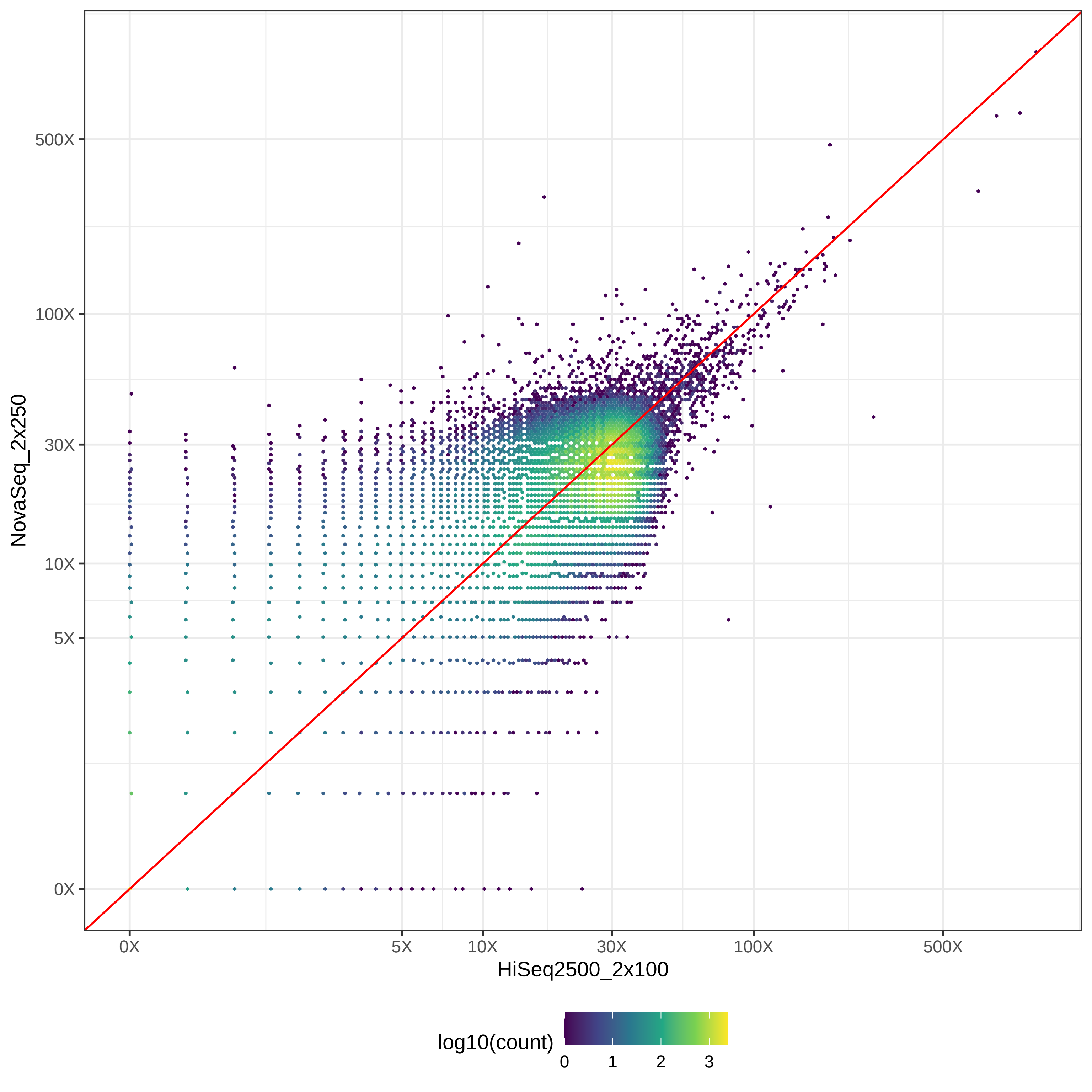

### GIAB_Low_Complexity_HiSeq2500_2x100-v-PacBio_CCS-15k_hex.pdf

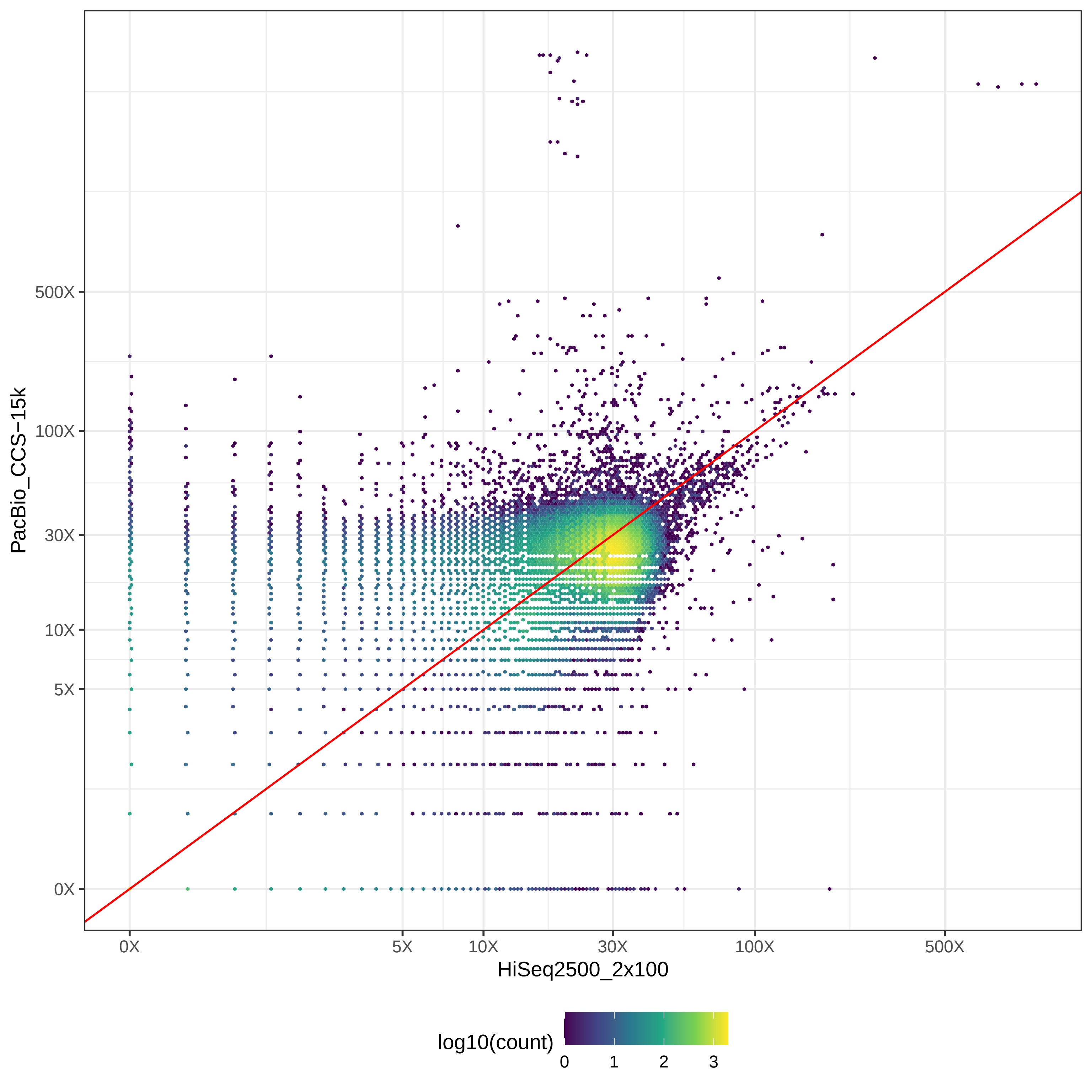

### GIAB_Low_Complexity_HiSeq2500_2x100-v-PacBio_CLR_hex.pdf

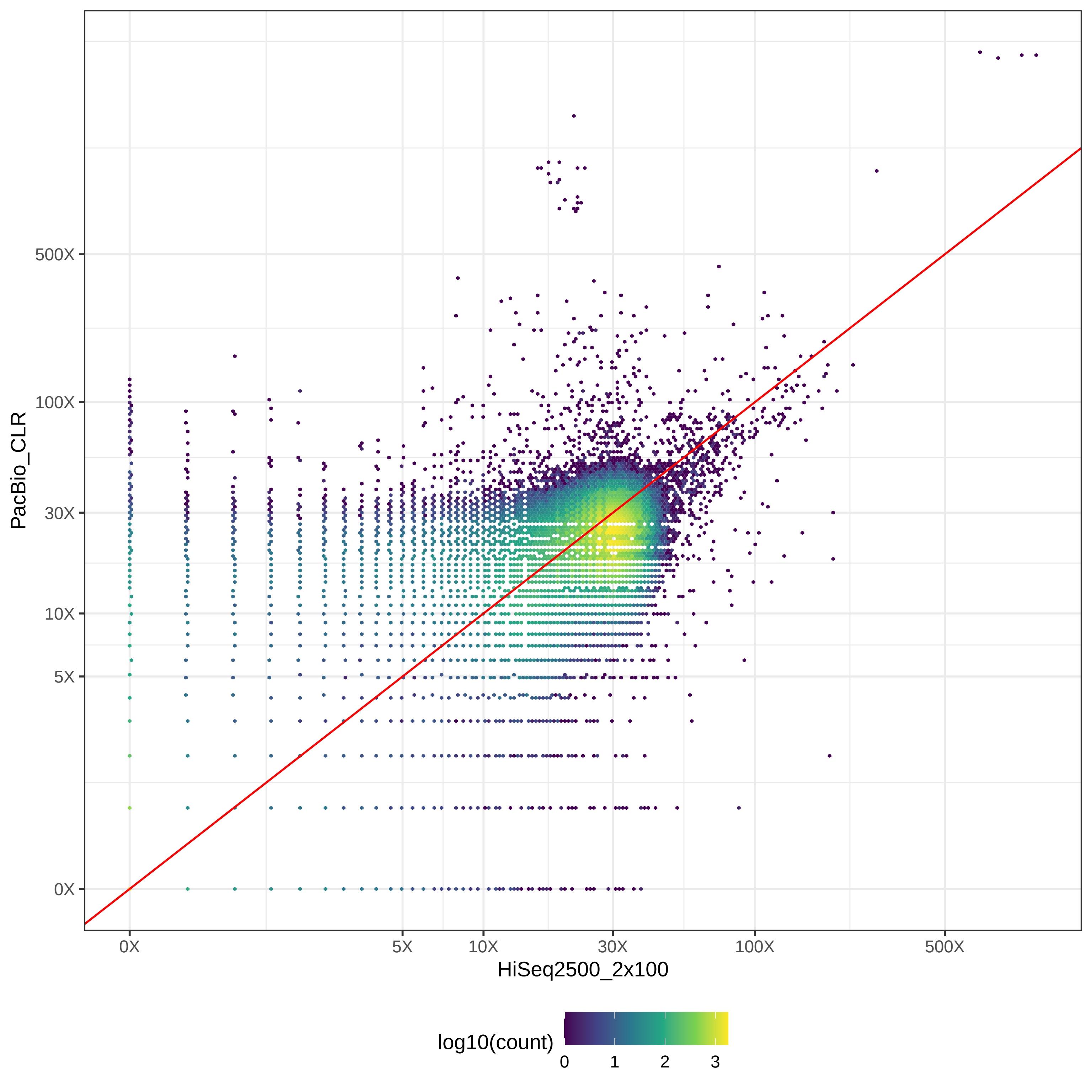

### GIAB_Low_Complexity_HiSeq2500_2x100-v-PromethION_μ=8092_hex.pdf

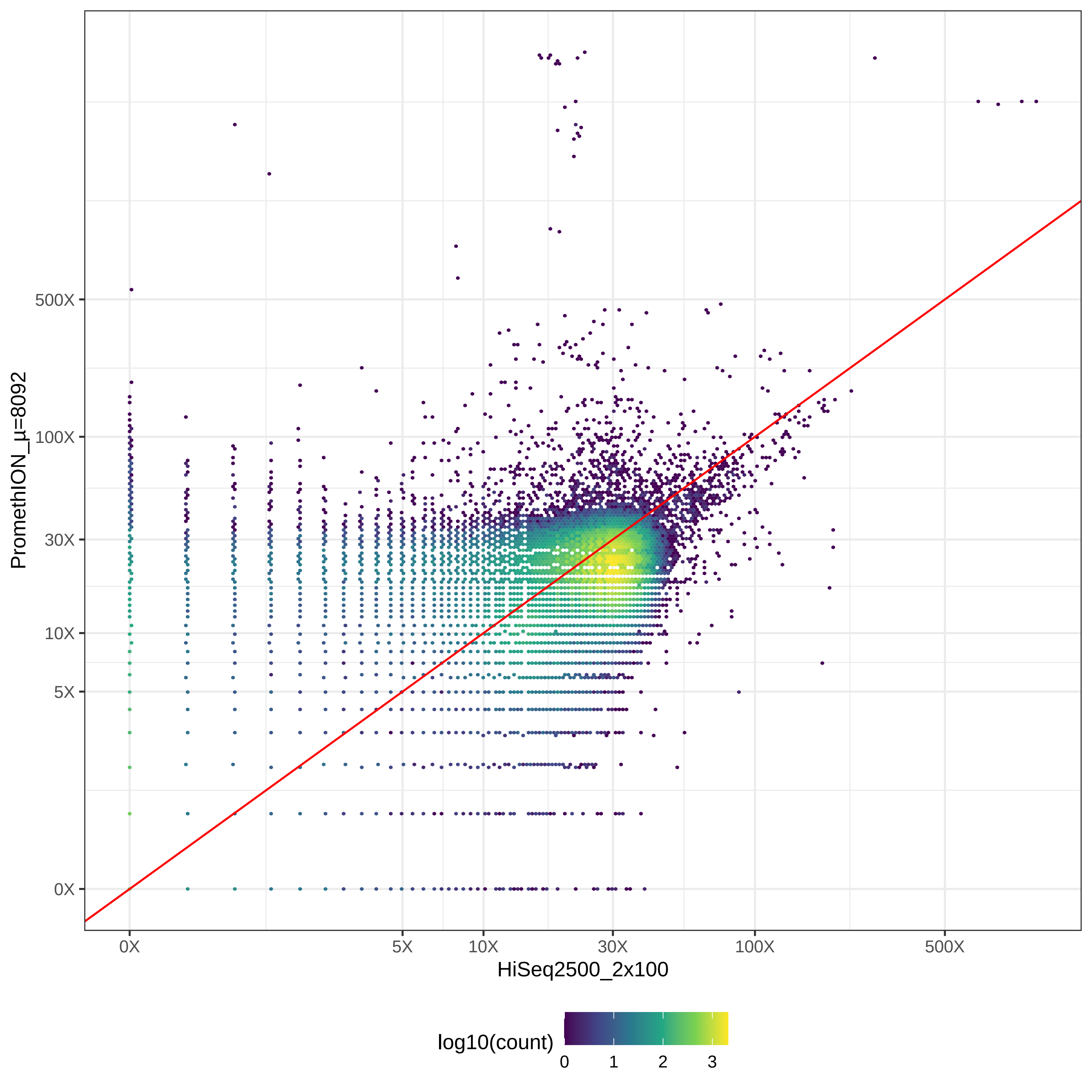

### GIAB_Low_Complexity_HiSeq2500_2x126-v-HiSeq4000_2x150_hex.pdf

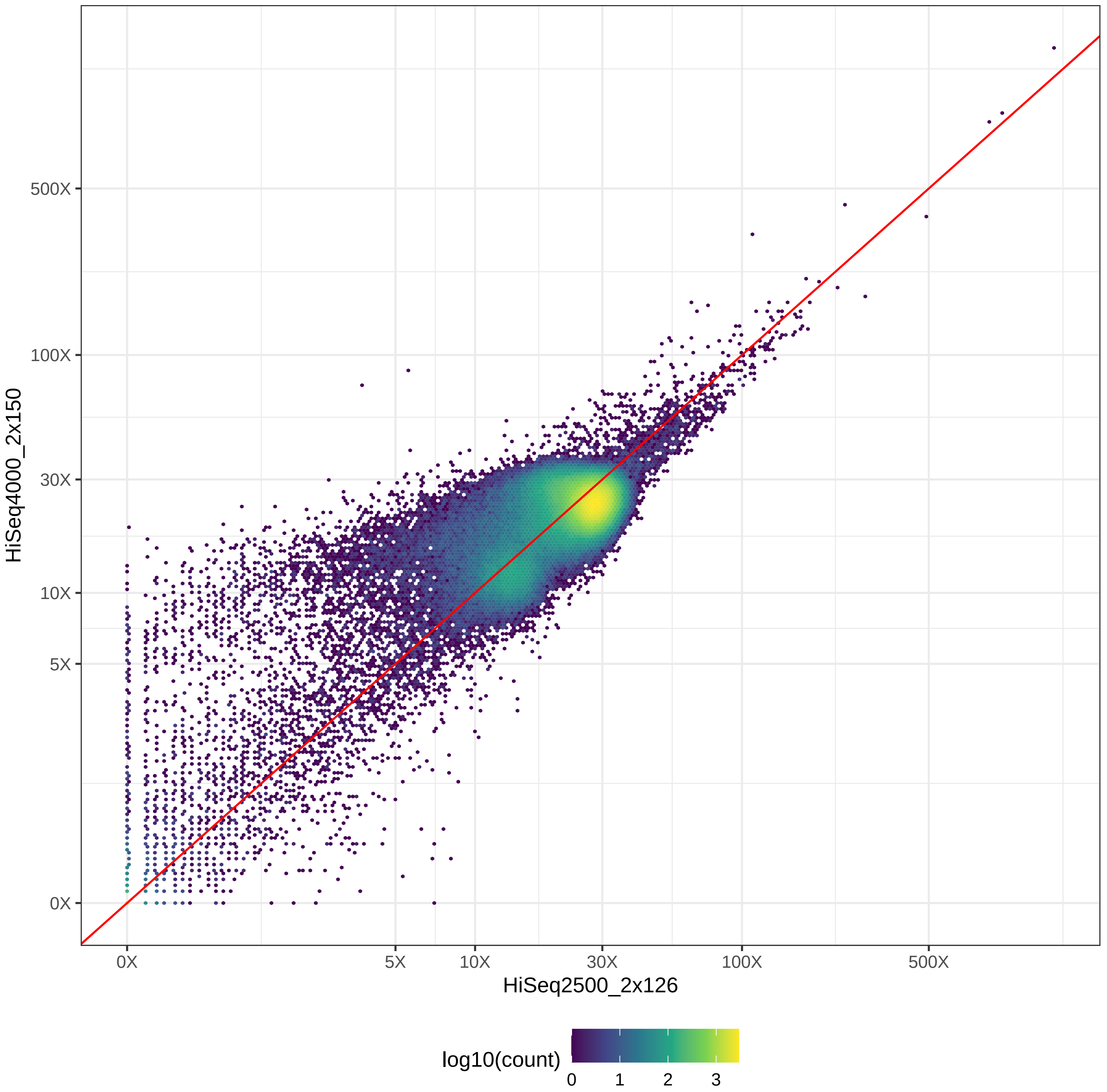

### GIAB_Low_Complexity_HiSeq2500_2x126-v-HiSeqX10_2x150_hex.pdf

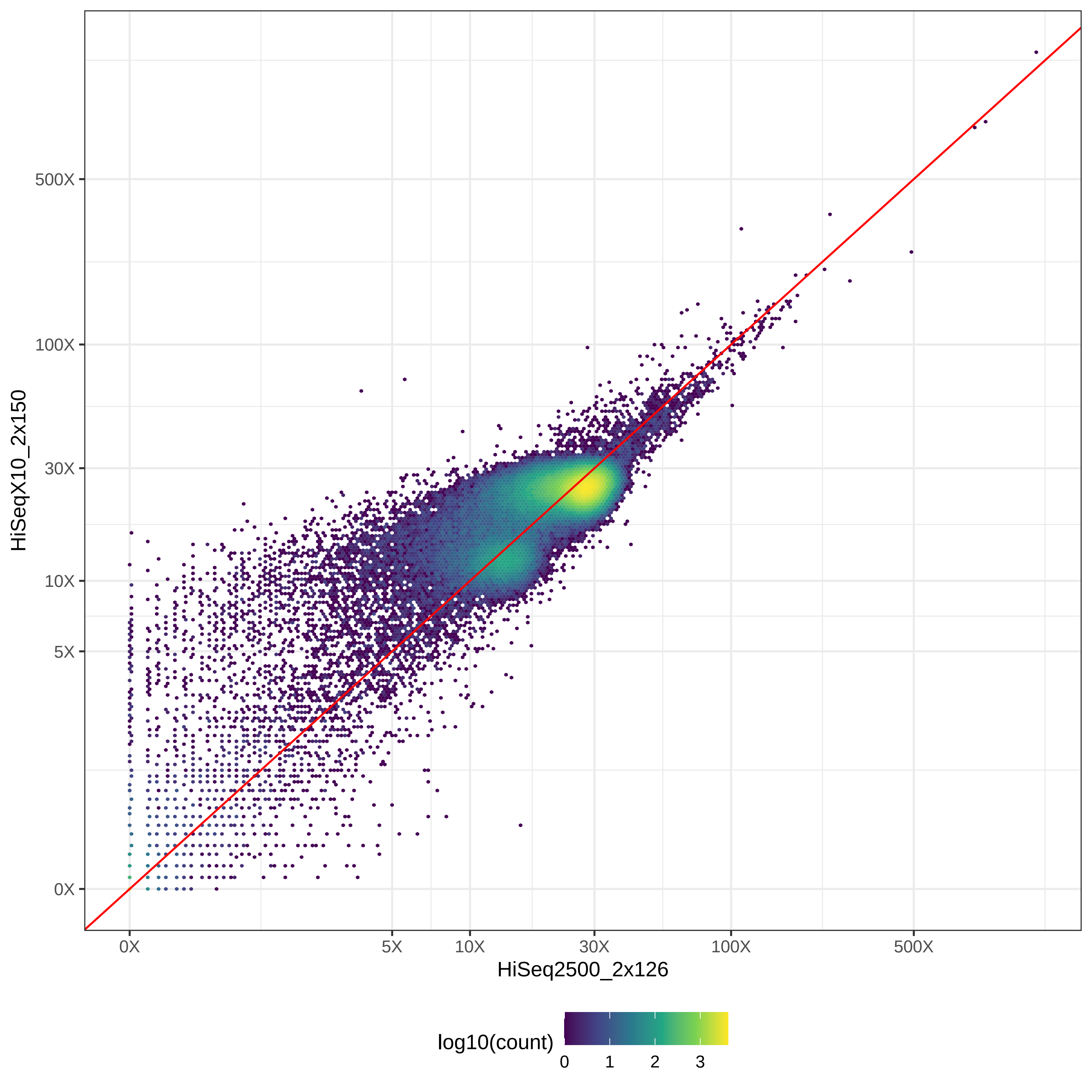

### GIAB_Low_Complexity_HiSeq2500_2x126-v-MGISEQ2000_2x150_hex.pdf

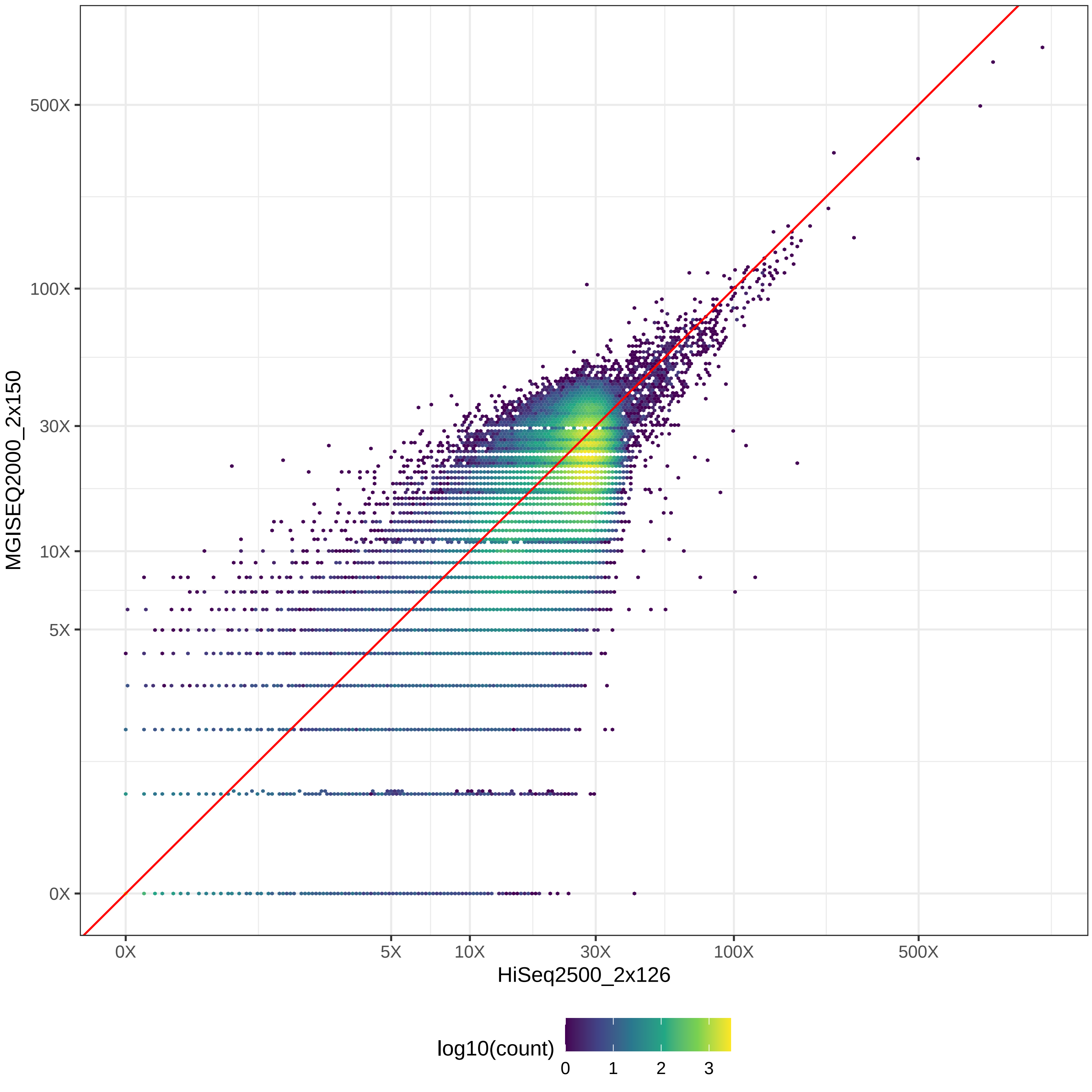

### GIAB_Low_Complexity_HiSeq2500_2x126-v-NovaSeq_2x150_hex.pdf

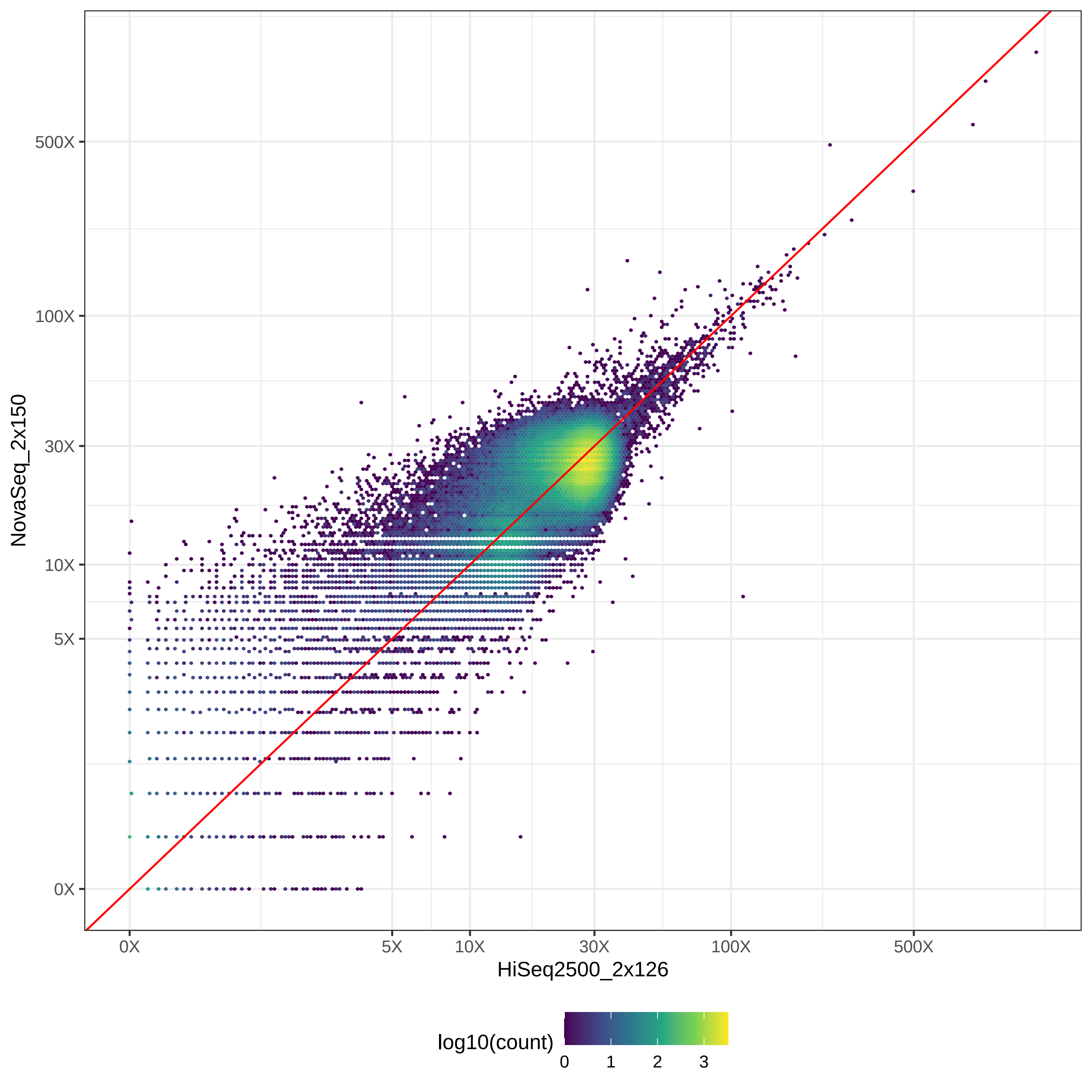

### GIAB_Low_Complexity_HiSeq2500_2x126-v-NovaSeq_2x250_hex.pdf

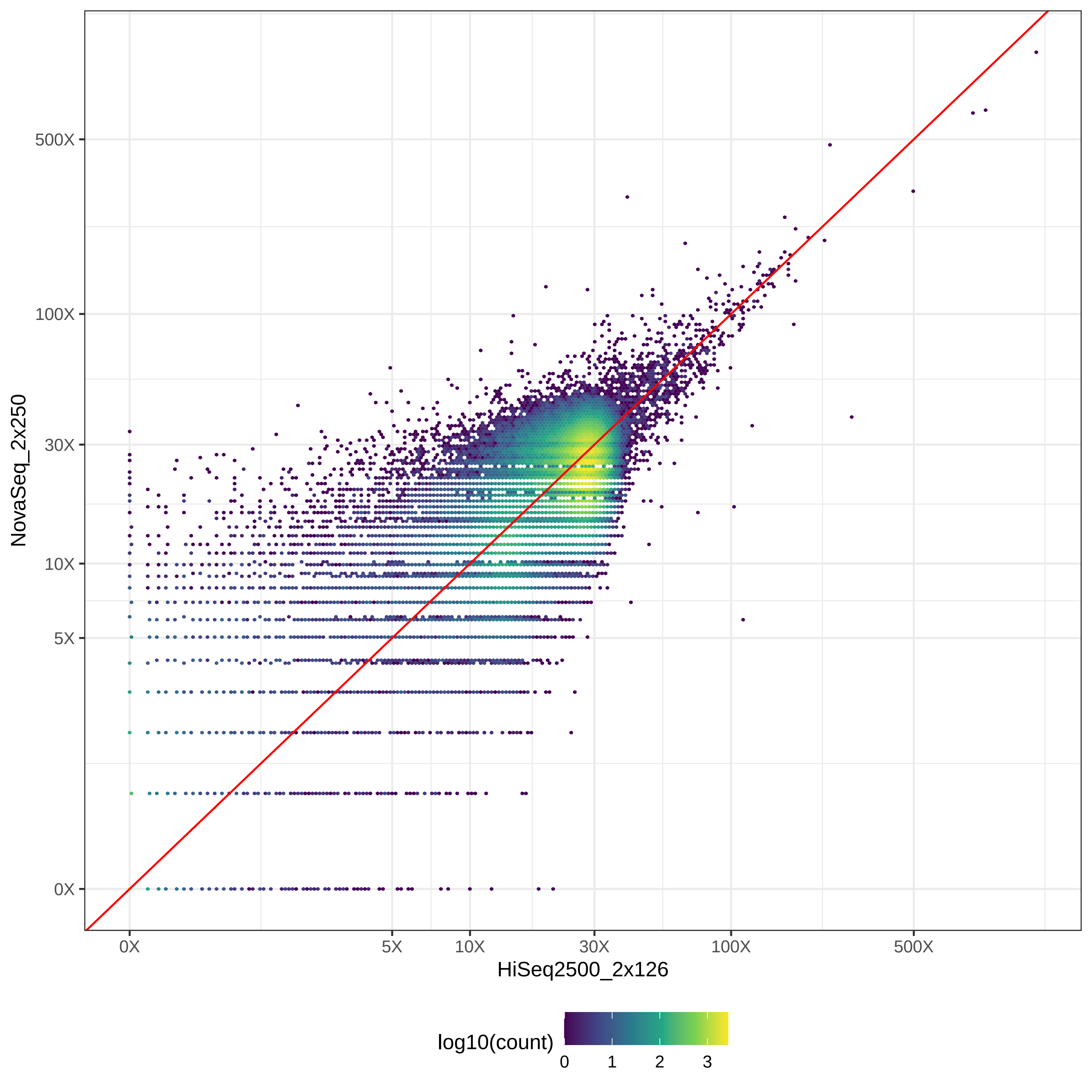

### GIAB_Low_Complexity_HiSeq2500_2x126-v-PacBio_CCS-15k_hex.pdf

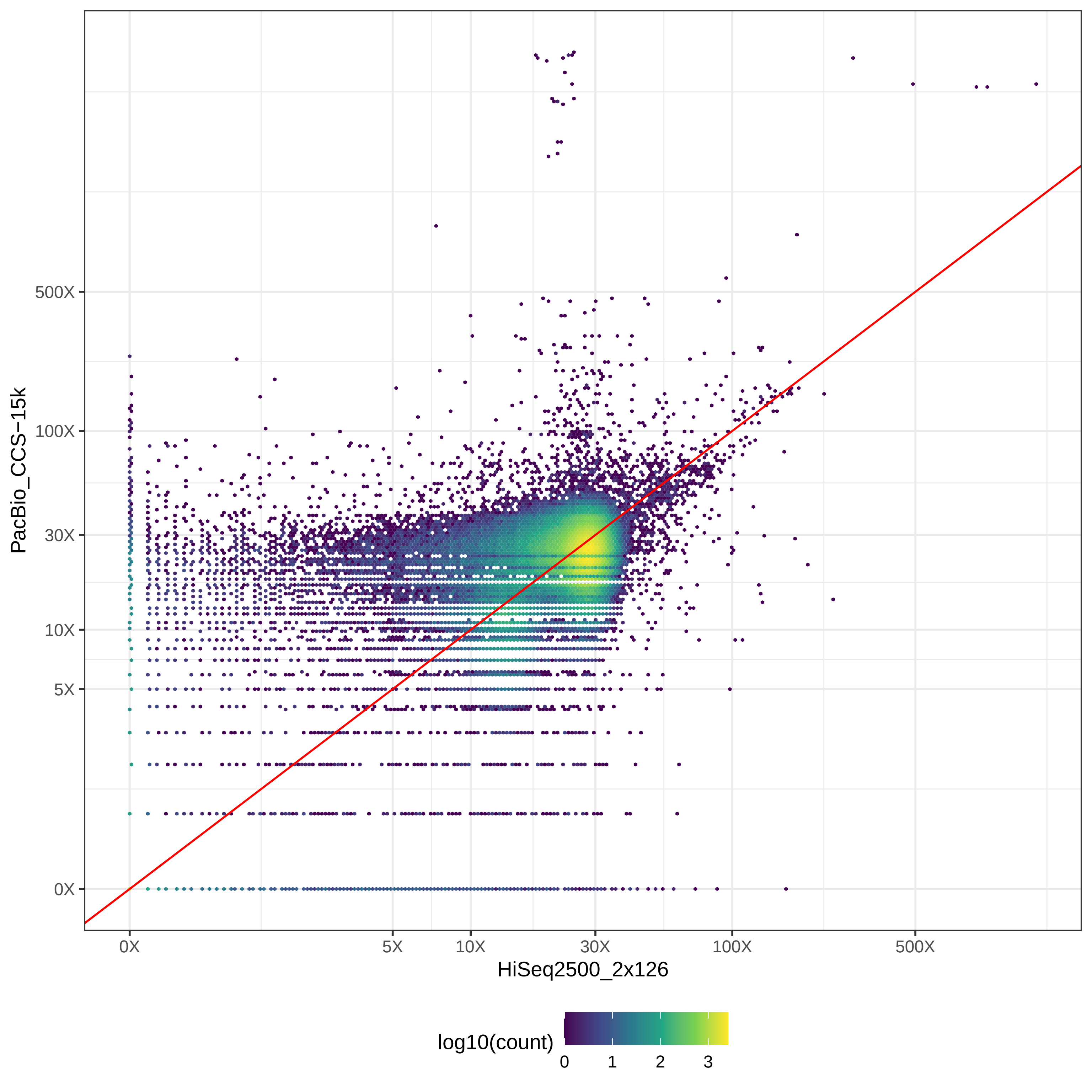

### GIAB_Low_Complexity_HiSeq2500_2x126-v-PacBio_CLR_hex.pdf

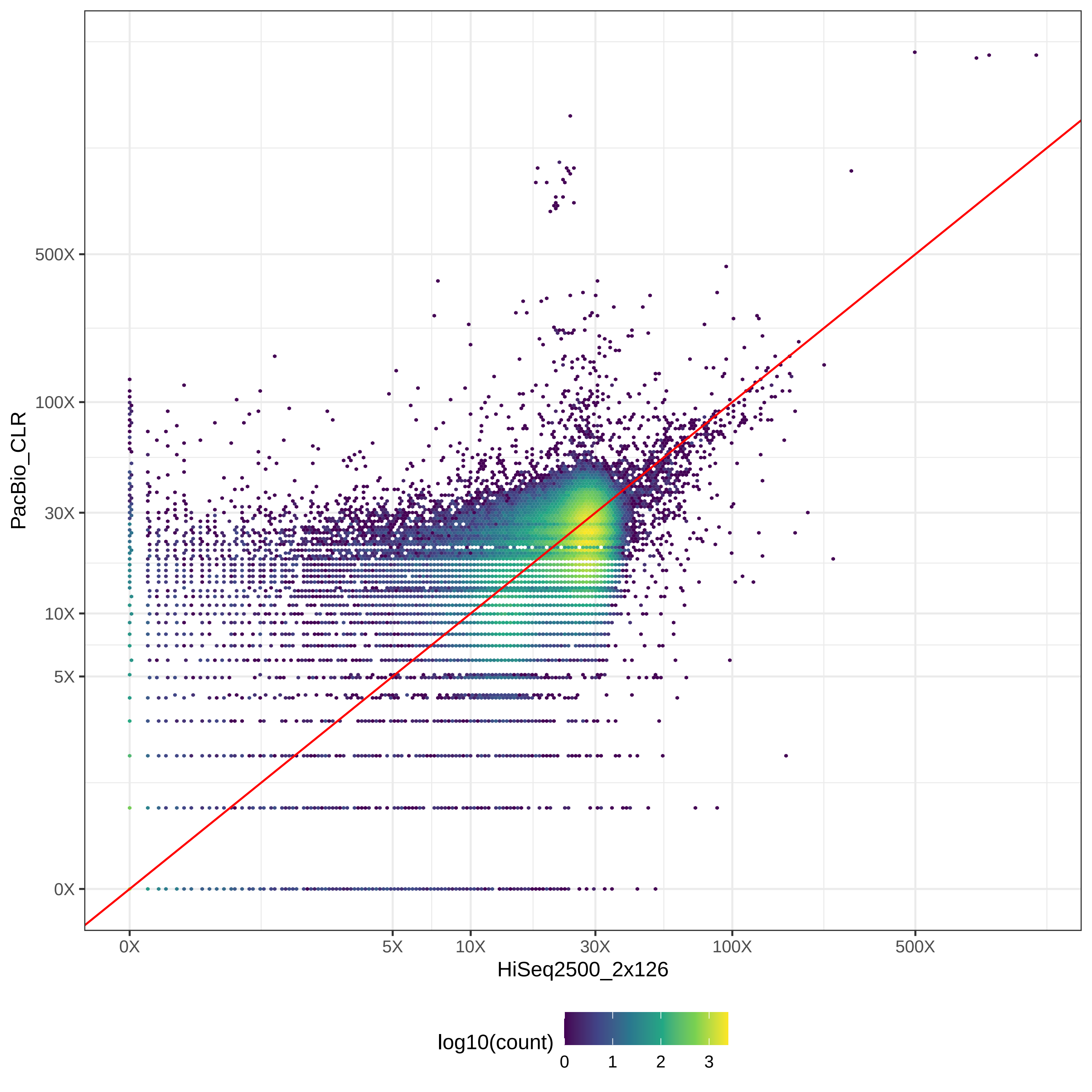

### GIAB_Low_Complexity_HiSeq2500_2x126-v-PromethION_μ=8092_hex.pdf

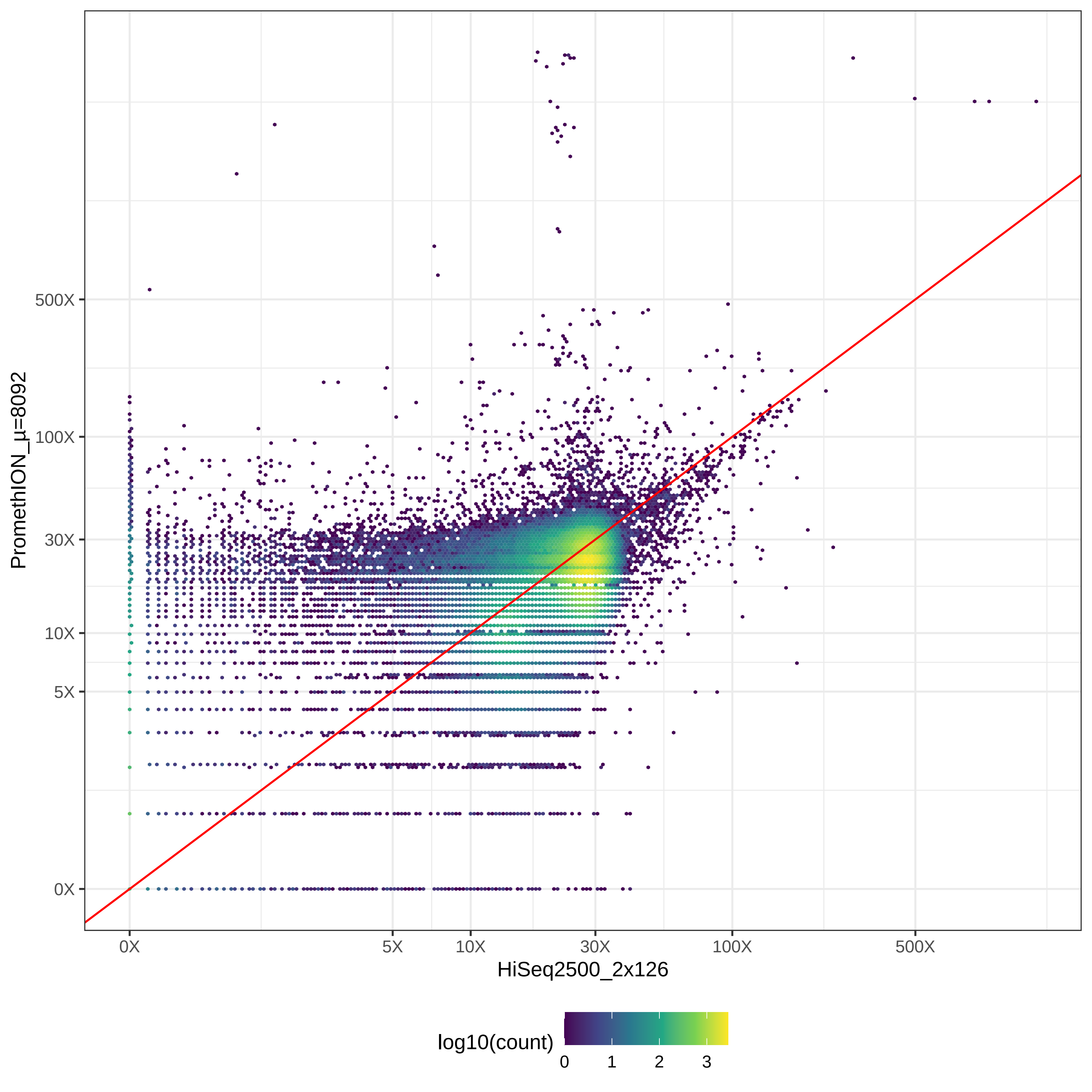

### GIAB_Low_Complexity_HiSeq4000_2x150-v-HiSeqX10_2x150_hex.pdf

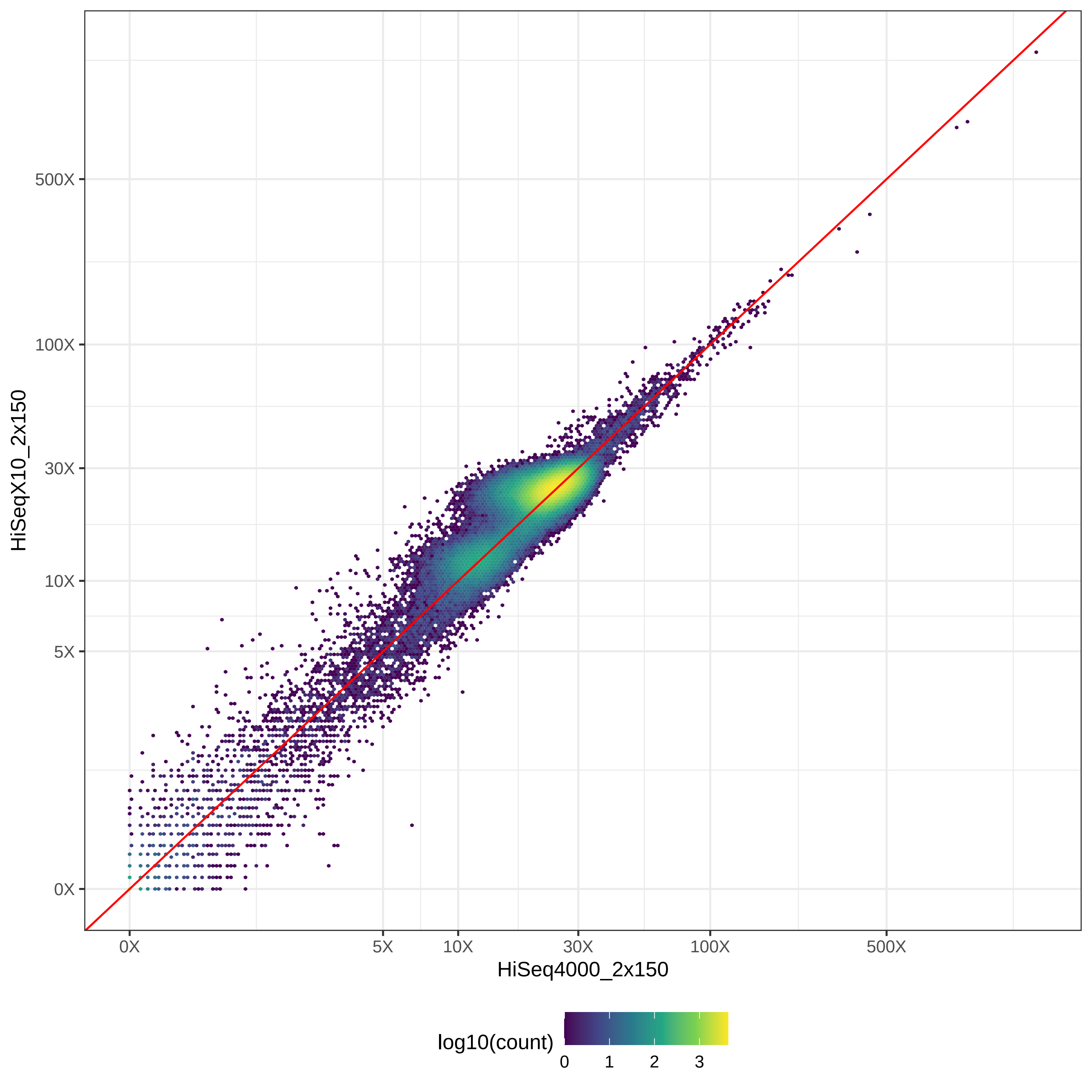

### GIAB_Low_Complexity_HiSeq4000_2x150-v-MGISEQ2000_2x150_hex.pdf

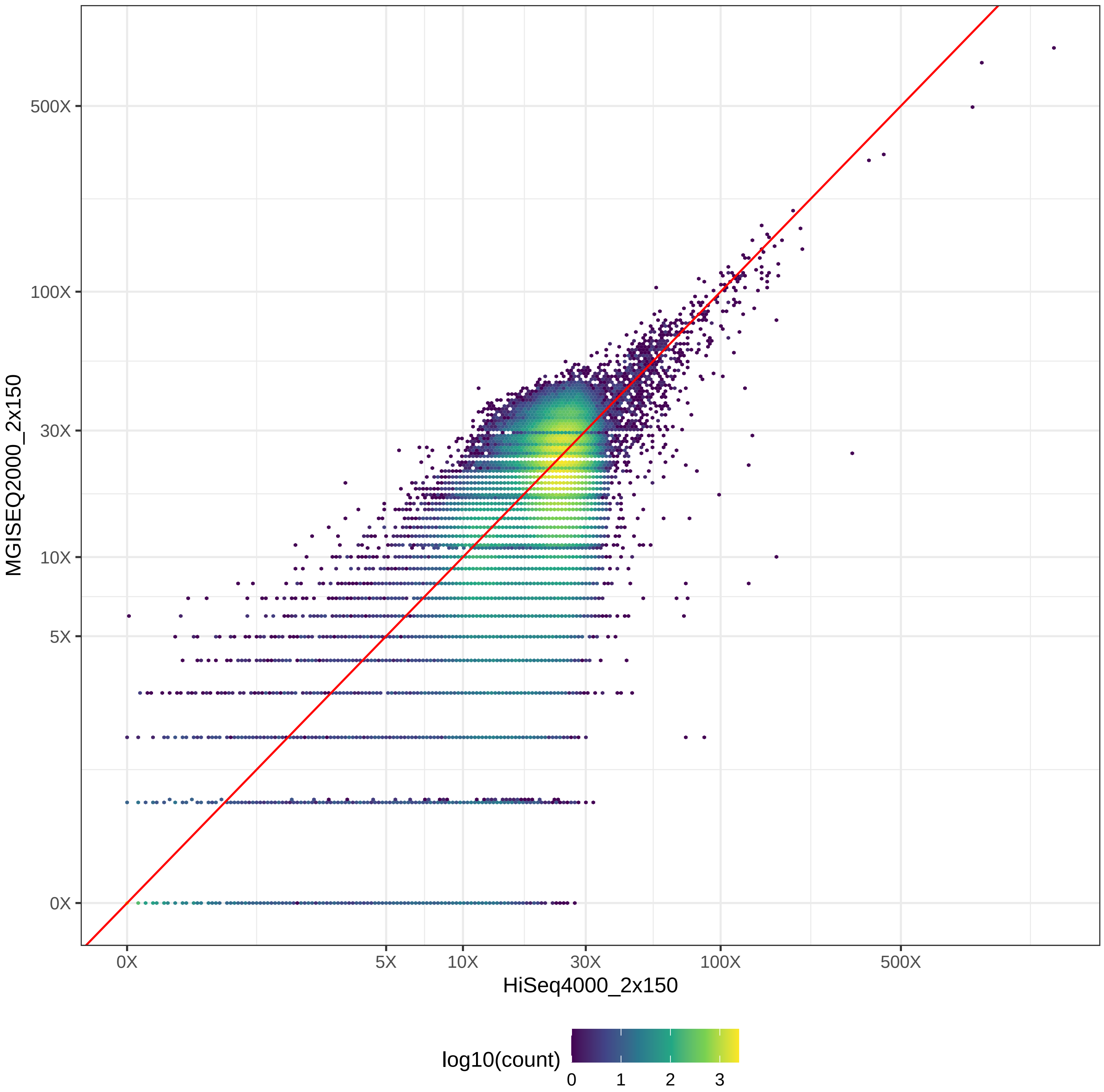

### GIAB_Low_Complexity_HiSeq4000_2x150-v-NovaSeq_2x150_hex.pdf

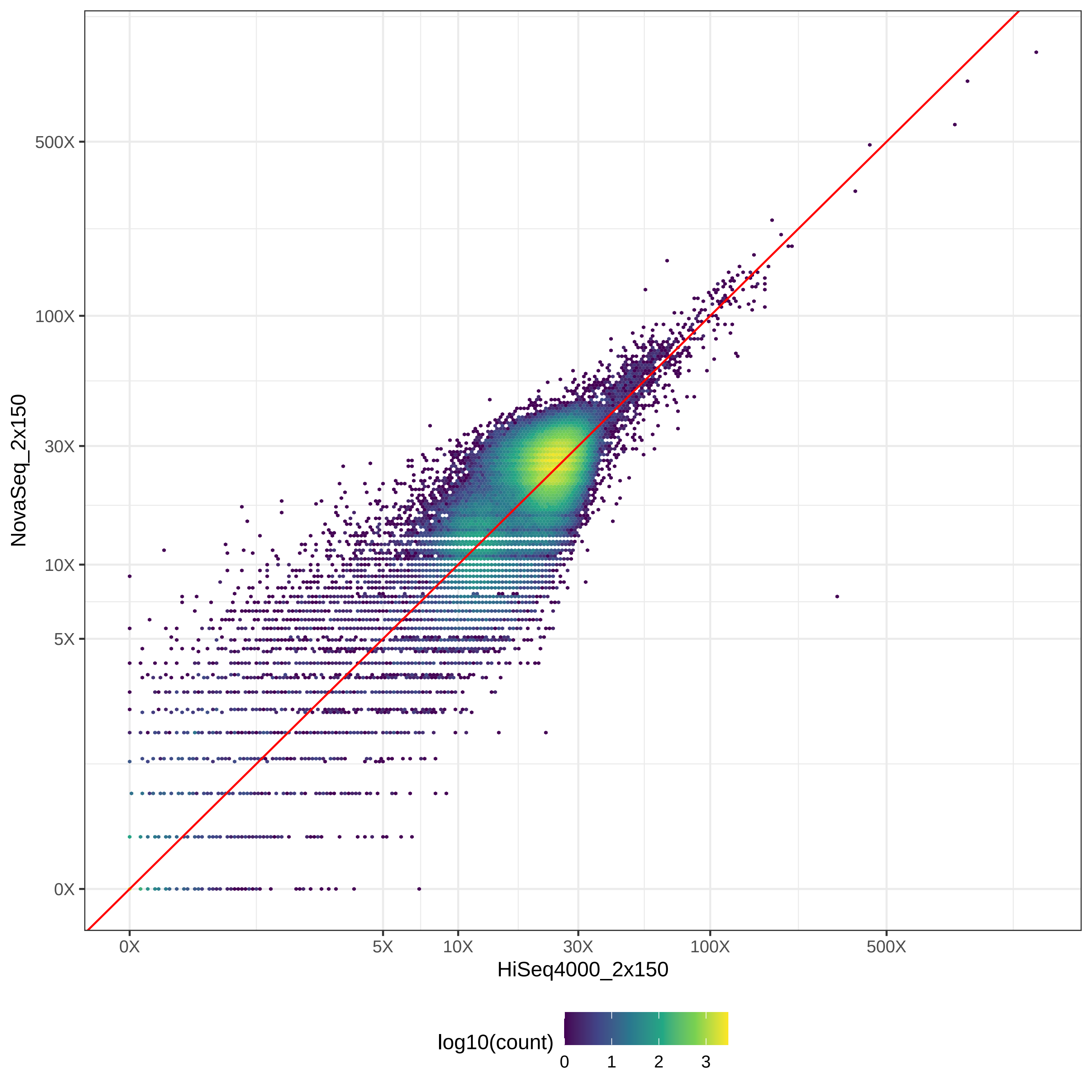

### GIAB_Low_Complexity_HiSeq4000_2x150-v-NovaSeq_2x250_hex.pdf

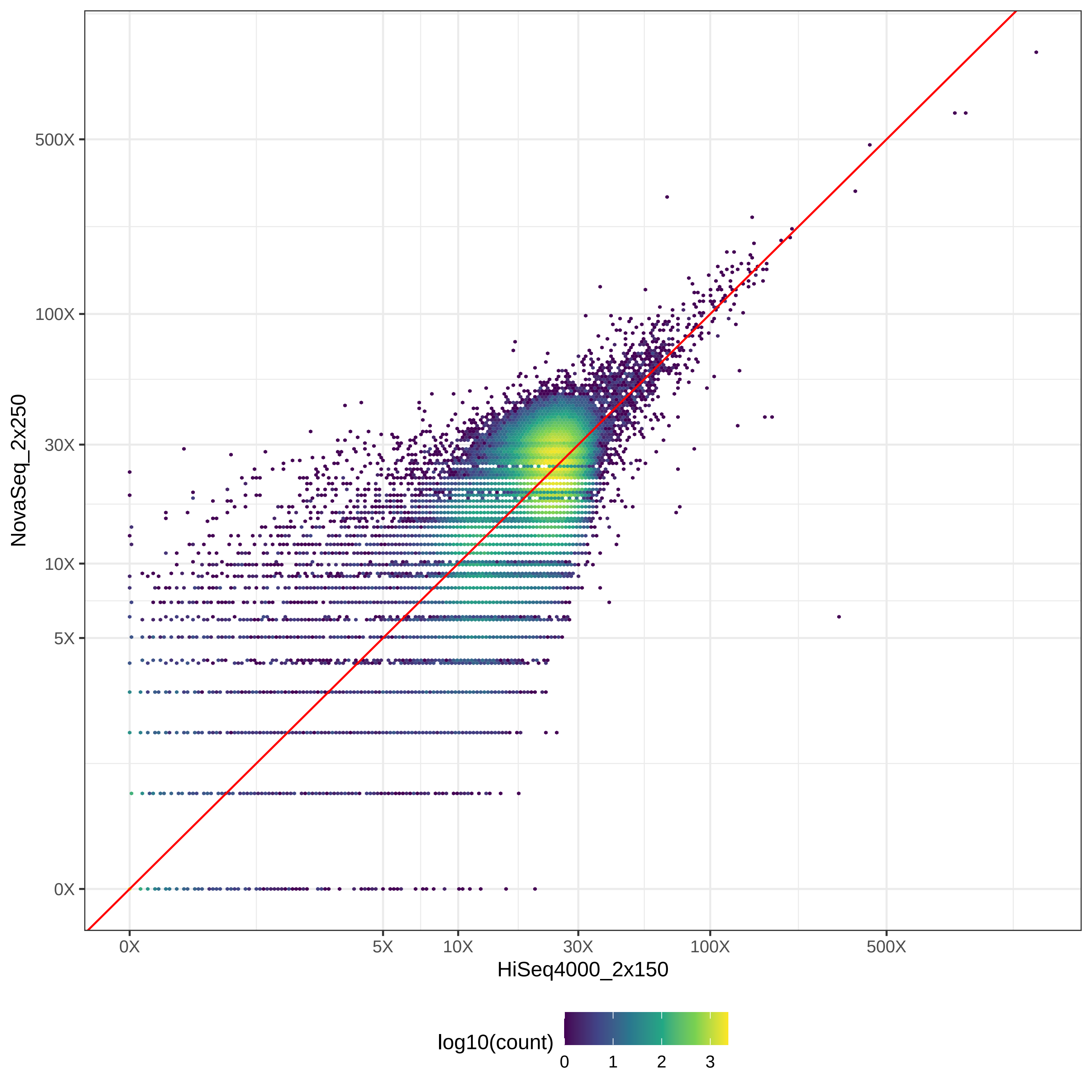

### GIAB_Low_Complexity_HiSeq4000_2x150-v-PacBio_CCS-15k_hex.pdf

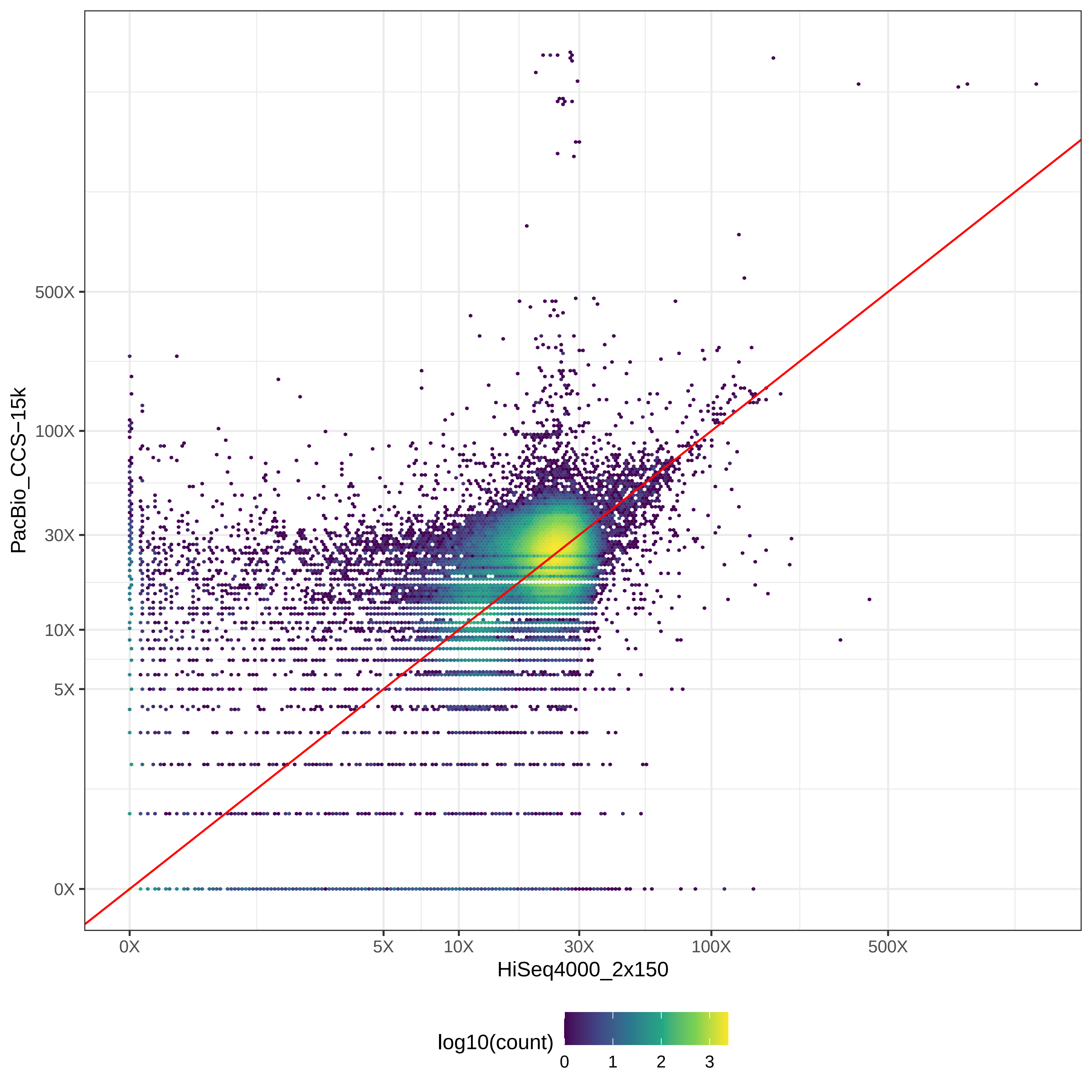

### GIAB_Low_Complexity_HiSeq4000_2x150-v-PacBio_CLR_hex.pdf

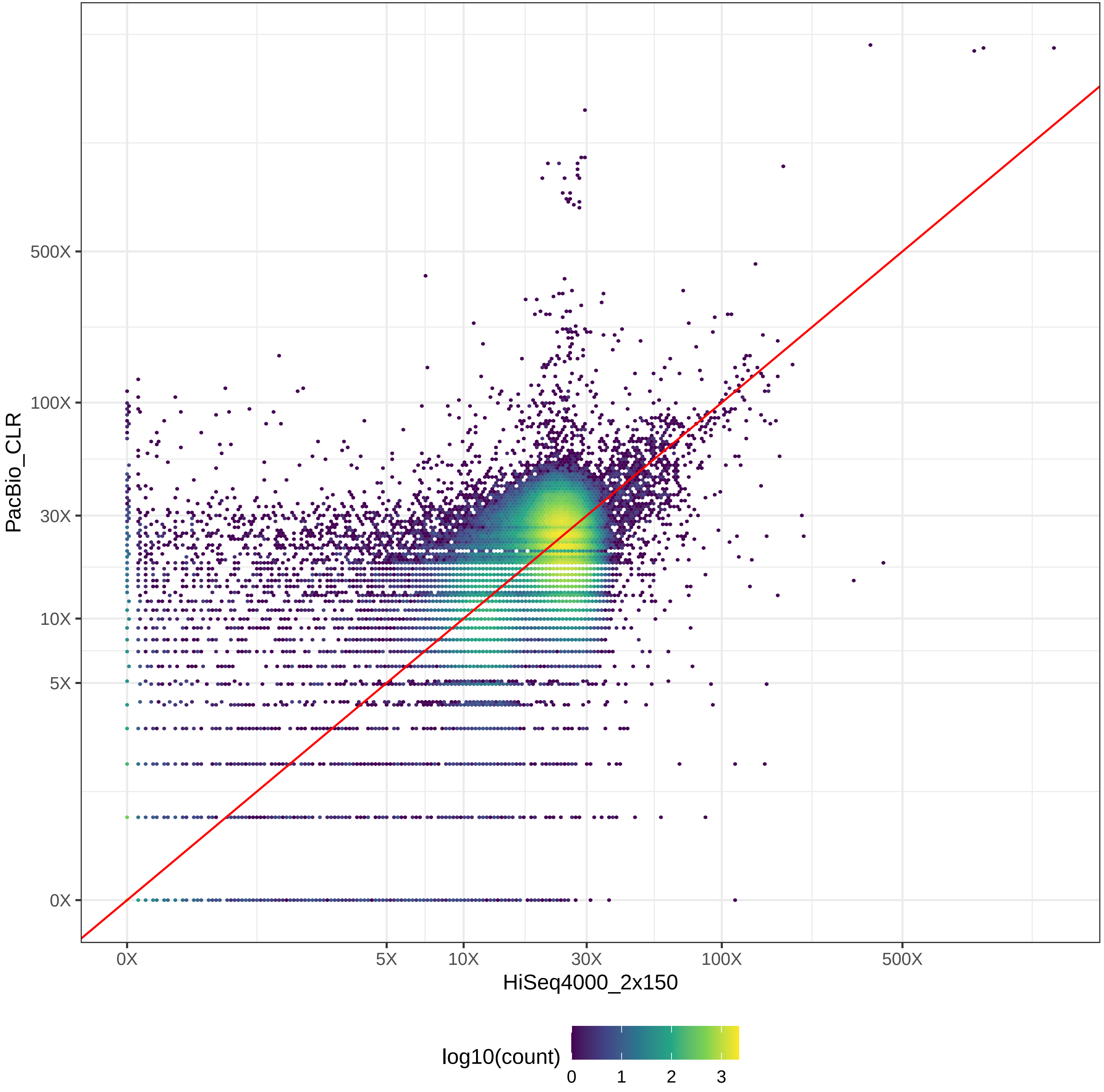

### GIAB_Low_Complexity_HiSeq4000_2x150-v-PromethION_μ=8092_hex.pdf

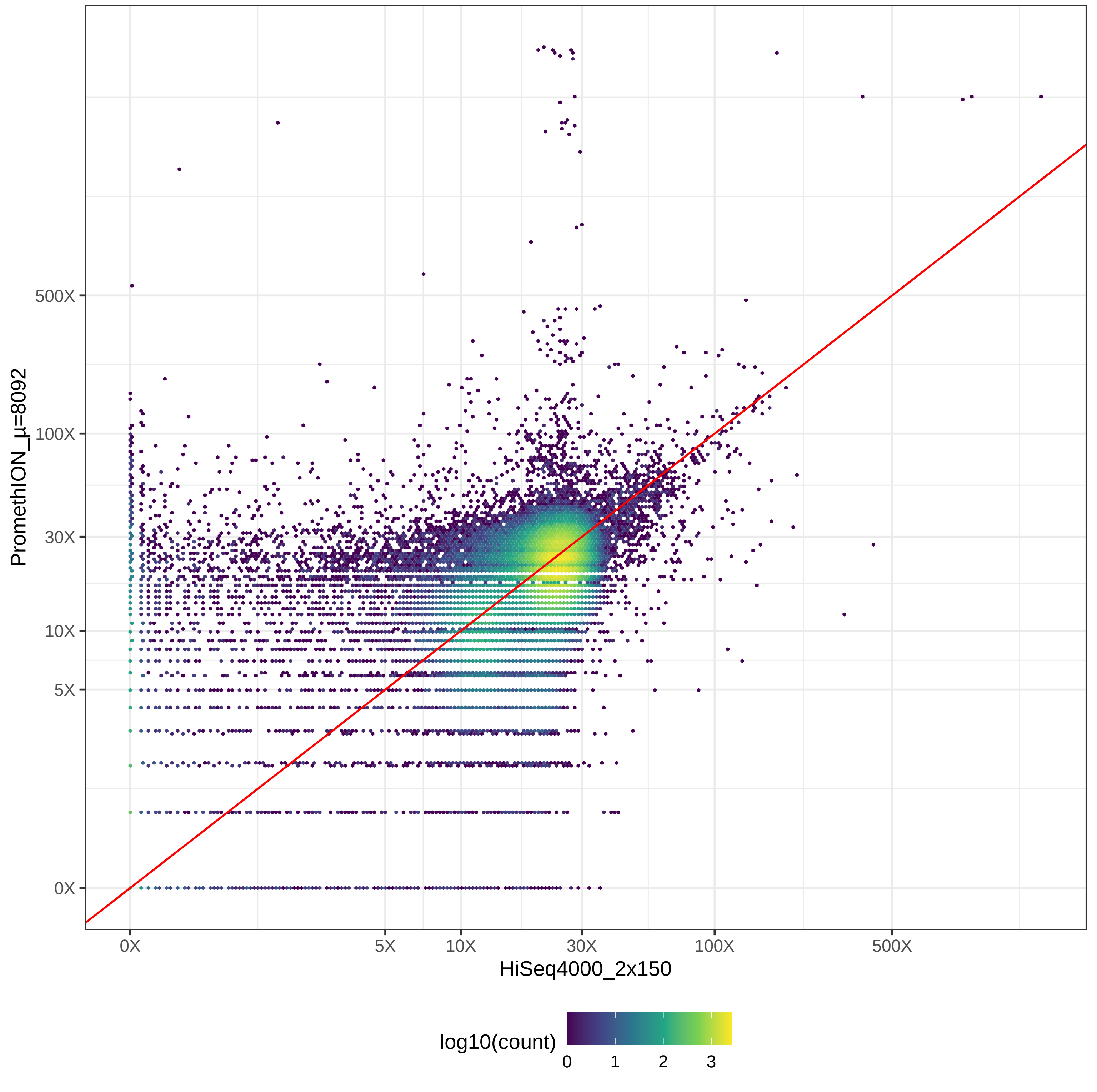

### GIAB_Low_Complexity_HiSeqX10_2x150-v-MGISEQ2000_2x150_hex.pdf

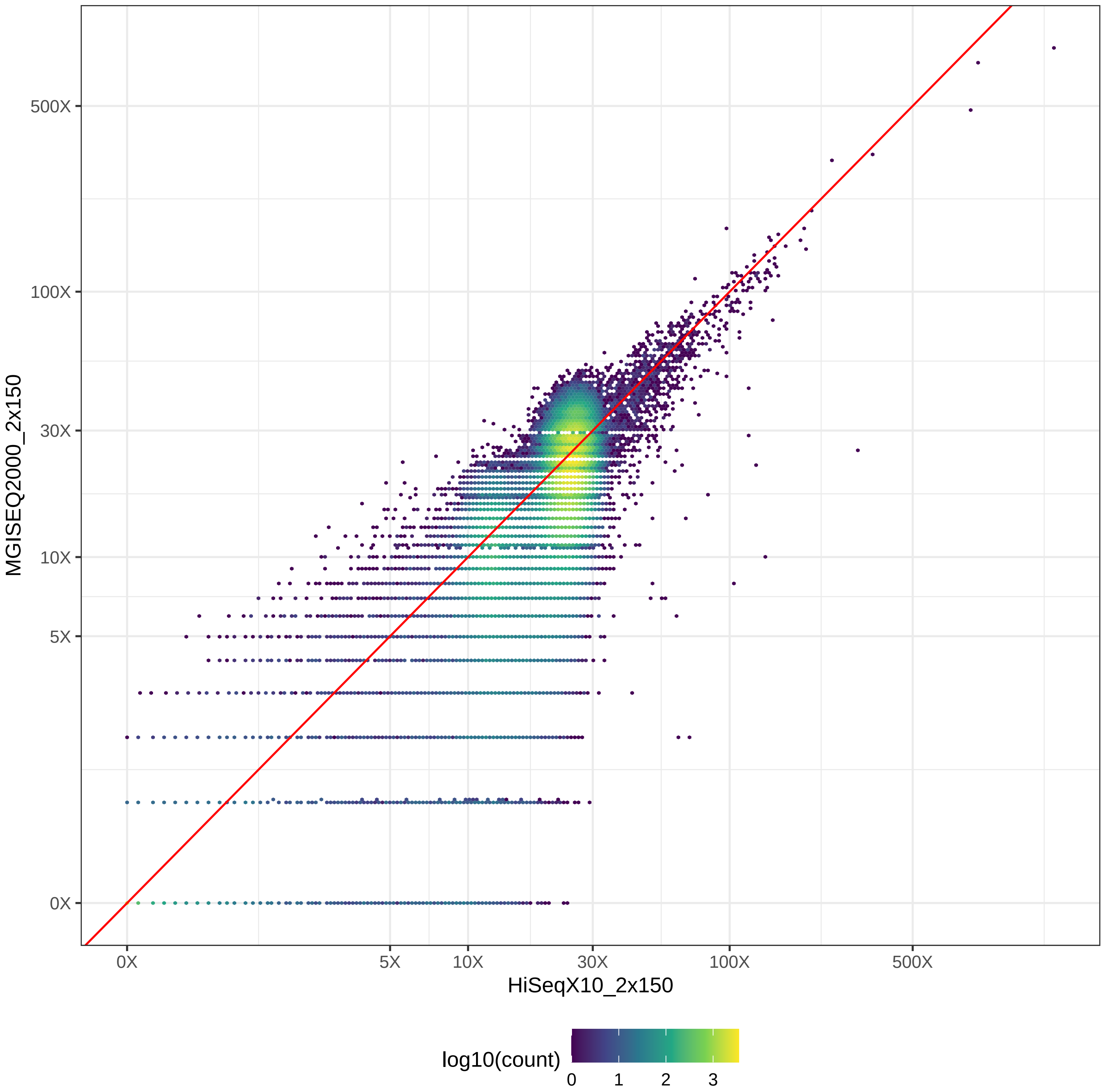
