## Supplementary Data 2 for "Multi-Platform Assessment of DNA Sequencing Performance using Human and Bacterial Reference Genomes in the ABRF Next-Generation Sequencing Study"

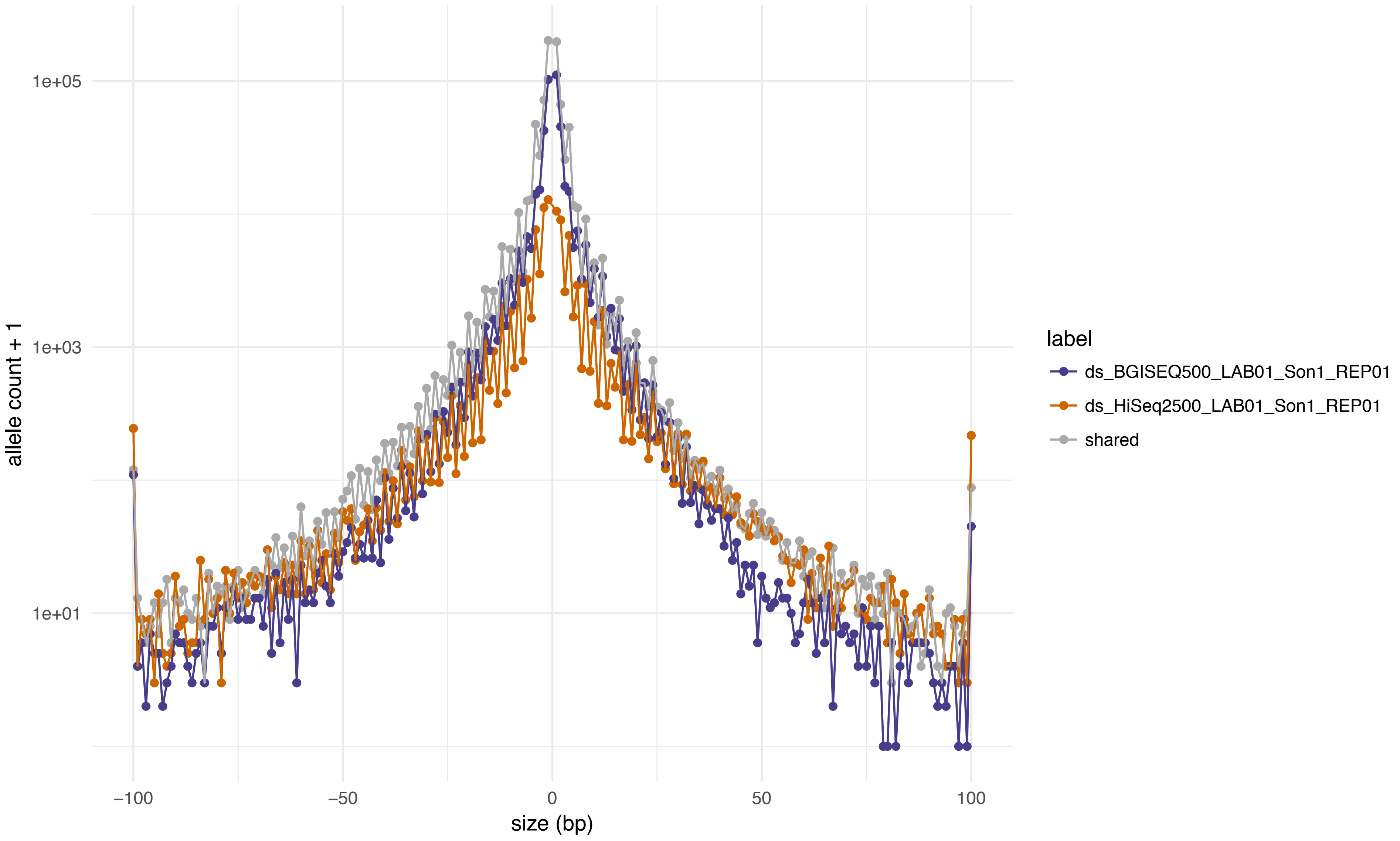
1e+05
1e+03
1e+01
allele count + 1
−100
−50
0
50
100
size (bp)
label
ds_BGISEQ500_LAB01_Son1_REP01
ds_HiSeq2500_LAB01_Son1_REP01
shared

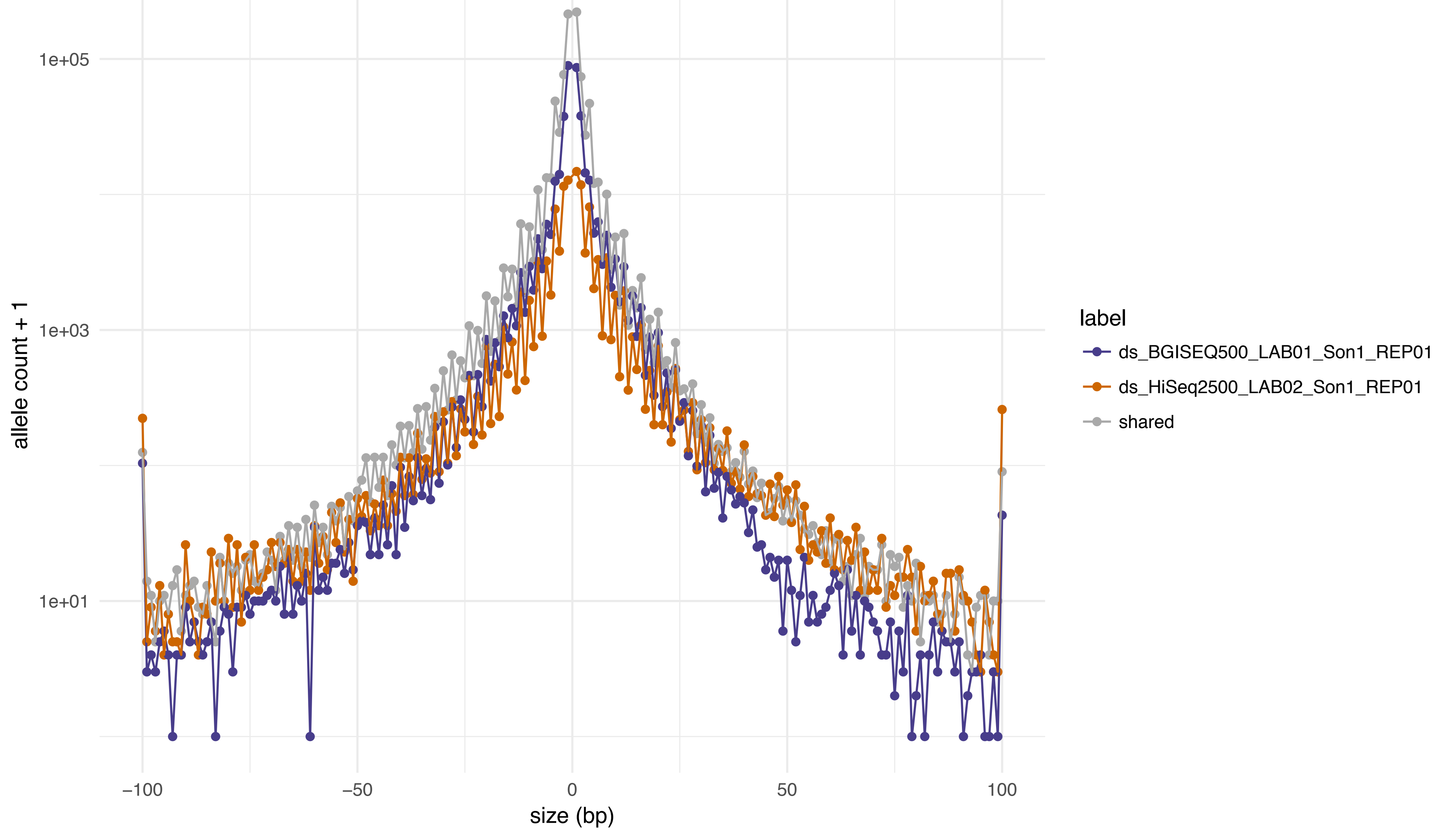
1e+05
1e+03
1e+01
allele count + 1
−100
−50
0
50
100
size (bp)
label
ds_BGISEQ500_LAB01_Son1_REP01
ds_HiSeq2500_LAB02_Son1_REP01
shared

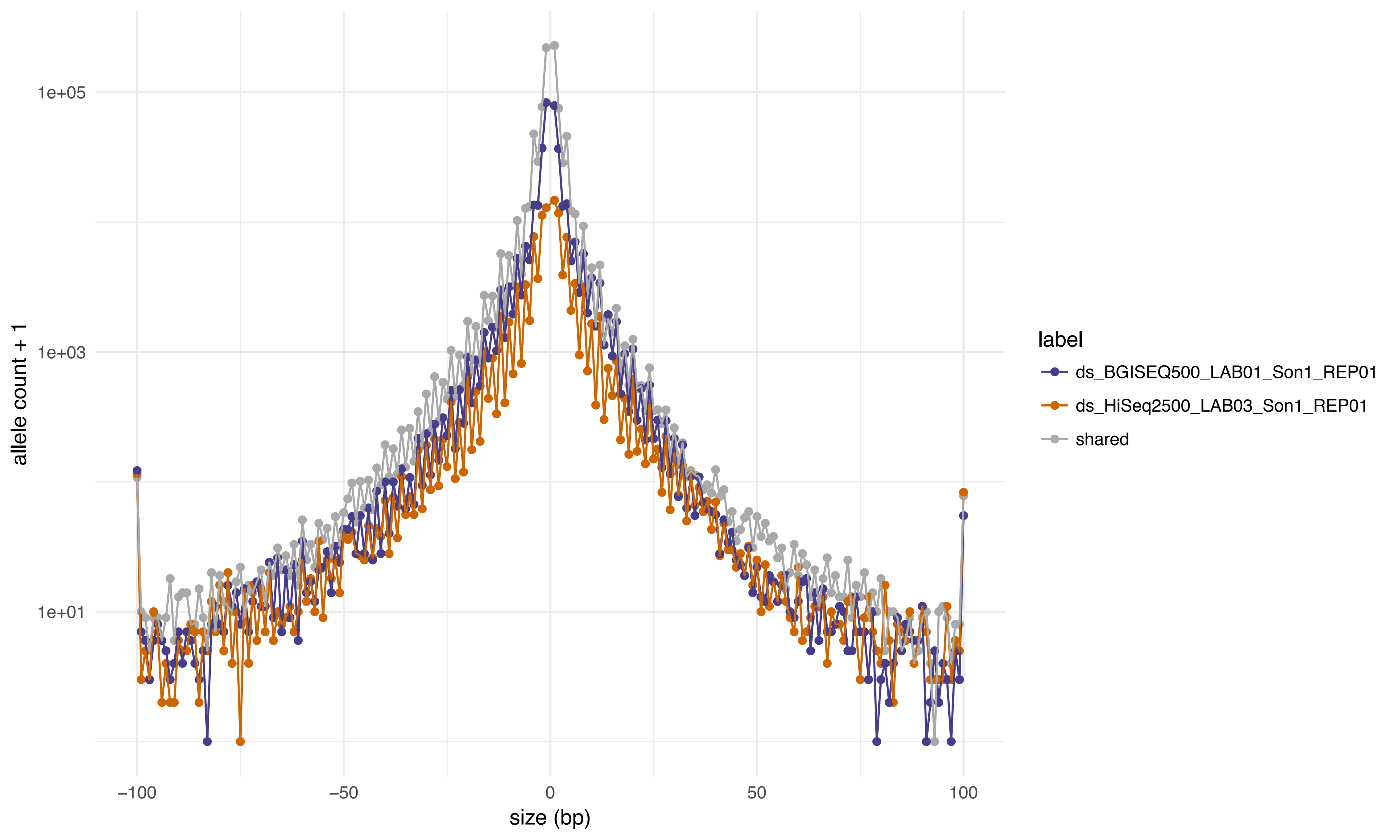
allele count + 1
1e+05
1e+03
1e+01
−100
−50
0
50
100
size (bp)
label
ds_BGISEQ500_LAB01_Son1_REP01
ds_HiSeq2500_LAB03_Son1_REP01
shared

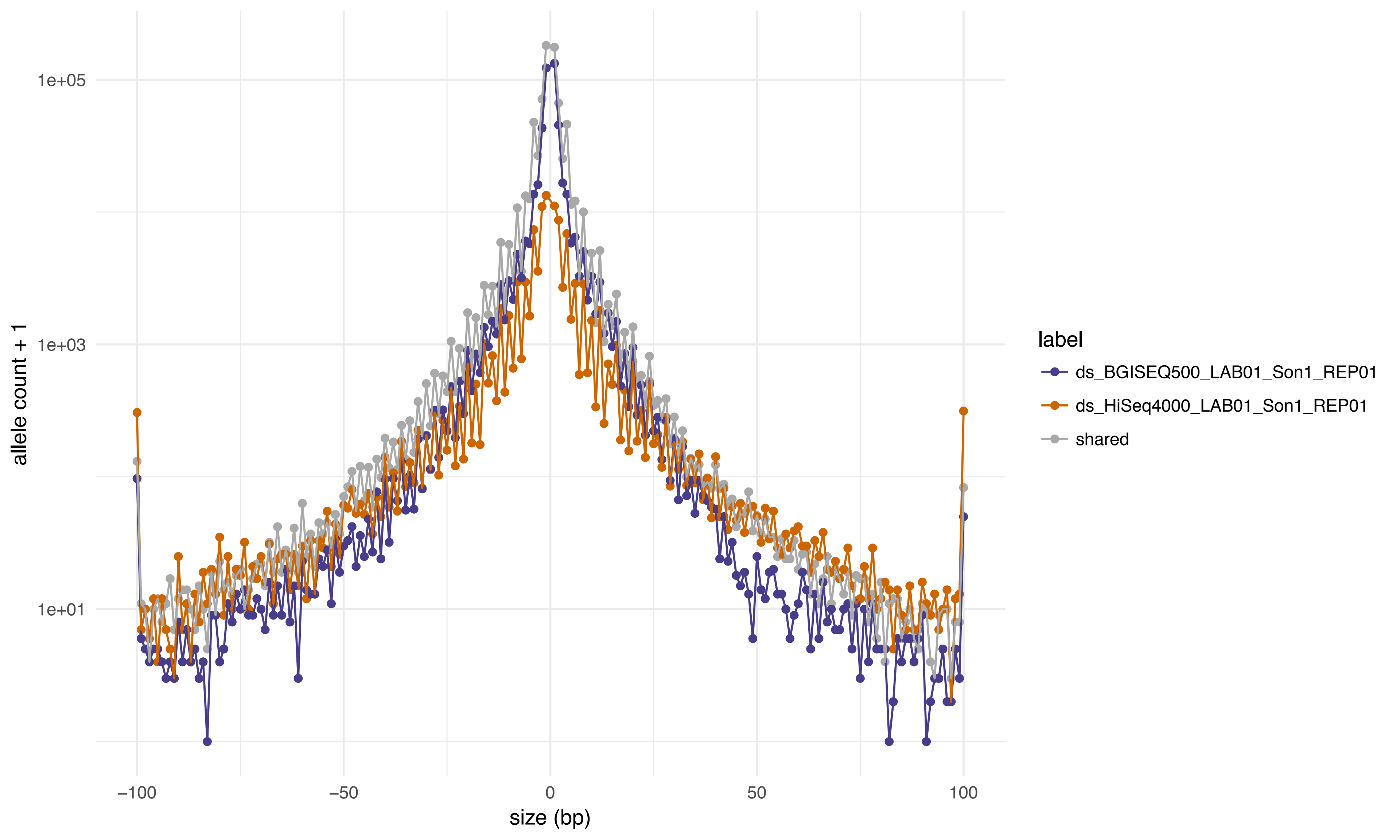

1e+05
1e+03
1e+01
allele count + 1
−100
−50
0
50
100
size (bp)
label
ds_BGISEQ500_LAB01_Son1_REP01
ds_HiSeq4000_LAB01_Son1_REP01
shared

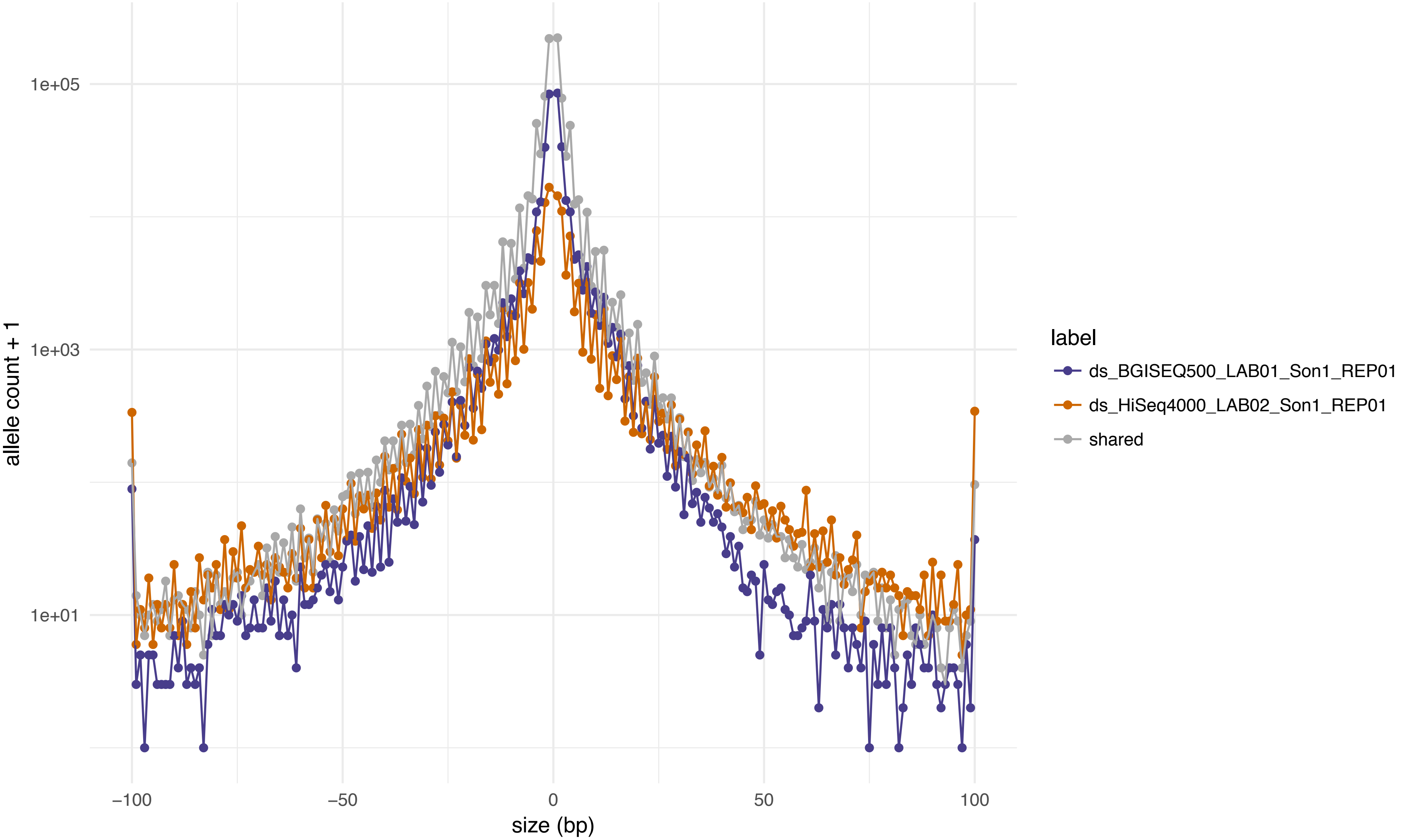
1e+05
1e+03
1e+01
allele count + 1
−100
−50
0
50
100
size (bp)
label
ds_BGISEQ500_LAB01_Son1_REP01
ds_HiSeq4000_LAB02_Son1_REP01
shared

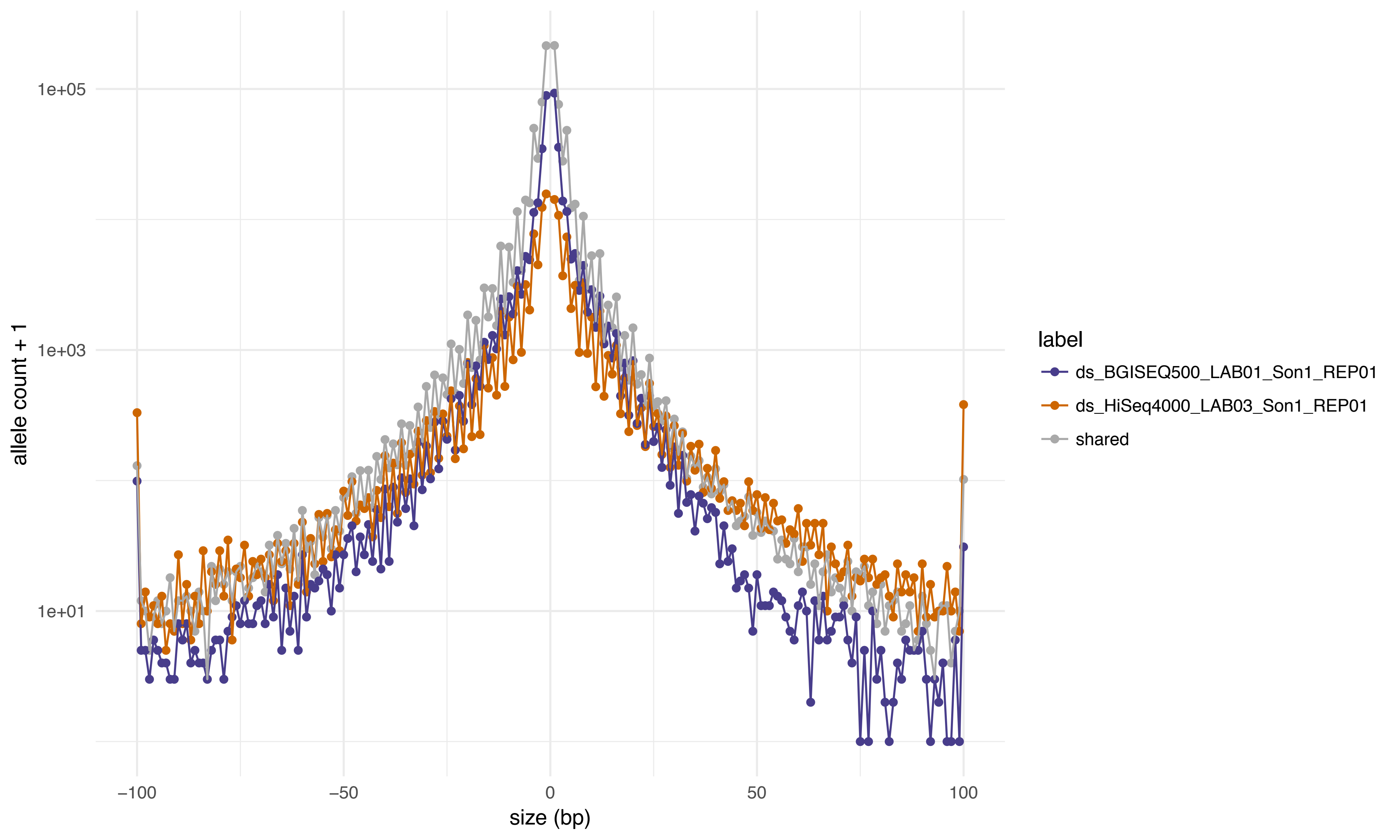

1e+05
1e+03
1e+01
allele count + 1
−100
−50
0
50
100
size (bp)
label
ds_BGISEQ500_LAB01_Son1_REP01
ds_HiSeq4000_LAB03_Son1_REP01
shared

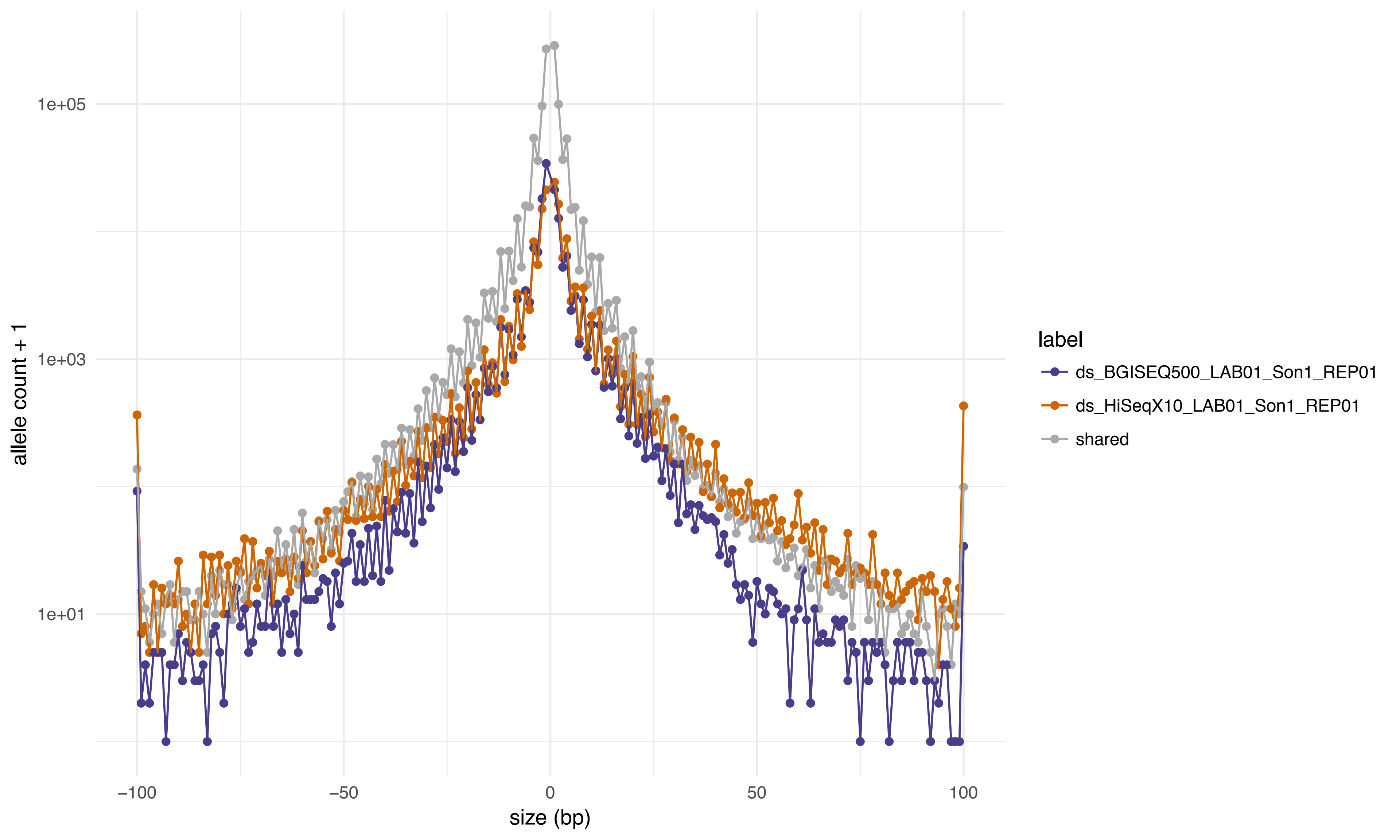

1e+05
1e+03
1e+01
allele count + 1
−100
−50
0
50
100
size (bp)
label
ds_BGISEQ500_LAB01_Son1_REP01
ds_HiSeqX10_LAB01_Son1_REP01
shared

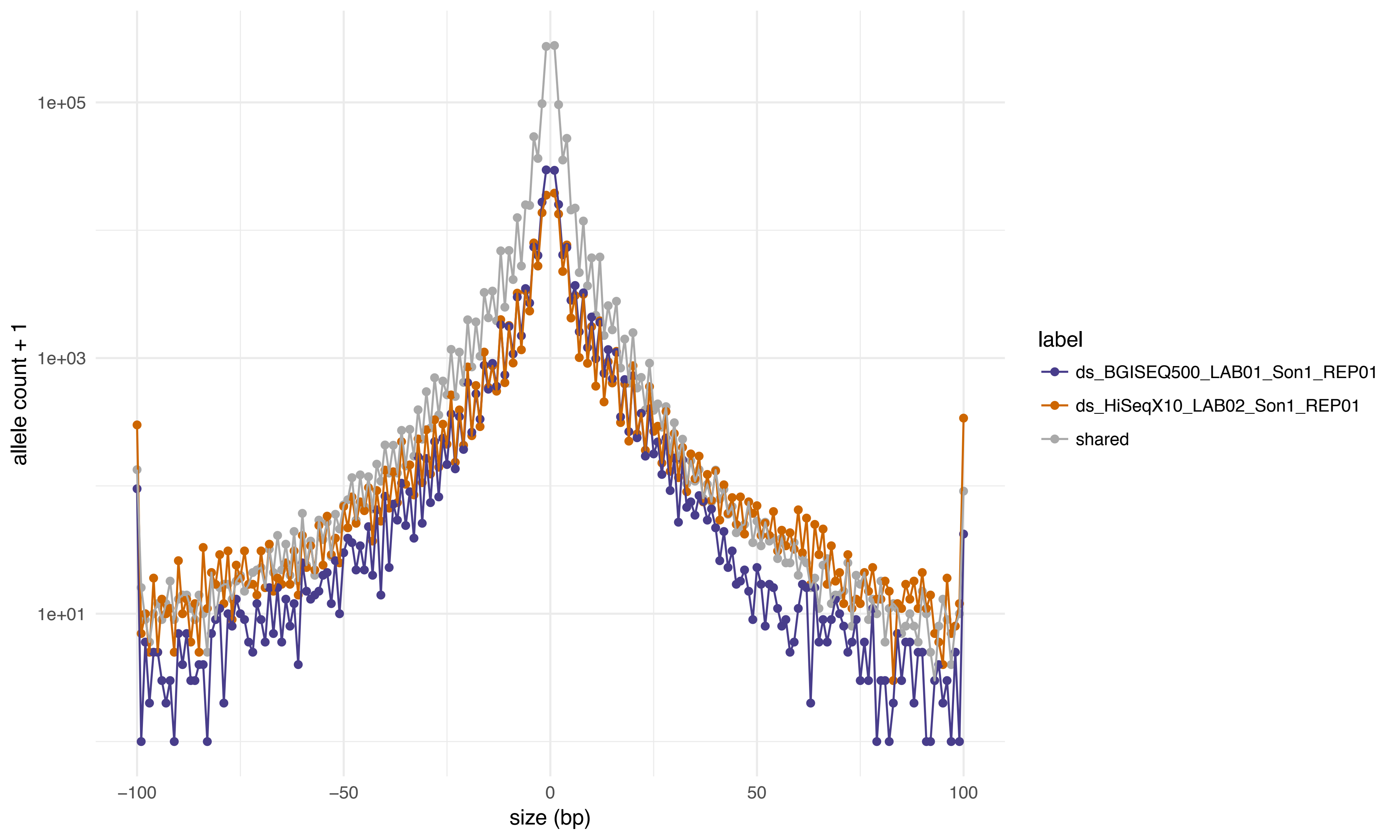

allele count + 1
1e+05
1e+03
1e+01
size (bp)
−100
−50
0
50
100
label
ds_BGISEQ500_LAB01_Son1_REP01
ds_HiSeqX10_LAB02_Son1_REP01
shared

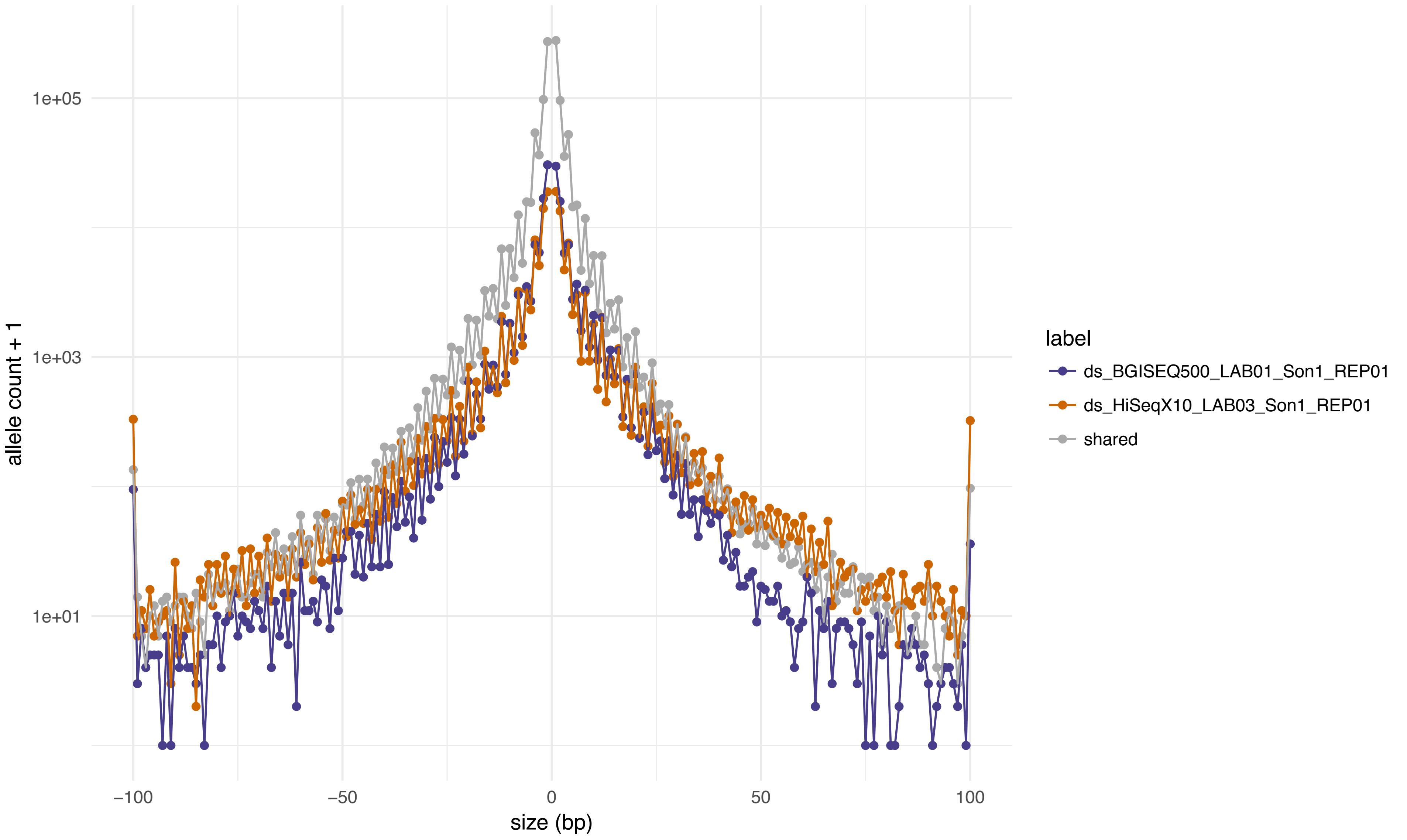

1e+05
1e+03
1e+01
allele count + 1
−100
−50
0
50
100
size (bp)
label
ds_BGISEQ500_LAB01_Son1_REP01
ds_HiSeqX10_LAB03_Son1_REP01
shared

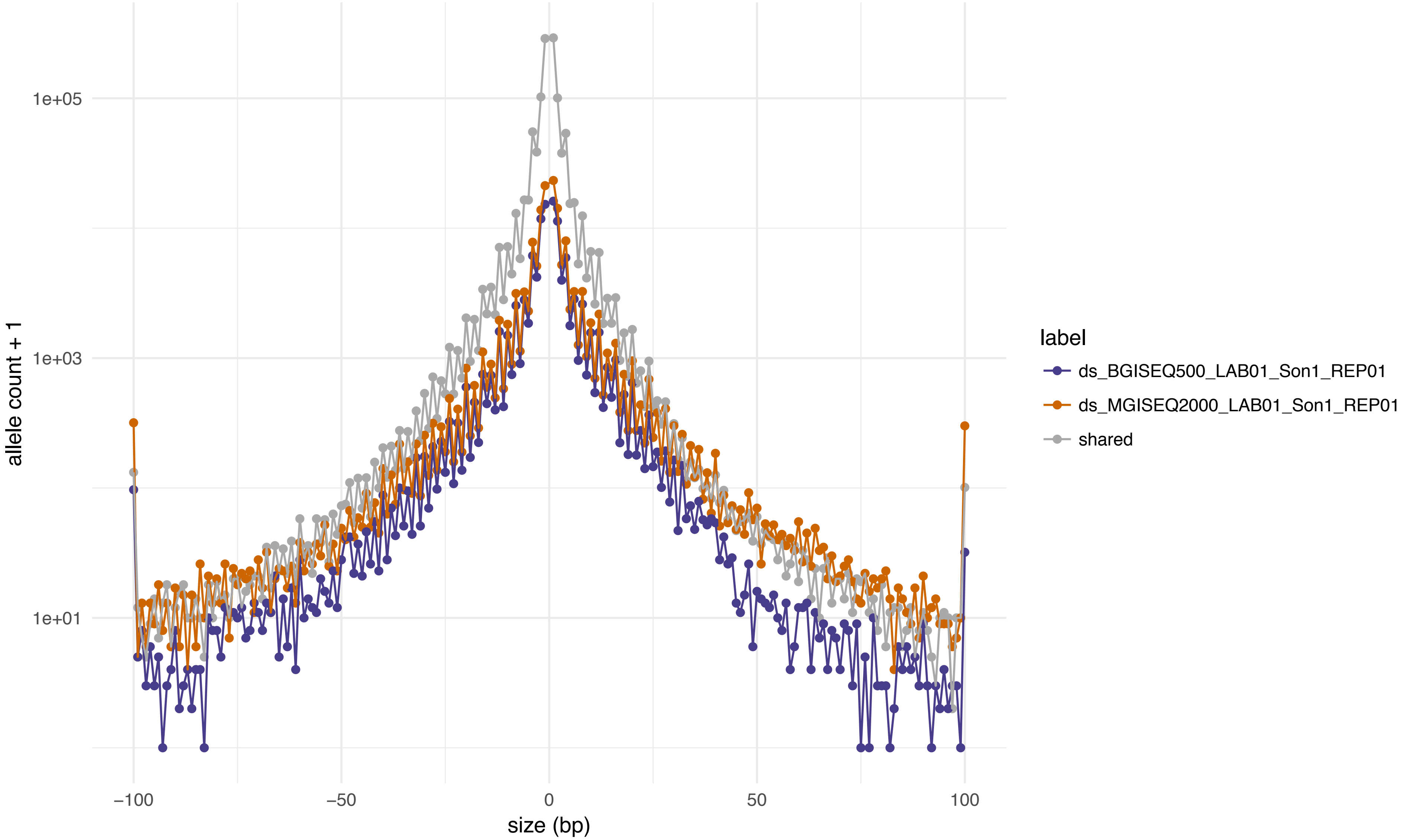
1e+05
1e+03
1e+01
allele count + 1
−100
−50
0
50
100
size (bp)
label
ds_BGISEQ500_LAB01_Son1_REP01
ds_MGISEQ2000_LAB01_Son1_REP01
shared

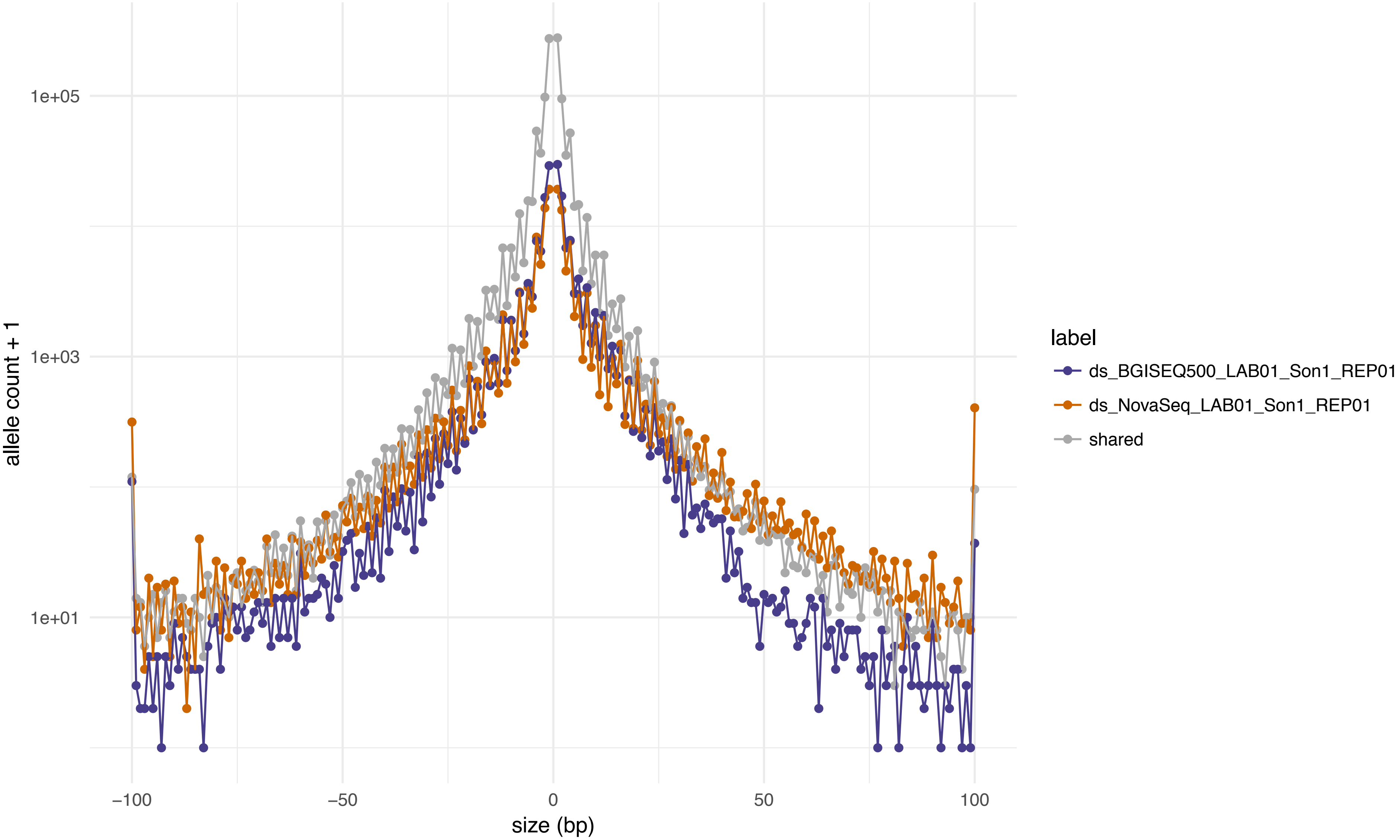

allele count + 1
1e+05
1e+03
1e+01
−100
−50
0
50
100
size (bp)
label
ds_BGISEQ500_LAB01_Son1_REP01
ds_NovaSeq_LAB01_Son1_REP01
shared

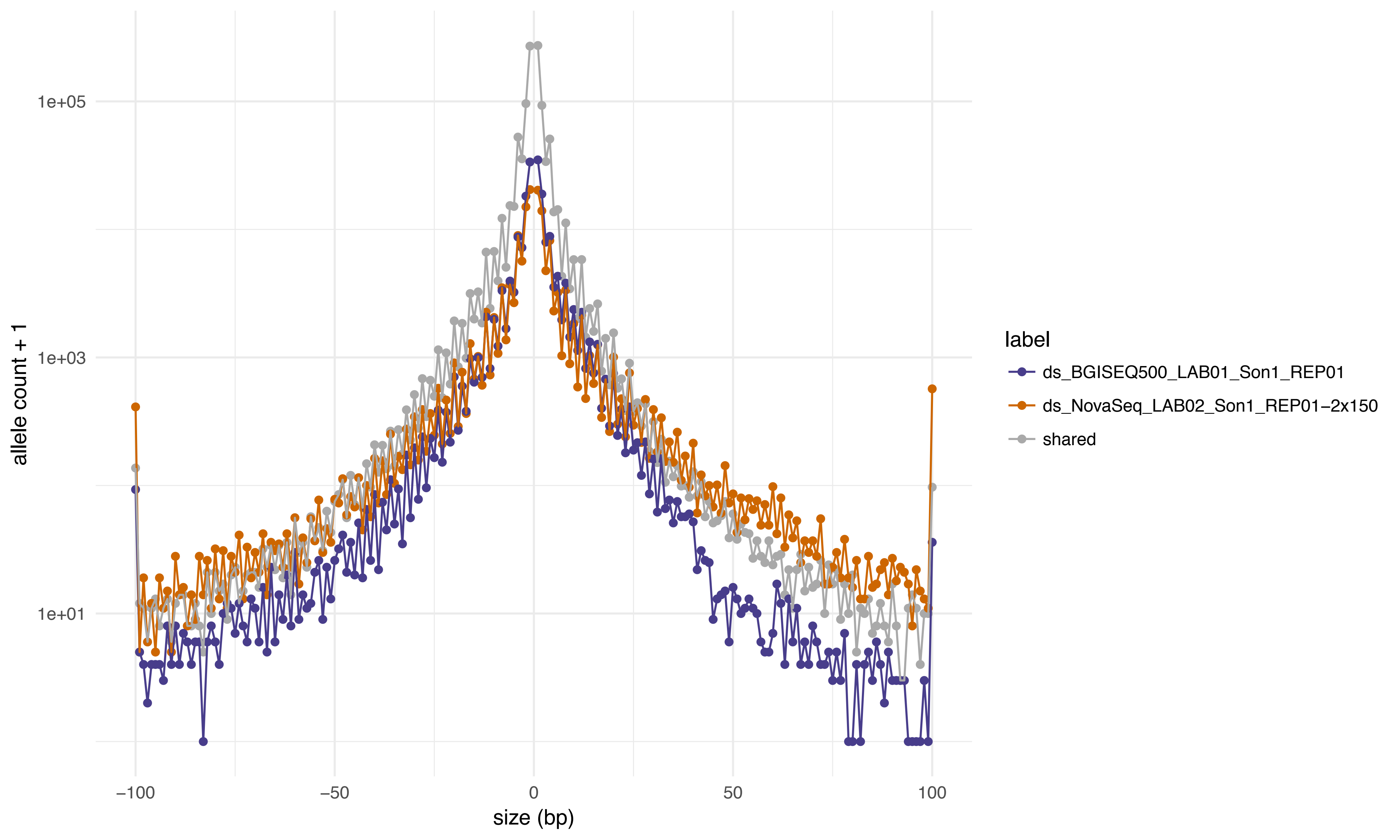

1e+05
1e+03
1e+01
allele count + 1
−100
−50
0
50
100
size (bp)
label
ds_BGISEQ500_LAB01_Son1_REP01
ds_NovaSeq_LAB02_Son1_REP01−2x150
shared

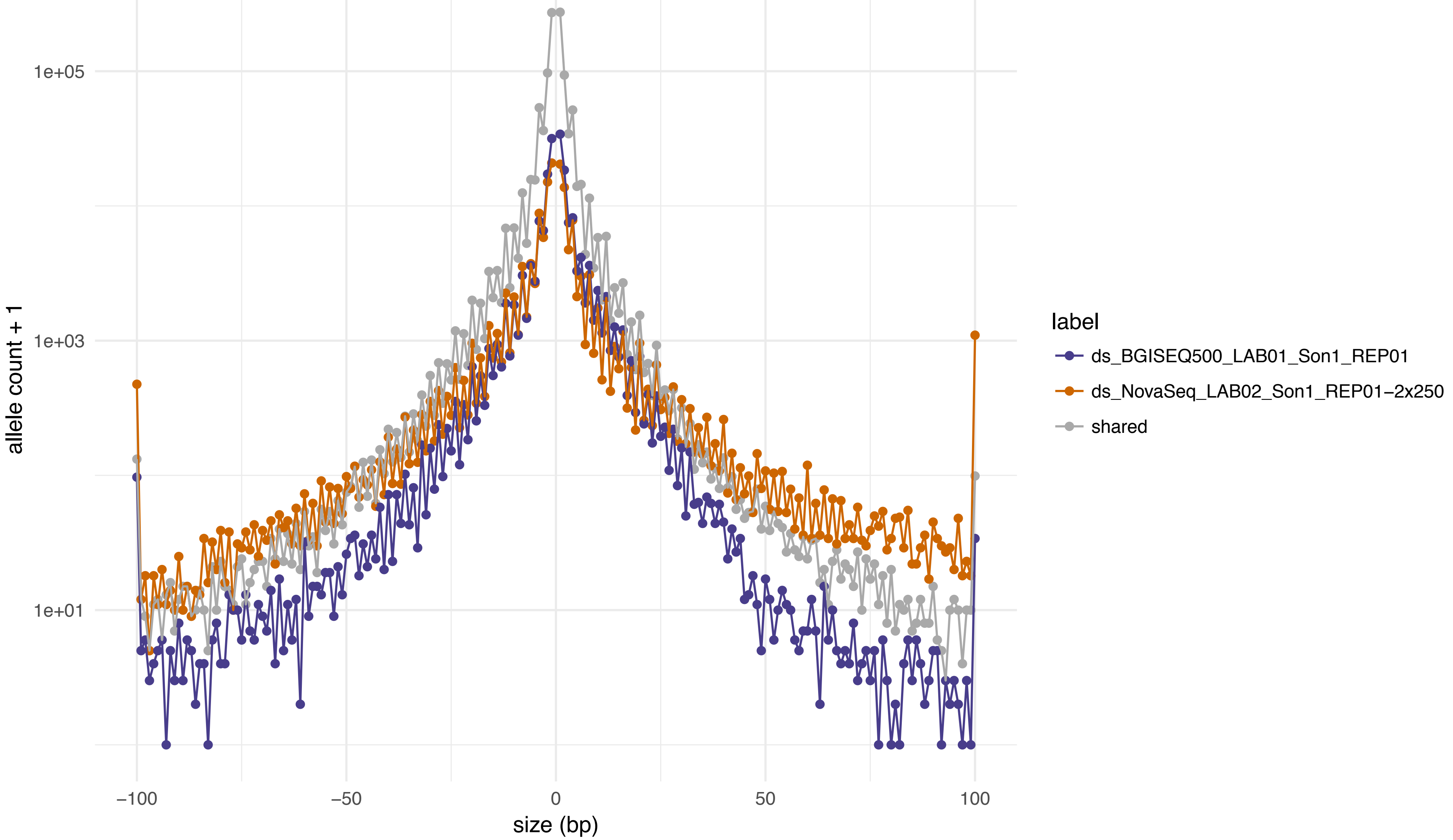
1e+05
1e+03
1e+01
allele count + 1
−100
−50
0
50
100
size (bp)
label
ds_BGISEQ500_LAB01_Son1_REP01
ds_NovaSeq_LAB02_Son1_REP01−2x250
shared

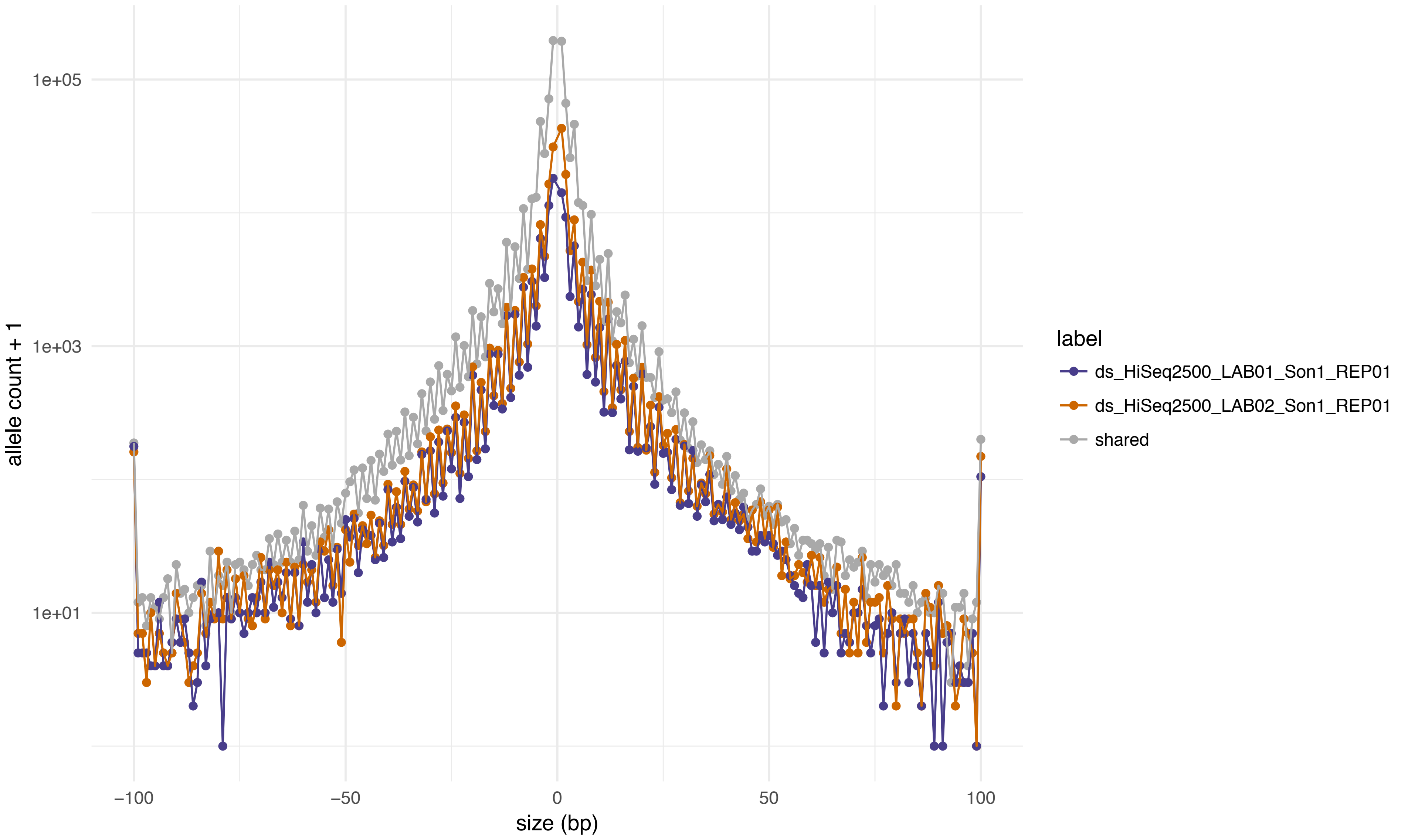

1e+05
1e+03
1e+01
allele count + 1
−100
−50
0
50
100
size (bp)
label
ds_HiSeq2500_LAB01_Son1_REP01
ds_HiSeq2500_LAB02_Son1_REP01
shared

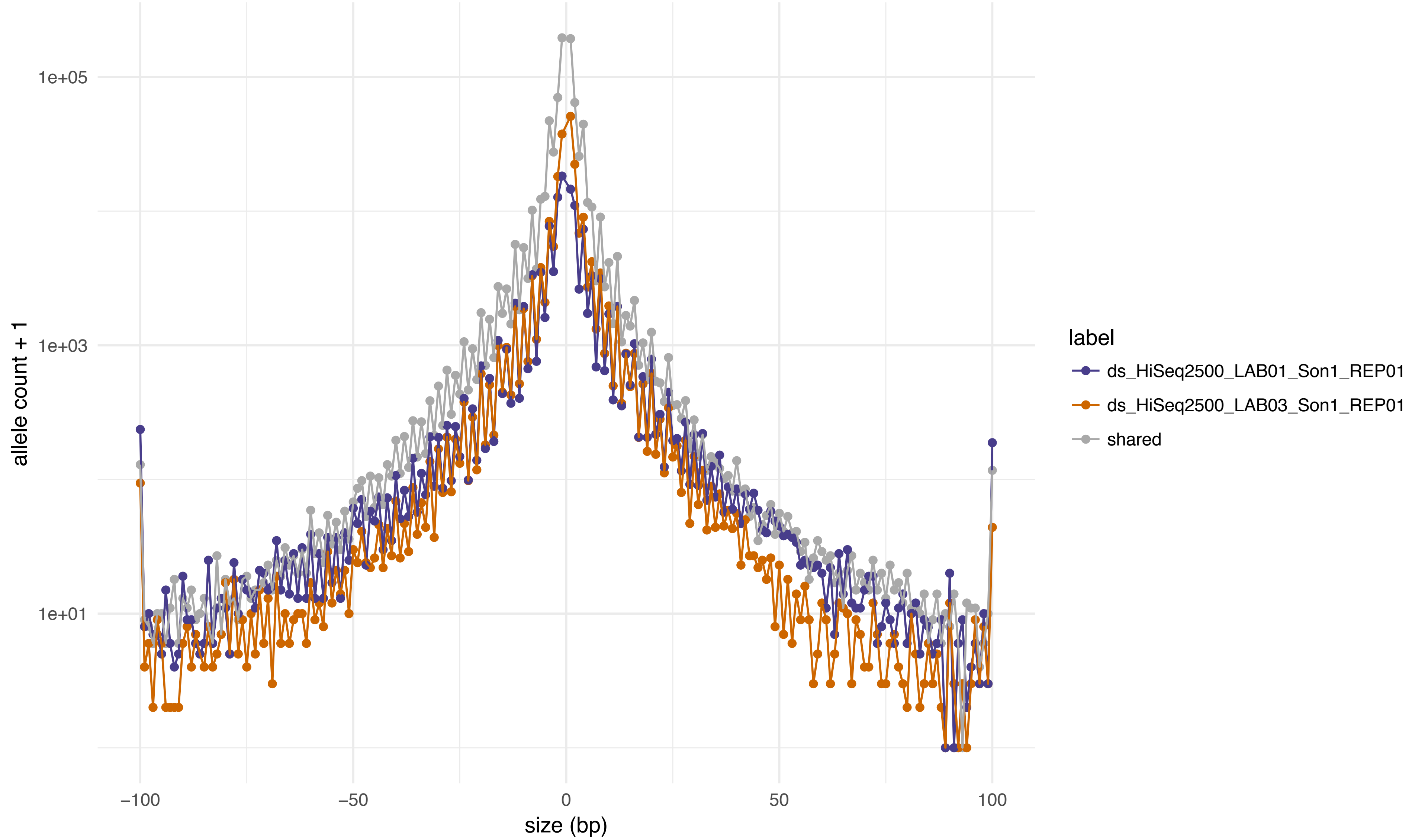

1e+05
1e+03
1e+01
allele count + 1
−100
−50
0
50
100
size (bp)
label
ds_HiSeq2500_LAB01_Son1_REP01
ds_HiSeq2500_LAB03_Son1_REP01
shared

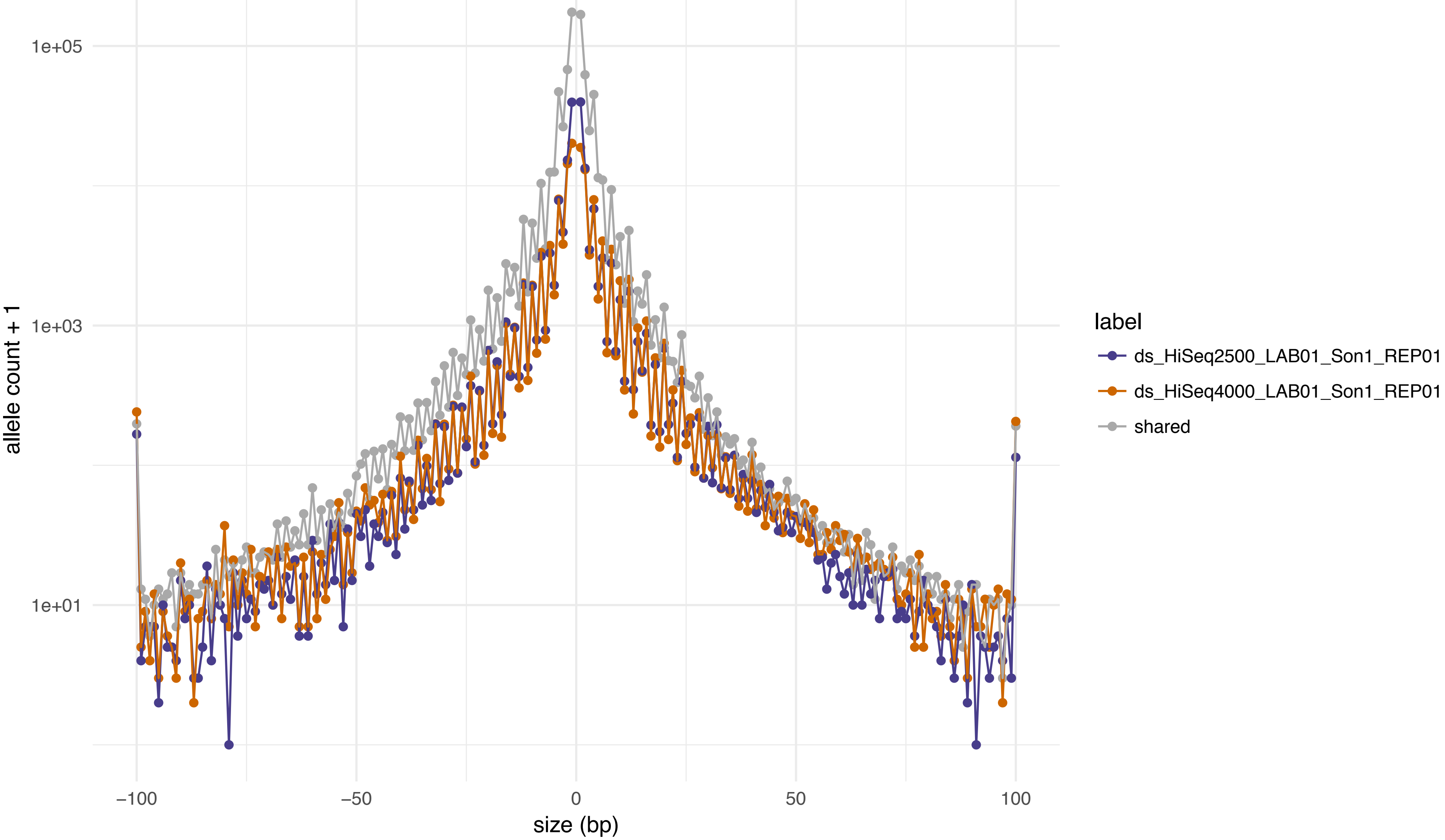

1e+05
1e+03
1e+01
allele count + 1
−100
−50
0
50
100
size (bp)
label
ds_HiSeq2500_LAB01_Son1_REP01
ds_HiSeq4000_LAB01_Son1_REP01
shared

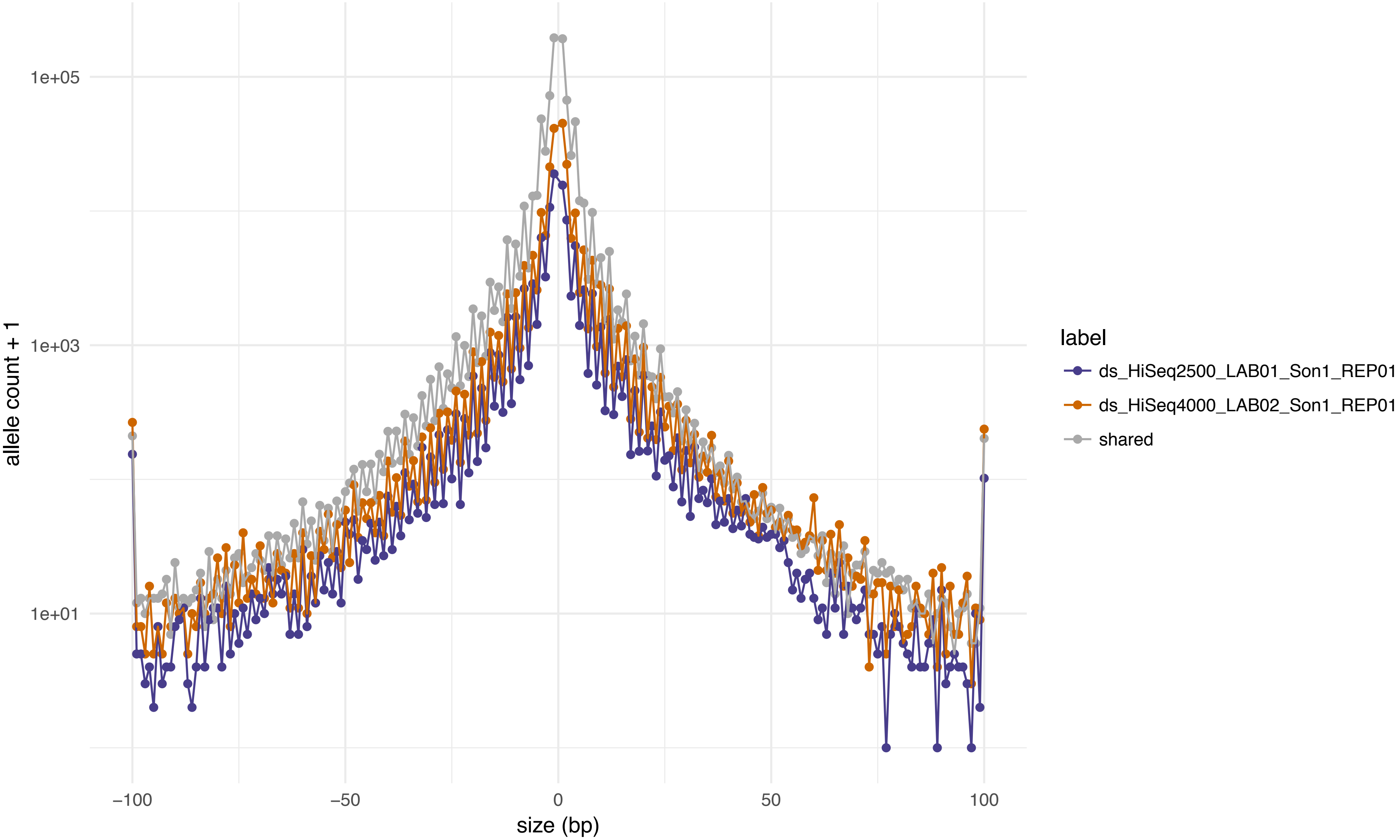
1e+05
1e+03
1e+01
allele count + 1
−100
−50
0
50
100
size (bp)
label
ds_HiSeq2500_LAB01_Son1_REP01
ds_HiSeq4000_LAB02_Son1_REP01
shared

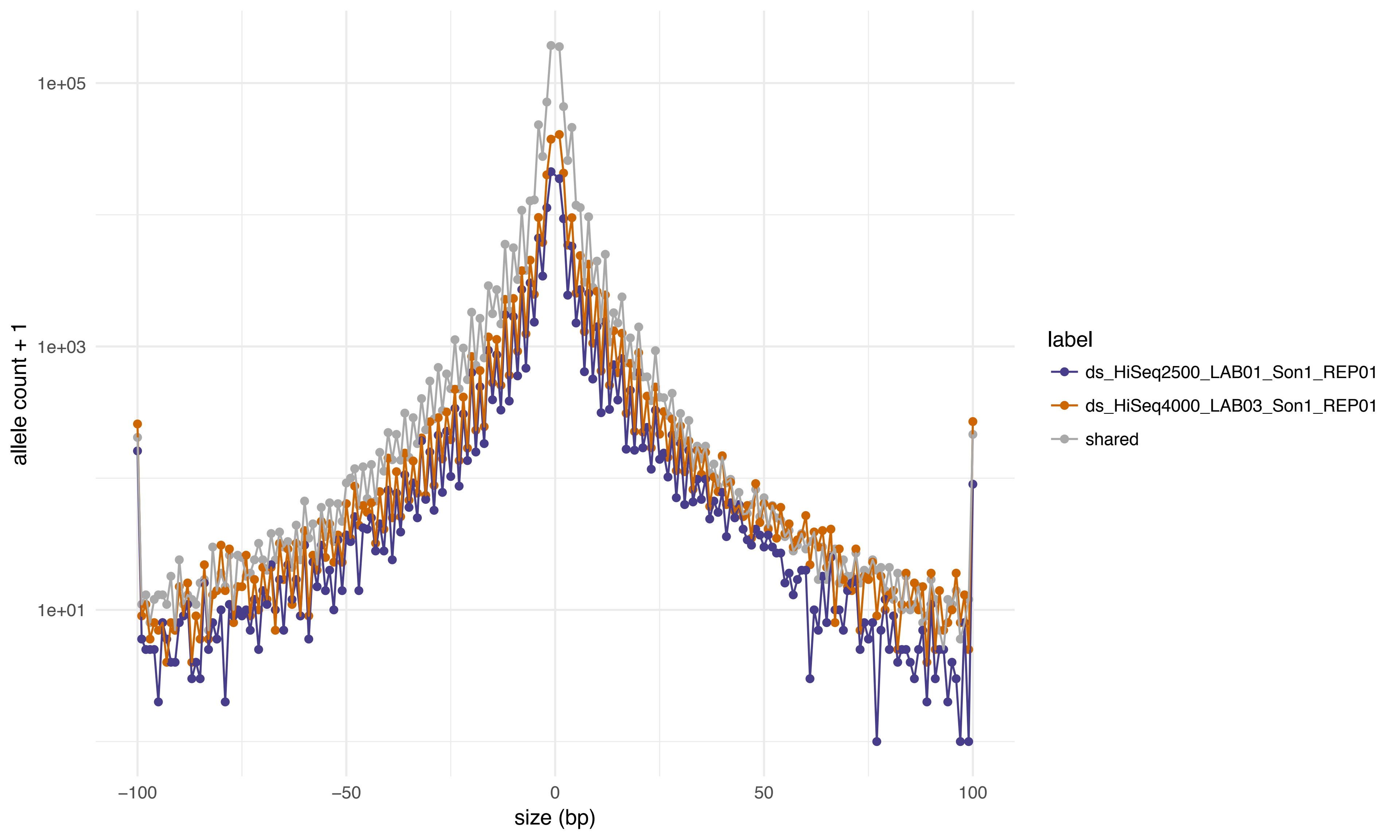
1e+05
1e+03
1e+01
allele count + 1
−100
−50
0
50
100
size (bp)
label
ds_HiSeq2500_LAB01_Son1_REP01
ds_HiSeq4000_LAB03_Son1_REP01
shared

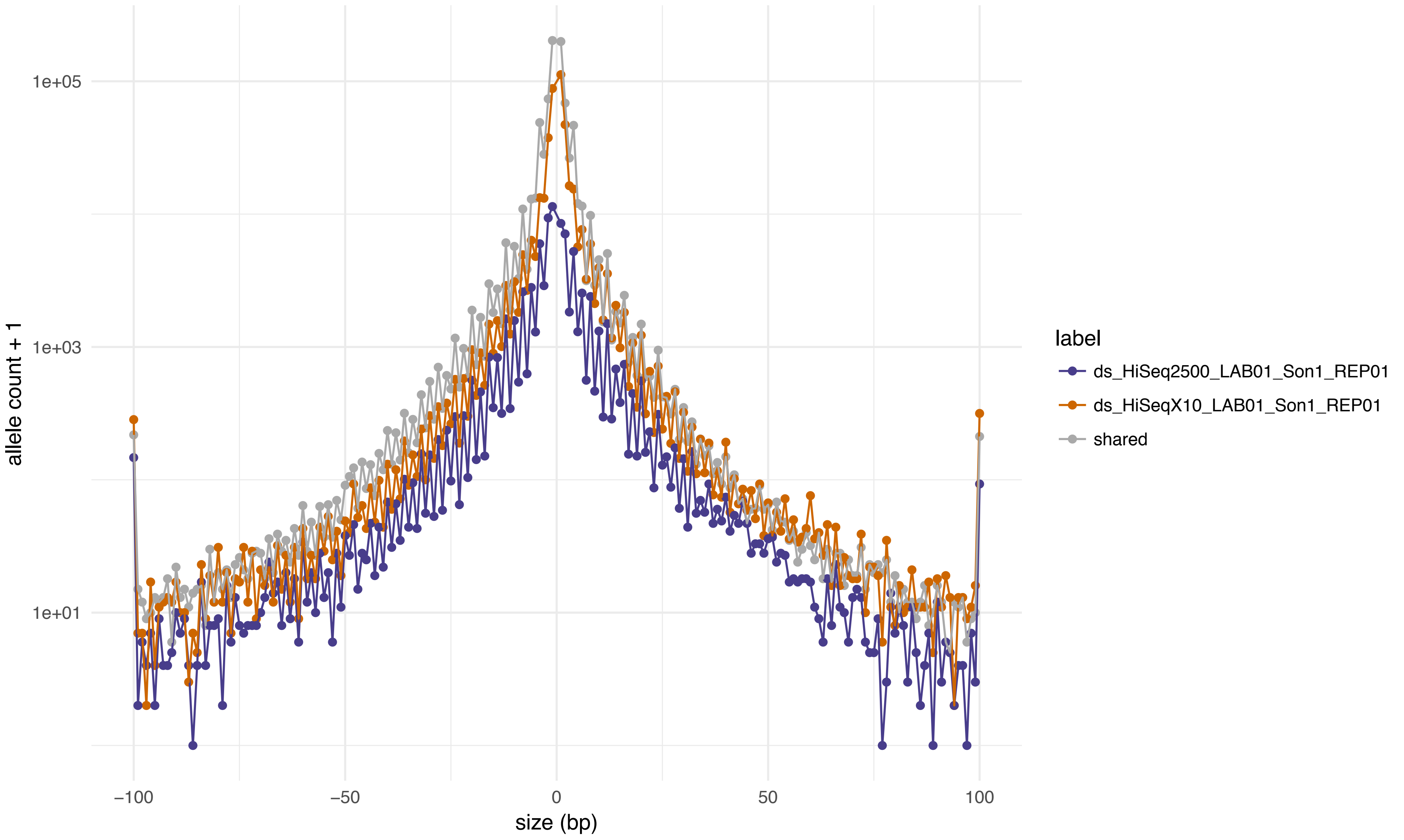

1e+05
1e+03
1e+01
allele count + 1
−100
−50
0
50
100
size (bp)
label
ds_HiSeq2500_LAB01_Son1_REP01
ds_HiSeqX10_LAB01_Son1_REP01
shared

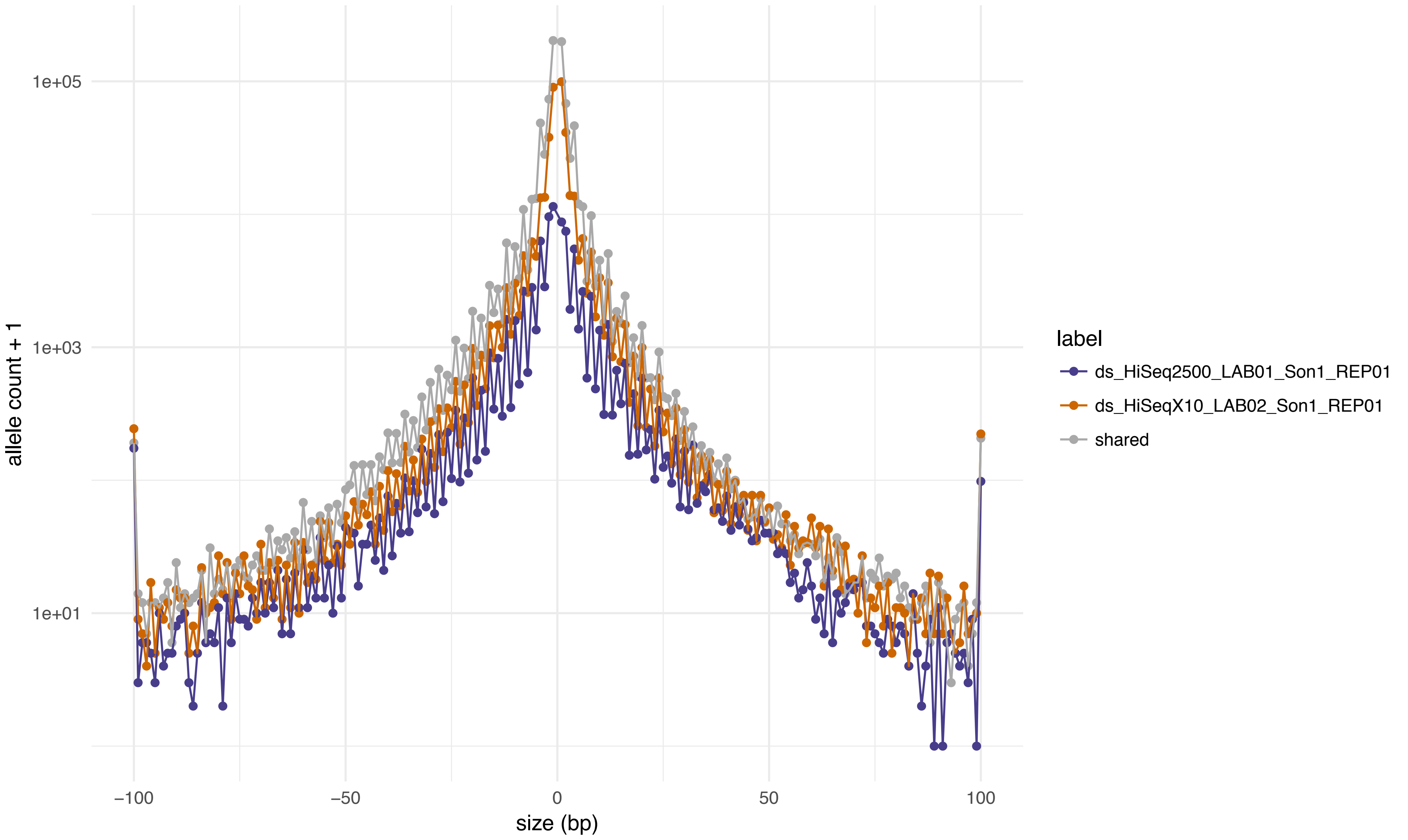

1e+05
1e+03
1e+01
allele count + 1
−100
−50
0
50
100
size (bp)
label
ds_HiSeq2500_LAB01_Son1_REP01
ds_HiSeqX10_LAB02_Son1_REP01
shared

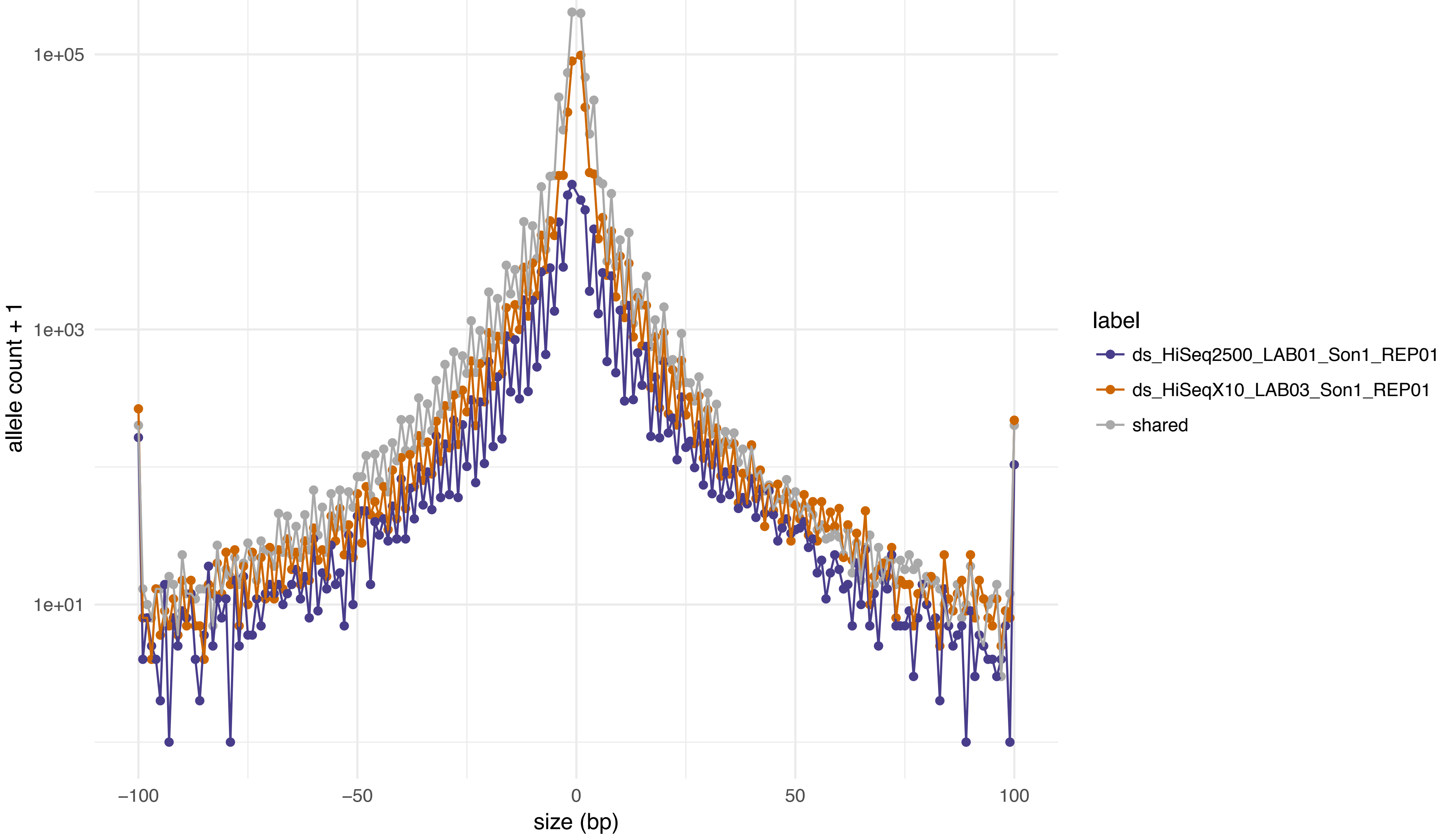

1e+05
1e+03
1e+01
allele count + 1
−100
−50
0
50
100
size (bp)
label
ds_HiSeq2500_LAB01_Son1_REP01
ds_HiSeqX10_LAB03_Son1_REP01
shared

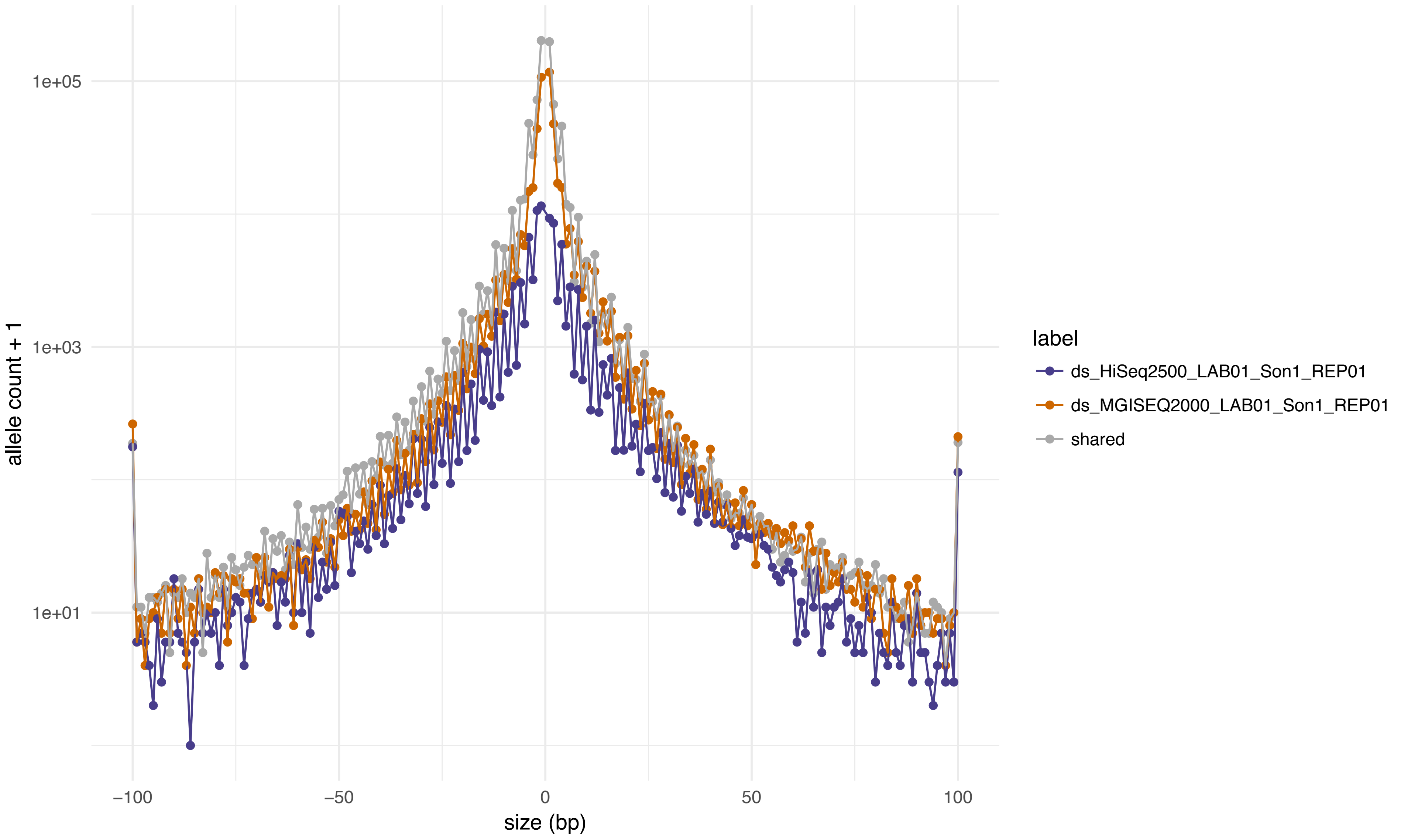

1e+05
1e+03
1e+01
allele count + 1
−100
−50
0
50
100
size (bp)
label
ds_HiSeq2500_LAB01_Son1_REP01
ds_MGISEQ2000_LAB01_Son1_REP01
shared

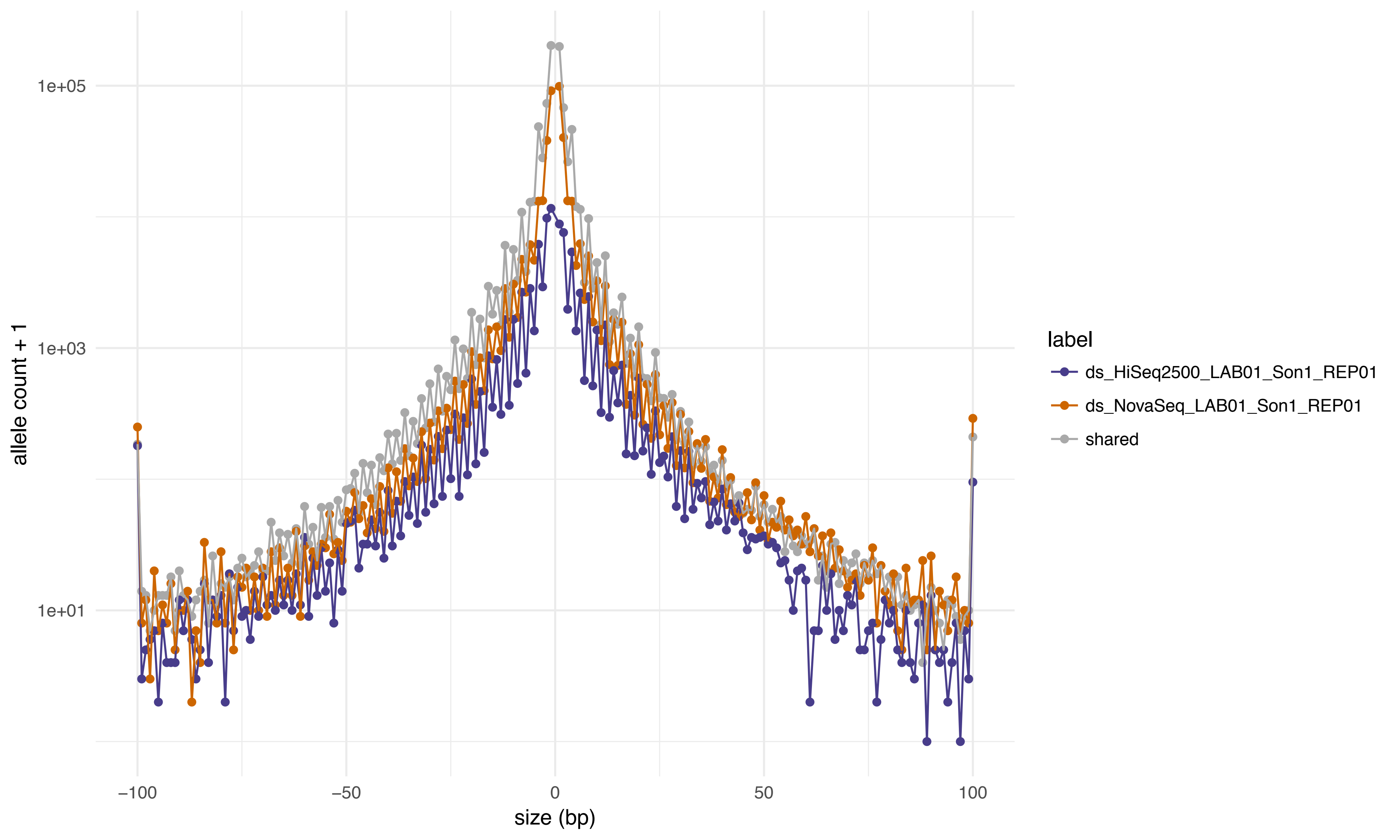
1e+05
1e+03
1e+01
allele count + 1
−100
−50
0
50
100
size (bp)
label
ds_HiSeq2500_LAB01_Son1_REP01
ds_NovaSeq_LAB01_Son1_REP01
shared

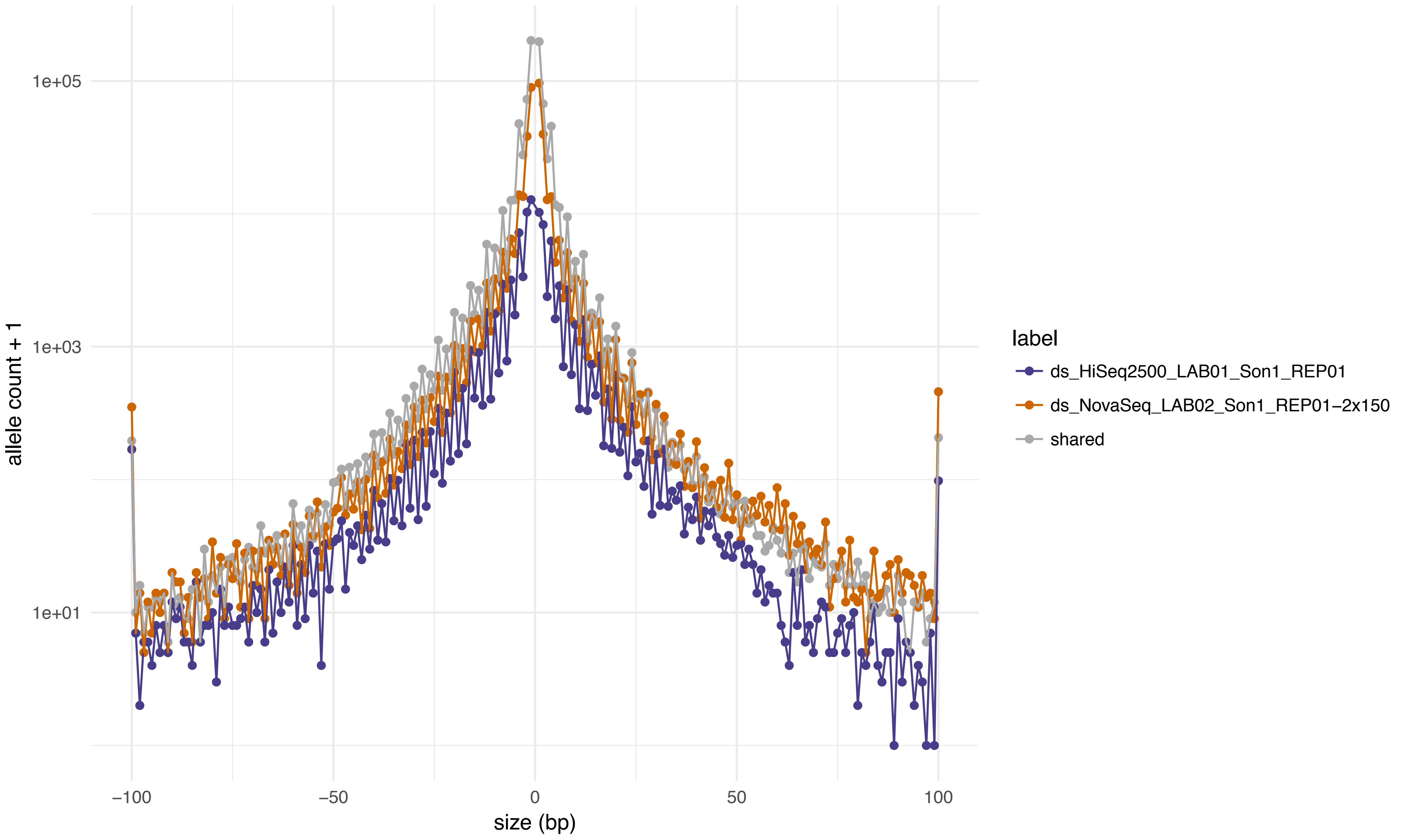

allele count + 1
1e+05
1e+03
1e+01
−100
−50
0
50
100
size (bp)
label
ds_HiSeq2500_LAB01_Son1_REP01
ds_NovaSeq_LAB02_Son1_REP01−2x150
shared

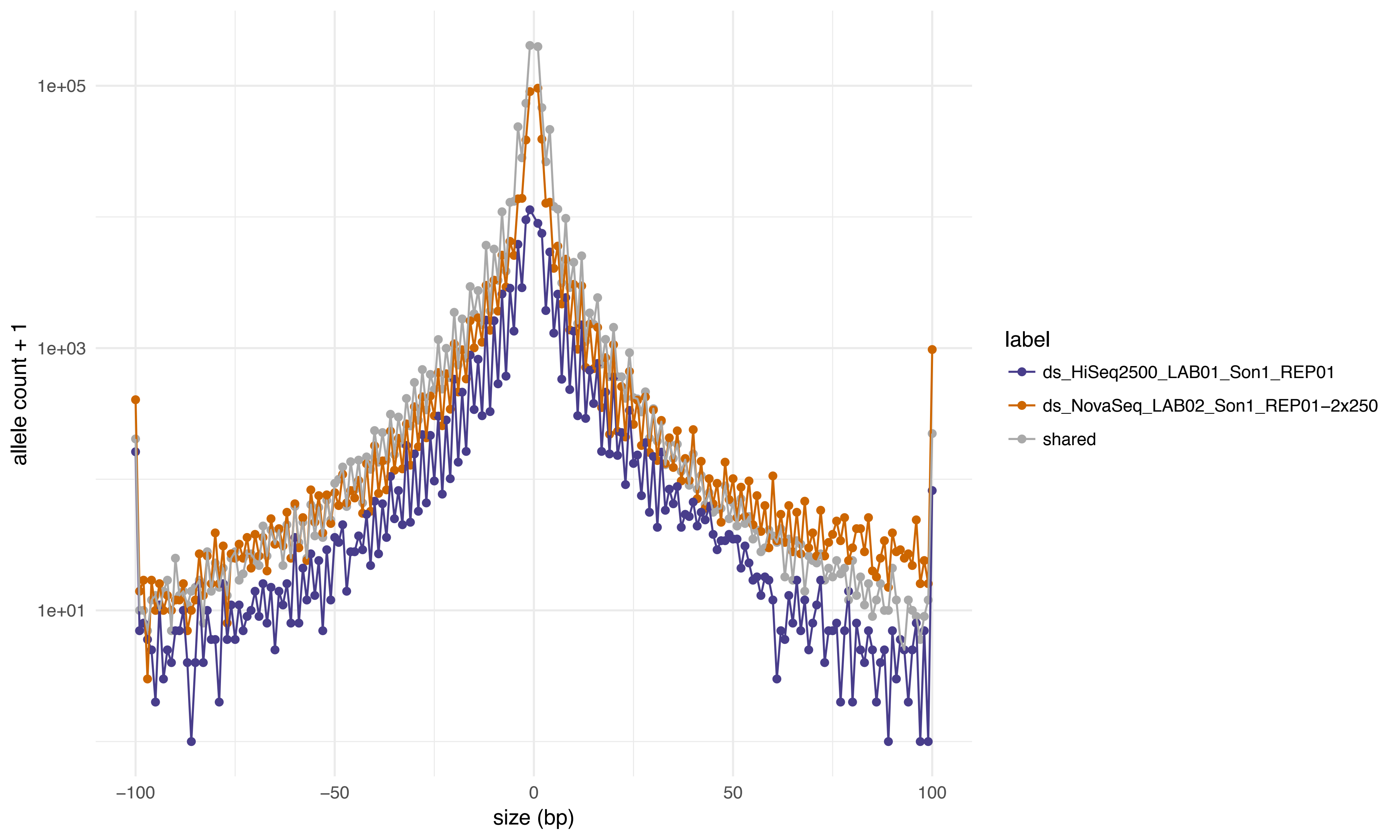

allele count + 1
1e+05
1e+03
1e+01
−100
−50
0
50
100
size (bp)
label
ds_HiSeq2500_LAB01_Son1_REP01
ds_NovaSeq_LAB02_Son1_REP01−2x250
shared

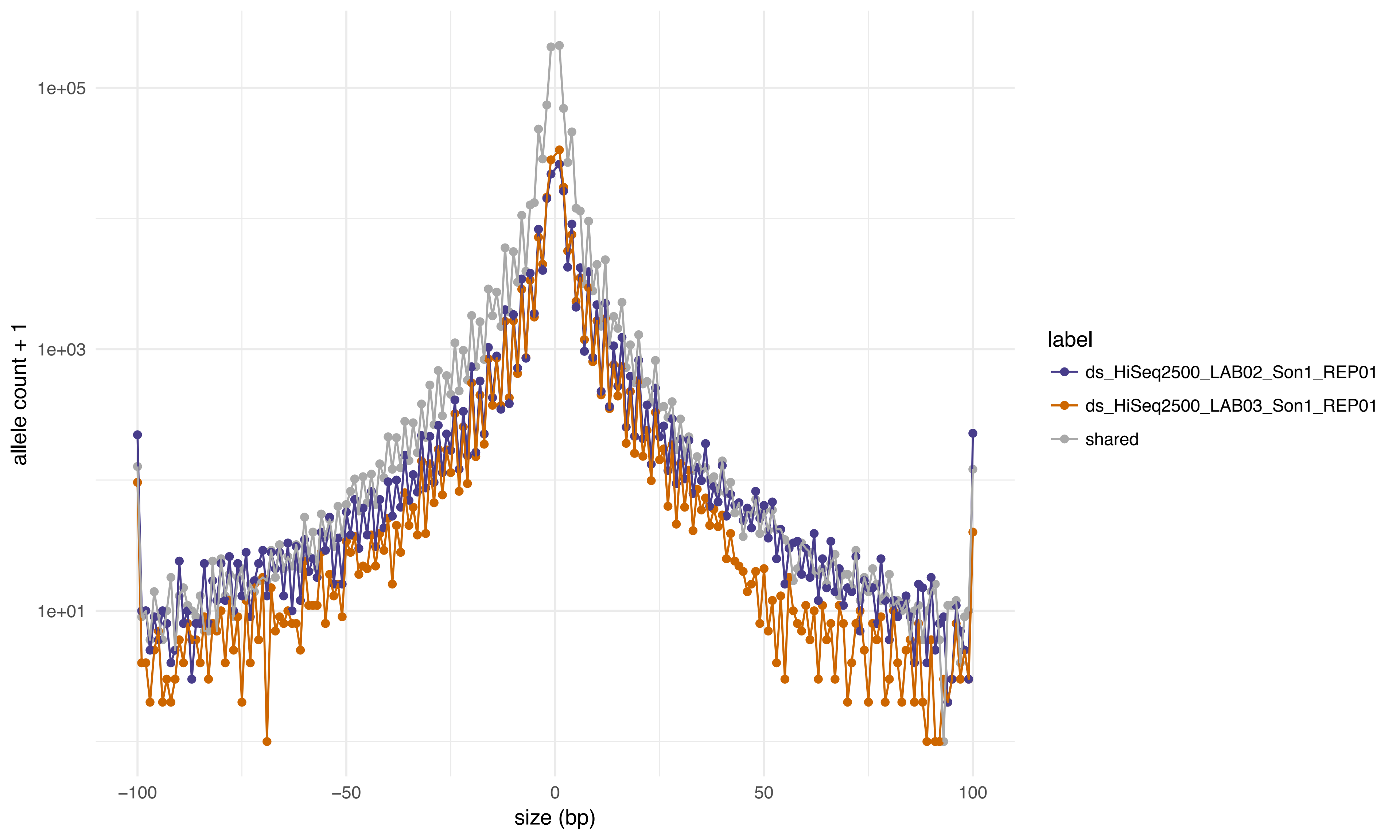
1e+05
1e+03
1e+01
allele count + 1
−100
−50
0
50
100
size (bp)
label
ds_HiSeq2500_LAB02_Son1_REP01
ds_HiSeq2500_LAB03_Son1_REP01
shared

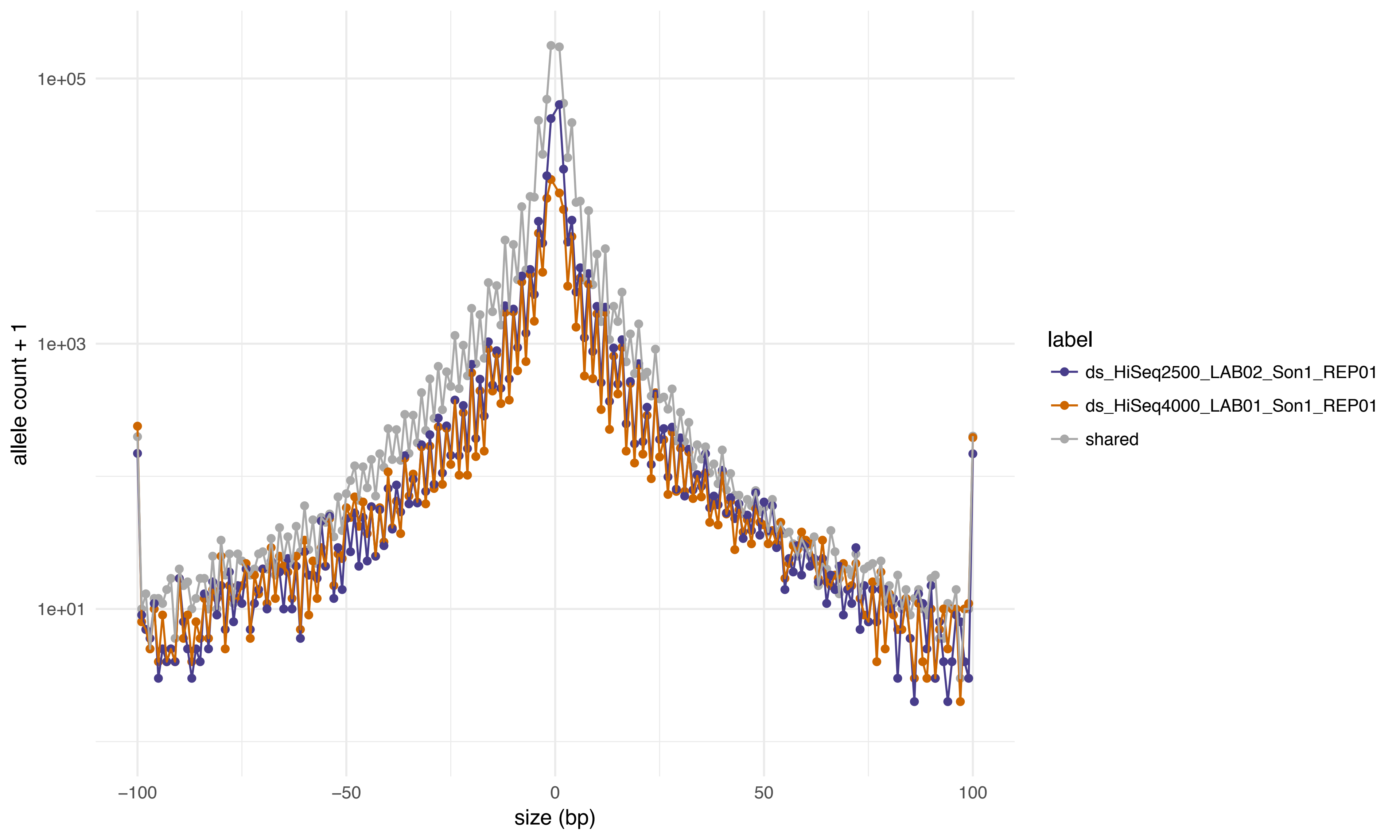

1e+05
1e+03
1e+01
allele count + 1
−100
−50
0
50
100
size (bp)
label
ds_HiSeq2500_LAB02_Son1_REP01
ds_HiSeq4000_LAB01_Son1_REP01
shared

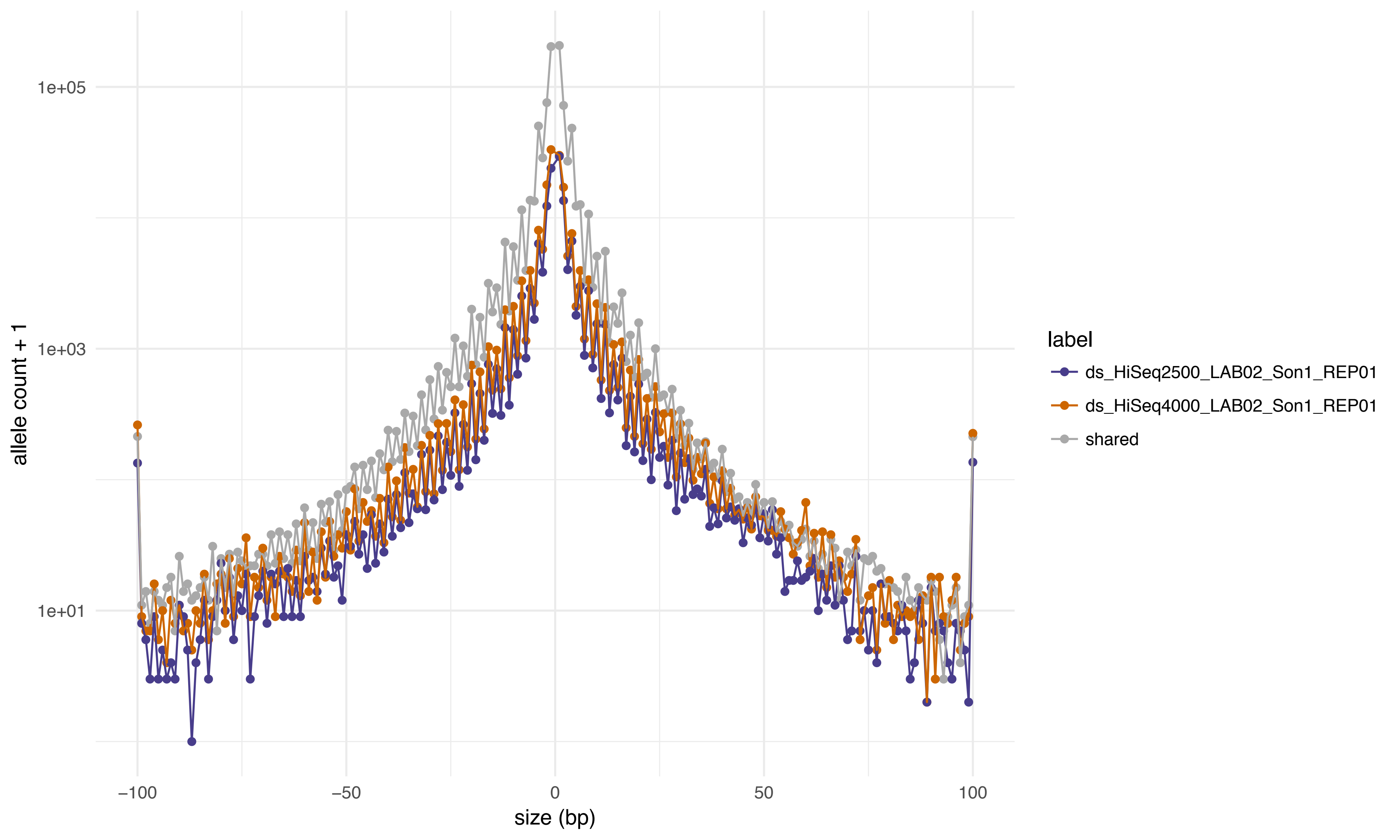
1e+05
1e+03
1e+01
allele count + 1
−100
−50
0
50
100
size (bp)
label
ds_HiSeq2500_LAB02_Son1_REP01
ds_HiSeq4000_LAB02_Son1_REP01
shared

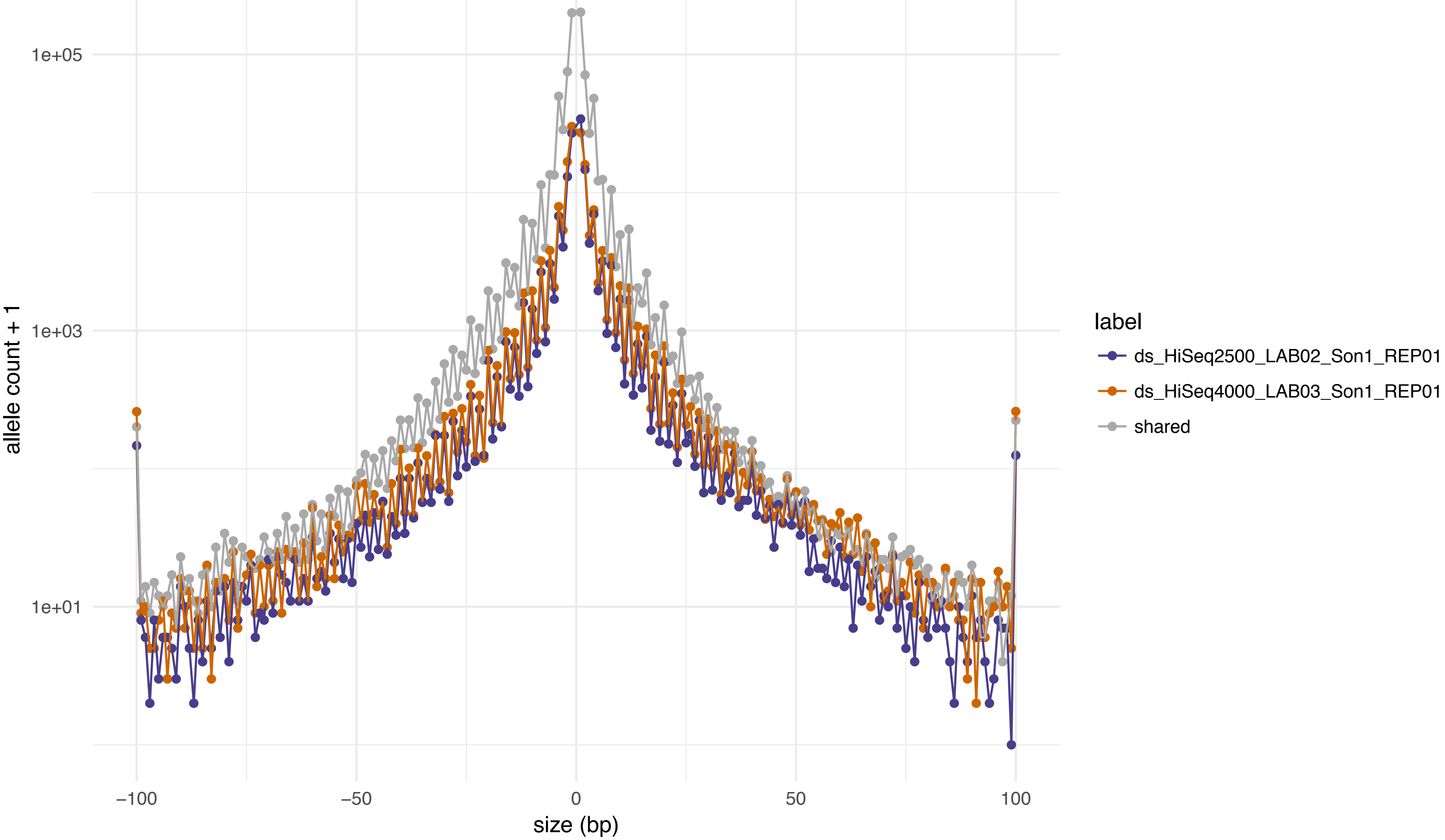

allele count + 1
1e+05
1e+03
1e+01
−100
−50
0
50
100
size (bp)
label
ds_HiSeq2500_LAB02_Son1_REP01
ds_HiSeq4000_LAB03_Son1_REP01
shared

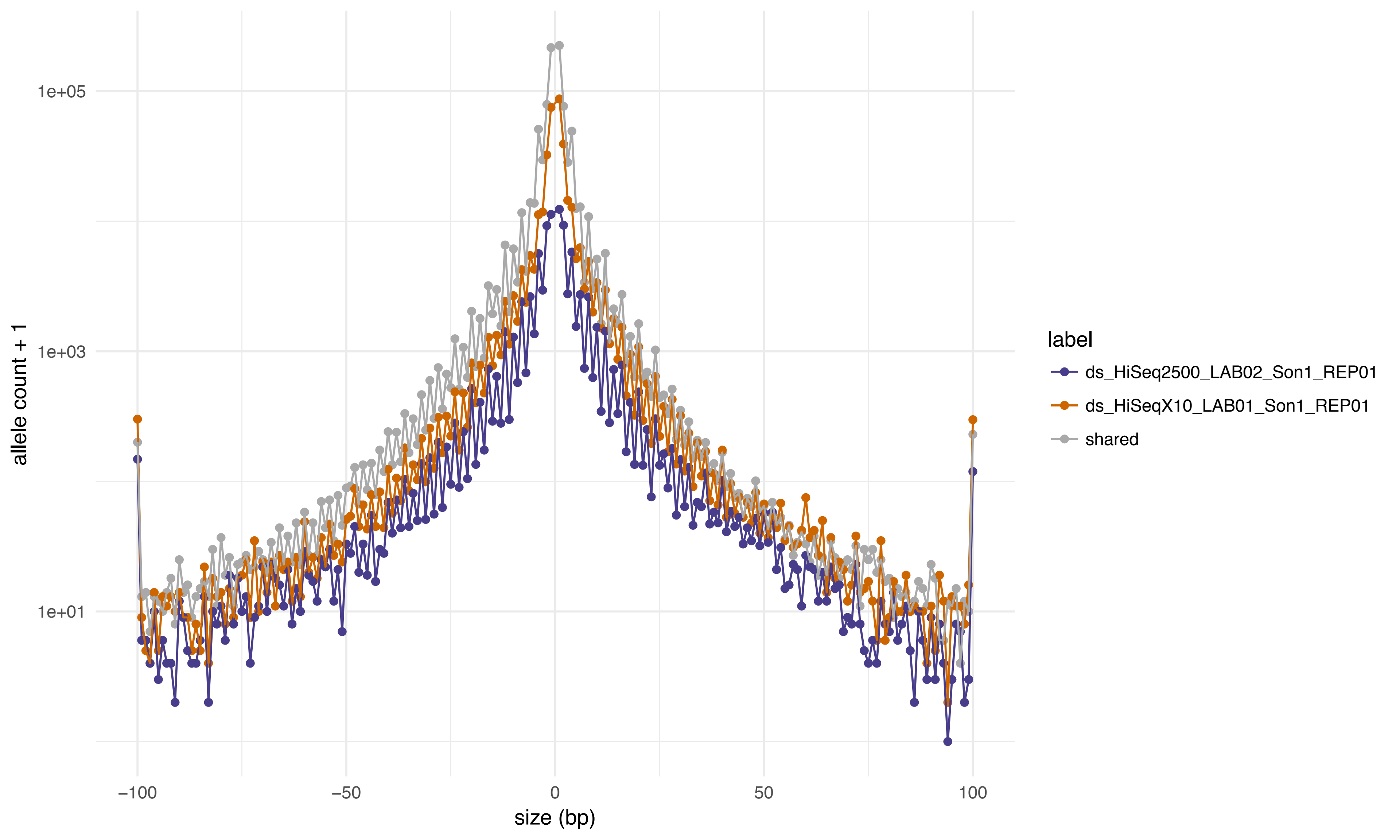

allele count + 1
1e+05
1e+03
1e+01
−100
−50
0
50
100
size (bp)
label
ds_HiSeq2500_LAB02_Son1_REP01
ds_HiSeqX10_LAB01_Son1_REP01
shared

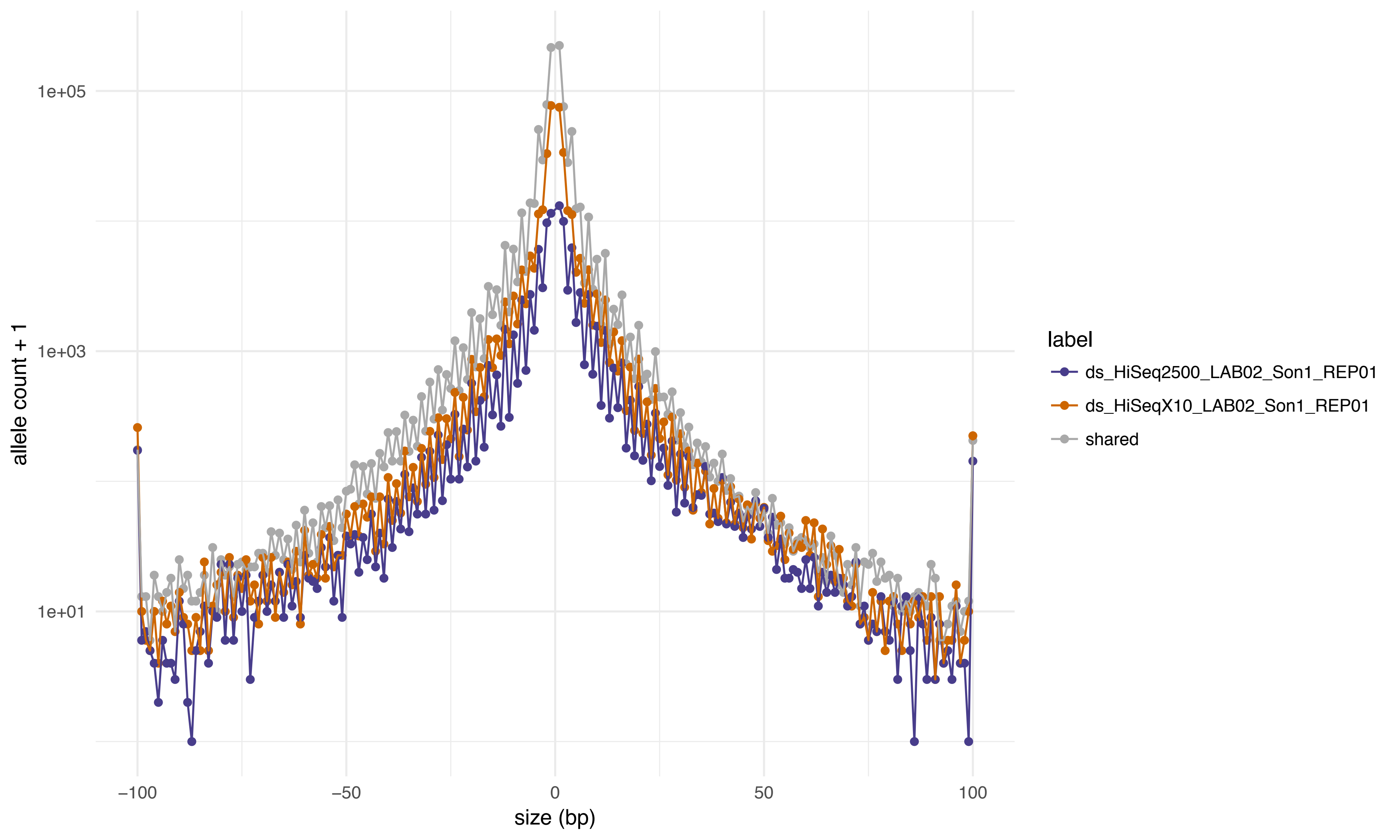

1e+05
1e+03
1e+01
allele count + 1
−100
−50
0
50
100
size (bp)
label
ds_HiSeq2500_LAB02_Son1_REP01
ds_HiSeqX10_LAB02_Son1_REP01
shared

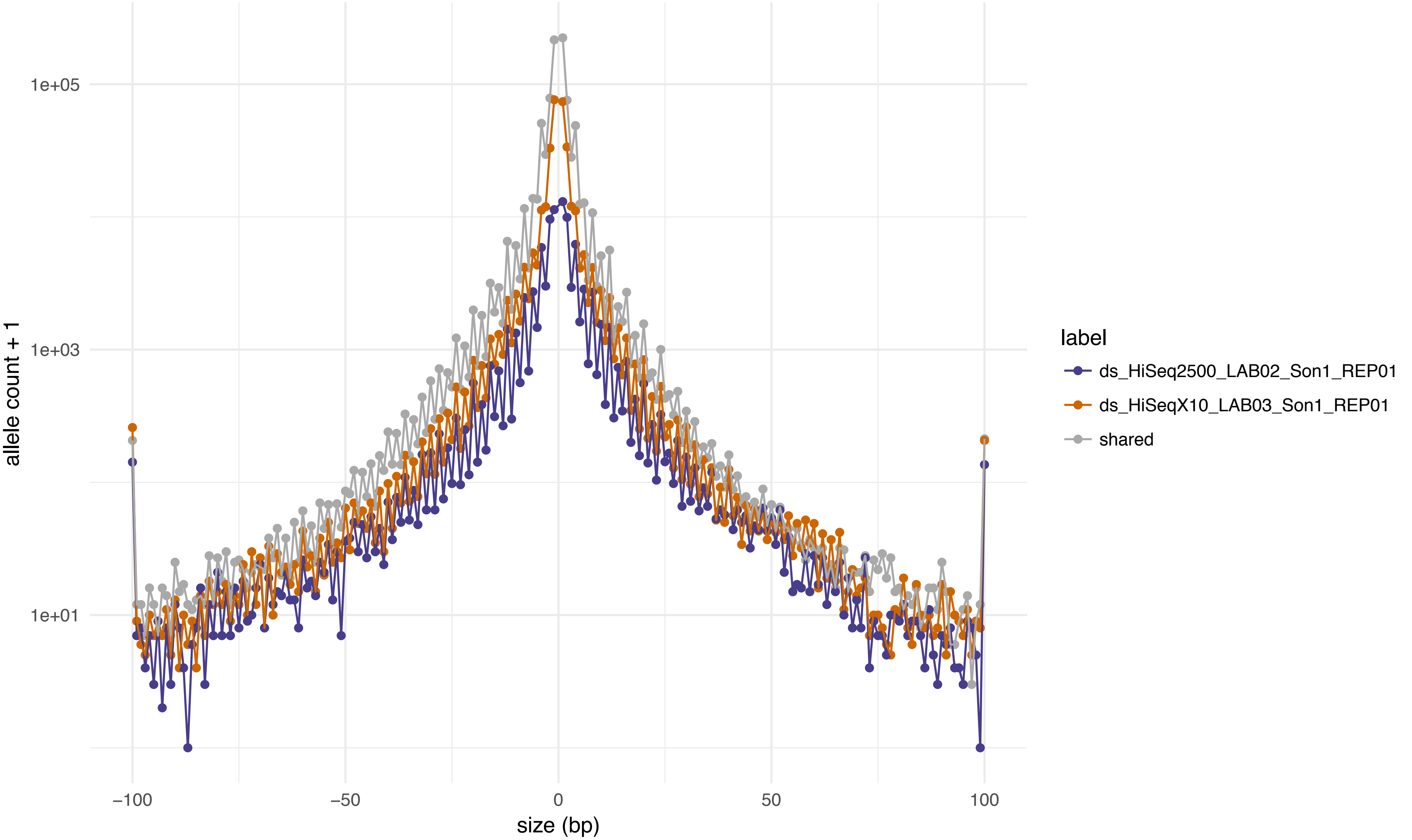

allele count + 1
1e+05
1e+03
1e+01
−100
−50
0
50
100
size (bp)
label
ds_HiSeq2500_LAB02_Son1_REP01
ds_HiSeqX10_LAB03_Son1_REP01
shared

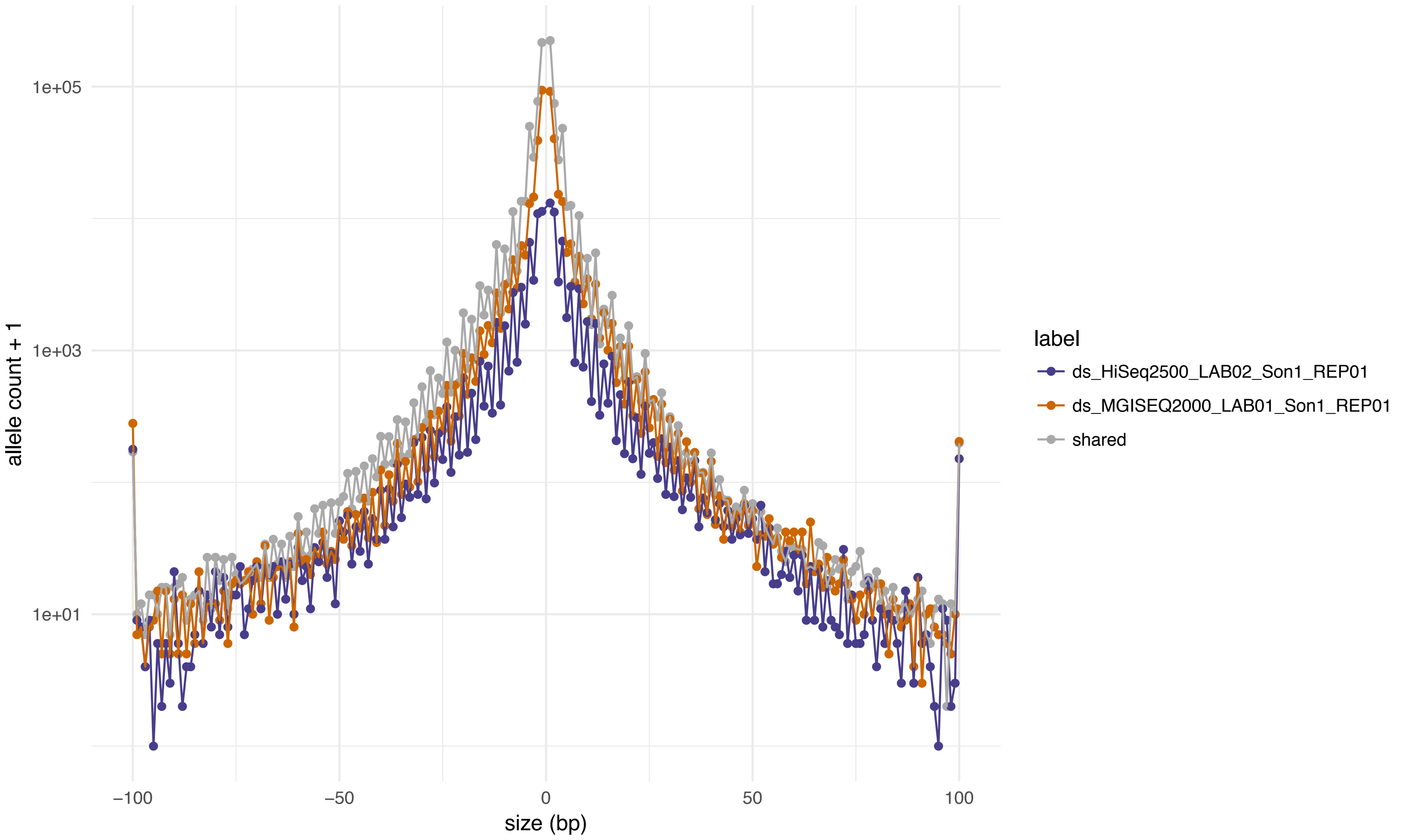
1e+05
1e+03
1e+01
allele count + 1
−100
−50
0
50
100
size (bp)
label
ds_HiSeq2500_LAB02_Son1_REP01
ds_MGISEQ2000_LAB01_Son1_REP01
shared

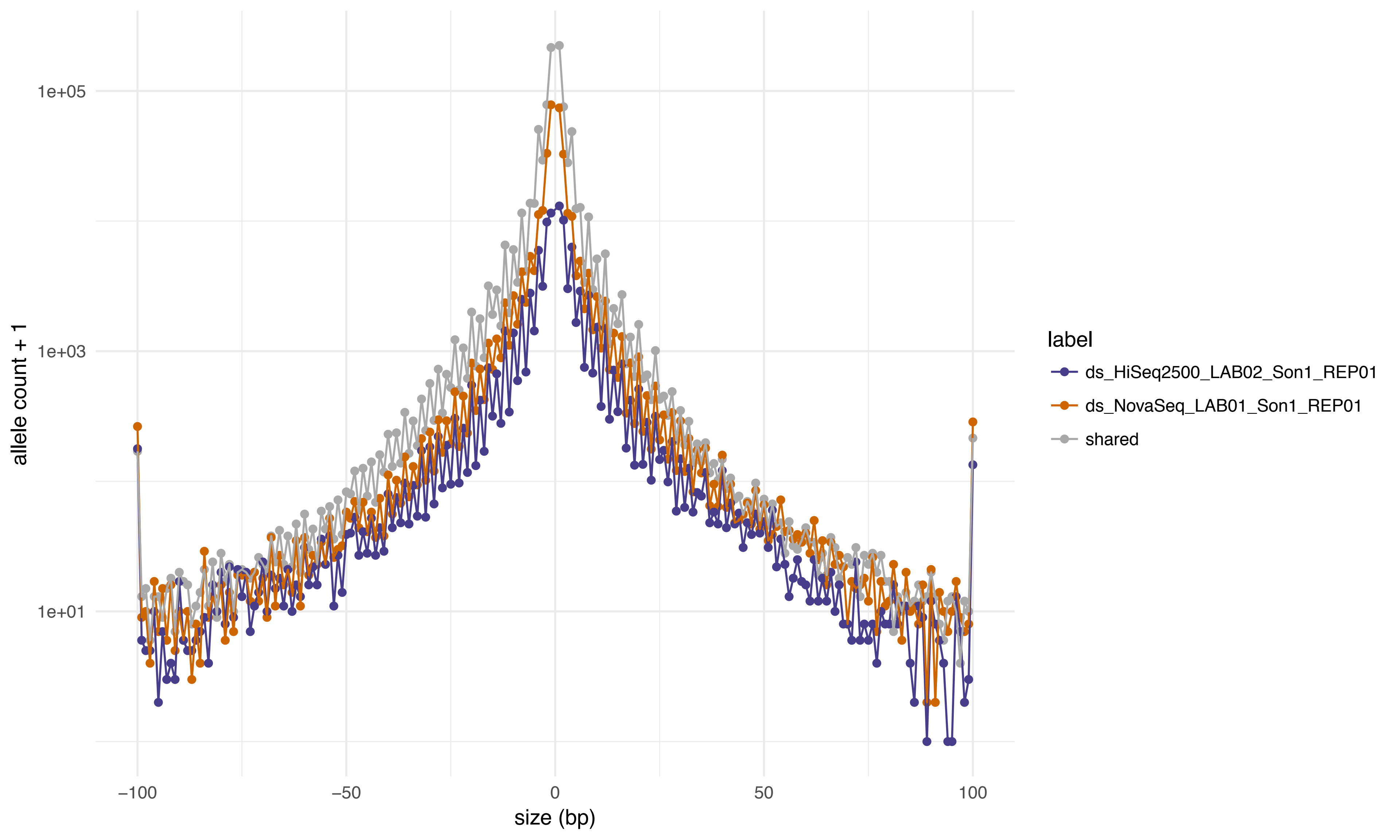
1e+05
1e+03
1e+01
allele count + 1
−100
−50
0
50
100
size (bp)
label
ds_HiSeq2500_LAB02_Son1_REP01
ds_NovaSeq_LAB01_Son1_REP01
shared

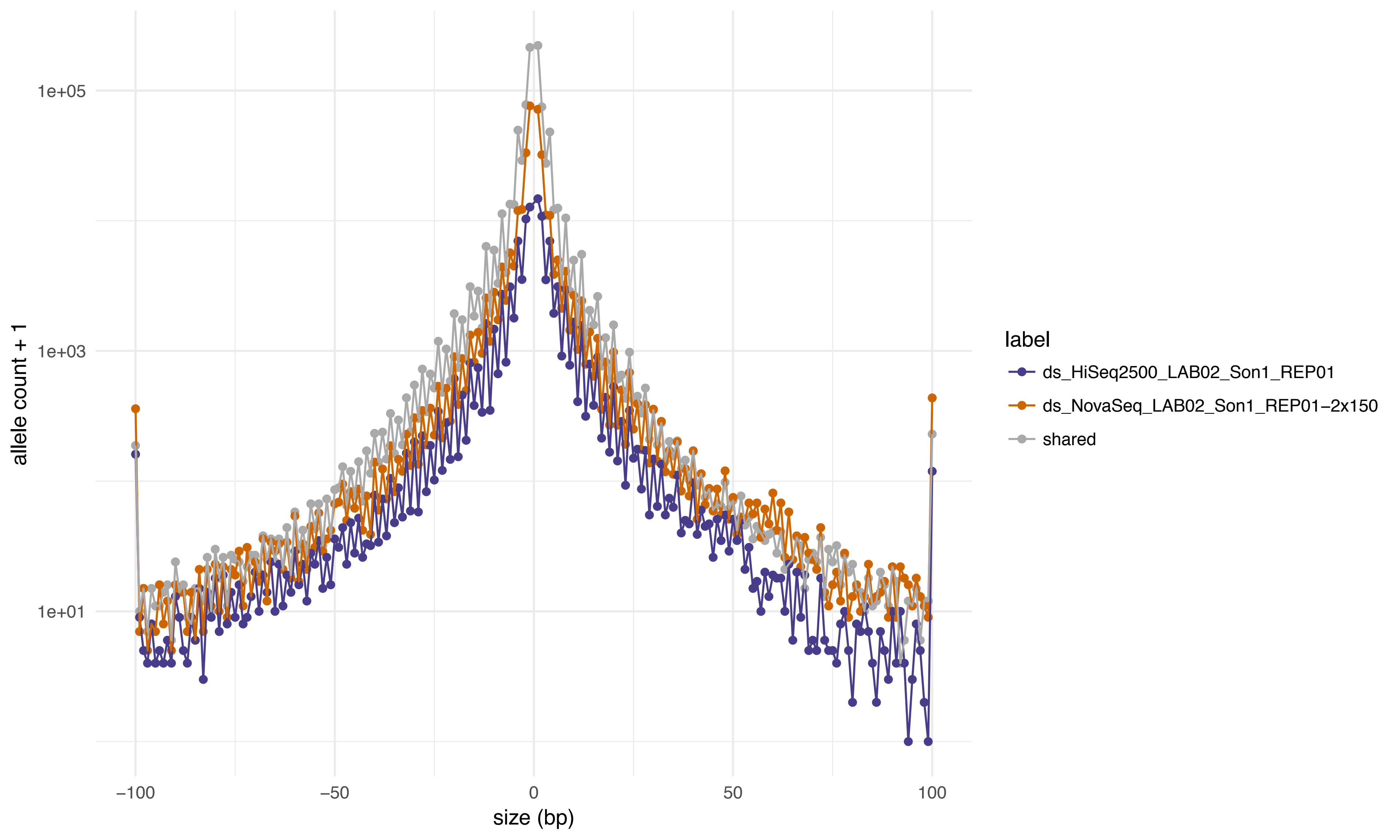
1e+05
1e+03
1e+01
allele count + 1
−100
−50
0
50
100
size (bp)
label
ds_HiSeq2500_LAB02_Son1_REP01
ds_NovaSeq_LAB02_Son1_REP01−2x150
shared

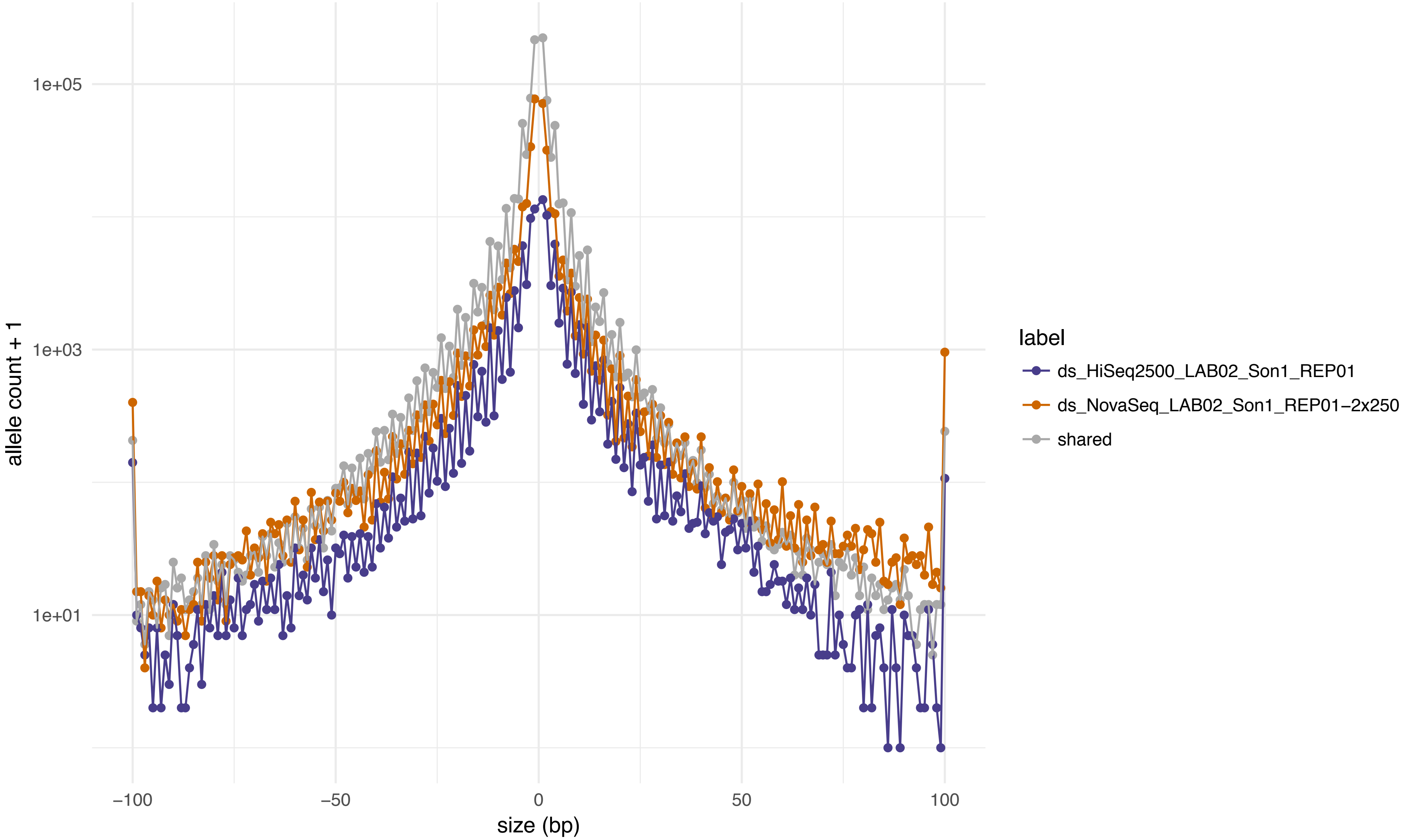

1e+05
1e+03
1e+01
allele count + 1
−100
−50
0
50
100
size (bp)
label
ds_HiSeq2500_LAB02_Son1_REP01
ds_NovaSeq_LAB02_Son1_REP01−2x250
shared

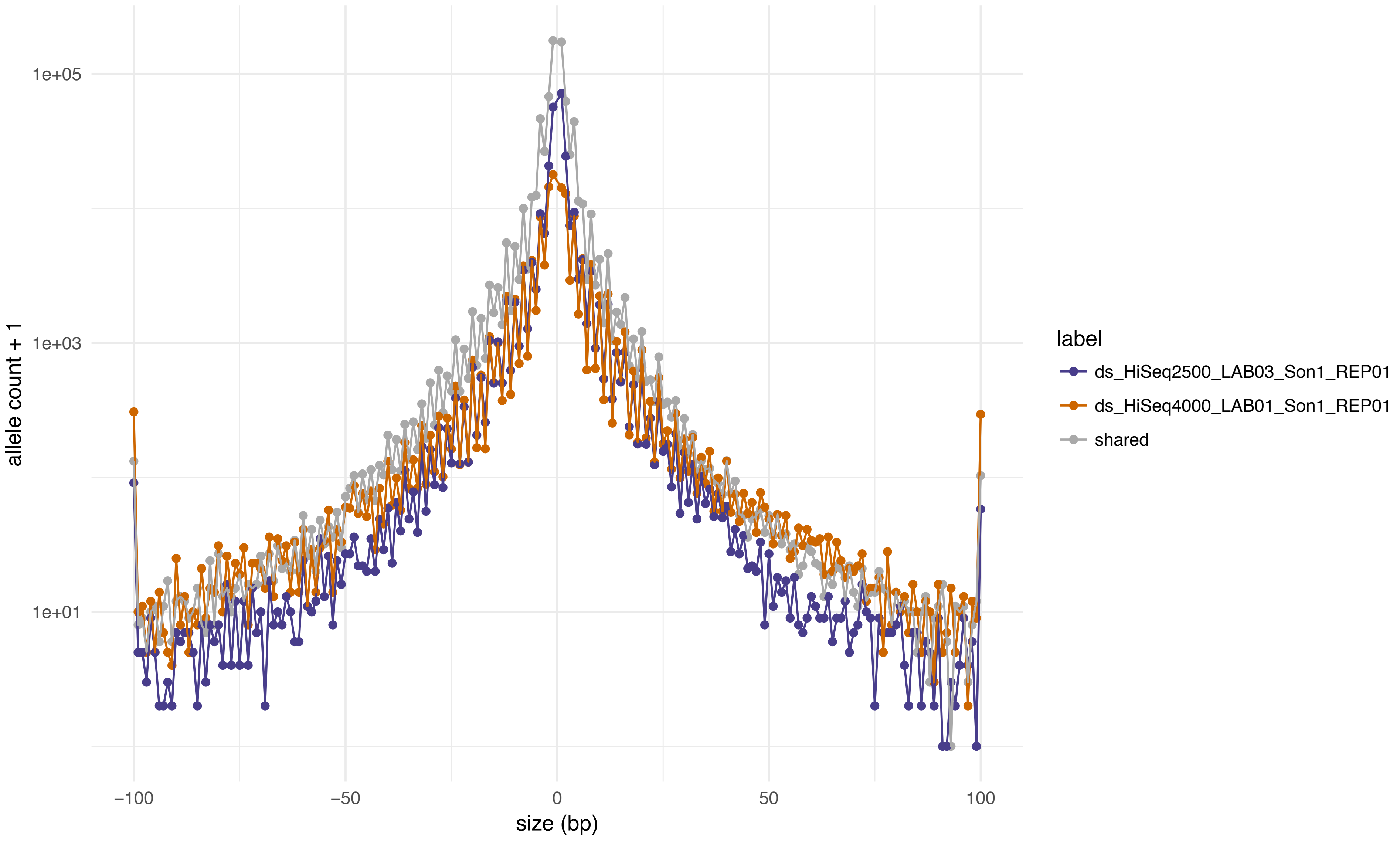
1e+05
1e+03
1e+01
allele count + 1
−100
−50
0
50
100
size (bp)
label
ds_HiSeq2500_LAB03_Son1_REP01
ds_HiSeq4000_LAB01_Son1_REP01
shared

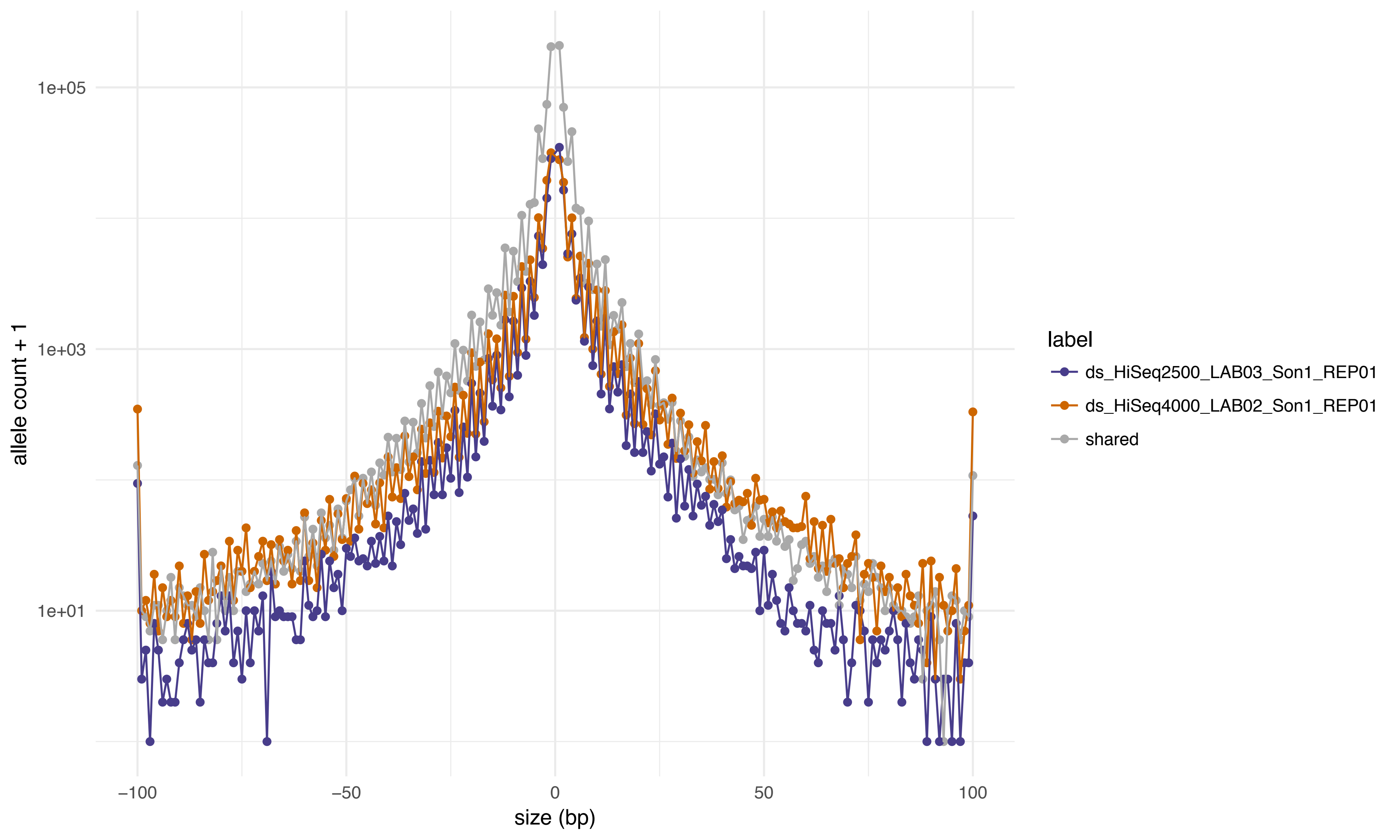
1e+05
1e+03
1e+01
allele count + 1
−100
−50
0
50
100
size (bp)
label
ds_HiSeq2500_LAB03_Son1_REP01
ds_HiSeq4000_LAB02_Son1_REP01
shared

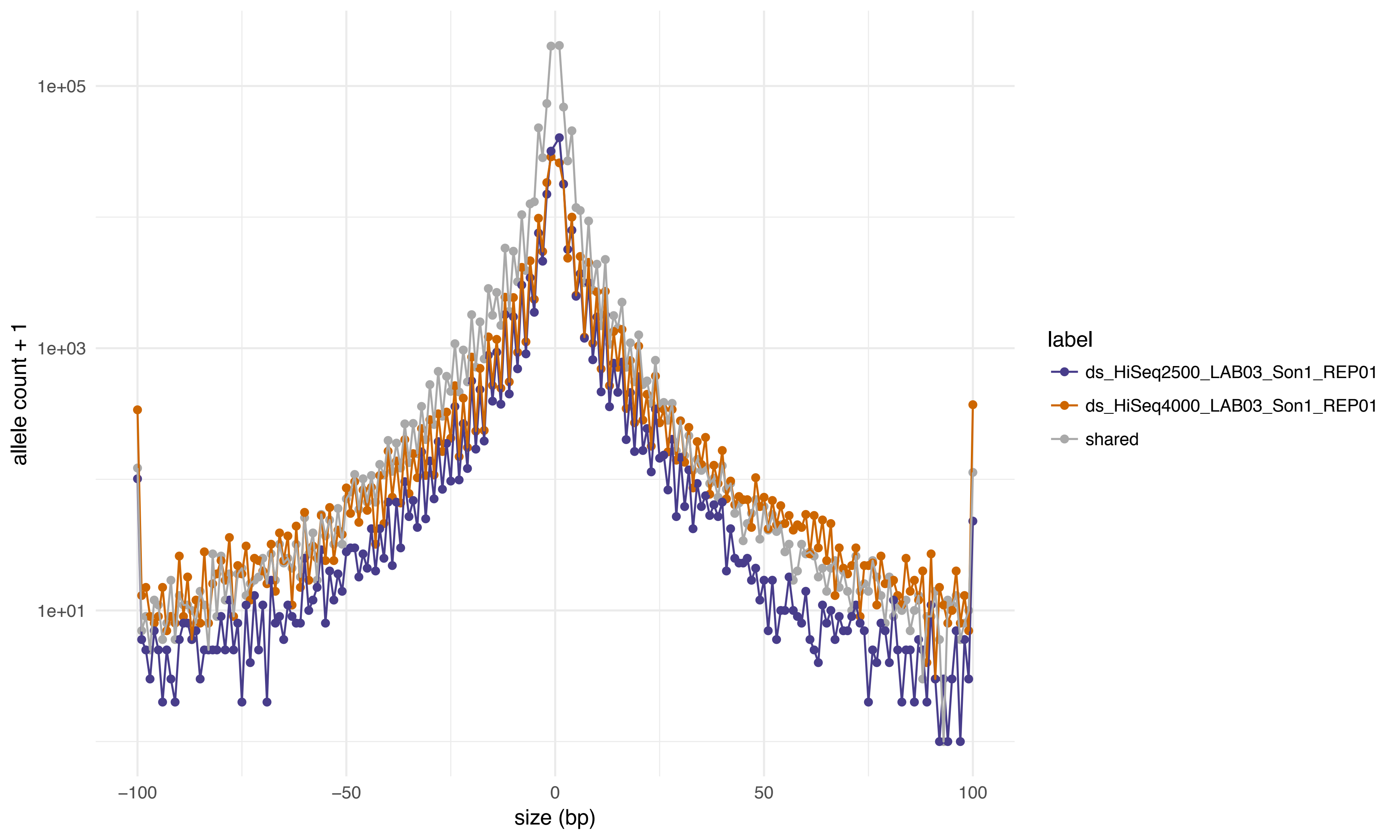
1e+05
1e+03
1e+01
allele count + 1
−100
−50
0
50
100
size (bp)
label
ds_HiSeq2500_LAB03_Son1_REP01
ds_HiSeq4000_LAB03_Son1_REP01
shared

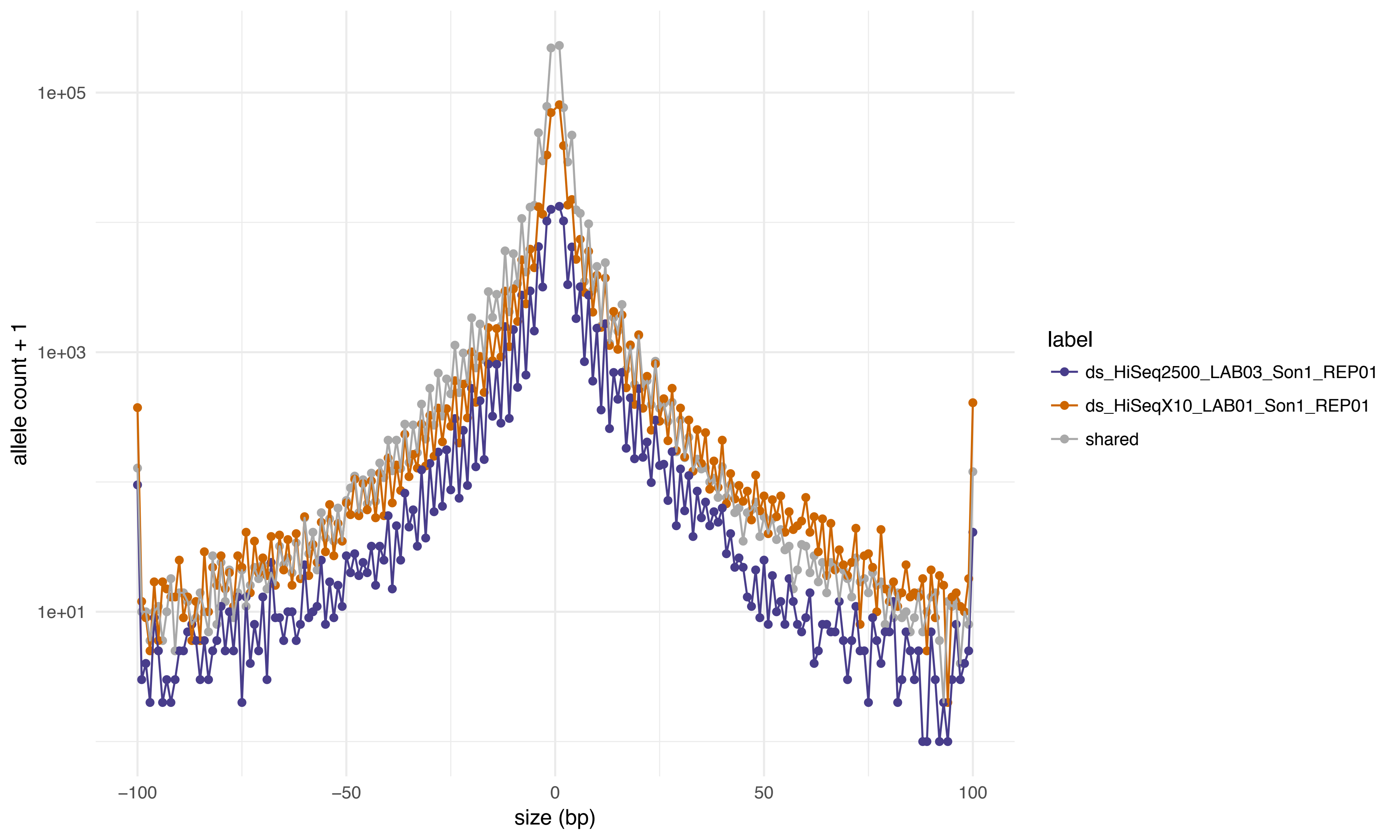

1e+05
1e+03
1e+01
allele count + 1
−100
−50
0
50
100
size (bp)
label
ds_HiSeq2500_LAB03_Son1_REP01
ds_HiSeqX10_LAB01_Son1_REP01
shared

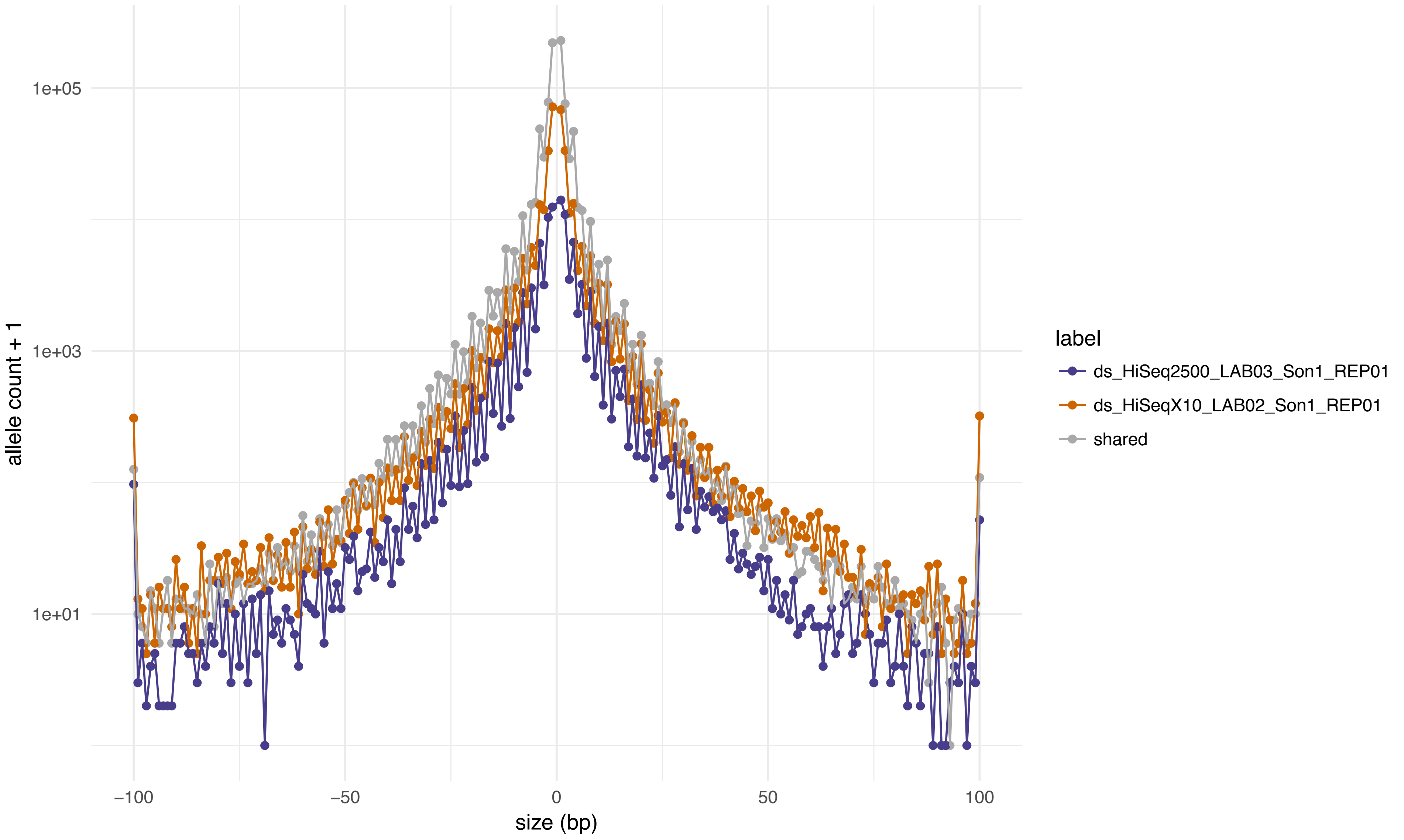
1e+05
1e+03
1e+01
allele count + 1
−100
−50
0
50
100
size (bp)
label
ds_HiSeq2500_LAB03_Son1_REP01
ds_HiSeqX10_LAB02_Son1_REP01
shared

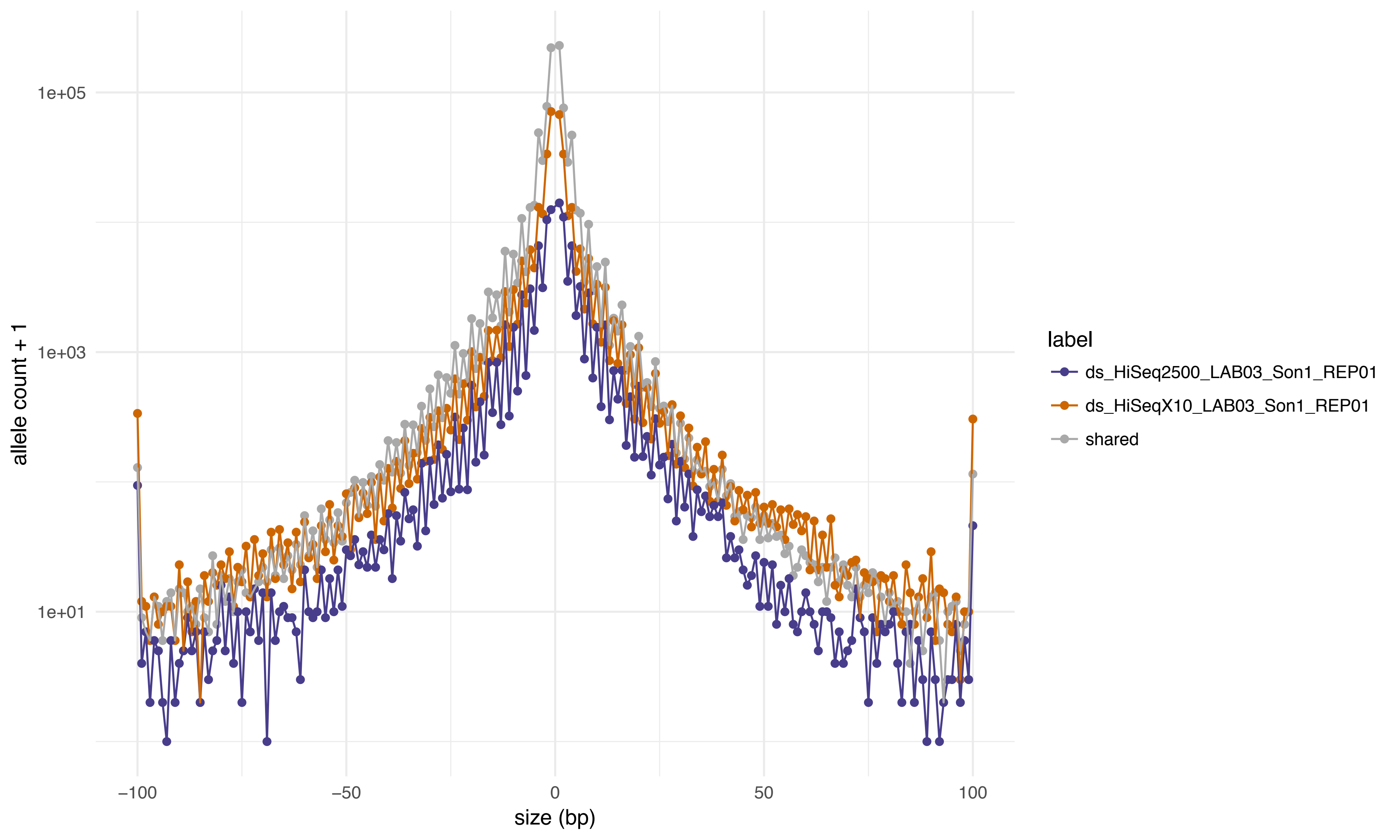

1e+05
1e+03
1e+01
allele count + 1
−100
−50
0
50
100
size (bp)
label
ds_HiSeq2500_LAB03_Son1_REP01
ds_HiSeqX10_LAB03_Son1_REP01
shared

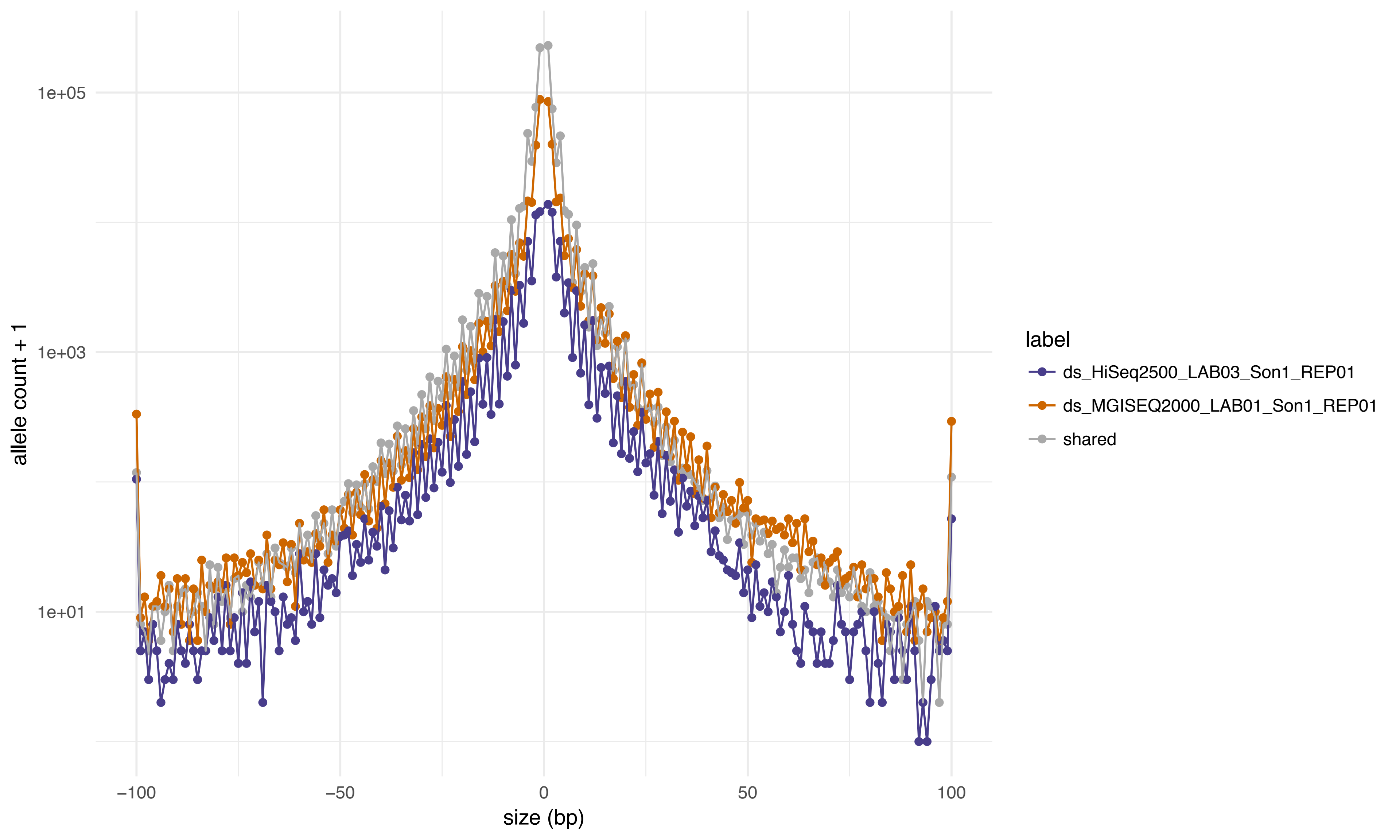

1e+05
1e+03
1e+01
allele count + 1
−100
−50
0
50
100
size (bp)
label
ds_HiSeq2500_LAB03_Son1_REP01
ds_MGISEQ2000_LAB01_Son1_REP01
shared

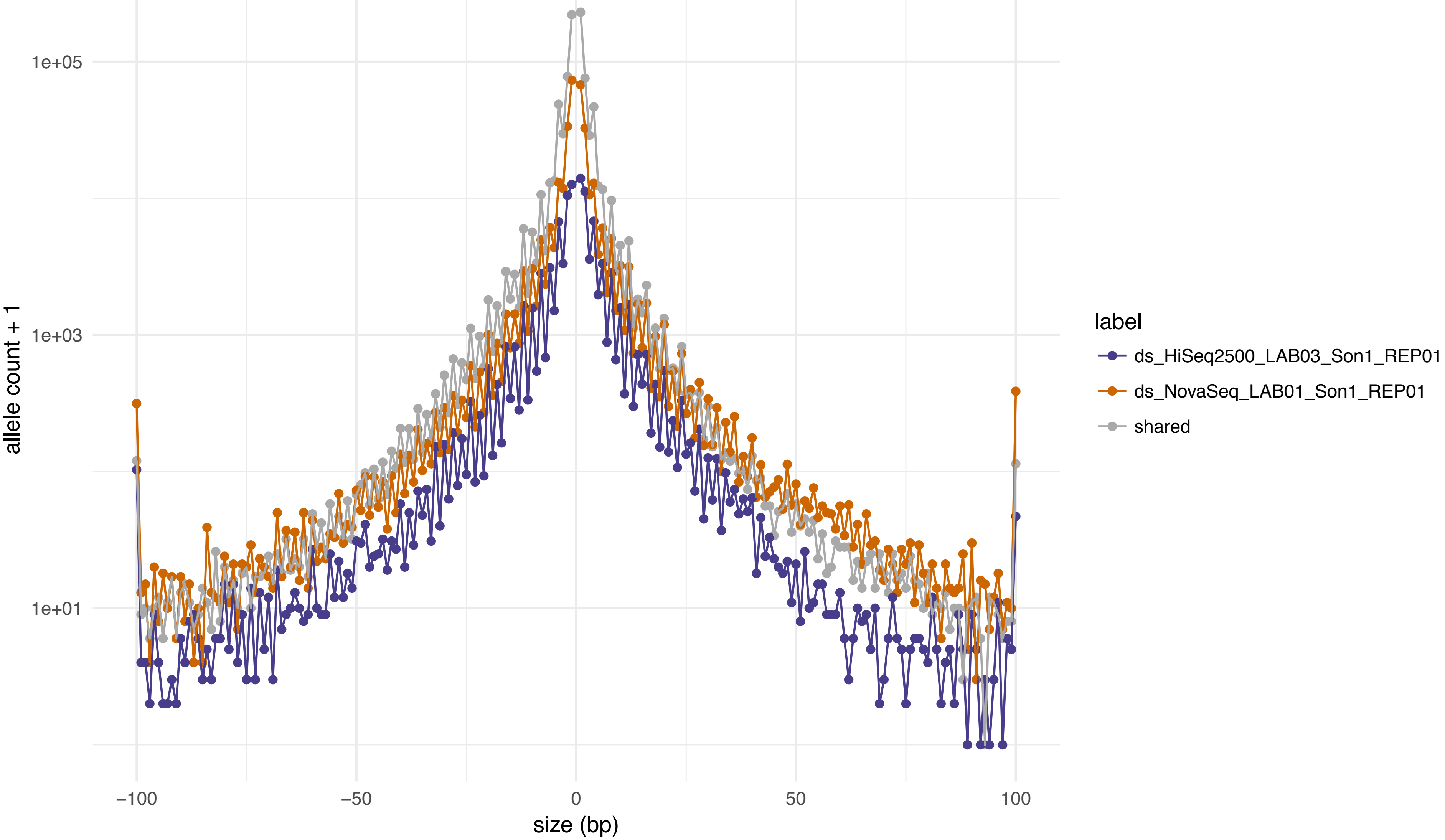
1e+05
1e+03
1e+01
allele count + 1
−100
−50
0
50
100
size (bp)
label
ds_HiSeq2500_LAB03_Son1_REP01
ds_NovaSeq_LAB01_Son1_REP01
shared

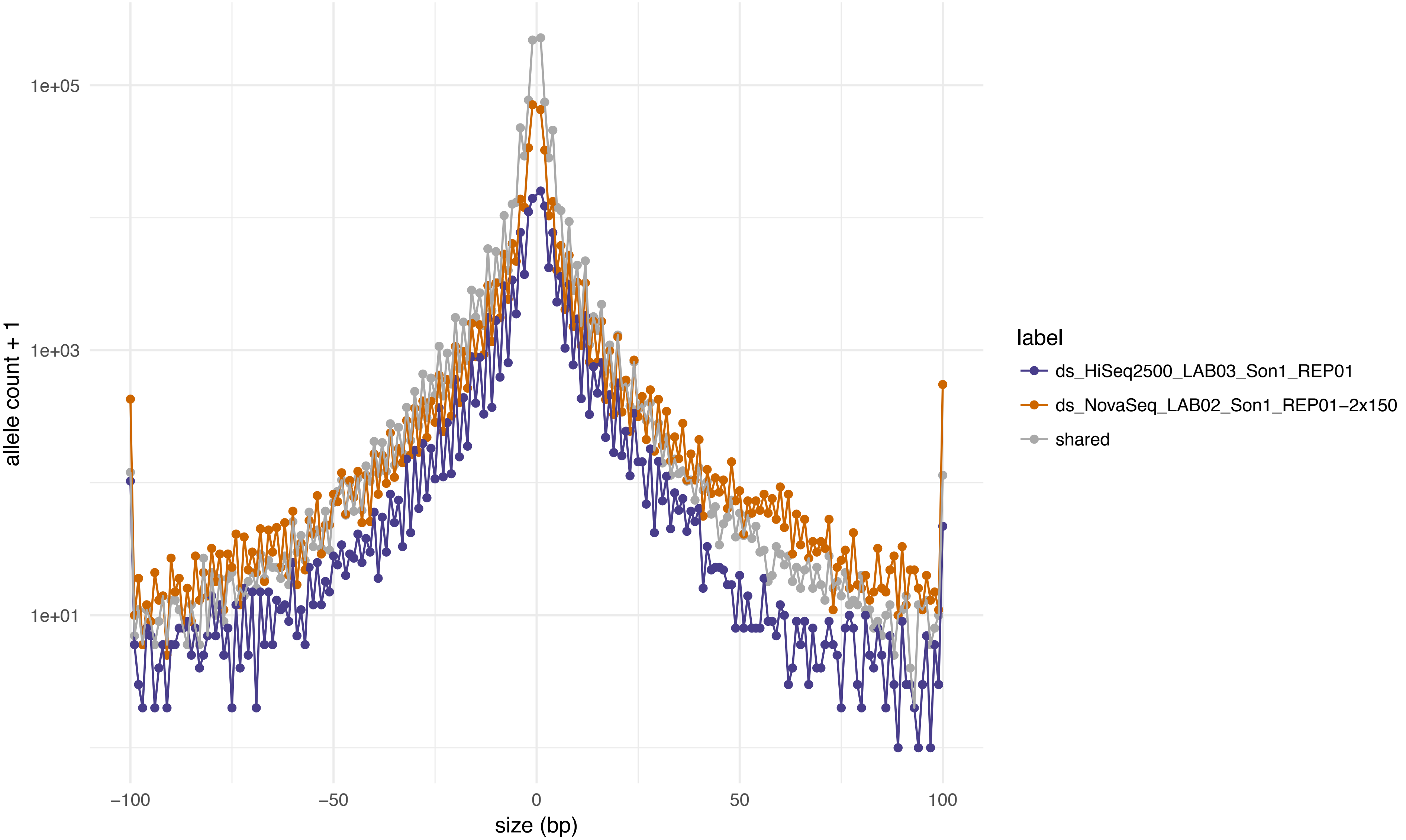

1e+05
1e+03
1e+01
allele count + 1
−100
−50
0
50
100
size (bp)
label
ds_HiSeq2500_LAB03_Son1_REP01
ds_NovaSeq_LAB02_Son1_REP01−2x150
shared

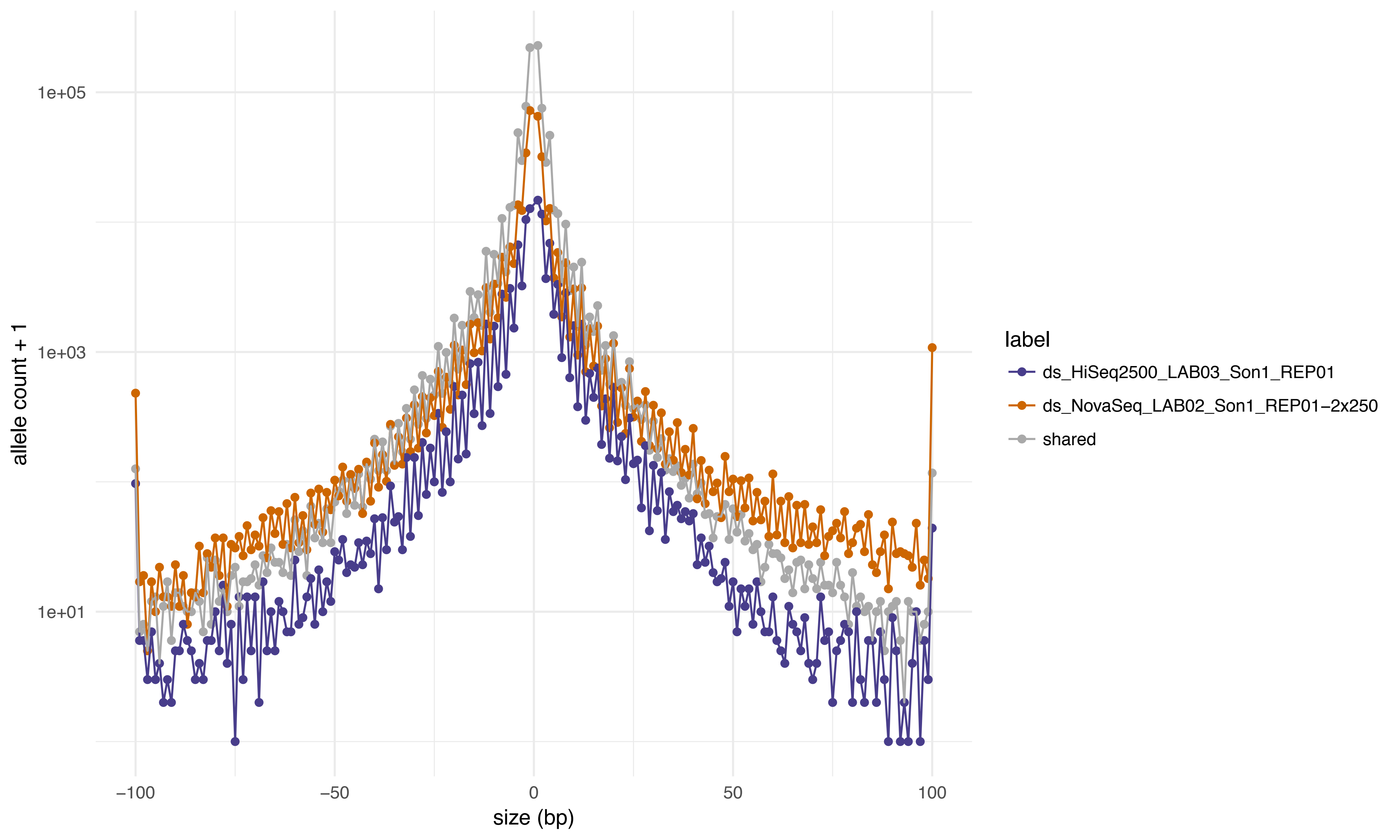

1e+05
1e+03
1e+01
allele count + 1
−100
−50
0
50
100
size (bp)
label
ds_HiSeq2500_LAB03_Son1_REP01
ds_NovaSeq_LAB02_Son1_REP01−2x250
shared

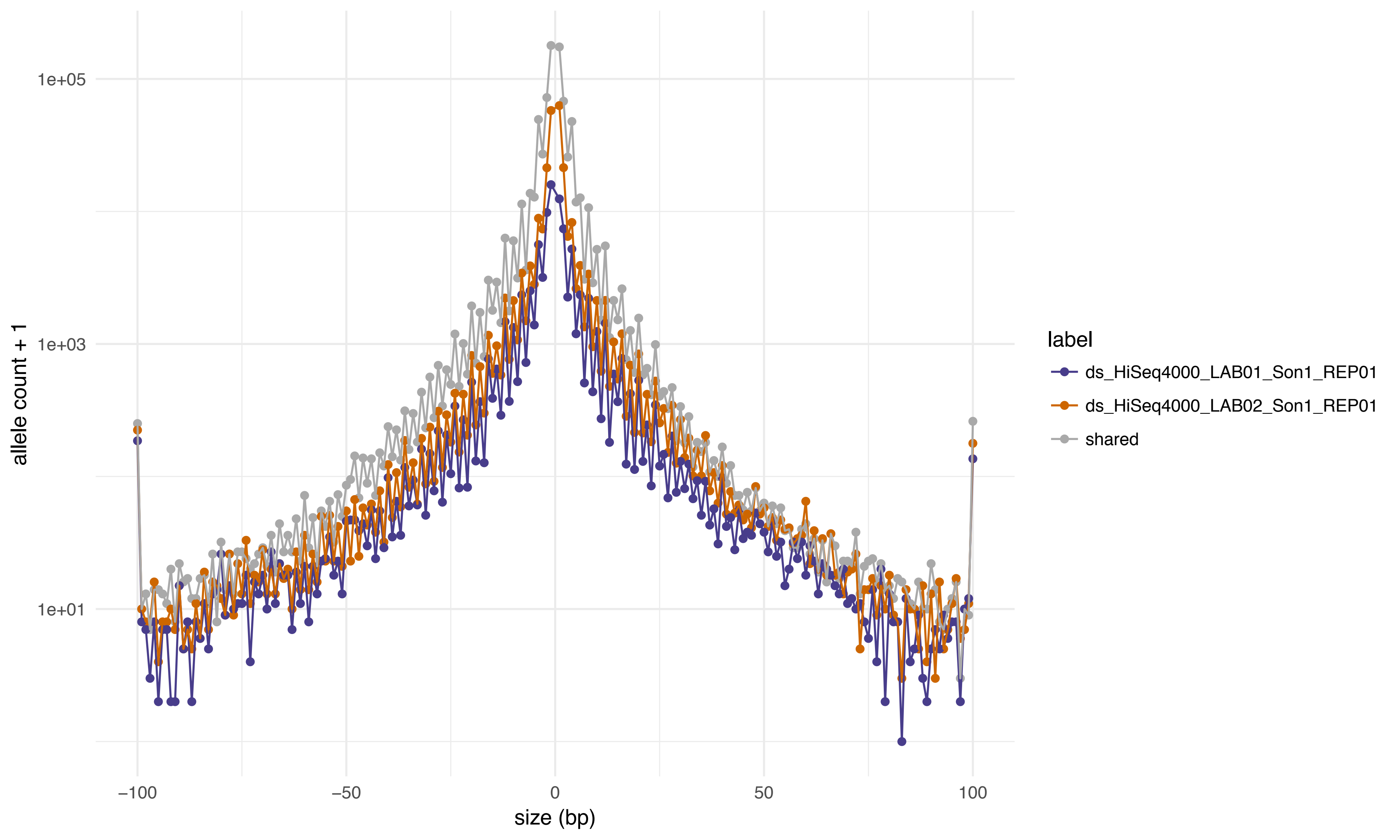

1e+05
1e+03
1e+01
allele count + 1
−100
−50
0
50
100
size (bp)
label
ds_HiSeq4000_LAB01_Son1_REP01
ds_HiSeq4000_LAB02_Son1_REP01
shared

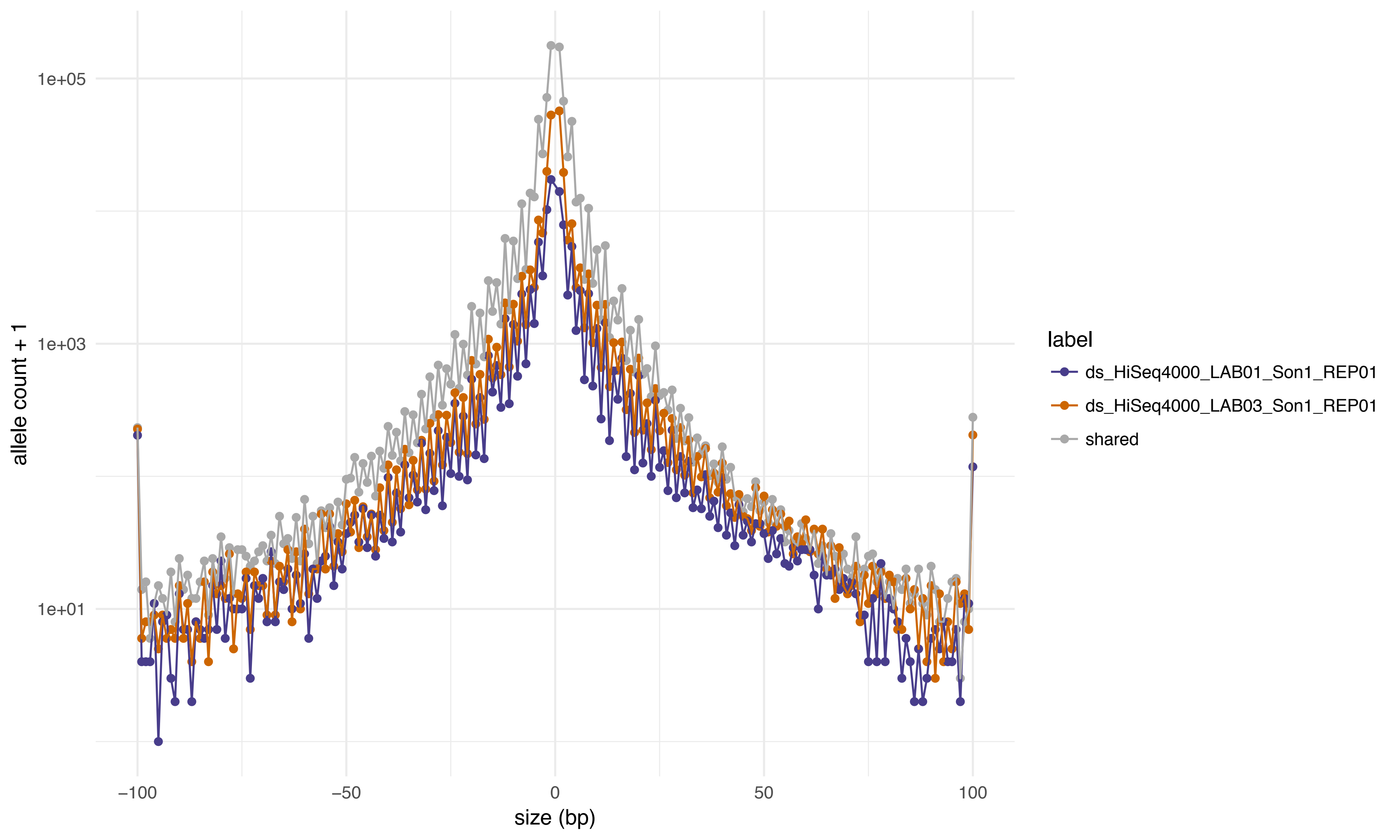
1e+05
1e+03
1e+01
allele count + 1
−100
−50
0
50
100
size (bp)
label
ds_HiSeq4000_LAB01_Son1_REP01
ds_HiSeq4000_LAB03_Son1_REP01
shared

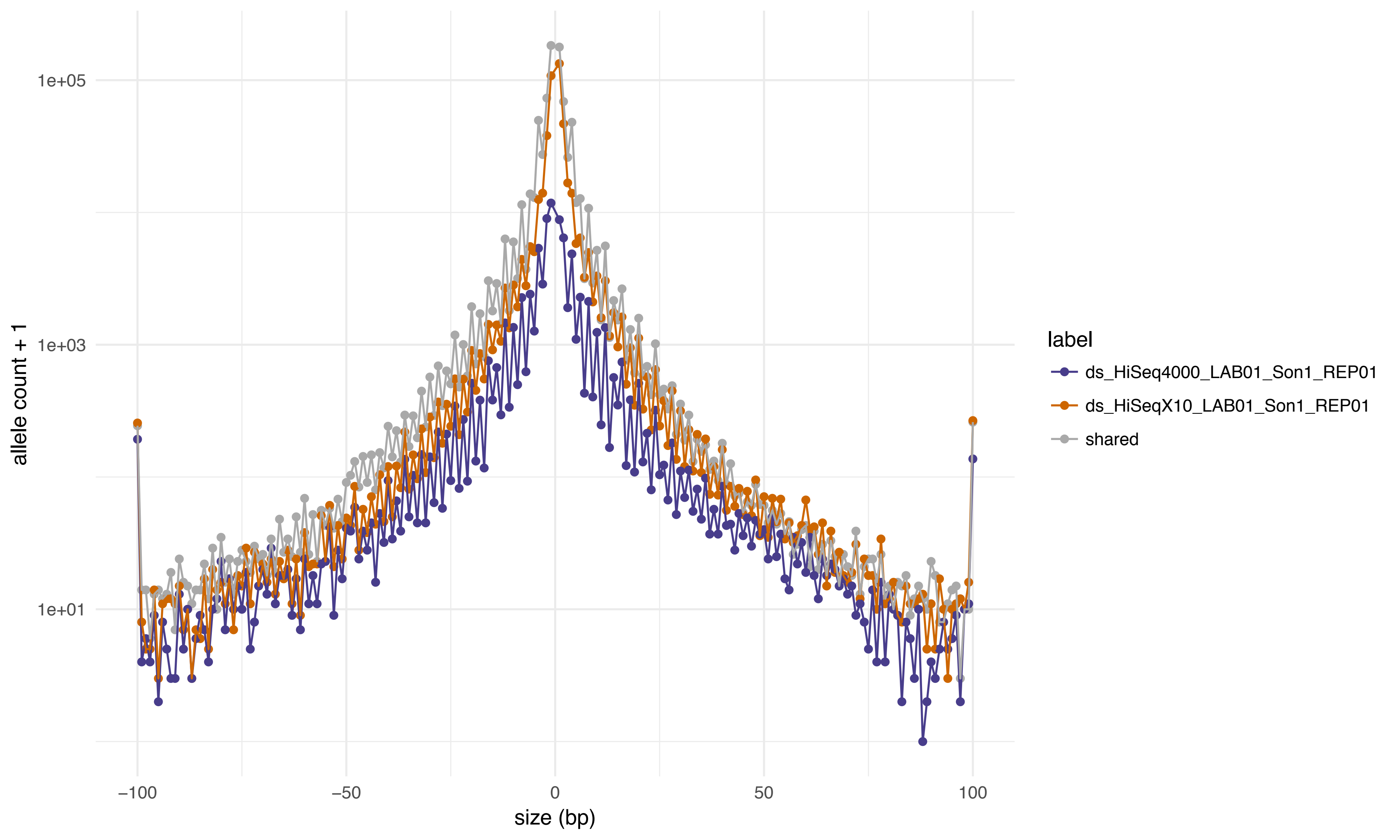
1e+05
1e+03
1e+01
allele count + 1
−100
−50
0
50
100
size (bp)
label
ds_HiSeq4000_LAB01_Son1_REP01
ds_HiSeqX10_LAB01_Son1_REP01
shared

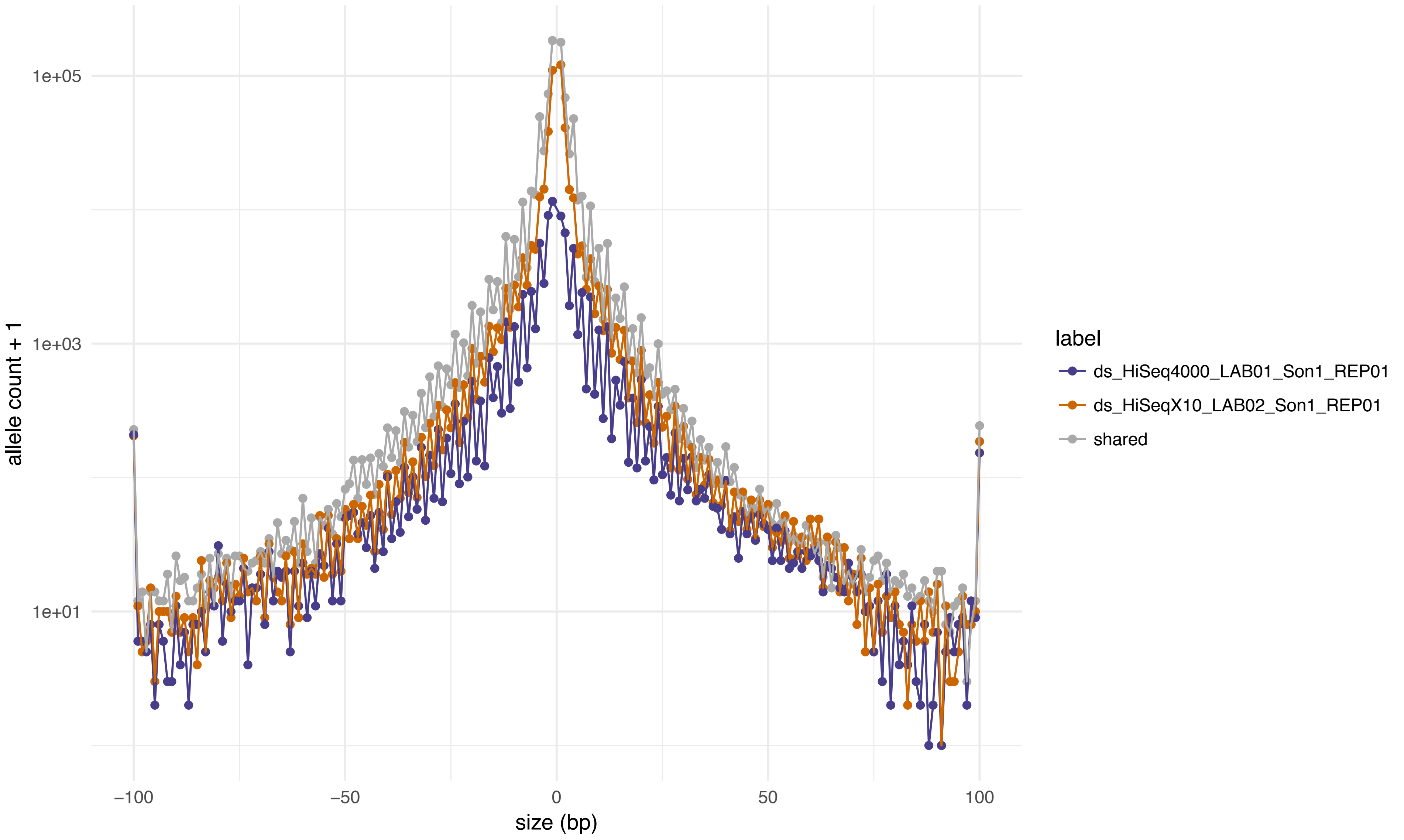
1e+05
1e+03
1e+01
allele count + 1
−100
−50
0
50
100
size (bp)
label
ds_HiSeq4000_LAB01_Son1_REP01
ds_HiSeqX10_LAB02_Son1_REP01
shared

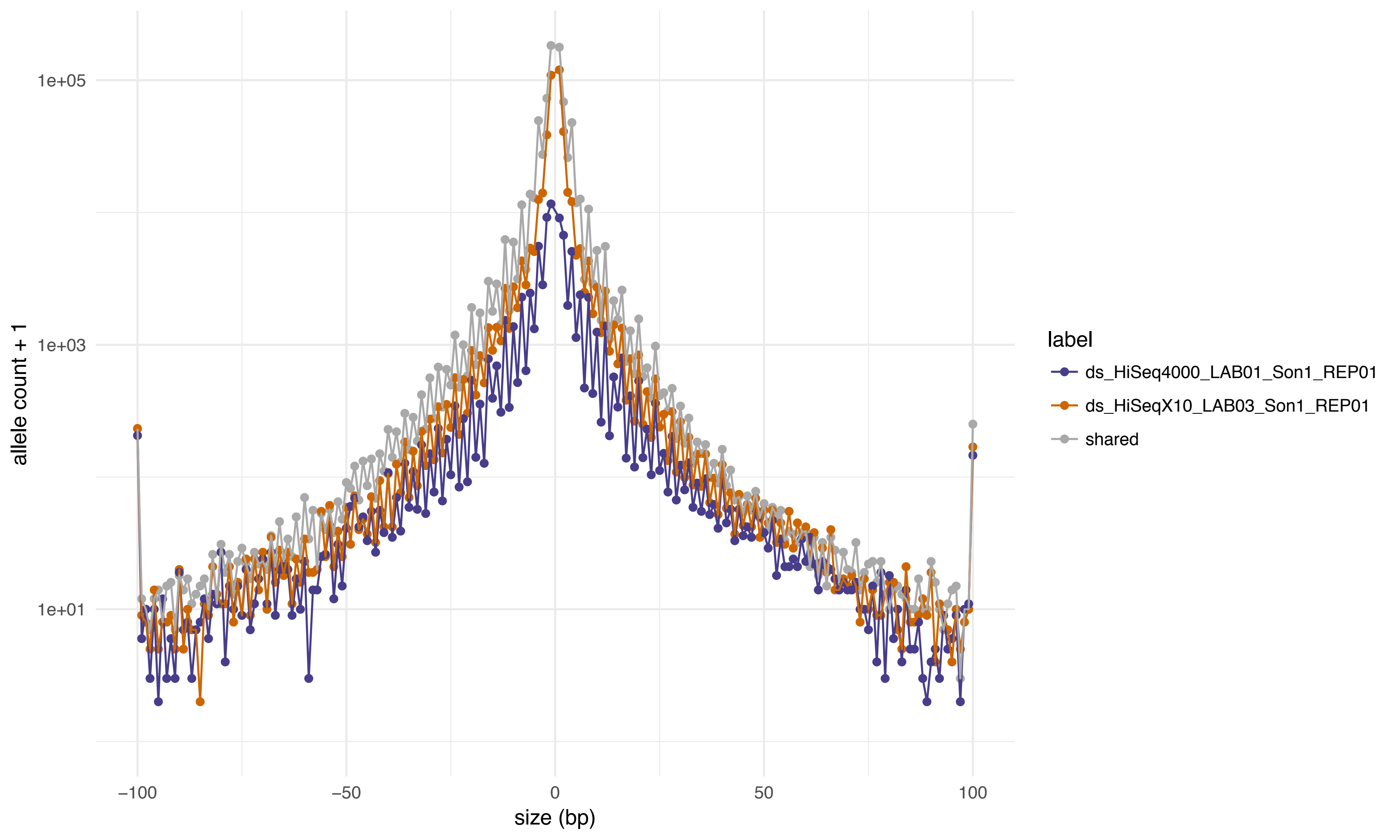

1e+05
1e+03
1e+01
allele count + 1
−100
−50
0
50
100
size (bp)
label
ds_HiSeq4000_LAB01_Son1_REP01
ds_HiSeqX10_LAB03_Son1_REP01
shared

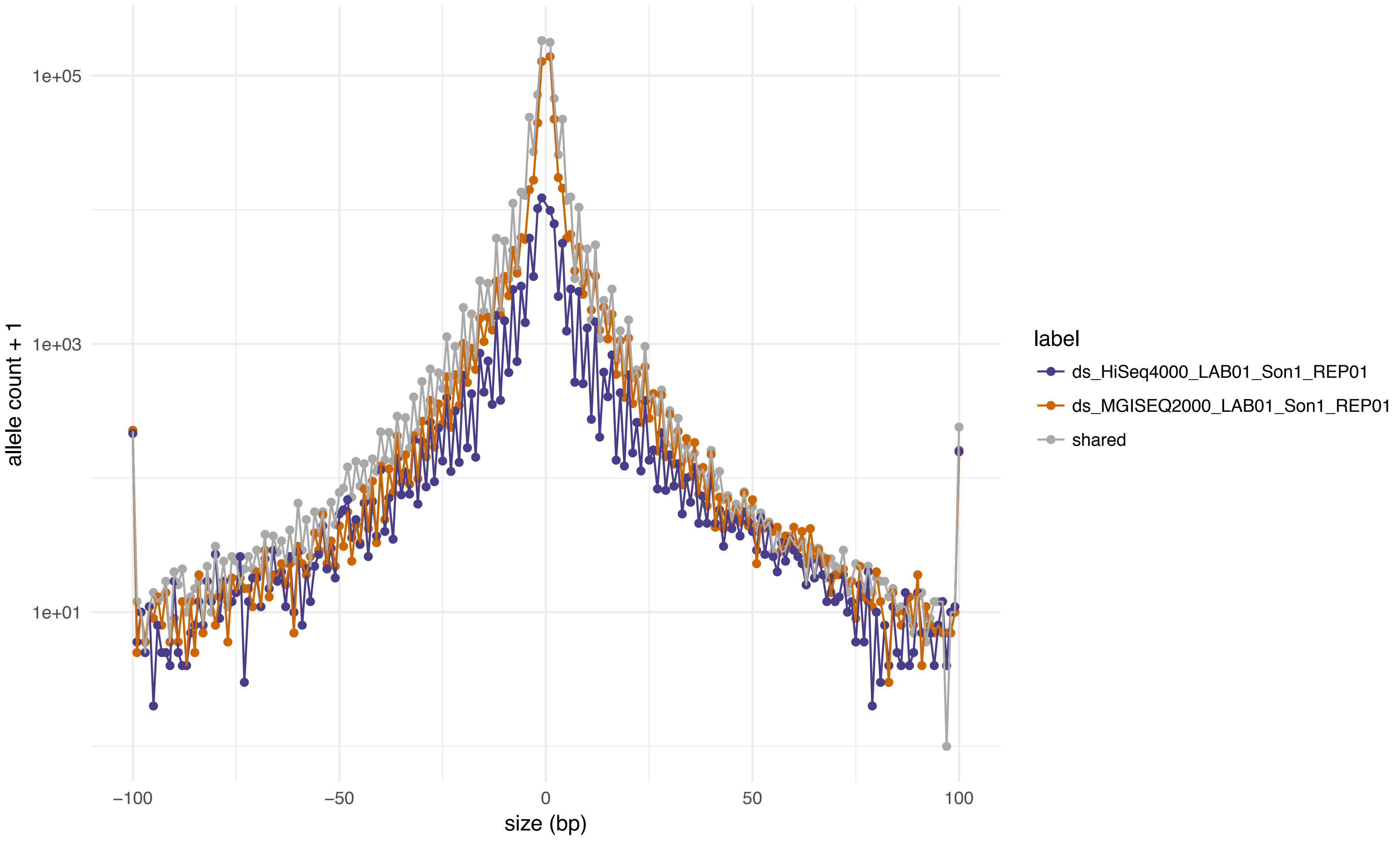

1e+05
1e+03
1e+01
allele count + 1
−100
−50
0
50
100
size (bp)
label
ds_HiSeq4000_LAB01_Son1_REP01
ds_MGISEQ2000_LAB01_Son1_REP01
shared

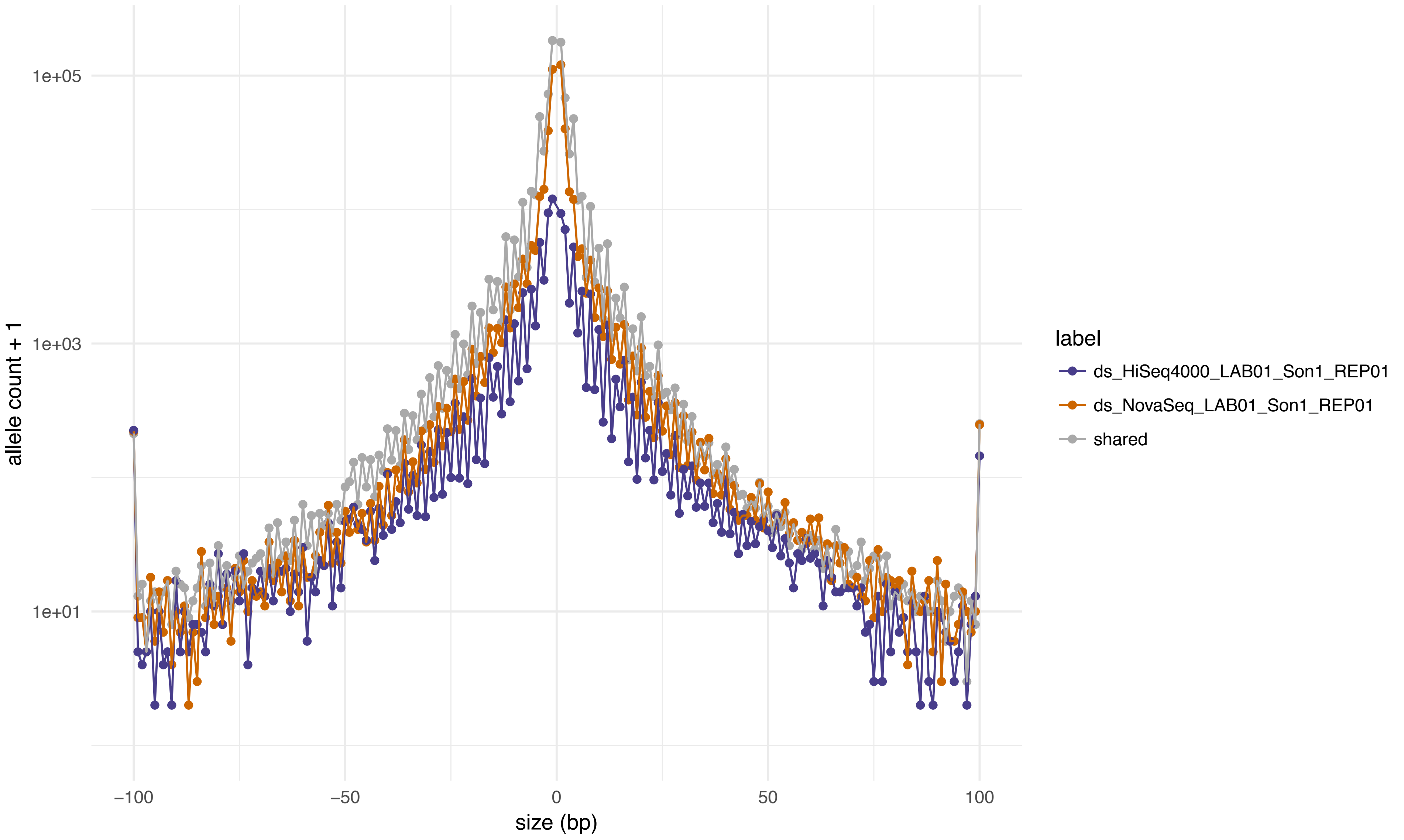
1e+05
1e+03
1e+01
allele count + 1
−100
−50
0
50
100
size (bp)
label
ds_HiSeq4000_LAB01_Son1_REP01
ds_NovaSeq_LAB01_Son1_REP01
shared

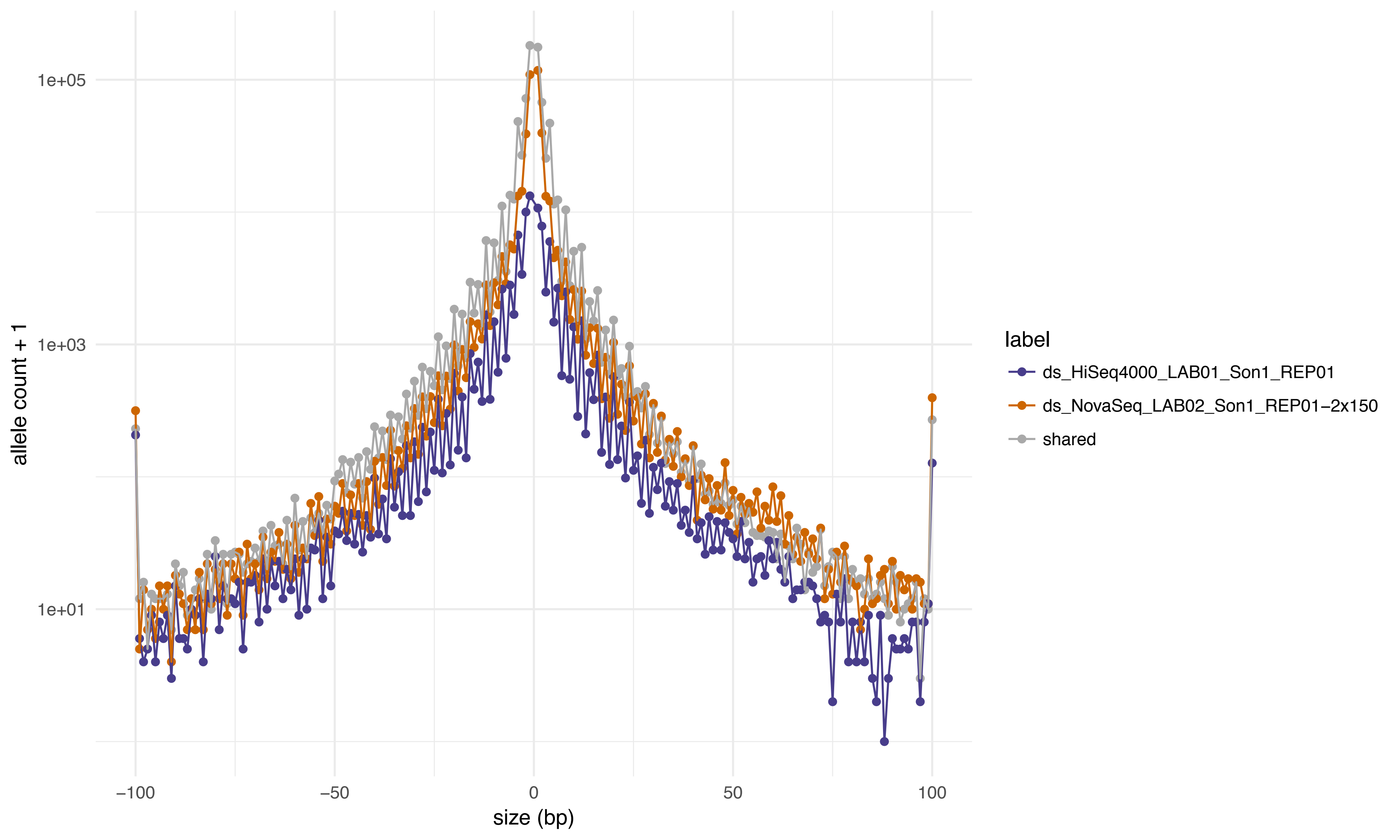
allele count + 1
1e+05
1e+03
1e+01
−100
−50
0
50
100
size (bp)
label
ds_HiSeq4000_LAB01_Son1_REP01
ds_NovaSeq_LAB02_Son1_REP01−2x150
shared

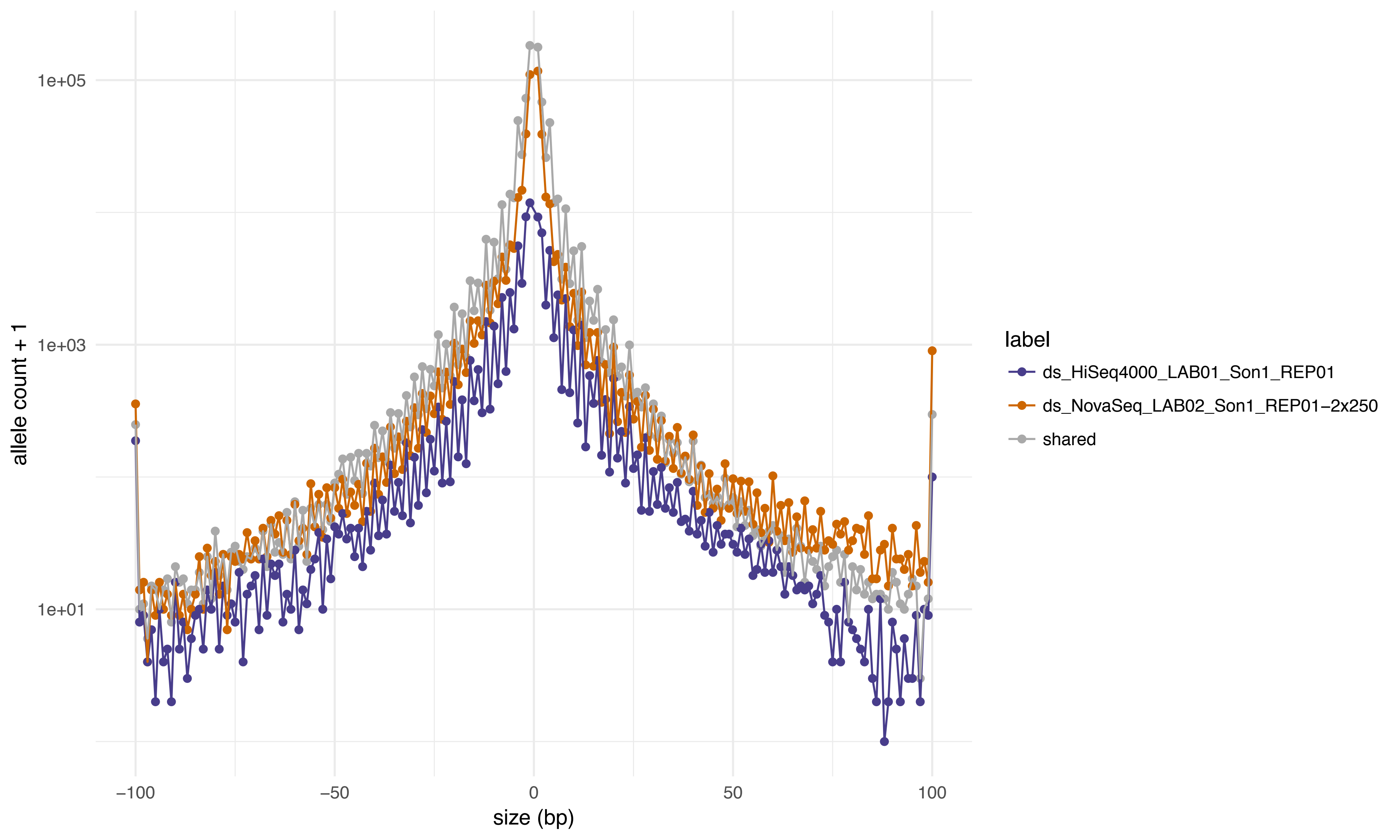
1e+05
1e+03
1e+01
allele count + 1
−100
−50
0
50
100
size (bp)
label
ds_HiSeq4000_LAB01_Son1_REP01
ds_NovaSeq_LAB02_Son1_REP01−2x250
shared

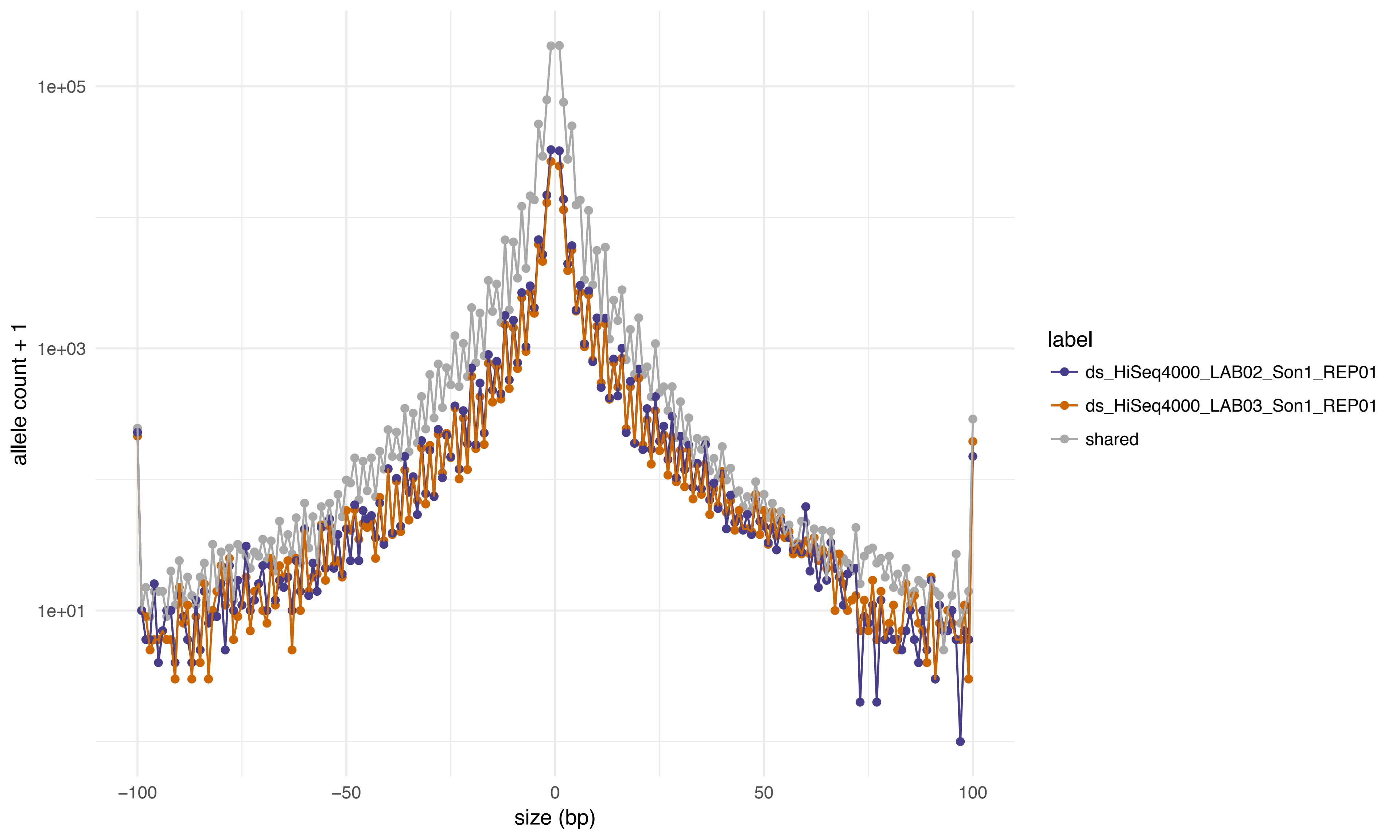

1e+05
1e+03
1e+01
allele count + 1
−100
−50
0
50
100
size (bp)
label
ds_HiSeq4000_LAB02_Son1_REP01
ds_HiSeq4000_LAB03_Son1_REP01
shared

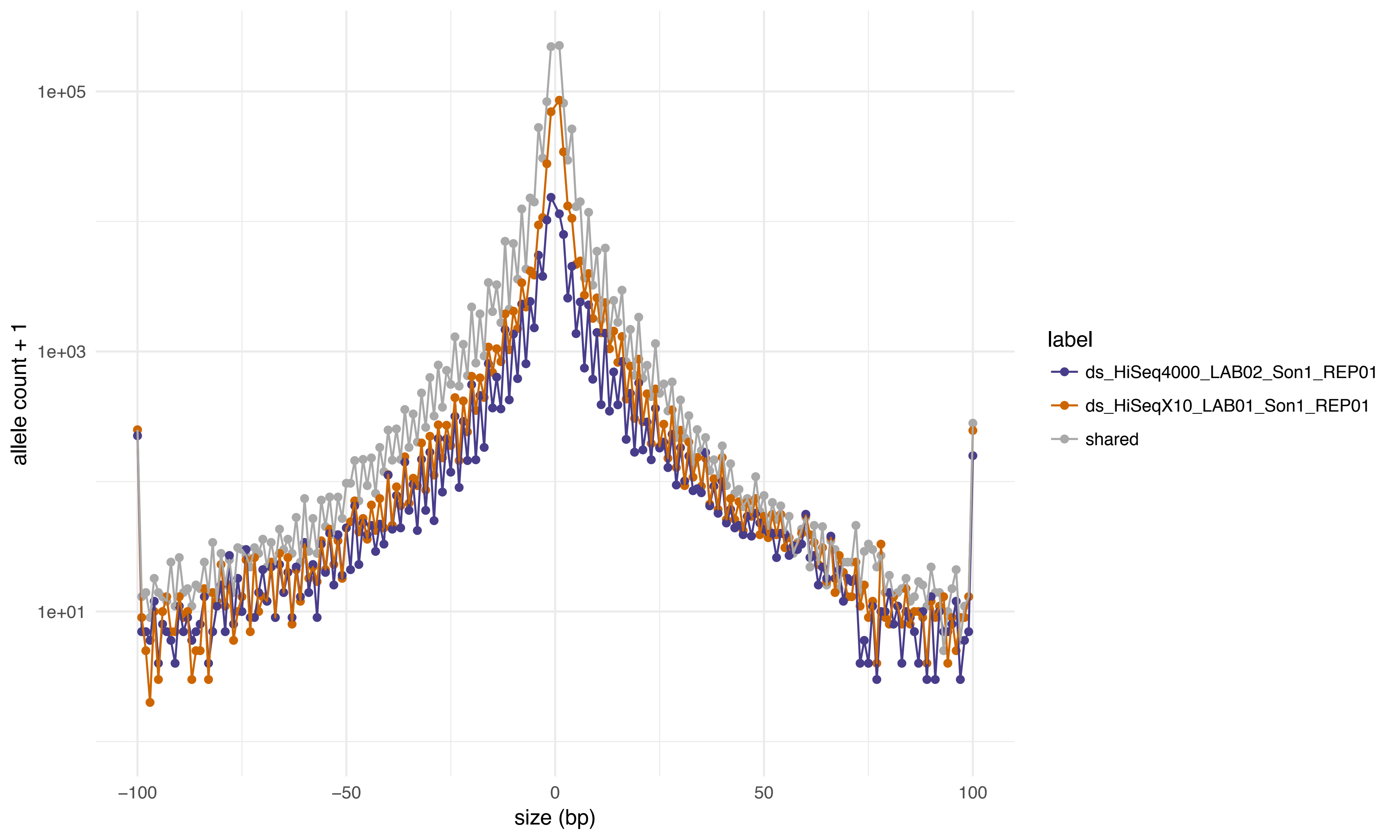
allele count + 1
1e+05
1e+03
1e+01
−100
−50
0
50
100
size (bp)
label
ds_HiSeq4000_LAB02_Son1_REP01
ds_HiSeqX10_LAB01_Son1_REP01
shared

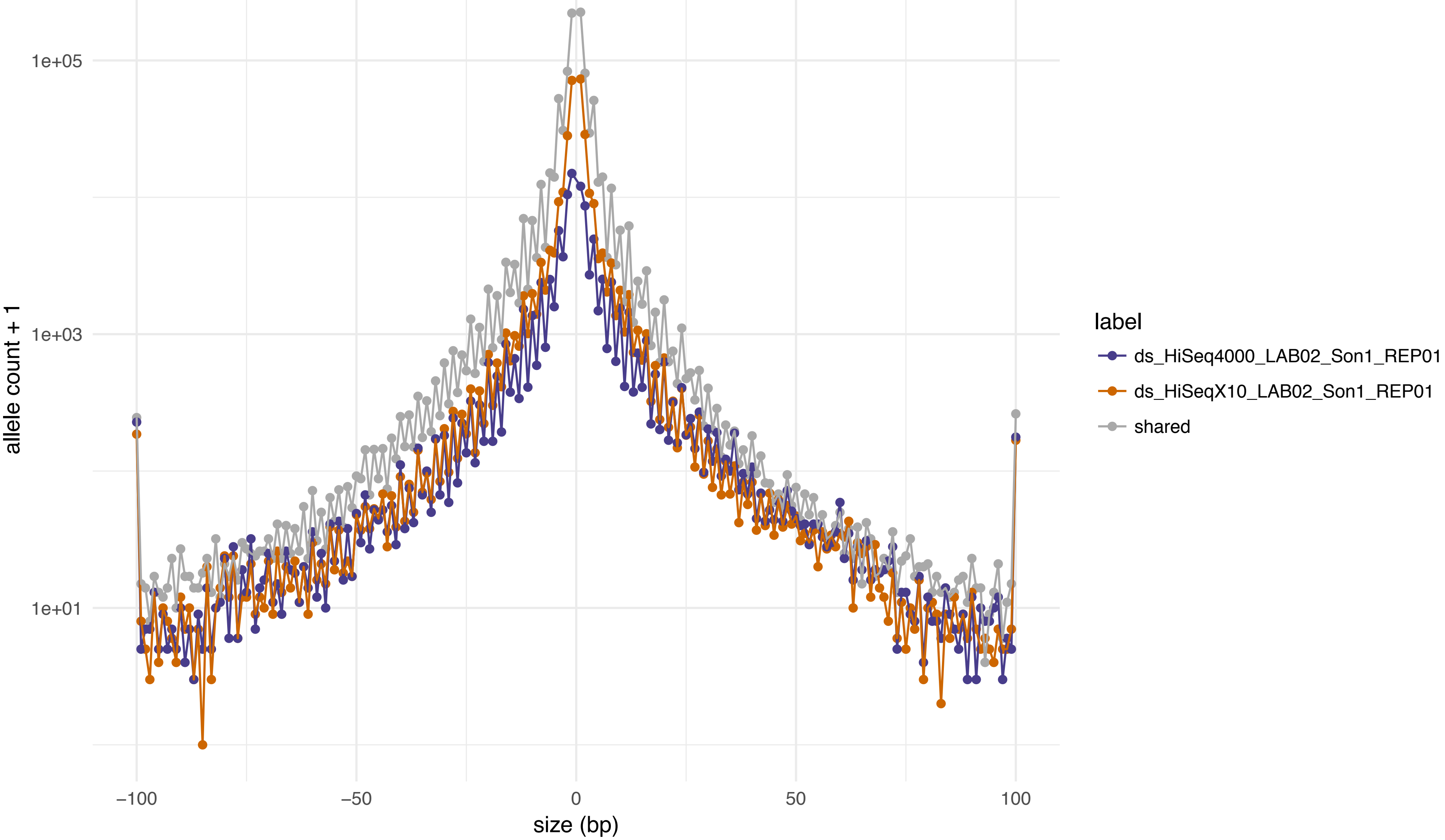

1e+05
1e+03
1e+01
allele count + 1
−100
−50
0
50
100
size (bp)
label
ds_HiSeq4000_LAB02_Son1_REP01
ds_HiSeqX10_LAB02_Son1_REP01
shared

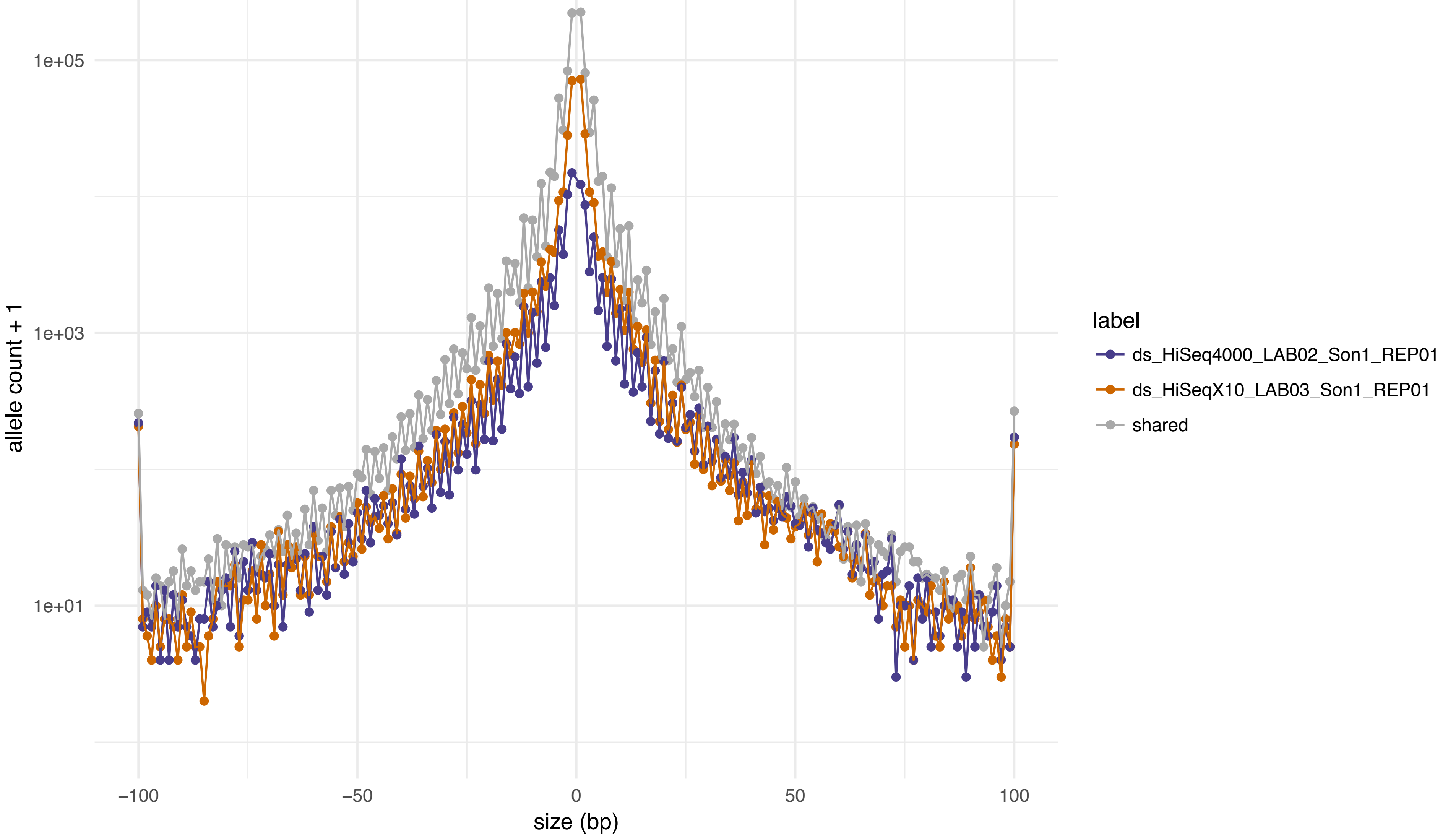
1e+05
1e+03
1e+01
allele count + 1
−100
−50
0
50
100
size (bp)
label
ds_HiSeq4000_LAB02_Son1_REP01
ds_HiSeqX10_LAB03_Son1_REP01
shared

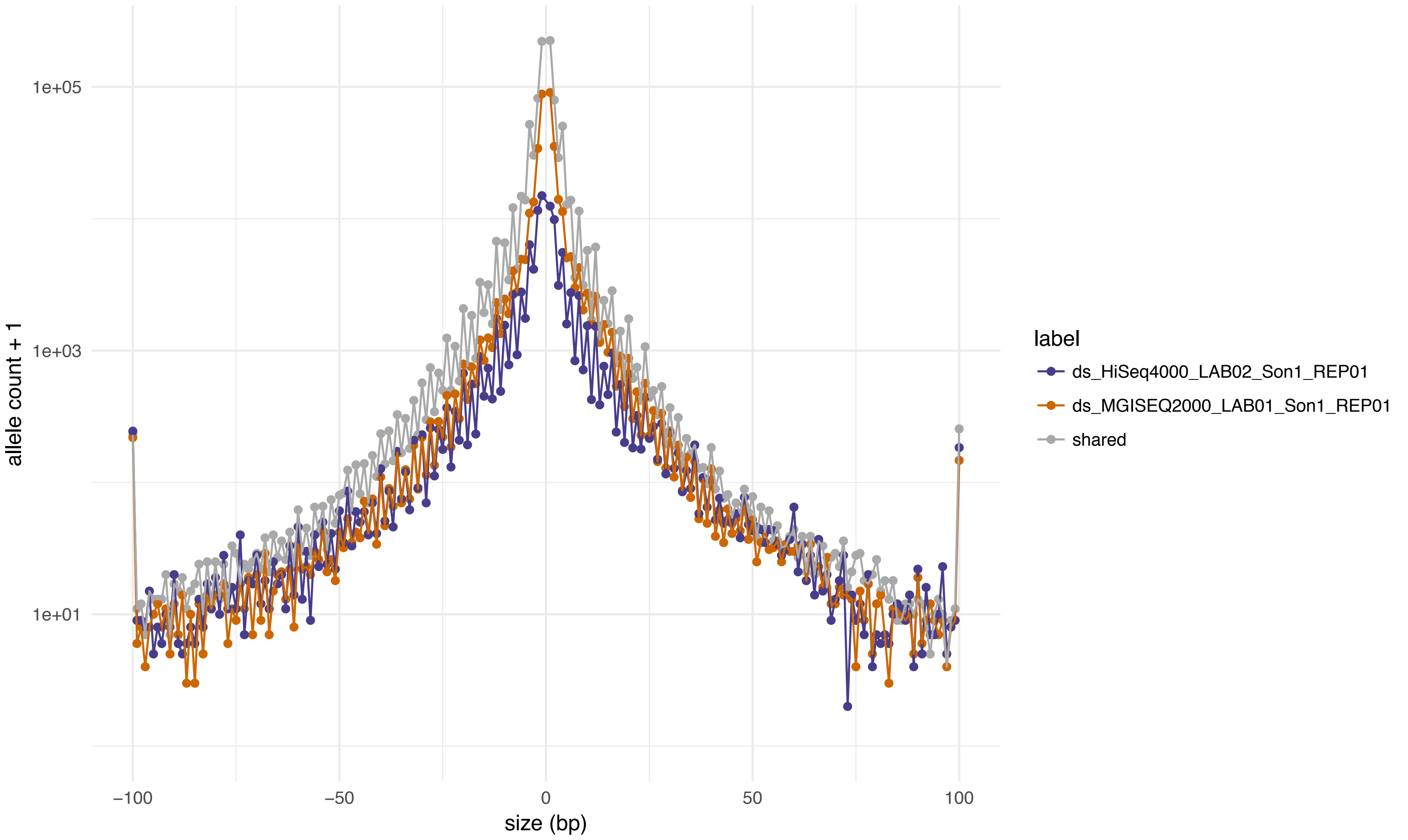

1e+05
1e+03
1e+01
allele count + 1
−100
−50
0
50
100
size (bp)
label
ds_HiSeq4000_LAB02_Son1_REP01
ds_MGISEQ2000_LAB01_Son1_REP01
shared

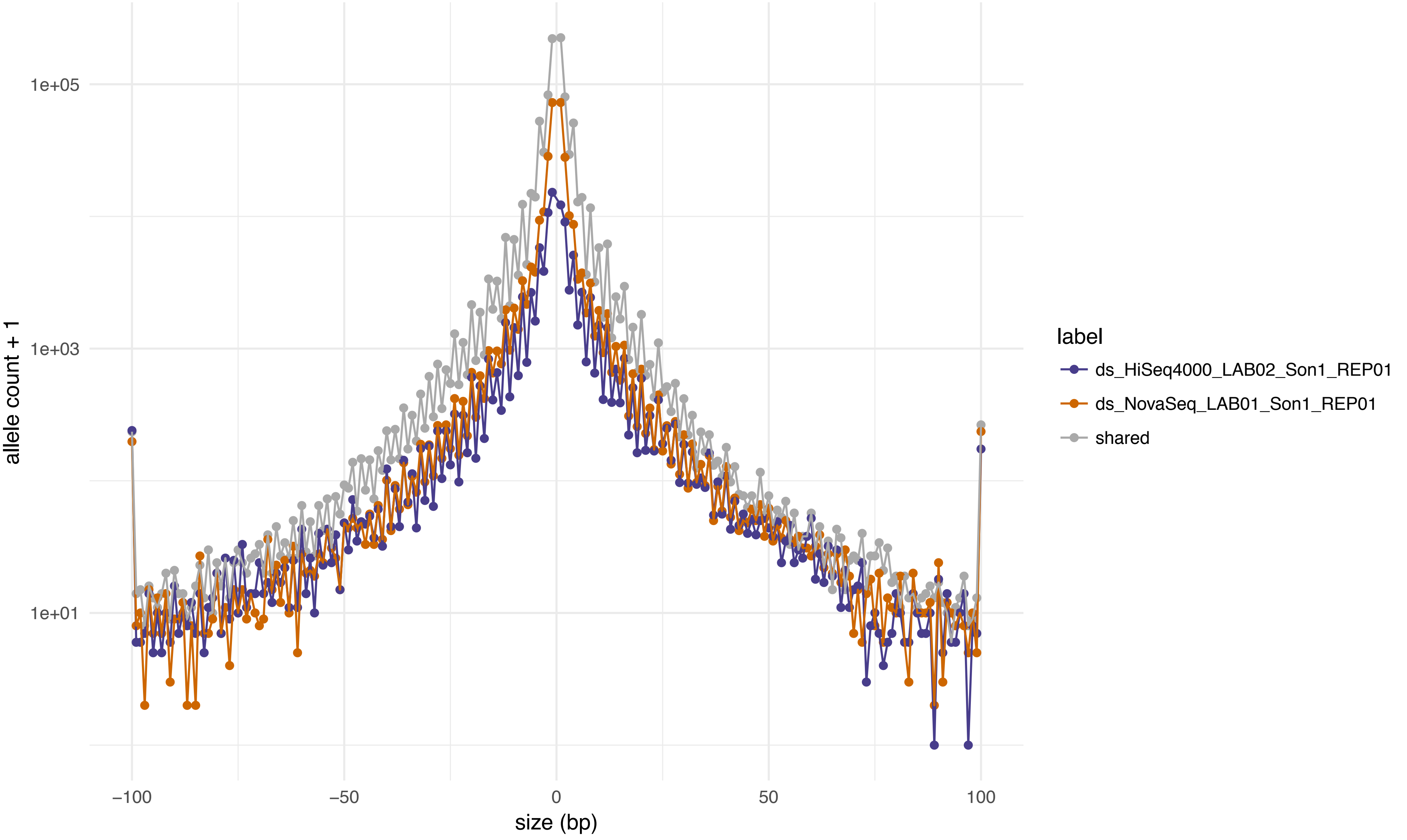

1e+05
1e+03
1e+01
allele count + 1
−100
−50
0
50
100
size (bp)
label
ds_HiSeq4000_LAB02_Son1_REP01
ds_NovaSeq_LAB01_Son1_REP01
shared

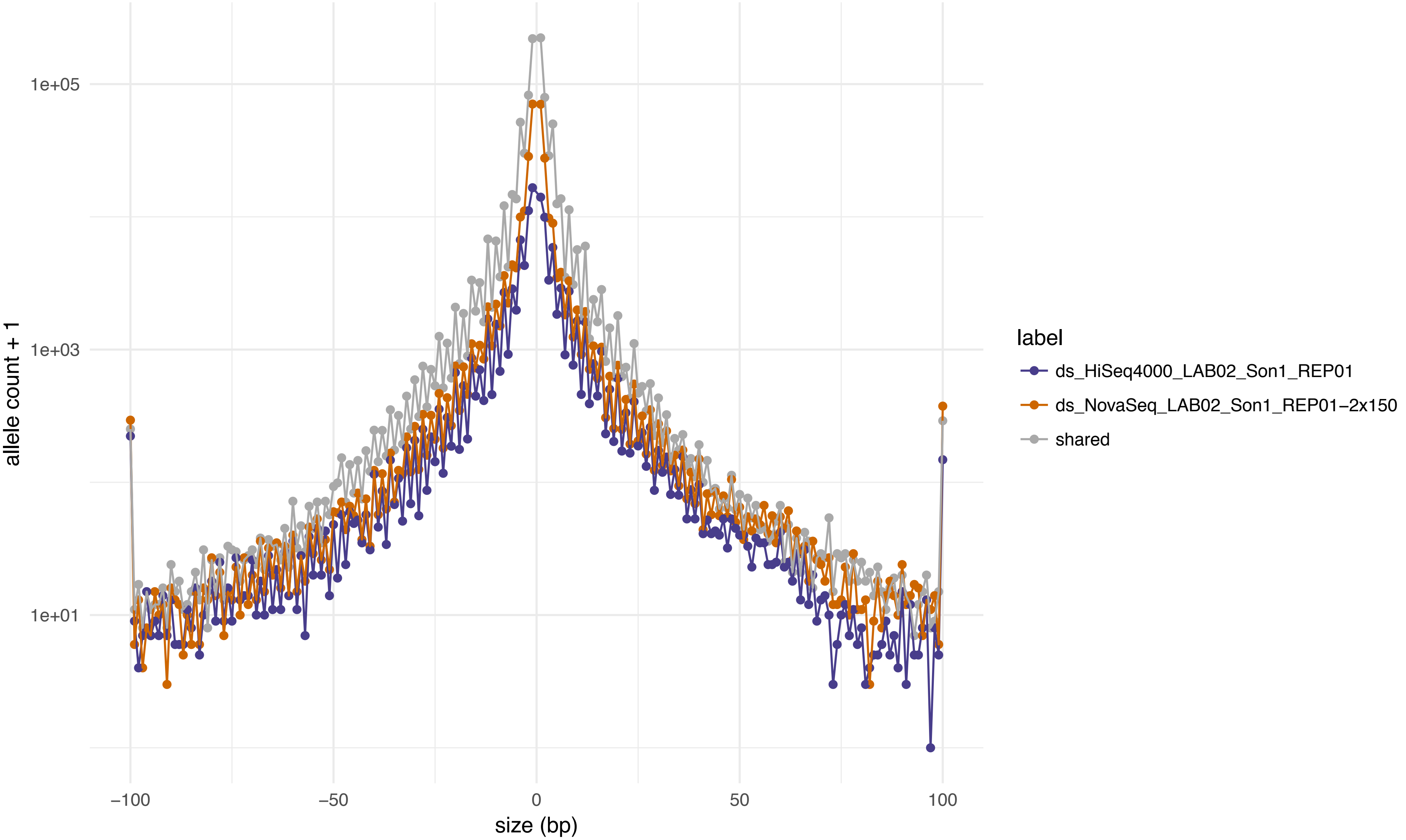
1e+05
1e+03
1e+01
allele count + 1
−100
−50
0
50
100
size (bp)
label
ds_HiSeq4000_LAB02_Son1_REP01
ds_NovaSeq_LAB02_Son1_REP01−2x150
shared

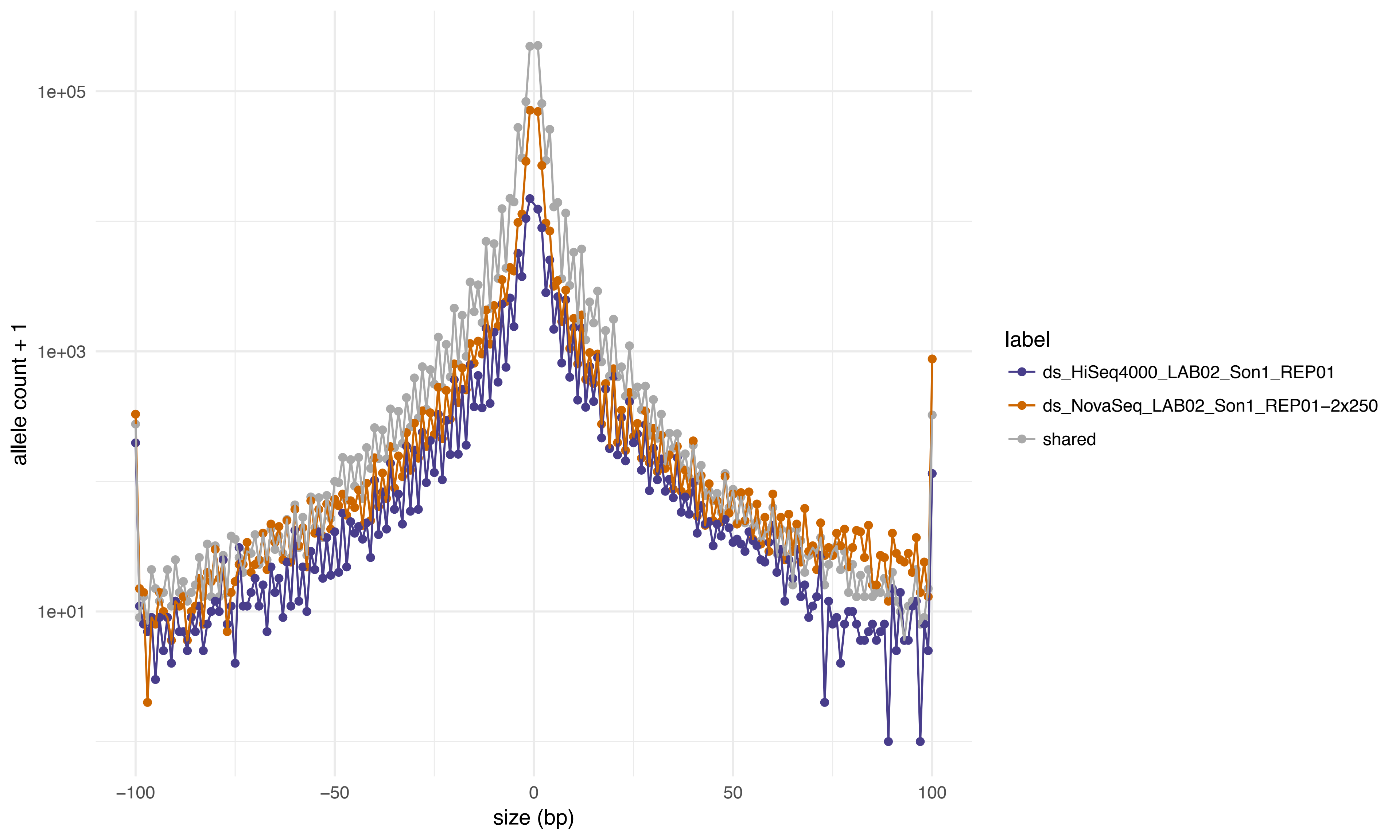
1e+05
1e+03
1e+01
allele count + 1
−100
−50
0
50
100
size (bp)
label
ds_HiSeq4000_LAB02_Son1_REP01
ds_NovaSeq_LAB02_Son1_REP01−2x250
shared

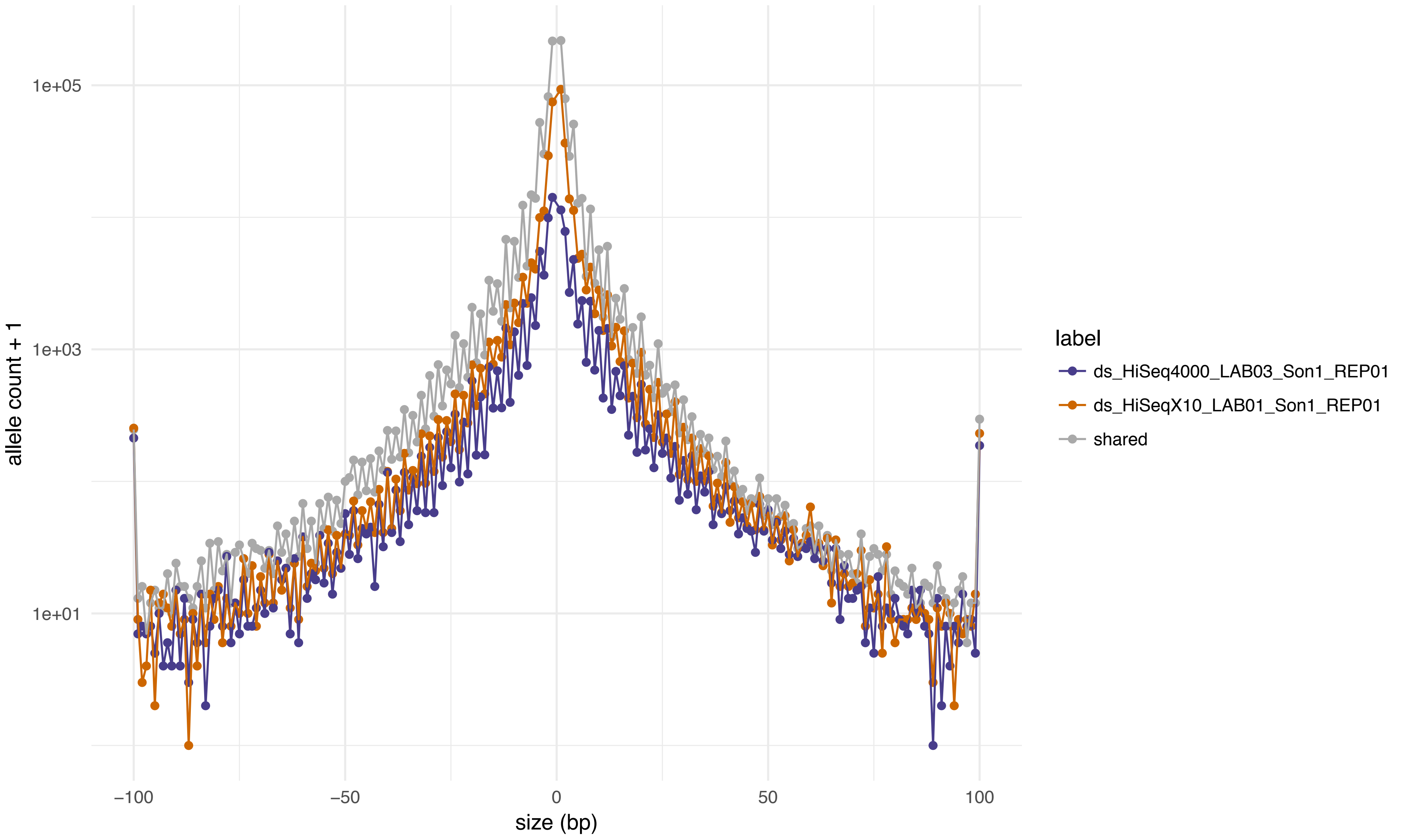

1e+05
1e+03
1e+01
allele count + 1
−100
−50
0
50
100
size (bp)
label
ds_HiSeq4000_LAB03_Son1_REP01
ds_HiSeqX10_LAB01_Son1_REP01
shared

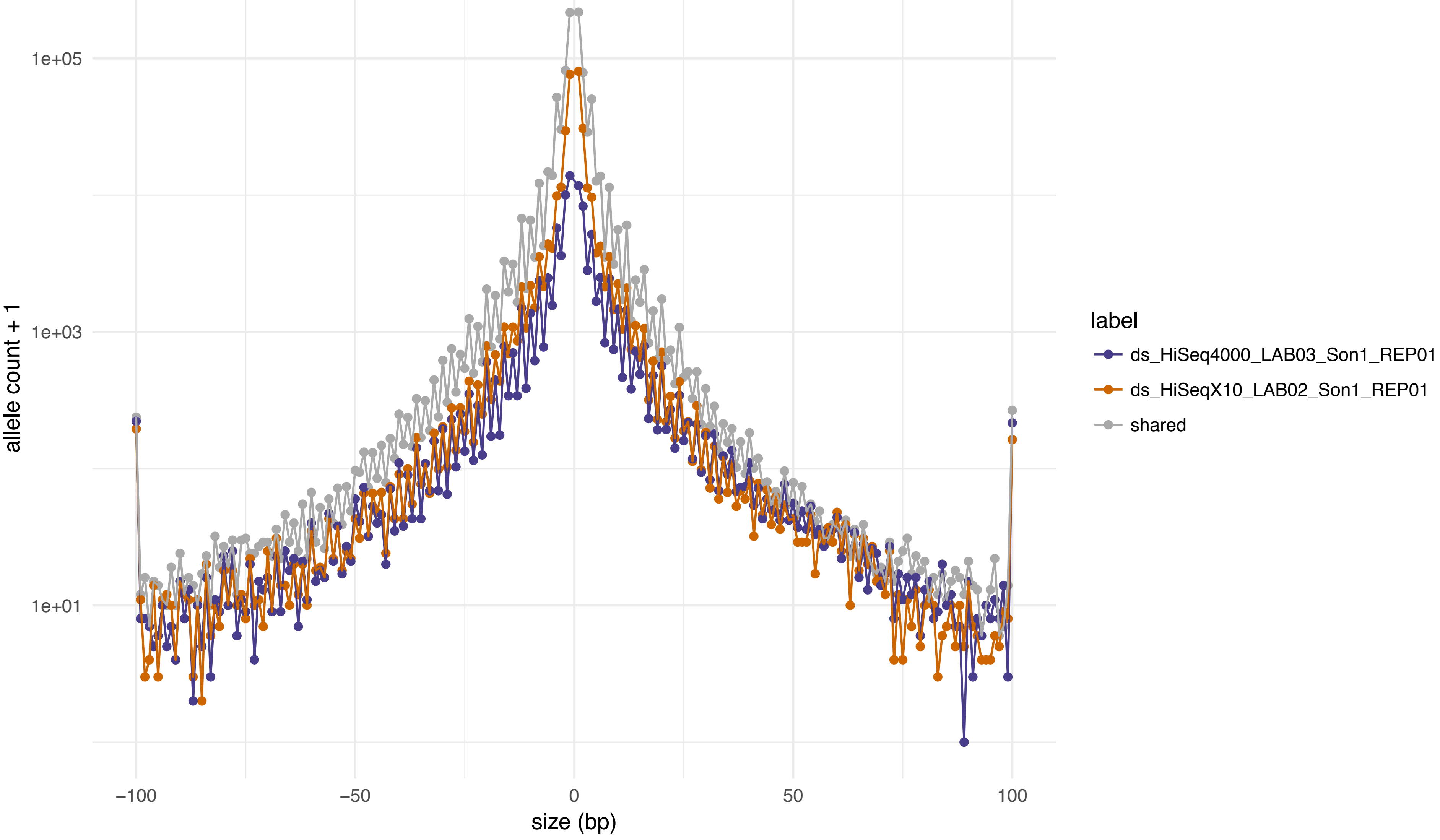
1e+05
1e+03
1e+01
allele count + 1
−100
−50
0
50
100
size (bp)
label
ds_HiSeq4000_LAB03_Son1_REP01
ds_HiSeqX10_LAB02_Son1_REP01
shared

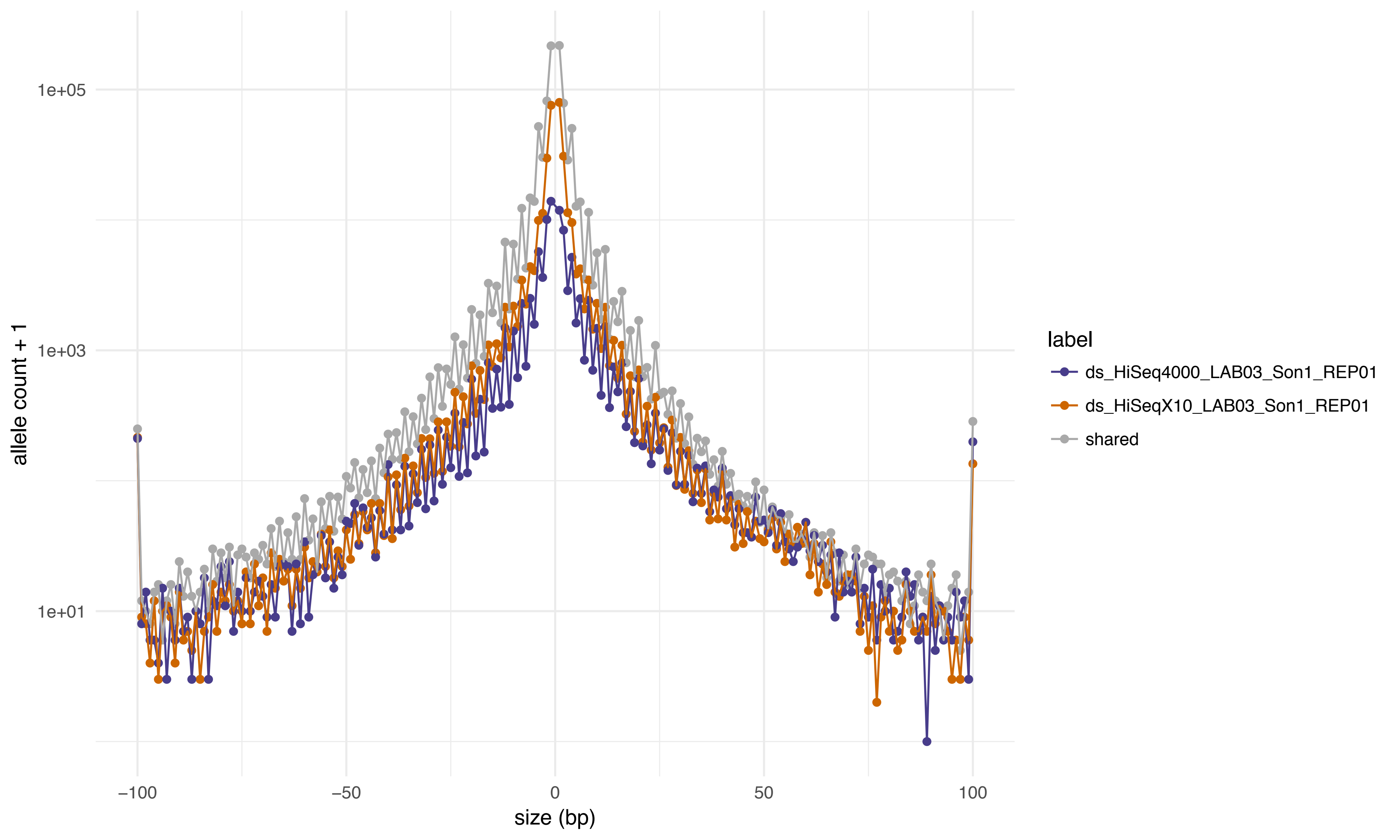

1e+05
1e+03
1e+01
allele count + 1
−100
−50
0
50
100
size (bp)
label
ds_HiSeq4000_LAB03_Son1_REP01
ds_HiSeqX10_LAB03_Son1_REP01
shared

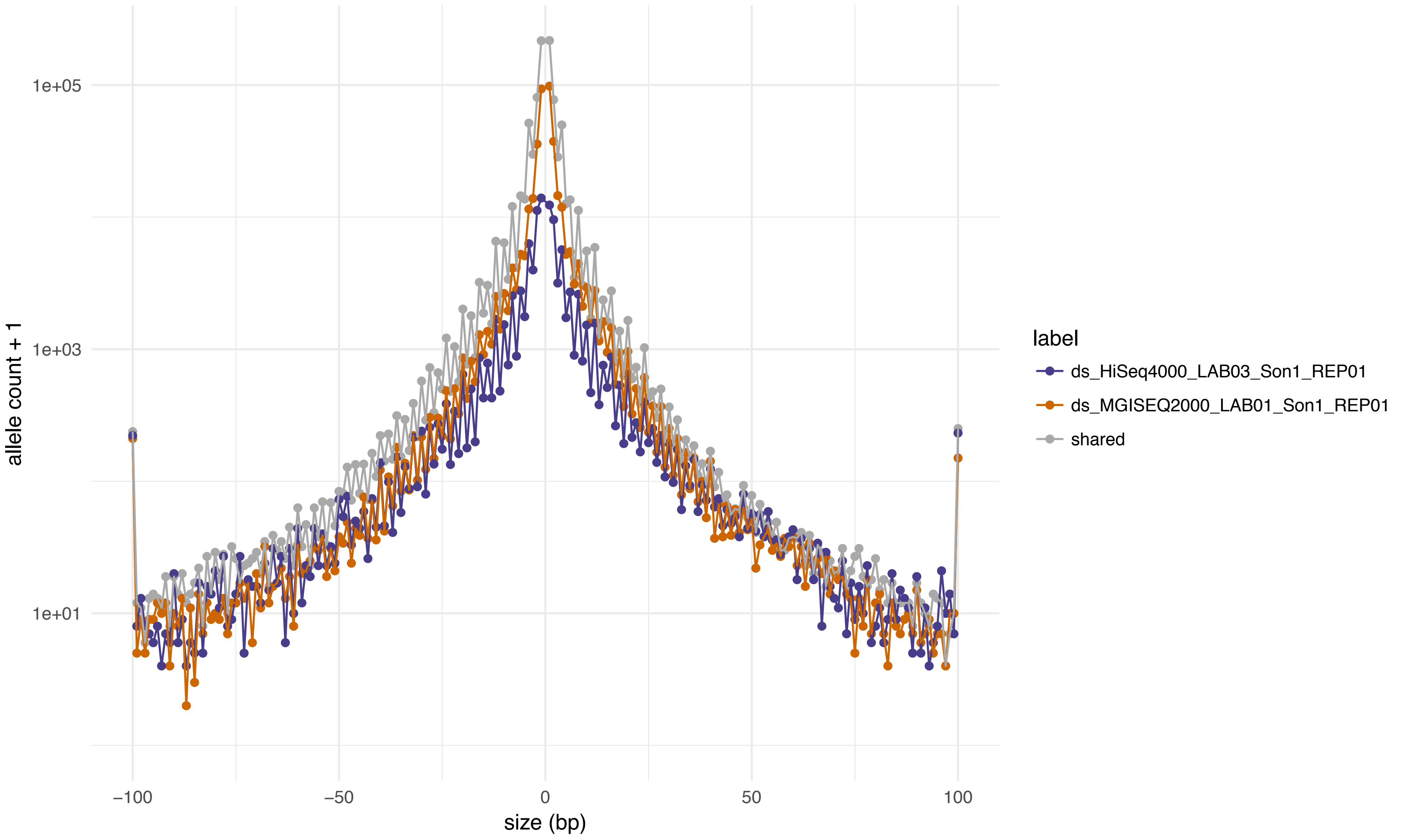
1e+05
1e+03
1e+01
allele count + 1
−100
−50
0
50
100
size (bp)
label
ds_HiSeq4000_LAB03_Son1_REP01
ds_MGISEQ2000_LAB01_Son1_REP01
shared

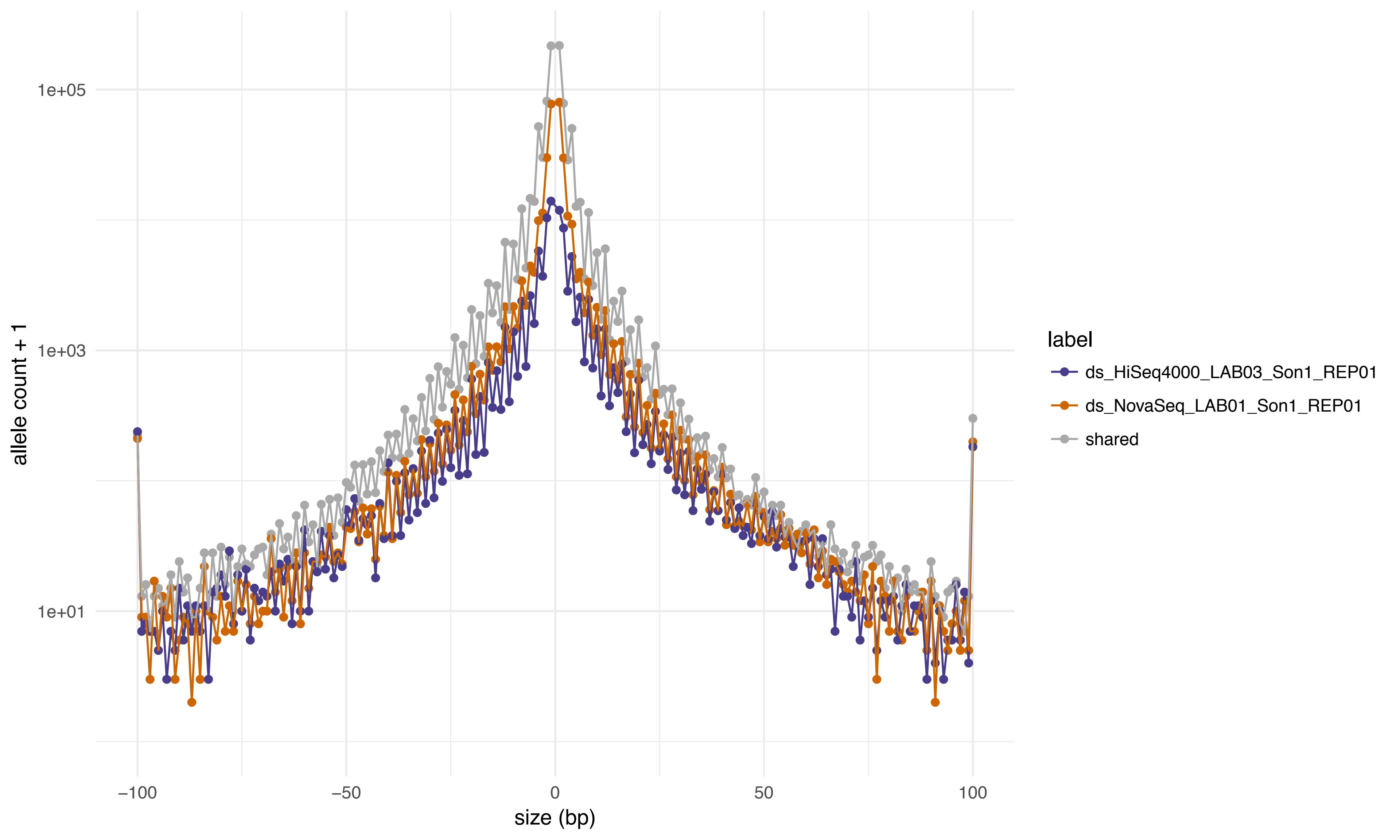
1e+05
1e+03
1e+01
allele count + 1
−100
−50
0
50
100
size (bp)
label
ds_HiSeq4000_LAB03_Son1_REP01
ds_NovaSeq_LAB01_Son1_REP01
shared

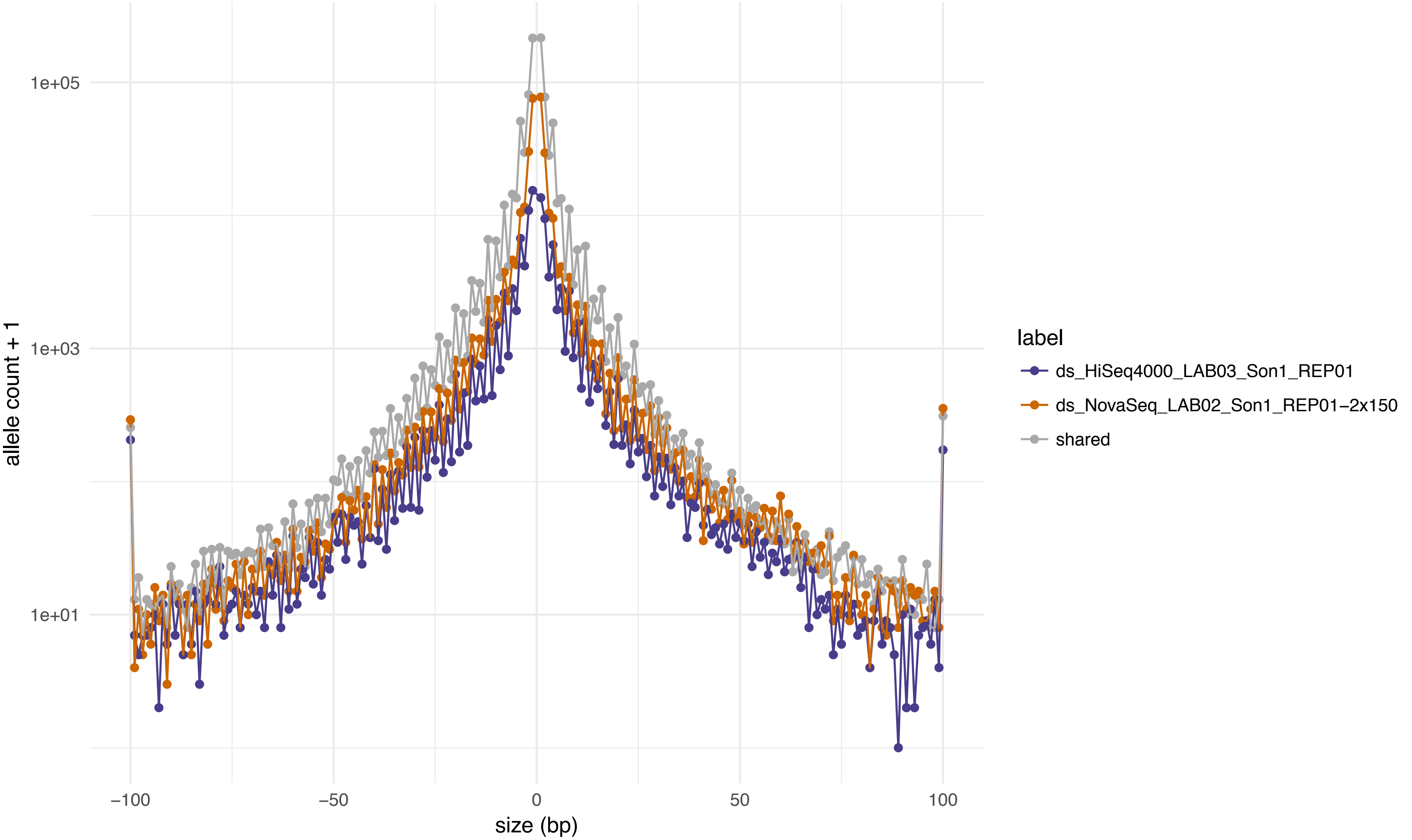
allele count + 1
1e+05
1e+03
1e+01
−100
−50
0
50
100
size (bp)
label
ds_HiSeq4000_LAB03_Son1_REP01
ds_NovaSeq_LAB02_Son1_REP01−2x150
shared

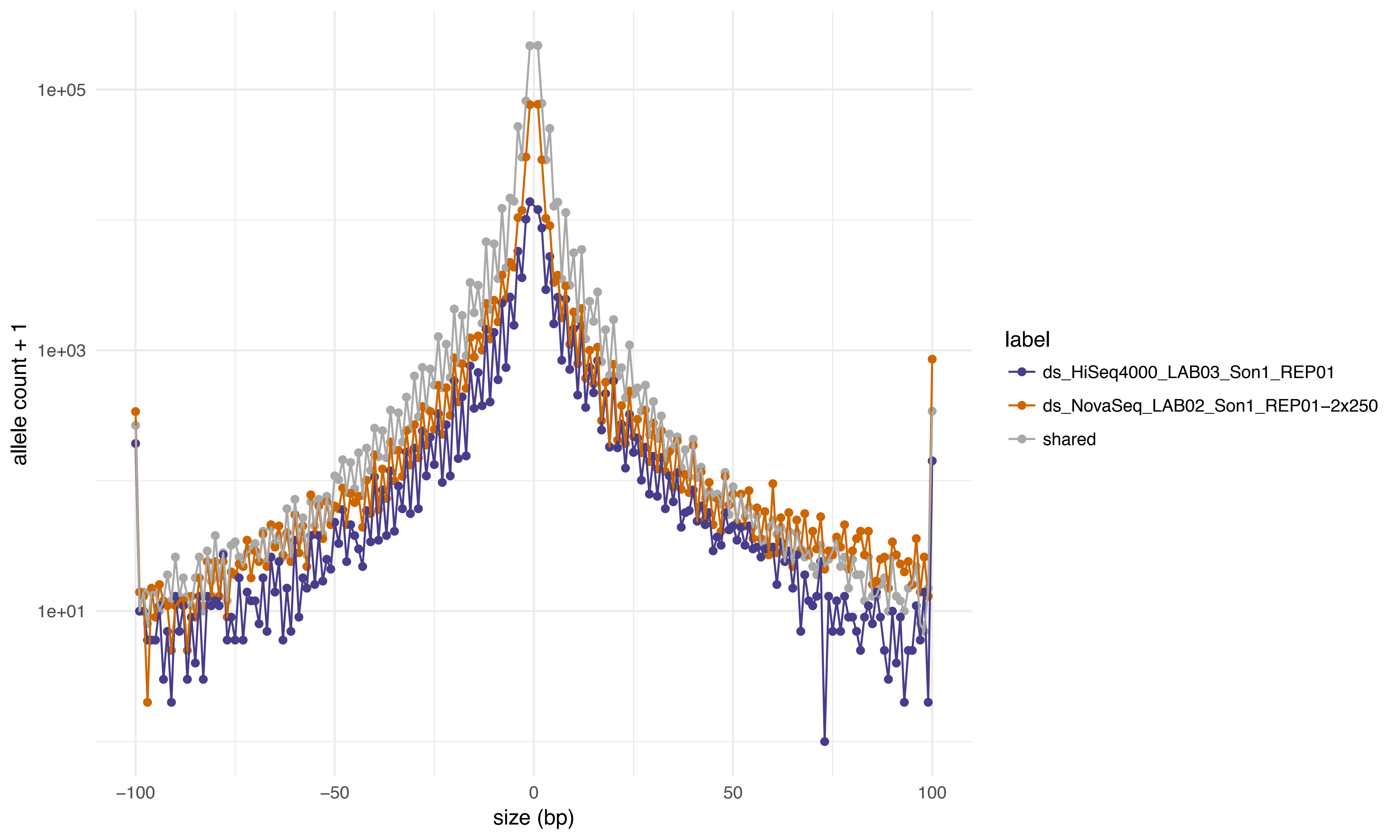
1e+05
1e+03
1e+01
allele count + 1
−100
−50
0
50
100
size (bp)
label
ds_HiSeq4000_LAB03_Son1_REP01
ds_NovaSeq_LAB02_Son1_REP01−2x250
shared

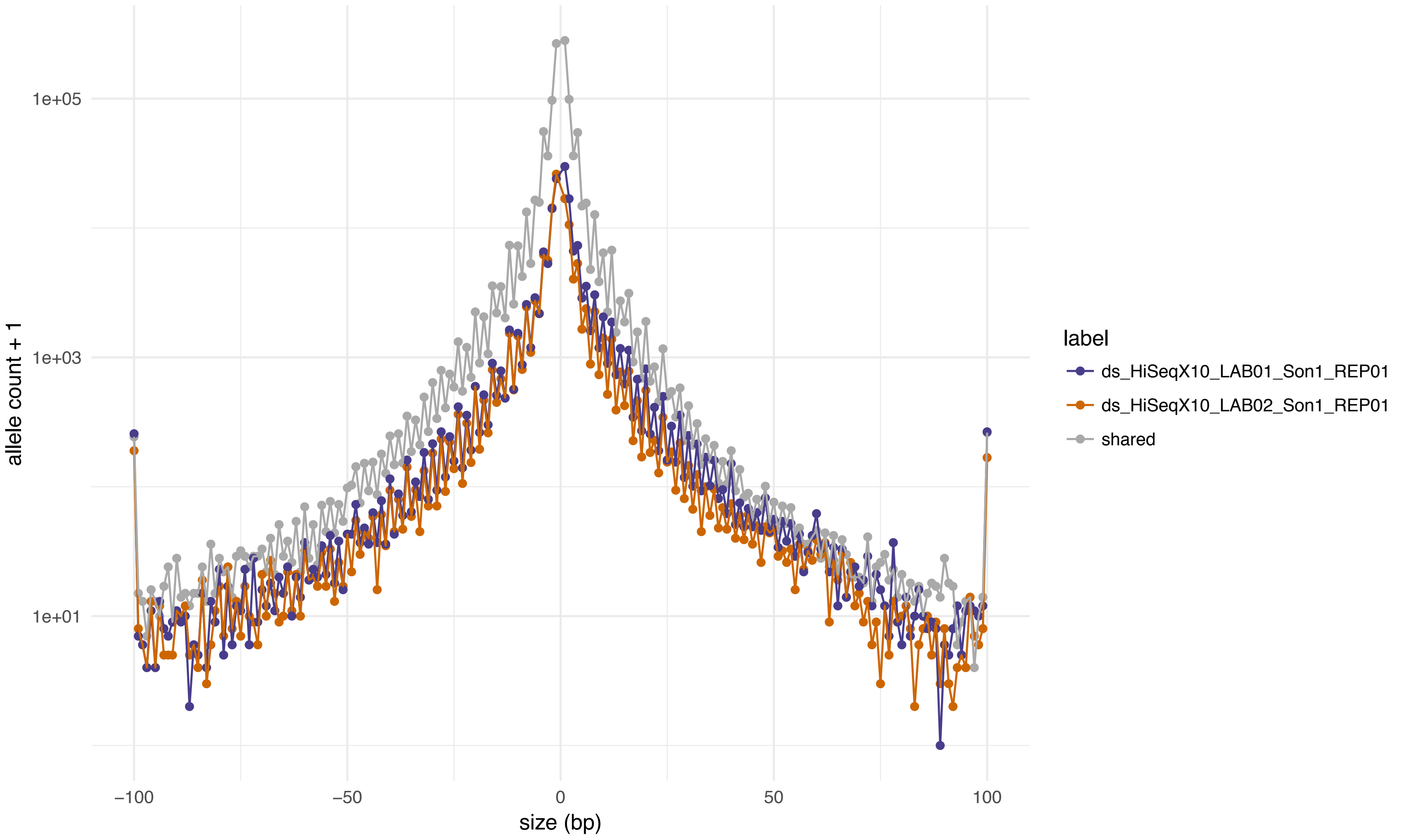

1e+05
1e+03
1e+01
allele count + 1
−100
−50
0
50
100
size (bp)
label
ds_HiSeqX10_LAB01_Son1_REP01
ds_HiSeqX10_LAB02_Son1_REP01
shared

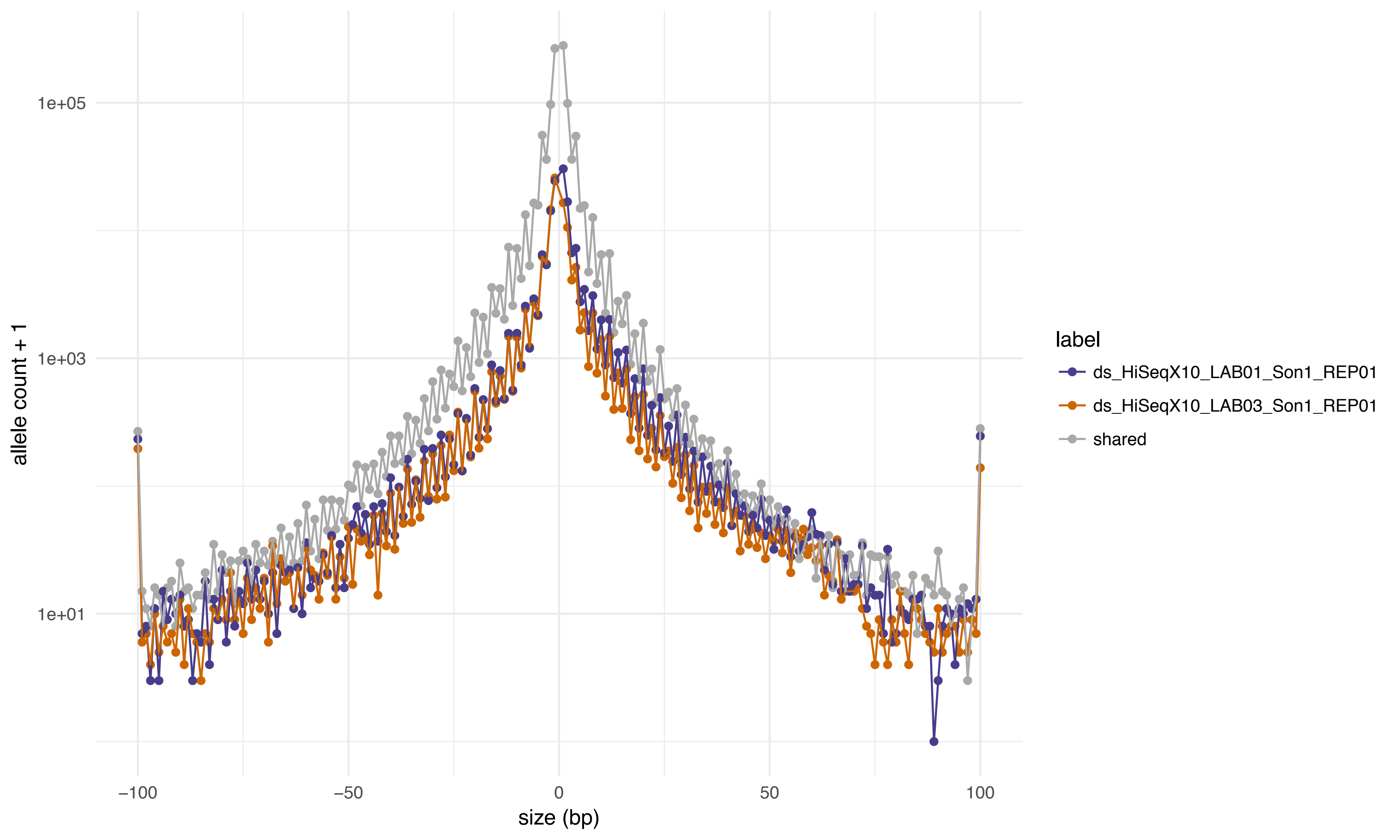
1e+05
1e+03
1e+01
allele count + 1
−100
−50
0
50
100
size (bp)
label
ds_HiSeqX10_LAB01_Son1_REP01
ds_HiSeqX10_LAB03_Son1_REP01
shared

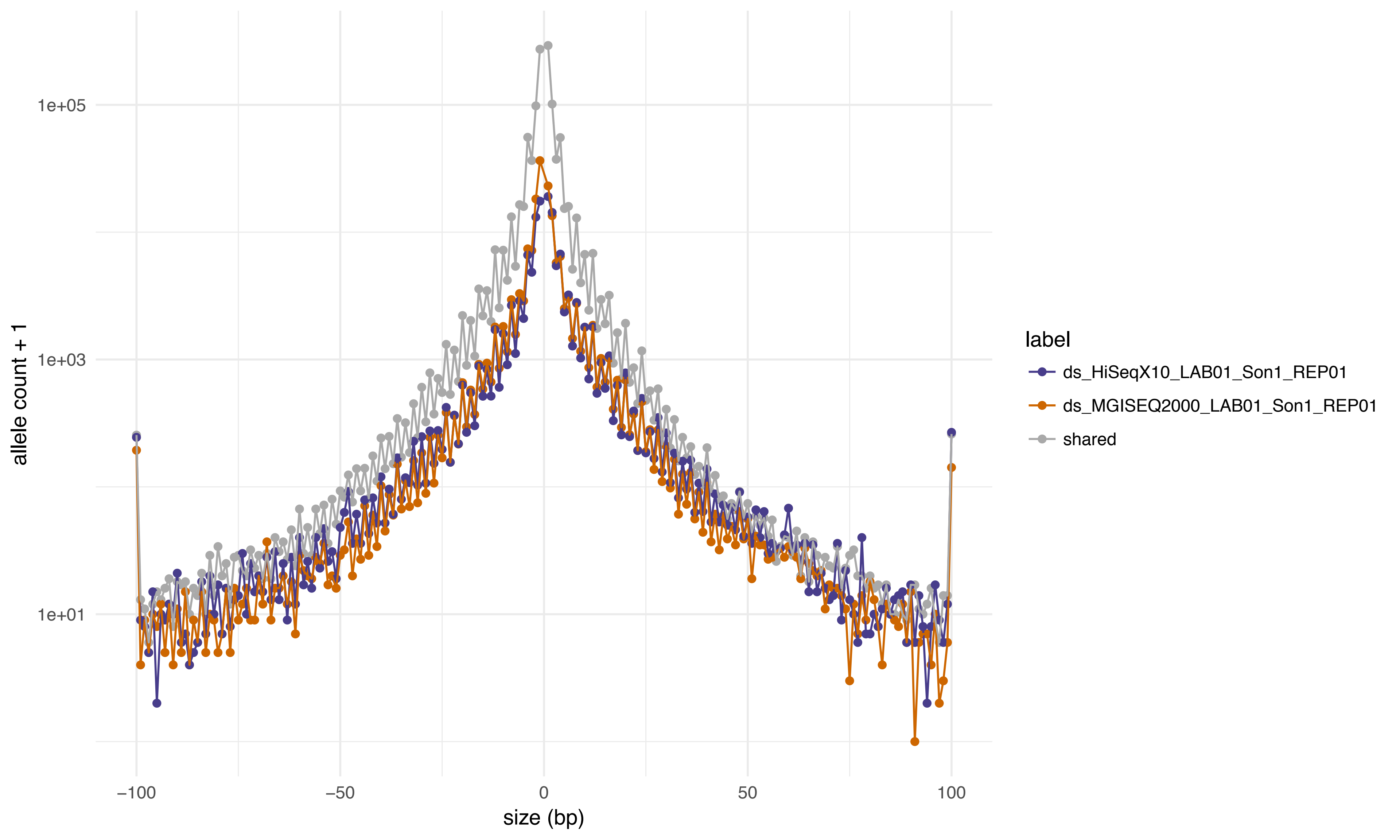

allele count + 1
1e+05
1e+03
1e+01
−100
−50
0
50
100
size (bp)
label
ds_HiSeqX10_LAB01_Son1_REP01
ds_MGISEQ2000_LAB01_Son1_REP01
shared

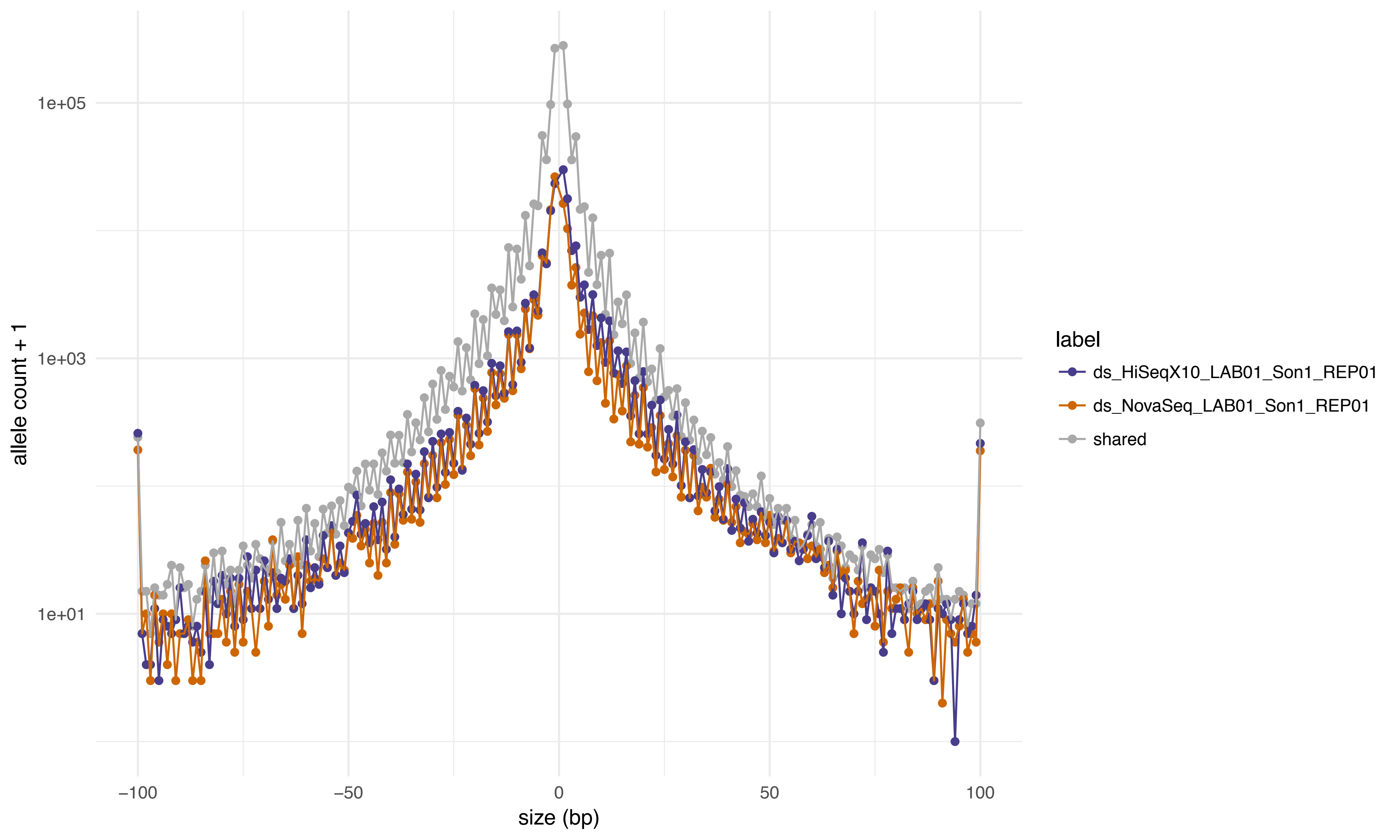

1e+05
1e+03
1e+01
allele count + 1
−100
−50
0
50
100
size (bp)
label
ds_HiSeqX10_LAB01_Son1_REP01
ds_NovaSeq_LAB01_Son1_REP01
shared

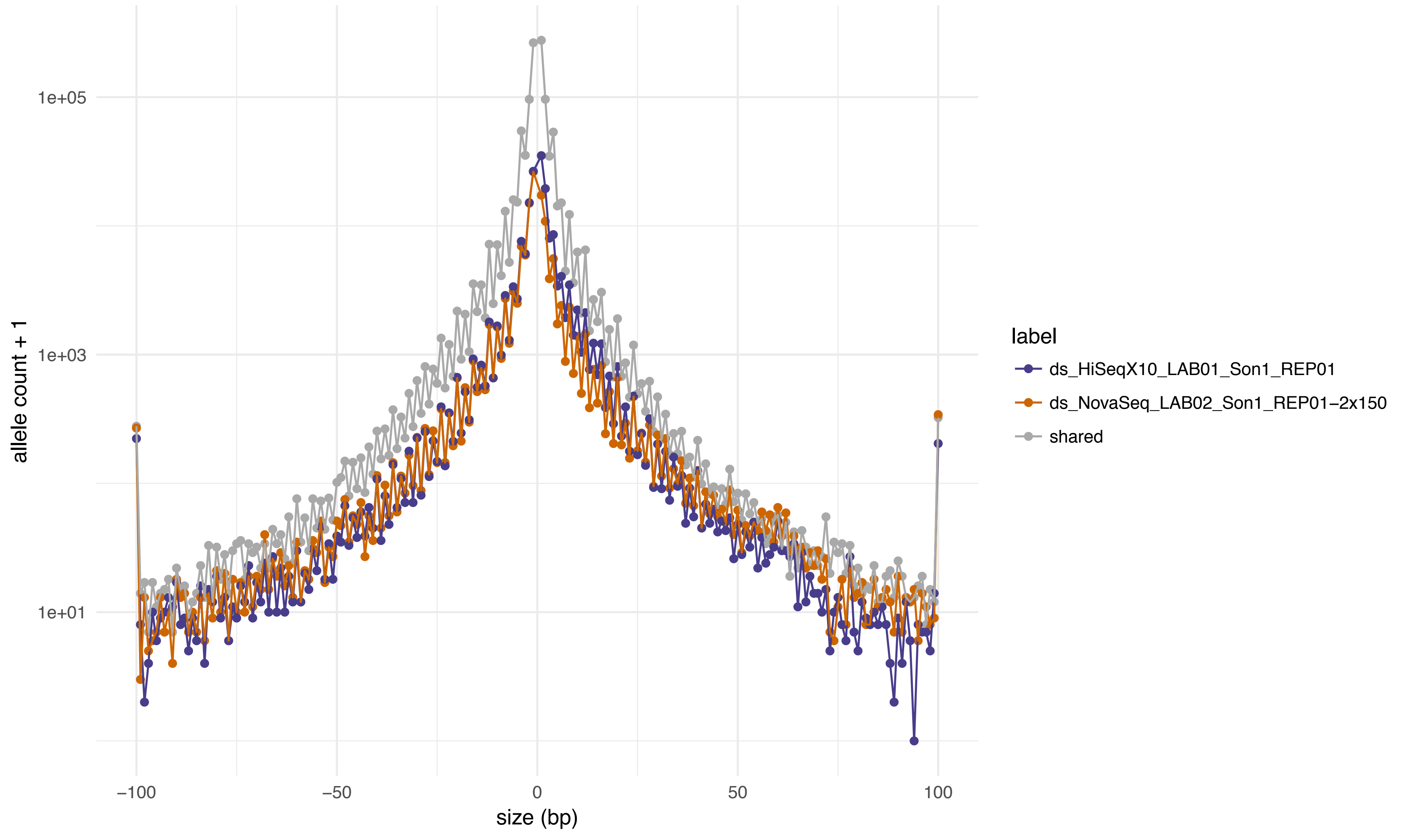

1e+05
1e+03
1e+01
allele count + 1
−100
−50
0
50
100
size (bp)
label
ds_HiSeqX10_LAB01_Son1_REP01
ds_NovaSeq_LAB02_Son1_REP01−2x150
shared

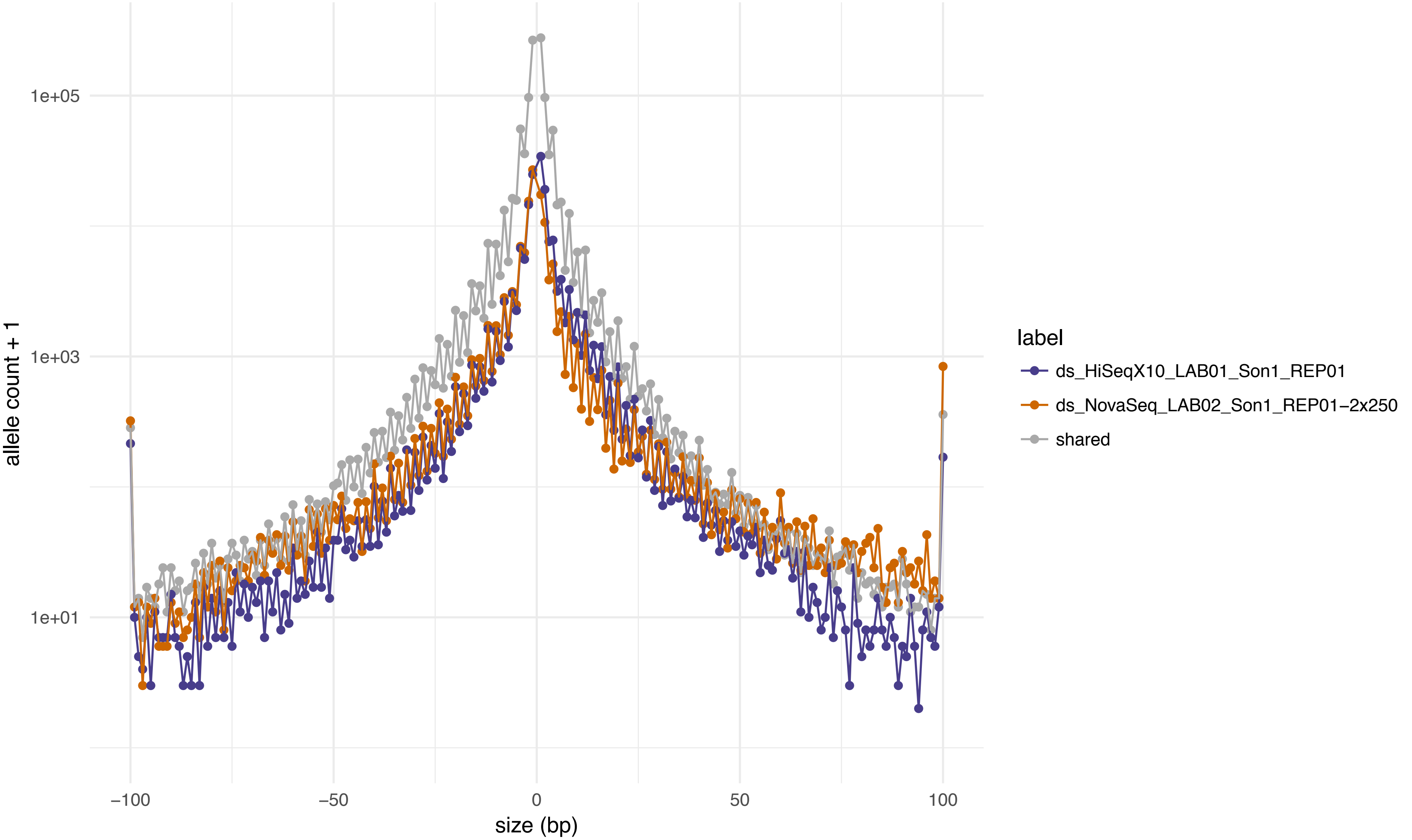

1e+05
1e+03
1e+01
allele count + 1
−100
−50
0
50
100
size (bp)
label
ds_HiSeqX10_LAB01_Son1_REP01
ds_NovaSeq_LAB02_Son1_REP01−2x250
shared

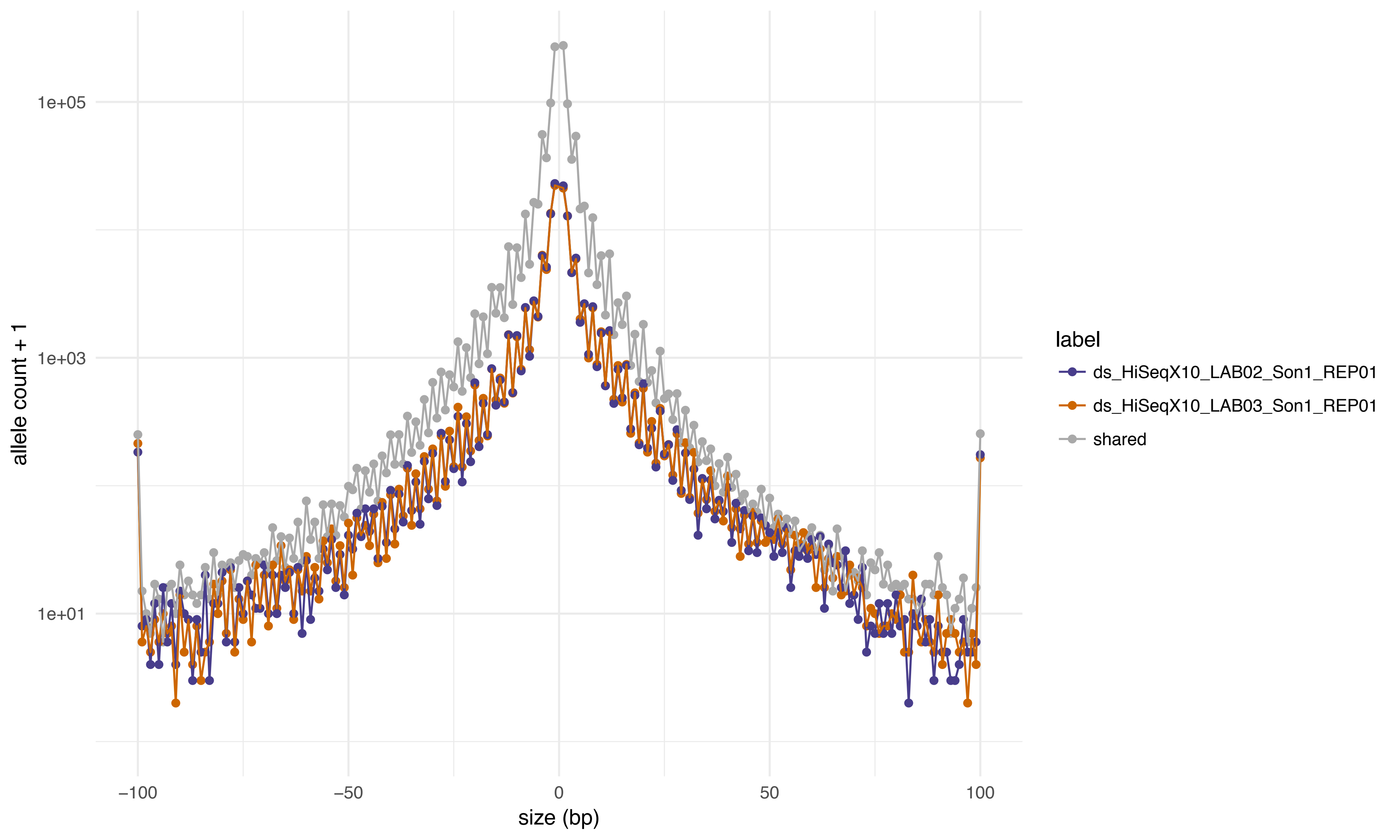

1e+05
1e+03
1e+01
allele count + 1
−100
−50
0
50
100
size (bp)
label
ds_HiSeqX10_LAB02_Son1_REP01
ds_HiSeqX10_LAB03_Son1_REP01
shared

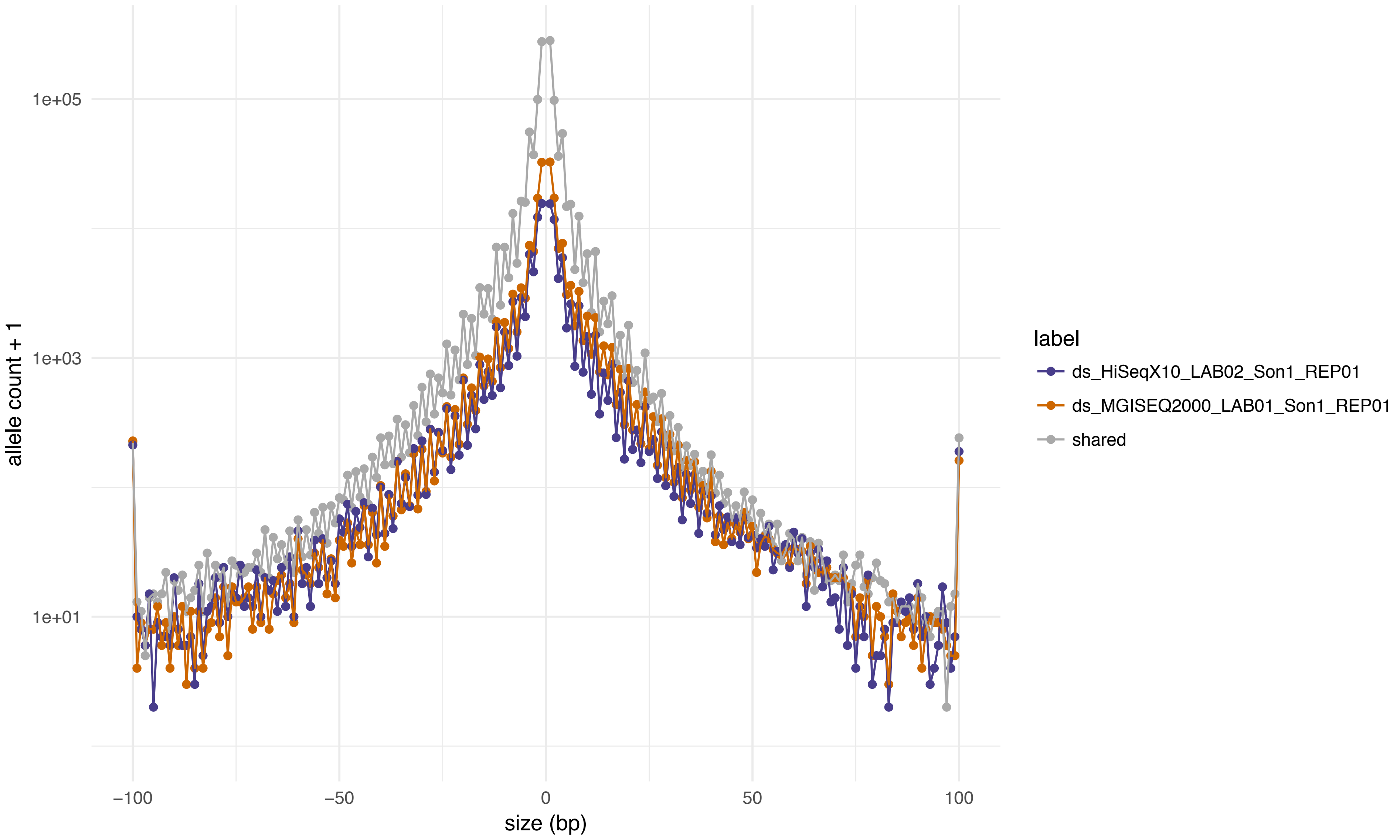

allele count + 1
1e+05
1e+03
1e+01
−100
−50
0
50
100
size (bp)
label
ds_HiSeqX10_LAB02_Son1_REP01
ds_MGISEQ2000_LAB01_Son1_REP01
shared

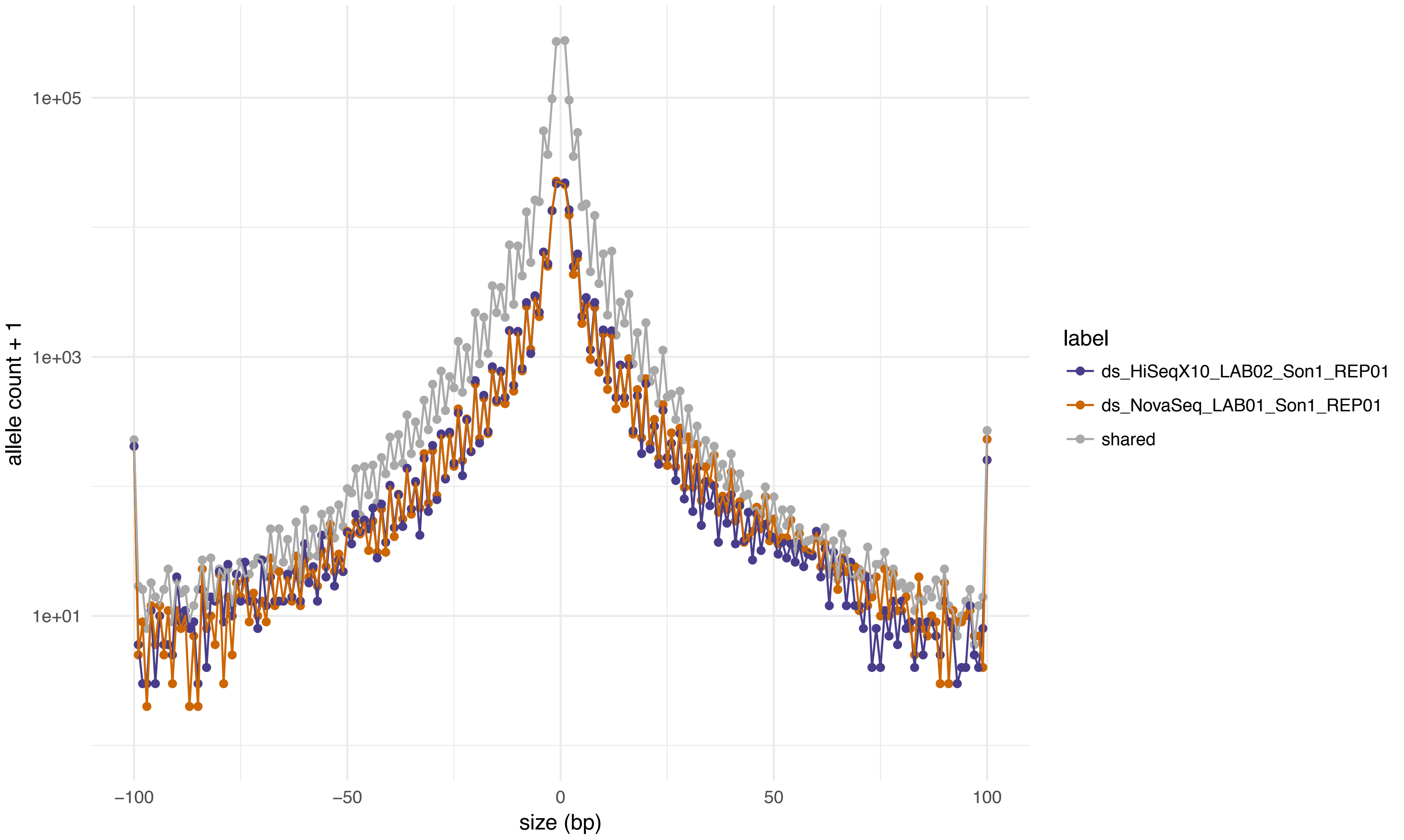

allele count + 1
1e+05
1e+03
1e+01
−100
−50
0
50
100
size (bp)
label
ds_HiSeqX10_LAB02_Son1_REP01
ds_NovaSeq_LAB01_Son1_REP01
shared

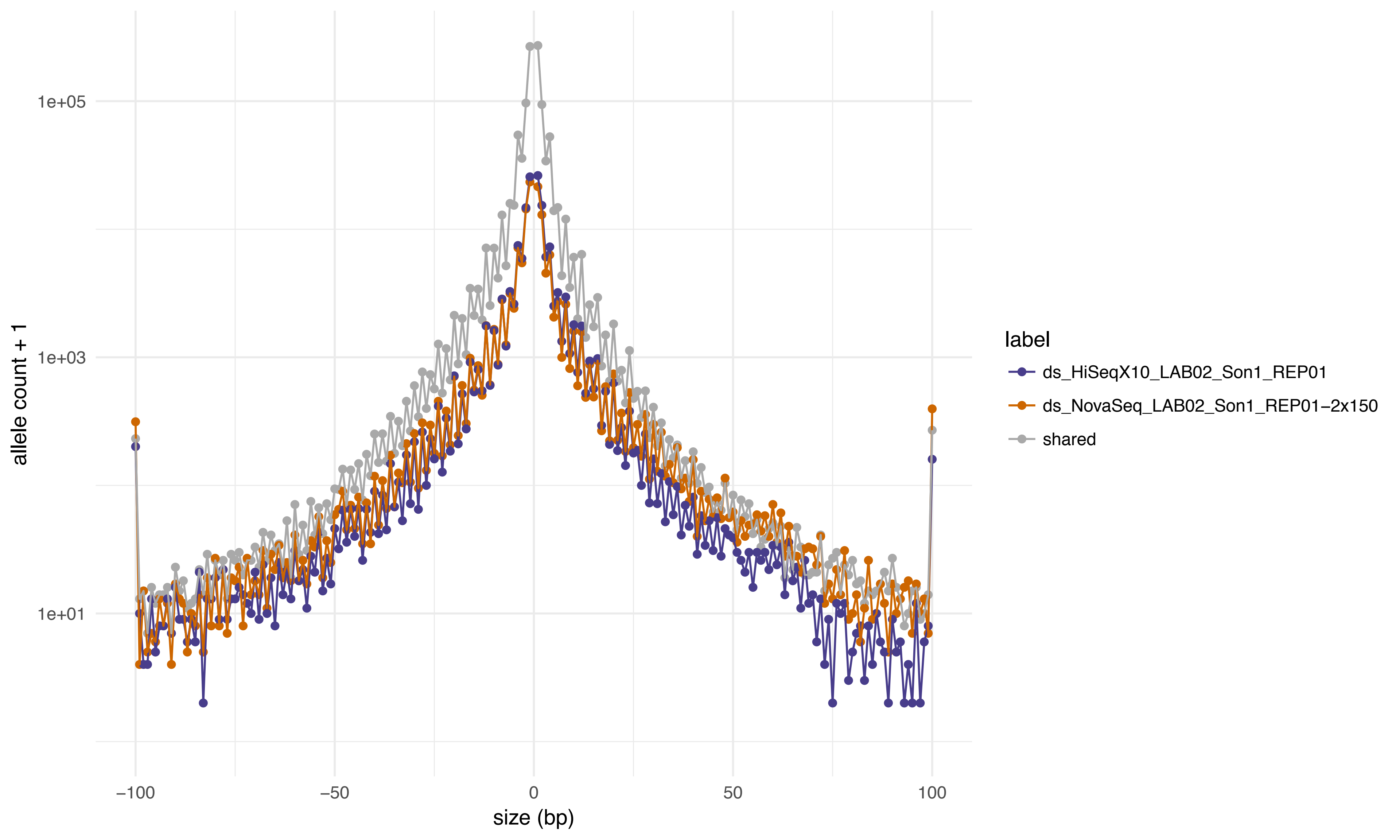

1e+05
1e+03
1e+01
allele count + 1
−100
−50
0
50
100
size (bp)
label
ds_HiSeqX10_LAB02_Son1_REP01
ds_NovaSeq_LAB02_Son1_REP01−2x150
shared

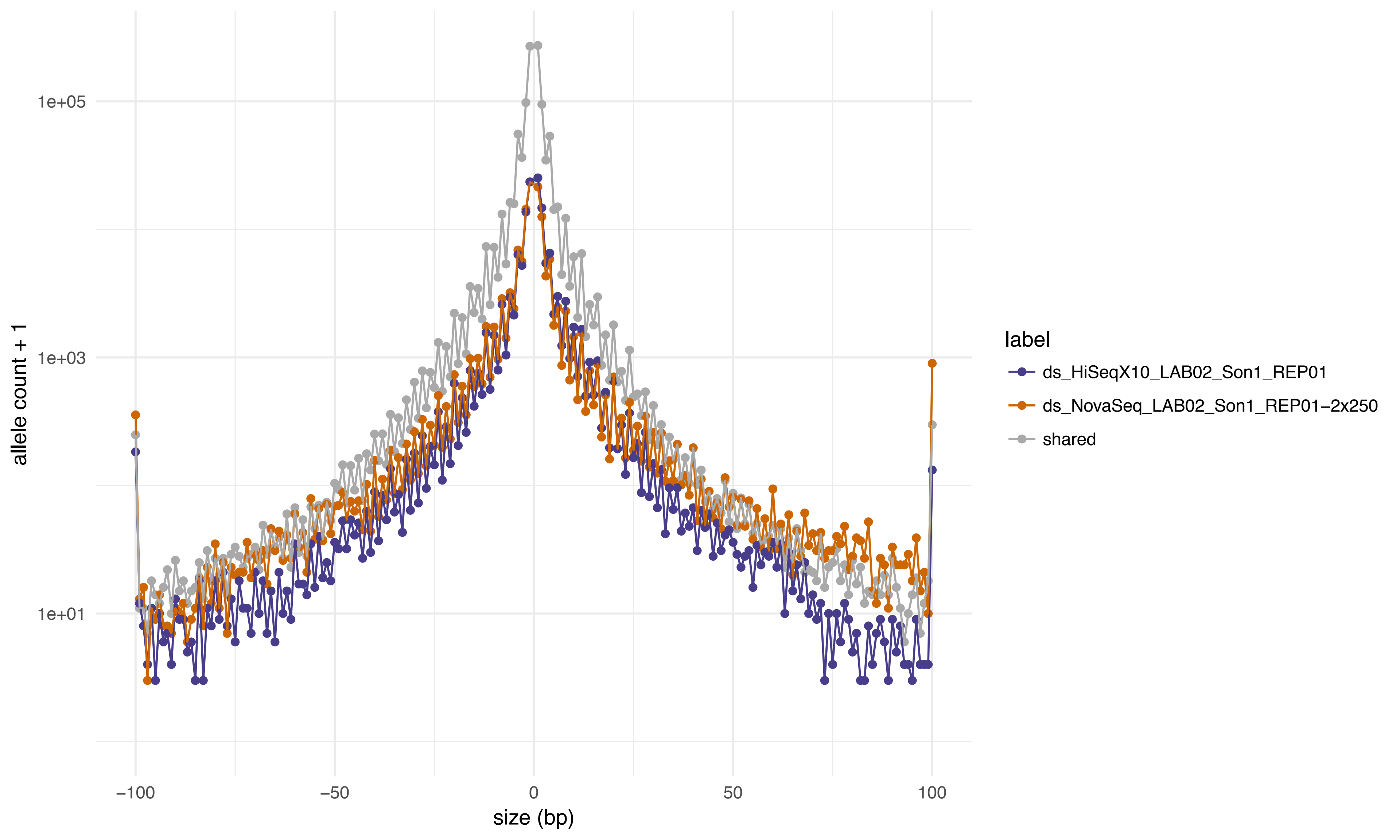
1e+05
1e+03
1e+01
allele count + 1
−100
−50
0
50
100
size (bp)
label
ds_HiSeqX10_LAB02_Son1_REP01
ds_NovaSeq_LAB02_Son1_REP01−2x250
shared

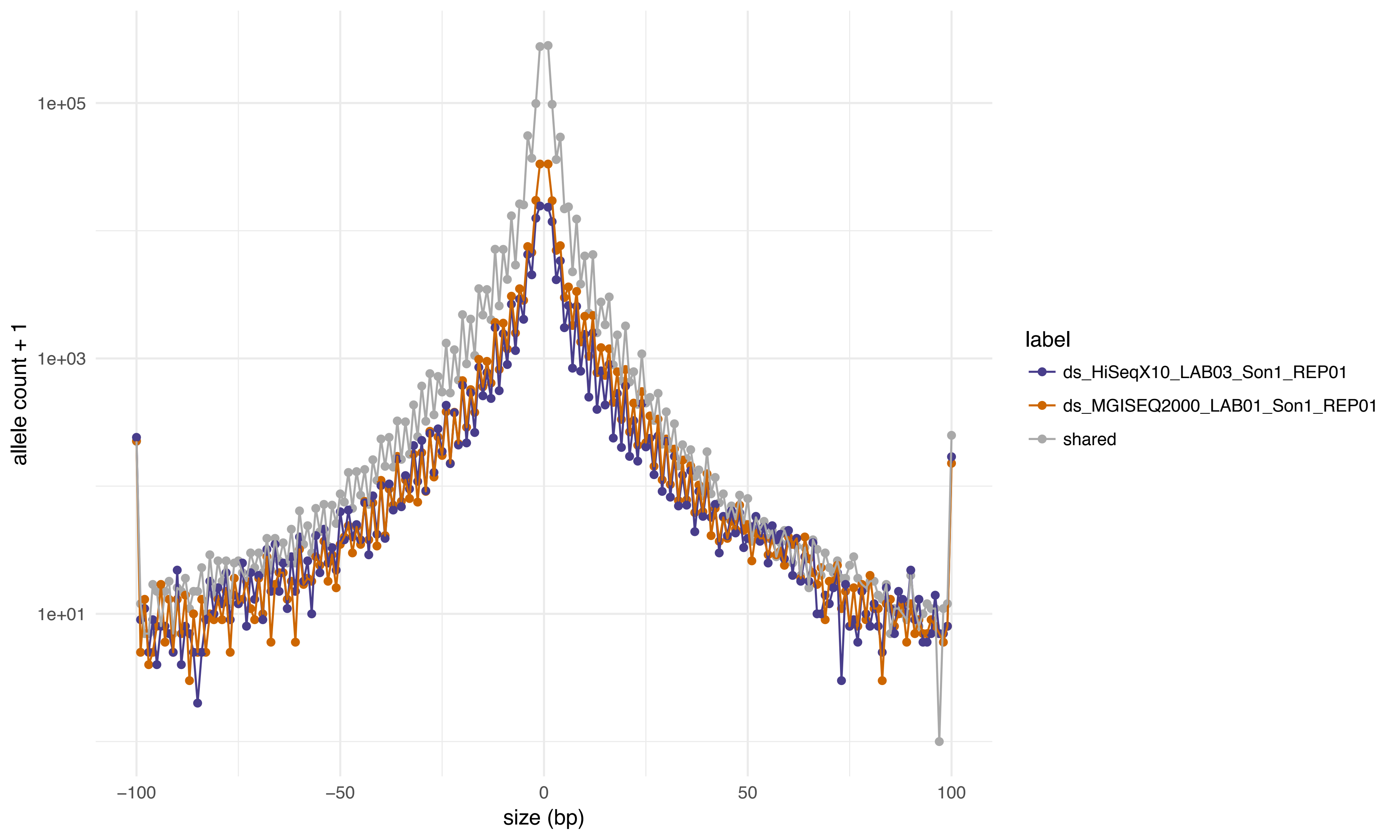

1e+05
1e+03
1e+01
allele count + 1
−100
−50
0
50
100
size (bp)
label
ds_HiSeqX10_LAB03_Son1_REP01
ds_MGISEQ2000_LAB01_Son1_REP01
shared

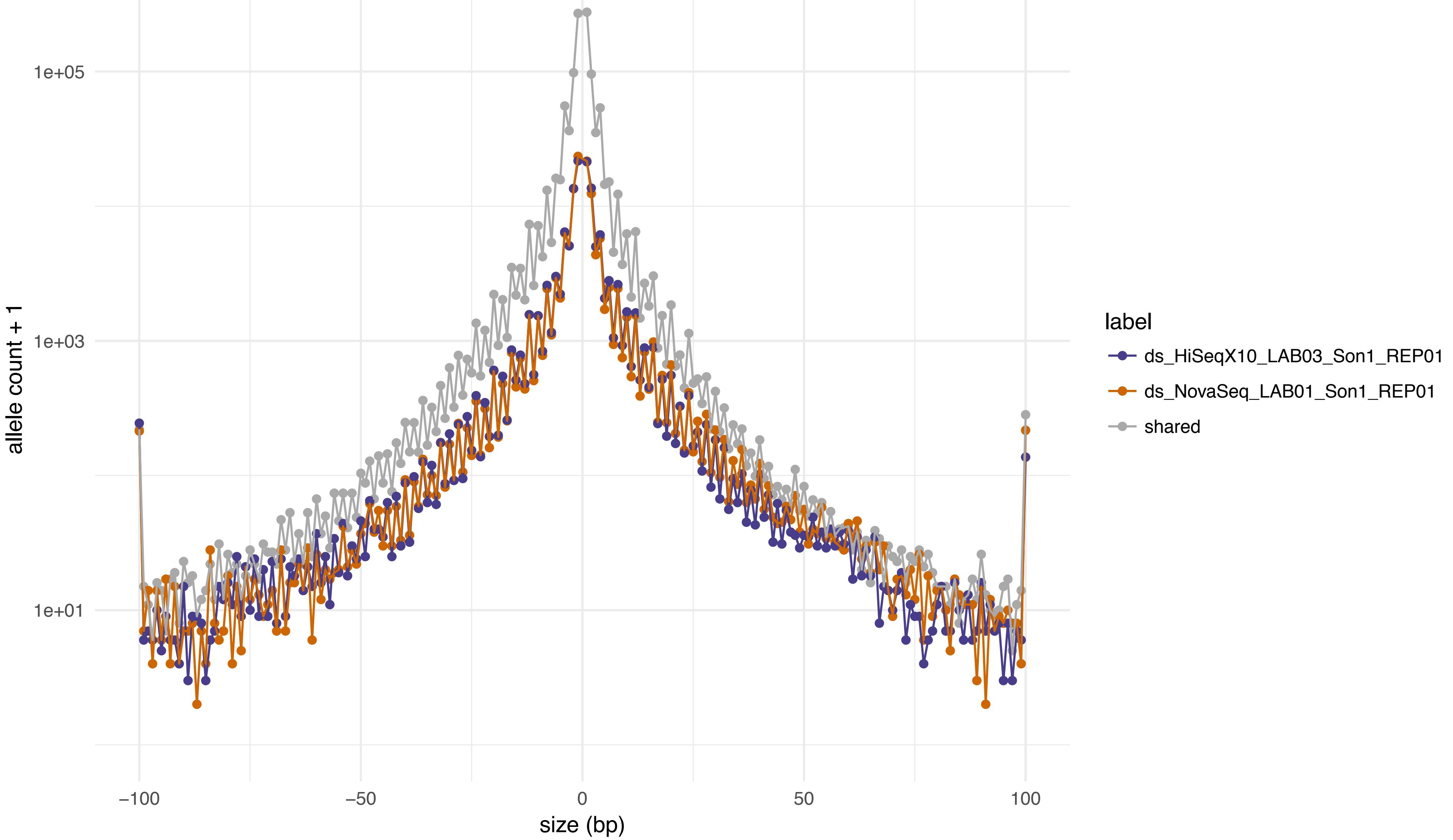

1e+05
1e+03
1e+01
allele count + 1
−100
−50
0
50
100
size (bp)
label
ds_HiSeqX10_LAB03_Son1_REP01
ds_NovaSeq_LAB01_Son1_REP01
shared

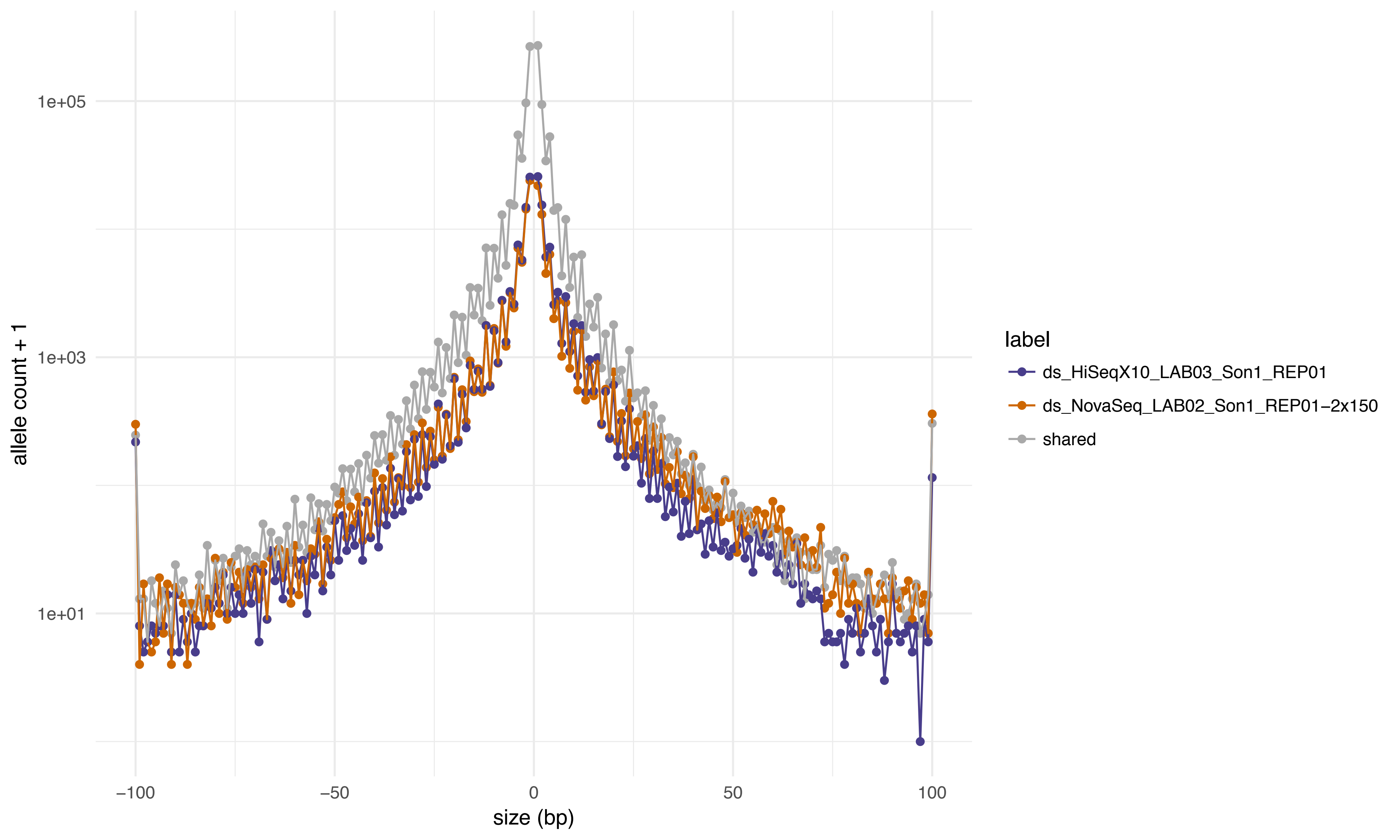
1e+05
1e+03
1e+01
allele count + 1
−100
−50
0
50
100
size (bp)
label
ds_HiSeqX10_LAB03_Son1_REP01
ds_NovaSeq_LAB02_Son1_REP01−2x150
shared

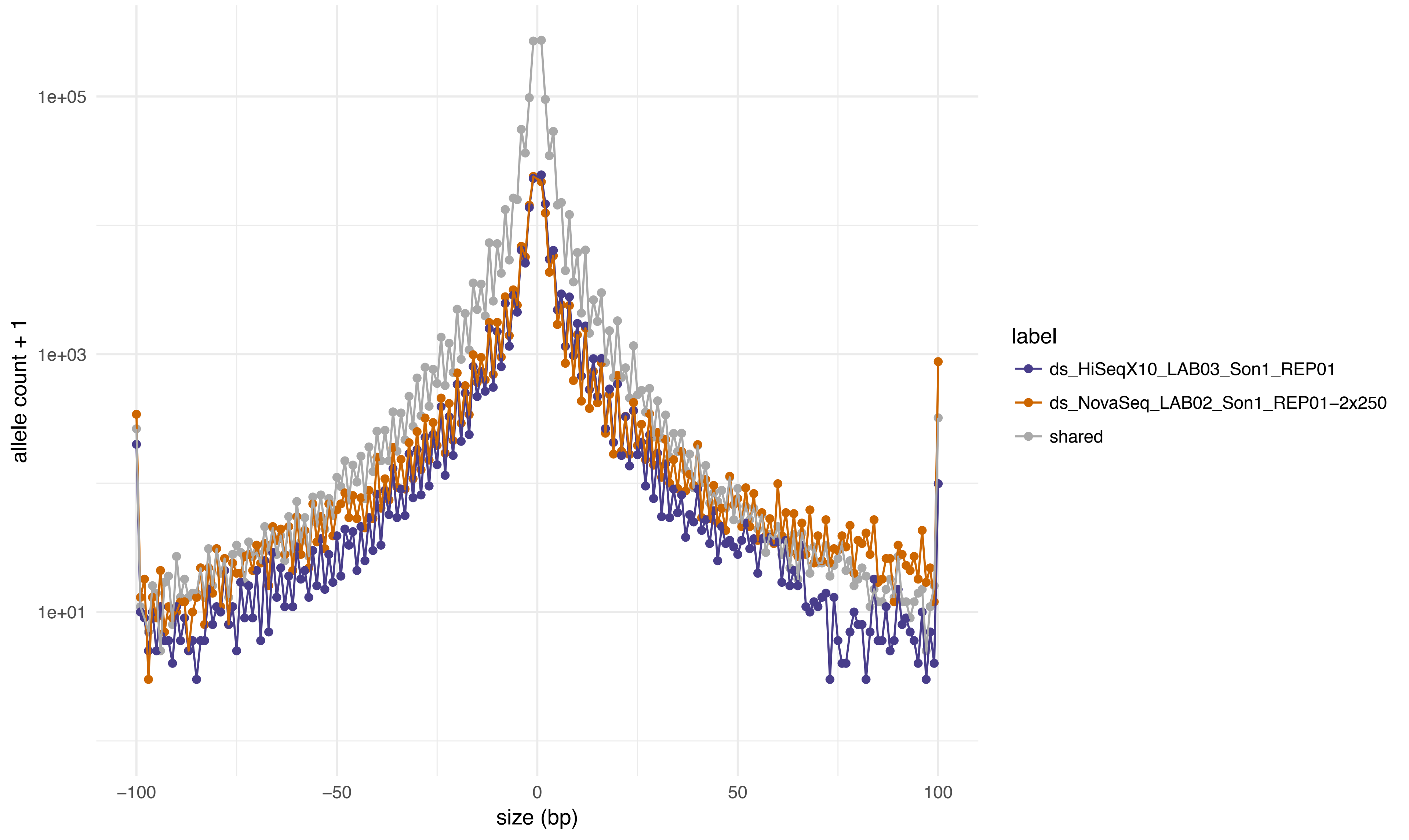
1e+05
1e+03
1e+01
allele count + 1
-100
-50
0
50
100
size (bp)
label
ds_HiSeqX10_LAB03_Son1_REP01
ds_NovaSeq_LAB02_Son1_REP01-2x250
shared

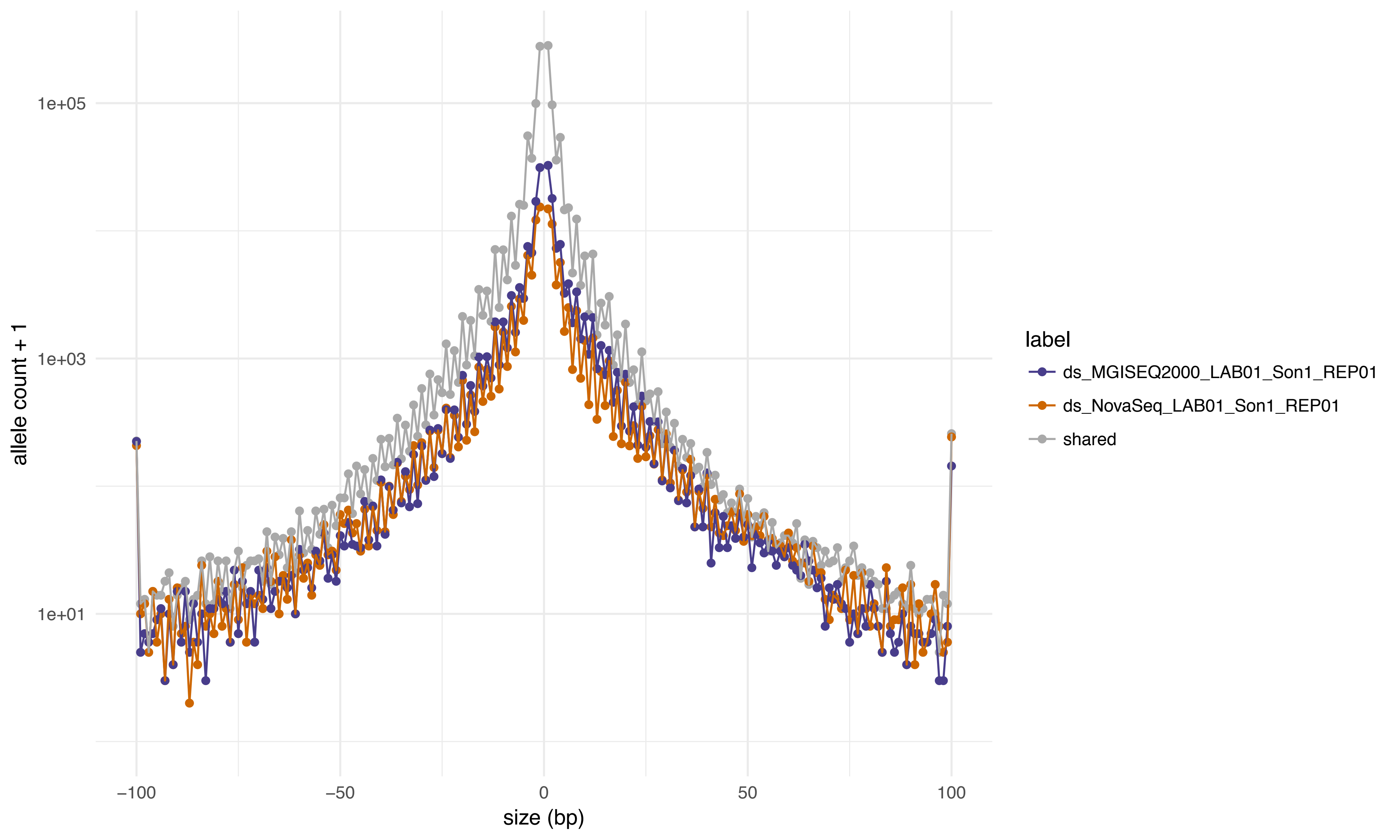

allele count + 1
1e+05
1e+03
1e+01
−100
−50
0
50
100
size (bp)
label
ds_MGISEQ2000_LAB01_Son1_REP01
ds_NovaSeq_LAB01_Son1_REP01
shared

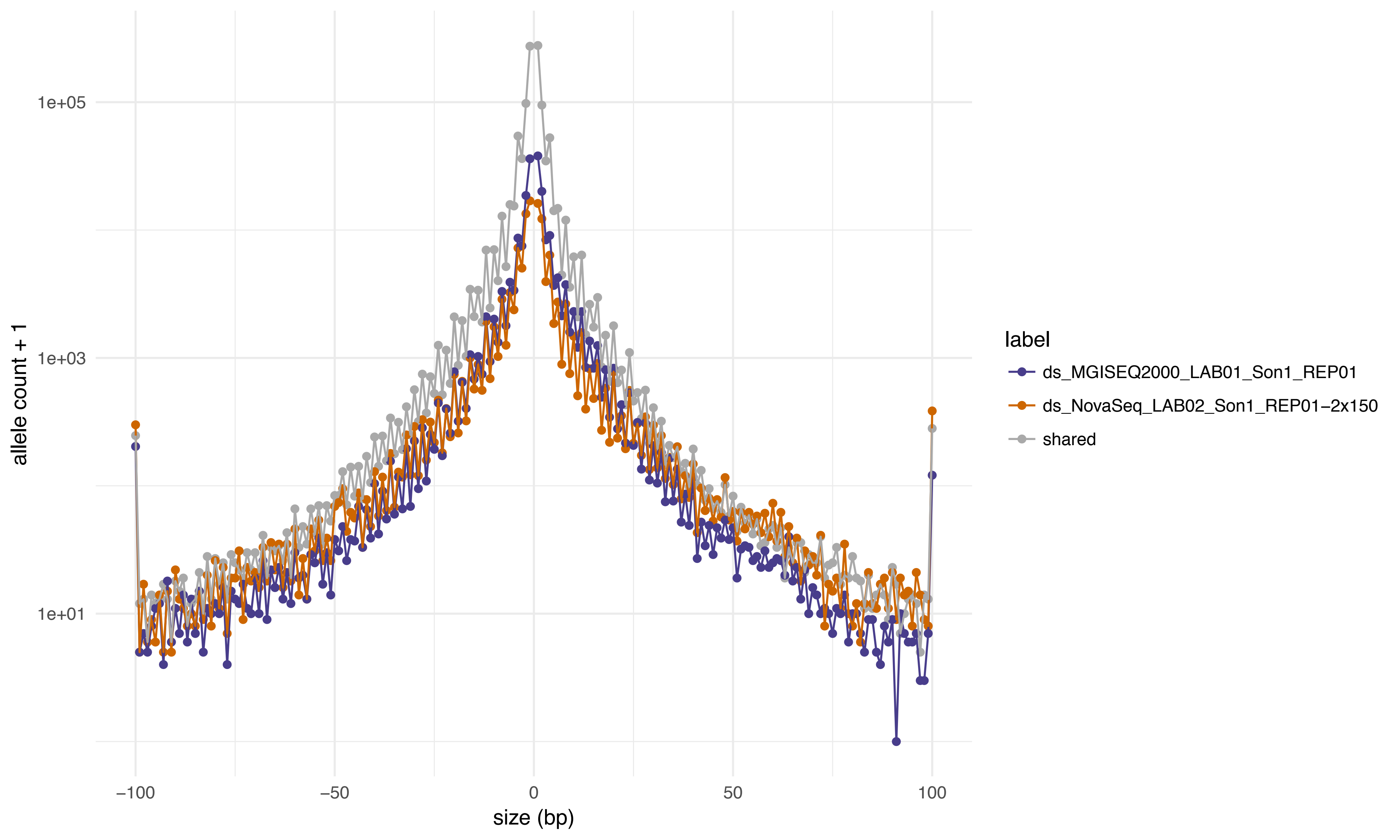
1e+05
1e+03
1e+01
allele count + 1
−100
−50
0
50
100
size (bp)
label
ds_MGISEQ2000_LAB01_Son1_REP01
ds_NovaSeq_LAB02_Son1_REP01−2x150
shared

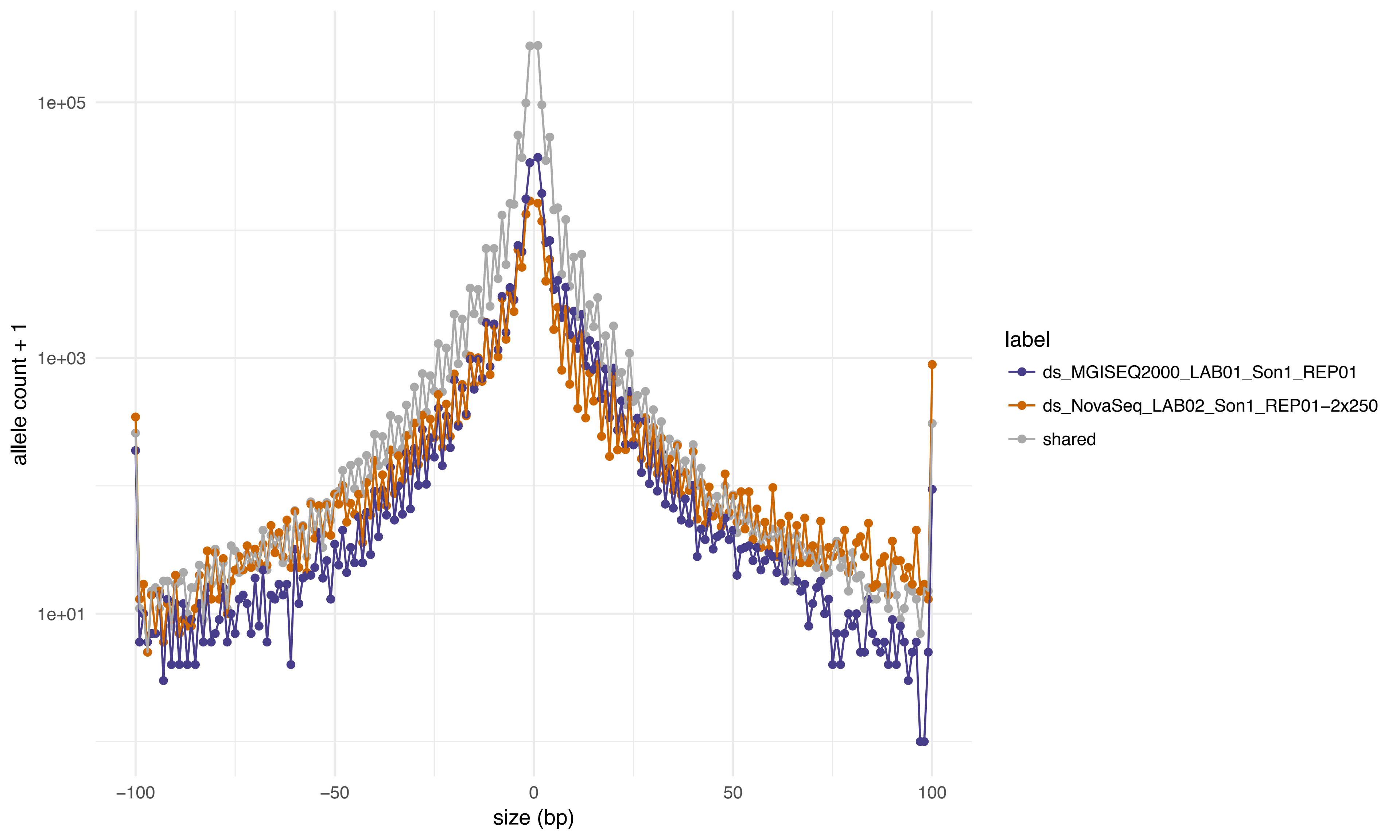

1e+05
1e+03
1e+01
allele count + 1
−100
−50
0
50
100
size (bp)
label
ds_MGISEQ2000_LAB01_Son1_REP01
ds_NovaSeq_LAB02_Son1_REP01−2x250
shared

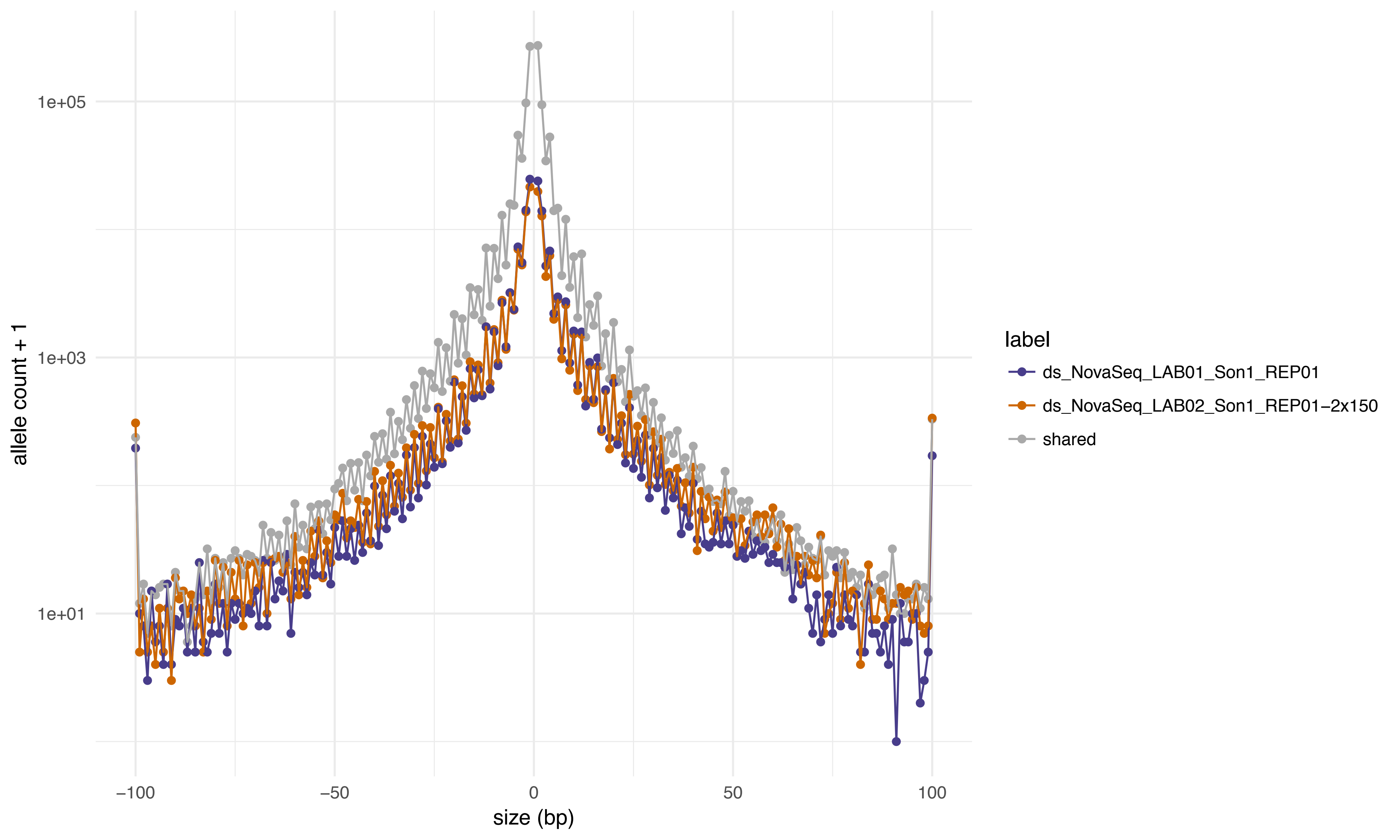
allele count + 1
1e+05
1e+03
1e+01
−100
−50
0
50
100
size (bp)
label
ds_NovaSeq_LAB01_Son1_REP01
ds_NovaSeq_LAB02_Son1_REP01−2x150
shared

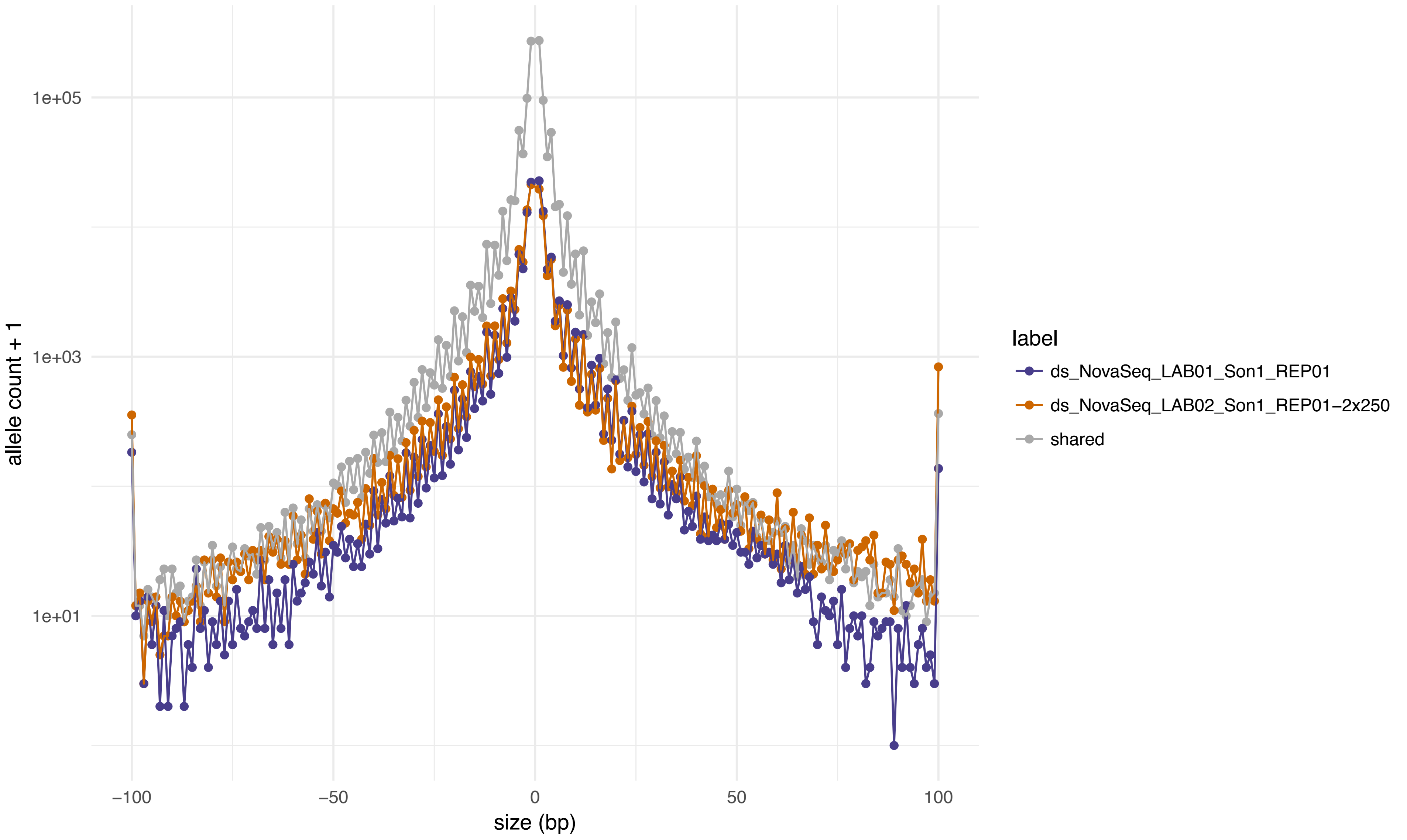

allele count + 1
1e+05
1e+03
1e+01
−100
−50
0
50
100
size (bp)
label
ds_NovaSeq_LAB01_Son1_REP01
ds_NovaSeq_LAB02_Son1_REP01−2x250
shared

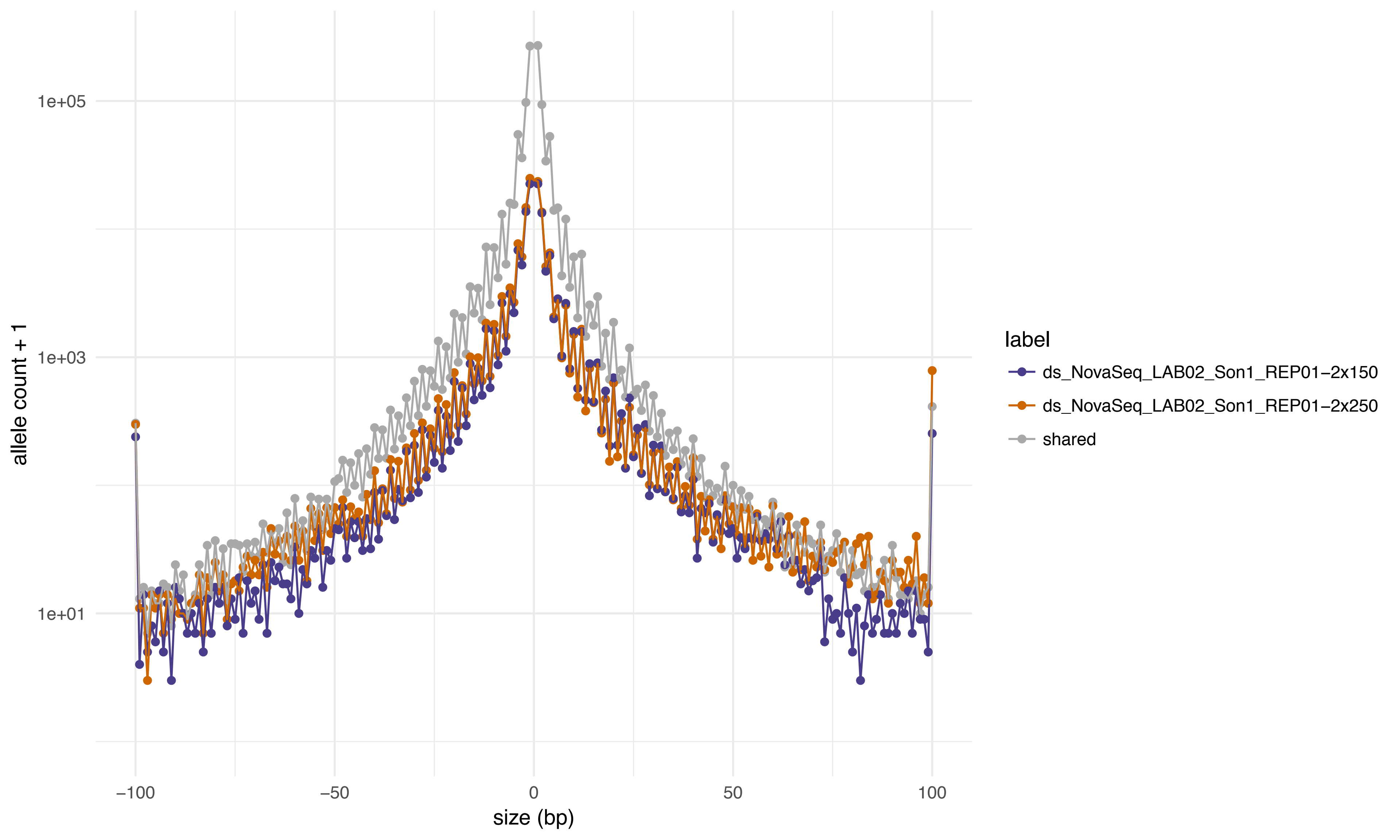

1e+05
1e+03
1e+01
allele count + 1
−100
−50
0
50
100
size (bp)
label
ds_NovaSeq_LAB02_Son1_REP01−2x150
ds_NovaSeq_LAB02_Son1_REP01−2x250
shared

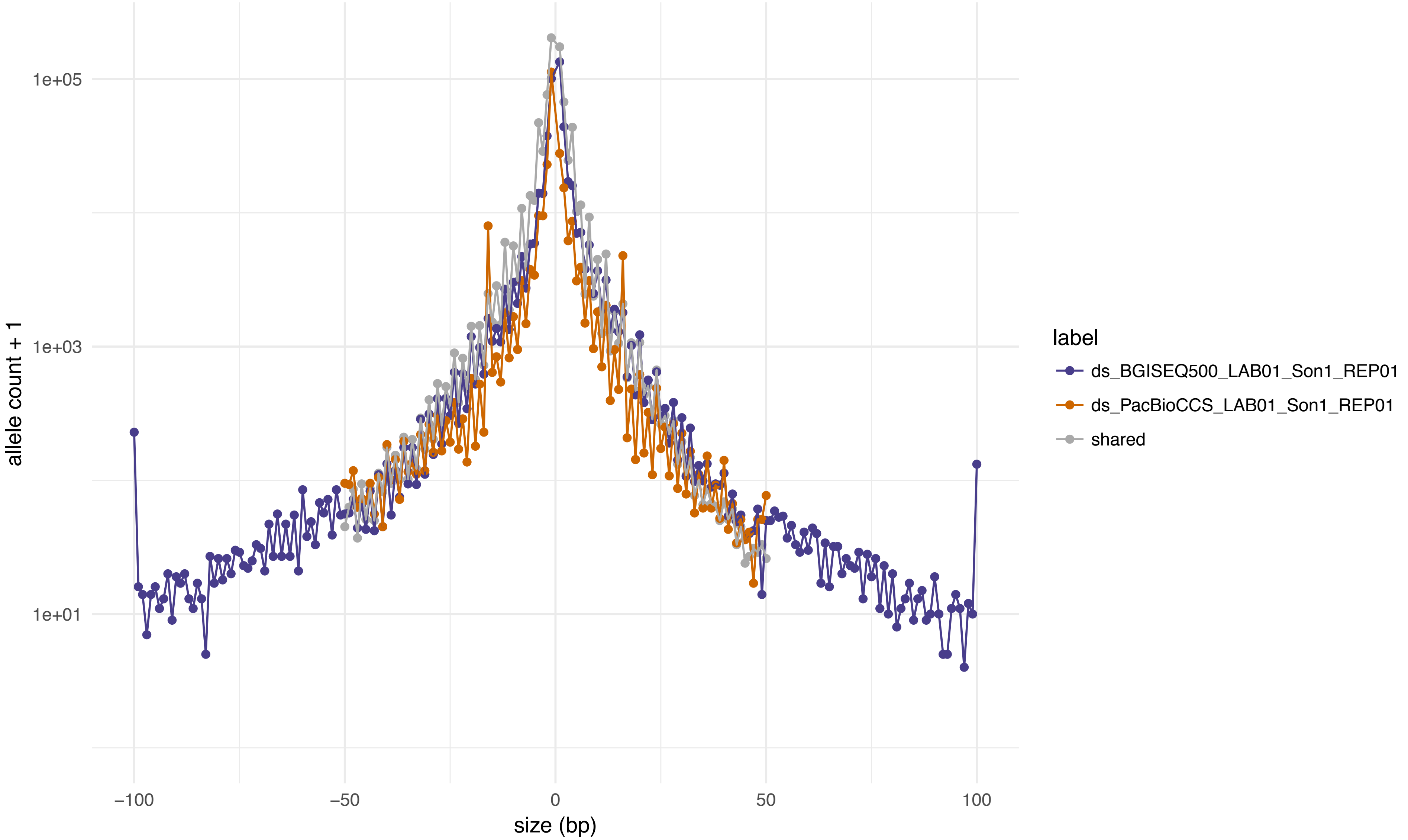
1e+05
1e+03
1e+01
allele count + 1
−100
−50
0
50
100
size (bp)
label
ds_BGISEQ500_LAB01_Son1_REP01
ds_PacBioCCS_LAB01_Son1_REP01
shared

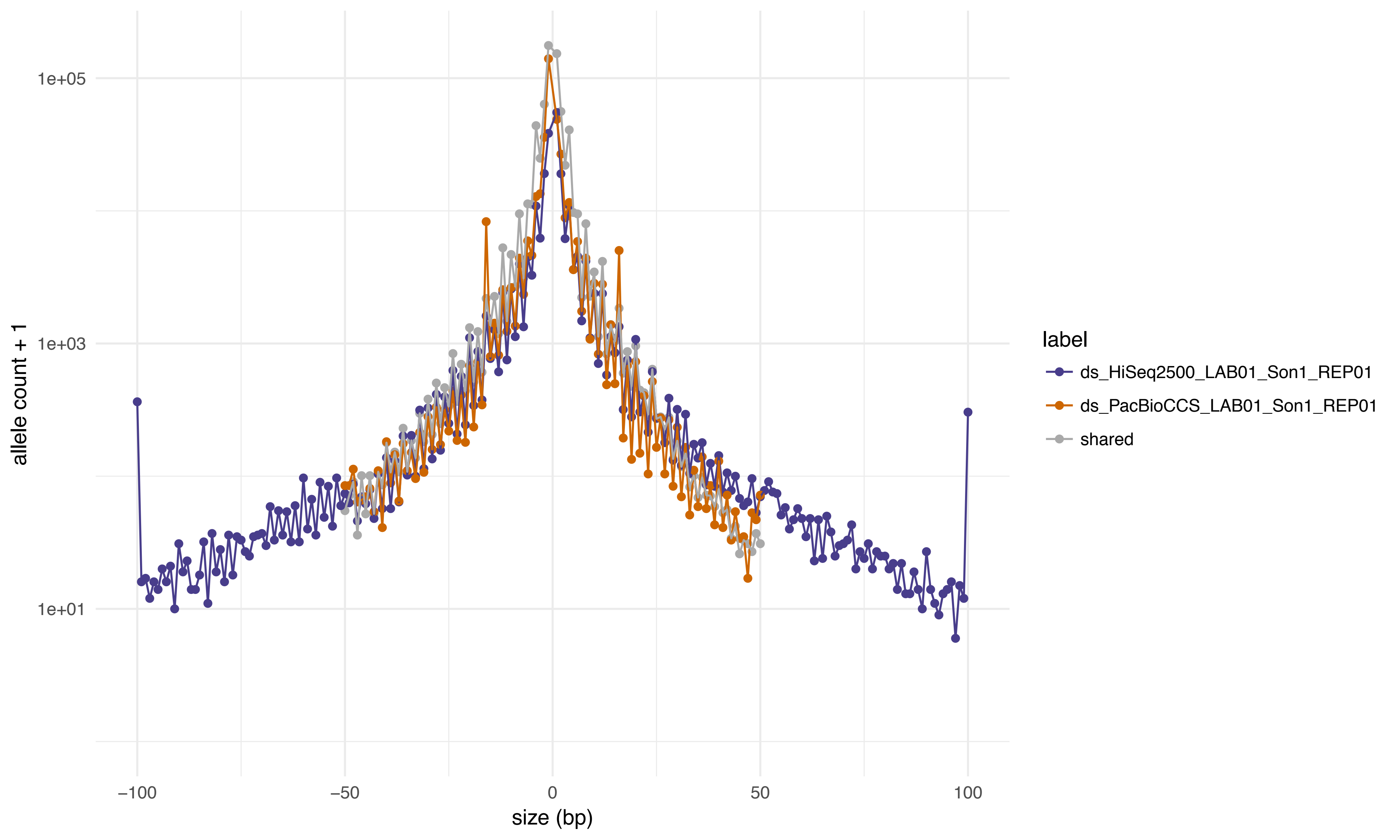
1e+05
1e+03
1e+01
allele count + 1
−100
−50
0
50
100
size (bp)
label
ds_HiSeq2500_LAB01_Son1_REP01
ds_PacBioCCS_LAB01_Son1_REP01
shared

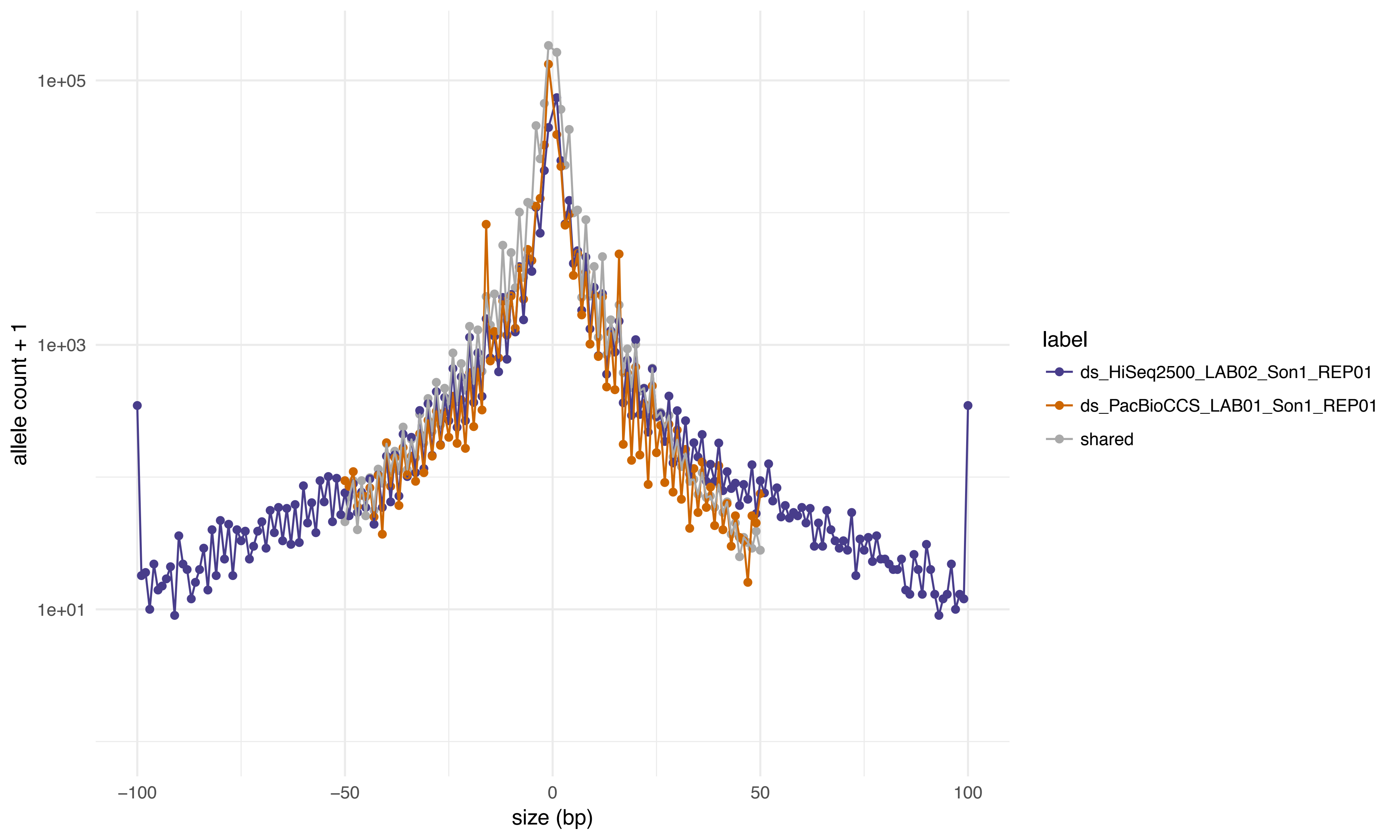
1e+05
1e+03
1e+01
allele count + 1
−100
−50
0
50
100
size (bp)
label
ds_HiSeq2500_LAB02_Son1_REP01
ds_PacBioCCS_LAB01_Son1_REP01
shared

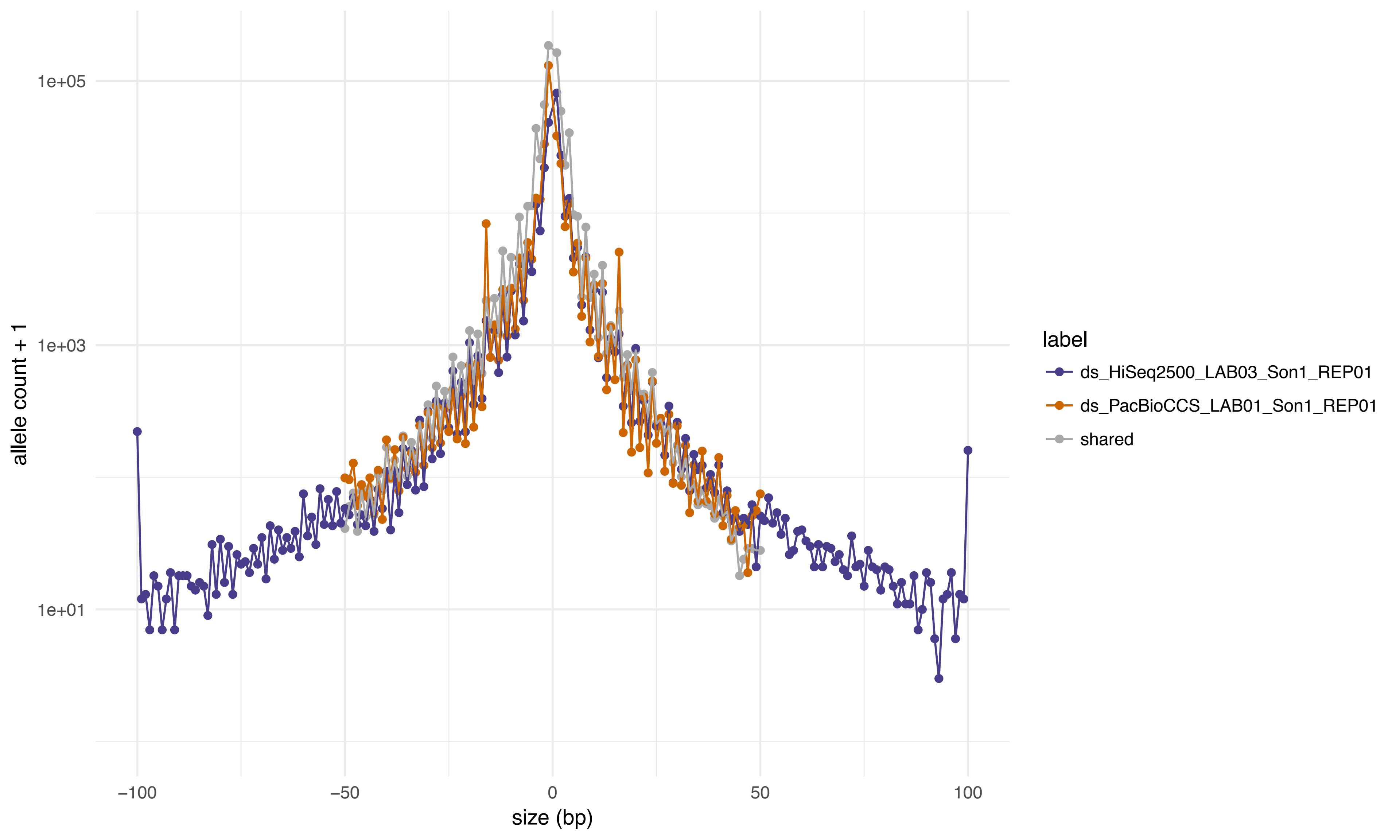

1e+05
1e+03
1e+01
allele count + 1
−100
−50
0
50
100
size (bp)
label
ds_HiSeq2500_LAB03_Son1_REP01
ds_PacBioCCS_LAB01_Son1_REP01
shared

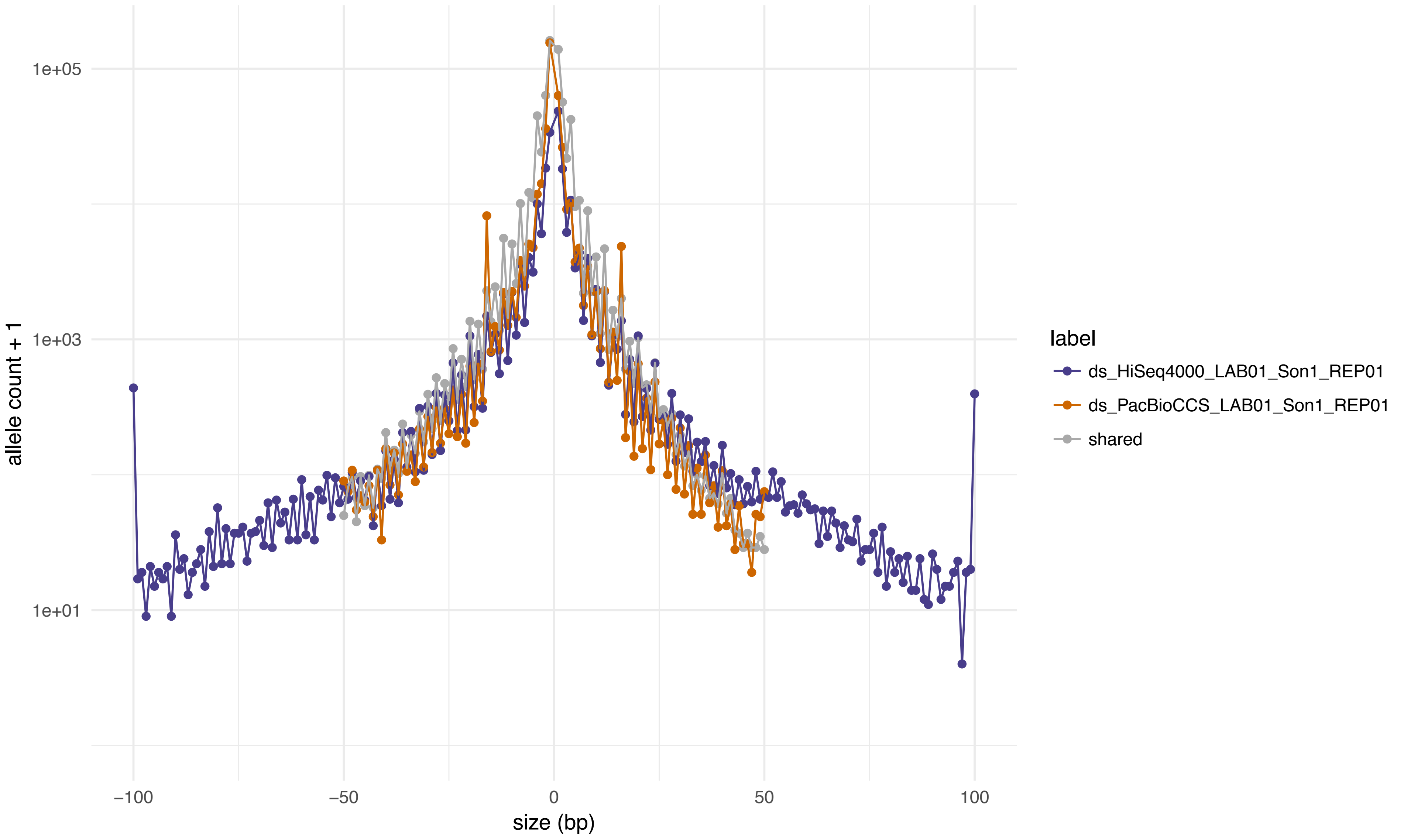

1e+05
1e+03
1e+01
allele count + 1
−100
−50
0
50
100
size (bp)
label
ds_HiSeq4000_LAB01_Son1_REP01
ds_PacBioCCS_LAB01_Son1_REP01
shared

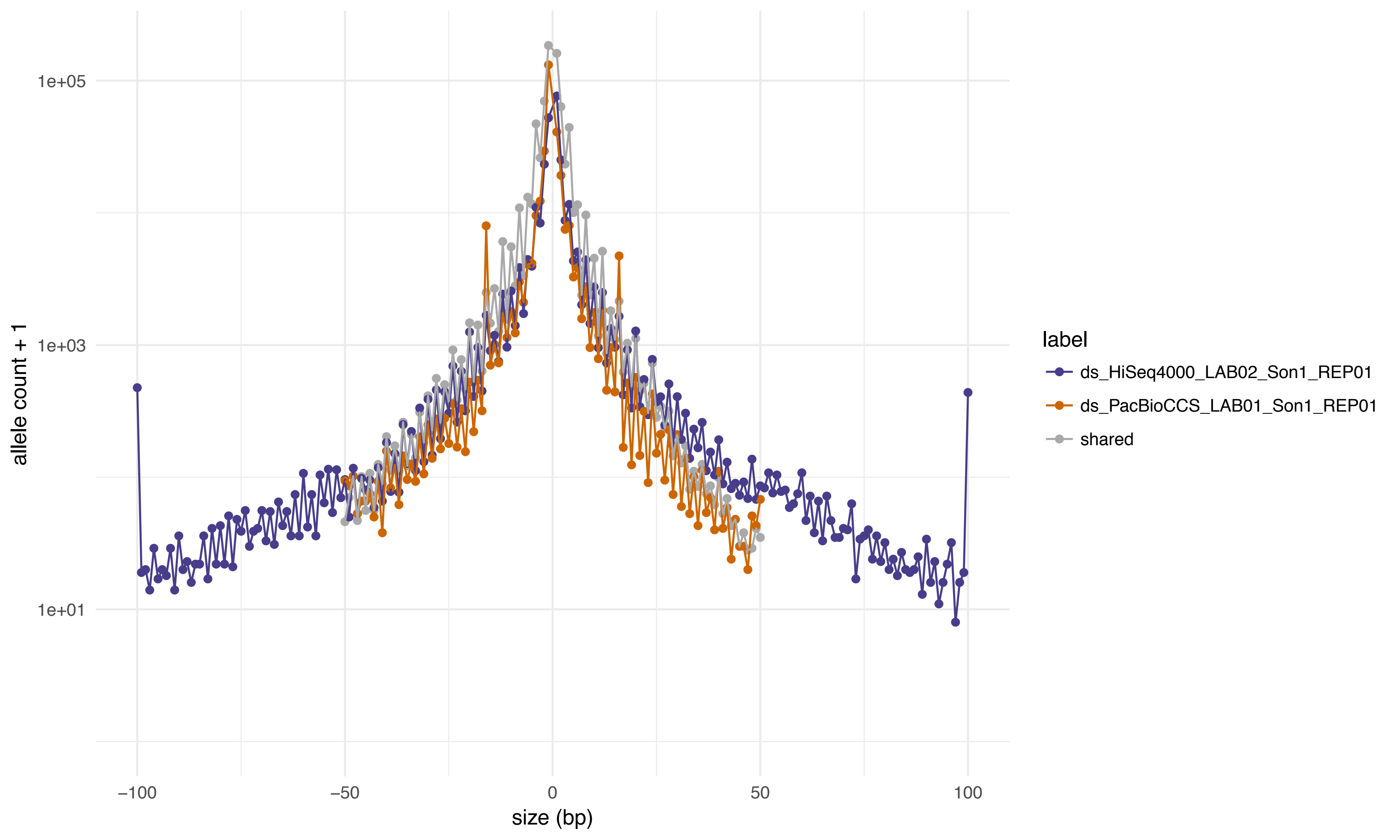

1e+05
1e+03
1e+01
allele count + 1
−100
−50
0
50
100
size (bp)
label
ds_HiSeq4000_LAB02_Son1_REP01
ds_PacBioCCS_LAB01_Son1_REP01
shared

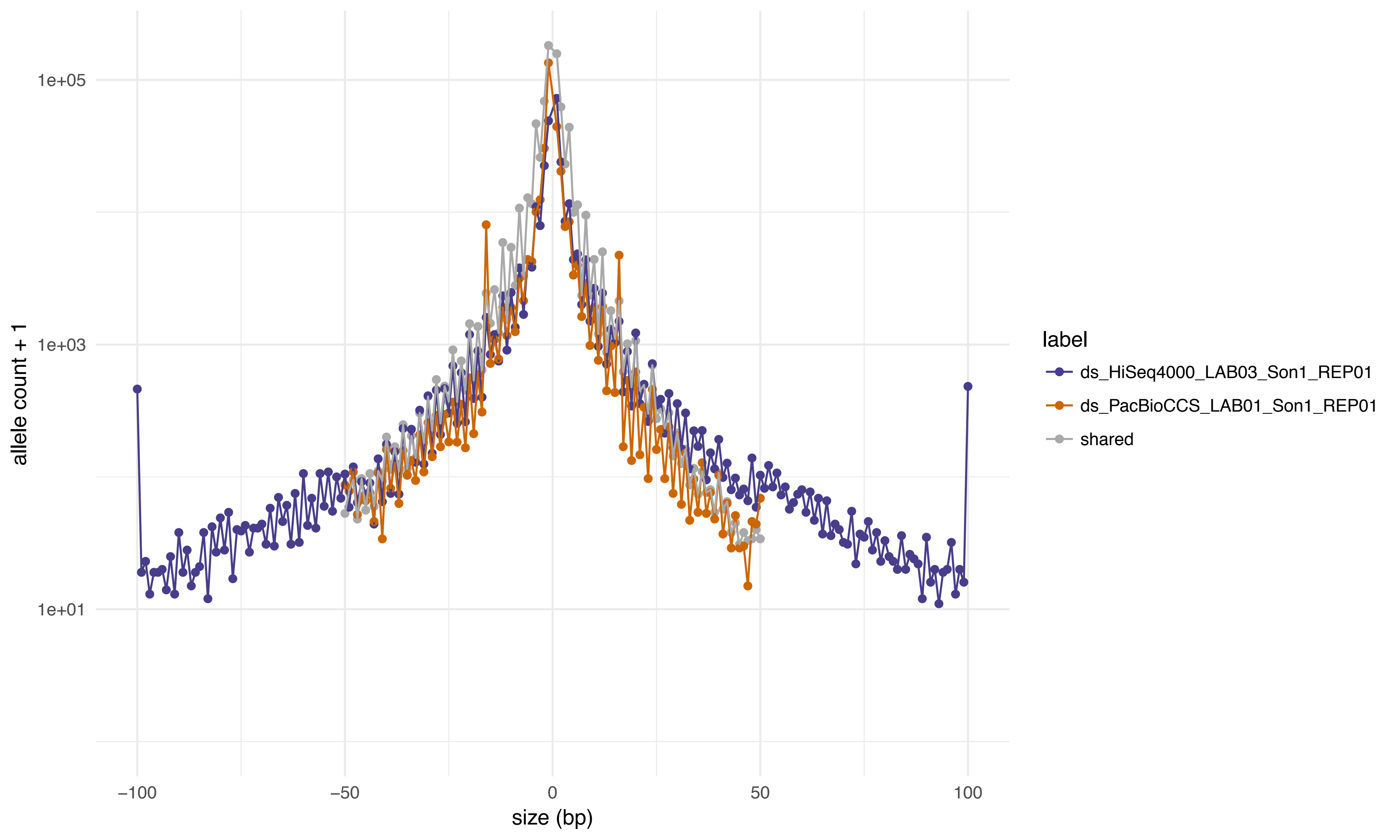

1e+05
1e+03
1e+01
allele count + 1
−100
−50
0
50
100
size (bp)
label
ds_HiSeq4000_LAB03_Son1_REP01
ds_PacBioCCS_LAB01_Son1_REP01
shared

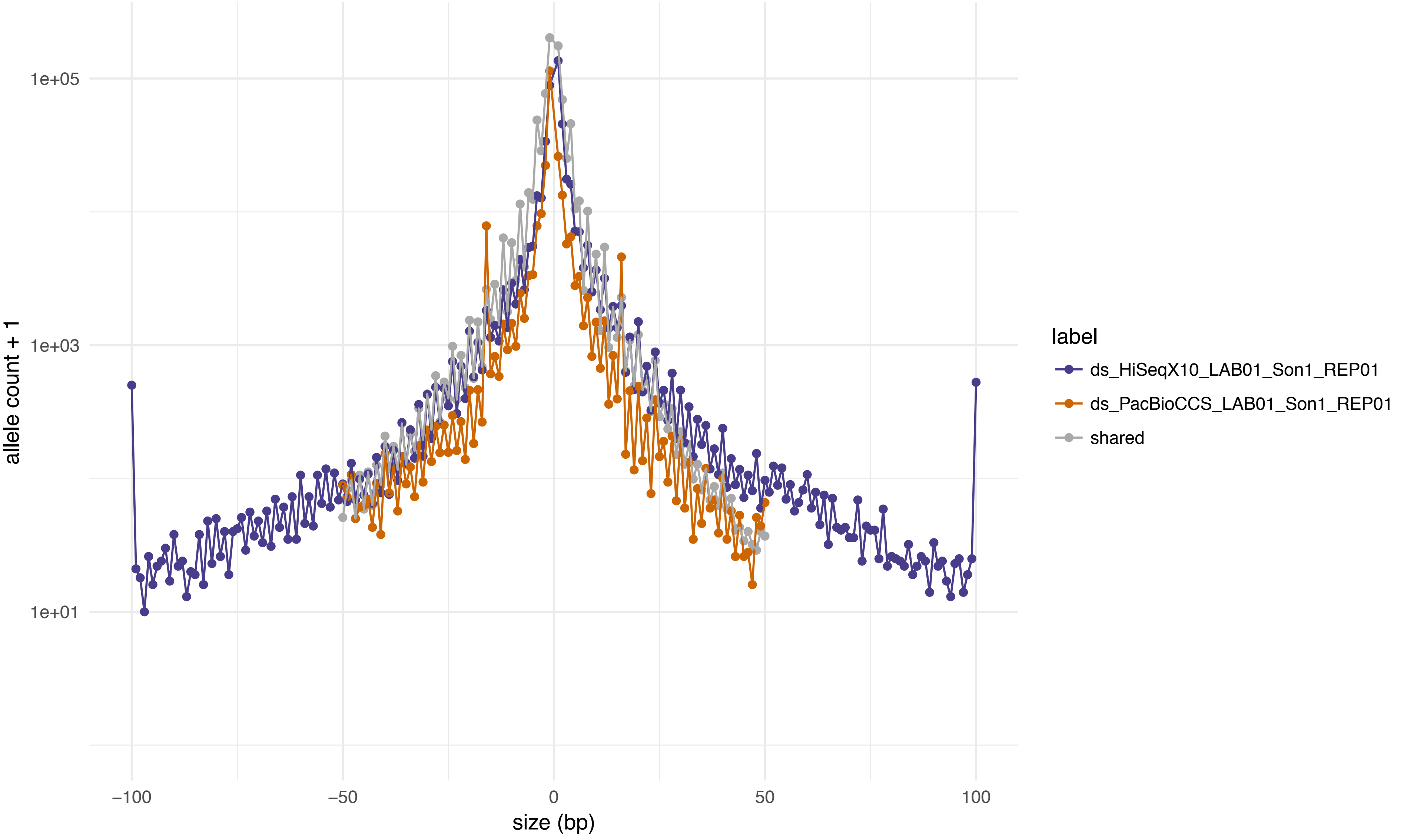
1e+05
1e+03
1e+01
allele count + 1
−100
−50
0
50
100
size (bp)
label
ds_HiSeqX10_LAB01_Son1_REP01
ds_PacBioCCS_LAB01_Son1_REP01
shared

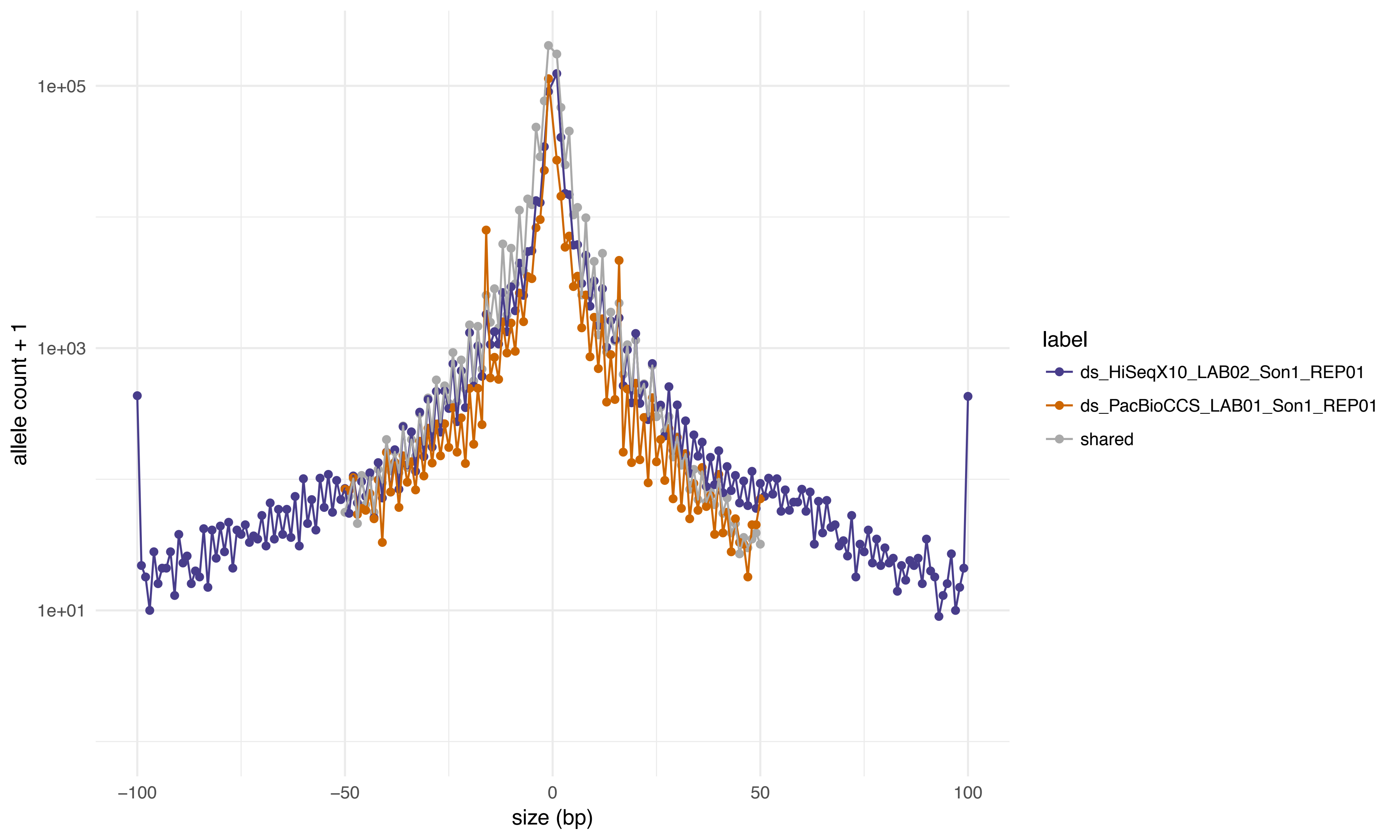

1e+05
1e+03
1e+01
allele count + 1
−100
−50
0
50
100
size (bp)
label
ds_HiSeqX10_LAB02_Son1_REP01
ds_PacBioCCS_LAB01_Son1_REP01
shared

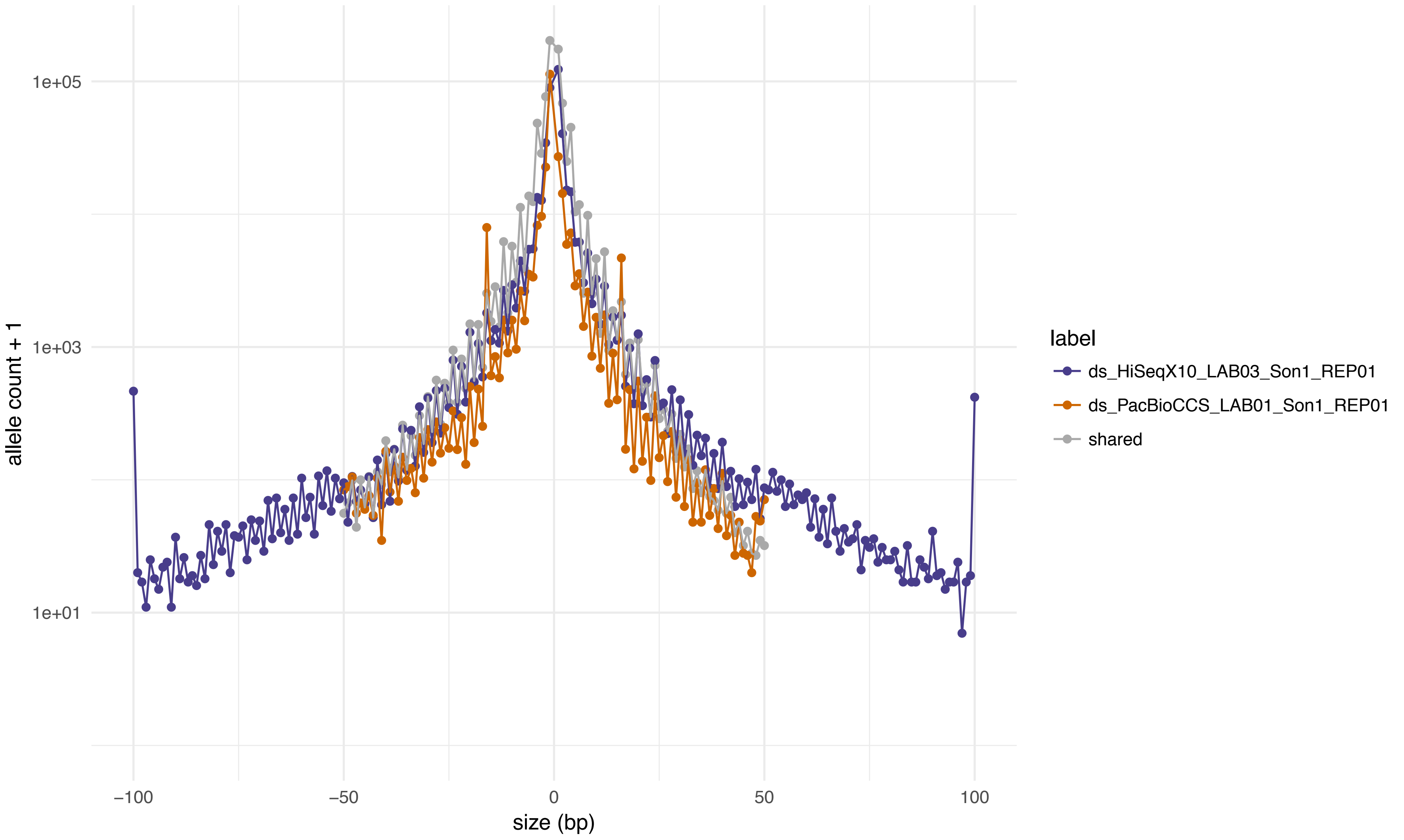

allele count + 1
1e+05
1e+03
1e+01
−100
−50
0
50
100
size (bp)
label
ds_HiSeqX10_LAB03_Son1_REP01
ds_PacBioCCS_LAB01_Son1_REP01
shared

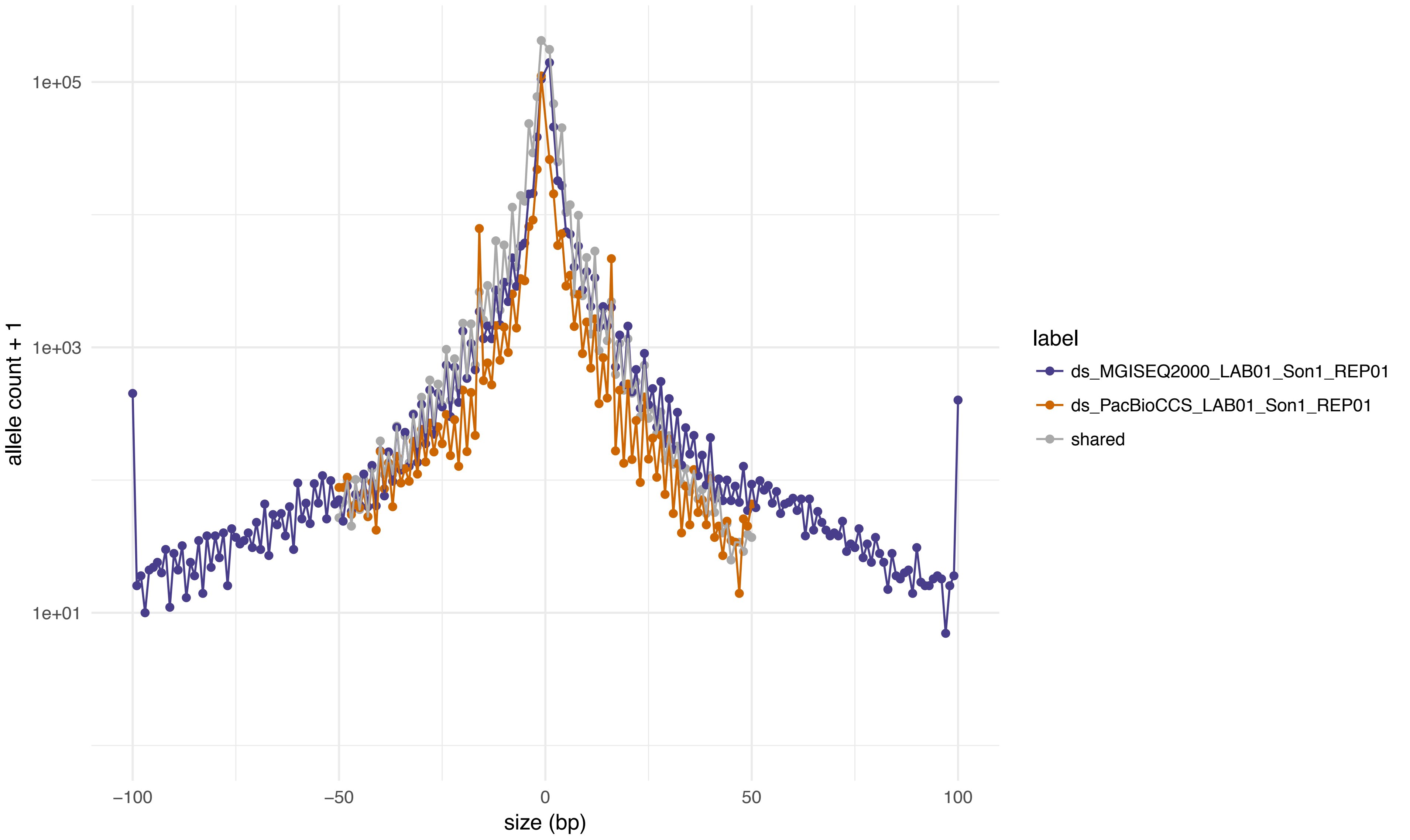

1e+05
1e+03
1e+01
allele count + 1
−100
−50
0
50
100
size (bp)
label
ds_MGISEQ2000_LAB01_Son1_REP01
ds_PacBioCCS_LAB01_Son1_REP01
shared

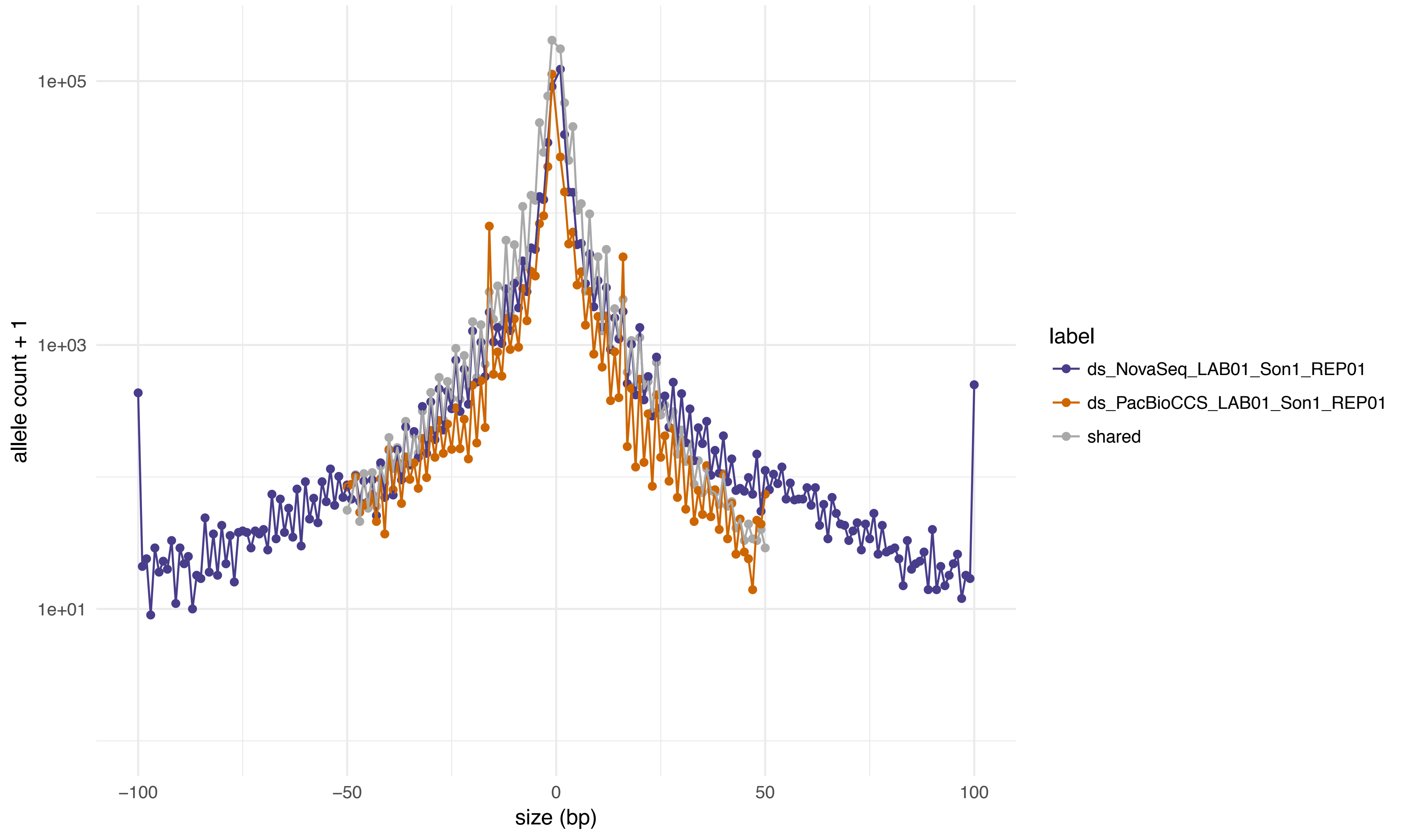
1e+05
1e+03
1e+01
allele count + 1
−100
−50
0
50
100
size (bp)
label
ds_NovaSeq_LAB01_Son1_REP01
ds_PacBioCCS_LAB01_Son1_REP01
shared

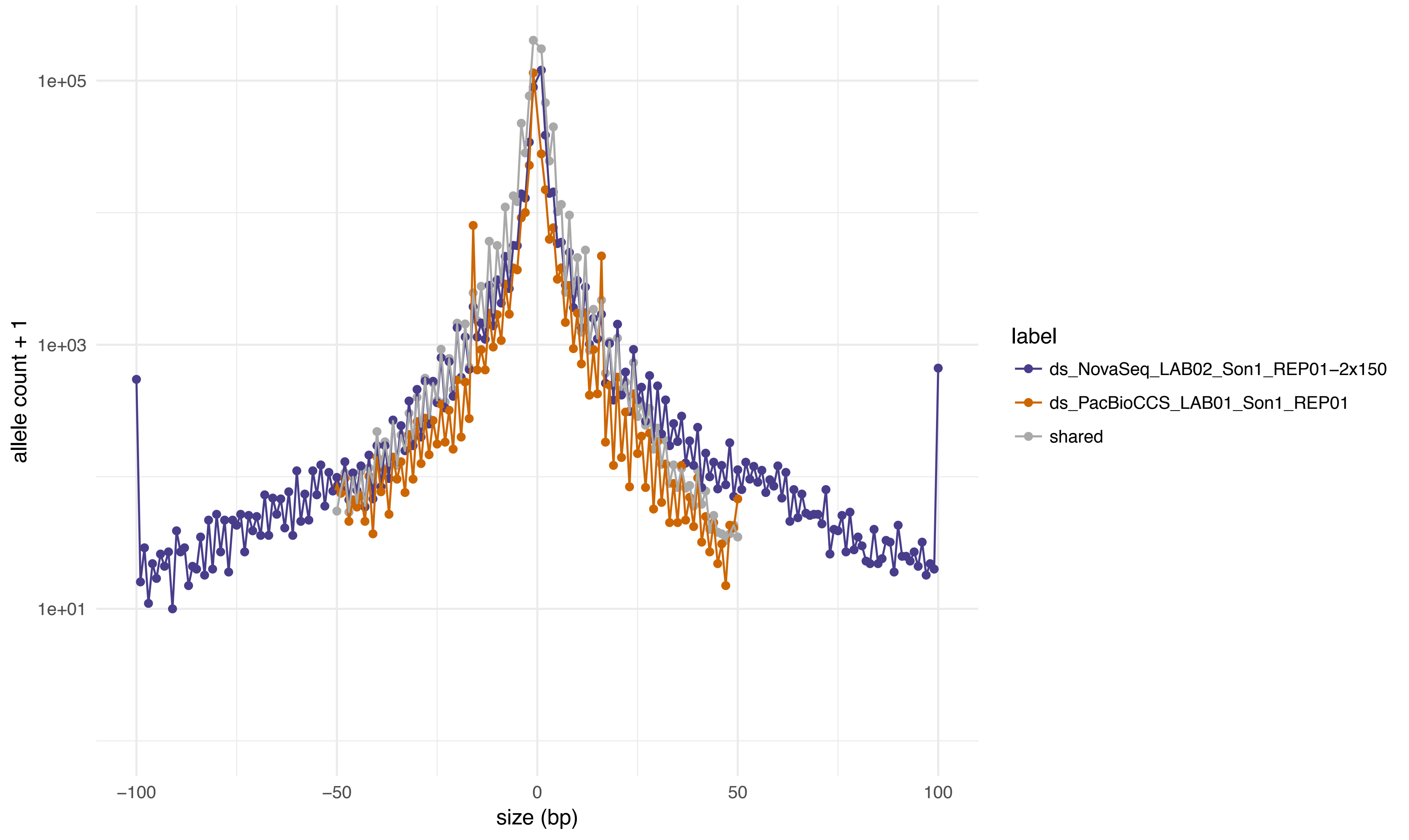

allele count + 1
1e+05
1e+03
1e+01
−100
−50
0
50
100
size (bp)
label
ds_NovaSeq_LAB02_Son1_REP01−2x150
ds_PacBioCCS_LAB01_Son1_REP01
shared

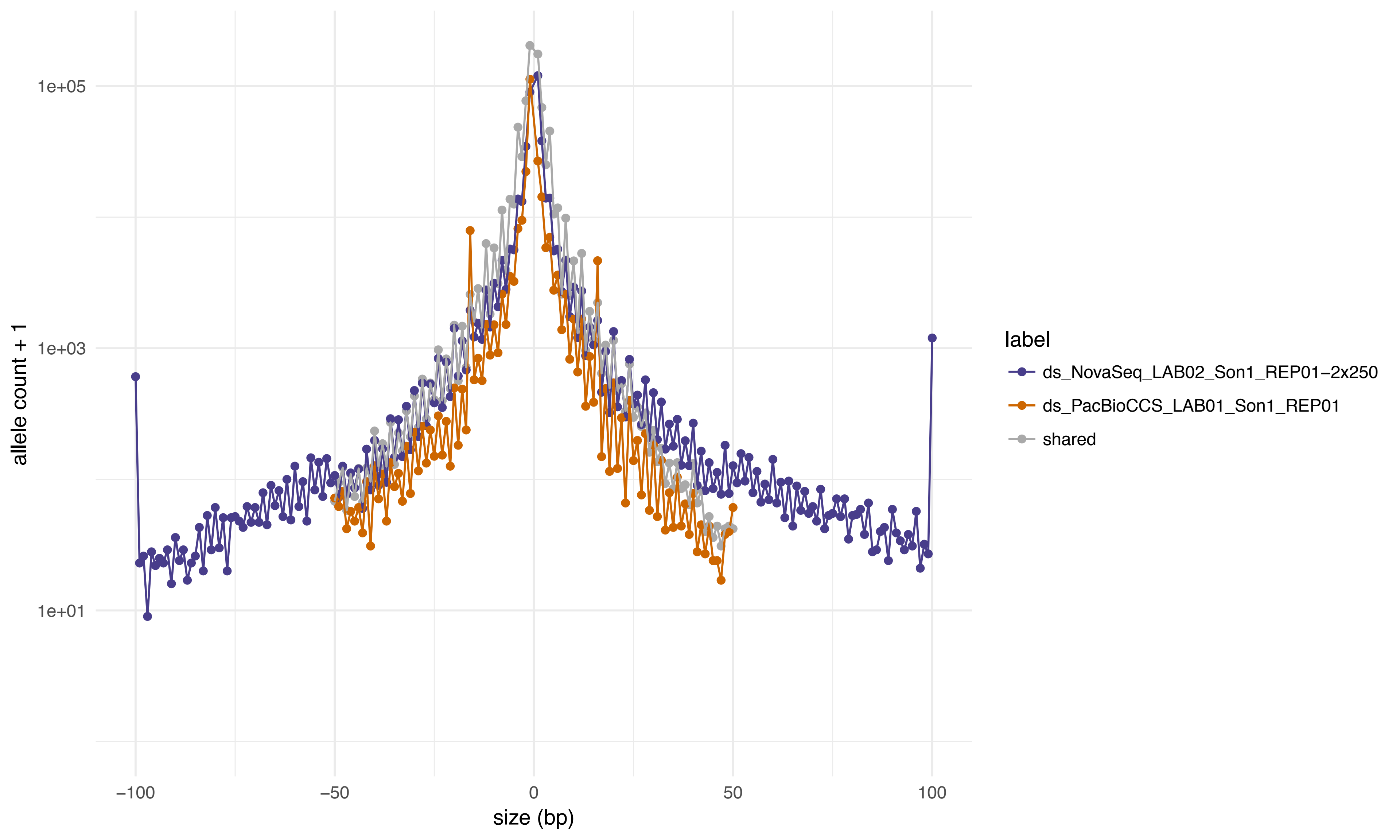

1e+05
1e+03
1e+01
allele count + 1
−100
−50
0
50
100
size (bp)
label
ds_NovaSeq_LAB02_Son1_REP01−2x250
ds_PacBioCCS_LAB01_Son1_REP01
shared

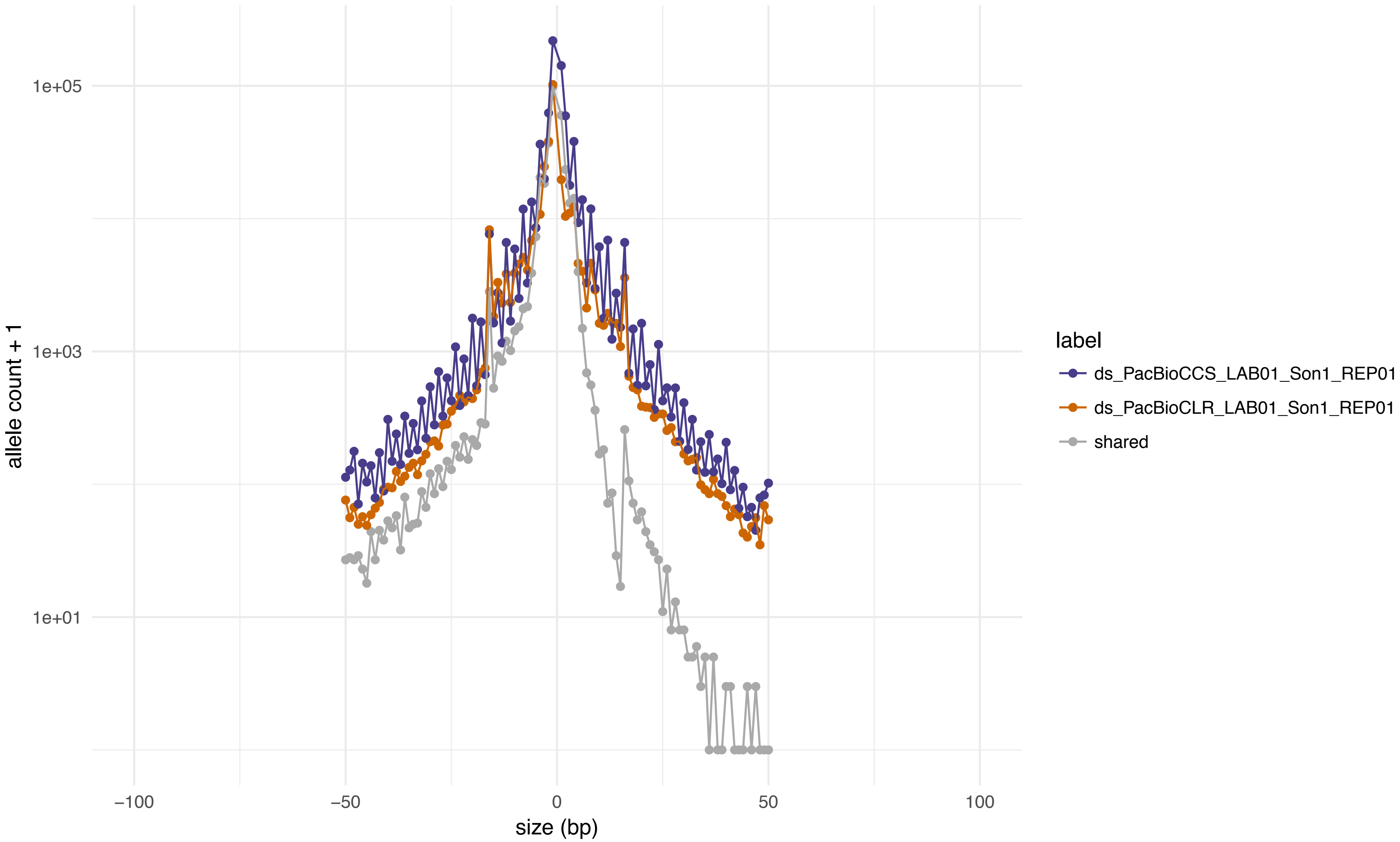

1e+05
1e+03
1e+01
allele count + 1
−100
−50
0
50
100
size (bp)
label
ds_PacBioCCS_LAB01_Son1_REP01
ds_PacBioCLR_LAB01_Son1_REP01
shared

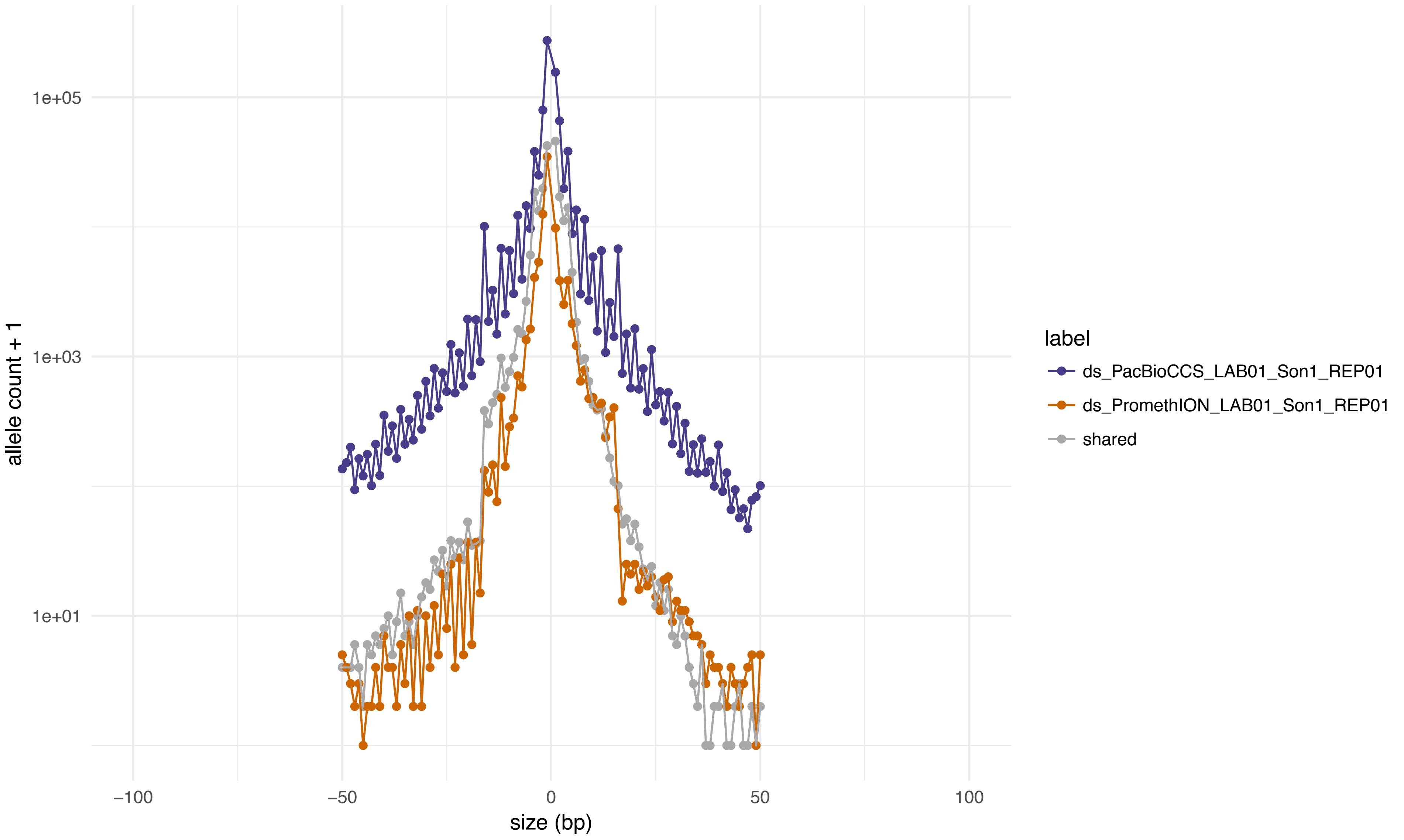

1e+05
1e+03
1e+01
allele count + 1
−100
−50
0
50
100
size (bp)
label
ds_PacBioCCS_LAB01_Son1_REP01
ds_PromethION_LAB01_Son1_REP01
shared

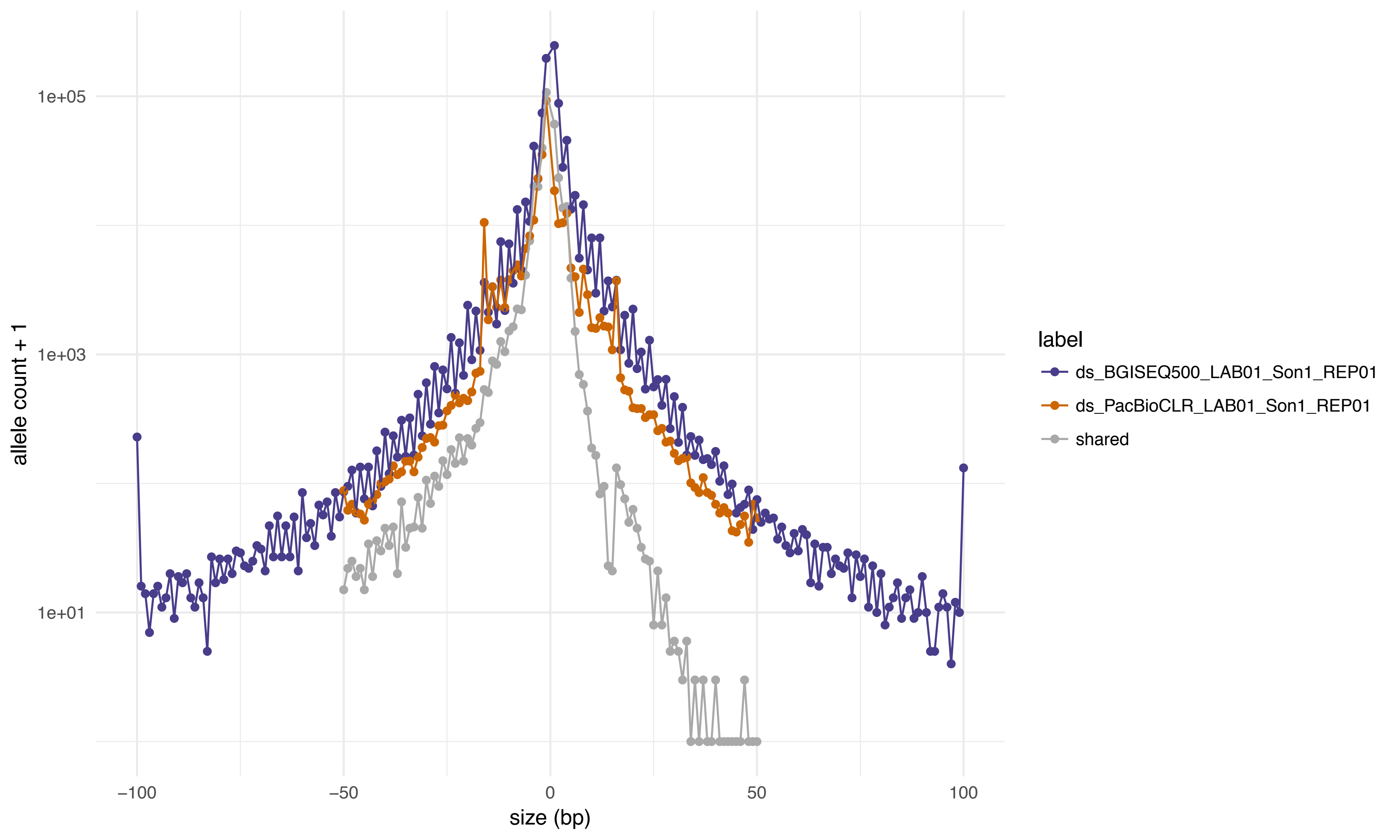

1e+05
1e+03
1e+01
allele count + 1
−100
−50
0
50
100
size (bp)
label
ds_BGISEQ500_LAB01_Son1_REP01
ds_PacBioCLR_LAB01_Son1_REP01
shared

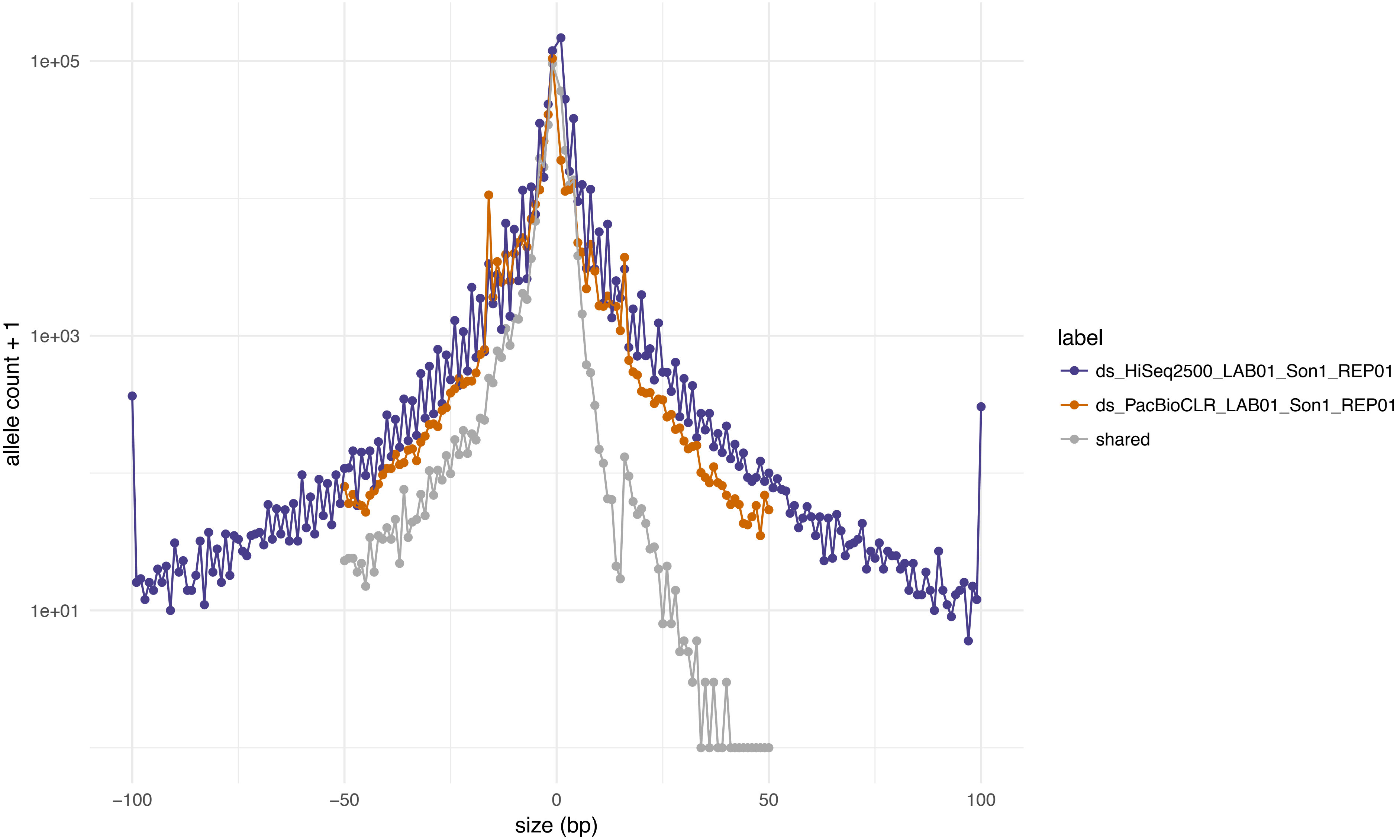
1e+05
1e+03
1e+01
allele count + 1
−100
−50
0
50
100
size (bp)
label
ds_HiSeq2500_LAB01_Son1_REP01
ds_PacBioCLR_LAB01_Son1_REP01
shared

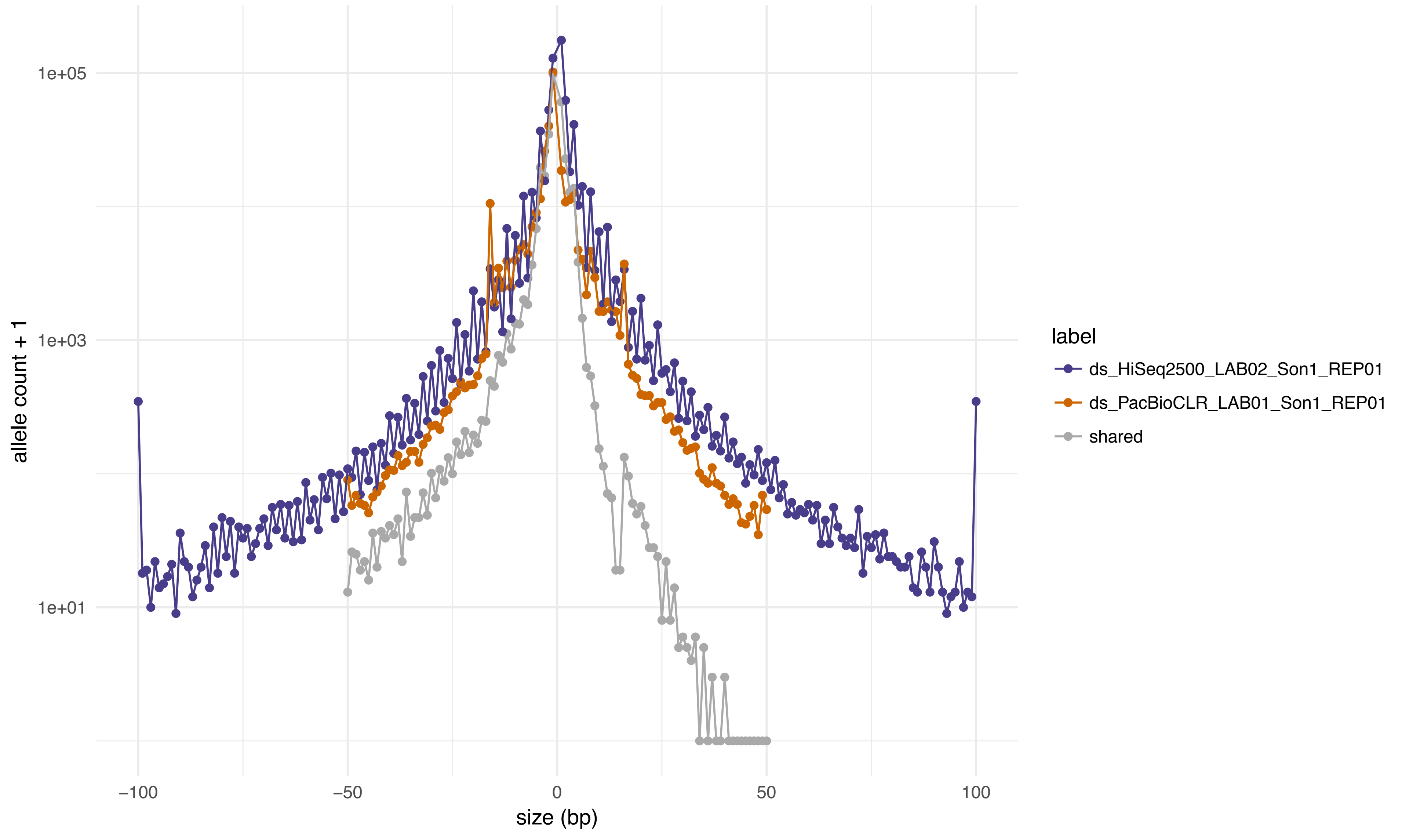

1e+05
1e+03
1e+01
allele count + 1
−100
−50
0
50
100
size (bp)
label
ds_HiSeq2500_LAB02_Son1_REP01
ds_PacBioCLR_LAB01_Son1_REP01
shared

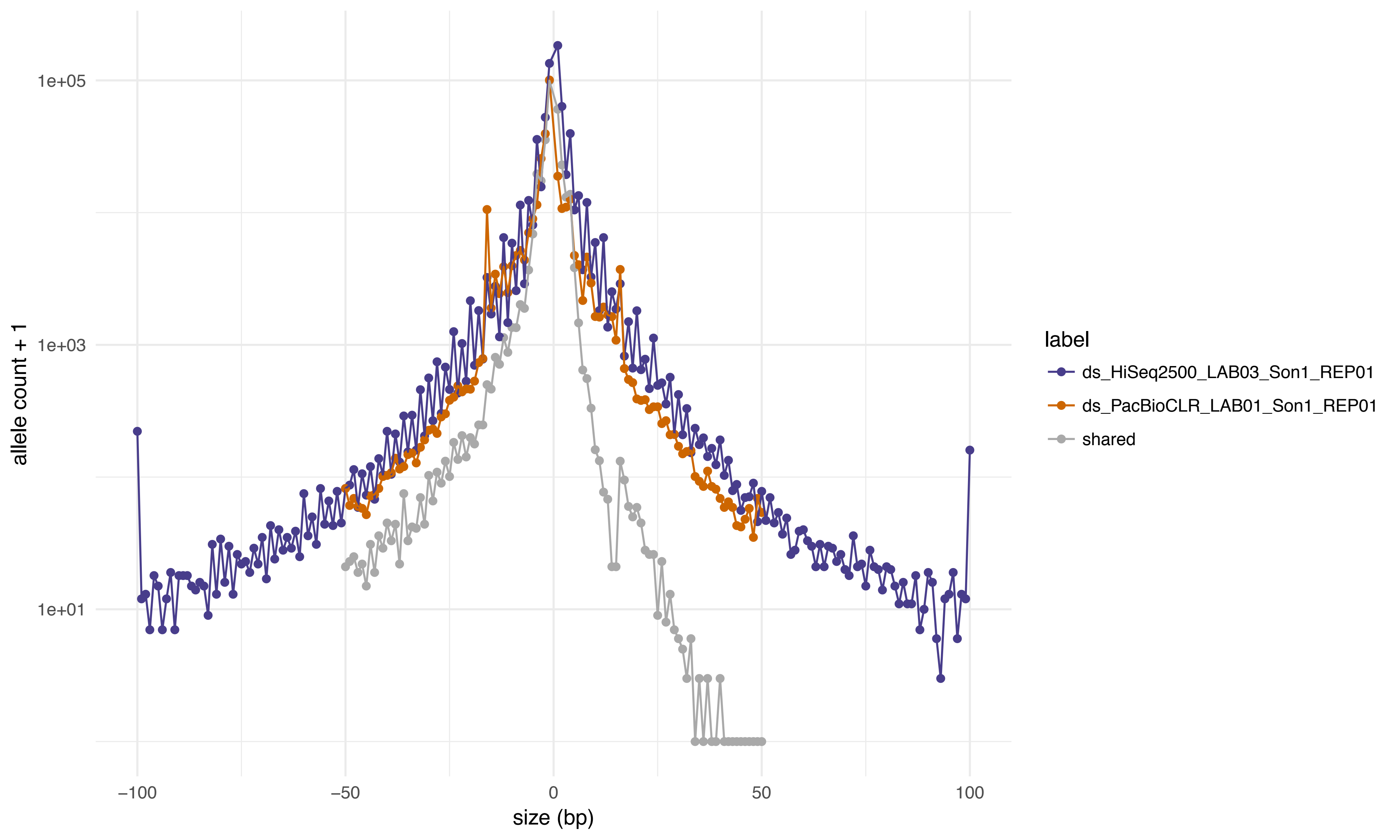

allele count + 1
1e+05
1e+03
1e+01
−100
−50
0
50
100
size (bp)
label
ds_HiSeq2500_LAB03_Son1_REP01
ds_PacBioCLR_LAB01_Son1_REP01
shared

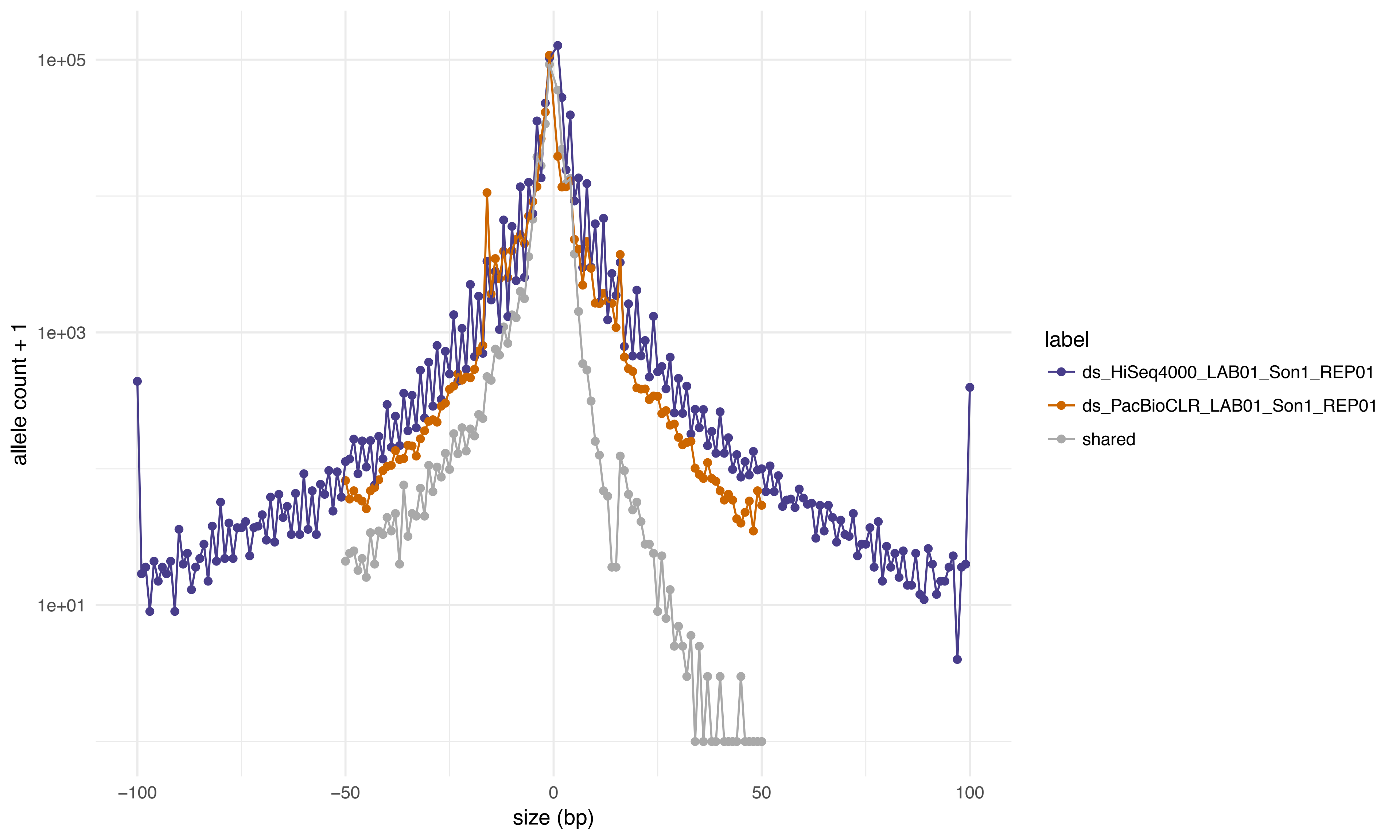

1e+05
1e+03
1e+01
allele count + 1
−100
−50
0
50
100
size (bp)
label
ds_HiSeq4000_LAB01_Son1_REP01
ds_PacBioCLR_LAB01_Son1_REP01
shared

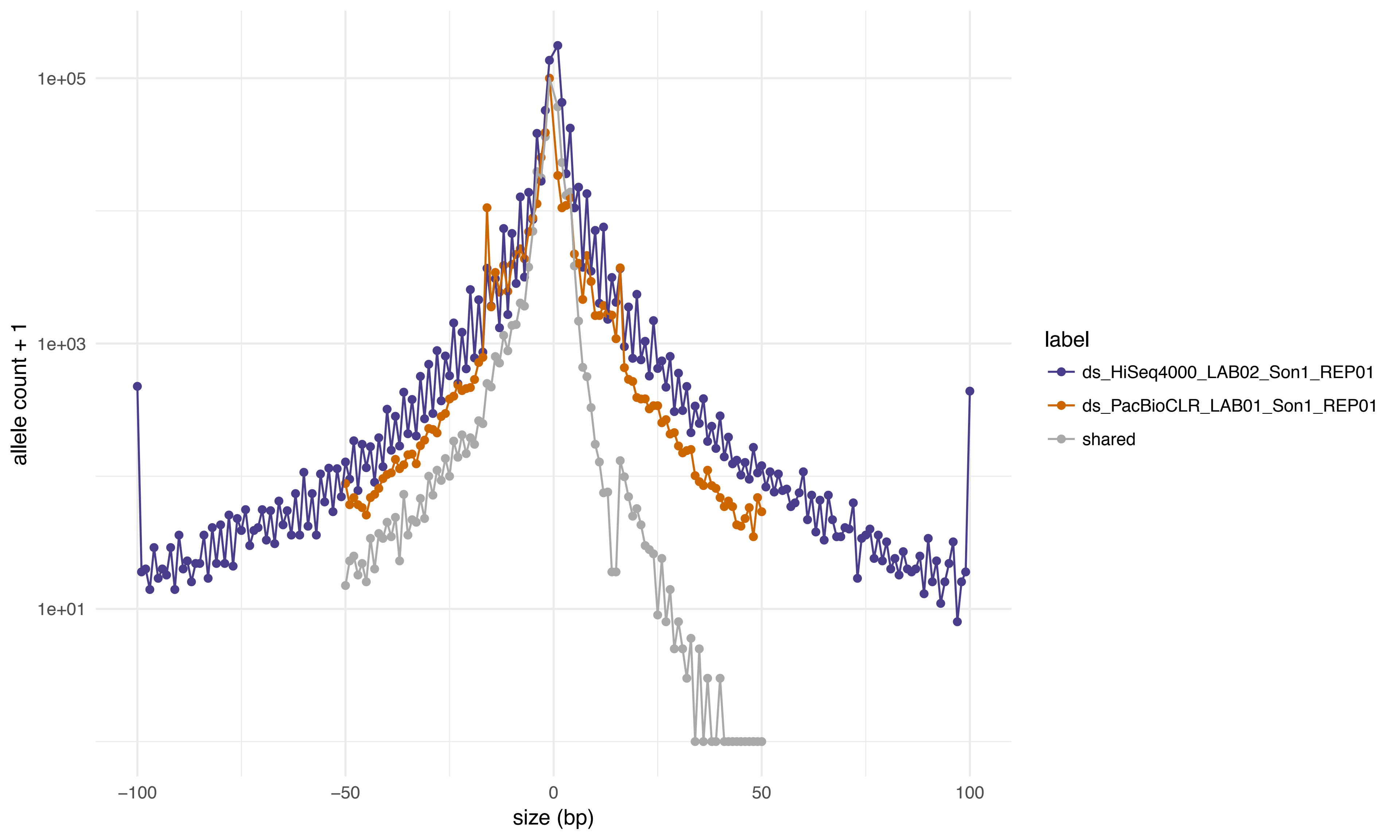

1e+05
1e+03
1e+01
allele count + 1
−100
−50
0
50
100
size (bp)
label
ds_HiSeq4000_LAB02_Son1_REP01
ds_PacBioCLR_LAB01_Son1_REP01
shared

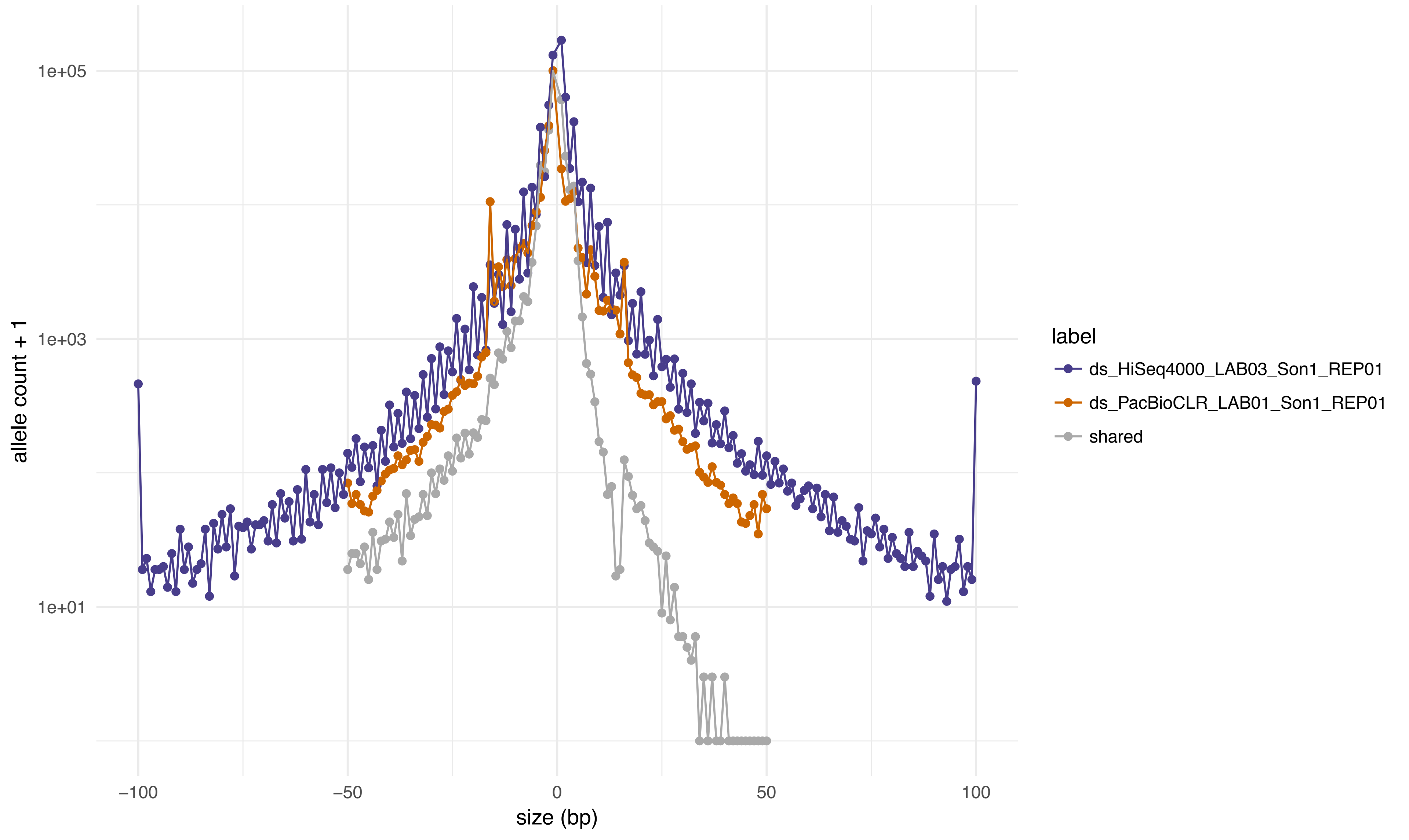
1e+05
1e+03
1e+01
allele count + 1
−100
−50
0
50
100
size (bp)
label
ds_HiSeq4000_LAB03_Son1_REP01
ds_PacBioCLR_LAB01_Son1_REP01
shared

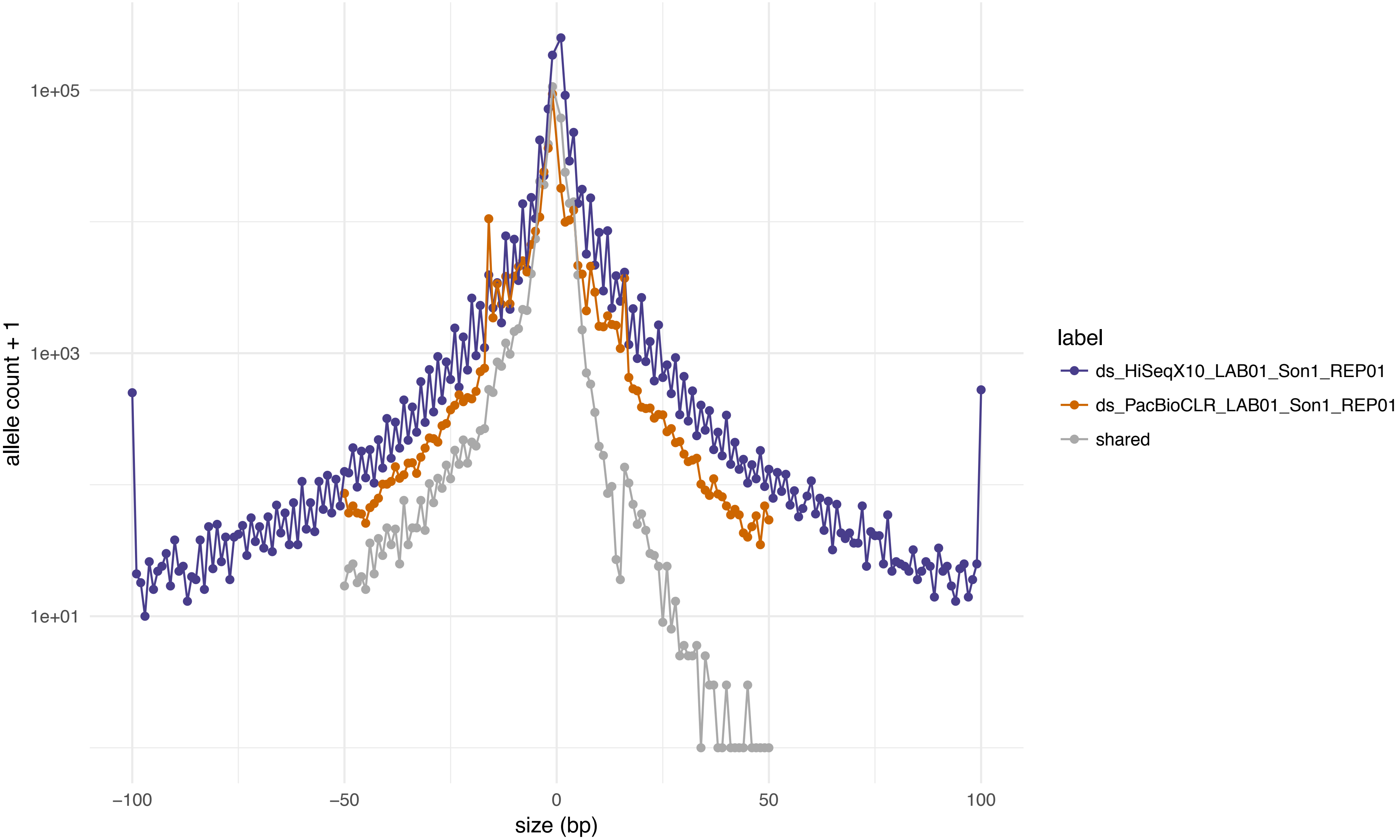

allele count + 1
1e+05
1e+03
1e+01
−100
−50
0
50
100
size (bp)
label
ds_HiSeqX10_LAB01_Son1_REP01
ds_PacBioCLR_LAB01_Son1_REP01
shared

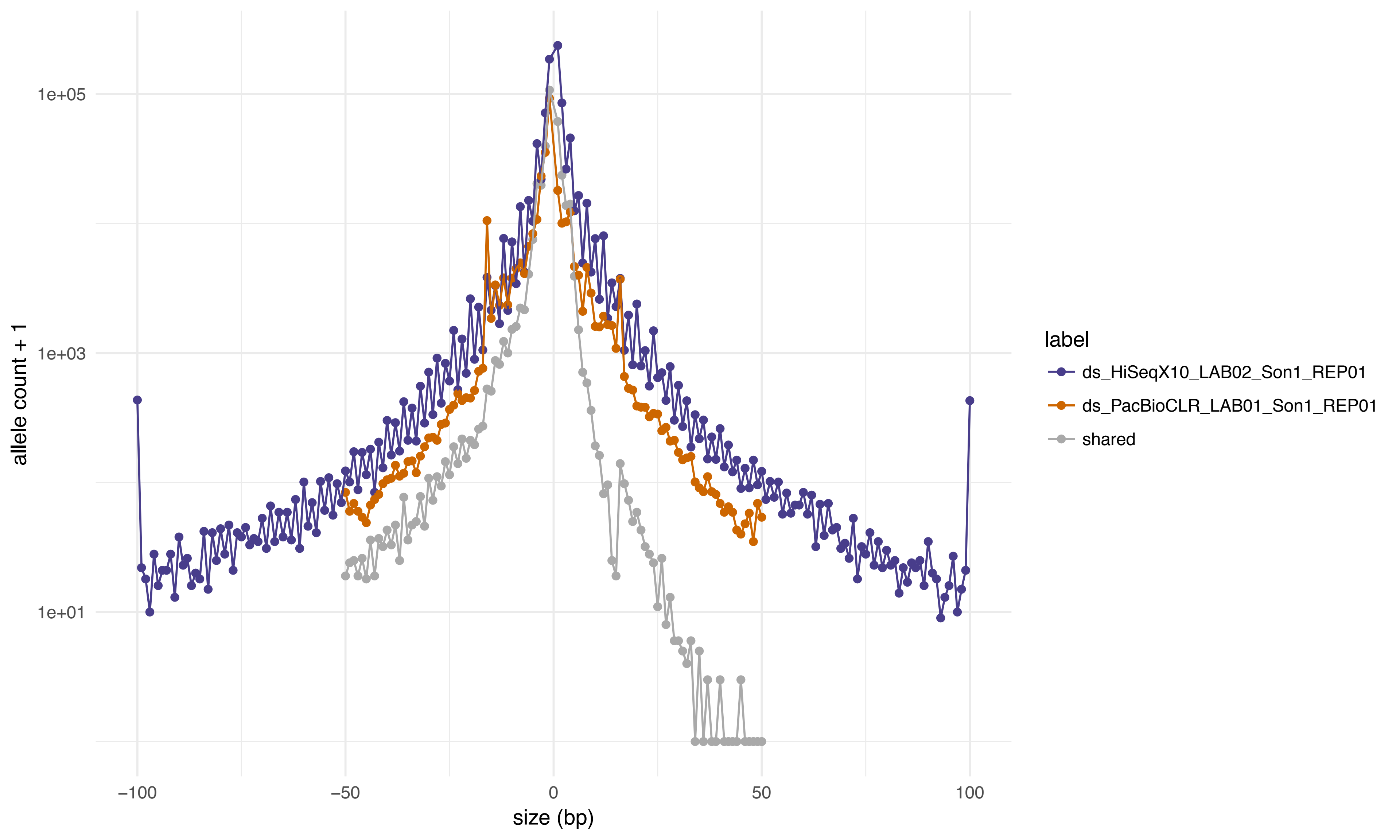

1e+05
1e+03
1e+01
allele count + 1
−100
−50
0
50
100
size (bp)
label
ds_HiSeqX10_LAB02_Son1_REP01
ds_PacBioCLR_LAB01_Son1_REP01
shared

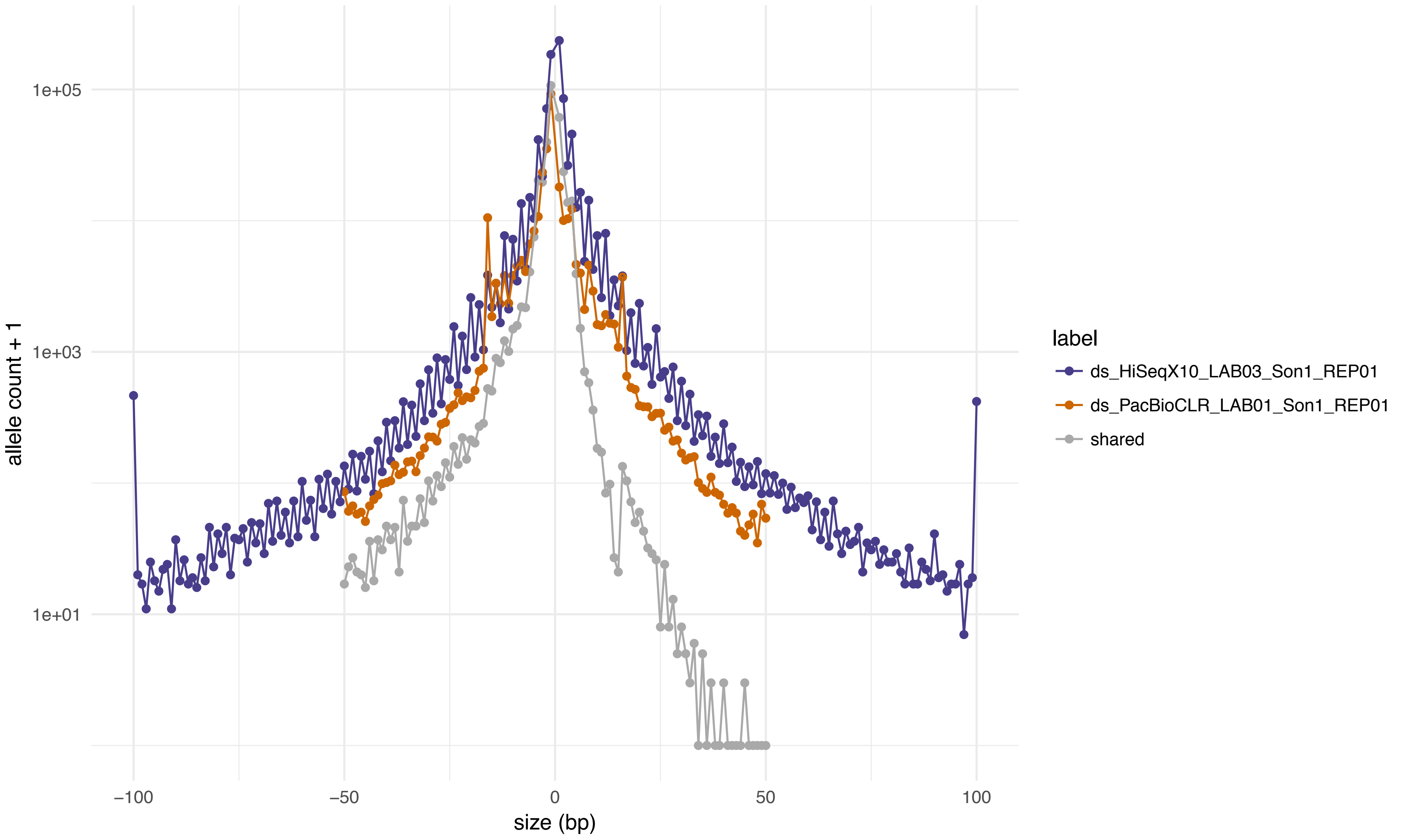
1e+05
1e+03
1e+01
allele count + 1
−100
−50
0
50
100
size (bp)
label
ds_HiSeqX10_LAB03_Son1_REP01
ds_PacBioCLR_LAB01_Son1_REP01
shared

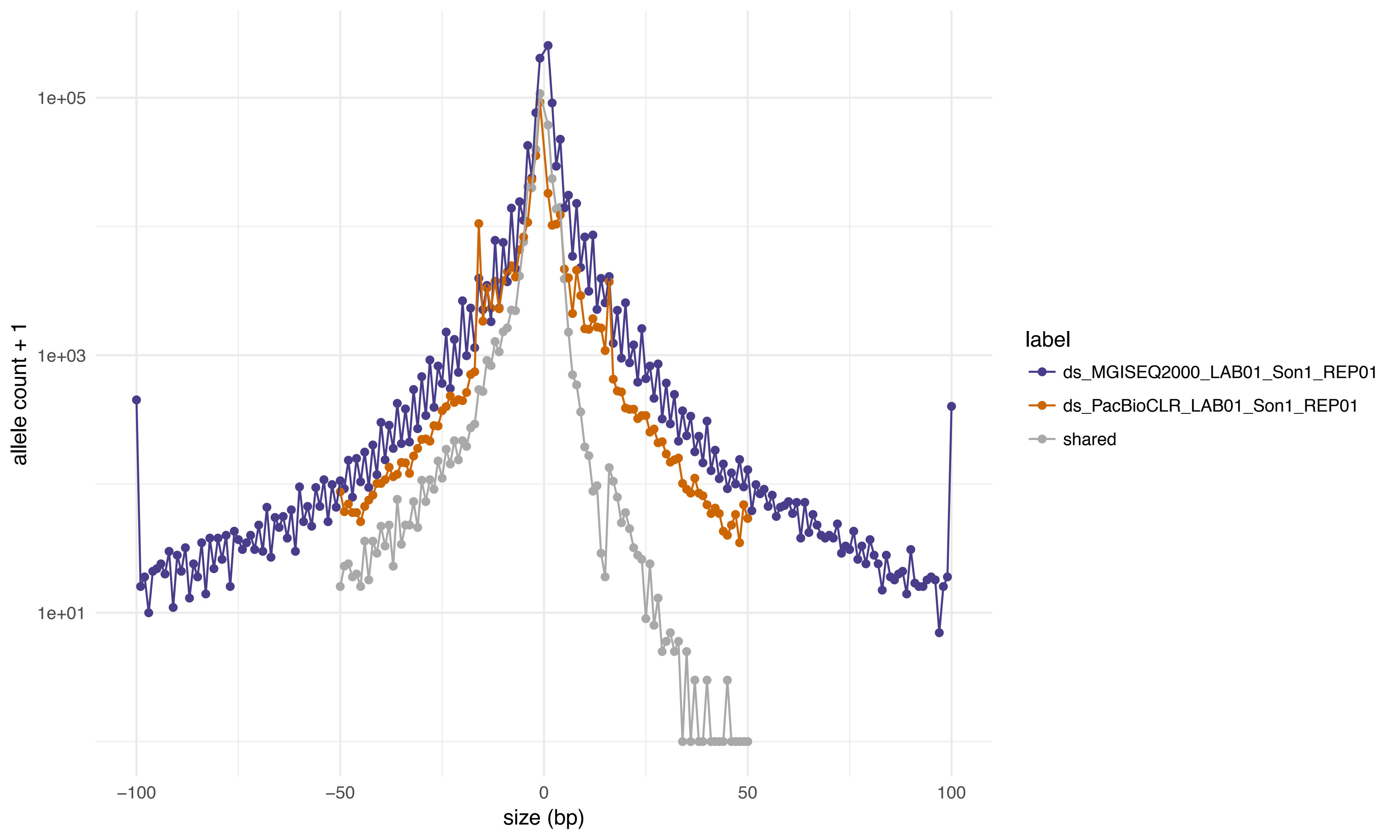
1e+05
1e+03
1e+01
allele count + 1
−100
−50
0
50
100
size (bp)
label
ds_MGISEQ2000_LAB01_Son1_REP01
ds_PacBioCLR_LAB01_Son1_REP01
shared

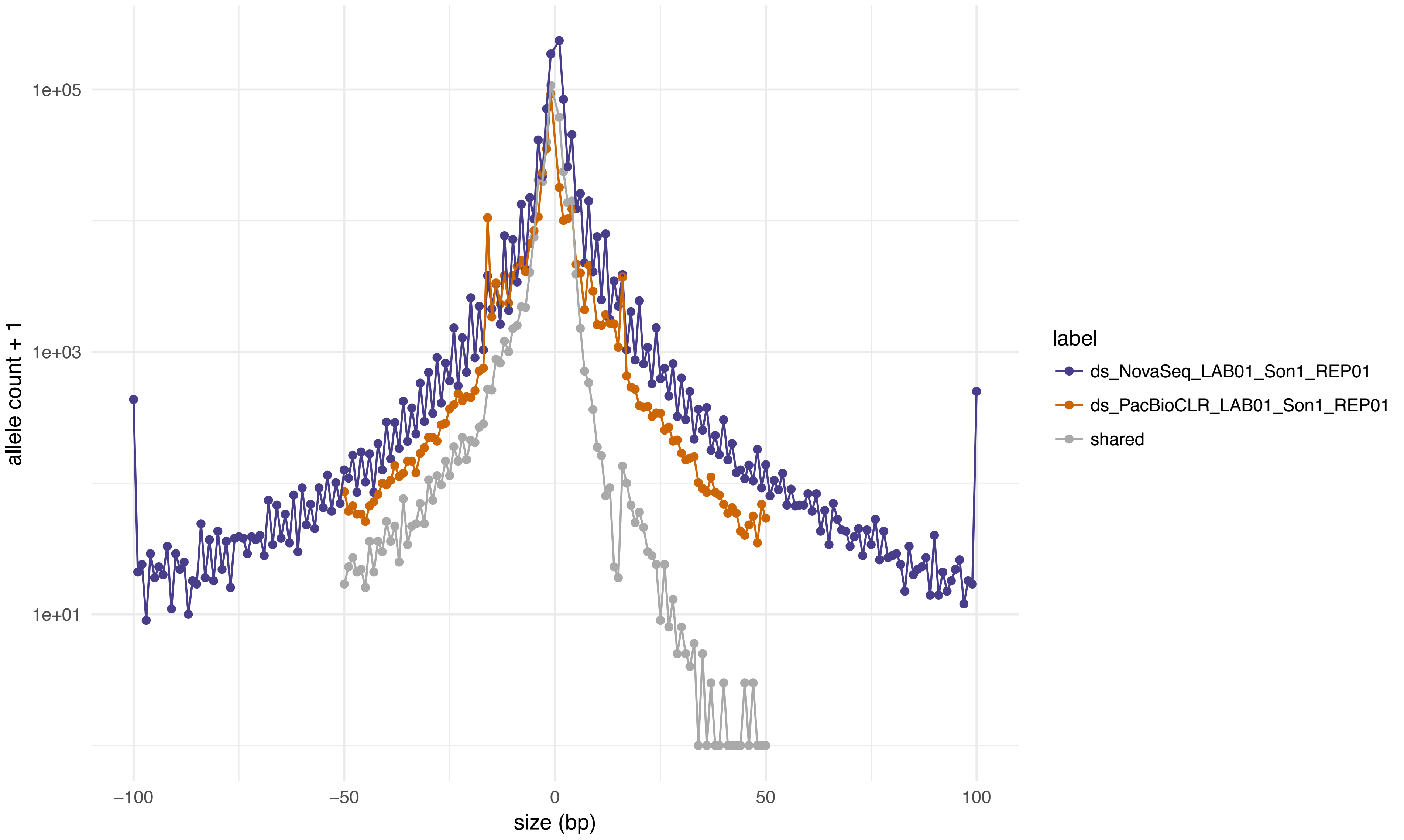
1e+05
1e+03
1e+01
allele count + 1
−100
−50
0
50
100
size (bp)
label
ds_NovaSeq_LAB01_Son1_REP01
ds_PacBioCLR_LAB01_Son1_REP01
shared

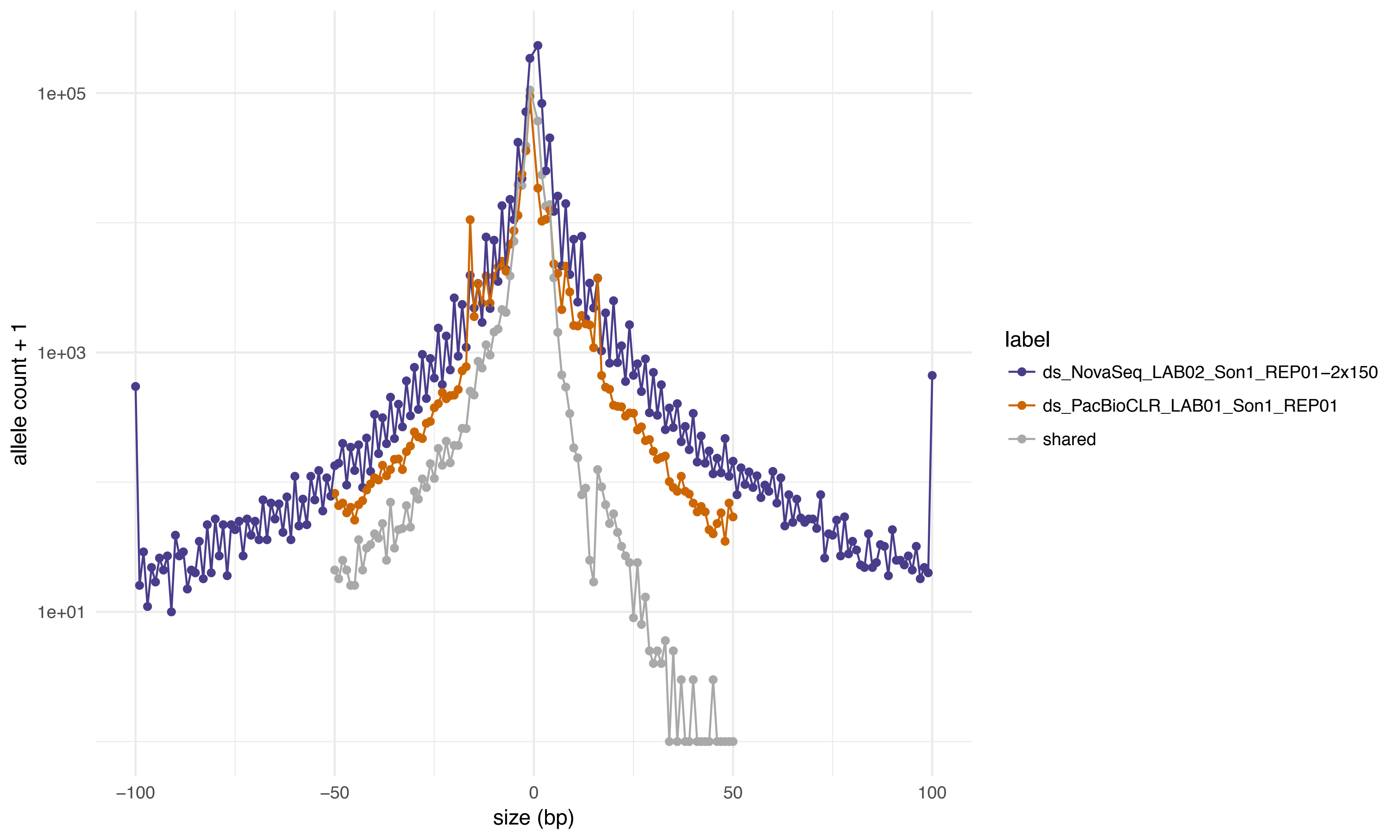

allele count + 1
1e+05
1e+03
1e+01
−100
−50
0
50
100
size (bp)
label
ds_NovaSeq_LAB02_Son1_REP01−2x150
ds_PacBioCLR_LAB01_Son1_REP01
shared

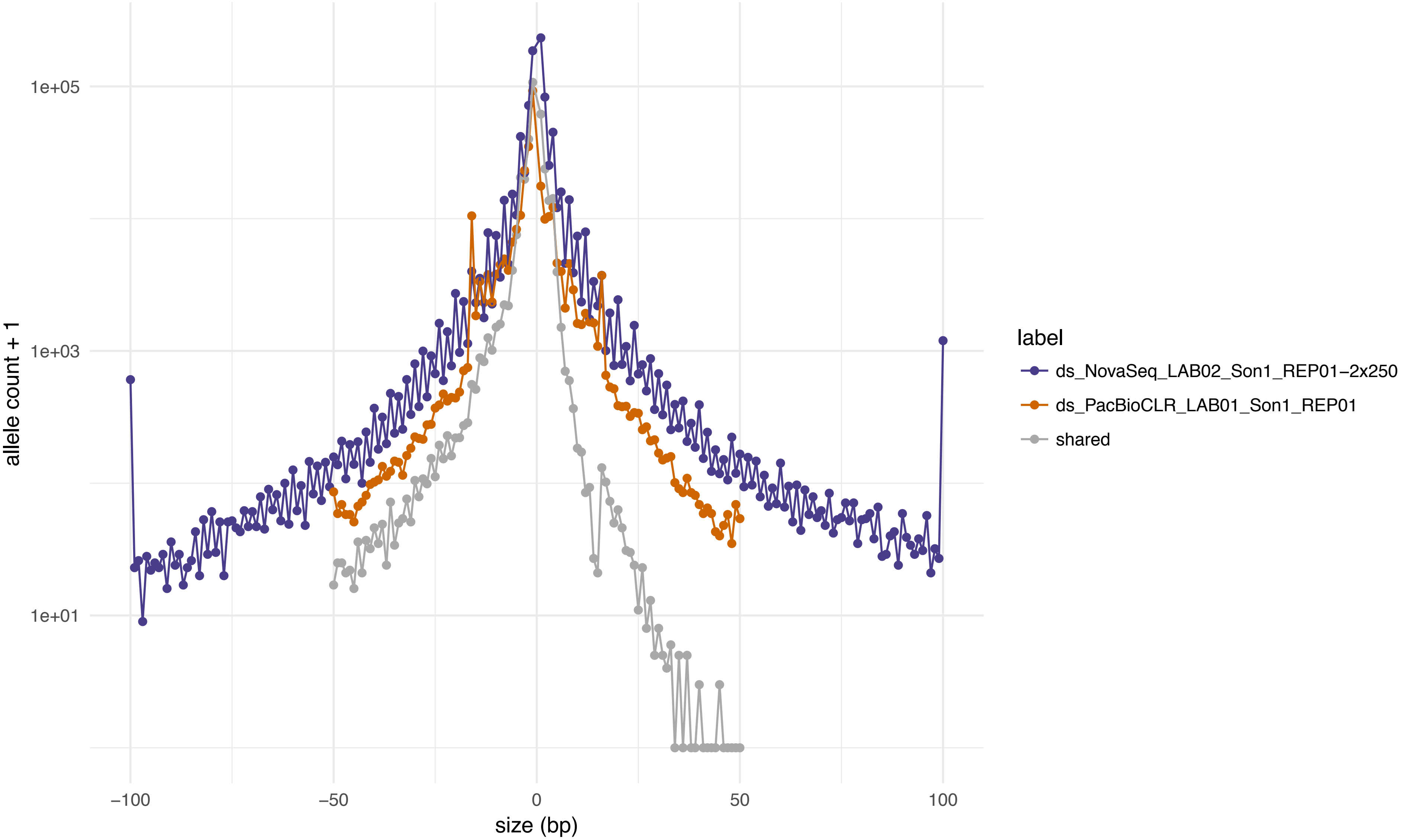

1e+05
1e+03
1e+01
allele count + 1
−100
−50
0
50
100
size (bp)
label
ds_NovaSeq_LAB02_Son1_REP01−2x250
ds_PacBioCLR_LAB01_Son1_REP01
shared

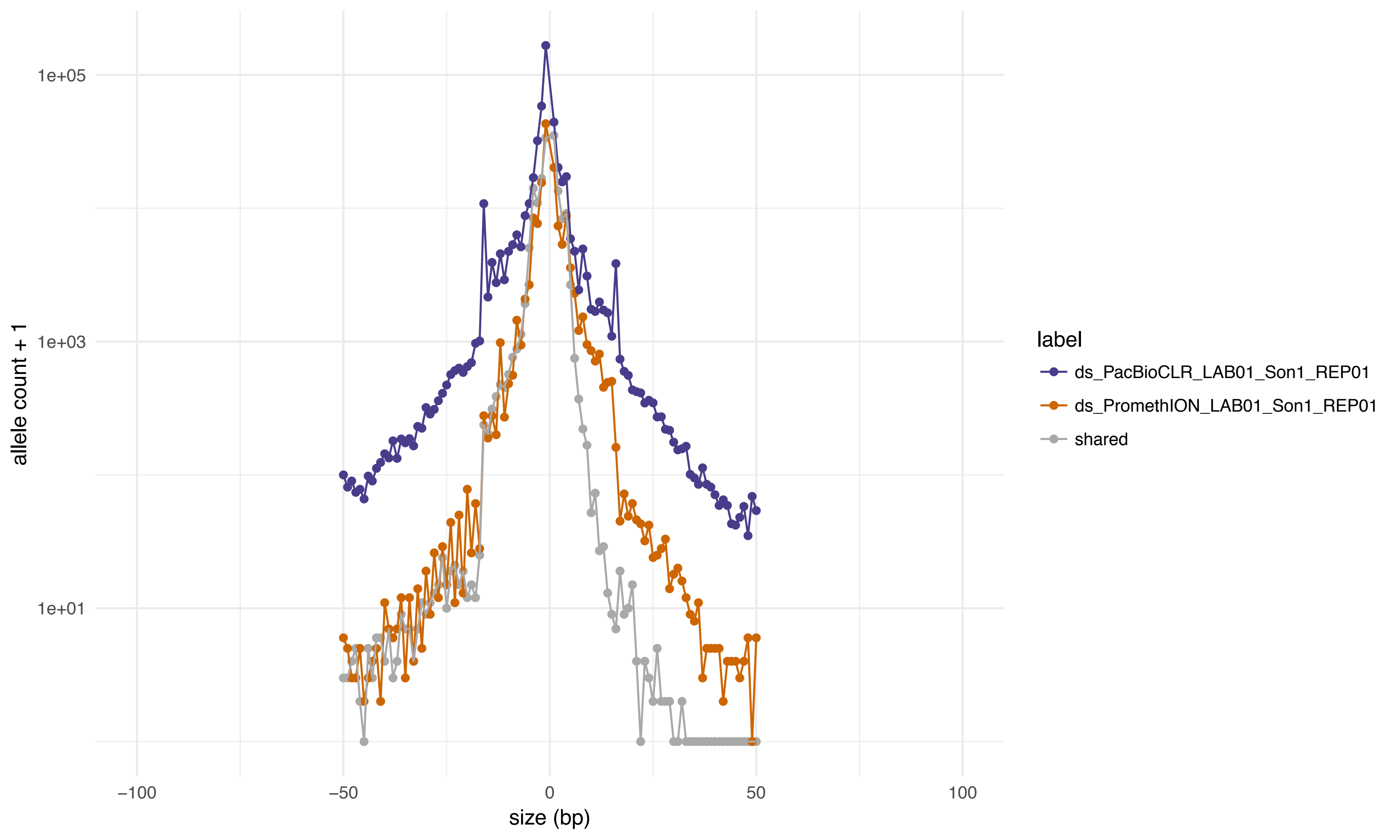
1e+05
1e+03
1e+01
allele count + 1
−100
−50
0
50
100
size (bp)
label
ds_PacBioCLR_LAB01_Son1_REP01
ds_PromethION_LAB01_Son1_REP01
shared

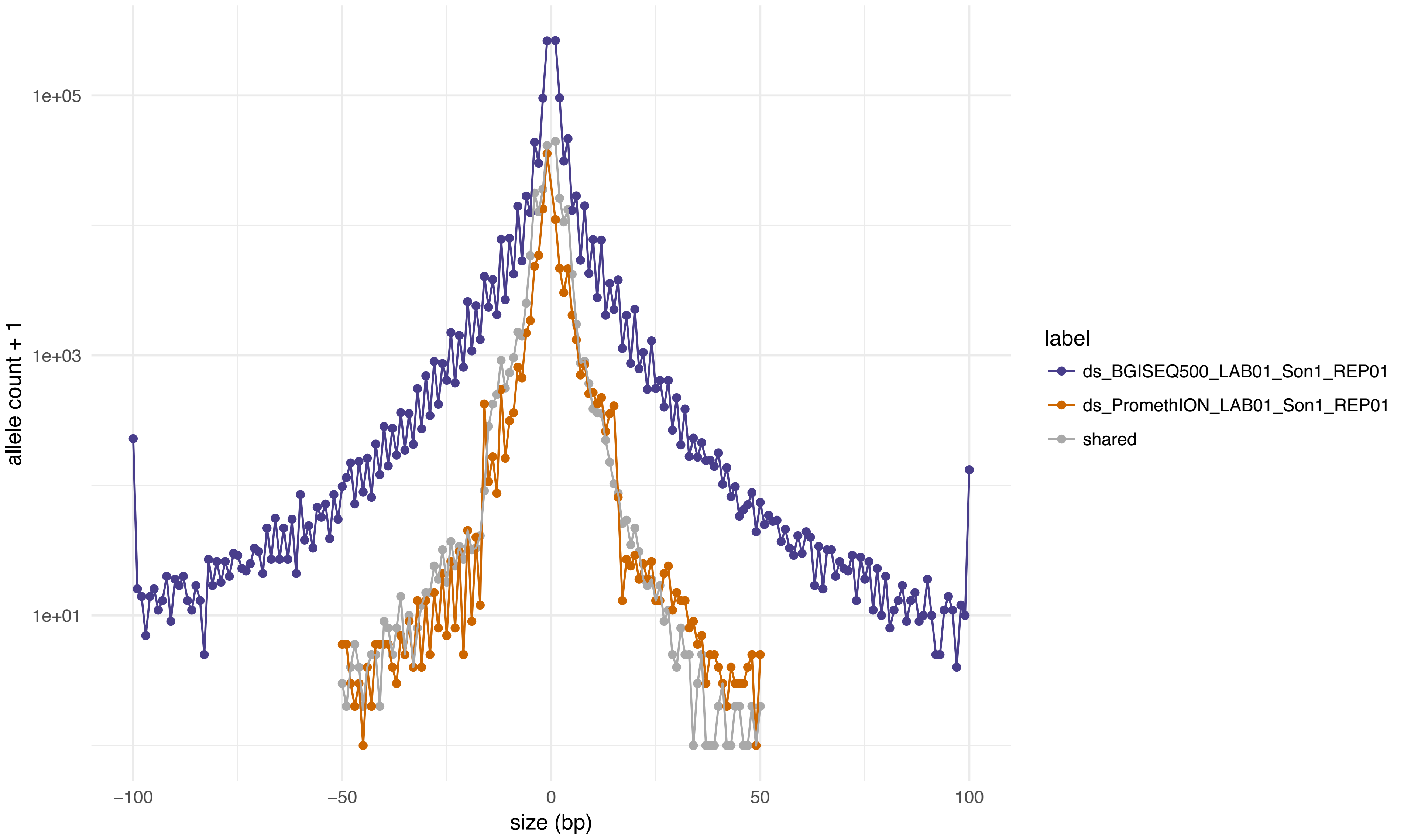

1e+05
1e+03
1e+01
allele count + 1
−100
−50
0
50
100
size (bp)
label
ds_BGISEQ500_LAB01_Son1_REP01
ds_PromethION_LAB01_Son1_REP01
shared

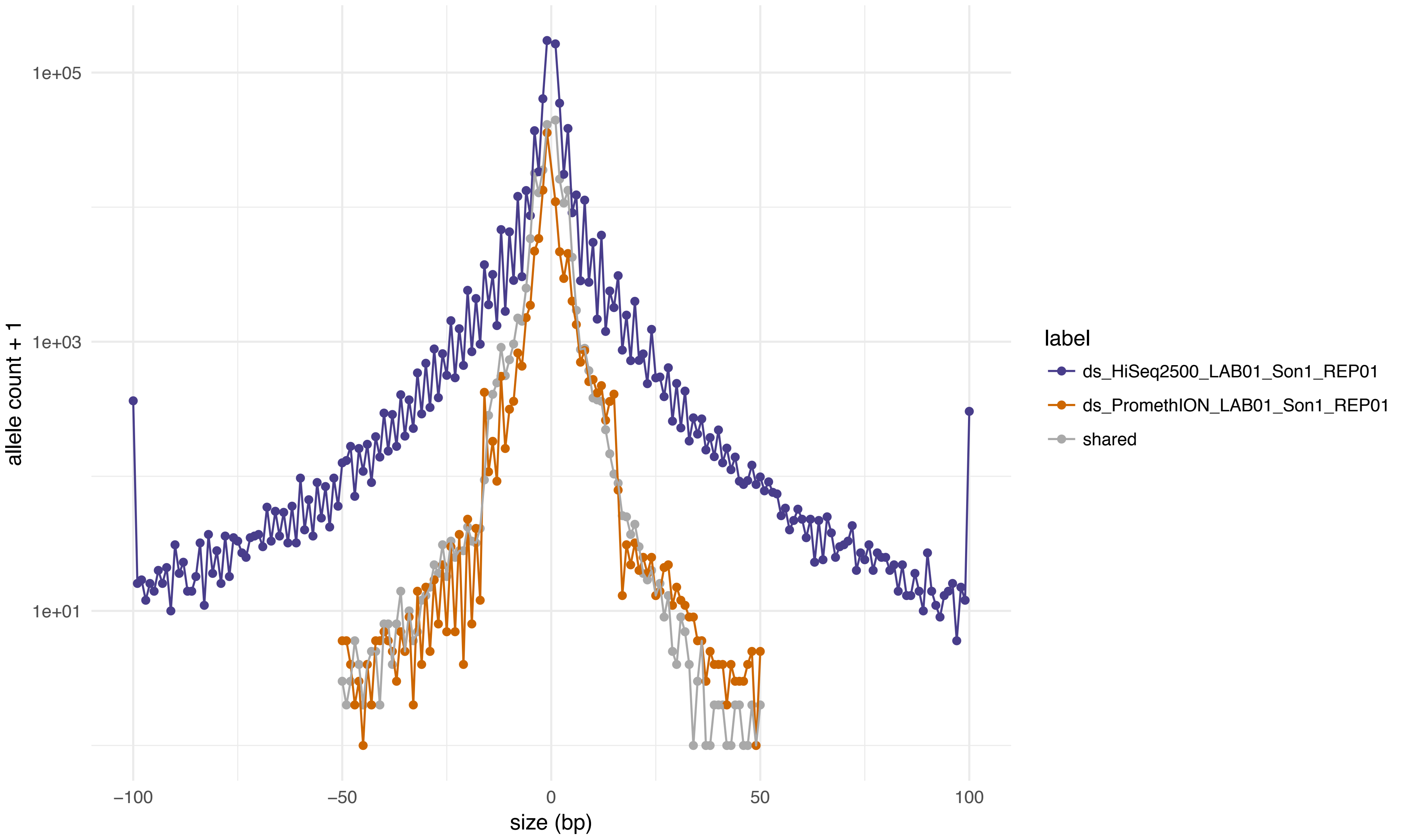
1e+05
1e+03
1e+01
allele count + 1
−100
−50
0
50
100
size (bp)
label
ds_HiSeq2500_LAB01_Son1_REP01
ds_PromethION_LAB01_Son1_REP01
shared

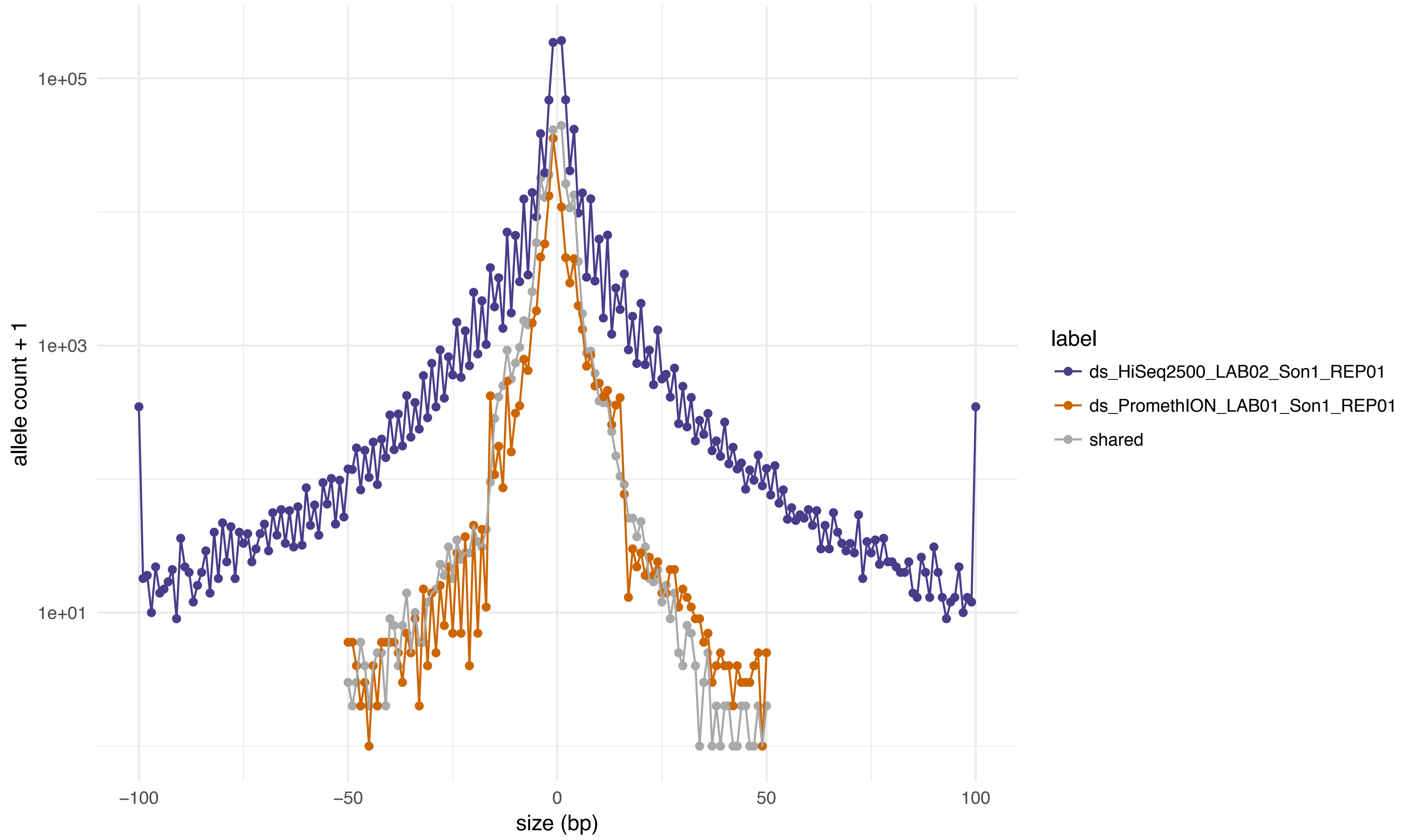

allele count + 1
1e+05
1e+03
1e+01
−100
−50
0
50
100
size (bp)
label
ds_HiSeq2500_LAB02_Son1_REP01
ds_PromethION_LAB01_Son1_REP01
shared

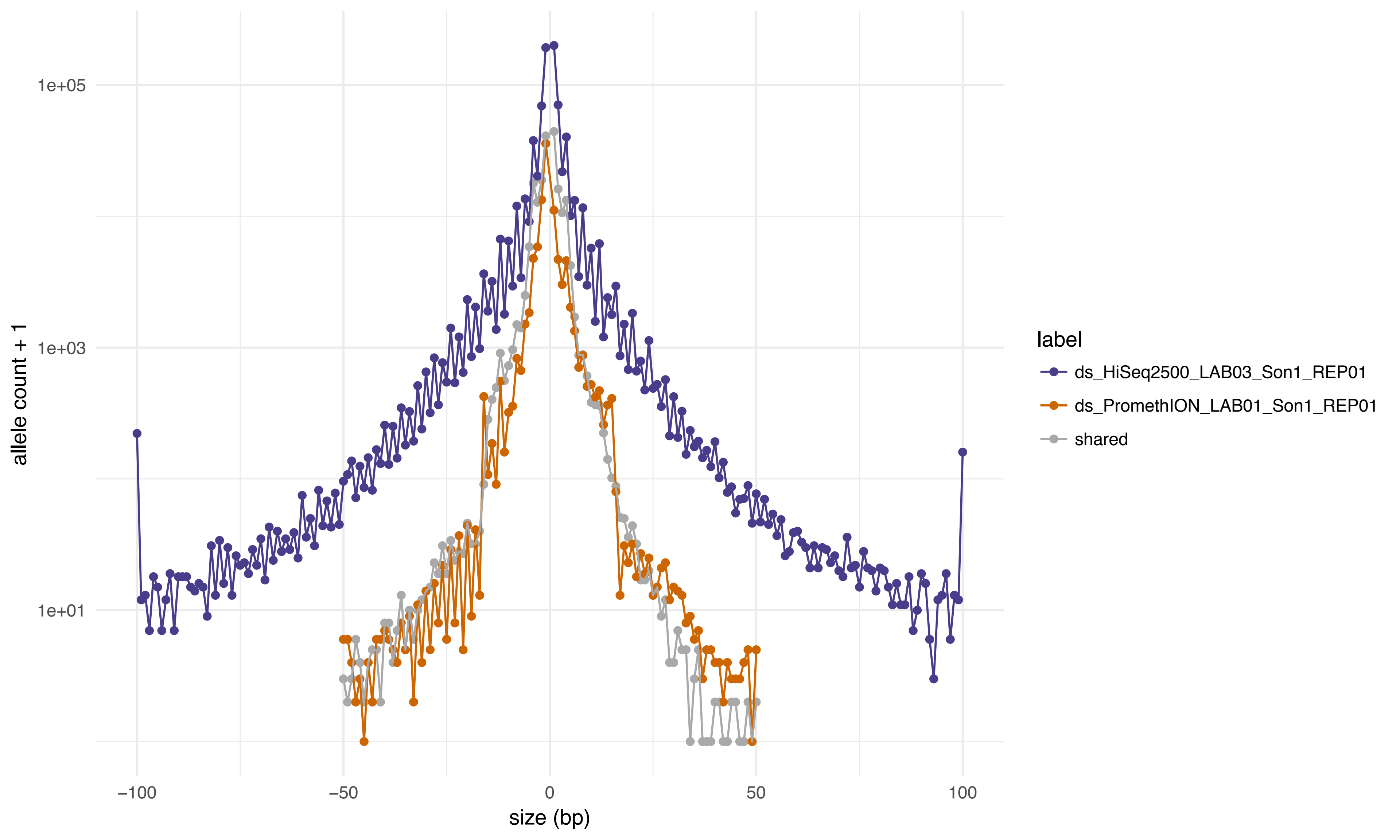
1e+05
1e+03
1e+01
allele count + 1
−100
−50
0
50
100
size (bp)
label
ds_HiSeq2500_LAB03_Son1_REP01
ds_PromethION_LAB01_Son1_REP01
shared

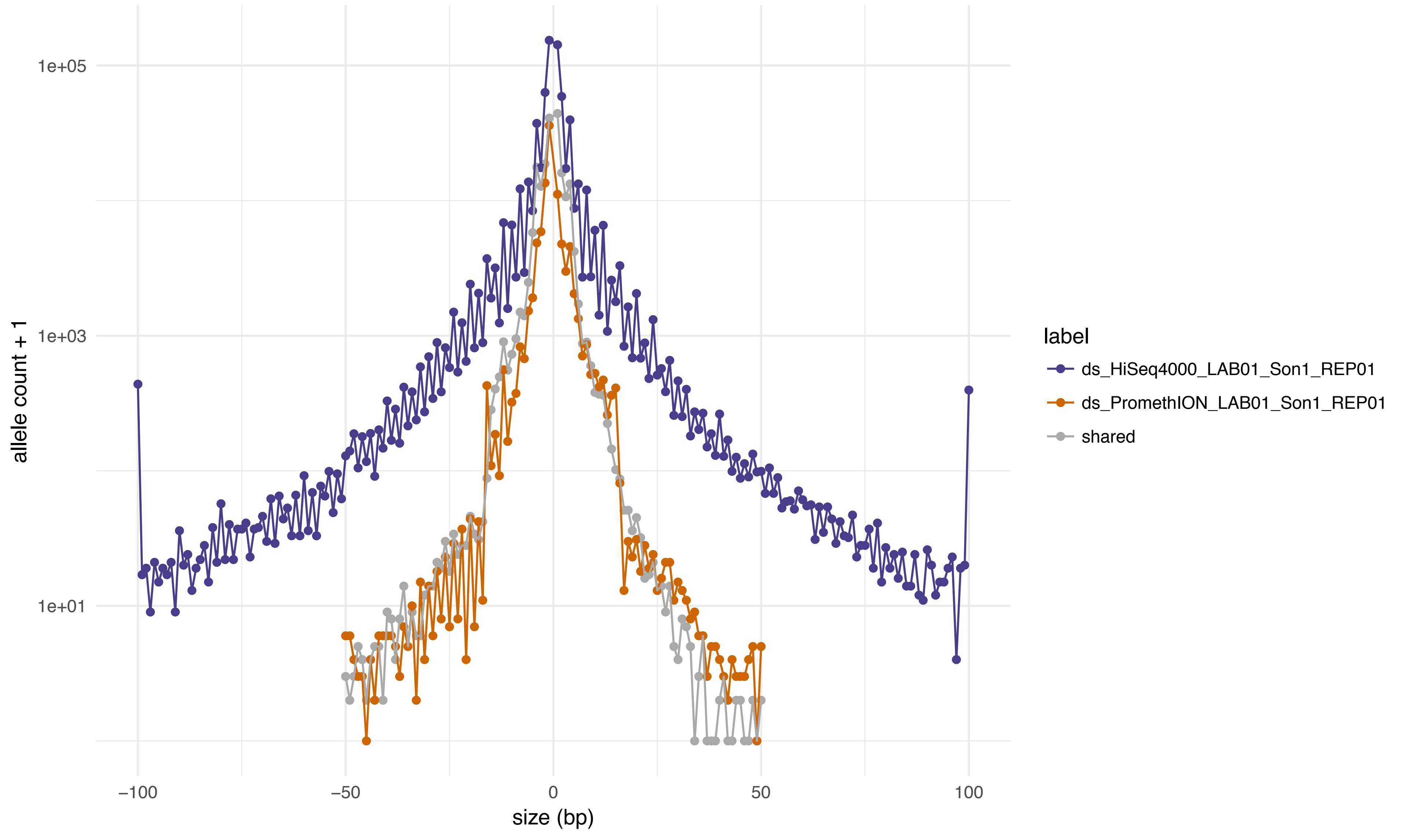

1e+05
1e+03
1e+01
allele count + 1
−100
−50
0
50
100
size (bp)
label
ds_HiSeq4000_LAB01_Son1_REP01
ds_PromethION_LAB01_Son1_REP01
shared

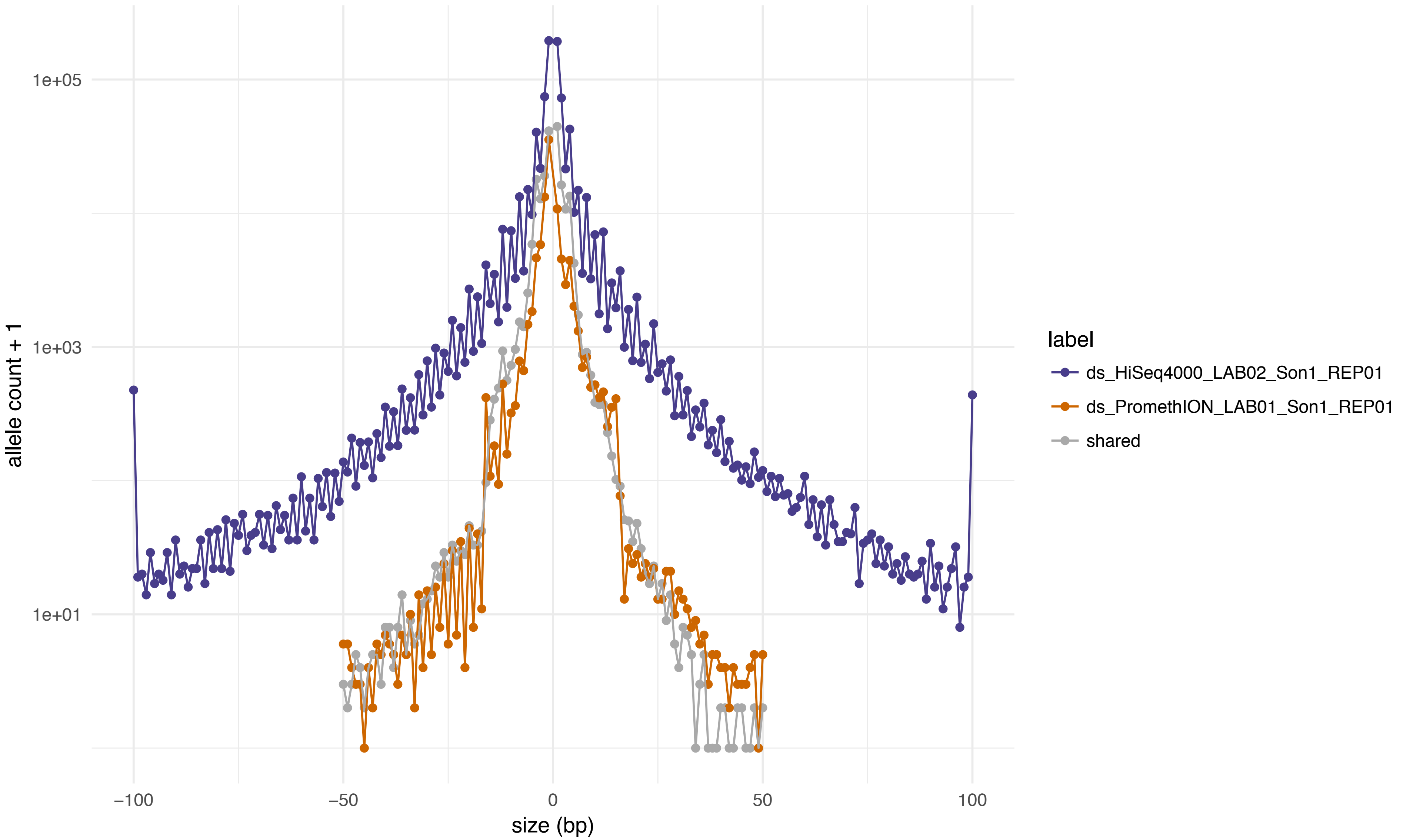
1e+05
1e+03
1e+01
allele count + 1
−100
−50
0
50
100
size (bp)
label
ds_HiSeq4000_LAB02_Son1_REP01
ds_PromethION_LAB01_Son1_REP01
shared

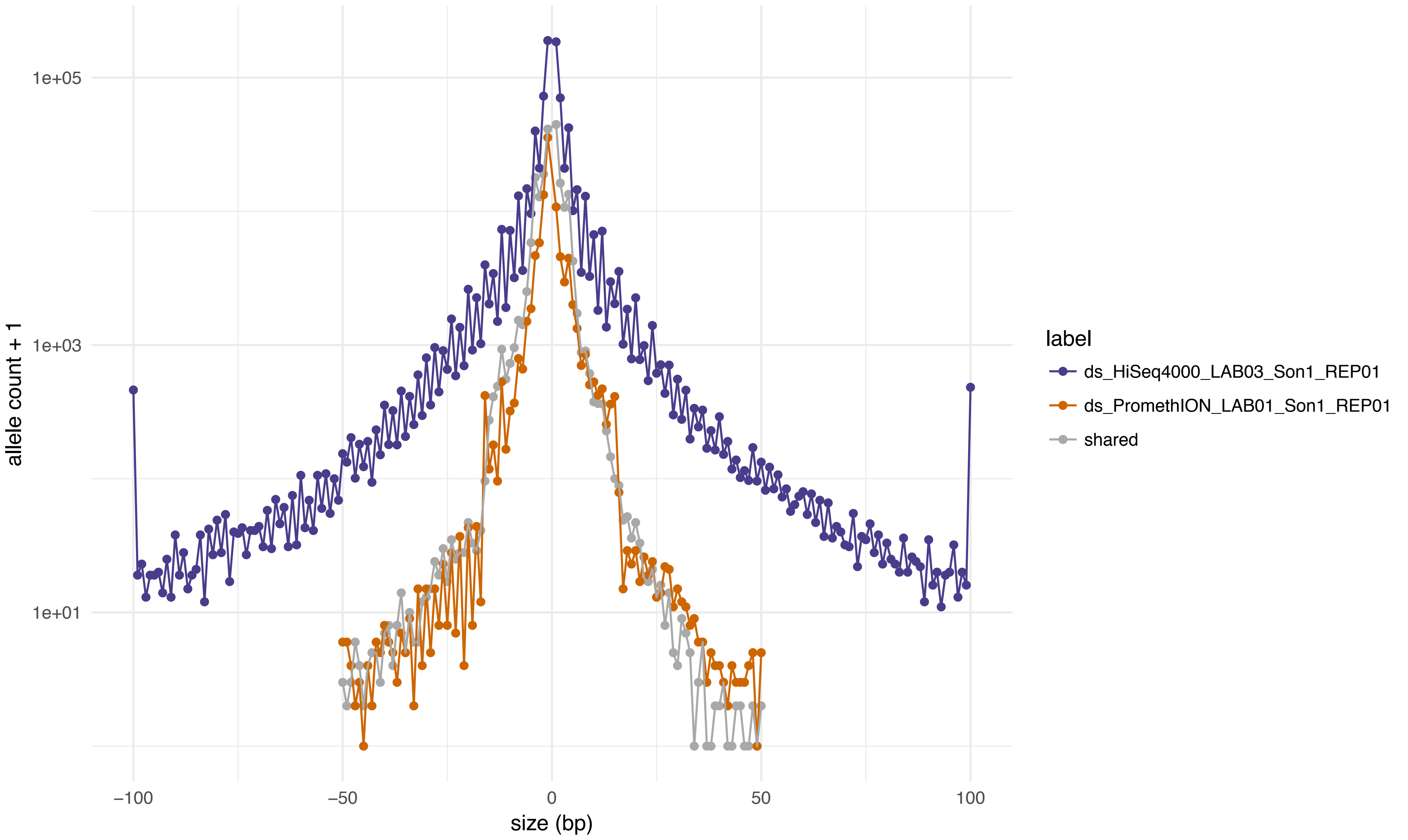
1e+05
1e+03
1e+01
allele count + 1
−100
−50
0
50
100
size (bp)
label
ds_HiSeq4000_LAB03_Son1_REP01
ds_PromethION_LAB01_Son1_REP01
shared

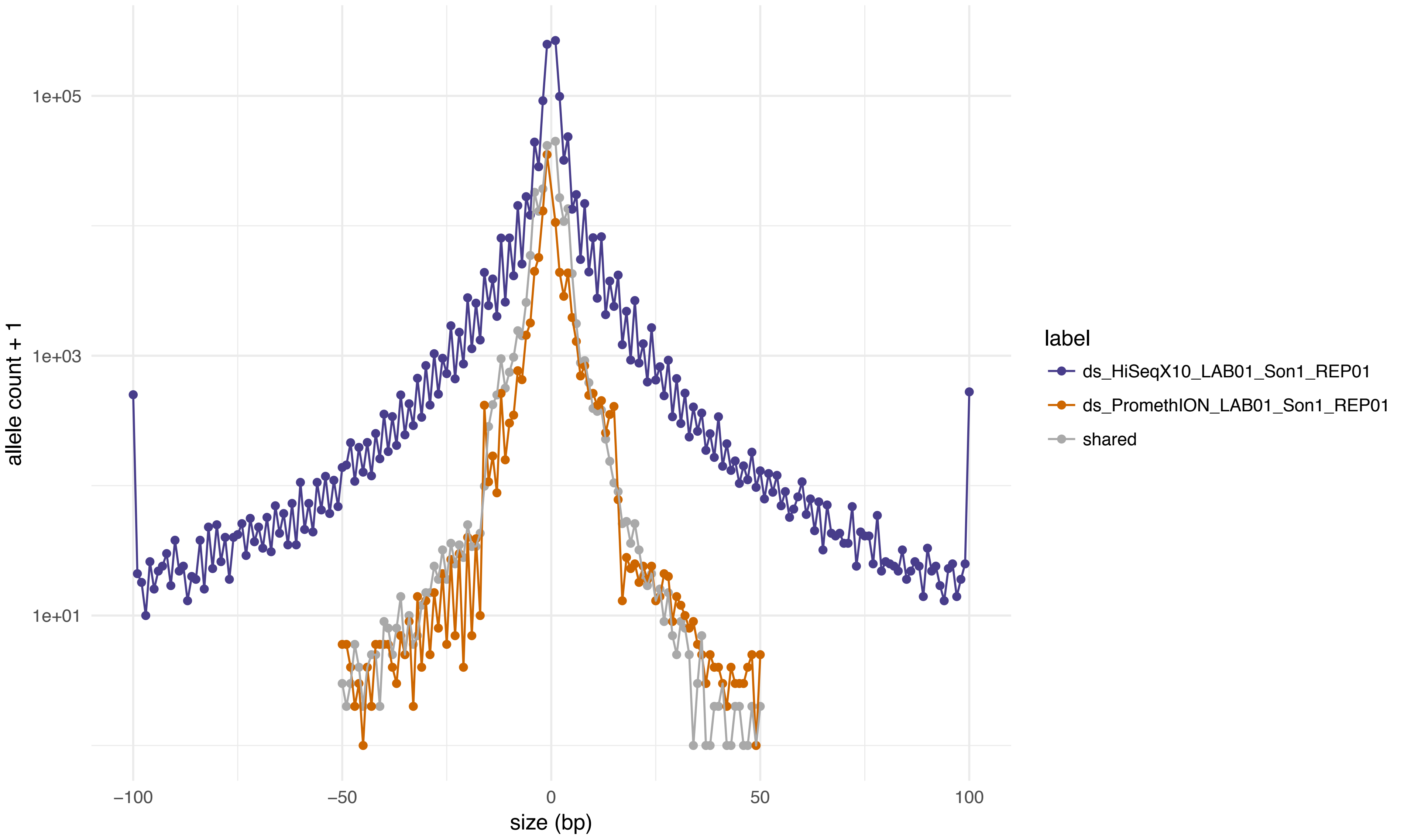

1e+05
1e+03
1e+01
allele count + 1
−100
−50
0
50
100
size (bp)
label
ds_HiSeqX10_LAB01_Son1_REP01
ds_PromethION_LAB01_Son1_REP01
shared

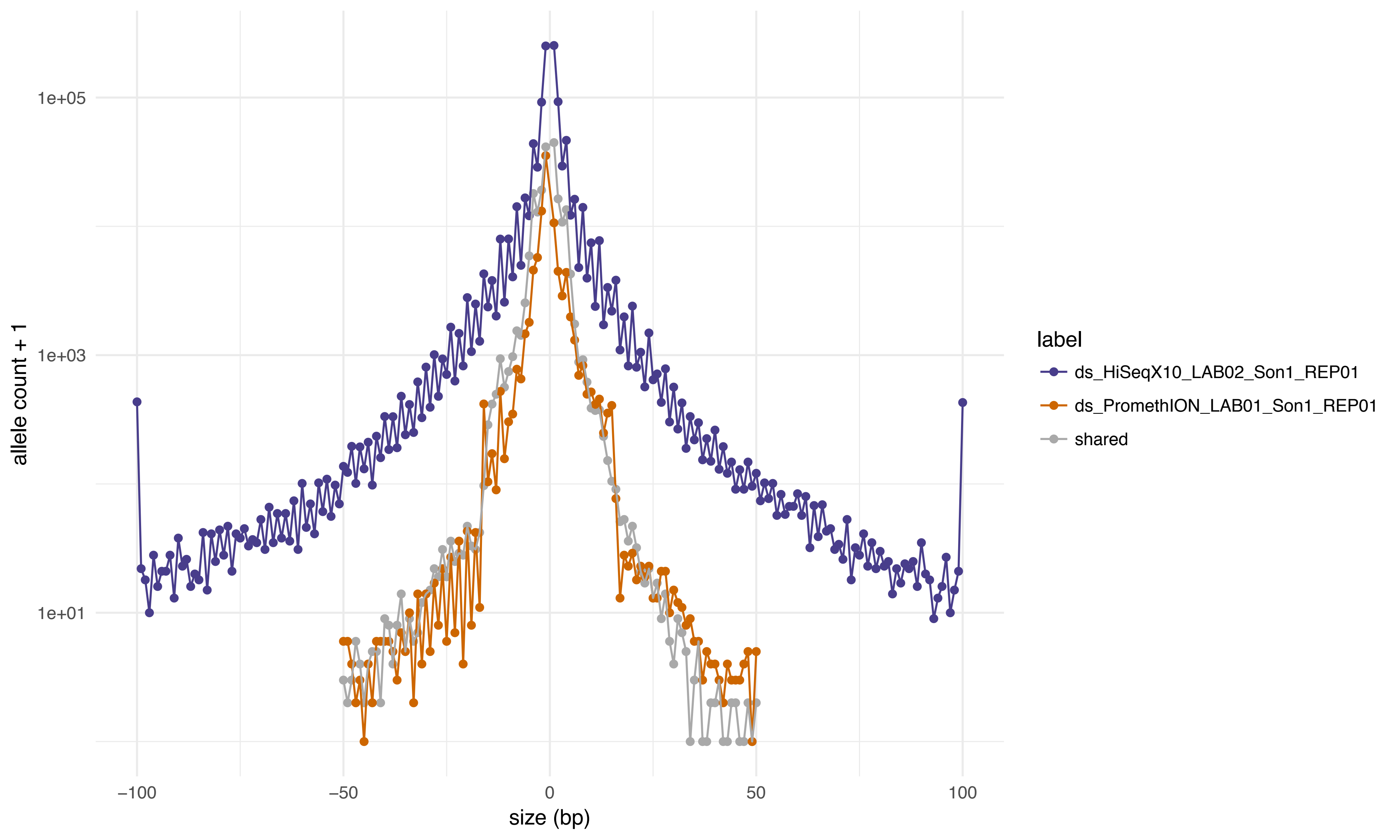

1e+05
1e+03
1e+01
allele count + 1
−100
−50
0
50
100
size (bp)
label
ds_HiSeqX10_LAB02_Son1_REP01
ds_PromethION_LAB01_Son1_REP01
shared

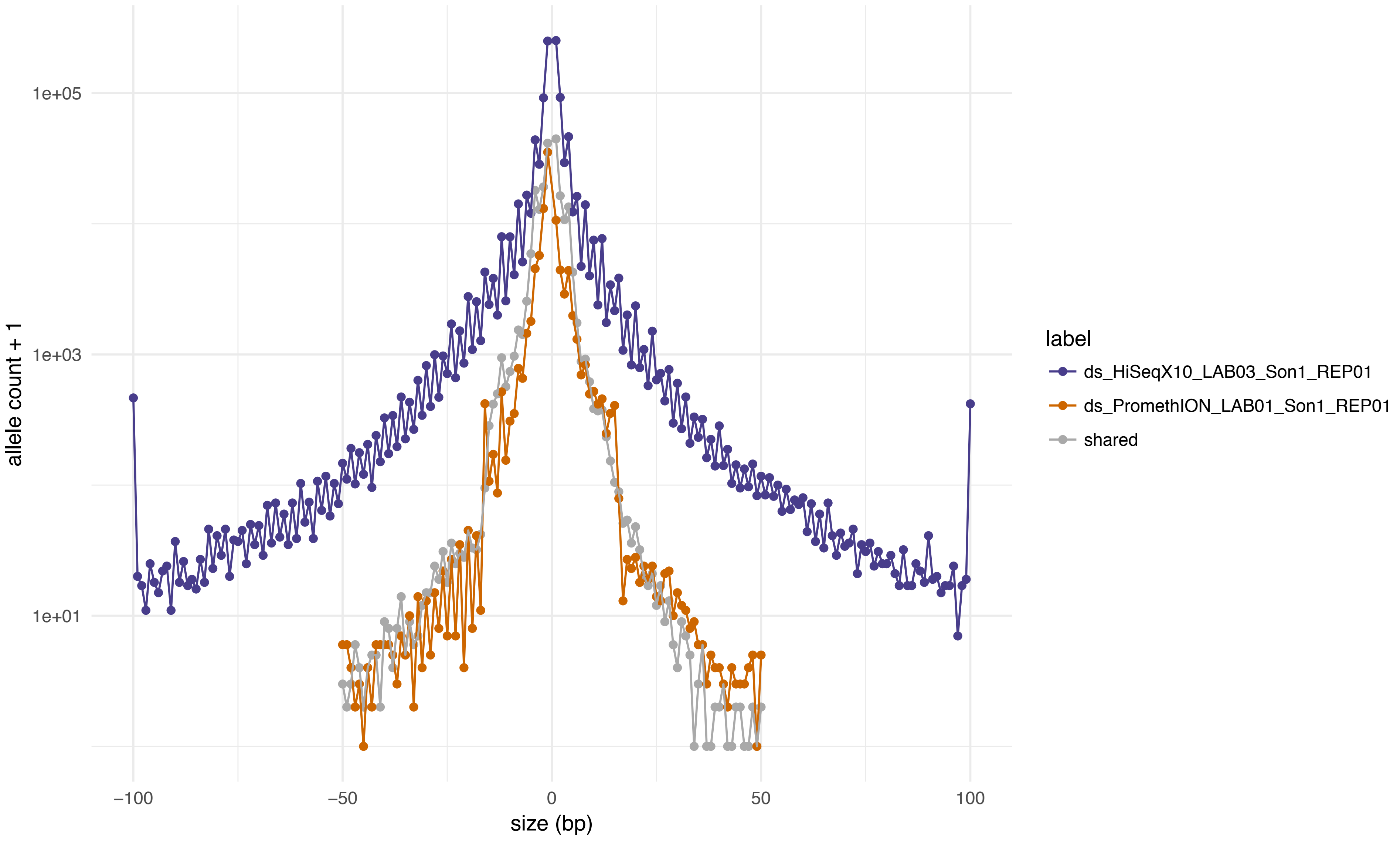

1e+05
1e+03
1e+01
allele count + 1
−100
−50
0
50
100
size (bp)
label
ds_HiSeqX10_LAB03_Son1_REP01
ds_PromethION_LAB01_Son1_REP01
shared

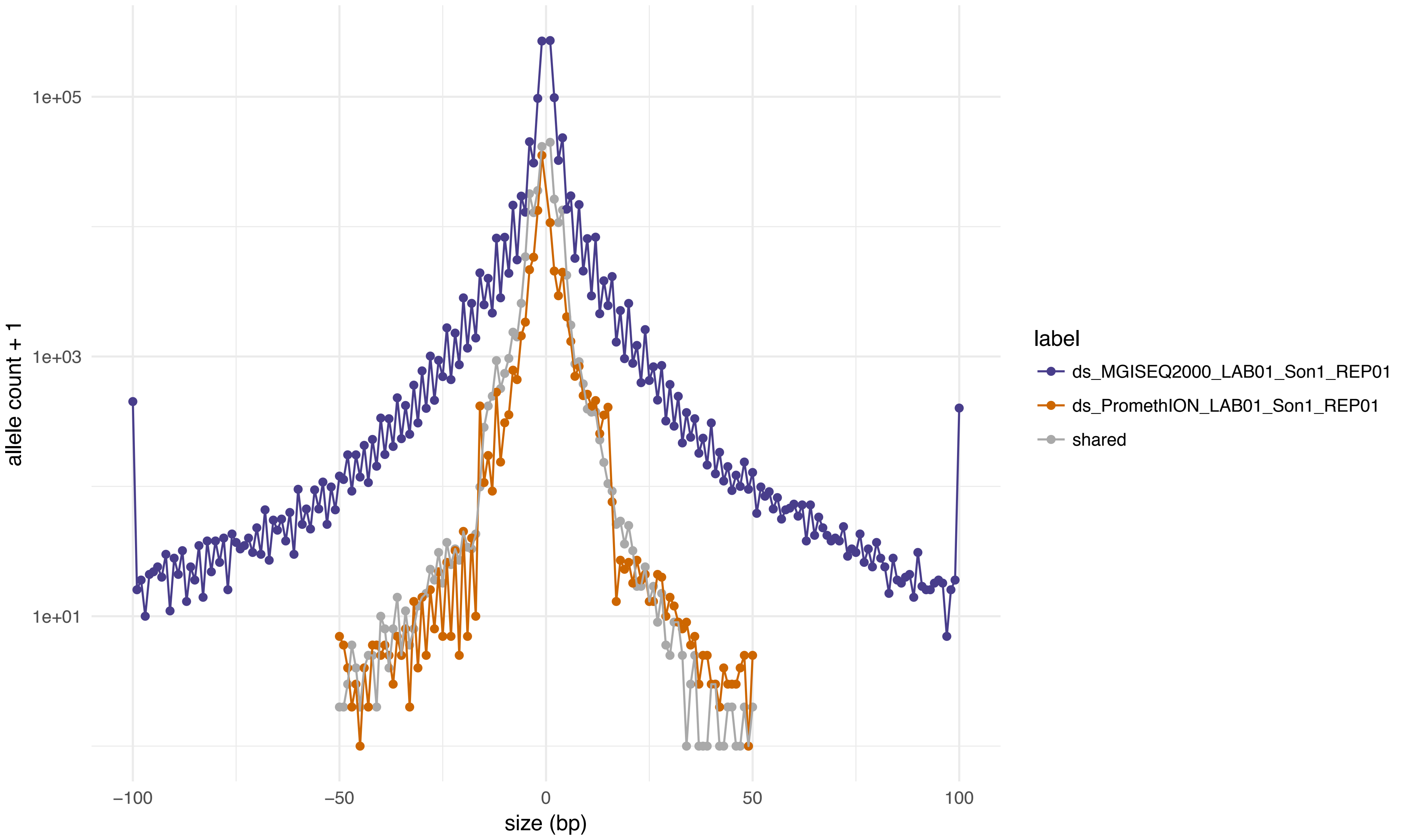
1e+05
1e+03
1e+01
allele count + 1
−100
−50
0
50
100
size (bp)
label
ds_MGISEQ2000_LAB01_Son1_REP01
ds_PromethION_LAB01_Son1_REP01
shared

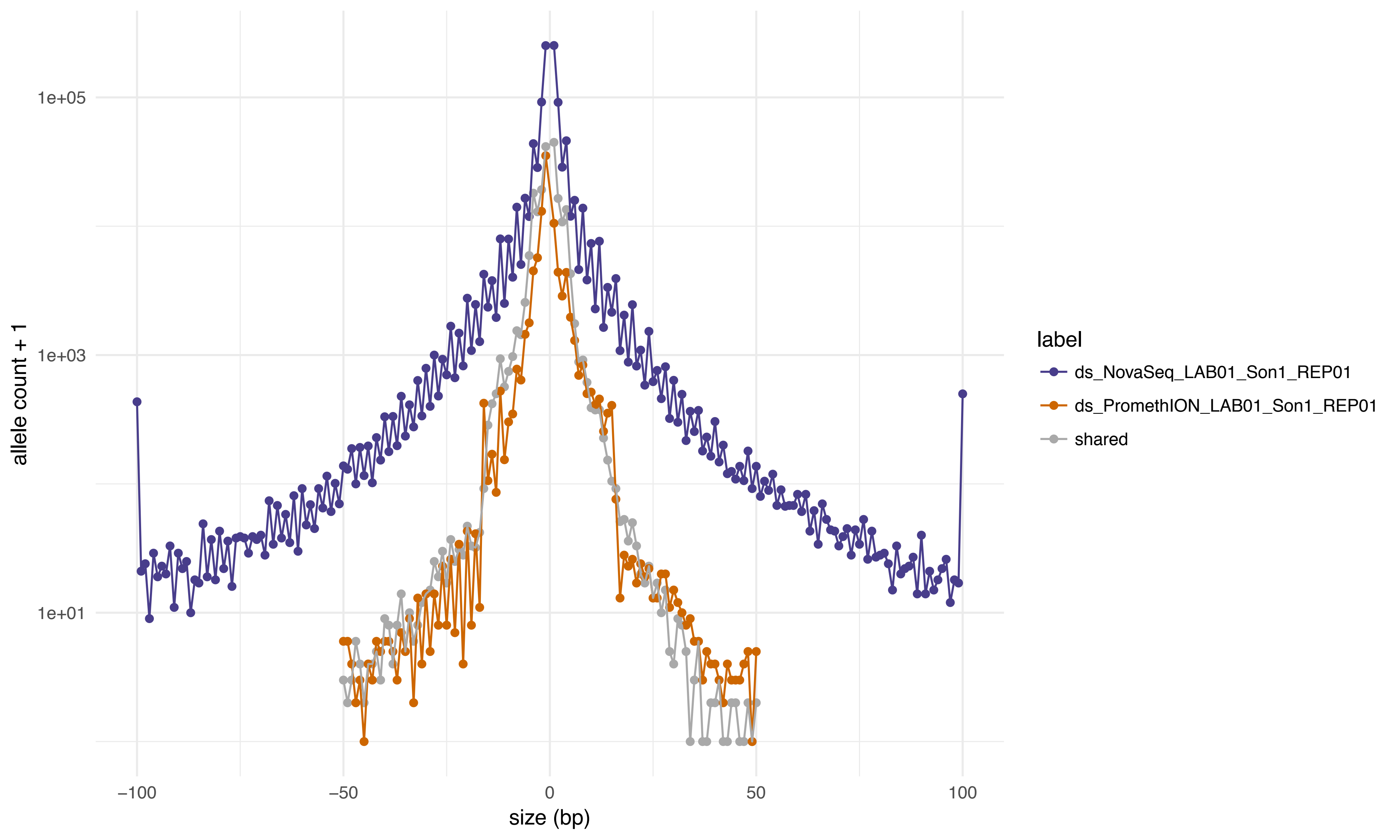

1e+05
1e+03
1e+01
allele count + 1
−100
−50
0
50
100
size (bp)
label
ds_NovaSeq_LAB01_Son1_REP01
ds_PromethION_LAB01_Son1_REP01
shared

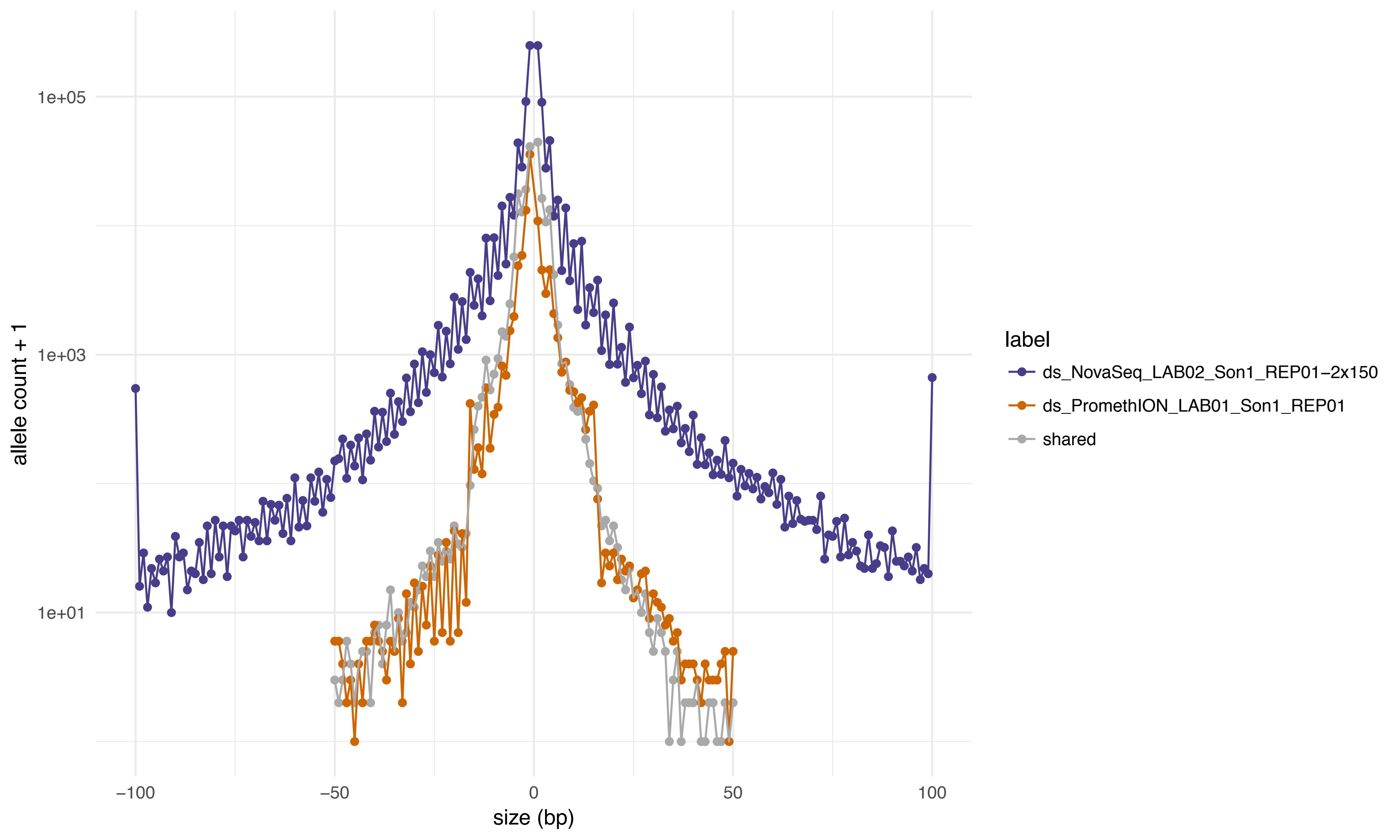

1e+05
1e+03
1e+01
allele count + 1
−100
−50
0
50
100
size (bp)
label
ds_NovaSeq_LAB02_Son1_REP01−2x150
ds_PromethION_LAB01_Son1_REP01
shared

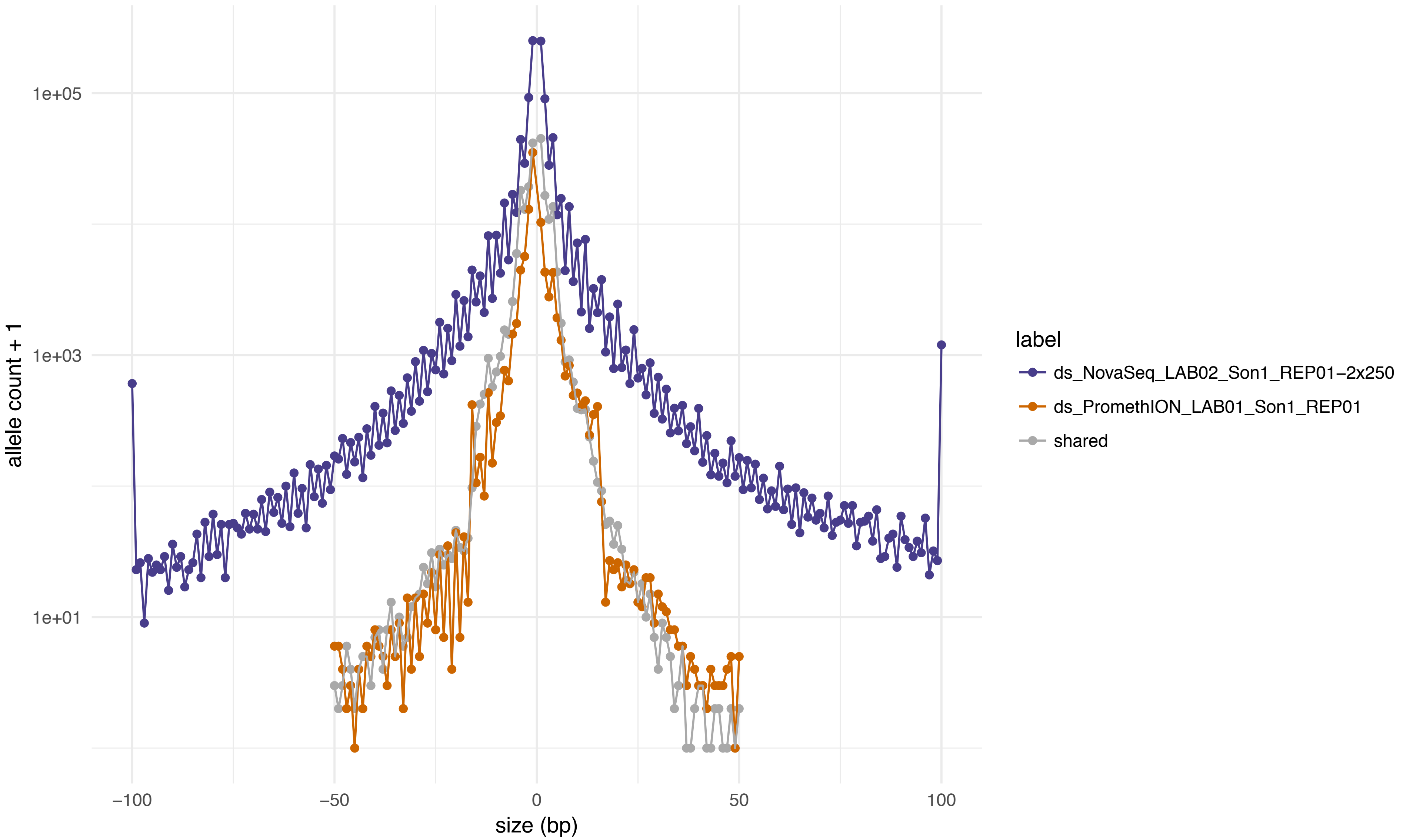

1e+05
1e+03
1e+01
allele count + 1
−100
−50
0
50
100
size (bp)
label
ds_NovaSeq_LAB02_Son1_REP01−2x250
ds_PromethION_LAB01_Son1_REP01
shared
