## Supplementary Data 3 for "Multi-Platform Assessment of DNA Sequencing Performance using Human and Bacterial Reference Genomes in the ABRF Next-Generation Sequencing Study": BacterialErrorRates.Rmd

````
---
title: "ABRF NGS2: BACTERIAL Error Rates by Context"
author: "Jonathan Foox"
date: "May 14, 2020"
output: 
  html_notebook: 
    code_folding: none
    highlight: tango
    toc: yes
    
---


```{r}
library(reshape2)
library(dplyr)
library(tidyr)
library(cowplot)
library(ggplot2)
library(ggridges)
library(ggpubr)
```


```{r}
df_mhist = read.csv("context/bacteria/mhists/bacteria_mhist_ALL_v2.csv")
head(df_mhist)
```

```{r}
df_mhist_melt = melt(df_mhist, id=c("sample","GC","base"))
head(df_mhist_melt)
```


```{r}
df_mhist_melt_basesAveraged = df_mhist_melt %>% 
  group_by(sample, GC, variable) %>% 
  summarize_at(vars(value), mean)

df_mhist_melt_basesAveraged = df_mhist_melt_basesAveraged[complete.cases(df_mhist_melt_basesAveraged), ] # to account for empty GCs

df_mhist_melt_basesAveraged = separate(df_mhist_melt_basesAveraged, sample, 
                                          c("platform", "lab", "genome", "rep"), sep="_", remove=F)

head(df_mhist_melt_basesAveraged)
```

```{r}
comp.edits = c("Sub1"="Substitutions", 
              "Del1"="Deletions",
              "Ins1"="Insertions")
```

```{r}
df_mhist_melt_basesAveraged$platform = factor(df_mhist_melt_basesAveraged$platform, levels=c("MiSeq", "Genapsys", "S5","PGM", "Flongle", "GridION"))
df_mhist_melt_basesAveraged$genome = factor(df_mhist_melt_basesAveraged$genome, levels=c("Ste", "Eco", "Pflu", "Pool"))
```


```{r, fig.width=10, fig.height=6}
errorRate_byGC = ggplot(subset(df_mhist_melt_basesAveraged, variable != "Other1" & variable != "N1"), 
       aes(x=GC, y=value, fill=variable)) +
  #geom_line(show.legend=F) +
  geom_bar(stat="identity", position="fill") +
  xlim(0, 100) +
  theme_bw() +
  theme(text = element_text(size=15),
        legend.position="bottom",
        axis.text.x = element_text(angle=0, size=11, hjust=0.5, vjust=0.5),
        axis.text.y = element_text(angle=0, size=11),
        strip.text.y = element_text(angle=0),
        #panel.grid.major = element_blank(), 
        panel.grid.minor = element_blank()) +
  scale_fill_manual(name = "", labels = c("Match", "Substitution", "Deletion", "Insertion"), 
                       values=c("Match1"="#b2df8a", "Sub1"="#e31a1c", "Del1"="#ff7f00", "Ins1"="#1f78b4")) +
  facet_grid(platform~genome,
             labeller = labeller(variable = comp.edits)) +
  coord_trans(y="sqrt") +
  xlab("GC%") + ylab("Error Rate") +#+ coord_cartesian(ylim = c(0,0.15))#+ ggtitle("Rates of Error Types across GC Compositions") #
  scale_y_continuous(breaks=c(0.01, 0.1, 0.25, 0.50, 1.0))
errorRate_byGC
```


```{r}
pdf("../figures/bacterial_GC_errorRate_v2.pdf", width=10, height=6)
errorRate_byGC
dev.off()
```


```{r}
df_mhist_melt_GCAveraged = df_mhist_melt %>% 
  group_by(sample, base, variable) %>% 
  summarize_at(vars(value), mean)

df_mhist_melt_GCAveraged = df_mhist_melt_GCAveraged[complete.cases(df_mhist_melt_GCAveraged), ] # to account for empty GCs


df_mhist_melt_GCAveraged = separate(df_mhist_melt_GCAveraged, sample, 
                                       c("platform", "lab", "genome", "rep"), sep="_", remove=F)
head(df_mhist_melt_GCAveraged)
```

```{r}
df_mhist_melt_GCAveraged$platform = factor(df_mhist_melt_GCAveraged$platform, levels=c("Flongle", "GridION", "Genapsys","MiSeq", "PGM", "S5"))
df_mhist_melt_GCAveraged$genome = factor(df_mhist_melt_GCAveraged$genome, levels=c("Ste", "Eco", "Pflu", "Pool"))
```


```{r, fig.width=12, fig.height=2}
errorRate_byCycle = ggplot(subset(df_mhist_melt_GCAveraged, variable != "Match1" & variable != "Other1" & variable != "N1"), 
       aes(x=base, y=value, fill=sample, color=variable)) +
  stat_smooth(show.legend=F, span=0.3, size=0.8, alpha=0.6, se = F) +
  #ylim(0, 0.03) +
  theme_bw() +
  theme(text = element_text(size=12),
        axis.text.x = element_text(angle=0, size=8, hjust=0.5, vjust=0.5),
        #panel.grid.major = element_blank(), 
        panel.grid.minor = element_blank()) +
  #theme(text = element_text(size=16)) +
  scale_color_discrete(name = "", labels = c("Substitution", "Deletion", "Insertion")) +
  facet_grid(genome~platform, scales="free",
             labeller = labeller(variable = comp.edits)) +
  xlab("Cycle") + ylab("Error Rate") #+ ggtitle("Rates of Error Types across Base")
errorRate_byCycle
```
````
