## Supplementary Data 3 for "Multi-Platform Assessment of DNA Sequencing Performance using Human and Bacterial Reference Genomes in the ABRF Next-Generation Sequencing Study": BacterialGCheatmaps.Rmd

````
---
title: "ABRF NGS2: Bacterial GC Mapping"
author: "Jonathan Foox"
date: "March 25, 2020"
output: 
  html_notebook: 
    code_folding: none
    highlight: tango
    toc: yes
    
---

```{r}
#library(devtools)
#install_github("jokergoo/ComplexHeatmap")
```


```{r}
library(tidyr)
library(ggplot2)
library(ggridges)
library("scales")
library(dplyr)
library(reshape2)

```

```{r}
df_bac_gc = read.csv("gcMetrics__table__ALL_byReadsUsed.tsv", sep="\t")
head(df_bac_gc)
```

```{r}
df_bac_gc_tmp = dcast(data = df_bac_gc, formula = ...~Sample, value.var="Reads_Used", fun.aggregate=sum)
df_bac_gc_tmp[1:10, 1:5]
```


```{r}
normalized<-function(y) {

  x<-y[!is.na(y)]

  x<-(x - min(x)) / (max(x) - min(x))

  y[!is.na(y)]<-x

  return(y)
  }
```

```{r}
df_bac_gc2 = apply(df_bac_gc_tmp[, colnames(df_bac_gc_tmp)[2:ncol(df_bac_gc_tmp)]],2,normalized)
df_bac_gc2[1:5, 1:5]
```

```{r}
df_bac_gc2 = melt(df_bac_gc2)
colnames(df_bac_gc2) = c("GC", "Sample", "Reads_Used")
df_bac_gc2$GC = df_bac_gc2$GC - 1
df_bac_gc2 = separate(df_bac_gc2, col=Sample, into=c("Platform", "Lab", "Genome", "Rep"), remove=F)
head(df_bac_gc2)
```


```{r}
df_bac_gc_unmelt = dcast(data = df_bac_gc2, formula = ...~GC, value.var="Reads_Used", fun.aggregate=mean)
df_bac_gc_unmelt[1:5, 1:16]
```

```{r}
df_expected = read.csv("bacteria_sentieon/gc_coverage/normalized_gc_coverage_ALL_windows.csv")
head(df_expected)
```


```{r}
library(ComplexHeatmap)
library(circlize)
```

```{r}
df_bac_gc_unmelt_matrix = as.matrix(df_bac_gc_unmelt[,6:ncol(df_bac_gc_unmelt)])
rownames(df_bac_gc_unmelt_matrix) = df_bac_gc_unmelt[,1]
df_bac_gc_unmelt_matrix[1:5, 1:10]
```

```{r}
#df_bac_gc_unmelt_matrix = df_bac_gc_unmelt_matrix[grepl("Eco", rownames(df_bac_gc_unmelt_matrix)), ]
```


```{r}
ha_names = HeatmapAnnotation(Platform = df_bac_gc_unmelt$Platform,
                             col = list(Platform = c("S5"="#ff7f00", "PGM"="#ffff33", "MiSeq"="#377eb8", "Genapsys"="#984ea3",
                                                     "Flongle"="#74c476", "GridION"="#005a32")),
                             #show_legend = T,
                             which="row")
```


```{r}
hm = Heatmap(df_bac_gc_unmelt_matrix,
                 col  = colorRamp2(c(0, max(df_bac_gc_unmelt_matrix)*0.25, 
                                     max(df_bac_gc_unmelt_matrix)*0.5, 
                                     max(df_bac_gc_unmelt_matrix)*0.75, 
                                     max(df_bac_gc_unmelt_matrix)), c( "#2c7bb6", "#abd9e9","#ffffbf","#fdae61","#d7191c")),
                 #col = colorRamp2(c(0,0.5,1,1.5,2), c( "#2c7bb6", "#abd9e9","#ffffbf","#fdae61","#d7191c")),
                 #rect_gp = gpar(col = "white", lwd = 0.25),
                 #column_title = "Metagenomic Pool",
                 cluster_columns = FALSE,
                 show_column_names = TRUE,
             column_title = "Bacterial Normalized Coverage across GC",
             cluster_rows = F,
             row_split = factor(df_bac_gc_unmelt$Genome, levels=c("Ste", "Eco", "Pflu", "Pool")),
                 column_title_gp = gpar(fontsize = 16),
                 column_names_side = "top",
                 column_labels = c("0",".",".",".",".",".",".",".",".",".",".","10",".",".",".",".",".",".",".",".",".","20",".",".",".",".",".",".",".",".",".","30",".",".",".",".",".",".",".",".",".","40",".",".",".",".",".",".",".",".",".","50",".",".",".",".",".",".",".",".",".","60",".",".",".",".",".",".",".",".",".","70",".",".",".",".",".",".",".",".",".","80",".",".",".",".",".",".",".",".",".","90",".",".",".",".",".",".",".",".","100"),
            
                 column_names_rot = 45,
                 column_names_gp = gpar(fontsize=12),
                 
                 show_row_names = TRUE,
                 #bottom_annotation = ha_dens,
                 show_heatmap_legend = T,
                 heatmap_legend_param = list(
                   title = "Coverage"
                 ))
```

```{r, fig.width=7, fig.height=6}
pdf("../figures/bacterial_GC_distribution.pdf", width=8, height=6.5)
draw(hm + ha_names, merge_legend = TRUE, heatmap_legend_side="right")
dev.off()
```
````
