## Supplementary Data 3 for "Multi-Platform Assessment of DNA Sequencing Performance using Human and Bacterial Reference Genomes in the ABRF Next-Generation Sequencing Study": BaterialMetagenomicMixtureMapping.Rmd

````
---
title: "ABRF NGS2: Bacterial Pool Taxon Mapping"
author: "Jonathan Foox"
date: "May 15, 2020"
output: 
  html_notebook: 
    code_folding: none
    highlight: tango
    toc: no
    
---


```{r}
setwd("/athena/masonlab/scratch/projects/abrf_ngs_phaseii/analysis")
```

```{r}
library(reshape2)
library(dplyr)
library(tidyr)
library(cowplot)
library(ggplot2)
library(ggridges)
library(ggpubr)
```


```{r}
df_bacmap = read.csv("bacterialPoolMapping_fromMosDepth.csv")
df_bacmap = separate(df_bacmap, col="sample", sep="_", into=c("platform","lab","pool","rep"), remove=F)

df_bacmap$platform = factor(df_bacmap$platform, levels=c("MiSeq","Genapsys","S5","PGM","Flongle","GridION"))
df_bacmap$taxon = factor(df_bacmap$taxon, levels=c("S_epidermidis","E_faecalis", "P_haloplanktis", "H_halophilus", "B_subtilis", "E_coli", "P_fluorescens",
                                                   "C_violaceum", "H_volcanii", "M_luteus"))

df_bacmap$frac = round(df_bacmap$pct / 100, 2)

head(df_bacmap)
```

```{r}
library(circlize)
```


```{r, fig.width=10, fig.height=6}
bac_frac = ggplot(df_bacmap, aes(x=sample, y=pct, fill=taxon)) +
  geom_bar(stat="identity", position = position_stack(reverse=TRUE)) +
  facet_grid(platform~., scales="free", space = "free") +
  theme_bw() +
    theme(text = element_text(size = 14),
          legend.position = "bottom",
        axis.text.y = element_blank(),
        axis.ticks.y = element_blank(),
        strip.text.y = element_text(angle=0)) +
  scale_fill_brewer(palette="Spectral") +
  xlab("") + ylab("Percentage") +
  coord_flip()
  #geom_hline(yintercept = seq(0, 1, 0.1), linetype="dashed", alpha=0.3) +
  #geom_text(size=3, position=position_stack(reverse=T, vjust=0.5), aes(label=frac))

bac_frac

  #scale_fill_manual(values=colorRamp2(c(0, 0.015, 0.025, 0.035, 0.05), c( "#2c7bb6", "#abd9e9","#ffffbf","#fdae61","#d7191c")))
```


```{r}
df_bacmap2 = df_bacmap[, c(2,6,8)] %>% group_by(platform, taxon) %>% summarise_all(funs(mean))
head(df_bacmap2)
```

```{r}
ggplot(df_bacmap2, aes(x=taxon, y=pct, fill=taxon)) +
  geom_bar(stat="identity") +
  facet_grid(.~platform) +
  theme_bw() +
    theme(text = element_text(size = 14),
        legend.position = "bottom",
        axis.text.x = element_blank(),
        axis.ticks.x = element_blank()) +
  geom_hline(yintercept = 10, linetype="dashed") +
  scale_fill_brewer(palette="Spectral")
```

```{r, fig.width=9, fig.height=4}
bac_distrib = ggplot(df_bacmap, aes(x=taxon, y=pct, fill=taxon)) +
  geom_boxplot() +
  geom_point() +
  theme_bw() +
    theme(text = element_text(size = 14),
        legend.position = "none",
        axis.text.x = element_text(angle=0, size=12, hjust=0.5, vjust=0.5),
        axis.ticks.x = element_blank()) +
  scale_x_discrete(labels=c("S_epidermidis"="S. epid\n28.0%\n(+)","E_faecalis"="E. faec\n37.2\n(+)%", "P_haloplanktis"="P. halo\n40.1%\n(-)", "H_halophilus"="H. halo\n41.6%\n(+)", "B_subtilis"="B. subt\n43.5%\n(+)", "E_coli"="E. coli\n50.8%\n(-)", "P_fluorescens"="P. fluo\n61.4%\n(-)","C_violaceum"="C. viol\n64.8%\n(-)", "H_volcanii"="H. volc\n65.5%\n(+)", "M_luteus"="M. lute\n72.0%\n(+)")) +
  geom_hline(yintercept = 10, linetype="dashed", alpha=0.5) +
  xlab("") + ylab("Percentage") +
  scale_fill_brewer(palette="Spectral")
bac_distrib
```


```{r, fig.width=8, fig.height=8}
pdf(file="../figures/bacterial_pool_mapping_distributions.pdf", width=8, height=8)
plot_grid(bac_frac, bac_distrib, ncol=1, align='v', axis='l', rel_heights = c(1, 0.5))
dev.off()
```
````
