## Supplementary Data 3 for "Multi-Platform Assessment of DNA Sequencing Performance using Human and Bacterial Reference Genomes in the ABRF Next-Generation Sequencing Study": HumanErrorRates.Rmd

````
---
title: "ABRF NGS2: Error Rates by Context"
author: "Jonathan Foox"
date: "May 13, 2020"
output: 
  html_notebook: 
    code_folding: none
    highlight: tango
    toc: yes
    
---


```{r}
library(reshape2)
library(dplyr)
library(tidyr)
library(cowplot)
library(ggplot2)
library(ggridges)
library(ggpubr)
```

# Stratified Error Rates

## bbmap-based

#### run_errorRates_per_GCwindow.slurm
```{bash}
#! /bin/bash -l
#SBATCH --job-name=NGS2__errorRates_per_GCwindow
#SBATCH --output=NGS2__errorRates_per_GCwindow_%A_%a.out
#SBATCH --cpus-per-task=1
#SBATCH --time=168:00:00   # HH/MM/SS
#SBATCH --mem=20G

source ~/.bashrc
echo 'Started at:' `date`

# copy data
rsync -av /athena/masonlab/scratch/databases/hs37d5/hs37d5.fa $TMPDIR
bam_path=$(cat sen_proj1_BAMs.txt | sed -n "${SLURM_ARRAY_TASK_ID}p")
rsync -av $bam_path $TMPDIR
cd $TMPDIR

# get variables
GCmax=$1
GCmin=$(bc <<< "$GCmax-0.01")
inbam=$(basename $bam_path)
outbam=$(echo $inbam | sed -e "s/.bam/_GC-$GCmax.bam/")
sample=$(echo $inbam | sed -e "s/sorted_dedup_realigned_//" | sed -e "s/.bam//")

# get pct GC for primary alignments
reformat.sh -in=$inbam -out=$outbam mingc=$GCmin maxgc=$GCmax mappedonly=t primaryonly=t -ref=hs37d5.fa -Xmx16g

# create histograms for that GC%
reformat.sh -in=$outbam ehist=ehist_${sample}_${GCmax}.txt mhist=mhist_${sample}_${GCmax}.txt indelhist=indelhist_${sample}_${GCmax}.txt -ref=hs37d5.fa -Xmx16g

# copy output
rsync -av *.txt /athena/masonlab/scratch/projects/abrf_ngs_phaseii/analysis/context/bbmap_hists

echo 'Finished at: ' `date`
```

#### submit on cluster
```{bash}
nums=$(for i in {01..99}; do printf 0.$i" "; done)
for num in $nums; do sbatch --array 1-76 run_errorRates_per_GCwindow.slurm $num; done
```

#### gather outputs
```{bash}
# outputs are in: /athena/masonlab/scratch/projects/abrf_ngs_phaseii/analysis/context/bbmap_hists/mhists
for i in `ls context/bbmap_hists/mhists/mhist_* | grep -v 'ALL'`; do \
  sample=$(echo $i | cut -d_ -f2-5); GC=$(echo $i | cut -d. -f2); \
  tail -n +2 $i | sed -e "s/^/${sample}\t${GC}\t/g"; done > context/bbmap_hists/mhists/mhist_ALL.txt
```


```{r}
df_mhist = read.csv("/athena/masonlab/scratch/projects/abrf_ngs_phaseii/analysis/context/ga4gh/bbmap_hists/mhists/mhist_ALL_v5.csv")
df_mhist_R1 = df_mhist#[, 1:9]
head(df_mhist_R1)
```


```{r}
df_mhist_R1_melt = melt(df_mhist_R1, id=c("sample","GC","base"))
head(df_mhist_R1_melt)
```


```{r}
df_mhist_R1_melt_basesAveraged = df_mhist_R1_melt %>% 
  group_by(sample, GC, variable) %>% 
  summarize_at(vars(value), mean)

df_mhist_R1_melt_basesAveraged = separate(df_mhist_R1_melt_basesAveraged, sample, 
                                          c("platform", "lab", "genome", "rep"), sep="_", remove=F)

head(df_mhist_R1_melt_basesAveraged)
```


```{r}
df_mhist_R1_melt_basesAveraged[grepl("CLR", df_mhist_R1_melt_basesAveraged$sample), ]$platform = "PacBio-CLR"
df_mhist_R1_melt_basesAveraged[grepl("CCS", df_mhist_R1_melt_basesAveraged$sample), ]$platform = "PacBio-CCS"
#df_mhist_R1_melt_basesAveraged[grepl("Nanopore", df_mhist_R1_melt_basesAveraged$platform), ]$platform = "PromethION"
df_mhist_R1_melt_basesAveraged[grepl("NovaSeq", df_mhist_R1_melt_basesAveraged$sample), ]$platform = "NovaSeq\n2x150"
df_mhist_R1_melt_basesAveraged[grepl("2x150", df_mhist_R1_melt_basesAveraged$sample), ]$platform = "NovaSeq\n2x150"
df_mhist_R1_melt_basesAveraged[grepl("2x250", df_mhist_R1_melt_basesAveraged$sample), ]$platform = "NovaSeq\n2x250"
```


```{r}
comp.edits = c("Sub1"="Substitutions", 
              "Del1"="Deletions",
              "Ins1"="Insertions")
```

```{r}
df_mhist_R1_melt_basesAveraged$platform = factor(df_mhist_R1_melt_basesAveraged$platform, levels=c("HiSeq2500", "HiSeq4000", "HiSeqX10", "NextSeq", "NovaSeq\n2x150", "NovaSeq\n2x250", "BGISEQ500", "MGISEQ2000", "Genapsys", "Flongle", "GridION", "PromethION", "PacBio-CLR", "PacBio-CCS"))
```


```{r, fig.width=12, fig.height=2}
errorRate_byGC = ggplot(subset(df_mhist_R1_melt_basesAveraged, variable != "Match1" & variable != "Other1" & variable != "N1"), 
       aes(x=GC, y=value, fill=sample, color=variable)) +
  #geom_rect(xmin=0, xmax=25, ymin=0, ymax=0.04, fill="#ffffe6", color=NA, alpha=0.01, show.legend=F) +
  #geom_rect(xmin=75, xmax=100, ymin=0, ymax=0.04, fill="#ffffe6", color=NA, alpha=0.01, show.legend=F) +
  #geom_point(size=0.7) +
  #stat_smooth(show.legend=F, span=0.1, size=0.8, alpha=0.7, se=F) +
  geom_line(show.legend=F) +
  xlim(0, 100) +
  theme_bw() +
  theme(text = element_text(size=12),
        axis.text.x = element_text(angle=0, size=8, hjust=0.5, vjust=0.5),
        #panel.grid.major = element_blank(), 
        panel.grid.minor = element_blank()) +
  scale_color_discrete(name = "", labels = c("Substitution", "Deletion", "Insertion")) +
  facet_grid(.~platform,
             labeller = labeller(variable = comp.edits)) +
  xlab("GC%") + ylab("Error Rate") #+ ggtitle("Rates of Error Types across GC Compositions") #+ coord_trans(y="sqrt") +scale_y_continuous(breaks=c(0.01, 0.02, 0.04, 0.06))
errorRate_byGC
```


```{r, fig.width=14, fig.height=3}
errorRate_byGC_stacked = ggplot(subset(df_mhist_R1_melt_basesAveraged, variable != "Other1" & variable != "N1"), 
       aes(x=GC, y=value, fill=variable)) +
  #geom_line(show.legend=F) +
  geom_bar(stat="identity", position="fill", show.legend=F) +
  xlim(0, 100) +
  theme_bw() +
  theme(text = element_text(size=15),
        #legend.position="bottom",
        axis.text.x = element_text(angle=90, size=10, hjust=1, vjust=0.5),
        axis.text.y = element_text(angle=0, size=10),
        strip.text.x = element_text(angle=0, size=10),
        #panel.grid.major = element_blank(), 
        panel.grid.minor = element_blank()) +
  scale_fill_manual(name = "", labels = c("Match", "Substitution", "Deletion", "Insertion"), 
                       values=c("Match1"="#b2df8a", "Sub1"="#e31a1c", "Del1"="#ff7f00", "Ins1"="#1f78b4")) +
  facet_grid(.~platform,
             labeller = labeller(variable = comp.edits)) +
  coord_trans(y="sqrt") +
  xlab("GC%") + ylab("Error Fraction") +#+ coord_cartesian(ylim = c(0,0.15))#+ ggtitle("Rates of Error Types across GC Compositions") #
  scale_y_continuous(breaks=c(0.01, 0.05, 0.13, 0.25, 0.50, 1.0))
errorRate_byGC_stacked
```


```{r}
pdf(file="../figures/errorRate_byGC_v2.pdf", width=12, height=2)
errorRate_byGC
dev.off()
```


```{r}
df_mhist_R1_melt_GCAveraged = df_mhist_R1_melt %>% 
  group_by(sample, base, variable) %>% 
  summarize_at(vars(value), mean)
df_mhist_R1_melt_GCAveraged = separate(df_mhist_R1_melt_GCAveraged, sample, 
                                       c("platform", "lab", "genome", "rep"), sep="_", remove=F)
head(df_mhist_R1_melt_GCAveraged)
```

```{r}
df_mhist_R1_melt_GCAveraged[grepl("CLR", df_mhist_R1_melt_GCAveraged$sample), ]$platform = "PacBio-CLR"
df_mhist_R1_melt_GCAveraged[grepl("CCS", df_mhist_R1_melt_GCAveraged$sample), ]$platform = "PacBio-CCS"
#df_mhist_R1_melt_GCAveraged[grepl("Nanopore", df_mhist_R1_melt_GCAveraged$platform), ]$platform = "PromethION"
df_mhist_R1_melt_GCAveraged[grepl("NovaSeq", df_mhist_R1_melt_GCAveraged$sample), ]$platform = "NovaSeq\n2x150"
df_mhist_R1_melt_GCAveraged[grepl("2x150", df_mhist_R1_melt_GCAveraged$sample), ]$platform = "NovaSeq\n2x150"
df_mhist_R1_melt_GCAveraged[grepl("2x250", df_mhist_R1_melt_GCAveraged$sample), ]$platform = "NovaSeq\n2x250"
```


```{r}
df_mhist_R1_melt_GCAveraged$platform = factor(df_mhist_R1_melt_GCAveraged$platform, levels=c("HiSeq2500", "HiSeq4000", "HiSeqX10", "NextSeq", "NovaSeq\n2x150", "NovaSeq\n2x250", "BGISEQ500", "MGISEQ2000", "Genapsys", "Flongle", "GridION", "PromethION", "PacBio-CLR", "PacBio-CCS"))
```


```{r, fig.width=12, fig.height=2}
errorRate_byCycle = ggplot(subset(df_mhist_R1_melt_GCAveraged, variable != "Match1" & variable != "Other1" & variable != "N1"), 
       aes(x=base, y=value, fill=sample, color=variable)) +
  stat_smooth(show.legend=F, span=0.3, size=0.8, alpha=0.6, se = F) +
  #ylim(0, 0.03) +
  theme_bw() +
  theme(text = element_text(size=12),
        axis.text.x = element_text(angle=0, size=8, hjust=0.5, vjust=0.5),
        #panel.grid.major = element_blank(), 
        panel.grid.minor = element_blank()) +
  #theme(text = element_text(size=16)) +
  scale_color_discrete(name = "", labels = c("Substitution", "Deletion", "Insertion")) +
  facet_grid(.~platform, scales="free",
             labeller = labeller(variable = comp.edits)) +
  xlab("Cycle") + ylab("Error Rate") #+ ggtitle("Rates of Error Types across Base")
errorRate_byCycle
```


```{r, fig.width=14, fig.height=2.5}
errorRate_byCycle_stacked = ggplot(subset(df_mhist_R1_melt_GCAveraged, variable != "Other1" & variable != "N1"), 
       aes(x=base, y=value, fill=variable)) +
  geom_bar(stat="identity", position="fill") +
  theme_bw() +
  theme(text = element_text(size=15),
        legend.position="bottom",
        axis.text.x = element_text(angle=90, size=10, hjust=1, vjust=0.5),
        axis.text.y = element_text(angle=0, size=10),
        strip.background = element_blank(),
        strip.text.x = element_blank(),
        #strip.text.x = element_text(angle=90, size=10),
        #panel.grid.major = element_blank(), 
        panel.grid.minor = element_blank()) +
  scale_fill_manual(name = "", labels = c("Match", "Substitution", "Deletion", "Insertion"), 
                       values=c("Match1"="#b2df8a", "Sub1"="#e31a1c", "Del1"="#ff7f00", "Ins1"="#1f78b4")) +
  facet_grid(.~platform,
             labeller = labeller(variable = comp.edits), scales="free_x") +
  coord_trans(y="sqrt") +
  xlab("Base Number") + ylab("Error Fraction") +#+ coord_cartesian(ylim = c(0,0.15))#+ ggtitle("Rates of Error Types across GC Compositions") #
  scale_y_continuous(breaks=c(0.01, 0.05, 0.13, 0.25, 0.50, 1.0))
errorRate_byCycle_stacked
```


```{r}
pdf(file="../figures/errorRate_byCycle_v2.pdf", width=12, height=2)
errorRate_byCycle
dev.off()
```


```{r}
df_error1 = read.csv("context/ucsc/ALL_errorRate_UCSC_withLongReads_v2.csv")

colorsList = list()
colorsList[["HiSeq2500"]] = "#9ecae1"
colorsList[["HiSeq4000"]] = "#6baed6"
colorsList[["HiSeqX10"]] = "#3182bd"
colorsList[["NovaSeq2x150"]] = "#ae017e"
colorsList[["NovaSeq2x250"]] = "#ae017e"
colorsList[["NextSeq"]] = "#d0d0ef"
colorsList[["BGISEQ500"]] = "#fc8d59"
colorsList[["MGISEQ2000"]] = "#e34a33"
colorsList[["Genapsys"]] = "#ffd92f"
colorsList[["Flongle"]] = "#a1d99b"
colorsList[["GridION"]] = "#41ab5d"
colorsList[["PromethION"]] = "#006d2c"
colorsList[["PacBioCLR"]] = "#ef3b2c"
colorsList[["PacBioCCS"]] = "#a50f15"
colorsList2 = as.data.frame(unlist(colorsList))
colnames(colorsList2)[1] = "plat_colors"

df_error = merge(df_error1, colorsList2, by.x="platform", by.y="row.names", sort=F)

df_error$context = factor(df_error$context, levels=c("Alu", "L1", "L2", "Low", "LTR", "Satellite", "Simple", "telo"))


df_error$platform = factor(df_error$platform, levels=c("HiSeq2500", "HiSeq4000", "HiSeqX10", "NovaSeq2x150", "NovaSeq2x250", "NextSeq", 
                                                       "BGISEQ500", "MGISEQ2000", "Genapsys", "Flongle", "GridION", "PromethION", "PacBioCLR", "PacBioCCS"))

head(df_error)
```


```{r}
context.names = c("Alu"="Alu", "L1"="L1", "L2"="L2", "Low"="Low Complexity", "LTR"="LTR",
               "Satellite"="Satellite", "Simple"="Simple Repeats", "telo"="Telomeres")
```


```{r, fig.width=12, fig.height=4}
error_barplot = ggplot(df_error, 
       aes(x=platform, y=error, fill=platform, group=allinfo)) +
  geom_bar(stat="identity", position="dodge") +
  #geom_point(size=0.2) +
  #geom_boxplot() +
  theme_bw() +
  theme(text = element_text(size = 14),
        legend.position = "bottom",
        axis.text.x = element_blank(),
        axis.ticks.x = element_blank(),
        strip.text.y = element_blank()) +
  scale_fill_manual(values=colorsList) +
  guides(fill=guide_legend(ncol=6)) +
  #scale_fill_manual(values=c("HiSeq2500"="#9ecae1", "HiSeq4000"="#6baed6", "HiSeqX10"="#3182bd", 
   #                          "NovaSeq"="#08519c", "BGISEQ500"="#fc8d59", "MGISEQ2000"="#e34a33", "NextSeq"="#756bb1")) +
  facet_grid(.~context, space="free", scales = "free",
             labeller = labeller(context = context.names)) +
  xlab("") + ylab("Relative Error Rate") +
  #scale_y_sqrt() +
  scale_y_continuous(breaks=c(0.01, 0.05, 0.1, 0.2)) +
  coord_trans(y="sqrt") 
  #geom_hline(yintercept = 0, linetype="dashed", alpha=0.5) #+ ylim(-0.02,0.02) 
error_barplot
```

```{r, fig.width=12, fig.height=8}
pdf(file="../figures/errorBy_GC_cycle_UCSCcontext_v2.pdf", width=13, height=8)
plot_grid(errorRate_byGC_stacked, errorRate_byCycle_stacked, error_barplot, ncol=1, align='v', axis='l', rel_heights = c(0.7, 0.7, 1))
dev.off()
```
````
