## Supplementary Data 3 for "Multi-Platform Assessment of DNA Sequencing Performance using Human and Bacterial Reference Genomes in the ABRF Next-Generation Sequencing Study": HumanErrorRates_UCSCfilter.Rmd

````
---
title: "ABRF NGS2: Error Rates by Context (UCSC)"
author: "Jonathan Foox"
date: "February 3, 2020"
output: 
  html_notebook: 
    code_folding: none
    highlight: tango
    toc: yes
    
---


```{r}
library(reshape2)
library(dplyr)
library(tidyr)
library(cowplot)
library(ggplot2)
library(ggridges)
library(ggpubr)
```


```{r}
setwd("/athena/masonlab/scratch/projects/abrf_ngs_phaseii/analysis")
```


```{r}
df_homo = read.csv("context/ucsc/ALL.homopolymers_v3.out")
head(df_homo)
```

```{r}
df_homo_m = melt(df_homo, id.vars = colnames(df_homo)[1:17], measure.vars = colnames(df_homo)[18:ncol(df_homo)], variable.name = "allinfo", value.name = "error" )
df_homo_m = separate(df_homo_m, col = "allinfo", into=c("platform", "lab", "genome", "rep"), sep = "_", remove=F)
df_homo_m[grepl("CCS",df_homo_m$allinfo), ]$platform = "PacBioCCS"
df_homo_m[grepl("CLR",df_homo_m$allinfo), ]$platform = "PacBioCLR"
#df_homo_m[grepl("Nanopore",df_homo_m$allinfo), ]$platform = "PromethION"
table(df_homo_m$platform)
```

```{r}
df_homo_hm = dcast(data = subset(df_homo_m[, c(7,18,seq(19,ncol(df_homo_m),1))], 
                                 copyNum < 100), 
                   formula=...~copyNum, value.var="error", fun.aggregate=mean)
head(df_homo_hm)
```


```{r}
library(ComplexHeatmap)
library(circlize)
```


```{r}
df_homo_hm_matrix = as.matrix(df_homo_hm[,6:ncol(df_homo_hm)])
rownames(df_homo_hm_matrix) = df_homo_hm[,1]
df_homo_hm_matrix = df_homo_hm_matrix[!grepl("PromethION_LAB01_Son1_REP01|Flongle|Genapsys", rownames(df_homo_hm_matrix)), ]
df_homo_hm_matrix[1:5, 1:5]
```


```{r}
ha_names_homo = HeatmapAnnotation(. = subset(df_homo_hm, allinfo != "PromethION_LAB01_Son1_REP01" & 
                                         allinfo != "Genapsys_LAB01_Son1_REP01" & 
                                         allinfo != "Flongle_LAB01_Son1_REP01.NativeBC")$platform,
                             col = list(. = c("BGISEQ500"="#fd8d3c", "HiSeq2500"="#bdd7e7", "HiSeq4000"="#6baed6", "NextSeq"="#d0d0ef",
                                                     "HiSeqX10"="#3182bd", "MGISEQ2000"="#e6550d", "NovaSeq"="#756bb1", "PromethION"="#006d2c",
                                                      "PacBioCLR" = "#ef3b2c", "PacBioCCS" = "#a50f15","GridION" = "#41ab5d")),
                             #"Genapsys"="#ffd92f","Flongle" = "#a1d99b", 
                             which="row",
                             show_legend = F)
```

```{r}
ha_labels_homo = columnAnnotation(foo = anno_mark(at = seq(1, 100, 5), 
                                             labels=colnames(df_homo_hm_matrix)[seq(1, 100, 5)],
                                             side="bottom", labels_gp = gpar(fontsize=8)))
```


```{r}
hm_homo = Heatmap(df_homo_hm_matrix,
             rect_gp = gpar(col = "white", lwd = 0.25),
             col = colorRamp2(c(0, 0.015, 0.025, 0.035, 0.05), c( "#2c7bb6", "#abd9e9","#ffffbf","#fdae61","#d7191c")),
             cluster_columns = FALSE,
             row_split = factor(subset(df_homo_hm, allinfo != "PromethION_LAB01_Son1_REP01" & 
                                         allinfo != "Genapsys_LAB01_Son1_REP01" & 
                                         allinfo != "Flongle_LAB01_Son1_REP01.NativeBC")$platform, 
                                levels=c("HiSeq2500", "HiSeq4000", "HiSeqX10", "NovaSeq", "NextSeq", 
                                         "BGISEQ500", "MGISEQ2000",  "PacBioCLR", "PacBioCCS", "GridION", "PromethION")),
             row_title_rot=0,
             row_title_gp = gpar(fontsize = 11),
             column_names_gp = gpar(fontsize=8),
             cluster_rows = F,
             show_column_names = F,
             bottom_annotation = ha_labels_homo,
             column_title = "Homopolymer Copy Number (Entropy=0)",
             column_title_side = "bottom",
             column_title_gp = gpar(fontsize=12),
             show_row_names = F,
             show_heatmap_legend = F,
             width = unit(14, "cm"))
```

```{r, fig.width=7, fig.height=7}
draw(hm_homo + ha_names_homo)
```


```{r}
df_STRs = read.csv("context/ucsc/ALL.STRs_v3.out")
head(df_STRs)
```

```{r}
df_STRs_m = melt(df_STRs, id.vars = colnames(df_STRs)[1:17], measure.vars = colnames(df_STRs)[18:ncol(df_STRs)], variable.name = "allinfo", value.name = "error" )
df_STRs_m = separate(df_STRs_m, col = "allinfo", into=c("platform", "lab", "genome", "rep"), sep = "_", remove=F)
df_STRs_m[grepl("CCS",df_STRs_m$allinfo), ]$platform = "PacBioCCS"
df_STRs_m[grepl("CLR",df_STRs_m$allinfo), ]$platform = "PacBioCLR"
head(df_STRs_m)
```

```{r}
df_STRs_hm = dcast(data = subset(df_STRs_m[, c(16,18,seq(19,ncol(df_STRs_m),1))]), formula=...~entropy, value.var="error", fun.aggregate=mean)
df_STRs_hm$platform = factor(df_STRs_hm$platform, levels=c("HiSeq2500", "HiSeq4000", "HiSeqX10", "NovaSeq", "NextSeq", "Genapsys","Flongle",
                                                              "BGISEQ500", "MGISEQ2000", "PacBioCLR", "PacBioCCS",  "GridION", "PromethION"))
head(df_STRs_hm)
```


```{r}
df_STRs_hm_matrix = as.matrix(df_STRs_hm[,6:ncol(df_STRs_hm)])
rownames(df_STRs_hm_matrix) = df_STRs_hm[,1]
df_STRs_hm_matrix = df_STRs_hm_matrix[!grepl("PromethION_LAB01_Son1_REP01|Flongle|Genapsys", rownames(df_STRs_hm_matrix)), ]
df_STRs_hm_matrix[1:5, 1:5]
```


```{r}
ha_names_STR = HeatmapAnnotation(. = subset(df_STRs_hm, allinfo != "PromethION_LAB01_Son1_REP01" & 
                                         allinfo != "Genapsys_LAB01_Son1_REP01" & 
                                         allinfo != "Flongle_LAB01_Son1_REP01.NativeBC")$platform,
                             col = list(. = c("BGISEQ500"="#fd8d3c", "HiSeq2500"="#bdd7e7", "HiSeq4000"="#6baed6", "NextSeq"="#d0d0ef",
                                                     "HiSeqX10"="#3182bd", "MGISEQ2000"="#e6550d", "NovaSeq"="#756bb1", "PromethION"="#006d2c",
                                                      "PacBioCLR" = "#ef3b2c", "PacBioCCS" = "#a50f15", "GridION" = "#41ab5d")),
                                              # "Flongle" = "#a1d99b", "Genapsys"="#ffd92f",
                             which="row")
```

```{r}
ha_labels_STR = columnAnnotation(foo = anno_mark(at = seq(1, 100, 5), 
                                             labels=colnames(df_STRs_hm_matrix)[seq(1, 100, 5)],
                                             side="bottom", labels_gp = gpar(fontsize=8)))
```


```{r}
hm_STR = Heatmap(df_STRs_hm_matrix[,1:100],
             rect_gp = gpar(col = "white", lwd = 0.25),
             #col = colorRamp2(c(0, 0.01, 0.02, 0.03, 0.04, 0.08, 0.12, 0.16), c( "#f2f0f7", "#dadaeb","#6a51a3","#7b3294","#d7191c", "#99000d", "#cb181d", "#fb6a4a")),
             col = colorRamp2(c(0, 0.01, 0.02, 0.03, 0.04), c( "#2c7bb6", "#abd9e9","#ffffbf","#fdae61","#d7191c")),
             # col  = colorRamp2(c(0, max(df_STRs_hm_matrix)*0.25, 
             #                        max(df_STRs_hm_matrix)*0.5, 
             #                        max(df_STRs_hm_matrix)*0.75, 
             #                        max(df_STRs_hm_matrix)), c( "#2c7bb6", "#abd9e9","#ffffbf","#fdae61","#d7191c")),
             show_row_names = F,
             row_split = subset(df_STRs_hm, allinfo != "PromethION_LAB01_Son1_REP01" & 
                                         allinfo != "Genapsys_LAB01_Son1_REP01" & 
                                         allinfo != "Flongle_LAB01_Son1_REP01.NativeBC")$platform,
             row_title_rot=0,
             row_title_gp = gpar(fontsize = 10),
             cluster_rows = F,
             
             cluster_columns = F,
             show_column_names = F,
             column_title_gp = gpar(fontsize=12),
             bottom_annotation = ha_labels_STR,
             column_title = "STR Entropy",
             column_title_side = "bottom",
             
             show_heatmap_legend = T,
             heatmap_legend_param = list(title = "Error Rate"),
             width = unit(14, "cm"))
```

```{r, fig.width=8, fig.height=7}
draw(hm_STR + ha_names_STR, merge_legend = TRUE, heatmap_legend_side="right")
```


```{r, fig.width=12, fig.height=5}
pdf(file="../figures/STR_heatmap_squat2.pdf", width=15, height=4)
draw(hm_homo + ha_names_homo + hm_STR + ha_names_STR, merge_legend = TRUE, heatmap_legend_side="right", ht_gap = unit(c(0.2,1,0.2), "cm"))
dev.off()
```


```{r, fig.width}
pdf(file="../figures/UCSC_error_STRs.pdf", width=10, height=5)
draw(hm + ha_names, merge_legend = TRUE, heatmap_legend_side="right")
dev.off()
```


################################ UCSC hap.py ##########################


```{r}
df_happy = read.csv("/athena/masonlab/scratch/projects/abrf_ngs_phaseii/analysis/happy/happy_by_UCSCcontext_v2.csv")
df_happy = separate(df_happy, col="sample", by="_", into=c("platform","lab","genome","rep"), remove=F)
df_happy$type = factor(df_happy$type, levels=c("SNP", "INDEL"))
head(df_happy)
```


```{r, fig.width=8, fig.height=12}
UCSC_happy = ggplot(df_happy, aes(x=recall, y=precision, color=platform)) +
  geom_point(size=2.5, alpha=0.7) +
  facet_grid(context~type) +
  #scale_color_manual(values=c("HiSeq2500"="#9ecae1", "HiSeq4000"="#6baed6", "HiSeqX10"="#3182bd", "NovaSeq"="#08519c", "BGISEQ500"="#fc8d59", "MGISEQ2000"="#e34a33", "NextSeq"="#756bb1", "PacBioCCS"="#74c476", "PacBioCLR"="#31a354", "PromethION"="#006d2c")) +  
  scale_color_manual(values=colorsList) +
  theme_bw() +
  theme(text = element_text(size = 14),
        legend.position="bottom",
        #legend.position = c(0.95, 0),
        #legend.justification = c(0.95, 0),
        #axis.text.x = element_text(angle = 90, hjust = 1),
        strip.text.y = element_text(angle=90))
UCSC_happy
```


```{r}
pdf(file="../figures/UCSC_happy_SNPsINDELs.pdf", width=8, height=12)
UCSC_happy
dev.off()
```


# Use vcfeval

```{r}
df_vcfeval_SNPs = read.csv("/athena/masonlab/scratch/projects/abrf_ngs_phaseii/analysis/context/ucsc/vcfeval/rtgeval_ALL_SNPs.csv")
df_vcfeval_INDELs = read.csv("/athena/masonlab/scratch/projects/abrf_ngs_phaseii/analysis/context/ucsc/vcfeval/rtgeval_ALL_INDELs.csv")
df_vcfeval_SNPs$VariantType = "SNPs"
df_vcfeval_INDELs$VariantType="INDELs"
df_vcfeval = rbind(df_vcfeval_SNPs, df_vcfeval_INDELs)
df_vcfeval$VariantType = factor(df_vcfeval$VariantType, levels=c("SNPs", "INDELs"))
df_vcfeval = subset(df_vcfeval, df_vcfeval$score %in% c(0,20,40,60,80,99))
df_vcfeval = separate(df_vcfeval, col="allinfo", sep="_", into=c("platform","lab","genome","rep","context"), remove=F)
df_vcfeval$platform = factor(df_vcfeval$platform, levels=c("BGISEQ500", "MGISEQ2000", "HiSeq2500", "HiSeq4000", "HiSeqX10", "NovaSeq"))

head(df_vcfeval)
```

```{r}
colors=c("BGISEQ500"="#fd8d3c", "HiSeq2500"="#6baed6", "HiSeq4000"="#3182bd", "HiSeqX10"="#08519c", "MGISEQ2000"="#e6550d", "NovaSeq"="#f768a1")
```


```{r, fig.width=7, fig.height=14}
ggplot(data=df_vcfeval, aes(x=sensitivity, y=precision, color=platform)) + 
  geom_point() + 
  #geom_line() + 
  theme_bw() +
  theme(axis.text.y = element_text(size=15)) +
  facet_grid(context~VariantType, scales="free") + 
  scale_color_manual(values=colors)  #+  geom_text(aes(label=score),hjust=-0.5,vjust=0)
  #coord_cartesian(xlim = c(0.5, 1), ylim=c(0.975, 1))
```
````
