## Supplementary Data 3 for "Multi-Platform Assessment of DNA Sequencing Performance using Human and Bacterial Reference Genomes in the ABRF Next-Generation Sequencing Study": HumanExomeComparisons.Rmd

````
---
title: "ABRF NGS2: Exome Analysis"
author: "Jonathan Foox"
date: "May 15, 2020"
output: 
  html_notebook: 
    code_folding: none
    highlight: tango
    toc: no
    
---

```{r}
setwd("/athena/masonlab/scratch/projects/abrf_ngs_phaseii/analysis")
```

```{r}
library(reshape2)
library(dplyr)
library(tidyr)
library(cowplot)
library(ggplot2)
library(ggridges)
library(ggpubr)
```

# Marta-based #
```{r}
snps =   read.csv("variants/non_filtered_vcfs_exome/rtgeval_ALL_SNPs.csv", header=F)
indels = read.csv("variants/non_filtered_vcfs_exome/rtgeval_ALL_INDELs.csv", header=F)
```

```{r}
colnames(snps) = c("Technology", "score", "true_positives_baseline", "false_positives", "true_positives_call", "false_negatives", "precision", "sensitivity", "f_measure")
colnames(indels) = c("Technology", "score", "true_positives_baseline", "false_positives", "true_positives_call", "false_negatives", "precision", "sensitivity", "f_measure")
snps$VariantType = "SNPs"
indels$VariantType="Indels"
```

```{r}
new_colors=c("BGISEQ500"="#fd8d3c", "HiSeq2500"="#6baed6", "HiSeq4000"="#3182bd", "HiSeqX10"="#08519c", "MGISEQ2000"="#e6550d", "NovaSeq"="#f768a1", "Proton"="#74c476", "S5"="#31a354", "NextSeq"="#f39a2a")
```

```{r}
combo = rbind(snps, indels)
combo$VariantType = factor(combo$VariantType, levels=c("SNPs", "Indels"))
combo = subset(combo, combo$score %in% c(0,20,40,60,80,99))

combo = separate(combo, col="Technology", sep="_", into=c("Platform","Lab","Genome","Rep"), remove=F)

combo$Platform = factor(combo$Platform, levels=c("BGISEQ500", "MGISEQ2000", "HiSeq2500", "HiSeq4000", "HiSeqX10", "NovaSeq", "NextSeq", "Proton", "S5"))
```

```{r}
combo$type = ifelse(combo$Platform %in% c("S5", "Proton"), "exome","genome")
```

```{r}
head(combo)
```


```{r}
combo2 = combo[, c(2,6,11,12,13,14,15)]
head(combo2)
```

```{r}
combo2 = combo2 %>% group_by(Platform, score,VariantType,type) %>% summarise_all(funs(mean))
```


```{r, fig.width=14, fig.height=7}
ggplot(data=combo2, aes(x=sensitivity, y=precision, color=Platform)) + 
  geom_point(size=3) + geom_line() + 
  theme_bw() +
  theme(axis.text.y = element_text(size=15)) +
  facet_wrap(VariantType~., scales="free") + 
  scale_color_manual(values=new_colors)  + 
  geom_text(aes(label=score),hjust=-0.5,vjust=0)
  #coord_cartesian(xlim = c(0.5, 1), ylim=c(0.975, 1))
```


```{r, fig.width=12, fig.height=12}
ggplot(data=combo, aes(x=Platform, y=f_measure, color=Platform)) +
  geom_point(size=3, shape=21) +
  theme_bw() + 
  theme(axis.text.y = element_text(size=15)) +
  facet_wrap(VariantType~score) + 
  scale_color_manual(values=new_colors) 
```


# Code from Marta
```{r}
require(ggplot2)
require(reshape)

args = commandArgs(trailingOnly=TRUE)
snps_file = args[1]
indels_file = args[2]

print("Loading data...")
snps = read.table(snps_file, header=F)
indels = read.table(indels_file, header=F)

print("Preprocessing...")
colnames(snps) = c("score", "true_positives_baseline", "false_positives", "true_positives_call", "false_negatives", "precision", "sensitivity", "f_measure", "Technology", "LAB")
colnames(indels) = c("score", "true_positives_baseline", "false_positives", "true_positives_call", "false_negatives", "precision", "sensitivity", "f_measure", "Technology", "LAB")

new_colors=c("BGISEQ500"="#fd8d3c", "HiSeq2500"="#6baed6", "HiSeq4000"="#3182bd", "HiSeqX10"="#08519c", "MGISEQ2000"="#e6550d", "NovaSeq-2x150"="#f768a1", "NovaSeq-2x250"="#ae017e", "PacBioCCS"="#74c476", "PacBioCLR"="#31a354", "PromethION" = "#006d2c")

snps$VariantType = "SNPs"
indels$VariantType="Indels"

combo = rbind(snps, indels)
combo$VariantType = factor(combo$VariantType, levels=c("SNPs", "Indels"))
combo = subset(combo, combo$score %in% c(0,20,40,60,80,99))
combo$Technology = factor(combo$Technology, levels=c("BGISEQ500", "MGISEQ2000", "HiSeq2500", "HiSeq4000", "HiSeqX10", "NovaSeq-2x150", "NovaSeq-2x250","PacBioCCS", "PacBioCLR", "PromethION"))

ggplot(data=combo, aes(x=sensitivity, y=precision, color=Technology)) + geom_point(size=3) + geom_line() + theme_minimal() + facet_wrap(~VariantType, scales="free") + scale_color_manual(values=new_colors) + geom_text(aes(label=score),hjust=-0.5,vjust=0)
ggsave("Sensitivity_vs_precision_snps_indels.pdf", device="pdf", height=5, width=12)

print("Done!")

```


# My own #

```{r}
df_happy = read.csv("happy/exome/happy_summary_exome.csv")
df_happy_melted = melt(df_happy, id=c("Sample","Platform","Lab","Genome","Rep"))
head(df_happy)
```

```{r}
df_happy_TPFNFP = subset(df_happy_melted, variable=="SNP_TP" | variable=="SNP_FN" | variable=="SNP_FP" | 
                         variable=="INDEL_TP" | variable=="INDEL_FN" | variable=="INDEL_FP")
df_happy_TPFNFP$calltype = ifelse(grepl("SNP",df_happy_TPFNFP$variable),'SNP','INDEL')
df_happy_TPFNFP$calltype = factor(df_happy_TPFNFP$calltype, levels=c('SNP','INDEL'))
df_happy_TPFNFP$variable = gsub(x = df_happy_TPFNFP$variable, pattern = ".+_", replacement = "")
tail(df_happy_TPFNFP)
```

```{r}
df_happy_precRec = subset(df_happy_melted, variable=="SNP_Recall" | variable=="SNP_Precision" |
                         variable=="INDEL_Recall" | variable=="INDEL_Precision")
df_happy_precRec$calltype = ifelse(grepl("SNP",df_happy_precRec$variable),'SNP','INDEL')
df_happy_precRec$calltype = factor(df_happy_precRec$calltype, levels=c('SNP','INDEL'))
df_happy_precRec$variable = gsub(x = df_happy_precRec$variable, pattern = ".+_", replacement = "")
df_happy_precRec = dcast(data = df_happy_precRec, formula = Sample+Platform+Lab+Genome+Rep+calltype~variable, 
                         fun.aggregate = sum, value.var = "value")


tail(df_happy_precRec, 20)
```

```{r}

colorsList = list()
colorsList[["HiSeq2500"]] = "#9ecae1"
colorsList[["HiSeq4000"]] = "#6baed6"
colorsList[["HiSeqX10"]] = "#3182bd"
colorsList[["NovaSeq"]] = "#ae017e"
colorsList[["NextSeq"]] = "#d0d0ef"
colorsList[["BGISEQ500"]] = "#fc8d59"
colorsList[["MGISEQ2000"]] = "#e34a33"
colorsList[["PromethION"]] = "#006d2c"
colorsList[["PacBioCLR"]] = "#ef3b2c"
colorsList[["PacBioCCS"]] = "#a50f15"

colorsList[["Proton"]] = "#a1d99b"
colorsList[["S5"]] = "#41ab5d"
```


```{r, fig.width=10, fig.height=6}
exome_happy = ggplot(subset(df_happy_precRec, Sample!="PromethION_LAB02_Son1_REP01"), 
       aes(Recall, Precision)) +
  geom_point(aes(color=Platform, shape=Genome), size=3) + 
  scale_shape_manual("Genome", values =c("Father"=15, "Mother"=16, "Son1"=17, "Son2"=17, "Son3"=17)) +
  scale_color_manual(values=colorsList) +
  #xlim(0.985,1) + ylim(0.985,1) +
  xlab("Specificity") + ylab("Sensitivity") +
  theme(text = element_text(size=16)) +
    facet_grid(.~calltype, scales = "free", switch="y") +
  theme_bw() +
  theme(text = element_text(size=14))
exome_happy
```

```{r}
pdf(file="../figures/exome_happy.pdf", width=10, height=6)
exome_happy
dev.off()
```
````
