## Supplementary Data 3 for "Multi-Platform Assessment of DNA Sequencing Performance using Human and Bacterial Reference Genomes in the ABRF Next-Generation Sequencing Study": HumanGCheatmaps.Rmd

````
---
title: "ABRF NGS2: Human GC Mapping"
author: "Jonathan Foox"
date: "June 4, 2020"
output: 
  html_notebook: 
    code_folding: none
    highlight: tango
    toc: yes
    
---

```{r}
#library(devtools)
#install_github("jokergoo/ComplexHeatmap")
```

```{r}
setwd("/athena/masonlab/scratch/projects/abrf_ngs_phaseii/analysis")
```


```{r}
library(tidyr)
library(ggplot2)
library(ggridges)
library("scales")
library(dplyr)
library(reshape2)
```

```{r}
df_gc = read.csv("gcCoverage_human_WGSandExome.csv")
#df_gc = separate(df_gc, col="sample", into=c("platform","lab","genome","rep"), sep="_", remove=F)
#df_gc = df_gc[grepl("exome", df_gc$type), ]
df_gc = df_gc[, 2:ncol(df_gc)]
head(df_gc)
```

```{r}
df_gc_tmp = dcast(data = df_gc, formula = ...~sample, value.var="numReads", fun.aggregate=sum)
df_gc_tmp[1:10, 1:5]
```


```{r}
normalized<-function(y) {
  x<-y[!is.na(y)]
  x<-(x - min(x)) / (max(x) - min(x))
  y[!is.na(y)]<-x
  return(y)
  }
```

```{r}
df_gc2 = apply(df_gc_tmp[, colnames(df_gc_tmp)[2:ncol(df_gc_tmp)]],2,normalized)
df_gc2[1:5, 1:5]
```

```{r}
df_gc2 = melt(df_gc2)
colnames(df_gc2) = c("GC", "Sample", "Reads_Used")
df_gc2$GC = df_gc2$GC - 1
df_gc2 = separate(df_gc2, col=Sample, into=c("Platform", "Lab", "Genome", "Rep"), remove=F)

df_gc2$type = ""
df_gc2[grepl("S5|Proton", df_gc2$Sample),  ]$type = "exome"
df_gc2[!grepl("S5|Proton", df_gc2$Sample), ]$type = "wgs"

head(df_gc2)
```


```{r}
df_gc_unmelt = dcast(data = df_gc2, formula = ...~GC, value.var="Reads_Used", fun.aggregate=mean)
df_gc_unmelt[1:5, 1:16]
```

```{r}
library(ComplexHeatmap)
library(circlize)
```

```{r}
df_gc_unmelt_matrix = as.matrix(df_gc_unmelt[,7:ncol(df_gc_unmelt)])
rownames(df_gc_unmelt_matrix) = df_gc_unmelt[,1]
df_gc_unmelt_matrix[1:5, 1:10]
```

```{r}
ha_names = HeatmapAnnotation(Platform = df_gc_unmelt$Platform,
                             col = list(Platform = c("HiSeq2500" = "#9ecae1", "HiSeq4000" = "#6baed6", "HiSeqX10" = "#3182bd", "NovaSeq" = "#ae017e", "NextSeq" = "#d0d0ef", "BGISEQ500" = "#fc8d59", "MGISEQ2000" = "#e34a33", "Genapsys" = "#ffd92f", "Flongle" = "#a1d99b", "GridION" = "#41ab5d", "PromethION" = "#006d2c", "PacBioCLR" = "#ef3b2c", "PacBioCCS" = "#a50f15",
                                                     "Proton" = "#fc8d59", "S5" = "#fee08b")),
                             #show_legend = T,
                             which="row")


```

```{r}
ha_genomes = HeatmapAnnotation(Genome = df_gc_unmelt$Genome,
                             col = list(Genome = c("Mother"="#FBB4AE", "Father"="#B3CDE3", "Son"="#CCEBC5", "Son1"="#CCEBC5", "Son2"="#CCEBC5", "Son3"="#CCEBC5")),
                             #show_legend = T,
                             which="row")

```

```{r}
ha_type = HeatmapAnnotation(Type = df_gc_unmelt$type,
                             col = list(Type = c("wgs"="#b2182b", "exome"="#2166ac")),
                             #show_legend = T,
                             which="row")

```


```{r, fig.width=12, fig.height=12}
hm = Heatmap(df_gc_unmelt_matrix,
                 col  = colorRamp2(c(0, max(df_gc_unmelt_matrix)*0.25, 
                                     max(df_gc_unmelt_matrix)*0.5, 
                                     max(df_gc_unmelt_matrix)*0.75, 
                                     max(df_gc_unmelt_matrix)), c( "#2c7bb6", "#abd9e9","#ffffbf","#fdae61","#d7191c")),
                 #col = colorRamp2(c(0,0.5,1,1.5,2), c( "#2c7bb6", "#abd9e9","#ffffbf","#fdae61","#d7191c")),
                 #rect_gp = gpar(col = "white", lwd = 0.25),
                 #column_title = "Metagenomic Pool",
                 cluster_columns = FALSE,
                 show_column_names = TRUE,
             column_title = "Human Normalized Coverage across GC (WGS and Exome)",
             row_title = "Samples",
             cluster_rows = T,
             row_split = factor(df_gc_unmelt$type, levels=c("wgs", "exome")),
                 column_title_gp = gpar(fontsize = 16),
                 column_names_side = "top",
                 column_labels = c("0",".",".",".",".",".",".",".",".",".",".","10",".",".",".",".",".",".",".",".",".","20",".",".",".",".",".",".",".",".",".","30",".",".",".",".",".",".",".",".",".","40",".",".",".",".",".",".",".",".",".","50",".",".",".",".",".",".",".",".",".","60",".",".",".",".",".",".",".",".",".","70",".",".",".",".",".",".",".",".",".","80",".",".",".",".",".",".",".",".",".","90",".",".",".",".",".",".",".",".","100"),
            
                 column_names_rot = 45,
                 column_names_gp = gpar(fontsize=12),
                 
                 show_row_names = TRUE,
                 #bottom_annotation = ha_dens,
                 show_heatmap_legend = T,
                 heatmap_legend_param = list(
                   title = "Coverage"
                 ))

draw(hm + ha_type + ha_names + ha_genomes, merge_legend = TRUE, heatmap_legend_side="right")
```

```{r, fig.width=7, fig.height=6}
pdf("../figures/human_GC_distribution.pdf", width=8, height=10)
draw(hm + ha_type + ha_names + ha_genomes, merge_legend = TRUE, heatmap_legend_side="right")
dev.off()
```
````
