## Supplementary Data 3 for "Multi-Platform Assessment of DNA Sequencing Performance using Human and Bacterial Reference Genomes in the ABRF Next-Generation Sequencing Study": HumanNormalizedCoverageAllVsAllComparisons.Rmd

````
---
title: "ABRF NGS2: Coverage Heatmap"
author: "Jonathan Foox"
date: "May 26, 2020"
output: 
  html_notebook: 
    code_folding: none
    highlight: tango
    toc: yes
    
---

```{r}
setwd("/athena/masonlab/scratch/projects/abrf_ngs_phaseii/analysis/")
```

```{r}
library(reshape2)
library(dplyr)
library(tidyr)
library(cowplot)
library(ggplot2)
library(ggridges)
library(ggpubr)
```


```{r}
df = read.csv("Heatmap-lt30X.csv")
head(df)
```

```{r, fig.width=8, fig.height=8}
ggplot(df, aes(x=Sample1, y=Sample2, fill=value)) +
  geom_tile() +
  geom_text(aes(label=round(value, digits = 2)), size=1.5) +
  facet_wrap(~name, nrow=2) +
  scale_fill_gradientn(colors=c('#00429d', '#5681b9', '#93c4d2', '#ffffff', '#ffa59e', '#dd4c65', '#93003a')) +
  theme_bw() +
  theme(axis.text.x = element_text(angle=90,vjust=.5,hjust=1), 
        legend.position = 'bottom', 
        axis.title = element_blank(), 
        strip.background = element_rect(fill="white"), 
        strip.text = element_text(colour = 'black'))
```


```{r}
df_jf = read.csv('Heatmap-lt30X.JF.redundant.csv')
df_jf_m = melt(df_jf, id.vars = c("context","sample1and2","sample1","sample2"), measure.vars = c("value1","value2","equal"))
df_jf_m$variable = factor(df_jf_m$variable, levels=c("value1","equal","value2"))
head(df_jf_m)
```


```{r, fig.width=14, fig.height=5}
jon_cov = ggplot(df_jf_m, aes(x=sample1, y=value, fill=variable)) +
  geom_bar(stat="identity", position="fill") +
  facet_grid(.~context, scales="free", space="free", switch = "y") +
  theme_bw() +
  theme(axis.text.x = element_text(angle=90, hjust=1, vjust=0.5),
        axis.ticks.x = element_blank(),
        strip.text.y = element_text(angle=0),
        text = element_text(size=16),
        legend.position="bottom") +
  scale_fill_manual(labels=c("Platform", "Equivalent", "All other platforms"),
    values=c("#3288bd","gray","#d53e4f")) +
  #scale_x_discrete(labels=df_jf_m$sample2, position = "top") +
  geom_hline(yintercept = 0.5, linetype="dashed") +
  xlab("") + ylab("") +
  #geom_text(aes(label=sample2)) +
  coord_flip()
jon_cov
```

```{r}
pdf(file="../figures/cov_jon.pdf", width=14, height=5)
jon_cov
dev.off()
```
````
