## Supplementary Data 3 for "Multi-Platform Assessment of DNA Sequencing Performance using Human and Bacterial Reference Genomes in the ABRF Next-Generation Sequencing Study": HumanNormalizedCoverageComparisons.rmd

````
---
title: "ABRF"
output: html_notebook
---

Setup working directory and functions

```{r}

library(reshape2)
library(ggplot2)
library(tidyr)
library(vroom)
library(dplyr)
library(ggridges)

colors <- c("HiSeq2500_126" = "#bdd7e7",
            "HiSeq2500_100" = "#bdd7e7",
            "HiSeq4000_150" = "#6baed6",
            "HiSeqX10_150"  = "#3182bd",
            "NovaSeq_150" = "#f768a1",
            "NovaSeq_250" = "#ae017e",
            "BGISEQ500_100" = "#fd8d3c",
            "MGISEQ2000_150" = "#e6550d",
            "PacBioCLR_7.8kb" = "#31a354",
            "PacBioCCS_13.5kb" = "#74c476",
            "PromethION_8kb" = "#006d2c")

techs <- c(
  "BGISEQ500_100",
  "MGISEQ2000_150",
  "HiSeq2500_100",
  "HiSeq2500_126",
  "HiSeq4000_150",
  "HiSeqX10_150",
  "NovaSeq_150",
  "NovaSeq_250",
  "PacBioCCS_13.5kb",
  "PacBioCLR_7.8kb",
  "PromethION_8kb")

ids <- c(
  "BGISEQ500_L1_100",
  "MGISEQ2000_L1_150",
  "HiSeq2500_L3_100",
  "HiSeq2500_L1_126",
  "HiSeq2500_L2_126",
  "HiSeq4000_L1_150", 
  "HiSeq4000_L2_150",
  "HiSeq4000_L3_150",
  "HiSeqX10_L1_150",
  "HiSeqX10_L2_150",
  "HiSeqX10_L3_150",
  "NovaSeq_L1_150",
  "NovaSeq_L2_150",
  "NovaSeq_L2_250",
  "PacBioCCS_L1_13.5kb",
  "PacBioCLR_L1_7.8kb",
  "PromethION_L1_8kb")

columns <- c(
  "Chr",
  "Start",
  "End",
  "Annotation",
  "BGISEQ500_L1_Father_100",
  "BGISEQ500_L1_Mother_100",
  "BGISEQ500_L1_Son_100",
  "HiSeq2500_L1_Father_126",
  "HiSeq2500_L1_Mother_126",
  "HiSeq2500_L1_Son_126",
  "HiSeq2500_L2_Father_126",
  "HiSeq2500_L2_Mother_126",
  "HiSeq2500_L2_Son_126",
  "HiSeq2500_L3_Father_100",
  "HiSeq2500_L3_Mother_100",
  "HiSeq2500_L3_Son_100",
  "HiSeq4000_L1_Father_150",
  "HiSeq4000_L1_Mother_150",
  "HiSeq4000_L1_Son_150",
  "HiSeq4000_L2_Father_150",
  "HiSeq4000_L2_Mother_150",
  "HiSeq4000_L2_Son_150",
  "HiSeq4000_L3_Father_150",
  "HiSeq4000_L3_Mother_150",
  "HiSeq4000_L3_Son_150",
  "HiSeqX10_L1_Father_150",
  "HiSeqX10_L1_Mother_150",
  "HiSeqX10_L1_Son_150",
  "HiSeqX10_L2_Father_150",
  "HiSeqX10_L2_Mother_150",
  "HiSeqX10_L2_Son_150",
  "HiSeqX10_L3_Father_150",
  "HiSeqX10_L3_Mother_150",
  "HiSeqX10_L3_Son_150",
  "MGISEQ2000_L1_Father_150",
  "MGISEQ2000_L1_Mother_150",
  "MGISEQ2000_L1_Son_150",
  "PromethION_L1_Son_8kb",
  "NovaSeq_L1_Father_150",
  "NovaSeq_L1_Mother_150",
  "NovaSeq_L1_Son_150",
  "NovaSeq_L2_Father_150",
  "NovaSeq_L2_Father_250",
  "NovaSeq_L2_Mother_150",
  "NovaSeq_L2_Mother_250",
  "NovaSeq_L2_Son_150",
  "NovaSeq_L2_Son_250",
  "PacBioCCS_L1_Son_13.5kb",
  "PacBioCLR_L1_Son_7.8kb"
)

ihs <- function(x) {
  y <- log(x + sqrt(x ^ 2 + 1))
  return(y)
}

dist_point_line <- function(r, slope, intercept) {
  x <- as.numeric(r[3])
  y <- as.numeric(r[4])
  return((-slope*x + y - intercept)/sqrt(sum((-slope)^2 + 1)))
}


```

Ridge plots of HG002 data. 

```{r}
dat <- vroom::vroom("per-sample_coverage_no-rep02_LR_merged-zero-filtered_repeat-only.txt")
names(dat) <- columns

m <- melt(dat, id=c("Chr", "Start", "End", "Annotation"), variable.name = "ID", value.name = "Avg_Cov")

m <- separate(m, ID, c("Tech","Lab","Ind","ReadLen"),sep="_")
m$ID <- paste(m$Tech, m$Lab, m$ReadLen, sep="_")
m$Tech <- paste(m$Tech, m$ReadLen, sep="_")
ms <- subset(m, Ind == "Son")

ms$ID <- factor(ms$ID,levels = ids)

pdf(file="GIAB_tech-by-repeat_coverage-son_LR_Ridge_0-100.pdf", width=12.5,height=6)
ggplot(data=ms, aes(y=ID, x=ihs(Avg_Cov),fill=Tech,color=Tech,alpha=0.85)) +
  geom_density_ridges() +
  facet_wrap(~Annotation,nrow=2) +
  scale_x_continuous(breaks=c(0,ihs(10), ihs(25), ihs(50),ihs(100)), labels = c("0X","10X","25X","50X","100X")) +
  coord_cartesian(xlim=c(0,ihs(100))) +
  scale_color_manual(values=colors) + 
  scale_fill_manual(values=colors) +
  xlab("Average Coverage") +
  ylab("Technology") +
  theme_ridges() +
  theme(axis.text.y = element_text(size=12,color="grey34"), axis.text.x = element_text(size=10), axis.title.y = element_blank(), axis.title.x = element_text(hjust=0.5,size=14), strip.background = element_rect(fill="white"), strip.text = element_text(colour = 'black', size=12), legend.position = "none")
dev.off()
```

Pairwise hex density plots

```{r}
Alu <- vroom::vroom("son-only_Alu_merge.txt")
L1 <- vroom::vroom("son-only_L1_merge.txt")
L2 <- vroom::vroom("son-only_L2_merge.txt")
Low_Complexity <- vroom::vroom("son-only_Low_complexity_merge.txt")
LTR <- vroom::vroom("son-only_LTR_merge.txt")
Satellite <- vroom::vroom("son-only_Satellite_merge.txt")
Simple_Repeat <- vroom::vroom("son-only_Simple_repeat_merge.txt")
Telomere <- vroom::vroom("son-only_telo_merge.txt")

for (repType in c("Alu", "L1", "L2", "Low_Complexity", "LTR", "Satellite", "Simple_Repeat", "Telomere")) {
  for (v1 in 1:11) {
    v2 <- v1+1
    if (v1 == 11) {
      break
    }
    for (v3 in v2:11) {
      samX <- techs[v1]
      samY <- techs[v3]
      
      name <- paste("GIAB_",repType,"_",samX,"-v-",samY,"_hex.pdf", sep="")
      
      p <- ggplot(get(repType), aes(x=ihs(get(samX)), y=ihs(get(samY)))) +
        geom_hex(aes(fill = stat(log10(count))), bins=250) +
        scale_color_viridis_c() + 
        scale_fill_viridis_c() +
        geom_abline(slope=1, intercept=0, color="red") +
        scale_y_continuous(breaks=c(0, ihs(5), ihs(10), ihs(30), ihs(100), ihs(500)), labels = c("0X", "5X","10X","30X","100X","500X")) +
        scale_x_continuous(breaks=c(0, ihs(5), ihs(10), ihs(30), ihs(100), ihs(500)), labels = c("0X", "5X","10X","30X","100X","500X")) +
        ylab(samY) +
        xlab(samX) +
        theme_bw() +
        theme(strip.background = element_rect(fill="black"), strip.text = element_text(colour = 'white'), legend.position = 'bottom')
      
      pdf(file=name, width=8,height=8)
      print(p)
      dev.off()
    }
  }
}
```

Restricted to <30X Pairwise heatmap data

```{r}
aluPair <- vroom::vroom("son-only_Alu_merge_pairwise.txt")
aluPair$dist <- apply(aluPair, 1, dist_point_line, 1, 0)

l1Pair <- vroom::vroom("son-only_L1_merge_pairwise.txt")
l1Pair$dist <- apply(l1Pair, 1, dist_point_line, 1, 0)

l2Pair <- vroom::vroom("son-only_L2_merge_pairwise.txt")
l2Pair$dist <- apply(l2Pair, 1, dist_point_line, 1, 0)

ltrPair <- vroom::vroom("son-only_LTR_merge_pairwise.txt")
ltrPair$dist <- apply(ltrPair, 1, dist_point_line, 1, 0)

lcPair <- vroom::vroom("son-only_Low_complexity_merge_pairwise.txt")
lcPair$dist <- apply(lcPair, 1, dist_point_line, 1, 0)

satPair <- vroom::vroom("son-only_Satellite_merge_pairwise.txt")
satPair$dist <- apply(satPair, 1, dist_point_line, 1, 0)

simPair <- vroom::vroom("son-only_Simple_repeat_merge_pairwise.txt")
simPair$dist <- apply(simPair, 1, dist_point_line, 1, 0)

teloPair <- vroom::vroom("son-only_telo_merge_pairwise.txt")
teloPair$dist <- apply(teloPair, 1, dist_point_line, 1, 0)


df <- data.frame()
for (name in ls(pattern= "Pair")) {
  mat <- matrix(, nrow = 11, ncol = 11)
  for (c1 in 1:11) {
    c2 <- c1+1
    if (c1 == 11) {
      break
    }
    for (c3 in c2:11) {
      colX <- techs[c1]
      colY <- techs[c3]
      sub <- subset(get(name), (Col1 == colX & Col2 == colY & Val1 <= 30 & Val2 <= 30))
      xbias <- sum(sub$dist < 0)
      ybias <- sum(sub$dist > 0)
      zcalls <- sum(sub$dist == 0)
      mat[c3,c1] <- xbias/(xbias + ybias + zcalls)
      mat[c1,c3] <- ybias/(xbias + ybias + zcalls)
    }
  }
  rownames(mat) <- techs
  colnames(mat) <- techs
  
  pm <- melt(mat)
  names(pm) <- c("Sample1", "Sample2", "value")
  pm$name <- name
  
  df <- rbind(df,pm)
}

df[df$name == "aluPair",]$name <- "Alu"
df[df$name == "l1Pair",]$name <- "L1"
df[df$name == "l2Pair",]$name <- "L2"
df[df$name == "lcPair",]$name <- "Low Complexity"
df[df$name == "simPair",]$name <- "Simple Repeat"
df[df$name == "satPair",]$name <- "Satellite"
df[df$name == "teloPair",]$name <- "Telomere"
df[df$name == "ltrPair",]$name <- "LTR"

pdf(file="GIAB_Mask_Distances_lt30X.pdf", width=8,height=6)
ggplot(df, aes(x=Sample1, y=Sample2, fill=value)) + 
  geom_tile() +
  geom_text(aes(label=round(value, digits = 2)), size=1.5) +
  facet_wrap(~name, nrow=2) +
  scale_fill_gradientn(colors=c('#00429d', '#5681b9', '#93c4d2', '#ffffff', '#ffa59e', '#dd4c65', '#93003a')) +
  theme_bw() +
  theme(axis.text.x = element_text(angle=90,vjust=.5,hjust=1), legend.position = 'bottom', axis.title = element_blank(), strip.background = element_rect(fill="white"), strip.text = element_text(colour = 'black'))
dev.off()

```
````
