## Supplementary Data 3 for "Multi-Platform Assessment of DNA Sequencing Performance using Human and Bacterial Reference Genomes in the ABRF Next-Generation Sequencing Study": HumanSNPandINDELrtg.pdf

### Sensitivity vs. precision of SNP/INDEL calls.

1. For each sequencing platform VCF, run RTG vcfeval against the GIAB SNV/INDEL truth set:

```
rtg RTG_MEM=20G vcfeval \
-b ${GIAB_SNV_truth} \
-c ${vcfeval} \
-e ${GIAB_SNV_bed} \
-o ${outdir}/${output_prefix} \
-t ${reference} \
-T 2
```

2. Postprocessing:
    - INDEL outputs: concatenated non_snp_roc.tsv.gz files (outputs of rtg vcfeval) from all platforms (excluding the headers and including a sequencing platform label in the last column) and saved the output to combo_indels.txt.

    - SNP outputs: concatenated snp_roc.tsv.gz files from all platforms (the same way as above) and saved the output to combo_snps.txt.

3. Plotting sensitivity vs. precision of SNP and INDEL calling for all sequencing platforms using the following R script:

```
require(ggplot2)
require(reshape)

args <- commandArgs(trailingOnly=TRUE)
snps_file <- args[1] ## path to combo_snps.txt from step 2.
indels_file <- args[2] ## path to combo_indels.txt from step 2.

snps <- read.table(snps_file, header=F)
indels <- read.table(indels_file, header=F)

colnames(snps) <- c("score", "true_positives_baseline", "false_positives",
"true_positives_call", "false_negatives", "precision", "sensitivity",
"f_measure", "Technology", "LAB")

colnames(indels) <- c("score", "true_positives_baseline",
"false_positives", "true_positives_call", "false_negatives", "precision",
"sensitivity", "f_measure", "Technology", "LAB")
```

```r
## Define colors for each sequencing platform:
custom_colors <- c("BGISEQ500"="#fd8d3c", "HiSeq2500"="#6baed6",
"HiSeq4000"="#3182bd", "HiSeqX10"="#08519c", "MGISEQ2000"="#e6550d",
"NovaSeq-2x150"="#f768a1", "NovaSeq-2x250"="#ae017e",
"PacBioCCS"="#74c476", "PacBioCLR"="#31a354", "PromethION" = "#006d2c")

## Add a variant type label:
snps$VariantType <- "SNPs"
indels$VariantType <-"Indels"

## Combine SNP and INDEL data frames:
combo <- rbind(snps, indels)

## Set the desired order in which variant types will appear on the plot:
combo$VariantType <- factor(combo$VariantType, levels=c("SNPs", "Indels"))

## Subset out rows with GQ scores=c(0,20,40,60,80,99) to be included in the
plot:
combo <- subset(combo, combo$score %in% c(0,20,40,60,80,99))

## Set the desired order in which sequencing platforms will be listed in
the plot:
combo$Technology <- factor(combo$Technology, levels=c("BGISEQ500",
"MGISEQ2000", "HiSeq2500", "HiSeq4000", "HiSeqX10", "NovaSeq-2x150",
"NovaSeq-2x250","PacBioCCS", "PacBioCLR", "PromethION"))

## Generate sensitivity vs. precision plot for all sequencing platforms,
stratified by variant type:
ggplot(data=combo, aes(x=sensitivity, y=precision, color=Technology)) + \
geom_point(size=3) + \
geom_line() + \
theme_minimal() + \
facet_wrap(~VariantType, scales="free") + \
scale_color_manual(values=custom_colors) + \
geom_text(aes(label=score),hjust=-0.5,vjust=0)
```

### Plotting counts of true positive SNP/INDEL calls per UCSC RepeatMask context.

1. Stratify high confidence true positive SNP/INDEL call sets for each sequencing platform (vcf="tp-baseline.vcf.gz" outputs of *RTG vcfeval* generated as described in step 1 of the "Sensitivity vs. precision of SNP/INDEL calls" section above):

```
rtg vcffilter \
--non-snps-only \
-i ${vcf} \
-o ${outdir}/${vcf_prefix}.nonSNPs.vcf.gz

rtg vcffilter \
--snps-only \
-i ${vcf} \
-o ${outdir}/${vcf_prefix}.SNPs.vcf.gz
```

2. Intersect high confidence TP SNP and INDEL calls with the UCSC RepeatMask bed files (one bed file for each repeat category) and, for each sequencing platform, output number of SNPs and INDEL overlapping each repeat category:

```
for variantType in SNPs nonSNPs; do
echo ${vcf_prefix},${variantType},${bed_prefix},$(bedtools intersect -u -wa
-a ${outdir}/${vcf_prefix}.${variantType}.vcf.gz -b ${bed} | wc -l);
done
```

3. Concatenate all the logs with counts of SNPs and INDELs from step 2.

4. Plotting counts of high confidence TP SNP/INDEL calls per UCSC RepeatMask for each sequencing platform using the following R script:

```
require(ggplot2)
args <- commandArgs(trailingOnly=TRUE)
Combo_file <- args[1]
combo <- read.table(combo_file, header=F)
```

```
colnames(combo) <- c("Technology", "variantType", "Repeat_category",
"Count")

combo$Count <- as.numeric(combo$Count)

## Add a variant type label to the data frame and set a desired order of
variant types to appear in the plot:
combo$VariantType <- ifelse(combo$variantType == "SNPs", "SNPs", "INDELs")
combo$VariantType <- factor(combo$VariantType, levels=c("SNPs", "INDELs"))

## Define colors for each sequencing platform:
custom_colors <- c("BGISEQ500"="#fd8d3c", "HiSeq2500"="#6baed6",
"HiSeq4000"="#3182bd", "HiSeqX10"="#08519c", "MGISEQ2000"="#e6550d",
"NovaSeq-2x150"="#f768a1", "NovaSeq-2x250"="#ae017e",
"PacBioCCS"="#74c476", "PacBioCLR"="#31a354", "PromethION" = "#006d2c")

## Set the desired order in which sequencing platforms will be listed in
the plot:
combo$Technology <- factor(combo$Technology, levels=c("BGISEQ500",
"MGISEQ2000", "HiSeq2500", "HiSeq4000", "HiSeqX10", "NovaSeq-2x150",
"NovaSeq-2x250", "PacBioCCS", "PacBioCLR", "PromethION"))

## Plot a barplot of high confidence TP SNP/INDEL counts per UCSC
RepeatMask regions:
ggplot(combo, aes(x=Repeat_category, y=Count, fill=Technology)) + \
geom_bar(stat="identity", position="dodge") + \
facet_wrap(~VariantType,ncol=2, scales="free_y") + \
theme_minimal() + \
theme(axis.text.x=element_text(angle=90,
hjust=1),axis.title.x=element_blank()) + \
scale_fill_manual(values=custom_colors)
```

#### Generation of genome-wide indel size distribution plots.

1. Run *RTG vcfstats* on every VCF to get a histogram of allele lengths:

```
vcf=${1} ## path to SNP/INDEL variant call set for a given sequencing
platform
```

```
output_prefix=${2}
seq_platform_label=${3} ## e.g. HiSeqX10, BGISEQ, etc.
outdir=${4}

rtg RTG_MEM=20G vcfstats \
--allele-lengths \
${vcf} > ${tmp}/${output_prefix}.vcfstats

## Postprocessing of the vcfstats output:
cat ${tmp}/${output_prefix}.vcfstats | sed '1,20d' | cut -f1,4- \
| awk -v var=${seq_platform_label} '{if($1 ~ "-") print substr($1, 1,
length($1)-4)"\t"$2"\t"$3"\t"$4"\t"var; else print $0"\t"var}' \
> ${outdir}/${output_prefix}.vcfstats_short
```

2. Concatenate all *${outdir}/${output_prefix}.vcfstats_short* files and remove header lines:

```
cat ${outdir}/*.vcfstats_short | grep -v "length" > combo
```

3. Plot genome-wide indel size distribution for all sequencing platforms using the following R script:

```
require(ggplot2)
require(reshape)

args <- commandArgs(trailingOnly=TRUE)

combo_file <- args[1] ## path to combo file from step 2 above.
combo <- read.table(combo_file, header=F)

## Label the columns:
colnames(combo) <- c("length","Delete","Insert", "Indel", "Technology")

## Select all columns except for "Indel" (rtg vcfstats classifies the vast
majority of INDELs as either "Insert" or "Delete" in cases where there is
pure addition or removal of bases. Cases where there is a length change
between REF and ALT that is not pure (e.g. ATT --> CTTT) are classified as
"Indel" and where skipped in this analysis) and convert the data frame into
a molten data frame::
combo_m <- melt(combo[,c(1,2,3,5)], id=c("length", "Technology"))

## Create a new Length column in which length of deletions is expressed as
```

```
a negative value (keeping length of insertions as a positive value), for
plotting purposes:
combo_m$Length <- ifelse(combo_m$variable == "Delete", -(combo_m$length),
combo_m$length)

## Define colors for each sequencing platform:
custom_colors <- c("BGISEQ500"="#fd8d3c", "HiSeq2500"="#6baed6",
"HiSeq4000"="#3182bd", "HiSeqX10"="#08519c", "MGISEQ2000"="#e6550d",
"NovaSeq_2x150"="#f768a1", "NovaSeq_2x250"="#ae017e",
"PacBioCCS"="#74c476", "PacBioCLR"="#31a354", "PromethION" = "#006d2c")

## Set the desired order in which sequencing platforms will be listed in
the plot:
combo_m$Technology <- factor(combo_m$Technology, levels=c("BGISEQ500",
"MGISEQ2000", "HiSeq2500", "HiSeq4000", "HiSeqX10", "NovaSeq_2x150",
"NovaSeq_2x250", "PacBioCCS", "PacBioCLR", "PromethION"))

## Plot the genome-wide size distribution of short deletions and insertions
for all platforms:
ggplot(data=combo_m, aes(x=Length, y=value+1, color=Technology)) + \
geom_point(size=1, alpha=0.3) + \
theme_minimal() + \
xlab("size (bp)") + \
ylab("count+1") + \
scale_y_continuous(trans='log10') + \
xlim(-100,100) + \
scale_color_manual(values=custom_colors) + \
stat_smooth(alpha=0.3, se=FALSE, span=0.30)
```

#### Generation of indel size distribution plots restricted to high confidence true positive indels only.

1. For every variant call set, identify high confidence true positive (TP) calls. Command below was executed separately for the GIAB SNV/INDEL (includes INDELs up to 50bp in size) and SV truth sets (includes INDELs > 50bp in size):

```
rtg RTG_MEM=20G vcfeval \
-b ${GIAB_truth} \
-c ${vcfeval} \
-e ${GIAB_bed} \
```

```
-o ${outdir}/${output_prefix} \
-t ${reference_sdf} \
-T 2
```

2. Run *RTG vcfstats* only on TP variant calls (identified separately for SNV/INDEL and SV truth sets from step1 above):

```
rtg RTG_MEM=20G vcfstats \
--allele-lengths \
${outdir}/${output_prefix}/tp-baseline.vcf.gz \
> ${tmp}/${output_prefix}.${category}.vcfstats

## Postprocessing of the output file (rtg vcfstats outputs cumulative
counts for indels of size > 100bp and it labels them as e.g. "100-199",
"200-299" etc. We modified the label to just say e.g. "100", "200", etc for
plotting purposes):
cat ${tmp}/${output_prefix}.${category}.vcfstats | sed '1,20d' \
| cut -f1,4- | grep -v Variant \
| awk '{if($1 ~ "-") print substr($1, 1, length($1)-4)"\t"$2"\t"$3"\t"$4;
else print $0}' > ${outdir}/${output_prefix}.tp-baseline.vcfstats_short
```

3. Run *RTG vcfstats* on the GIAB SNV/INDEL and SV truth sets:

```
## Subset GIAB truth sets to high confidence regions only:
bcftools view \
-R ${high_conf_SV_bed} \
-Oz \
-o ${GIAB_SV_truth_set_prefix}.highConfRegions.vcf.gz \
${GIAB_SV_truth_set}

tabix -p vcf ${GIAB_SV_truth_set_prefix}.highConfRegions.vcf.gz

bcftools view \
-R ${high_conf_SNVindel_bed} \
-Oz \
-o ${GIAB_SNV_truth_set_prefix}.highConfRegions.vcf.gz \
${GIAB_SNV_truth_set}

tabix -p vcf HG002_GRCh37_GIAB_highconf_nonSNPs_highConfRegions.vcf.gz

## Run RTG vcfstats on GIAB truth sets (high confidence variants only):
```

```
for vcf in ${GIAB_SV_truth_set_prefix}.highConfRegions.vcf.gz
${GIAB_SNV_truth_set_prefix}.highConfRegions.vcf.gz; do

prefix=$(echo ${vcf} | sed 's/.vcf.gz//g');
label="GIAB_truth_set";

rtg RTG_MEM=20G vcfstats \
--allele-lengths \
${vcf} \
> ${tmp}/${prefix}.tp-baseline.vcfstats;

cat ${tmp}/${prefix}.tp-baseline.vcfstats | sed -n '/^length/,$p' | cut
-f1,4- | awk -v var=${label} '{if($1 ~ "-") print substr($1, 1,
length($1)-4)"\t"$2"\t"$3"\t"$4"\t"var; else print $0"\t"var}' \
> ${outdir}/${prefix}.tp-baseline.vcfstats_short;
done
```

4. For each sequencing platform (from step 2) and for the GIAB truth sets (from step 3) combine counts of TP INDELs falling into appropriate size bins, as defined using the SNV/INDEL and SV truth sets (to cover the entire range of INDEL sizes):

```
SNVIndel_dir=${1} ## path to directory containing
*tp-baseline.vcfstats_short outputs for TP calls, as defined by the GIAB
SNV/INDEL truth set.
SV_dir=${2} ## path to directory containing *tp-baseline.vcfstats_short
outputs for TP calls, as defined by the GIAB SV truth set.
prefix=${3}
outdir=${4}

paste -d '\t' ${SV_dir}/${prefix}.tp-baseline.vcfstats_short \
${SNVIndel_dir}/${prefix}.tp-baseline.vcfstats_short \
| awk '{print $1"\t"$2+$7"\t"$3+$8"\t"$4+$9}' | grep -v length \
> ${outdir}/${prefix}.tp-baseline.vcfstats_short
```

5. Concatenate all *${outdir}/${output_prefix}.vcfstats_short* files and remove header lines:

```
cat ${outdir}/${prefix}.tp-baseline.vcfstats_short | grep -v "length" > combo
```

6. Plot indel size distribution of high confidence TP INDEL calls for all sequencing platforms (including INDEL size distribution of the combined GIAB SNV/SV truth sets) using the following R script:

```
require(ggplot2)
require(reshape)

args <- commandArgs(trailingOnly=TRUE)
combo_file <- args[1] ## path to the combo file from step 5.
combo <- read.table(combo_file, header=F)

## Label the columns:
colnames(combo) <- c("length","Delete","Insert", "Indel", "Technology")

## Select all columns except for "Indel" and convert the data frame into a
molten data frame:
Combo_m <- melt(combo[,c(1,2,3,5)], id=c("length", "Technology"))

## Create a new Length column in which length of deletions is expressed as
a negative value, for plotting purposes:
combo_m$Length <- ifelse(combo_m$variable == "Delete", -(combo_m$length),
combo_m$length)

## Set the desired order in which sequencing platforms will be listed in
the plot:
combo_m$Technology <- factor(combo_m$Technology, levels=c("BGISEQ500",
"MGISEQ2000", "HiSeq2500", "HiSeq4000", "HiSeqX10","NovaSeq-2x150",
"NovaSeq-2x250", "PacBioCCS", "PacBioCLR", "PromethION",
"GIAB_Indel_truth_set", "GIAB_SV_truth_set"))

## Define colors for each sequencing platform:
Custom_colors <- c("BGISEQ500"="#fd8d3c", "HiSeq2500"="#6baed6",
"HiSeq4000"="#3182bd", "HiSeqX10"="#08519c", "MGISEQ2000"="#e6550d",
"NovaSeq-2x150"="#f768a1", "NovaSeq-2x250"="#ae017e",
"PacBioCCS"="#74c476", "PacBioCLR"="#31a354", "PromethION" = "#006d2c")

## Subset out GIAB truth sets from the data frame:
Combo_m_truth <- subset(combo_m, combo_m$Technology ==
"GIAB_Indel_truth_set" | combo_m$Technology=="GIAB_SV_truth_set")

## Generate a barplot with counts of INDELs falling into each size bin in
the combined GIAB truth set:
p <- ggplot(data=combo_m_truth, aes(x=Length, y=value+1)) + \
geom_bar(stat="identity",color=alpha("gray", 0.4), fill="transparent") + \
scale_y_continuous(trans='log10') + \
xlim(-100,100) + \
```

```
theme_minimal()

## Plot indel size distribution for high confidence TP INDEL calls:
p + geom_point(data = combo_m_wo_truth, aes(x=Length, y=value+1,
color=Technology), size=1, alpha=0.3) + \
geom_line(data = combo_m_wo_truth, aes(x=Length, y=value+1,
color=Technology),alpha=0.3) + \
theme_minimal() + \
xlab("size (bp)") + \
ylab("count+1") + \
scale_y_continuous(trans='log10') + \
xlim(-100,100) + \
scale_color_manual(values=custom_colors)
```
