## Supplementary Data 3 for "Multi-Platform Assessment of DNA Sequencing Performance using Human and Bacterial Reference Genomes in the ABRF Next-Generation Sequencing Study": InsertSizes.Rmd

````
---
title: "ABRF NGS2: Insert Sizes"
author: "Jonathan Foox"
date: "June 3, 2020"
output: 
  html_notebook: 
    code_folding: none
    highlight: tango
    toc: yes
    
---


```{r}
library(reshape2)
library(dplyr)
library(tidyr)
library(cowplot)
library(ggplot2)
library(ggridges)
library(ggpubr)
```

```{r}
df_insert = read.csv("qc_insert/insertsize_matrix_ALL.csv")
df_insert$sample = "sample"
head(df_insert)
```

```{r}
df_insert2 = group_by(df_insert, allinfo) %>% mutate(percent = num/sum(num)*100)
head(df_insert2)
```

```{r}
df_insert2$platform = factor(df_insert2$platform, levels=c("HiSeq2500","HiSeq4000","HiSeqX10","NextSeq","NovaSeq","BGISEQ500","MGISEQ2000"))
df_insert2$rep = factor(df_insert2$rep, levels=c("REP01", "REP02", "REP01-2x150", "REP01-2x250"))
```


```{r, fig.height=14, fig.width=6}
inserts = ggplot(df_insert2, aes(insertsize, allinfo, height=percent, fill=platform)) +
  geom_ridgeline(scale=5, show.legend = F) + #, binwidth=5
  #stat_bin(scale=10, binwidth=5) +
  facet_grid(platform+lab~., scales="free_y", space="free_y", switch="y") + 
  #guides(colour=FALSE) +
  scale_fill_discrete(guide = guide_legend(reverse=TRUE)) +
  theme_classic() + 
  theme(#axis.text.y=element_blank(), 
    axis.title.y=element_blank(), 
    axis.ticks.y=element_blank(), 
    panel.spacing.y=unit(0,"pt"), 
    strip.text.y=element_text(size=10, angle=20), 
    strip.text.x = element_blank(),
    axis.text.y=element_blank(), 
    text = element_text(size=16),
    legend.text=element_text(size=10)) + 
  xlab("Insert size")
inserts
```

```{r}
pdf(file="../figures/insert_sizes_v2.pdf", width=6, height=14)
inserts
#dev.off()
```
````
