## Supplementary Data 3 for "Multi-Platform Assessment of DNA Sequencing Performance using Human and Bacterial Reference Genomes in the ABRF Next-Generation Sequencing Study": MappingStatsALL.Rmd

````
---
title: "ABRF NGS2: BAM Stats for ALL Samples"
author: "Jonathan Foox"
date: "May 15, 2020"
output: 
  html_notebook: 
    code_folding: none
    highlight: tango
    toc: no
    
---

```{r}
setwd("/athena/masonlab/scratch/projects/abrf_ngs_phaseii/analysis")
```


```{r}
library(reshape2)
library(dplyr)
library(tidyr)
library(cowplot)
library(ggplot2)
library(ggridges)
library(ggpubr)
```


```{r}
df_md = read.csv("mosdepth_ALL.csv")
df_md = separate(df_md, col="sample", by="_", into=c("platform","lab","genome","rep"), remove=F)
```

```{r}
bac_taxa = c("Eco", "Ste", "Pflu", "Pool")
df_md$kingdom = ifelse(df_md$genome %in% bac_taxa, "bacteria", "human")
df_md$kingdom = factor(df_md$kingdom, levels=c("human", "bacteria"))

longs = c("Flongle", "GridION", "PromethION", "PacBioCLR", "PacBioCCS", "Proton", "S5", "PGM")
df_md$readtype = ifelse(df_md$platform %in% longs, "long", "short")
df_md$readtype = factor(df_md$readtype, levels=c("short", "long"))

df_md$ome = ifelse((df_md$platform %in% c("Proton", "S5") & df_md$kingdom == "human"), "exome", "genome")
df_md$ome = factor(df_md$ome, levels=c("genome", "exome"))


df_md$platform = factor(df_md$platform, levels=c("HiSeq2500", "HiSeq4000", "HiSeqX10", "NovaSeq", "NextSeq", "Genapsys",
                                                 "MiSeq", "Proton", "S5", "PGM", "BGISEQ500", "MGISEQ2000", 
                                                 "Flongle", "GridION", "PromethION", "PacBioCCS", "PacBioCLR"))

head(df_md)
```


```{r}
df_md_tot = subset(df_md, chrom=="total")
head(df_md_tot)
```


```{r, fig.width=15, fig.height=4}
mosdepth_human = ggplot(subset(df_md_tot, kingdom=="human"), 
       aes(x=sample, y=mean)) +
  geom_bar(stat="identity", aes(fill=genome)) +
  #geom_point(aes(color=genome)) +
  theme_bw() +
  theme(text = element_text(size=12),
        axis.text.x = element_blank(),
        axis.ticks.x = element_blank(),
        panel.grid.minor = element_blank(),
        strip.text.x = element_text(size=0),
        strip.background = element_blank(),
        panel.spacing=unit(0, "lines"),
        plot.margin = unit(c(0, 0, 0, 0), "cm")) +
  #scale_fill_manual(values=c("Mother"="#FBB4AE", "Father"="#B3CDE3", "Son"="#CCEBC5", "Son1"="#CCEBC5", "Son2"="#CCEBC5", "Son3"="#CCEBC5")) +
  scale_fill_manual(values=c("#E41A1C", "#377EB8", "#4DAF4A", "#4DAF4A", "#3b9639", "#2c7f2b")) +
  facet_grid(.~kingdom+ome+readtype+platform, scales="free", space = "free") +
  scale_y_continuous(breaks=c(1, 10, 20, 40, 60, 80)) +
  xlab("") + ylab("Mean Depth") +
  coord_trans(y="sqrt")
mosdepth_human
```

```{r}
mosdepth_bacteria = ggplot(subset(df_md_tot, kingdom=="bacteria"), 
       aes(x=sample, y=mean)) +
  geom_bar(stat="identity", aes(fill=genome)) +
  #geom_point(aes(color=genome)) +
  theme_bw() +
  theme(text = element_text(size=12),
        axis.text.x = element_blank(),
        axis.ticks.x = element_blank(),
        panel.grid.minor = element_blank(),
        strip.text.x = element_text(size=0),
        strip.background = element_blank(),
        panel.spacing=unit(0, "lines"),
        plot.margin = unit(c(0, 0, 0, 0), "cm")) +
  #scale_fill_manual(values=c("#E41A1C", "#377EB8", "#4DAF4A", "#4DAF4A", "#3b9639", "#2c7f2b")) +
  facet_grid(.~kingdom+ome+readtype+platform, scales="free", space = "free") +
  xlab("") + ylab("") +
  scale_y_continuous(breaks=c(10, 100, 200, 400, 600)) +
  coord_trans(y="sqrt")
mosdepth_bacteria
```


```{r, fig.width=15, fig.height=3}
print(plot_grid(mosdepth_human, mosdepth_bacteria, ncol=2, rel_widths=c(1,0.6)))
```


```{r}
df_ss = read.csv("samstats_ALL_v2.csv")
df_ss = separate(df_ss, col="sample", by="_", into=c("platform","lab","genome","rep"), remove=F)
```

```{r}
bac_taxa = c("Eco", "Ste", "Pflu", "Pool")
df_ss$kingdom = ifelse(df_ss$genome %in% bac_taxa, "bacteria", "human")
df_ss$kingdom = factor(df_ss$kingdom, levels=c("human", "bacteria"))

longs = c("Flongle", "GridION", "PromethION", "PacBioCLR", "PacBioCCS", "Proton", "S5", "PGM")
df_ss$readtype = ifelse(df_ss$platform %in% longs, "long", "short")
df_ss$readtype = factor(df_ss$readtype, levels=c("short", "long"))

df_ss$ome = ifelse((df_ss$platform %in% c("Proton", "S5") & df_ss$kingdom == "human"), "exome", "genome")
df_ss$ome = factor(df_ss$ome, levels=c("genome", "exome"))


df_ss$platform = factor(df_ss$platform, levels=c("HiSeq2500", "HiSeq4000", "HiSeqX10", "NovaSeq", "NextSeq", "Genapsys",
                                                 "MiSeq", "Proton", "S5", "PGM", "BGISEQ500", "MGISEQ2000", 
                                                 "Flongle", "GridION", "PromethION", "PacBioCCS", "PacBioCLR"))

head(df_ss)
```

```{r}
df_ss_m = melt(df_ss, id.vars = c("sample", "platform", "lab", "genome", "rep", "kingdom", "readtype", "ome"))
head(df_ss_m)
```


```{r, fig.width=15, fig.height=4}
ss_human = ggplot(subset(df_ss_m, variable!="allreads" & variable!="basesmappedcigar" & kingdom=="human"), 
             aes(sample, value, fill=variable)) + 
  geom_bar(stat="identity", position = position_fill()) +
  xlab("") + ylab("Read Mapping Proportions") + 
 theme_bw() +
  theme(text = element_text(size=12),
        axis.text.x = element_blank(),
        axis.ticks.x = element_blank(),
        panel.grid.minor = element_blank(),
        strip.text.x = element_text(size=0),
        strip.background = element_blank(),
        panel.spacing=unit(0, "lines"),
        plot.margin = unit(c(0, 0, 0, 0), "cm")) +
  #scale_fill_manual(values=c("#E41A1C", "#377EB8", "#4DAF4A", "#4DAF4A", "#3b9639", "#2c7f2b")) +
  facet_grid(.~kingdom+ome+readtype+platform, scales="free", space = "free") +
  scale_fill_manual("legend", values = c("properlymapped" = "#4DAF4A", 
                                         "duplicates" = "#377EB8", 
                                         "multimapped"="#984EA3", 
                                         "unmapped" = "#E41A1C"))
ss_human
```


```{r}
ss_bacteria = ggplot(subset(df_ss_m, variable!="allreads" & variable!="basesmappedcigar" & kingdom=="bacteria"), 
             aes(sample, value, fill=variable)) + 
  geom_bar(stat="identity", position = position_fill(), show.legend=F) +
  xlab("") + ylab("") + 
 theme_bw() +
  theme(text = element_text(size=12),
        #axis.text.x = element_blank(),
        axis.text.x = element_text(angle=90, size=6),
        axis.ticks.x = element_blank(),
        panel.grid.minor = element_blank(),
        strip.text.x = element_text(size=0),
        strip.background = element_blank(),
        panel.spacing=unit(0, "lines"),
        plot.margin = unit(c(0, 0, 0, 0), "cm")) +
  #scale_fill_manual(values=c("#E41A1C", "#377EB8", "#4DAF4A", "#4DAF4A", "#3b9639", "#2c7f2b")) +
  facet_grid(.~kingdom+ome+readtype+platform, scales="free", space = "free") +
  scale_fill_manual("legend", values = c("properlymapped" = "#4DAF4A", 
                                         "duplicates" = "#377EB8", 
                                         "multimapped"="#984EA3", 
                                         "unmapped" = "#E41A1C"))
ss_bacteria
```


```{r, fig.width=15, fig.height=6}
pdf(file="../figures/Fig1__StatsALL_small.pdf", width=10, height=4)
print(plot_grid(mosdepth_human, mosdepth_bacteria, ss_human, ss_bacteria, ncol=2, nrow=2, rel_widths=c(1,0.6), align = 'v', axis = 'l'))
dev.off()
```


```{r, fig.width=15, fig.height=4}
mosdepth_human_facetRot = ggplot(subset(df_md_tot, kingdom=="human"), 
       aes(x=sample, y=mean)) +
  geom_bar(stat="identity", aes(fill=genome), show.legend=F) +
  #geom_point(aes(color=genome)) +
  theme_bw() +
  theme(text = element_text(size=12),
        axis.text.x = element_blank(),
        axis.ticks.x = element_blank(),
        panel.grid.minor = element_blank(),
        strip.text.x = element_text(angle=90, size=6),
        panel.spacing=unit(0, "lines"),
        plot.margin = unit(c(0, 0, 0, 0), "cm")) +
  #scale_fill_manual(values=c("Mother"="#FBB4AE", "Father"="#B3CDE3", "Son"="#CCEBC5", "Son1"="#CCEBC5", "Son2"="#CCEBC5", "Son3"="#CCEBC5")) +
  scale_fill_manual(values=c("#E41A1C", "#377EB8", "#4DAF4A", "#4DAF4A", "#3b9639", "#2c7f2b")) +
  facet_grid(.~kingdom+ome+readtype+platform, scales="free", space = "free") +
  scale_y_continuous(breaks=c(1, 10, 20, 40, 60, 80)) +
  xlab("") + ylab("Mean Depth") +
  coord_trans(y="sqrt")
mosdepth_human_facetRot
```

```{r}
mosdepth_bacteria_facetRot = mosdepth_bacteria = ggplot(subset(df_md_tot, kingdom=="bacteria"), 
       aes(x=sample, y=mean)) +
  geom_bar(stat="identity", aes(fill=genome), show.legend=F) +
  #geom_point(aes(color=genome)) +
  theme_bw() +
  theme(text = element_text(size=12),
        axis.text.x = element_blank(),
        axis.ticks.x = element_blank(),
        panel.grid.minor = element_blank(),
        strip.text.x = element_text(angle=90, size=6),
        panel.spacing=unit(0, "lines"),
        plot.margin = unit(c(0, 0, 0, 0), "cm")) +
  #scale_fill_manual(values=c("#E41A1C", "#377EB8", "#4DAF4A", "#4DAF4A", "#3b9639", "#2c7f2b")) +
  facet_grid(.~kingdom+ome+readtype+platform, scales="free", space = "free") +
  xlab("") + ylab("") +
  scale_y_continuous(breaks=c(10, 100, 200, 400, 600)) +
  coord_trans(y="sqrt")
mosdepth_bacteria_facetRot
```


```{r, fig.width=15, fig.height=3}
pdf(file="../figures/Fig1__StatsALL_facetRot.pdf", width=15, height=6)
print(plot_grid(mosdepth_human_facetRot, mosdepth_bacteria, ncol=2, rel_widths=c(1,0.6)))
dev.off()
```


# mosdepth plots

```{r}
colorsList = list()
colorsList[["HiSeq2500"]] = "#9ecae1"
colorsList[["HiSeq4000"]] = "#6baed6"
colorsList[["HiSeqX10"]] = "#3182bd"
colorsList[["NovaSeq"]] = "#ae017e"
colorsList[["NextSeq"]] = "#d0d0ef"
colorsList[["BGISEQ500"]] = "#fc8d59"
colorsList[["MGISEQ2000"]] = "#e34a33"
colorsList[["Genapsys"]] = "#ffd92f"
colorsList[["Flongle"]] = "#a1d99b"
colorsList[["GridION"]] = "#41ab5d"
colorsList[["PromethION"]] = "#006d2c"
colorsList[["PacBioCLR"]] = "#ef3b2c"
colorsList[["PacBioCCS"]] = "#a50f15"
colorsList2 = as.data.frame(unlist(colorsList))
colnames(colorsList2)[1] = "plat_colors"
```


```{r}
df_md_human_distrib = subset(df_md, kingdom=="human" & ome=="genome" & chrom!="total")
df_md_human_distrib = df_md_human_distrib %>% group_by(sample, platform, lab, genome, rep) %>% mutate(pct = bases / sum(bases) * 100)
```

```{r}
df_md_human_distrib$platform = factor(df_md_human_distrib$platform, levels=c("HiSeq2500", "HiSeq4000", "HiSeqX10", "NovaSeq", "NextSeq", 
                                                       "BGISEQ500", "MGISEQ2000", "Genapsys", "Flongle", "GridION", "PromethION", "PacBioCLR", "PacBioCCS",
                                                       "MiSeq", "Proton", "S5", "PGM"))

df_md_human_distrib$chrom = factor(df_md_human_distrib$chrom, levels=c(c("1", "2", "3", "4", "5", "6", "7", "8", "9", "10", 
"11", "12", "13", "14", "15", "16", "17", "18", "19", "20", "21", "22", "X", "Y", "MT",  "AE015929.1", "AE016825.1", "AL009126.3", "CP001628.1", "GL000191.1", "GL000192.1", "GL000193.1", "GL000194.1", "GL000195.1", "GL000196.1", "GL000197.1", "GL000198.1", "GL000199.1", "GL000200.1", "GL000201.1", "GL000202.1", "GL000203.1", "GL000204.1", "GL000205.1", "GL000206.1", "GL000207.1", "GL000208.1", "GL000209.1", "GL000210.1", "GL000211.1", "GL000212.1", "GL000213.1", "GL000214.1", "GL000215.1", "GL000216.1", "GL000217.1", "GL000218.1", "GL000219.1", "GL000220.1", "GL000221.1", "GL000222.1", "GL000223.1", "GL000224.1", "GL000225.1", "GL000226.1", "GL000227.1", "GL000228.1", "GL000229.1", "GL000230.1", "GL000231.1", "GL000232.1", "GL000233.1", "GL000234.1", "GL000235.1", "GL000236.1", "GL000237.1", "GL000238.1", "GL000239.1", "GL000240.1", "GL000241.1", "GL000242.1", "GL000243.1", "GL000244.1", "GL000245.1", "GL000246.1", "GL000247.1", "GL000248.1", "GL000249.1", "NC_007605", "NC_013967.1", "NC_016830.1", "NC_017316.1", "NC_017668.1", "NZ_AHCA01000001.1", "NZ_AHCA01000002.1", "NZ_AHCA01000003.1", "NZ_AHCA01000004.1", "NZ_AHCA01000005.1", "NZ_AHCA01000006.1", "NZ_AHCA01000007.1", "NZ_AHCA01000008.1", "NZ_AHCA01000009.1", "NZ_AHCA01000010.1", "NZ_AHCA01000011.1", "NZ_AHCA01000012.1", "NZ_AHCA01000013.1", "NZ_AHCA01000014.1", "NZ_AHCA01000015.1", "NZ_AHCA01000016.1", "NZ_AHCA01000017.1", "NZ_AHCA01000018.1", "NZ_AHCA01000019.1", "NZ_AHCA01000020.1", "NZ_AHCA01000021.1", "NZ_AHCA01000022.1", "NZ_AHCA01000023.1", "NZ_AHCA01000024.1", "NZ_AHCA01000025.1", "NZ_AHCA01000026.1", "NZ_AHCA01000027.1", "NZ_AHCA01000028.1", "NZ_AHCA01000029.1", "NZ_AHCA01000030.1", "NZ_AHCA01000031.1", "NZ_AHCA01000032.1", "NZ_AHCA01000033.1", "NZ_AHCA01000037.1", "NZ_AHCA01000040.1", "NZ_AHCA01000042.1", "NZ_AHCA01000045.1", "NZ_AHCA01000052.1", "NZ_AHCA01000055.1", "U00096.3",  "hs37d5", "total")))

```


```{r, fig.width=13, fig.height=15}
cov_by_chrom = ggplot(df_md_human_distrib, aes(x=sample, y=pct, fill=platform)) +
  geom_bar(stat="identity") +
  theme_bw() +
  theme(text = element_text(size=12),
        axis.text.x = element_blank(),
        axis.ticks.x = element_blank(),
        panel.grid.minor = element_blank(),
        strip.text.y = element_text(angle=0, size=6)) +
  facet_wrap(chrom~.) +
  scale_fill_manual(values = colorsList) +
  xlab("") + ylab("Percentage")
cov_by_chrom
```


```{r}
pdf(file="../figures/cov_by_chrom.pdf", width=13, height=15)
cov_by_chrom
dev.off()
```
````
